# TBCK Deficiency Alters Ribosomal Function, RNA Splicing, and miRNA Networks: Insights from Multi-Omics Analyses

**DOI:** 10.1101/2025.09.23.677540

**Authors:** Abdias Diaz-Rosado, Kelly Clark, Rajesh Angireddy, Michael Gilbert, Annabel Sangree, Jan Oppelt, Nicholas Vrettos, Emily L. Durham, Elizabeth Gonzalez, Emily E. Lubin, Ashley Melendez, Dana Layo-Carris, Kaitlin A Katsura, Elizabeth Bhoj

## Abstract

TBC1 domain-containing kinase (TBCK) is an important protein with implications in brain development. Biallelic variants in the *TBCK* gene are known to cause TBCK-related neurodevelopmental disorder (OMIM #616900) [1], a rare genetic multisystemic disease characterized by developmental delay, variable developmental regression, seizures, and premature death in late childhood for which no cure is currently available. Though previous work has provided a better understanding of the protein’s role, the mechanism for how *TBCK* variants affect gene expression and protein regulation has remained understudied. To better understand the impact of these alterations, and using an unbiased approach, we employed the power of multi-omics to define the cellular consequences at the transcript and protein level. Our comprehensive analysis uncovered significant disruptions in ribosomal and translation-related pathways with widespread alternative splicing defects, and key miRNA changes that validate previously reported molecular findings. This work provides a clearer molecular framework for TBCK dysfunction in *TBCK-/-* cells and offers a valuable foundation to identify potential therapeutic targets.

## Background

TBCK-related neurodevelopmental disorder (TBCK Syndrome) is a rare multi-systemic Mendelian disorder marked by severe progressive neurodevelopmental impairments that manifest in early childhood. Affected individuals commonly present with developmental delays, seizures, hypotonia, variable regression and other systemic complications [2–4]. The disease is caused by biallelic variants in the *TBC1 domain-containing kinase* (*TBCK*) gene, which encodes a protein implicated in cellular growth and mTOR signaling [2, 5, 6]. Variants in *TBCK* have also been shown to impair autophagy, mitochondrial respiration, and lysosomal degradation [7, 8], all of which are key in maintaining cellular homeostasis on metabolically demanding tissues such as the brain. These molecular irregularities have been consistently observed in *TBCK-/-* cells and are thought to contribute to the multisystemic manifestations of the syndrome [2, 7–9].

Structurally, the TBCK protein comprises three distinct domains: a Protein Kinase (PK) domain, a Tre-2/Bub2/Cdc16 (TBC) domain, and a Rhodanese (R) domain [3, 10]. Even though the precise function of TBCK remains to be fully characterized, its TBC domain is believed to mediate the protein’s catalytic function, whereas the PK and R domains remain poorly studied [10, 11]. TBC domains are evolutionarily conserved and function as GTPase-Activating Proteins (GAPs) for Rab GTPases, using a characteristic dual-finger mechanism to catalyze the hydrolysis of GTP to GDP [12].

Similar to other well-characterized TBC domains, TBCK’s TBC domain is predicted, based on *in-silico* analysis, to function as a GAP for members of the Rab family of small GTPases [11]. Though there is no direct biochemical evidence of TBCK’s GAP activity, a prior study showed that TBCK preferentially binds to the active GTP-bound form of Rab5 over its inactive GDP-bound state [13]. This preferential interaction suggests a potential regulatory role in Rab5-dependent pathways, which are important for early endosome formation and maturation [14, 15]. Therefore, variants in *TBCK* may lead to irregularities in vesicle trafficking. This proposed mechanism is supported by studies in *TBCK-/-* cells, which have revealed impaired lysosomal function [8, 9], and defects in the endolysosomal trafficking of neurons [16] (*bioRxiv*).

In more recent studies, TBCK has been implicated in mRNA transport via its role in the FERRY complex (<u>F</u>ive-subunit <u>E</u>ndosomal <u>R</u>ab5 and <u>R</u>NA/ribosome intermediar<u>Y</u>), which is important for the localized translation of mRNAs in neurons, one in which Rab5 plays a key role in transporting the complex [17, 18]. Localized mRNA translation is crucial for maintaining the specialized functions of polarized cells, particularly in neurons, enabling them to meet metabolic demands and respond dynamically to their microenvironment [19]. Central to this process is the precise synthesis and assembly of ribosomes, which are responsible for translating genetic information into functional proteins [20, 21]. Disruptions in any component of this finely tuned system, including Rab5-associated transport, can lead to severe impairments in cellular homeostasis and contribute to disease progression [22]. While TBCK has been implicated in diverse cellular pathways, the extent to which pathogenic variants disrupt gene expression across multiple domains, including transcription, isoform splicing and protein synthesis, remains unknown.

To investigate how TBCK loss affects different pathways, we conducted a comprehensive multi-omics analysis using primary *TBCK-/-* fibroblasts. These fibroblasts retain the patient’s native genetic background and endogenous *TBCK* expression levels, allowing for the detection of disease-relevant molecular alterations without the confounding effects of immortalization. Since TBCK Syndrome is a multisystemic disorder, dermal fibroblasts serve as a practical and clinically accessible model to capture systemic cellular defects that may reflect broader pathophysiological mechanisms beyond the central nervous system [23]. Notably, the specific fibroblast lines used in this study have been characterized and published previously [7, 8], providing a validated foundation for our analyses. We applied RNA sequencing (RNA-seq) to profile global gene expression changes, mass spectrometry-based proteomics to assess alterations in the proteome, splicing analysis to examine RNA processing dynamics, and miRNA-sequencing (miRNA-seq) to explore post-transcriptional regulation by small non-coding RNAs in the context of the disease. Our analyses revealed novel insights into disrupted molecular pathways such as ribosome biogenesis, translation, splicing, and miRNAs, reinforcing the critical role of TBCK in supporting key regulators of cellular physiology. The findings of this work expand our understanding of TBCK’s molecular function and highlight previously unrecognized regulatory networks that may serve as potential targets for future therapeutic intervention.

## Results

### *TBCK* variants disrupt ribosomal gene expression and impair translational capacity

To investigate how loss of *TBCK* influenced global transcriptional landscapes, we performed bulk RNA-sequencing on skin-derived fibroblasts from three affected individuals harboring the homozygous Boricua variant p.R126X [7]. This variant is associated with the severe, neurodegenerative spectrum of the disease (genotypes shown in **Fig. S1F**). Differential expression (DE) analysis of age and sex matched samples identified 1,018 significantly dysregulated genes in *TBCK-/-* cells relative to controls (527 upregulated, 491 downregulated; **Fig. S1A**). Gene ontology (GO) enrichment using MSigDB C5 ontology revealed 488 affected biological pathways (**Fig. S1B**). From the top 40 enriched terms, three major functional categories emerged: ribosome-related processes, RNA splicing, and non-coding RNA regulation (**Fig. 1A**). Though additional pathways were also enriched, overrepresented categories highlighted central roles in ribosome regulation and splicing control. Principal component analysis (PCA) distinguished *TBCK-/-* cells from controls, showing discrete clustering with minimal intragroup variance despite the small sample size (*n=3*) (**Fig. 1B**). Notably, fibroblasts from individuals 2 and 3, siblings who share a consanguineous background, clustered closely while individual 1 exhibited a more distant transcriptional profile.

**Figure 1.**
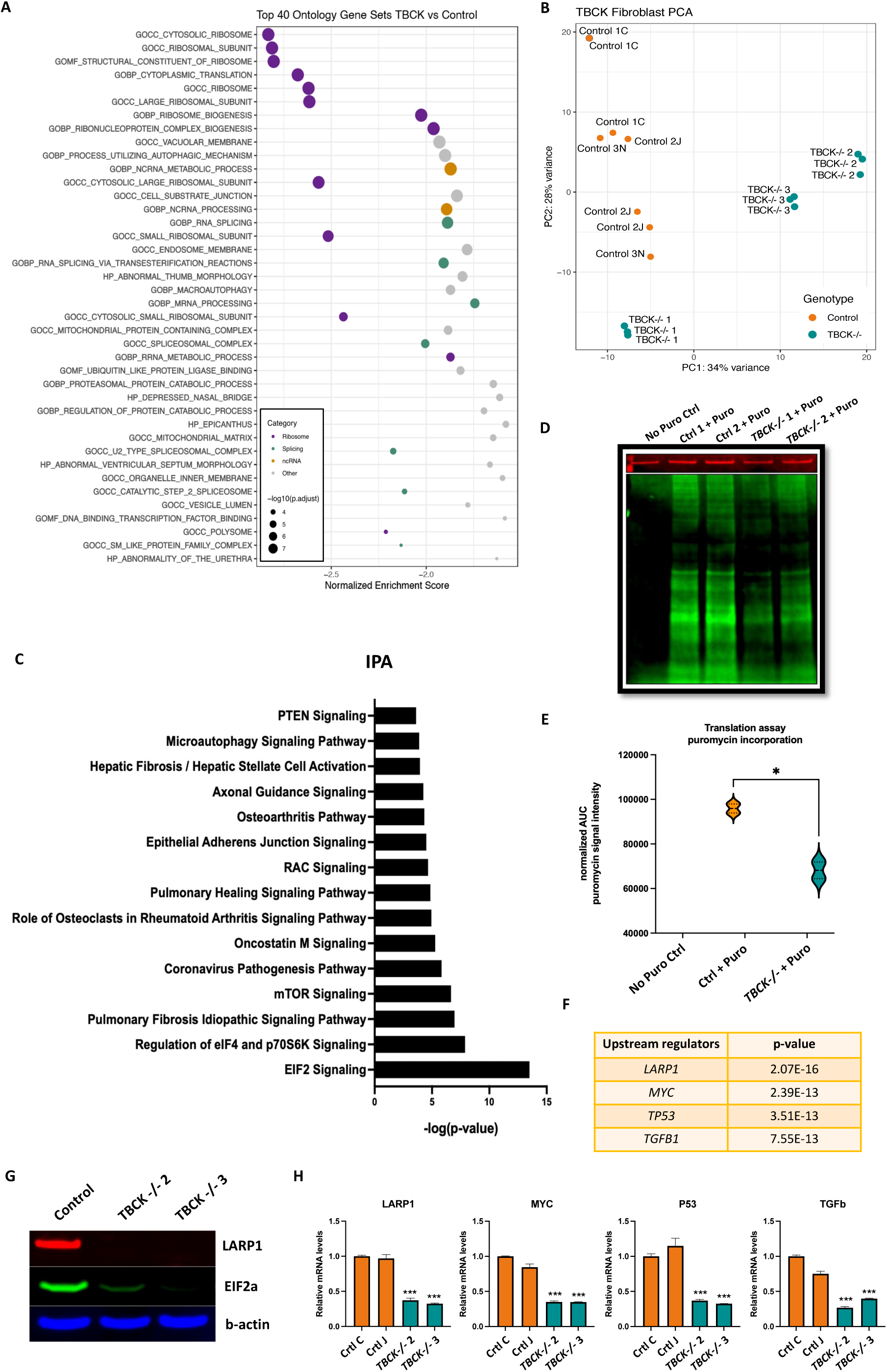
TBCK mutations disrupt ribosomal gene expression and impair translational capacity. – RNA-seq and functional analyses of *TBCK-/-* fibroblasts (n = 3 individuals; n = 3 controls): **A**) Dot plot showing the top 40 significantly enriched gene ontology pathways (MSigDB C5), categorized by ribosomal function, RNA splicing, and non-coding RNA regulation. **B**) Principal component analysis (PCA) depicting transcriptomic separation between *TBCK-/-* and control samples. Each biological replicate was sequenced with technical triplicates. **C**) Ingenuity Pathway Analysis (IPA) identifying top dysregulated pathways in *TBCK-/-* cells. **D**) Puromycin incorporation (SUnSET) assay following 3hr serum starvation and 10% FBS refeeding in the presence of puromycin. Loading control, vinculin, is represented in red, and puromycin in green. **E**) Violin plot quantifying puromycin incorporation (AUC/fluorescence) in control vs. *TBCK-/-* samples. **F**) Predicted upstream regulators identified by IPA based on differentially expressed genes. **G**) Western blot validation of protein expression, for key translational regulators identified in IPA. **H**) RT-qPCR analysis of transcript levels for upstream regulators showing reduced levels in *TBCK-/-* cells.

To complement our GO findings with a more statistically powerful model, we employed Ingenuity Pathway Analysis (IPA) to identify signaling cascades perturbed in *TBCK-/-* cells. This analysis uncovered 528 altered pathways (**Fig. S1C**), with eIF2 and eIF4/p70S6K as the most affected (**Fig. 1C**), both playing essential roles in translation initiation and elongation [24, 25].

To validate the results from our GO and IPA, we conducted a puromycin incorporation assay, SUrface Sensing of Translation (SUnSET), as a proxy for nascent protein synthesis. *TBCK-/-* cells showed reduced puromycin levels as compared to controls, indicating globally impaired translation (**Fig. 1D–E**). To identify potential upstream regulators driving these transcriptomic and translational abnormalities, we leveraged IPA’s predictive algorithm, which identified four upstream regulators significantly affected in *TBCK-/-* cells: LARP1, MYC, TP53, and TGFB1 (**Fig. 1F**). Strikingly, LARP1, a well-established mediator of ribosomal protein mRNA stability and translational control [26, 27], emerged as the most significantly affected regulator. To validate these IPA-based predictions, we assessed the protein and transcript levels of the top upstream regulators, LARP1, MYC, TP53, TGFB1, as well as EIF2, and found that all five targets were consistently downregulated in *TBCK-/-* fibroblasts compared to controls (**Fig. 1G– H**), supporting their involvement in the observed transcriptomic and translational changes of *TBCK-/-* samples.

Together, these results reveal that biallelic loss-of-function in *TBCK* exert broad transcriptional effects with a marked impact on ribosomal gene networks and translational capacity. The consistent disruption of upstream regulators of translation pathways and core components of the protein synthesis machinery highlights TBCK’s role in maintaining ribosomal homeostasis. These transcriptomic alterations raised questions of whether TBCK loss also perturbs protein abundance, which we addressed next through global proteomics analysis.

### Proteomic profiling of *TBCK-/-* fibroblasts reveals dysregulation of RNA splicing and other disease-relevant pathways

While transcript-level data provides valuable insights into gene expression, protein abundance is shaped by multiple layers of regulation, including mRNA stability, splicing variation, post-translational modification, and degradation [28, 29]. To complement our transcriptomic analysis and determine whether alterations in *TBCK-/-* cells extend to the proteome, we performed mass spectrometry-based proteomic profiling on *TBCK-/-* dermal fibroblasts and control cells.

Although multiple hypothesis correction analysis did not yield statistically significant DE proteins (*adjusted p < 0.05*) (**Fig. S2A**), our exploratory analysis identified 877 proteins with nominal significance (*unadjusted p < 0.05*) in *TBCK-/-* cells (**Fig. S2B**). PCA showed clear separation between *TBCK-/-* and control groups, indicating distinct proteomic signatures (**Fig. 2A**). This clustering was further supported by unsupervised hierarchical analysis, which revealed high intra-group similarity and consistent inter-group separation (**Fig. 2B**).

**Figure 2.**
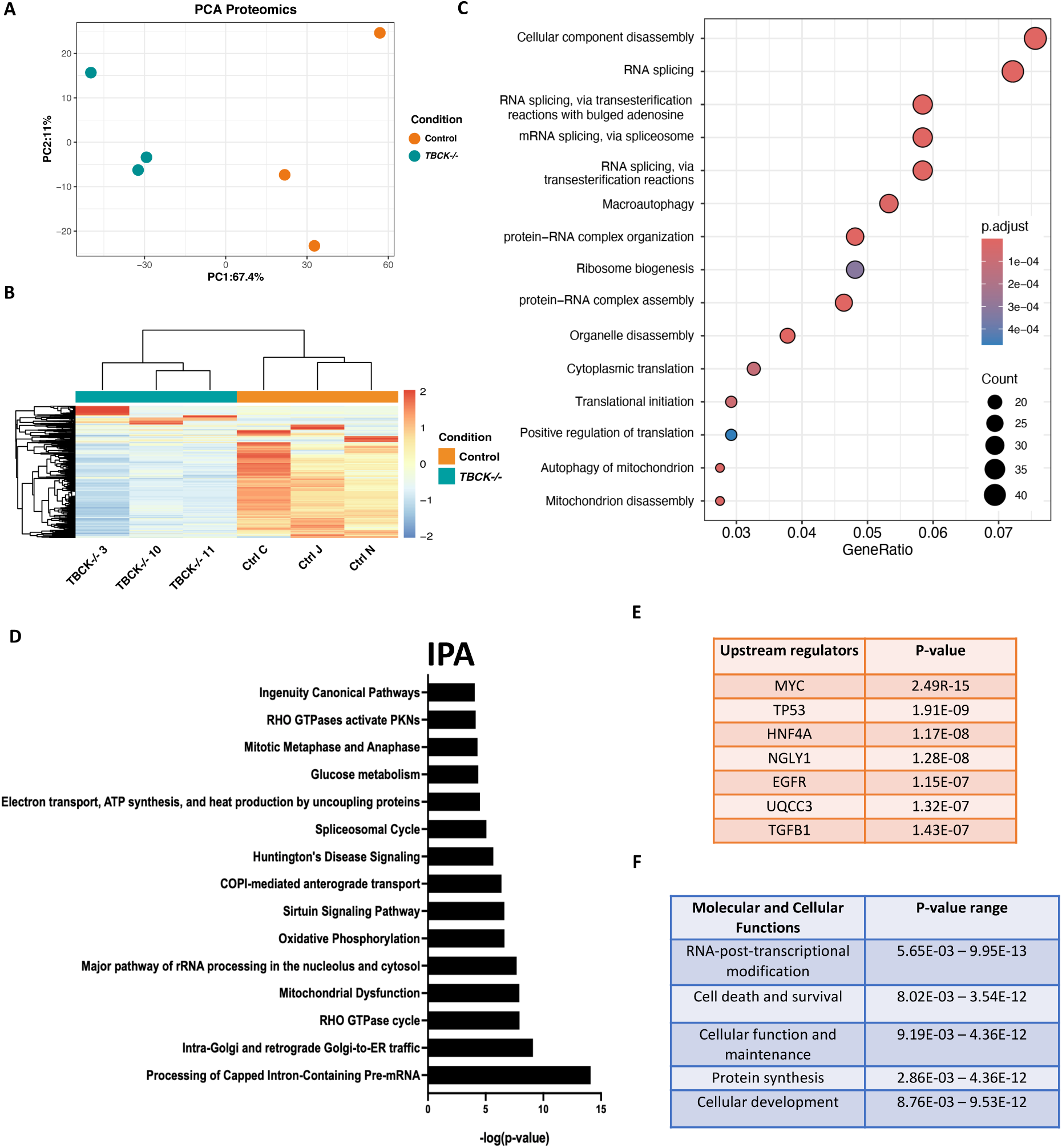
Proteomic profiling of TBCK-deficient fibroblasts reveals dysregulation of RNA splicing and other disease-relevant pathways. – **A**) Principal Component Analysis (PCA) of proteomic data from *TBCK-/-* fibroblasts and controls (n = 3 per group), showing distinct clustering by condition. **B**) Heatmap of protein abundance with unsupervised hierarchical clustering, illustrating consistent intra-group similarity and inter-group separation. Color scale represents relative protein expression. **C**) Gene ontology enrichment analysis of dysregulated proteins, highlighting pathways related to RNA splicing, ribosome biogenesis, and mitochondrial function. Circle size reflects gene count; color indicates adjusted p-value. **D**) Ingenuity Pathway Analysis (IPA) of proteomic data identifying top dysregulated canonical pathways in TBCK-deficient cells. **E**) Upstream regulators predicted by IPA to be altered in *TBCK-/-* samples, ranked by p-value. Regulators shown have known roles in transcriptional and post-transcriptional control. **F**) Summary of molecular and cellular functions impacted by TBCK loss, based on IPA categorization. p-value ranges reflect the statistical significance of the observed functional disruptions.

Concordant with our RNA-seq results, our pathway enrichment analysis highlighted defects in RNA splicing and ribosomal related proteins (**Fig. 2C**). Notably, pathways associated with autophagy and mitochondrial homeostasis were also enriched. These findings align with previous reports describing autophagic, lysosomal, and mitochondrial abnormalities in *TBCK-/-* cells [7, 8].

To elucidate top canonical pathways disrupted in affected individuals, we employed IPA and identified significant perturbations in pathways related to ‘Processing of Capped Intron-Containing Pre-mRNA’, ‘Intra-Golgi and retrograde Golgi-to-ER traffic’, ‘Major pathway of rRNA processing in the nucleus and cytosol’, and ‘Spliceosomal Cycle’ (**Fig. 2D**). Using IPA’s mapping network tool, our overlay analysis revealed widespread dysregulation across multiple components of the spliceosomal cycle (**Fig. S2D**), further supporting the central role of splicing defects in TBCK pathology. These findings are consistent with our transcriptomic data, in which RNA splicing emerged as the second most affected category (**Fig. 1A**).

We next identified upstream regulators contributing to the observed proteomic changes and obtained several proteins with key roles in cell growth and stress response, including MYC, TP53, EGFR, and TGFB1 (**Fig. 2E**). Of these, MYC, TP53, and TGFB1 also showed dysregulation at the transcript level (**Fig. 1H**), suggesting a consistent perturbation of core regulatory networks across both omics’ layers.

Finally, we examined broader molecular functions impacted by loss of TBCK, and identified ‘RNA post-transcriptional modification’, ‘Cell death & survival’, ‘Protein synthesis’, and ‘Cellular development’ as the most disrupted biological processes (**Fig. 2F**). Our results not only validated the findings of our transcriptomic analysis, but they also revealed a converging dysfunction across multiple levels of regulation at the protein level.

### TBCK loss alters global splicing fidelity and disrupts exon integrity of its own isoforms

To further interrogate the basis of the persistent splicing defects identified in both our transcriptomic and proteomic datasets, we performed a multivariate analysis of alternative splicing using rMATS (*FDR < 0.05; Δ<u>P</u>ercent <u>S</u>pliced In (PSI) > 0.1*). *TBCK-/-* cells exhibited widespread disruption across all major classes of splicing events, including Exon Skipping, Intron Retention, Mutually Exclusive Exons, and Alternative 5’ and 3’ Splice Sites, suggesting a broad impairment of spliceosome function (**Fig. 3A**). Comprehensive annotation of these events revealed several genes with altered exon usage in *TBCK-/-* cells (**Fig. S2C**). GO analysis of differentially spliced transcripts highlighted the enrichment of two main pathways associated with ‘catalytic activity acting on tRNA’ and ‘cytoskeletal anchor activity’ (**Fig. 3B**), as well as ribosomal biogenesis and RNA splicing factor genes (**Fig. 3C-D; Fig. S2E-F**), suggesting a global disruption of RNA maturation processes.

**Figure 3.**
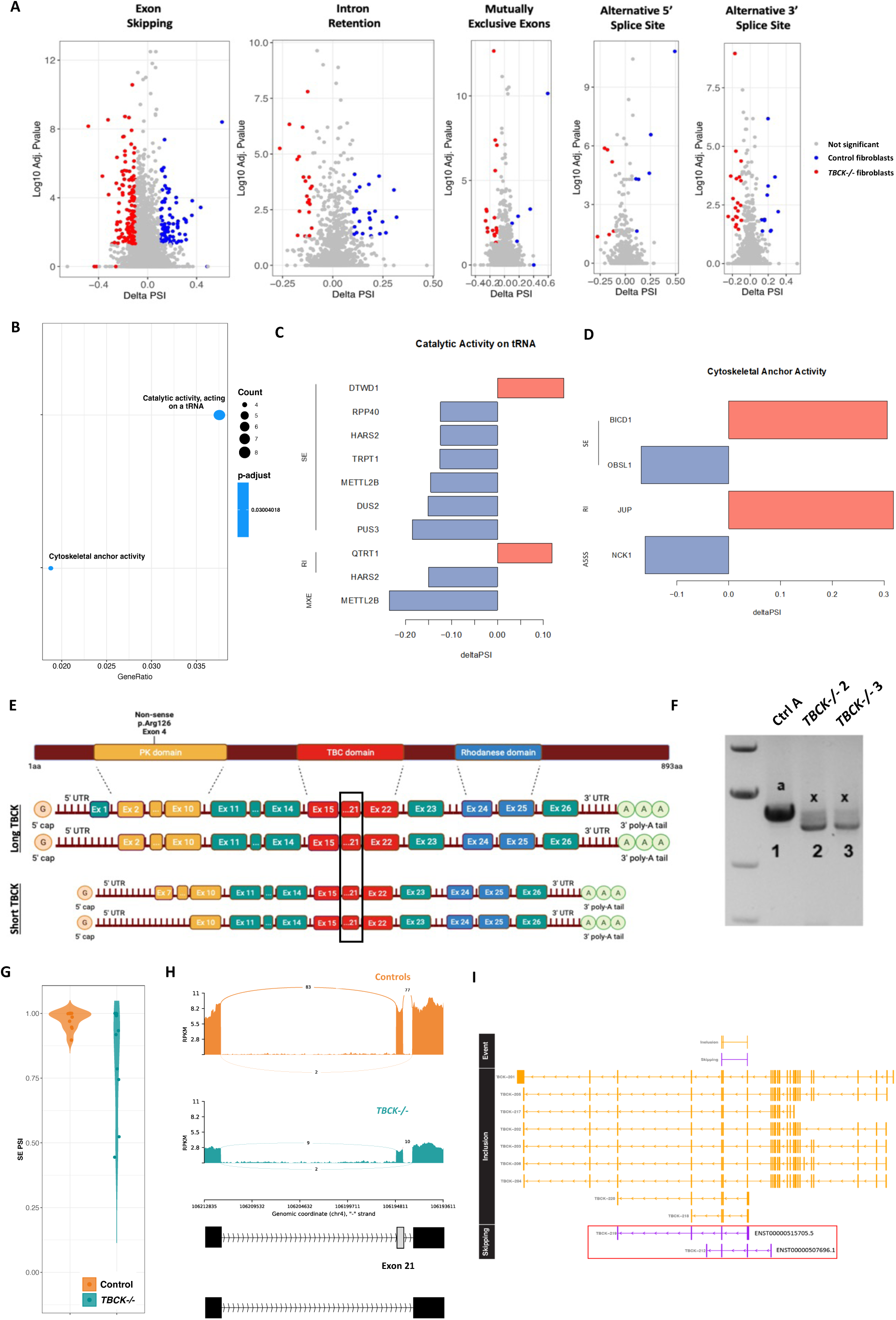
TBCK loss alters global splicing fidelity and disrupts exon integrity of its own isoforms. **A**) Volcano plots representing five major categories of alternative splicing events, revealing significant dysregulated splicing on *TBCK-/-* vs control cells (n = 3 per group; ΔPSI ≥ 0.1, FDR < 0.05). Upregulated events in *TBCK-/-* samples are shown in blue; downregulated in red. **B**) Gene ontology analysis of differentially spliced genes reveals enrichment with dot size reflecting gene count and color intensity adjusted p-value. **C-D**) Bar plots showing representative genes contributing to the top enriched splicing-related pathways, with the direction and magnitude of splicing change indicated by delta PSI values. (SE) – Skipped Exon, (RI) – Retained Intron, (MXE) - Mutual Exclusion and (A5SS) – Alternative 5’ Splice Site **E**) Schematic of TBCK gene structure showing long and short isoforms, annotated with PK, TBC, and Rhodanese domains. Exon 21, more frequently skipped in *TBCK-/-* samples, is in red. **F**) RT-PCR analysis demonstrating the presence of smaller TBCK isoforms in *TBCK-/-* fibroblasts. (a) denotes control sample and (x) *TBCK-/-* individuals **G**) Violin plot showing increased exon skipping in *TBCK-/-* fibroblasts compared to controls. **H**) Sashimi plots depicting read coverage and exon-exon junctions across Exon 21 in control and *TBCK-/-*fibroblasts. Increased exon skipping in *TBCK-/-* samples is supported by junction read counts. **I**) Transcriptome mapping of Exon 21 skipping events to Ensembl TBCK isoforms.

Given the extensive alterations in spliceosomal dynamics, we next asked whether TBCK deficiency could affect the splicing patterns of its own transcripts. The *TBCK* gene encodes multiple annotated isoforms, including both long and short transcripts (**Fig. 3E**) [3]. Even though the full-length canonical isoform of *TBCK* is lost in samples with the p.R126X variant, residual expression of smaller *TBCK* transcripts persists (**Fig. 3F**), raising the possibility of either benign compensatory or dysfunctional poison isoform production. A detailed inspection of the splicing landscape in *TBCK-/-* samples revealed global increased exon skipping relative to controls (**Fig. 3G**). Sashimi plot visualization localized this aberrant splicing to Exon 21, in the functional TBC domain, which was more frequently excluded in *TBCK-/-* samples as compared to controls (**Fig. 3H**).

To determine the biological identity and potential protein-coding capacity of the exon-skipped transcripts, we mapped the splicing junctions to known *TBCK* isoforms. Our analysis uncovered two candidate transcripts: a non-coding isoform with unknown coding sequence (ENST00000515705.5, isoform 219), and a small protein-coding transcript with a partial coding sequence (ENST00000507696.1, isoform 212) (**Fig. 3I**). RT-PCR validation confirmed the presence of isoform 212 in *TBCK-/-* cells, while isoform 219 was undetectable, likely due to its incomplete or non-consensus CDS (**Fig. S2G**). These results indicate that despite loss of full-length *TBCK*, *TBCK-/-* cells continue to produce alternate isoforms with potentially distinct functional consequences.

These results indicate that loss of canonical *TBCK* leads not only to widespread defects in splicing activity but also to exon-level alternate splicing within its own gene. The preferential skipping of Exon 21, located within the TBC domain, underscores a mechanism by which residual *TBCK* transcripts may lose critical functional domains. The emergence of both coding and non-coding isoforms from this splicing disruption highlights the complex interplay between splicing regulation and disease pathology, including this previously unreported mechanism of TBCK regulation. These findings reveal that loss of *TBCK* can impact the alternative splicing of multiple genes, including itself.

### TBCK deficiency alters miRNA expression and impacts key regulatory pathways

MicroRNAs (miRNAs) are short non-coding RNAs that fine-tune gene expression by promoting mRNA degradation or repressing translation, thereby contributing to the robustness of developmental and stress response programs [30]. Increasing evidence suggests that components of the RNA splicing machinery are closely involved in miRNA biogenesis, influencing processes from transcriptional regulation to precursor processing and maturation [31–33]. Given the widespread splicing perturbations observed in *TBCK-/-* cells (**Fig. 3**), we next examined whether these defects also impacted miRNA regulation, which is essential for post-transcriptional gene control.

Rigorous quality control measures were applied to filter out low-confidence reads and sample contaminants based on transcript size distribution, read alignment fidelity, biotype classification, and relative abundance of top-expressed miRNAs (**Fig. S3A–D**). Global miRNA expression levels revealed no differences between control and *TBCK-/-* cells (**Fig. 4A, B**), and sample variability remained low (**Fig. 4C**).

**Figure 4:**
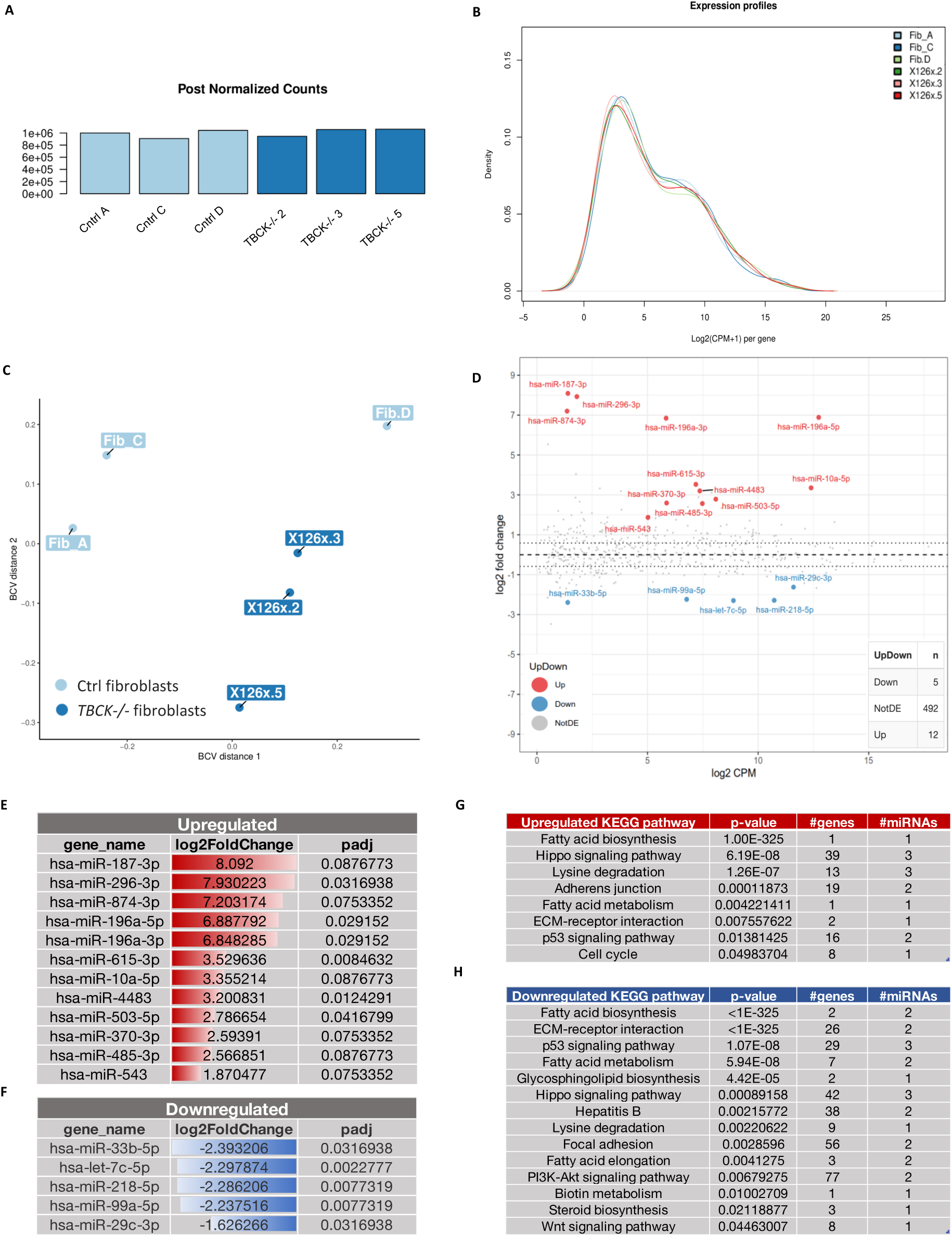
TBCK deficiency alters miRNA expression and impacts key regulatory pathways. **A**) Bar graph showing post-normalized counts of miRNA transcripts in control vs. TBCK-/-fibroblasts, indicating overall miRNA abundance. **B**) Density plot of log₂(CPM + 1) values illustrating the distribution of miRNA expression levels across all samples. X126 nomenclature represents a sample from a TBCK-/- individual. **C**) PCA showing separation between control and TBCK-deficient samples based on global miRNA expression profiles. **D**) MA plot displaying log₂ fold-change (y-axis) versus log₂ counts per million (CPM, x-axis) for each miRNA. Differentially expressed miRNAs (adjusted p-value ≤ 0.1) are color-coded to indicate upregulation or downregulation. **E**) Table summary of significantly upregulated miRNAs (red bars) in TBCK-/- cells, including gene name, log₂ fold-change, and adjusted p-value. **F**) Table summary of significantly downregulated miRNAs (blue bars) with corresponding gene name, log₂ fold-change, and adjusted p-value. **G**) KEGG pathways significantly enriched among targets of upregulated miRNAs, identified using the Pathways Union method in DIANA mirPath v3.0. Columns include pathway name, enrichment p-value, number of genes involved, and number of contributing miRNAs. **H**) Corresponding table of enriched KEGG pathways associated with downregulated miRNAs, also identified via Pathways Union analysis.

Subsequent DE analysis identified 17 miRNAs significantly altered in *TBCK-/-* fibroblasts (fold change ≥1.5; adjusted p < 0.1), comprising 12 upregulated and 5 downregulated (**Fig. 4D–F**). To assess the functional relevance of these dysregulated miRNAs, we performed enrichment analysis using DIANA mirPath v3.0, focusing exclusively on experimentally validated miRNA-target interactions. To evaluate the collective biological impact of multiple dysregulated miRNAs, we applied the Pathways Union method within DIANA mirPath. This approach identifies KEGG pathways significantly enriched by the union of all experimentally validated targets of the selected miRNAs and captures combinatorial regulatory effects that may be missed when evaluating individual miRNAs.

This analysis revealed 35 enriched KEGG pathways (**Fig. S3E**), several of which are directly related to terms that were previously identified in our transcriptomic and proteomic analyses. Interestingly, our results showed that the p53 pathway was enriched in both upregulated and downregulated miRNA targets (**Fig. 4G, H**), mirroring the altered expression of TP53 across our transcriptomic and proteomic datasets (**Figs. 1F, 1H**; **Fig. 2E**). We also observed an impact on several regulatory pathways, including in the PI3K-Akt signaling pathway (77 genes), Hippo signaling (42 genes), and ECM-receptor interaction (26 genes) (**Fig. 4G, 4H**), each of which plays critical roles in neurodevelopment, cellular growth, and structural organization [34–37].

Fatty acid biosynthesis and metabolism pathways were affected as well by both up- and downregulated miRNAs, suggesting altered lipid homeostasis, which has been previously confirmed in *TBCK-/-* individuals [4, 7, 9].

Together, these findings demonstrate that miRNA alterations in *TBCK-/-* cells are not isolated anomalies but rather part of a broader disruption of interconnected signaling networks. This convergence on key developmental, structural, and metabolic pathways highlights the extensive scope of post-transcriptional dysregulation driven by TBCK loss. Our results support a model in which TBCK deficiency perturbs miRNA-mediated regulatory circuits, contributing to the multisystem pathology characteristic of the syndrome.

## Discussion

While the severe progressive neurologic phenotype of individuals with TBCK Syndrome has been well-described, the underlying molecular mechanisms are still poorly understood [2, 4, 7, 9]. This limits our ability to identify and validate disease biomarkers as well as to design and test targeted therapies. To help fill this gap, we applied a comprehensive multi-omics approach to define the molecular landscape of *TBCK-/-* dermal fibroblasts and to uncover new insights into the regulatory roles of TBCK.

Our RNA-seq data revealed a striking downregulation of ribosomal and splicing pathways in *TBCK-/-* cells (**Fig. 1A**), prompting us to further investigate whether these transcriptomic abnormalities also manifested at the protein level. Functional validation through puromycin incorporation assays supported the transcriptomic findings, confirming reduced translation in *TBCK-/-* cells (**Fig. 1D–E**). These results suggest that TBCK plays a previously underappreciated role in regulating ribosome biogenesis and protein synthesis, either by a primary or secondary mechanism. Similar findings were observed using ReNcell neural progenitor cell-derived neurons, in which GO analysis revealed significant enrichment of pathways associated with ribosome biogenesis, rRNA metabolic process, rRNA processing, and eukaryotic translation termination [38] (*bioRxiv)*. These results support the idea that TBCK deficiency can disrupt core components of the translational machinery across different cell types.

Adding to this, recent structural and functional work identified TBCK as a core component of FERRY [17], a Rab5 effector complex that links early endosomes to the translation machinery by facilitating the transport of specific mRNAs and ribosomes within the cell [18]. In this model, TBCK interacts with FERRY subunits to tether mRNAs to endosomal membranes, enabling spatial regulation of translation, particularly for transcripts involved in mitochondrial function and neurodevelopment. Interestingly, our findings expand upon this model by showing that TBCK deficiency not only disrupts transcription at a global level but also affecting the expression of upstream regulators such as LARP1, MYC, TP53, and TGFB1, all of which are known to, directly or indirectly, modulate ribosome biogenesis, translation and stress responses (**Fig. 1F– H**) [39–47]. Likewise, our IPA analysis revealed consistent downregulation of eIF2 and eIF4/p70S6K signaling (**Fig. 1C**), supporting a broader translational repression phenotype.

Beyond the observed defects in translation-related pathways, our data also revealed altered protein levels of MYC, TGFβ, and TP53, mirroring changes seen in our transcriptomic analysis (**Fig. 1F, H**; **Fig. 2E**). MYC is a well-established transcription factor that governs cellular growth, ribosome production, and metabolism [48, 49], while TGFβ plays a central role in cell proliferation, differentiation, and extracellular matrix remodeling [50]. p53 is a key tumor suppressor that mediates stress responses by promoting DNA repair, cell cycle arrest, or apoptosis [51–54]. Ribosomal biogenesis defects typically activate p53, linking translational stress to cell fate regulation [55, 56]. However, in *TBCK-/-* cells, we found significantly reduced p53 levels, suggesting a reversal of this canonical response. This paradox raises the possibility that p53 downregulation may precede ribosomal dysfunction or that an alternate regulatory mechanism is interfering with its expression. One such mechanism may involve the combinatorial effect of dysregulated miRNAs, which could modulate p53 targets or its upstream regulators. This convergence across omics platforms suggests that TBCK loss disrupts not only basal translational machinery but also higher-order regulatory circuits that govern cellular identity and stress responses.

Interestingly, this finding contrasts with the role of TBCK in cancer biology. In renal carcinoma cells, TBCK has been shown to function as an anti-apoptotic factor, where its downregulation by siRNA or by miR-1208 sensitizes tumor cells to cisplatin and TRAIL through caspase-dependent apoptosis [57]. In that context, TBCK promotes cell survival, and its loss facilitates programmed cell death. By contrast, in *TBCK-/-* fibroblasts we observed significantly reduced levels of the tumor suppressor p53, suggesting that TBCK loss in patient cells does not activate canonical apoptotic surveillance pathways but instead blunts them. This divergence underscores the context-dependent role of TBCK: in tumors, high TBCK expression may protect cells from apoptosis, whereas in neurodevelopmental disease, TBCK deficiency disrupts ribosomal and stress-response circuits in a way that suppresses p53 activity. The failure to engage p53-mediated apoptosis in *TBCK-/-* cells may contribute to the accumulation of cellular damage and the progressive degenerative phenotype characteristic of TBCK Syndrome. It is also possible that these findings reflect the specific biology of dermal fibroblasts, and further validation in additional patient-derived systems (e.g., neural progenitors or neurons) will be required to determine whether p53 downregulation is a universal feature of TBCK deficiency. Together, these observations highlight how TBCK loss alters canonical stress pathways in a manner that is distinct from cancer biology. This paradox is further underscored when TBCK Syndrome is considered alongside classical ribosomopathies, where ribosomal stress typically activates, rather than suppresses, p53 signaling.

Given the significant impact of TBCK loss on ribosome biogenesis and translation, it is reasonable to compare TBCK Syndrome with other disorders in the broader category of ribosomopathies, a group of diseases characterized by defects in ribosomal biology and overlapping clinical features. Diamond-Blackfan Anemia (DBA) and Shwachman-Diamond Syndrome (SDS) are rare disorders that similarly exhibit impaired ribosomal function [58–60]. Several molecular features observed in *TBCK-/-* cells mirror those reported in these syndromes. For instance, MYC, which is downregulated in our TBCK model, is also significantly reduced in SDS, where its repression is linked to defective ribosome production and impaired cell proliferation [61]. Moreover, elevated p53 expression, a key indicator of ribosomal stress, is commonly observed in the bone marrow of individuals with SDS and may contribute to hematopoietic dysfunction [62]. In DBA, aberrant activation of the p53 pathway has also been reported as a central disease mechanism [63]. In parallel, dysregulation of the TGFβ pathway, another hallmark of our TBCK data, has been documented in DBA and implicated in its pathophysiology [64].

Together, these overlapping disease features reinforce the idea that TBCK Syndrome shares important pathogenic mechanisms with classical ribosomopathies, particularly those involving MYC repression, p53 dysregulation, and altered TGFβ levels. This alignment may provide a foundation for cross-disease therapeutic insights that has the potential to benefit multiple rare disease communities.

Following our transcriptomic analysis, proteomics offered a crucial layer of validation and expansion, enabling us to determine whether mRNA-level changes translate to functional differences in protein abundance and activity. This step is particularly important when studying post-transcriptional processes like splicing, which are not always predictable from transcript data alone. Proteomic analysis extended our observations by identifying abnormal levels of splicing-related proteins. IPA highlighted disruptions in the processing of ‘capped intron-containing pre-mRNAs’ as the most affected pathway in *TBCK-/-* cells (**Fig. 2D**), indicating that proteins involved in RNA processing are significantly affected in the absence of functional TBCK.

In addition to the well-established effects that TBCK loss has upon mitochondrial function and autophagy [7, 8], both of which were also evident in our transcriptomic and proteomic datasets, we observed an additional impact on Golgi-associated pathways at the protein level (**Fig. 2D**: ‘Intra-Golgi and retrograde Golgi-to-ER traffic’; ‘COPI-mediated anterograde transport’). Even though this change was not as pronounced as those related to RNA splicing, they suggest that Golgi disruption is an additional organelle-level defect in TBCK Syndrome. Although previous work suggested altered ER-Golgi trafficking in human NPCs, the exact function of TBCK in Golgi-ER function remains poorly understood [6]. Notably, TBC1D23, another protein with high TBC-sequence similarity to TBCK, is known to act in the endosome-Golgi trafficking pathway [65].

Our alternative splicing analysis also showed that all major types of splicing events were affected in *TBCK-/-* cells (**Fig. 3A**), which not only matches our proteomic and transcriptomic data but indicates as well that TBCK plays an upstream role in the post-transcriptional regulatory network. This was an interesting discovery, especially as we also saw *TBCK* itself undergoing alternative splicing without its canonical transcript (**Fig. 3E-I**). Isoform-specific analysis identified exon skipping in the TBC domain, particularly on Exon 21, which is believed to harbor TBCK’s catalytic function [3, 10, 66]. We hypothesize that in the absence of functional TBCK, a cascade of stress events sends a signal to the nucleus and disrupts the splicing machinery, leading to altered splicing patterns across multiple transcripts, including *TBCK* itself. Alternatively, if TBCK plays a direct role in regulating splicing, its loss could result in changes to the exon makeup of its own transcript. Another possibility is that this observation arises from the technical limitations of short-read RNA sequencing, which may misrepresent transcript structures due to incomplete alignment or read-through artifacts. In this case, long-read sequencing would be a valuable method for resolving the full-length isoforms and determining the biological relevance of these splicing changes.

Although TBCK is recognized as a predominantly cytoplasmic protein, our data suggest that alterations to this gene can affect splicing, which is a well-known nuclear process. One interesting observation supporting this claim comes from prior reports showing co-localization of TBCK to the nuclei of neurons and myotubes [16, 67], strengthening the idea that TBCK may carry out nuclear functions. In fact, our IPA analysis revealed that loss of TBCK affects many regulatory hubs in the spliceosomal cycle (**Fig. S2D**), which adds to the idea that TBCK may be playing a role in the processing of RNA in the nucleus. However, it is unclear from our data whether this is a primary, or more secondary effect.

Given these widespread splicing perturbations, and the known role of splicing factors in miRNA biogenesis, including those of p53 and MYC in pri-miRNAs production [32, 68–71], we examined whether miRNA expression was also altered in *TBCK-/-* cells. Due to this mechanistic overlap, miRNA profiling became a logical next step to understand how TBCK loss may influence gene regulation at multiple layers. Though no overall differences in total miRNA levels were observed between control and *TBCK-/-* cells, several individual miRNAs were enriched in pathways that echoed those detected in our transcriptomic and proteomic analyses. Some of those were related to cytoskeletal regulation and cell adhesion (**Fig. 1A**: ‘Cell substrate junction’, **Fig. 1C**: ‘RAC signaling’; **Fig. 2D**: ‘mitotic metaphase and anaphase’; **Fig. 3D**: ‘cytoskeletal anchor activity; **Fig. 4G**: ‘Adherens junction’, ‘ECM-receptor interaction’; **Fig. 4H**: ‘ECM-receptor interaction’, ‘Focal adhesion’) aligning with previous studies showing that TBCK influences F-actin structure, proliferation, and cell size [5]. Additionally, we observed dysregulation of miRNAs associated with lipid metabolism, which is consistent with previous reports of lipofuscin accumulation in human tissues and dyslipidemia in *TBCK-/-* individuals [7, 66].

Interestingly, the *p53* pathway was enriched among both upregulated and downregulated miRNA targets. Since ribosomal stress is known to activate *p53* signaling [55, 56], its observed reduction at both transcript and protein levels in TBCK cells is unexpected. This discrepancy raises the possibility that miRNAs may be actively suppressing *p53*, offering a potential mechanism by which TBCK loss alters response signaling.

In this context, our findings reveal significant disruptions of processes not previously associated with the syndrome, such as ribosomal deregulation, splicing, and miRNA expression changes. Interestingly, this layered disruption suggests that many of the previously reported downstream effects of TBCK Syndrome, like autophagy, lysosomal defects, mitochondrial abnormalities, and mTOR pathway dysregulation [2, 7, 8, 72], may stem from earlier upstream disturbances in ribosomal biogenesis, translation, and abnormal splicing. While it is possible that some of these observations reflect downstream consequences or compensatory mechanisms, the consistency across datasets supports the notion that TBCK occupies a more central role in gene regulatory networks than previously recognized. These results should be interpreted with caution, as they may reflect variant-specific consequences of the p.R126X pathogenic allele rather than universal features across all individuals with TBCK Syndrome.

Together, these unbiased findings provide a novel perspective on TBCK-related dysfunction and reveal novel dysregulation of ribosomes, protein synthesis, splicing-related, and miRNA dysregulated networks that play a central role in TBCK pathology. In the search for new therapeutic interventions, our findings provide a roadmap to identify mechanistic nodes that may serve as strategic targets for therapeutic development.

### Conclusions

In summary, our integrative multi-omics analysis reveals that pathogenic variants in *TBCK* can profoundly alter ribosomal homeostasis, splicing fidelity, and miRNA regulatory networks, uncovering a previously unrecognized role for *TBCK* in maintaining translational capacity and post-transcriptional gene regulation. The consistent convergence of transcriptomic, proteomic, splicing, and miRNA perturbations highlights TBCK as a central node in cellular stress and growth pathways, with disruptions in key regulators such as LARP1, MYC, TP53, and TGFB1 offering mechanistic insight into the multisystemic nature of TBCK Syndrome. By identifying both shared and novel pathways of dysfunction, including ribosome biogenesis, spliceosomal integrity, and miRNA-mediated signaling, this work not only expands the molecular framework of TBCK pathology but also provides a roadmap for the discovery of therapeutic targets that could benefit individuals affected by this devastating disorder.

## Methods

### Cell culture

*TBCK-/-* fibroblasts were obtained from arm skin biopsies under IRB-approved protocols (#13278) at the Children’s Hospital of Philadelphia, and control fibroblasts were sourced from the Coriell Institute for Medical Research biobank. Cells were cultured at 37°C with 5% CO₂ in DMEM/High Glucose medium [Cytvia - SH30243] supplemented with 10% FBS.

For each omics dataset included in the study, five biological replicates from TBCK individuals were initially processed, each with three technical replicates. However, two biological replicates were excluded from downstream analysis, one due to confirmed mycoplasma contamination and another due to its classification as an outlier during quality control assessment.

### RNA-seq

Library preparation was performed using the Illumina TruSeq Stranded mRNA Library Prep Kit with paired-end indexing. Sequencing was performed to a depth of ∼25 million reads per sample. The quality of raw reads was assessed using FastQC to ensure high-quality input for alignment. Paired-end reads were aligned to the human GRCh38 transcriptome using kallisto. Illumina adaptors were trimmed using Cutadapt [73]. DESeq2 was used to identify differentially expressed genes (DEGs) between *TBCK-/-* and control fibroblasts [74]. Gene set enrichment analysis (GSEA) was conducted using the clusterProfiler R package [75], focusing on GO Biological Processes. Results were plotted using ggplot2 [76].

### Proteomics

Proteome preparation for mass spectrometry: samples were lysed in a buffer of 8 M urea, 100 mM NaCl, 50 mL Tris-HCl (pH 8) and a cocktail of protease inhibitors (Thermo Fisher Scientific). Samples were subjected to 2 x 30 s of sonication with a Biorupter (Diagenode), 20 min of incubation on ice, followed by 10 min of centrifugation at 21,000 × g and 4°C. Protein content of the supernatant was estimated with using BCA. The proteins were reduced with 5 mM dithiothreitol for 30 min at 55°C and alkylated with 10 mM iodoacetamide in the dark for 30 min at room temperature. The proteins were diluted to 2 M urea with Tris-HCl and digested with sequencing-grade trypsin (Promega) at an enzyme:protein ratio of 1:25 for 12 hr at 37°C. The digestion was quenched with 1% trifluoroacetic acid. Samples were then desalted using Pierce C18 Tips, 10 µL bed (Thermo Scientific). Peptides were brought to 1 μg/3 μl in 0.1% formic acid (buffer A) prior to mass spectrometry acquisition.

Bottom-up nanoLC-MS/MS and data analysis: *TBCK-/-* and control fibroblast samples were analyzed with nanoLC-MS/MS. Peptides were separated using an UltiMate3000 (Dionex) HPLC system (Thermo Fisher Scientific, San Jose, CA, USA) using a 75 μm ID fused capillary pulled in-house and packed with 2.4 μm ReproSil-Pur C18 beads to 20 cm. The HPLC gradient was 0%–35% solvent B (A = 0.1% formic acid, B = 95% acetonitrile, 0.1% formic acid) over 20 min and from 45% to 95% solvent B in 40 min at a flow rate of 300nl/min. The QExactive HF (Thermo Fisher Scientific, San Jose, CA, USA) mass spectrometer was configured following a data-dependent acquisition (DDA) method. All solvents used in analysis of MS samples were LC-MS grade. Full MS scans from 300-1500 m/z were analyzed in the Orbitrap at 120,000 FWHM resolution and 5E5 AGC target value, for 50 ms maximum injection time. Ions were selected for MS2 analysis with an isolation window of 2 m/z, for a maximum injection time of 50 ms and a target AGC of 5E4. MS raw files were processed with Proteome Discoverer version 2.3 and MS spectra were searched against a target + reverse database with the SEQUEST search engine using *H. sapiens* FASTA databases (reviewed, canonical entries, downloaded 9/27/21). For global proteome samples, iBAQ quantification was performed on unique + razor peptides. Trypsin cleavage was specific with up to 2 missed cleavages allowed. Match between runs was enabled but restricted to matches within a single biological replicate by separation of replicates into independent searches. Match between runs parameters included a retention time alignment window of 20 min and a match time window of 0.7 min. False discovery rate (FDR) was set to 0.01.

### Splicing Analysis

Reads were aligned to the human genome (GRCh38) using Spliced Transcripts Alignment to a Reference (STAR) [77]. rMATs v 4.1.2 was used to quantify differential splicing between control and *TBCK-/-* groups [78]. rMATs outputs were processed in R v 4.3.2 using maser [79]. Events were filtered for an average of 10 reads across samples and considered significant with a delta percent spliced in (PSI) > 0.1 and FDR < 0.05. Volcano plots were made using the volcano function in maser. Molecular function GO analysis was performed in R using enrich plot. Sashimi plots were made on rmats2sashimiplot (https://github.com/Xinglab/rmats2sashimiplot), and transcript plots in XI were made using maser function plotTranscripts using the gencode.v29 gtf.

### miRNA-seq

Small RNA isolation from control and *TBCK-/-* dermal fibroblasts was done using the miRNeasy Mini Kit [Cat.No. 217004]. RNA purity was verified using the Thermo Scientific NanoDrop 8000, and library preparation was performed with QIAseq miRNA Library Kit (12) [Cat. No. / ID: 331502]. Sequencing was done with the NextSeq 2000 Illumina system. All samples were processed through adapter and low-quality ends trimming with reads shorter than 25 nucleotides discarded. In the case of paired-end sequencing, both reads of the pair were discarded if at least one was filtered out. Following the Quiaseq protocol, trailing adapter was removed and collapsed over RNA+adapter+UMI, from which adapter+UMI was removed.

We removed rRNA, tRNA, snoRNA, and snRNA as potential contamination, but we did not remove mRNA and other non-coding RNA fragments in small RNA libraries. All collapsed reads were then mapped at the same time to rRNA, tRNA, snRNA, and snoRNA contamination database and microRNA stem loop sequences. Reads mapping with equal or fewer mismatches to the contamination database were removed. For human, microRNA-length reads are filtered to 17-27 nt. For genome mapping of miRNA-seq, we used end-to-end mapping (no soft-clipping at either of the ends) and a maximum of 2 mismatches. Multi mapped reads were not filtered. Differential expression analysis was done using the DIANA mirPath v3.0 Pathways Union.

### PCR and qRT-qPCR

Cells were lysed in TRIzol (Life Technologies; Cat: 15596018), and RNA was extracted using the Direct-zol RNA Miniprep Kit (Zymo Research; Cat: R2050) following the manufacturer’s instructions. 1 µg of total RNA was reverse transcribed with SuperScript™ VILO™ MasterMix (ThermoFisher Scientific; Cat: 11755050) according to the protocol. Reverse transcription reactions were carried out in 20 µL volumes with incubation at 25°C for 10 min, 42°C for 60 min, and 85°C for 5 min. RT-qPCR was conducted on the Quant Studio-3 PCR System using Power-SYBR Green master mix (Applied Biosystems; Cat: 100029284). A melt-curve analysis was performed at the end of each run to confirm the specificity of amplification. The primers employed are listed in Supplementary Table 1. Relative mRNA levels were determined with the 2^−ΔΔCt^ method, using GAPDH as the reference gene.

### Western Blot

Total protein was extracted from *TBCK-/-* fibroblasts (∼2 × 10⁶ cells) using 1X RIPA lysis buffer (Cell Signaling Technologies, #9806S) supplemented with protease and phosphatase inhibitors. Lysates were spun down by centrifugation to remove cellular debris, and protein concentration was quantified using the Pierce™ BCA Protein Assay Kit (Thermo Scientific, #23225) according to the manufacturer’s instructions. Equal amounts of protein (15 μg per sample) were separated by SDS-PAGE on NuPAGE™ 4–12% Bis-Tris Mini Gels (Invitrogen, #NP0321BOX) using 1X MES SDS running buffer (Invitrogen, #11509166) at 180 V for 40 minutes. Proteins were then transferred to PVDF membranes (0.2 μm, Invitrogen, #88520) for 1 hour at 30 V. Membranes were blocked for 30 minutes in 5% non-fat milk in TBS-T and subsequently incubated overnight at 4°C with primary antibodies using the manufacturer recommended concentrations: anti-eIF2α (Santa Cruz Biotechnology D-3 sc-133132), anti-LARP1 (Invitrogen, #PA5-35912), and anti-β-actin (Santa Cruz Biotechnology, #sc-69879).

### Puromycin Assay (SUnSET)

Puromycin incorporation assay (SUnSET) was performed on fibroblasts derived from TBCK individuals and control cells. All cells were serum-starved for 3 hours in DMEM lacking fetal bovine serum (FBS) to synchronize translation activity, followed by stimulation with 10% FBS in the presence of 1 μg/mL puromycin for 30 minutes. Following treatment, cells were lysed in RIPA buffer supplemented with protease inhibitors. Lysates were resolved by SDS-PAGE and transferred to nitrocellulose membranes. Incorporated puromycin was detected by immunoblotting using an anti-puromycin antibody, with vinculin as loading controls. Quantification of the results was done using an unpaired student’s t-test.

### Ingenuity Pathway Analysis

#### Canonical Pathway and DEGs

To identify significantly enriched canonical pathways, a right-tailed Fisher’s Exact Test was used to evaluate the overlap between the supplied dataset and curated pathway gene sets in the Ingenuity Knowledge Base. The values were adjusted for multiple testing using the Benjamini-Hochberg method.

#### Upstream Regulator analysis

Overlap was evaluated between the differentially expressed genes in the dataset and known downstream targets of each upstream molecule in the database. The statistical significance of the overlap was determined using a Fisher’s Exact Test.

#### Biological Processes GO/Functional Enrichment

Enrichment of biological functions and processes was assessed by comparing the proportion of genes involved in a specific biological function using a right-tailed Fisher’s Exact Test. FDR correction using the Benjamini-Hochberg method was applied to account for multiple comparisons.

**Table 1:**
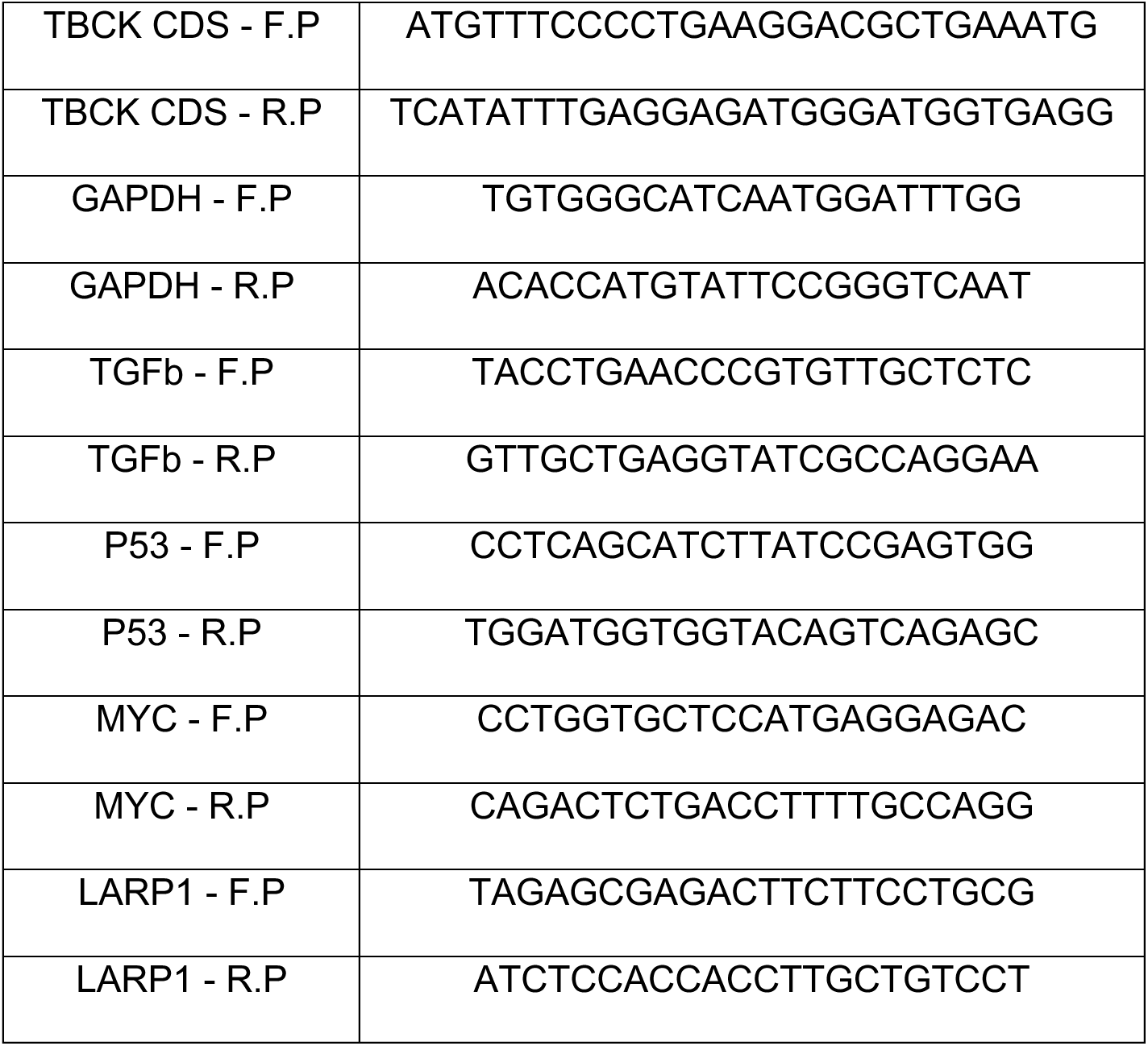
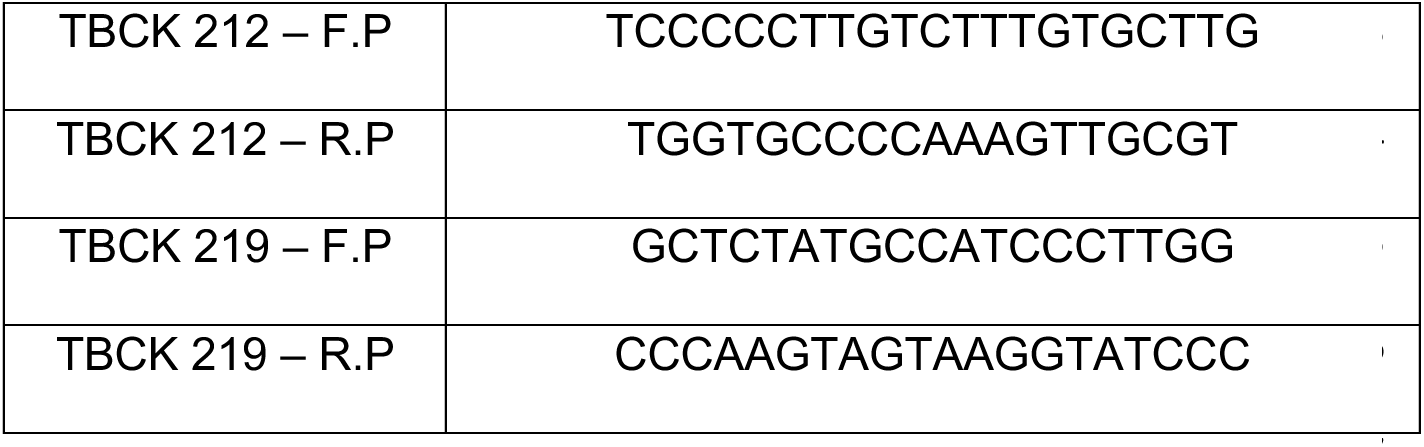
Primers used in the study.

## Supplemental Figures

### Supplemental 1

A. All DE genes (attached file)
B. RNA-seq enriched pathways (attached file)
C. IPA pathways (attached file)
D. Ribosome biogenesis term (attached file)
E. Splicing term (attached file)
F. Samples Genotype

**Supplemental Fig 1:**
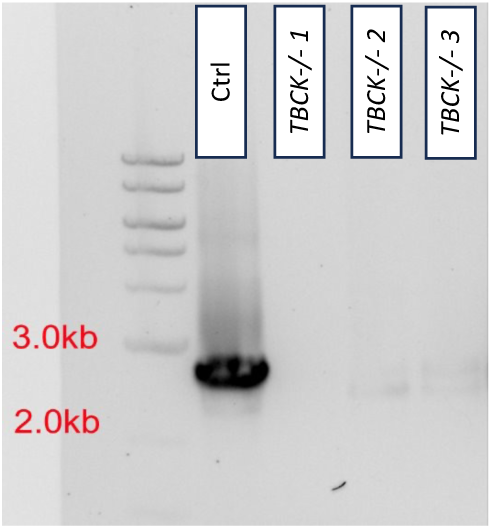
RNA-seq dataset and genotype. **A**) List of all DE genes in *TBCK-/-* fibroblasts vs control. **B**) List of total enrichment analysis with all GO terms. **C**) total IPA pathways detected in RNA-seq dataset. **D-E**) ribosome biogenesis and splicing terms with affected genes **F**) PCR genotype of *TBCK-/-* fibroblasts.

### Supplemental 2

**Supplemental Fig 2:**
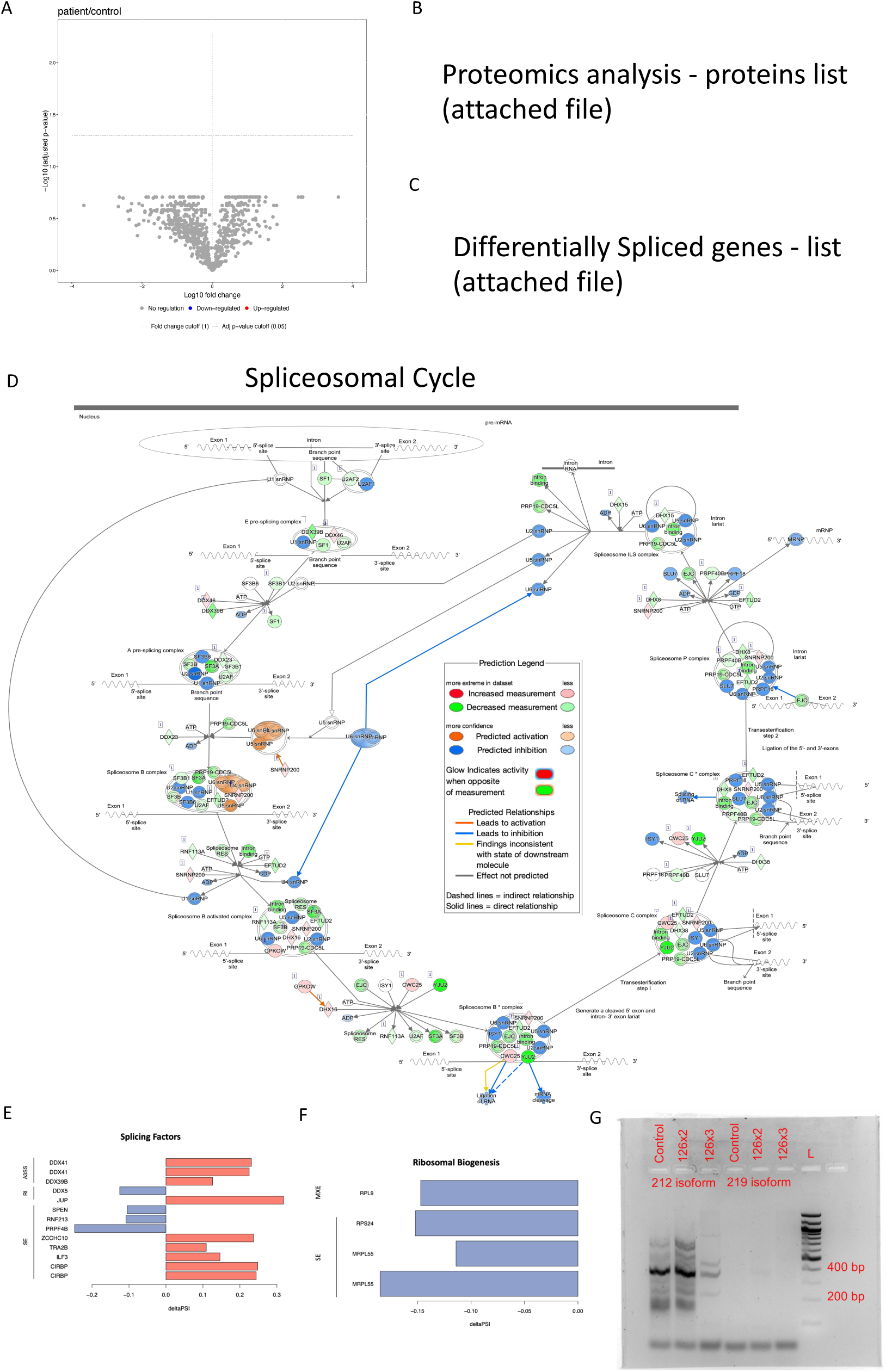
Proteomics Dataset and Splicing Analysis. **A**) volcano plot from proteomic samples showing a DE analysis for control vs *TBCK-/-* samples (FC=1; adj.p-val<0.05). **B**) List of all statistically significant (unadjusted p-val<0.05) proteins from proteomics assay. **C**) total list of maser filtered rMAT events **D**) Network overlay diagram of showing all regulatory nodes of the spliceosome cycle and their relative abundance (green-downregulated in dataset; red-upregulated in dataset). **E**) and **F**) Change in Percent Spliced In (Delta PSI): Illustrating delta PSI values of genes in affected splicing pathways, indicating changes in splicing efficiency of additional terms. **G**) RT-PCR validating the expression of isoforms in *TBCK-/-* samples.

### Supplemental 3

**Supplemental Fig 3:**
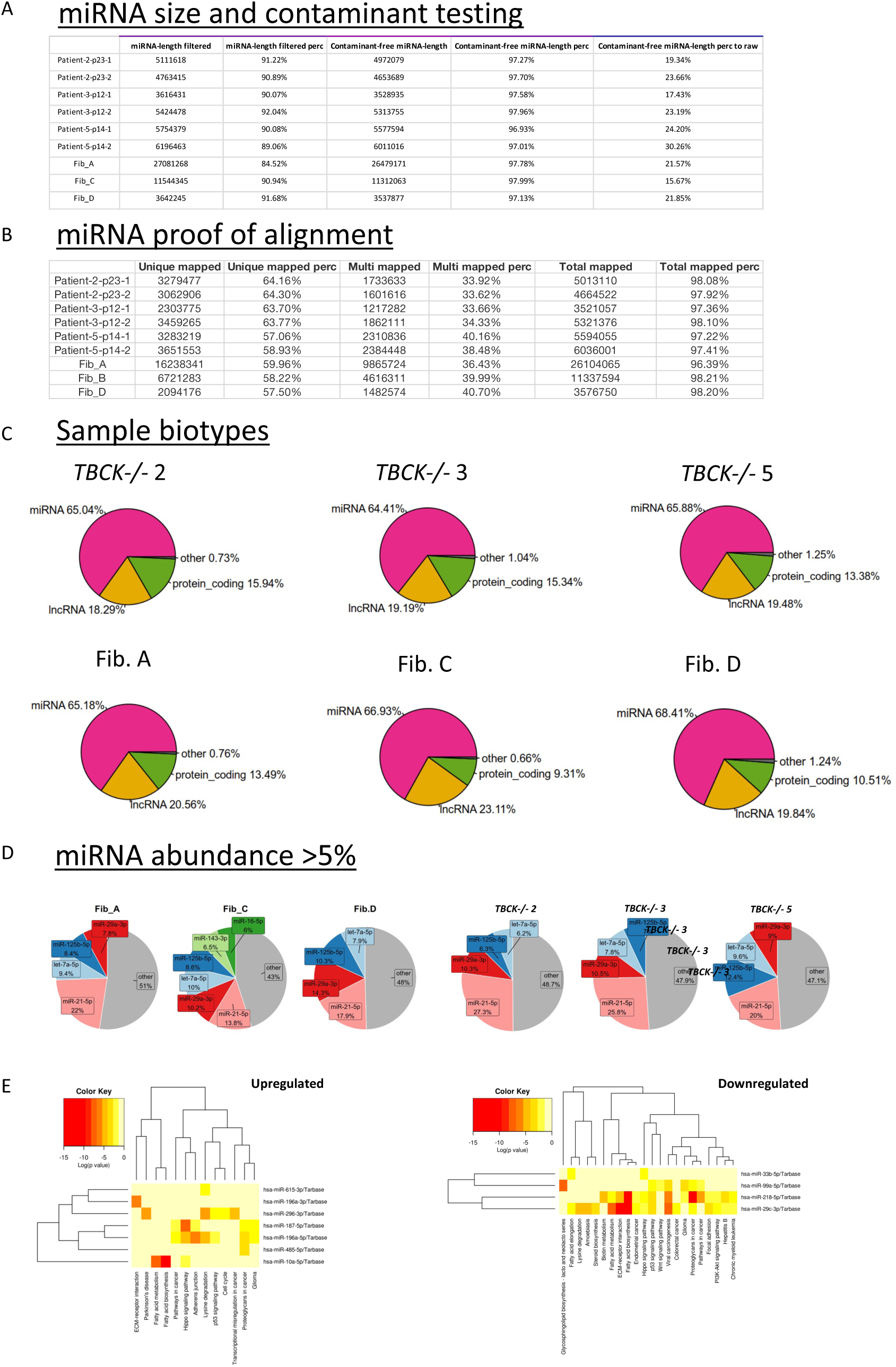
miRNA-seq dataset. **A)** miRNA Size and Contaminant Testing: Tabulated data presenting the total counts of miRNA-length filtered reads, the percentage of contaminant-free miRNA-length reads, and the comparison of contaminant-free miRNA-length reads to raw reads from *TBCK-/-* samples 2, 3, and 5, along with fibroblasts A, C, and D. **B**) miRNA Proof of Alignment: Summary of miRNA sequencing alignment statistics indicating the number and percentage of uniquely mapped reads, multi-mapped reads, and the total mapped reads for each TBCK-/- sample and controls. **C**) Sample Biotypes: Pie charts displaying the distribution of biotypes, including miRNA, protein-coding, lncRNA, and others for *TBCK-/-* samples (*TBCK-/-* 2, *TBCK-/-* 3, *TBCK-/-* 5) and fibroblast controls (Fib. A, Fib. C, Fib. D), demonstrating the proportion of miRNA reads relative to other biotypes. **D**) miRNA Abundance >5%: Pie charts illustrating the relative abundance of miRNAs, with a focus on those constituting more than 5% of the miRNA population in each sample. Each chart represents the distribution of miRNAs for individual *TBCK-/-* samples and controls. **E**). Heatmaps of miRNA Differential Expression: Two heatmaps showing the log fold change of upregulated and downregulated miRNAs across *TBCK-/-* and fibroblast samples. The left heatmap corresponds to miRNAs with increased expression, while the right heatmap shows miRNAs with decreased expression. Color intensity represents the magnitude of log fold change.

## Supporting information

Splicing_term_gene_list

RNAseq_enriched_pathways_go_patient_vs_control

Ribosome_biogenesis_gene_list

IPA_RNAseq_TotalPathways

All_DE_genes_RNAseq_curated

splicing_maser-filtered-rMATs-events

proteomics_analysis_protein_list

## Abbreviations

AGC: Automatic Gain Control
BCA: Bicinchoninic Acid
CDS: Coding Sequence
COPI: Coat Protein Complex I
Ct: Cycle threshold (qPCR)
DDA: Data Dependent Acquisition
DE: Differential Expression
DEGs: Differentially Expressed Genes
DMEM: Dulbecco’s Modified Eagle Medium
DMSO: Dimethyl Sulfoxide
ECM: Extracellular Matrix
EIF2: Eukaryotic Translation Initiation Factor 2
eIF4: Eukaryotic Translation Initiation Factor 4
FBS: Fetal Bovine Serum
FDR: False Discovery Rate
FERRY: Five-subunit Endosomal Rab5 and RNA/ribosome intermediarY complex
FP: Forward Primer
GAPDH: Glyceraldehyde 3-Phosphate Dehydrogenase
GDP: Guanosine Diphosphate
GO: Gene Ontology
GRCh38: Genome Reference Consortium Human Build 38
GSEA: Gene Set Enrichment Analysis
GTP: Guanosine Triphosphate
GTPase: Guanosine Triphosphatase
HPLC: High Performance Liquid Chromatography
iBAQ: Intensity-Based Absolute Quantification
IPA: Ingenuity Pathway Analysis
IRB: Institutional Review Board
KEGG: Kyoto Encyclopedia of Genes and Genomes
LARP1: La-Related Protein 1
LC-MS: Liquid Chromatography–Mass Spectrometry
MES: 2-(N-morpholino) ethanesulfonic acid
MSigDB: Molecular Signatures Database
mTOR: Mechanistic Target of Rapamycin
MYC: MYC Proto-Oncogene
nt: Nucleotide
OMIM: Online Mendelian Inheritance in Man
PCA: Principal Component Analysis
PCR: Polymerase Chain Reaction
PDCD4: Programmed Cell Death Protein 4
PI3K: Phosphoinositide 3-Kinase
PK: Protein Kinase
PSI: Percent Spliced In
PVDF: Polyvinylidene Difluoride
qPCR: Quantitative Polymerase Chain Reaction
R: Rhodanese domain
RNA: Ribonucleic Acid
RNA-seq: RNA Sequencing
rMATS: replicate Multivariate Analysis of Transcript Splicing
RT-PCR: Reverse Transcription Polymerase Chain Reaction
RT-qPCR: Reverse Transcription Quantitative PCR
SDS-PAGE: Sodium Dodecyl Sulfate Polyacrylamide Gel Electrophoresis
SF3B1: Splicing Factor 3b Subunit 1
siRNA: Small Interfering RNA
SUnSET: SUrface SEnsing of Translation
TBC: Tre-2/Bub2/Cdc16 domain
TBCK: TBC1 Domain-Containing Kinase
TBS-T: Tris-Buffered Saline with Tween 20
TGFβ: Transforming Growth Factor Beta
TRAIL: TNF-Related Apoptosis-Inducing Ligand
Tris-HCl: Tris(hydroxymethyl)aminomethane Hydrochloride
UTR: Untranslated Region

## Declarations

### Ethics approval and consent to participate

Patient-derived fibroblasts were obtained under protocols approved by the Institutional Review Board (IRB - #13278) of the Children’s Hospital of Philadelphia. Written informed consent was obtained from all participating families prior to sample collection.

### Consent for publication

Not applicable

### Availability of data and materials

All data generated or analyzed during this study are included in this published article and its supplementary information files.

### Author Contributions

A.D.-R. conceived the study, performed the majority of experiments, analyzed the data, and wrote the manuscript. K.C. collaborated on the analysis of RNA-seq data. R.A. performed RT-qPCR validations of upstream regulators identified through IPA. M.G. collaborated on the proteomics analysis. A.S. contributed to the splicing analysis. J.O. collaborated on the analysis of the miRNA-seq dataset. N.V. provided guidance on troubleshooting miRNA experiments. E.D. contributed intellectual input to the grant supporting the IPA analysis. E.G. assisted with the SUnSET assay for assessing global translation. E.E.L. contributed intellectual input to the grant supporting the IPA analysis. A.M. contributed intellectual input to the grant supporting the IPA analysis. D.L.-C. contributed intellectual input to the grant supporting the IPA analysis. K.A.K. assisted with sample collection and RNA extraction. J.T.-H. collected and provided patient-derived fibroblasts. X.O.-G. collected and provided patient-derived fibroblasts. E.B. supervised the study, provided overall guidance, and contributed to manuscript preparation.

### Funding and Acknowledgements

This work was supported by (5K08NS109281-02). We are deeply grateful to the TBCK Foundation and the families for their continued support and advocacy. We also thank the Ortiz-Gonzalez Lab for generously providing the cells used in this study.

