## Supplementary material for "TBCK Deficiency Alters Ribosomal Function, RNA Splicing, and miRNA Networks: Insights from Multi-Omics Analyses": Splicing_term_gene_list

| baseMean | log2FoldChange | lfcSE | stat | pvalue | padj |
|---|---|---|---|---|---|
| 1.834342428 | -0.171793648 | 0.720777534 | -0.238344898 | 0.811613601 | 0.000671936 |
| 57.94839011 | 0.02408886 | 0.41728538 | 0.057727544 | 0.953965653 | 0.001992167 |
| 5.56425208 | 0.319538411 | 0.447503616 | 0.714046545 | 0.475198398 | 0.002707011 |
| 9.897606838 | -0.093461314 | 0.417184127 | -0.224028932 | 0.822734791 | 0.005120489 |
| 151.2837819 | 0.434180527 | 0.175186627 | 2.478388531 | 0.013197734 | 0.010856995 |
| 12.20376829 | -0.193002522 | 0.335408762 | -0.575424807 | 0.565004031 | 0.012706 |
| 1.935763322 | -1.233193526 | 0.677076444 | -1.821350509 | 0.068553593 | 0.013405636 |
| 0.149519532 | 0.937570085 | 2.429288757 | 0.385944274 | 0.69953794 | 0.016495126 |
| 5.616693207 | -0.631019855 | 0.456614966 | -1.381951759 | 0.166986514 | 0.031838972 |
| 15.90716194 | -0.645979717 | 0.416135129 | -1.552331615 | 0.120582897 | 0.032838242 |
| 0.731452833 | -0.070076099 | 1.117597795 | -0.062702431 | 0.950003461 | 0.039347008 |
| 2.364193509 | 0.085295029 | 0.575551496 | 0.148197041 | 0.882187267 | 0.041327614 |
| 0.999366626 | 0.463371045 | 0.910331216 | 0.509013683 | 0.610742633 | 0.045822086 |
| 4.268410856 | -0.152922675 | 0.440875432 | -0.346861413 | 0.728695437 | 0.047622276 |

| ensg | gene |
|---|---|
| ENSG00000003756 | RBM5 |
| ENSG00000004487 | KDM1A |
| ENSG00000004534 | RBM6 |
| ENSG00000005007 | UPF1 |
| ENSG00000005436 | GCFC2 |
| ENSG00000006712 | PAF1 |
| ENSG00000007392 | LUC7L |
| ENSG00000008128 | CDK11A |
| ENSG00000011243 | AKAP8L |
| ENSG00000011304 | PTBP1 |
| ENSG00000013441 | CLK1 |
| ENSG00000014164 | ZC3H3 |
| ENSG00000021776 | AQR |
| ENSG00000023734 | STRAP |
