## Supplementary material for "TBCK Deficiency Alters Ribosomal Function, RNA Splicing, and miRNA Networks: Insights from Multi-Omics Analyses": Ribosome_biogenesis_gene_list

| baseMean | log2FoldChange | lfcSE | stat | pvalue |
|---|---|---|---|---|
| 733.512007 | 1.06773697 | 0.14859448 | 7.18557644 | 6.69E-13 |
| 43.3273678 | -1.145458 | 0.21920649 | -5.2254746 | 1.74E-07 |
| 165.240762 | -0.9306643 | 0.18336411 | -5.0754985 | 3.86E-07 |
| 58.9696581 | -0.9542205 | 0.20016921 | -4.7670692 | 1.87E-06 |
| 46.6366389 | -0.9012922 | 0.19226264 | -4.687818 | 2.76E-06 |
| 15.1789441 | -1.2129795 | 0.26704587 | -4.5422142 | 5.57E-06 |
| 87.4370064 | 0.80328704 | 0.18296447 | 4.39039906 | 1.13E-05 |
| 3.50924945 | 2.60105181 | 0.62045351 | 4.19217843 | 2.76E-05 |
| 18.3830134 | -1.1925152 | 0.28679265 | -4.1581094 | 3.21E-05 |
| 15.3188823 | -1.2526916 | 0.31392922 | -3.9903633 | 6.60E-05 |
| 236.151439 | 0.81705384 | 0.20617257 | 3.96296099 | 7.40E-05 |
| 25.2471506 | -1.1195197 | 0.28720787 | -3.8979421 | 9.70E-05 |
| 38.9650518 | -0.715515 | 0.1841276 | -3.8859737 | 0.00010192 |
| 28.8239801 | -1.4633501 | 0.38272512 | -3.8235016 | 0.00013157 |
| 58.1564938 | -1.0948082 | 0.28971172 | -3.7789573 | 0.00015749 |
| 93.8920765 | 0.98081988 | 0.26080114 | 3.76079591 | 0.00016937 |
| 21.3269755 | -0.9479506 | 0.2568605 | -3.6905268 | 0.00022379 |
| 218.233175 | 0.55822784 | 0.15187375 | 3.67560444 | 0.00023729 |
| 206.342439 | -1.6712468 | 0.45761985 | -3.6520419 | 0.00026016 |
| 23.765833 | -1.1667095 | 0.32443576 | -3.5961187 | 0.000323 |
| 14.6997167 | -1.3459584 | 0.37824611 | -3.5584197 | 0.00037309 |
| 21.6434426 | -1.08055 | 0.30796724 | -3.5086523 | 0.00045038 |
| 46.0277023 | -0.7037506 | 0.20376373 | -3.4537578 | 0.00055283 |
| 1.71275759 | -2.9524314 | 0.86312673 | -3.4206233 | 0.00062478 |
| 20.9175705 | -0.9670426 | 0.29197038 | -3.3121259 | 0.0009259 |
| 43.3310679 | -0.7711244 | 0.23525848 | -3.277775 | 0.00104629 |
| 28.8595601 | -1.0019572 | 0.30685632 | -3.2652324 | 0.00109374 |
| 197.521326 | 0.44175174 | 0.1353019 | 3.26493372 | 0.0010949 |
| 19.1793454 | 1.06644359 | 0.3297433 | 3.23416302 | 0.00122 |
| 8956.85788 | 0.74414909 | 0.23014113 | 3.23344669 | 0.00122306 |
| 51.783071 | 0.66611202 | 0.20651061 | 3.22555829 | 0.00125727 |
| 1.59881952 | 2.63871784 | 0.82120803 | 3.21321486 | 0.00131258 |
| 31.136532 | -0.9457474 | 0.29527286 | -3.2029607 | 0.00136023 |
| 3.61087597 | -1.7370142 | 0.54804522 | -3.1694725 | 0.00152716 |
| 43.9304658 | -1.1385215 | 0.36018366 | -3.160947 | 0.00157257 |
| 2.03058764 | -2.5770652 | 0.81897243 | -3.1467057 | 0.00165121 |
| 1.68008847 | -2.6119164 | 0.84444328 | -3.0930632 | 0.00198102 |
| 75.7657763 | -0.5576852 | 0.18148968 | -3.0728207 | 0.00212046 |
| 2.84411508 | -2.1971318 | 0.7167781 | -3.0652887 | 0.0021746 |
| 528.02067 | -0.2676271 | 0.08765541 | -3.0531728 | 0.00226435 |
| 2.96618797 | -1.7751641 | 0.583356 | -3.0430202 | 0.00234217 |
| 11.042606 | -1.2694953 | 0.41827932 | -3.0350419 | 0.00240502 |

| | | | | |
|---|---|---|---|---|
| 20.6583058 | -0.792365 | 0.26379032 | -3.0037682 | 0.00266658 |
| 9.10927571 | -1.2802978 | 0.42774525 | -2.9931315 | 0.00276131 |
| 15.0871029 | -1.117458 | 0.37618669 | -2.9704879 | 0.00297327 |
| 71.4319226 | -0.6146497 | 0.20722657 | -2.9660759 | 0.00301626 |
| 19.8560428 | -0.8418189 | 0.28469924 | -2.9568709 | 0.00310778 |
| 15.4433011 | -1.1695722 | 0.39560424 | -2.9564197 | 0.00311233 |
| 14.9062379 | -0.8081861 | 0.27496016 | -2.9392844 | 0.00328971 |
| 7.2404271 | -1.2041715 | 0.41315669 | -2.9145637 | 0.00356186 |
| 2.76536318 | -1.7851171 | 0.61701076 | -2.8931701 | 0.00381375 |
| 7.27612877 | -1.0020734 | 0.34911599 | -2.8703164 | 0.00410061 |
| 11.2093012 | -0.8954685 | 0.31256792 | -2.8648765 | 0.00417172 |

| padj | ensg | gene |
|---|---|---|
| 7.26E-10 | ENSG00000122085 | MTERF4 |
| 2.32E-05 | ENSG00000148303 | RPL7A |
| 4.59E-05 | ENSG00000179218 | CALR |
| 0.00016911 | ENSG00000108298 | RPL19 |
| 0.00022773 | ENSG00000118181 | RPS25 |
| 0.00039235 | ENSG00000170889 | RPS9 |
| 0.00067194 | ENSG00000135250 | SRPK2 |
| 0.0013077 | ENSG00000125630 | POLR1B |
| 0.00143549 | ENSG00000136942 | RPL35 |
| 0.00254707 | ENSG00000161970 | RPL26 |
| 0.00276128 | ENSG00000170854 | RIOX2 |
| 0.0033507 | ENSG00000197756 | RPL37A |
| 0.00345135 | ENSG00000125457 | MIF4GD |
| 0.0041864 | ENSG00000063177 | RPL18 |
| 0.00485804 | ENSG00000140988 | RPS2 |
| 0.00512049 | ENSG00000164985 | PSIP1 |
| 0.0063388 | ENSG00000188846 | RPL14 |
| 0.00662818 | ENSG00000177971 | IMP3 |
| 0.00699326 | ENSG00000188976 | NOC2L |
| 0.00831366 | ENSG00000123144 | TRIR |
| 0.0091931 | ENSG00000083845 | RPS5 |
| 0.01060193 | ENSG00000164587 | RPS14 |
| 0.01236532 | ENSG00000100316 | RPL3 |
| 0.01340564 | ENSG00000105248 | YJU2 |
| 0.01745672 | ENSG00000162244 | RPL29 |
| 0.01924757 | ENSG00000142541 | RPL13A |
| 0.01993008 | ENSG00000161016 | RPL8 |
| 0.01993008 | ENSG00000167721 | TSR1 |
| 0.02152844 | ENSG00000148824 | MTG1 |
| 0.02155102 | ENSG00000149716 | LTO1 |
| 0.0218993 | ENSG00000104131 | EIF3J |
| 0.02250695 | ENSG00000058729 | RIOK2 |
| 0.02303075 | ENSG00000105372 | RPS19 |
| 0.02508248 | ENSG00000168028 | RPSA |
| 0.02555533 | ENSG00000142937 | RPS8 |
| 0.02651341 | ENSG00000112578 | BYSL |
| 0.03001988 | ENSG00000173141 | MRPL57 |
| 0.03138615 | ENSG00000166441 | RPL27A |
| 0.03183897 | ENSG00000183431 | SF3A3 |
| 0.03271844 | ENSG00000108592 | FTSJ3 |
| 0.03344414 | ENSG00000173545 | ZNF622 |
| 0.03414062 | ENSG00000198755 | RPL10A |

| | | |
|---|---|---|
| 0.03702978 | ENSG00000231500 | RPS18 |
| 0.03778253 | ENSG00000105193 | RPS16 |
| 0.03974086 | ENSG00000105373 | NOP53 |
| 0.04009087 | ENSG00000130255 | RPL36 |
| 0.04117557 | ENSG00000229117 | RPL41 |
| 0.04119084 | ENSG00000149273 | RPS3 |
| 0.04251723 | ENSG00000142676 | RPL11 |
| 0.0445052 | ENSG00000145592 | RPL37 |
| 0.04665393 | ENSG00000116251 | RPL22 |
| 0.04951367 | ENSG00000136718 | IMP4 |
| 0.04991971 | ENSG00000145425 | RPS3A |
