## Supplementary material for "TBCK Deficiency Alters Ribosomal Function, RNA Splicing, and miRNA Networks: Insights from Multi-Omics Analyses": IPA_RNAseq_TotalPathways

¬© 2000-2023 QIAGEN. All rights reserved.

| Ingenuity Canonical Pathways | -log(p-value) |
|---|---|
| EIF2 Signaling | 1.35E+01 |
| Regulation of eIF4 and p70S6K Signaling | 7.88E+00 |
| Pulmonary Fibrosis Idiopathic Signaling Pathway | 6.96E+00 |
| mTOR Signaling | 6.66E+00 |
| Coronavirus Pathogenesis Pathway | 5.83E+00 |
| Oncostatin M Signaling | 5.28E+00 |
| Role of Osteoclasts in Rheumatoid Arthritis Signaling Pathway | 4.94E+00 |
| Pulmonary Healing Signaling Pathway | 4.86E+00 |
| RAC Signaling | 4.66E+00 |
| Epithelial Adherens Junction Signaling | 4.50E+00 |
| Osteoarthritis Pathway | 4.33E+00 |
| Axonal Guidance Signaling | 4.25E+00 |
| Hepatic Fibrosis / Hepatic Stellate Cell Activation | 3.95E+00 |
| Microautophagy Signaling Pathway | 3.88E+00 |
| PTEN Signaling | 3.61E+00 |
| LPS-stimulated MAPK Signaling | 3.55E+00 |
| Wound Healing Signaling Pathway | 3.49E+00 |
| NGF Signaling | 3.47E+00 |
| Mouse Embryonic Stem Cell Pluripotency | 3.44E+00 |
| IGF-1 Signaling | 3.40E+00 |
| BMP signaling pathway | 3.31E+00 |
| Human Embryonic Stem Cell Pluripotency | 3.28E+00 |
| Ephrin Receptor Signaling | 3.26E+00 |
| Sertoli Cell-Sertoli Cell Junction Signaling | 3.25E+00 |
| Actin Nucleation by ARP-WASP Complex | 3.23E+00 |
| Role of Macrophages, Fibroblasts and Endothelial Cells in Rheumatoid Arthritis | 3.17E+00 |
| Hepatic Fibrosis Signaling Pathway | 3.14E+00 |
| Prostate Cancer Signaling | 3.10E+00 |
| GŒ±12/13 Signaling | 3.07E+00 |
| Leukocyte Extravasation Signaling | 3.01E+00 |
| IL-10 Signaling | 3.01E+00 |
| PEDF Signaling | 2.95E+00 |
| Inhibition of Matrix Metalloproteases | 2.94E+00 |
| MSP-RON Signaling in Cancer Cells Pathway | 2.88E+00 |
| Tumor Microenvironment Pathway | 2.87E+00 |
| PI3K/AKT Signaling | 2.85E+00 |
| Xenobiotic Metabolism General Signaling Pathway | 2.80E+00 |
| Signaling by Rho Family GTPases | 2.78E+00 |
| FAT10 Signaling Pathway | 2.75E+00 |

| | |
|---|---|
| IL-8 Signaling | 2.65E+00 |
| Angiopoietin Signaling | 2.63E+00 |
| GDNF Family Ligand-Receptor Interactions | 2.63E+00 |
| Toll-like Receptor Signaling | 2.56E+00 |
| Neurotrophin/TRK Signaling | 2.56E+00 |
| TNFR2 Signaling | 2.56E+00 |
| Antioxidant Action of Vitamin C | 2.56E+00 |
| Thyroid Cancer Signaling | 2.53E+00 |
| Activin Inhibin Signaling Pathway | 2.51E+00 |
| Bladder Cancer Signaling | 2.50E+00 |
| Virus Entry via Endocytic Pathways | 2.45E+00 |
| 4-1BB Signaling in T Lymphocytes | 2.44E+00 |
| MSP-RON Signaling in Macrophages Pathway | 2.42E+00 |
| Activation of IRF by Cytosolic Pattern Recognition Receptors | 2.42E+00 |
| Role of Chondrocytes in Rheumatoid Arthritis Signaling Pathway | 2.36E+00 |
| Sirtuin Signaling Pathway | 2.36E+00 |
| Apoptosis Signaling | 2.32E+00 |
| Acute Phase Response Signaling | 2.31E+00 |
| Agrin Interactions at Neuromuscular Junction | 2.28E+00 |
| IL-17A Signaling in Fibroblasts | 2.27E+00 |
| GP6 Signaling Pathway | 2.22E+00 |
| Germ Cell-Sertoli Cell Junction Signaling | 2.18E+00 |
| Regulation of Cellular Mechanics by Calpain Protease | 2.18E+00 |
| Antigen Presentation Pathway | 2.18E+00 |
| Sertoli Cell Germ Cell Junction Signaling Pathway (Enhanced) | 2.18E+00 |
| Glioma Invasiveness Signaling | 2.15E+00 |
| Regulation of the Epithelial-Mesenchymal Transition Pathway | 2.12E+00 |
| HGF Signaling | 2.11E+00 |
| Factors Promoting Cardiogenesis in Vertebrates | 2.10E+00 |
| CD27 Signaling in Lymphocytes | 2.10E+00 |
| Role of IL-17A in Arthritis | 2.10E+00 |
| April Mediated Signaling | 2.04E+00 |
| Multiple Sclerosis Signaling Pathway | 2.04E+00 |
| TGF-Œ≤ Signaling | 2.02E+00 |
| B Cell Activating Factor Signaling | 2.00E+00 |
| NF-Œ∫B Activation by Viruses | 1.99E+00 |
| Colorectal Cancer Metastasis Signaling | 1.99E+00 |
| Endometrial Cancer Signaling | 1.99E+00 |
| PCP (Planar Cell Polarity) Pathway | 1.99E+00 |
| Renal Cell Carcinoma Signaling | 1.96E+00 |
| Protein Ubiquitination Pathway | 1.96E+00 |

| | |
|---|---|
| Semaphorin Signaling in Neurons | 1.96E+00 |
| Role of Osteoblasts, Osteoclasts and Chondrocytes in Rheumatoid Arthritis | 1.94E+00 |
| Senescence Pathway | 1.93E+00 |
| Parkinson's Signaling | 1.92E+00 |
| Inhibition of ARE-Mediated mRNA Degradation Pathway | 1.91E+00 |
| HOTAIR Regulatory Pathway | 1.91E+00 |
| Chemokine Signaling | 1.91E+00 |
| Molecular Mechanisms of Cancer | 1.89E+00 |
| Transcriptional Regulatory Network in Embryonic Stem Cells | 1.89E+00 |
| Myelination Signaling Pathway | 1.88E+00 |
| Glucocorticoid Receptor Signaling | 1.86E+00 |
| iNOS Signaling | 1.84E+00 |
| Integrin Signaling | 1.84E+00 |
| Role of NANOG in Mammalian Embryonic Stem Cell Pluripotency | 1.84E+00 |
| Huntington's Disease Signaling | 1.81E+00 |
| BAG2 Signaling Pathway | 1.80E+00 |
| 14-3-3-mediated Signaling | 1.77E+00 |
| IL-17A Signaling in Airway Cells | 1.77E+00 |
| Paxillin Signaling | 1.75E+00 |
| PPAR Signaling | 1.75E+00 |
| PDGF Signaling | 1.75E+00 |
| Regulation of the Epithelial Mesenchymal Transition in Development Pathway | 1.75E+00 |
| G Beta Gamma Signaling | 1.73E+00 |
| IL-6 Signaling | 1.73E+00 |
| Melanoma Signaling | 1.73E+00 |
| FAT10 Cancer Signaling Pathway | 1.73E+00 |
| D-myo-inositol-5-phosphate Metabolism | 1.73E+00 |
| WNT/Œ≤-catenin Signaling | 1.72E+00 |
| Actin Cytoskeleton Signaling | 1.71E+00 |
| Neuroinflammation Signaling Pathway | 1.71E+00 |
| TNFR1 Signaling | 1.70E+00 |
| UVC-Induced MAPK Signaling | 1.70E+00 |
| Ceramide Signaling | 1.65E+00 |
| RANK Signaling in Osteoclasts | 1.65E+00 |
| ID1 Signaling Pathway | 1.65E+00 |
| Thrombin Signaling | 1.65E+00 |
| Ephrin B Signaling | 1.63E+00 |
| Ovarian Cancer Signaling | 1.61E+00 |
| ERBB Signaling | 1.61E+00 |
| RAR Activation | 1.59E+00 |
| TWEAK Signaling | 1.58E+00 |

| | |
|---|---|
| ERK5 Signaling | 1.58E+00 |
| Role of Tissue Factor in Cancer | 1.57E+00 |
| PAK Signaling | 1.55E+00 |
| Macropinocytosis Signaling | 1.53E+00 |
| Cardiac Hypertrophy Signaling (Enhanced) | 1.52E+00 |
| Cholecystokinin/Gastrin-mediated Signaling | 1.51E+00 |
| UVA-Induced MAPK Signaling | 1.50E+00 |
| Vitamin-C Transport | 1.49E+00 |
| NRF2-mediated Oxidative Stress Response | 1.49E+00 |
| Cancer Drug Resistance by Drug Efflux | 1.48E+00 |
| Serotonin Receptor Signaling | 1.46E+00 |
| GNRH Signaling | 1.46E+00 |
| Production of Nitric Oxide and Reactive Oxygen Species in Macrophages | 1.46E+00 |
| ERK/MAPK Signaling | 1.46E+00 |
| CXCR4 Signaling | 1.45E+00 |
| Synaptogenesis Signaling Pathway | 1.45E+00 |
| Regulation of the Epithelial Mesenchymal Transition by Growth Factors Pathway | 1.44E+00 |
| Autophagy | 1.43E+00 |
| RHOA Signaling | 1.42E+00 |
| Xenobiotic Metabolism Signaling | 1.40E+00 |
| Sumoylation Pathway | 1.40E+00 |
| RHOGDI Signaling | 1.39E+00 |
| FLT3 Signaling in Hematopoietic Progenitor Cells | 1.39E+00 |
| Estrogen-Dependent Breast Cancer Signaling | 1.39E+00 |
| JAK/STAT Signaling | 1.39E+00 |
| BEX2 Signaling Pathway | 1.39E+00 |
| IL-2 Signaling | 1.38E+00 |
| Semaphorin Neuronal Repulsive Signaling Pathway | 1.37E+00 |
| Purine Nucleotides De Novo Biosynthesis II | 1.36E+00 |
| Natural Killer Cell Signaling | 1.36E+00 |
| IL-17A Signaling in Gastric Cells | 1.35E+00 |
| VEGF Family Ligand-Receptor Interactions | 1.35E+00 |
| MIF Regulation of Innate Immunity | 1.34E+00 |
| ILK Signaling | 1.32E+00 |
| HER-2 Signaling in Breast Cancer | 1.31E+00 |
| HIPPO signaling | 1.31E+00 |
| Induction of Apoptosis by HIV1 | 1.30E+00 |
| ERB2-ERBB3 Signaling | 1.30E+00 |
| Pyridoxal 5'-phosphate Salvage Pathway | 1.30E+00 |
| fMLP Signaling in Neutrophils | 1.30E+00 |
| Chronic Myeloid Leukemia Signaling | 1.30E+00 |

| | |
|---|---|
| Xenobiotic Metabolism AHR Signaling Pathway | 1.29E+00 |
| Opioid Signaling Pathway | 1.28E+00 |
| D-myo-inositol (1,4,5,6)-Tetrakisphosphate Biosynthesis | 1.28E+00 |
| D-myo-inositol (3,4,5,6)-tetrakisphosphate Biosynthesis | 1.28E+00 |
| WNT/Ca+ pathway | 1.28E+00 |
| Role of RIG1-like Receptors in Antiviral Innate Immunity | 1.28E+00 |
| Protein Kinase A Signaling | 1.28E+00 |
| 3-phosphoinositide Biosynthesis | 1.26E+00 |
| CD40 Signaling | 1.26E+00 |
| Mitotic Roles of Polo-Like Kinase | 1.26E+00 |
| Role of IL-17F in Allergic Inflammatory Airway Diseases | 1.25E+00 |
| STAT3 Pathway | 1.24E+00 |
| HIF1Œ± Signaling | 1.24E+00 |
| ERBB4 Signaling | 1.23E+00 |
| Superpathway of Inositol Phosphate Compounds | 1.23E+00 |
| Lipoate Biosynthesis and Incorporation II | 1.22E+00 |
| IL-33 Signaling Pathway | 1.22E+00 |
| SPINK1 General Cancer Pathway | 1.21E+00 |
| Cardiac Hypertrophy Signaling | 1.21E+00 |
| Th2 Pathway | 1.21E+00 |
| Acute Myeloid Leukemia Signaling | 1.21E+00 |
| Role of MAPK Signaling in Promoting the Pathogenesis of Influenza | 1.20E+00 |
| Reelin Signaling in Neurons | 1.20E+00 |
| Regulation of Actin-based Motility by Rho | 1.19E+00 |
| CDK5 Signaling | 1.19E+00 |
| Urate Biosynthesis/Inosine 5'-phosphate Degradation | 1.17E+00 |
| Colanic Acid Building Blocks Biosynthesis | 1.17E+00 |
| G-Protein Coupled Receptor Signaling | 1.16E+00 |
| Methylglyoxal Degradation III | 1.16E+00 |
| Neuregulin Signaling | 1.16E+00 |
| FcŒ≥ Receptor-mediated Phagocytosis in Macrophages and Monocytes | 1.15E+00 |
| Apelin Endothelial Signaling Pathway | 1.15E+00 |
| 3-phosphoinositide Degradation | 1.14E+00 |
| Fc Epsilon RI Signaling | 1.14E+00 |
| Role of Osteoblasts in Rheumatoid Arthritis Signaling Pathway | 1.12E+00 |
| IL-1 Signaling | 1.12E+00 |
| Death Receptor Signaling | 1.12E+00 |
| Assembly of RNA Polymerase III Complex | 1.12E+00 |
| PPARŒ±/RXRŒ± Activation | 1.11E+00 |
| Salvage Pathways of Pyrimidine Ribonucleotides | 1.10E+00 |
| Renin-Angiotensin Signaling | 1.10E+00 |

| | |
|---|---|
| Caveolar-mediated Endocytosis Signaling | 1.09E+00 |
| Glioblastoma Multiforme Signaling | 1.08E+00 |
| Hypoxia Signaling in the Cardiovascular System | 1.07E+00 |
| VEGF Signaling | 1.07E+00 |
| ATM Signaling | 1.05E+00 |
| Lymphotoxin Œ≤ Receptor Signaling | 1.05E+00 |
| Ascorbate Recycling (Cytosolic) | 1.05E+00 |
| Tetrahydrobiopterin Biosynthesis I | 1.05E+00 |
| Biotin-carboxyl Carrier Protein Assembly | 1.05E+00 |
| Adenine and Adenosine Salvage I | 1.05E+00 |
| 1D-myo-inositol Hexakisphosphate Biosynthesis V (from Ins(1,3,4)P3) | 1.05E+00 |
| Tetrahydrobiopterin Biosynthesis II | 1.05E+00 |
| RAN Signaling | 1.02E+00 |
| 1D-myo-inositol Hexakisphosphate Biosynthesis II (Mammalian) | 1.02E+00 |
| IL-3 Signaling | 1.02E+00 |
| Role of JAK family kinases in IL-6-type Cytokine Signaling | 1.02E+00 |
| Endocannabinoid Developing Neuron Pathway | 1.01E+00 |
| MIF-mediated Glucocorticoid Regulation | 1.01E+00 |
| Interferon Signaling | 1.01E+00 |
| CNTF Signaling | 1.01E+00 |
| Glutathione-mediated Detoxification | 9.80E-01 |
| Ferroptosis Signaling Pathway | 9.57E-01 |
| Pathogen Induced Cytokine Storm Signaling Pathway | 9.52E-01 |
| GADD45 Signaling | 9.46E-01 |
| Synaptic Long Term Potentiation | 9.45E-01 |
| Purine Nucleotides Degradation II (Aerobic) | 9.40E-01 |
| Telomerase Signaling | 9.34E-01 |
| P2Y Purigenic Receptor Signaling Pathway | 9.32E-01 |
| Role of MAPK Signaling in the Pathogenesis of Influenza | 9.31E-01 |
| 2-ketoglutarate Dehydrogenase Complex | 9.30E-01 |
| Heme Degradation | 9.30E-01 |
| Methionine Salvage II (Mammalian) | 9.30E-01 |
| Aryl Hydrocarbon Receptor Signaling | 9.26E-01 |
| CCR3 Signaling in Eosinophils | 9.20E-01 |
| MicroRNA Biogenesis Signaling Pathway | 9.05E-01 |
| IL-17 Signaling | 9.05E-01 |
| Role of PKR in Interferon Induction and Antiviral Response | 8.96E-01 |
| Thrombopoietin Signaling | 8.90E-01 |
| Macrophage Alternative Activation Signaling Pathway | 8.85E-01 |
| Iron homeostasis signaling pathway | 8.84E-01 |
| Insulin Receptor Signaling | 8.49E-01 |

| | |
|---|---|
| HMGB1 Signaling | 8.40E-01 |
| Arsenate Detoxification I (Glutaredoxin) | 8.39E-01 |
| Pentose Phosphate Pathway (Oxidative Branch) | 8.39E-01 |
| Galactose Degradation I (Leloir Pathway) | 8.39E-01 |
| Role of PI3K/AKT Signaling in the Pathogenesis of Influenza | 8.37E-01 |
| BER (Base Excision Repair) Pathway | 8.15E-01 |
| ABRA Signaling Pathway | 8.12E-01 |
| GŒ±q Signaling | 8.10E-01 |
| S100 Family Signaling Pathway | 8.06E-01 |
| TCA Cycle II (Eukaryotic) | 8.04E-01 |
| Gap Junction Signaling | 7.99E-01 |
| Aldosterone Signaling in Epithelial Cells | 7.90E-01 |
| Th1 and Th2 Activation Pathway | 7.90E-01 |
| Role of JAK1 and JAK3 in Œ≥c Cytokine Signaling | 7.88E-01 |
| Non-Small Cell Lung Cancer Signaling | 7.85E-01 |
| IL-22 Signaling | 7.75E-01 |
| Endocannabinoid Cancer Inhibition Pathway | 7.73E-01 |
| Stearate Biosynthesis I (Animals) | 7.72E-01 |
| GM-CSF Signaling | 7.72E-01 |
| Prolactin Signaling | 7.72E-01 |
| CLEAR Signaling Pathway | 7.67E-01 |
| Neutrophil Extracellular Trap Signaling Pathway | 7.65E-01 |
| Th1 Pathway | 7.58E-01 |
| Ephrin A Signaling | 7.55E-01 |
| Ethanol Degradation IV | 7.47E-01 |
| Basal Cell Carcinoma Signaling | 7.42E-01 |
| PFKFB4 Signaling Pathway | 7.37E-01 |
| Melanocyte Development and Pigmentation Signaling | 7.34E-01 |
| D-myo-inositol (1,4,5)-Trisphosphate Biosynthesis | 7.21E-01 |
| Pancreatic Adenocarcinoma Signaling | 7.14E-01 |
| Type II Diabetes Mellitus Signaling | 7.13E-01 |
| Estrogen Receptor Signaling | 7.06E-01 |
| Thioredoxin Pathway | 7.06E-01 |
| Glycogen Biosynthesis II (from UDP-D-Glucose) | 7.06E-01 |
| Glycoaminoglycan-protein Linkage Region Biosynthesis | 7.06E-01 |
| NAD Salvage Pathway II | 6.96E-01 |
| Relaxin Signaling | 6.95E-01 |
| Glutathione Redox Reactions I | 6.72E-01 |
| Antiproliferative Role of Somatostatin Receptor 2 | 6.72E-01 |
| Role of WNT/GSK-3Œ≤ Signaling in the Pathogenesis of Influenza | 6.59E-01 |
| Superoxide Radicals Degradation | 6.55E-01 |

| | |
|---|---|
| Spliceosomal Cycle | 6.51E-01 |
| Atherosclerosis Signaling | 6.44E-01 |
| Macrophage Classical Activation Signaling Pathway | 6.41E-01 |
| PD-1, PD-L1 cancer immunotherapy pathway | 6.31E-01 |
| Hepatic Cholestasis | 6.26E-01 |
| Sphingosine and Sphingosine-1-phosphate Metabolism | 6.10E-01 |
| Œ±-Adrenergic Signaling | 6.10E-01 |
| Cachexia Signaling Pathway | 6.09E-01 |
| Triacylglycerol Biosynthesis | 6.06E-01 |
| Oxytocin Signaling Pathway | 5.96E-01 |
| Embryonic Stem Cell Differentiation into Cardiac Lineages | 5.71E-01 |
| FGF Signaling | 5.64E-01 |
| Retinoic acid Mediated Apoptosis Signaling | 5.51E-01 |
| Inhibition of Angiogenesis by TSP1 | 5.50E-01 |
| CDX Gastrointestinal Cancer Signaling Pathway | 5.46E-01 |
| Pentose Phosphate Pathway | 5.36E-01 |
| Glycine Betaine Degradation | 5.36E-01 |
| Erythropoietin Signaling Pathway | 5.24E-01 |
| Apelin Adipocyte Signaling Pathway | 5.12E-01 |
| Clathrin-mediated Endocytosis Signaling | 5.07E-01 |
| Hematopoiesis from Multipotent Stem Cells | 5.05E-01 |
| Oleate Biosynthesis II (Animals) | 5.05E-01 |
| Complement System | 5.00E-01 |
| Neuroprotective Role of THOP1 in Alzheimer's Disease | 4.99E-01 |
| NAD Signaling Pathway | 4.93E-01 |
| Cell Cycle Regulation by BTG Family Proteins | 4.85E-01 |
| Notch Signaling | 4.85E-01 |
| Assembly of RNA Polymerase I Complex | 4.76E-01 |
| NAD Phosphorylation and Dephosphorylation | 4.76E-01 |
| Guanosine Nucleotides Degradation III | 4.76E-01 |
| Remodeling of Epithelial Adherens Junctions | 4.57E-01 |
| Pyrimidine Ribonucleotides Interconversion | 4.56E-01 |
| DNA Double-Strand Break Repair by Non-Homologous End Joining | 4.50E-01 |
| Role of IL-17A in Psoriasis | 4.50E-01 |
| Acyl-CoA Hydrolysis | 4.50E-01 |
| Leukotriene Biosynthesis | 4.50E-01 |
| p53 Signaling | 4.48E-01 |
| Mechanisms of Viral Exit from Host Cells | 4.42E-01 |
| Apelin Cardiomyocyte Signaling Pathway | 4.39E-01 |
| Acetone Degradation I (to Methylglyoxal) | 4.29E-01 |
| Telomere Extension by Telomerase | 4.27E-01 |

| | |
|---|---|
| Choline Biosynthesis III | 4.27E-01 |
| Neuropathic Pain Signaling in Dorsal Horn Neurons | 4.23E-01 |
| Retinoate Biosynthesis I | 4.16E-01 |
| Pyrimidine Ribonucleotides De Novo Biosynthesis | 4.16E-01 |
| Endothelin-1 Signaling | 4.10E-01 |
| Chondroitin Sulfate Degradation (Metazoa) | 4.05E-01 |
| Bile Acid Biosynthesis, Neutral Pathway | 4.05E-01 |
| D-myo-inositol (1,4,5)-trisphosphate Degradation | 4.05E-01 |
| Adenosine Nucleotides Degradation II | 4.05E-01 |
| D-myo-inositol (1,3,4)-trisphosphate Biosynthesis | 3.85E-01 |
| Oxytocin in Brain Signaling Pathway | 3.83E-01 |
| Adrenomedullin signaling pathway | 3.83E-01 |
| DHCR24 Signaling Pathway | 3.83E-01 |
| tRNA Splicing | 3.81E-01 |
| Apelin Pancreas Signaling Pathway | 3.81E-01 |
| IL-23 Signaling Pathway | 3.81E-01 |
| Neurovascular Coupling Signaling Pathway | 3.74E-01 |
| TREM1 Signaling | 3.72E-01 |
| Ribonucleotide Reductase Signaling Pathway | 3.71E-01 |
| ISGylation Signaling Pathway | 3.70E-01 |
| Adipogenesis pathway | 3.70E-01 |
| GŒ±i Signaling | 3.64E-01 |
| Kinetochore Metaphase Signaling Pathway | 3.56E-01 |
| Role of MAPK Signaling in Inhibiting the Pathogenesis of Influenza | 3.56E-01 |
| Hereditary Breast Cancer Signaling | 3.52E-01 |
| Oxidative Phosphorylation | 3.43E-01 |
| MYC Mediated Apoptosis Signaling | 3.39E-01 |
| Cell Cycle: G2/M DNA Damage Checkpoint Regulation | 3.39E-01 |
| Valine Degradation I | 3.32E-01 |
| Inflammasome pathway | 3.32E-01 |
| Breast Cancer Regulation by Stathmin1 | 3.32E-01 |
| Amyloid Processing | 3.29E-01 |
| Phagosome Formation | 3.21E-01 |
| UVB-Induced MAPK Signaling | 3.20E-01 |
| IL-13 Signaling Pathway | 3.18E-01 |
| Amyotrophic Lateral Sclerosis Signaling | 3.18E-01 |
| Histamine Degradation | 3.17E-01 |
| Fatty Acid Œ±-oxidation | 3.17E-01 |
| Endoplasmic Reticulum Stress Pathway | 3.17E-01 |
| Mitochondrial Dysfunction | 3.05E-01 |
| Superpathway of D-myo-inositol (1,4,5)-trisphosphate Metabolism | 3.03E-01 |

| | |
|---|---|
| Pregnenolone Biosynthesis | 3.03E-01 |
| Putrescine Degradation III | 3.03E-01 |
| Corticotropin Releasing Hormone Signaling | 2.98E-01 |
| Nitric Oxide Signaling in the Cardiovascular System | 2.95E-01 |
| Pyrimidine Deoxyribonucleotides De Novo Biosynthesis I | 2.90E-01 |
| Histidine Degradation VI | 2.90E-01 |
| Oxidative Ethanol Degradation III | 2.86E-01 |
| EGF Signaling | 2.86E-01 |
| Necroptosis Signaling Pathway | 2.83E-01 |
| eNOS Signaling | 2.78E-01 |
| NOD1/2 Signaling Pathway | 2.78E-01 |
| Granulocyte Adhesion and Diapedesis | 2.78E-01 |
| Crosstalk between Dendritic Cells and Natural Killer Cells | 2.72E-01 |
| MSP-RON Signaling Pathway | 2.70E-01 |
| LPS/IL-1 Mediated Inhibition of RXR Function | 2.70E-01 |
| Role of NFAT in Cardiac Hypertrophy | 2.70E-01 |
| Glioma Signaling | 2.68E-01 |
| Bupropion Degradation | 2.66E-01 |
| Glycolysis I | 2.66E-01 |
| Gluconeogenesis I | 2.66E-01 |
| GŒ±s Signaling | 2.63E-01 |
| Polyamine Regulation in Colon Cancer | 2.63E-01 |
| Nicotine Degradation III | 2.63E-01 |
| Pyroptosis Signaling Pathway | 2.60E-01 |
| Acetylcholine Receptor Signaling Pathway | 2.57E-01 |
| Xenobiotic Metabolism CAR Signaling Pathway | 2.57E-01 |
| Role of JAK1, JAK2 and TYK2 in Interferon Signaling | 2.54E-01 |
| Ubiquinol-10 Biosynthesis (Eukaryotic) | 2.54E-01 |
| TR/RXR Activation | 2.53E-01 |
| Cardiomyocyte Differentiation via BMP Receptors | 2.44E-01 |
| Tryptophan Degradation X (Mammalian, via Tryptamine) | 2.44E-01 |
| Apelin Liver Signaling Pathway | 2.44E-01 |
| Small Cell Lung Cancer Signaling | 2.44E-01 |
| Melatonin Degradation I | 2.35E-01 |
| CDP-diacylglycerol Biosynthesis I | 2.34E-01 |
| Phosphatidylglycerol Biosynthesis II (Non-plastidic) | 2.15E-01 |
| Dopamine Degradation | 1.99E-01 |
| Dilated Cardiomyopathy Signaling Pathway | 0.00E+00 |
| Oxytocin in Spinal Neurons Signaling Pathway | 0.00E+00 |
| SNARE Signaling Pathway | 0.00E+00 |
| Immunogenic Cell Death Signaling Pathway | 0.00E+00 |

| | |
|---|---|
| Circadian Rhythm Signaling | 0.00E+00 |
| Synaptic Long Term Depression | 0.00E+00 |
| Chaperone Mediated Autophagy Signaling Pathway | 0.00E+00 |
| WNK Renal Signaling Pathway | 0.00E+00 |
| Adrenergic Receptor Signaling Pathway (Enhanced) | 0.00E+00 |
| GABAergic Receptor Signaling Pathway (Enhanced) | 0.00E+00 |
| Glutaminergic Receptor Signaling Pathway (Enhanced) | 0.00E+00 |
| NFKBIE Signaling Pathway | 0.00E+00 |
| Orexin Signaling Pathway | 0.00E+00 |
| Coagulation System | 0.00E+00 |
| LXR/RXR Activation | 0.00E+00 |
| VDR/RXR Activation | 0.00E+00 |
| FXR/RXR Activation | 0.00E+00 |
| PXR/RXR Activation | 0.00E+00 |
| Tight Junction Signaling | 0.00E+00 |
| Role of BRCA1 in DNA Damage Response | 0.00E+00 |
| IL-12 Signaling and Production in Macrophages | 0.00E+00 |
| Role of Pattern Recognition Receptors in Recognition of Bacteria and Viruses | 0.00E+00 |
| Role of NFAT in Regulation of the Immune Response | 0.00E+00 |
| FcŒ≥RIIB Signaling in B Lymphocytes | 0.00E+00 |
| CCR5 Signaling in Macrophages | 0.00E+00 |
| Calcium-induced T Lymphocyte Apoptosis | 0.00E+00 |
| Cytotoxic T Lymphocyte-mediated Apoptosis of Target Cells | 0.00E+00 |
| Airway Pathology in Chronic Obstructive Pulmonary Disease | 0.00E+00 |
| CTLA4 Signaling in Cytotoxic T Lymphocytes | 0.00E+00 |
| IL-15 Production | 0.00E+00 |
| T Helper Cell Differentiation | 0.00E+00 |
| IL-9 Signaling | 0.00E+00 |
| CD28 Signaling in T Helper Cells | 0.00E+00 |
| IL-15 Signaling | 0.00E+00 |
| Dendritic Cell Maturation | 0.00E+00 |
| Melatonin Signaling | 0.00E+00 |
| Cellular Effects of Sildenafil (Viagra) | 0.00E+00 |
| Docosahexaenoic Acid (DHA) Signaling | 0.00E+00 |
| ICOS-ICOSL Signaling in T Helper Cells | 0.00E+00 |
| Lipid Antigen Presentation by CD1 | 0.00E+00 |
| Role of CHK Proteins in Cell Cycle Checkpoint Control | 0.00E+00 |
| DNA Methylation and Transcriptional Repression Signaling | 0.00E+00 |
| Androgen Signaling | 0.00E+00 |
| Role of OCT4 in Mammalian Embryonic Stem Cell Pluripotency | 0.00E+00 |
| Growth Hormone Signaling | 0.00E+00 |

| | |
|---|---|
| CREB Signaling in Neurons | 0.00E+00 |
| Type I Diabetes Mellitus Signaling | 0.00E+00 |
| Primary Immunodeficiency Signaling | 0.00E+00 |
| Allograft Rejection Signaling | 0.00E+00 |
| Autoimmune Thyroid Disease Signaling | 0.00E+00 |
| Graft-versus-Host Disease Signaling | 0.00E+00 |
| p70S6K Signaling | 0.00E+00 |
| G Protein Signaling Mediated by Tubby | 0.00E+00 |
| Communication between Innate and Adaptive Immune Cells | 0.00E+00 |
| Sphingosine-1-phosphate Signaling | 0.00E+00 |
| Systemic Lupus Erythematosus Signaling | 0.00E+00 |
| CDC42 Signaling | 0.00E+00 |
| FAK Signaling | 0.00E+00 |
| AMPK Signaling | 0.00E+00 |
| Phospholipase C Signaling | 0.00E+00 |
| Altered T Cell and B Cell Signaling in Rheumatoid Arthritis | 0.00E+00 |
| Leptin Signaling in Obesity | 0.00E+00 |
| B Cell Development | 0.00E+00 |
| Regulation of IL-2 Expression in Activated and Anergic T Lymphocytes | 0.00E+00 |
| Granzyme A Signaling | 0.00E+00 |
| NUR77 Signaling in T Lymphocytes | 0.00E+00 |
| PKCŒ∏ Signaling in T Lymphocytes | 0.00E+00 |
| Role of Hypercytokinemia/hyperchemokinemia in the Pathogenesis of Influenza | 0.00E+00 |
| Antiproliferative Role of TOB in T Cell Signaling | 0.00E+00 |
| OX40 Signaling Pathway | 0.00E+00 |
| PI3K Signaling in B Lymphocytes | 0.00E+00 |
| Cyclins and Cell Cycle Regulation | 0.00E+00 |
| Cell Cycle Control of Chromosomal Replication | 0.00E+00 |
| Assembly of RNA Polymerase II Complex | 0.00E+00 |
| Role of JAK2 in Hormone-like Cytokine Signaling | 0.00E+00 |
| Dopamine-DARPP32 Feedback in cAMP Signaling | 0.00E+00 |
| Hematopoiesis from Pluripotent Stem Cells | 0.00E+00 |
| nNOS Signaling in Neurons | 0.00E+00 |
| Netrin Signaling | 0.00E+00 |
| Heparan Sulfate Biosynthesis | 0.00E+00 |
| Thyroid Hormone Metabolism II (via Conjugation and/or Degradation) | 0.00E+00 |
| Phospholipases | 0.00E+00 |
| Estrogen Biosynthesis | 0.00E+00 |
| Chondroitin Sulfate Biosynthesis | 0.00E+00 |
| Dermatan Sulfate Biosynthesis | 0.00E+00 |
| Nicotine Degradation II | 0.00E+00 |

| | |
|---|---|
| Serotonin Degradation | 0.00E+00 |
| Superpathway of Melatonin Degradation | 0.00E+00 |
| Noradrenaline and Adrenaline Degradation | 0.00E+00 |
| Ethanol Degradation II | 0.00E+00 |
| Superpathway of Methionine Degradation | 0.00E+00 |
| Agranulocyte Adhesion and Diapedesis | 0.00E+00 |
| Sperm Motility | 0.00E+00 |
| TEC Kinase Signaling | 0.00E+00 |
| Unfolded protein response | 0.00E+00 |
| SAPK/JNK Signaling | 0.00E+00 |
| Cardiac Œ≤-adrenergic Signaling | 0.00E+00 |
| GABA Receptor Signaling | 0.00E+00 |
| IL-4 Signaling | 0.00E+00 |
| B Cell Receptor Signaling | 0.00E+00 |
| Phototransduction Pathway | 0.00E+00 |
| Dopamine Receptor Signaling | 0.00E+00 |
| cAMP-mediated signaling | 0.00E+00 |
| p38 MAPK Signaling | 0.00E+00 |
| NF-Œ∫B Signaling | 0.00E+00 |
| T Cell Receptor Signaling | 0.00E+00 |
| GPCR-Mediated Integration of Enteroendocrine Signaling Exemplified by an L Cell | 0.00E+00 |
| GPCR-Mediated Nutrient Sensing in Enteroendocrine Cells | 0.00E+00 |
| Gustation Pathway | 0.00E+00 |
| Phagosome Maturation | 0.00E+00 |
| IL-7 Signaling Pathway | 0.00E+00 |
| Th17 Activation Pathway | 0.00E+00 |
| SPINK1 Pancreatic Cancer Pathway | 0.00E+00 |
| NER (Nucleotide Excision Repair, Enhanced Pathway) | 0.00E+00 |
| Endocannabinoid Neuronal Synapse Pathway | 0.00E+00 |
| T Cell Exhaustion Signaling Pathway | 0.00E+00 |
| Systemic Lupus Erythematosus in T Cell Signaling Pathway | 0.00E+00 |
| Systemic Lupus Erythematosus in B Cell Signaling Pathway | 0.00E+00 |
| White Adipose Tissue Browning Pathway | 0.00E+00 |
| Xenobiotic Metabolism PXR Signaling Pathway | 0.00E+00 |
| Insulin Secretion Signaling Pathway | 0.00E+00 |
| Coronavirus Replication Pathway | 0.00E+00 |
| Calcium Signaling | 0.00E+00 |

| Ratio | z-score | Molecules |
|---|---|---|
| 1.48E-01 | -2.828 | FAU,MAPK1,MT-TM,PDPK1,RAP1B,RASD2,RPL10A,RPL18,RPL18A,RPL |
| 1.25E-01 | 0 | FAU,ITGA7,ITGB2,MAPK1,MT-TM,PDPK1,PPP2R2D,RAP1B,RASD2,RPS |
| 9.20E-02 | 0 | CAV1,CCN4,CDH1,COL11A1,COL15A1,COL4A1,COL4A2,COL4A3,COL7. |
| 1.07E-01 | -0.816 | FAU,HMOX1,MAPK1,PDPK1,PPP2R2D,RAP1B,RASD2,RHOB,RPS12,RPS |
| 1.03E-01 | 3.71 | CHUK,FAU,IL17RA,MAPK1,MAPK8,NFKBIA,REL,RPS12,RPS14,RPS16,R |
| 2.09E-01 | -0.707 | MAPK1,MMP1,MMP3,MT2A,PLAU,RAP1B,RASD2,RRAS2,TIMP3 |
| 8.12E-02 | -0.209 | ADAM12,ADAM19,CDH1,CHUK,COL11A1,COL15A1,COL4A1,COL4A2,( |
| 9.55E-02 | -1.147 | CDH1,FZD1,FZD4,MAPK1,MAPK8,MMP1,MMP17,MMP3,NFKBIA,NOT |
| 1.09E-01 | -0.302 | ARPC5,BAIAP2,CYFIP2,IQGAP2,ITGA7,ITGB2,MAPK1,MAPK8,MCF2L,P |
| 1.01E-01 | -0.775 | ARPC5,BAIAP2,CDH1,FGF1,NOTCH3,PAK1,PPP2R2D,RAP1B,RASD2,RC |
| 8.47E-02 | 0.243 | ACAN,CCN4,CHUK,CREB1,DKK1,FZD1,FZD4,ITGA7,ITGB2,MMP1,MMF |
| 6.47E-02 | NaN | ADAM12,ADAM19,ARPC5,BAIAP2,EPHB1,FZD1,FZD4,ITGA7,ITGB2,M/ |
| 8.76E-02 | NaN | COL11A1,COL15A1,COL4A1,COL4A2,COL4A3,COL7A1,COL8A2,CSF1,C |
| 9.38E-02 | -2.828 | CHMP6,COL11A1,COL15A1,COL4A1,COL4A2,COL4A3,COL7A1,COL8A. |
| 9.27E-02 | -1 | CHUK,INPPL1,ITGA7,ITGB2,MAGIX,MAPK1,PDPK1,RAP1B,RASD2,REL, |
| 1.18E-01 | 1.414 | CHUK,CREB1,MAPK1,MAPK8,NFKBIA,PAK1,RAP1B,RASD2,REL,RRAS2 |
| 7.54E-02 | 0.688 | CHUK,COL11A1,COL15A1,COL4A1,COL4A2,COL4A3,COL7A1,COL8A2,I |
| 1.00E-01 | 1 | CHUK,CREB1,MAP3K10,MAP3K4,MAPK1,MAPK8,PDPK1,RAP1B,RASD |
| 1.06E-01 | -1.265 | FZD1,FZD4,ID1,ID3,ID4,MAPK1,RAP1B,RASD2,RRAS2,SMAD1,TCF7L1 |
| 1.05E-01 | 0.447 | IGFBP2,IGFBP4,IGFBP5,MAPK1,MAPK8,PDPK1,RAP1B,RASD2,RRAS2,Y |
| 1.10E-01 | 0 | CREB1,FST,MAPK1,MAPK8,RAP1B,RASD2,REL,RRAS2,SMAD1,SMAD7 |
| 7.96E-02 | -1.5 | FZD1,FZD4,ID1,ID3,ID4,INHBA,KLF2,MAPK1,NTF4,PDPK1,RAP1B,RASD |
| 7.92E-02 | 1.897 | ARPC5,CREB1,DOK1,EPHB1,FGF1,ITGA7,ITGB2,MAPK1,NGEF,PAK1,R/ |
| 7.41E-02 | 0.943 | CDH1,CDH13,CDH17,COL4A3,CREB1,GJA1,LAMA2,MAP3K10,MAP3K4 |
| 1.08E-01 | -1.134 | ARPC5,BAIAP2,ITGA7,ITGB2,RAP1B,RASD2,RHOB,ROCK2,RRAS2,VASP |
| 6.63E-02 | NaN | CHUK,CREB1,CSF1,DKK1,FZD1,FZD4,IL16,IL17RA,IRAK1,MAPK1,MMP: |
| 6.15E-02 | 0.655 | CHUK,CREB1,FZD1,FZD4,IL17RA,IRAK1,ITGA7,ITGB2,MAPK1,MAPK8,N |
| 9.65E-02 | NaN | CHUK,CREB1,GSTP1,MAPK1,NFKBIA,NKX3-1,PDPK1,RAP1B,RASD2,RE |
| 9.02E-02 | 1 | CDH1,CDH13,CDH17,CHUK,MAPK1,MAPK8,NFKBIA,RAP1B,RASD2,REI |
| 7.77E-02 | 0.258 | CYBA,EDIL3,EZR,ITGB2,MAP3K4,MAPK1,MAPK8,MMP1,MMP17,MMI |
| 8.44E-02 | 0.832 | BHLHE40,CCN4,CHUK,CREB1,HLA-A,HLA-B,HLA-DPA1,HLA-DPB1,HMC |
| 1.07E-01 | 0 | CHUK,MAPK1,NFKBIA,RAP1B,RASD2,REL,ROCK2,RRAS2,TCF7L1 |
| 1.54E-01 | 0.816 | ADAM12,MMP1,MMP17,MMP3,TFPI2,TIMP3 |
| 8.57E-02 | -0.302 | CREB1,MAPK1,NFKBIA,RAP1B,RASD2,REL,RRAS2,TCF7L1,VEGFC,VHL,Y |
| 7.82E-02 | 0.277 | CSF1,FGF1,HLA-A,HLA-B,MAPK1,MMP1,MMP17,MMP3,PLAU,RAP1B, |
| 7.50E-02 | -0.333 | CHUK,IL17RA,INPPL1,ITGA7,ITGB2,MAPK1,NFKBIA,PDPK1,PPP2R2D,R |
| 8.39E-02 | 0 | GSTO1,GSTP1,HMOX1,MAP3K10,MAP3K4,MAPK1,MAPK8,MGST2,NC |
| 6.74E-02 | -0.258 | ARHGEF16,ARPC5,BAIAP2,CDC42EP5,CDH1,CDH13,CDH17,EZR,ITGA7 |
| 1.23E-01 | NaN | NUB1,PSMB8,PSMC3,PSMC4,PSMD11,PSMD8,UBE2Z |

| | | |
|---|---|---|
| 7.14E-02 | 0.302 | CDH1,CHUK,HMOX1,IRAK1,ITGB2,MAPK1,MAPK8,NFKBIA,RAP1B,RAS |
| 1.05E-01 | 0 | ANGPTL1,CHUK,NFKBIA,PAK1,RAP1B,RASD2,REL,RRAS2 |
| 1.05E-01 | 0.816 | CREB1,DOK1,MAPK1,MAPK8,PDLIM7,RAP1B,RASD2,RRAS2 |
| 1.03E-01 | 0 | CHUK,IRAK1,MAPK1,MAPK8,NFKBIA,REL,TIRAP,UBB |
| 1.03E-01 | 0.378 | CREB1,MAPK1,MAPK8,NTF4,PDPK1,RAP1B,RASD2,RRAS2 |
| 1.56E-01 | 0 | CHUK,MAPK8,NFKBIA,REL,TBK1 |
| 8.77E-02 | -0.816 | CHUK,GSTO1,HMOX1,MAPK1,MAPK8,NFKBIA,NXN,PLCB4,REL,SLC23/ |
| 1.01E-01 | -0.707 | CDH1,MAPK1,NTF4,PDK1,RAP1B,RASD2,RRAS2,TCF7L1 |
| 6.91E-02 | 1.291 | CDH1,CHUK,FST,FSTL3,GATA2,INHBA,MAPK1,MAPK8,NFKBIA,ROCK2, |
| 8.62E-02 | 1 | CDH1,FGF1,MAPK1,MMP1,MMP17,MMP3,RAP1B,RASD2,RRAS2,VEG |
| 8.47E-02 | NaN | AP2S1,AP3S2,CAV1,CLTA,HLA-A,HLA-B,ITGB2,RAP1B,RASD2,RRAS2 |
| 1.47E-01 | NaN | CHUK,MAPK1,MAPK8,NFKBIA,REL |
| 8.40E-02 | 1 | CHUK,CREB1,HLA-DPA1,HLA-DPB1,ITGB2,MAPK1,RAP1B,RASD2,REL,I |
| 1.08E-01 | 0.816 | CHUK,ISG15,MAPK8,NFKBIA,PPIB,REL,TBK1 |
| 7.80E-02 | -0.905 | CHUK,IL17RA,IRAK1,MAPK1,MMP1,MMP17,MMP3,PGAM5,PLAU,TN |
| 6.16E-02 | -1.941 | ATG101,ATG4B,CDH1,DUSP6,GTF3C2,MAP1LC3A,MAPK1,NDRG1,ND |
| 8.65E-02 | -1.134 | CAPN5,CHUK,MAPK1,MAPK8,NFKBIA,RAP1B,RASD2,REL,RRAS2 |
| 7.03E-02 | 0.333 | CHUK,CRABP2,HMOX1,IRAK1,MAPK1,MAPK8,NFKBIA,PDPK1,RAP1B,I |
| 1.01E-01 | 1.342 | LAMA2,MAPK1,MAPK8,PAK1,RAP1B,RASD2,RRAS2 |
| 9.20E-02 | 0 | CHUK,IL17RA,MAPK1,MAPK8,MMP1,MMP3,NFKBIA,REL |
| 7.87E-02 | 0 | COL11A1,COL15A1,COL4A1,COL4A2,COL4A3,COL7A1,COL8A2,LAMA2 |
| 7.06E-02 | NaN | CDH1,MAP3K10,MAP3K4,MAPK1,MAPK8,PAK1,PDPK1,RAP1B,RASD2 |
| 8.89E-02 | 0 | CAPN5,EZR,ITGA7,ITGB2,MAPK1,RAP1B,RASD2,RRAS2 |
| 1.28E-01 | NaN | HLA-A,HLA-B,HLA-DPA1,HLA-DPB1,PSMB8 |
| 6.36E-02 | -0.775 | ARPC5,CDH1,GJA1,LAMA2,MAP3K10,MAP3K4,MAPK1,MAPK8,PDPK1 |
| 9.59E-02 | 0.816 | MAPK1,PLAU,RAP1B,RASD2,RHOB,RRAS2,TIMP3 |
| 6.67E-02 | NaN | CDH1,FGF1,FZD1,FZD4,MAPK1,NOTCH3,RAP1B,RASD2,RBPJ,REL,RRA |
| 7.58E-02 | 1.134 | ITGA7,ITGB2,MAP3K10,MAP3K4,MAPK1,MAPK8,PAK1,RAP1B,RASD2, |
| 7.19E-02 | 1.508 | CREB1,DKK1,FZD1,FZD4,MAPK8,PLCB4,ROCK2,SMAD1,TCF7L1,TGFBR |
| 1.05E-01 | 1 | CHUK,MAP3K10,MAP3K4,MAPK8,NFKBIA,REL |
| 1.05E-01 | NaN | IL17RA,MAPK1,MAPK8,MMP1,NFKBIA,REL |
| 1.19E-01 | 2 | CHUK,MAPK1,MAPK8,NFKBIA,REL |
| 6.31E-02 | -1.604 | CAPN5,DUSP6,HLA-A,HLA-B,HLA-DPA1,HLA-DPB1,IL17RA,JARID2,MA |
| 8.33E-02 | 0.378 | INHBA,MAPK1,MAPK8,RAP1B,RASD2,RRAS2,SMAD1,SMAD7 |
| 1.16E-01 | NaN | CHUK,MAPK1,MAPK8,NFKBIA,REL |
| 8.97E-02 | 0.447 | CHUK,MAPK1,NFKBIA,RAP1B,RASD2,REL,RRAS2 |
| 5.90E-02 | -1.069 | CDH1,FZD1,FZD4,MAPK1,MAPK8,MMP1,MMP17,MMP3,RAP1B,RASI |
| 1.00E-01 | -0.447 | CDH1,MAPK1,PDPK1,RAP1B,RASD2,RRAS2 |
| 1.00E-01 | 1.633 | FZD1,FZD4,MAPK8,ROCK2,RSPO3,WNT16 |
| 8.86E-02 | 1 | MAPK1,PAK1,RAP1B,RASD2,RRAS2,UBB,VHL |
| 5.86E-02 | NaN | HLA-A,HLA-B,HSPA4L,HSPB9,PSMB8,PSMC3,PSMC4,PSMD11,PSMD8, |

| | | |
|---|---|---|
| 9.84E-02 | NaN | MAPK1,PAK1,PLXNA2,RHOB,ROCK2,SEMA7A |
| 6.14E-02 | NaN | CHUK,CSF1,DKK1,FZD1,FZD4,MAPK1,MAPK8,MMP1,MMP3,NFKBIA,S |
| 5.69E-02 | 0 | BHLHE40,CAPN5,CDC26,CHUK,ITSN2,MAPK1,MAPKAPK3,PDK1,PPP2I |
| 1.88E-01 | NaN | MAPK1,MAPK8,UCHL1 |
| 6.75E-02 | -0.447 | CNOT7,MAPK1,PPP2R2D,PSMB8,PSMC3,PSMC4,PSMD11,PSMD8,TNI |
| 6.75E-02 | -1.265 | CDH1,HOTAIR,JARID2,MMP1,MMP17,MMP3,NFKBIA,REL,ROCK2,TCF |
| 8.64E-02 | 1.633 | MAPK1,MAPK8,PLCB4,RAP1B,RASD2,ROCK2,RRAS2 |
| 5.11E-02 | NaN | ARHGEF16,CDH1,FZD1,FZD4,ITGA7,ITGB2,MAPK1,MAPK8,NFKBIA,PA |
| 6.71E-02 | 0.302 | FZD1,FZD4,INHBA,MAPK1,RAP1B,RASD2,RRAS2,SMAD1,TCF7L1,TGFE |
| 5.50E-02 | 1.886 | CREB1,FZD1,FZD4,ID4,ITGB2,LAMA2,MAPK1,NDRG1,PDK1,RAP1B,RA |
| 4.81E-02 | NaN | CAV1,CHUK,CREB1,GJA1,HLA-A,HLA-B,HLA-DPA1,HLA-DPB1,IL17RA,K |
| 1.06E-01 | 0 | CHUK,IRAK1,MAPK1,NFKBIA,REL |
| 6.13E-02 | 1.155 | ARPC5,CAPN5,CAV1,ITGA7,ITGB2,MAPK1,MAPK8,PAK1,RAP1B,RASD: |
| 7.26E-02 | NaN | FZD1,FZD4,MAPK1,RAP1B,RASD2,RRAS2,SMAD1,TCF7L1,WNT16 |
| 5.63E-02 | -1.134 | CAPN5,CLTA,CREB1,MAP3K10,MAPK1,MAPK8,PDPK1,PENK,PLCB4,PS |
| 8.24E-02 | NaN | MAPK1,PSMB8,PSMC3,PSMC4,PSMD11,PSMD8,REL |
| 7.09E-02 | 0.378 | MAPK1,MAPK8,PLCB4,RAP1B,RASD2,RRAS2,TUBB2A,YWHAB,YWHAH |
| 8.96E-02 | 1.342 | CHUK,IL17RA,MAPK1,MAPK8,NFKBIA,REL |
| 7.48E-02 | 1.342 | ITGA7,ITGB2,MAPK1,MAPK8,PAK1,RAP1B,RASD2,RRAS2 |
| 7.48E-02 | -0.816 | CHUK,MAPK1,NFKBIA,RAP1B,RASD2,REL,RRAS2,TNFRSF11B |
| 8.05E-02 | 0 | CAV1,INPPL1,MAPK1,MAPK8,RAP1B,RASD2,RRAS2 |
| 8.05E-02 | 0.447 | CDH1,FZD1,FZD4,RBPJ,REL,TCF7L1,WNT16 |
| 6.98E-02 | 1.414 | CAV1,CAV2,KCNJ6,MAPK1,PAK1,PDPK1,RAP1B,RASD2,RRAS2 |
| 6.98E-02 | 1.134 | CHUK,MAPK1,MAPK8,NFKBIA,RAP1B,RASD2,REL,RRAS2,TNFRSF11B |
| 1.00E-01 | NaN | CDH1,MAPK1,RAP1B,RASD2,RRAS2 |
| 1.00E-01 | 1 | CHUK,NFKBIA,REL,TGFBR3,TNFRSF11B |
| 6.12E-02 | -1 | DUSP6,G6PC3,INPPL1,ITPK1,PGAM5,PIP4K2A,PLCB4,PPP4C,PPP5C,PT |
| 6.32E-02 | 0.707 | CDH1,DKK1,FZD1,FZD4,GJA1,PPP2R2D,SOX4,TCF7L1,TGFBR3,UBB,WI |
| 5.74E-02 | 0.333 | ARPC5,BAIAP2,CYFIP2,EZR,FGF1,IQGAP2,ITGA7,ITGB2,MAPK1,PAK1,I |
| 5.36E-02 | -0.535 | CHUK,CREB1,FZD1,HLA-A,HLA-B,HLA-DPA1,HLA-DPB1,HMOX1,IRAK1, |
| 9.80E-02 | 1 | CHUK,MAPK8,NFKBIA,PAK1,REL |
| 9.80E-02 | 1 | MAPK1,MAPK8,RAP1B,RASD2,RRAS2 |
| 7.69E-02 | 1 | MAPK8,PPP2R2D,RAP1B,RASD2,REL,RRAS2,TNFRSF11B |
| 7.69E-02 | 1.342 | CHUK,MAP3K10,MAP3K4,MAPK1,MAPK8,NFKBIA,REL |
| 5.97E-02 | -1.155 | BHLHE40,CAV1,ID1,MAPK1,RAP1B,RASD2,RRAS2,SMAD1,STMN3,TGI |
| 5.78E-02 | 0 | ARHGEF16,ARHGEF26,CREB1,GATA2,MAPK1,PDPK1,PLCB4,RAP1B,RA |
| 8.33E-02 | 2.449 | EPHB1,ITSN2,MAPK1,PAK1,RGS3,ROCK2 |
| 6.33E-02 | -0.447 | FZD1,FZD4,GJA1,MAPK1,RAP1B,RASD2,RRAS2,TCF7L1,VEGFC,WNT16 |
| 7.53E-02 | 0.816 | MAPK1,MAPK8,PAK1,PDPK1,RAP1B,RASD2,RRAS2 |
| 4.90E-02 | -0.655 | CRABP2,CREB1,DKK1,GNGT1,HOXB5,HOXD3,KLF2,MAPK1,MAPK8,MI |
| 1.08E-01 | NaN | CHUK,NFKBIA,REL,TNFRSF12A |

| | | |
|---|---|---|
| 8.11E-02 | -0.447 | CREB1,RAP1B,RASD2,RRAS2,YWHAB,YWHAH |
| 5.80E-02 | 0.577 | CHUK,CSF1,MAPK1,MAPK8,MMP1,PPP2R2D,RAP1B,RASD2,RRAS2,T[illegible] |
| 6.84E-02 | 1.342 | ITGA7,ITGB2,MAPK1,MAPK8,PAK1,RAP1B,RASD2,RRAS2 |
| 7.89E-02 | 1 | CSF1,ITGB2,PAK1,RAP1B,RASD2,RRAS2 |
| 4.61E-02 | 1.528 | CHUK,FGF1,FZD1,FZD4,HAND2,IL17RA,ITGA7,ITGB2,MAP3K10,MAP3[illegible] |
| 6.72E-02 | 1.134 | MAPK1,MAPK8,PLCB4,RAP1B,RASD2,RHOB,ROCK2,RRAS2 |
| 7.14E-02 | 1 | MAPK1,MAPK8,PARP9,PLCB4,RAP1B,RASD2,RRAS2 |
| 1.30E-01 | NaN | GSTO1,NXN,SLC23A1 |
| 5.49E-02 | -0.378 | CYP2E1,GSTO1,GSTP1,HMOX1,MAPK1,MAPK8,MGST2,NQO2,PPIB,R[illegible] |
| 8.62E-02 | 0.447 | MAPK1,PDK1,RAP1B,RASD2,RRAS2 |
| 4.69E-02 | 1.279 | CAV1,CAV2,CDH1,CPE,CREB1,FBP2,G6PC3,HMOX1,KCNJ6,MAPK1,NF[illegible] |
| 5.76E-02 | 1.667 | CREB1,MAP3K10,MAP3K4,MAPK1,MAPK8,PAK1,PLCB4,RAP1B,RASD[illegible] |
| 5.76E-02 | 0.333 | CHUK,CYBA,MAP3K10,MAP3K4,MAPK1,MAPK8,NFKBIA,RAP1B,REL,R[illegible] |
| 5.58E-02 | 1.134 | CREB1,DUSP6,ITGA7,ITGB2,MAPK1,PAK1,RAP1B,RAPGEF3,RASD2,RR[illegible] |
| 5.95E-02 | 1.414 | ELMO2,MAPK1,MAPK8,PAK1,PLCB4,RAP1B,RASD2,RHOB,ROCK2,RRA[illegible] |
| 5.08E-02 | 0.775 | AP2S1,ARPC5,CDH1,CDH13,CDH17,COMP,CREB1,EPHB1,ITSN2,MAP[illegible] |
| 5.73E-02 | 0.333 | CDH1,CHUK,FGF1,MAPK1,MAPK8,MMP1,RAP1B,RASD2,REL,RRAS2,T[illegible] |
| 5.53E-02 | 0.905 | ATG101,ATG4B,CREB1,DRAM1,MAP1LC3A,MAPK1,MAPK8,PPP2R2D,[illegible] |
| 6.45E-02 | -1.134 | ARPC5,BAIAP2,CDC42EP5,EZR,NGEF,NRP2,PIP4K2A,ROCK2 |
| 5.10E-02 | NaN | ALDH1B1,GSTO1,GSTP1,HMOX1,MAP3K10,MAP3K4,MAPK1,MAPK8,[illegible] |
| 6.80E-02 | 1.633 | CDH1,MAPK8,NFKBIA,PIAS1,RCC1,RHOB,SP3 |
| 5.45E-02 | 0.632 | ARHGEF16,ARPC5,CDH1,CDH13,CDH17,EZR,ITGA7,ITGB2,PAK1,PIP4K[illegible] |
| 7.32E-02 | 0.447 | CREB1,MAPK1,PDPK1,RAP1B,RASD2,RRAS2 |
| 7.32E-02 | 1 | CREB1,MAPK1,RAP1B,RASD2,REL,RRAS2 |
| 7.32E-02 | 1 | MAPK1,PIAS1,RAP1B,RASD2,REL,RRAS2 |
| 7.32E-02 | 0 | MAPK8,NFKBIA,PPP2R2D,REL,TCF7L1,VEGFC |
| 8.06E-02 | NaN | MAPK1,MAPK8,RAP1B,RASD2,RRAS2 |
| 6.00E-02 | -0.707 | ACAN,ITGA7,ITGB2,NRP2,PAK1,PDE4C,PLXNA2,ROCK2,SMC3 |
| 1.82E-01 | NaN | GMPS,IMPDH1 |
| 5.56E-02 | 0.632 | HLA-A,HLA-B,MAP3K10,MAP3K4,MAPK1,PAK1,PVR,RAP1B,RASD2,RE[illegible] |
| 1.15E-01 | NaN | IL17RA,MAPK1,MAPK8 |
| 7.14E-02 | 0 | MAPK1,NRP2,RAP1B,RASD2,RRAS2,VEGFC |
| 9.09E-02 | NaN | MAPK1,MAPK8,NFKBIA,REL |
| 5.47E-02 | 0.333 | CDH1,CREB1,ITGB2,KRT18,MAPK1,MAPK8,PDPK1,PPP2R2D,REL,RHO[illegible] |
| 5.29E-02 | 1.897 | CHUK,COX15,ITGB2,MAPK1,MTCL1,NFKBIA,PDK1,RAP1B,RASD2,REL,[illegible] |
| 6.98E-02 | 2 | PPP2R2D,SAV1,SMAD1,STK3,YWHAB,YWHAH |
| 7.69E-02 | 2 | CHUK,MAPK8,NFKBIA,REL,TNFRSF11B |
| 7.69E-02 | 0 | MAPK1,PDPK1,RAP1B,RASD2,RRAS2 |
| 7.69E-02 | 0.447 | IRAK1,MAPK1,MAPK8,PAK1,PRPF4B |
| 6.11E-02 | 0.816 | ARPC5,MAPK1,NFKBIA,PLCB4,RAP1B,RASD2,REL,RRAS2 |
| 5.04E-02 | 0.535 | CHUK,FZD1,FZD4,MAPK1,NOTCH3,PLCB4,RAP1B,RASD2,RCC1,REL,RR[illegible] |

| | | |
|---|---|---|
| 6.90E-02 | -0.447 | ALDH1B1,GSTO1,GSTP1,MGST2,NQO2,REL |
| 5.00E-02 | 1.155 | AP2S1,CLTA,CREB1,KCNJ6,MAPK1,NFKBIA,PDK1,PENK,RAP1B,RASD2, |
| 5.56E-02 | -1.134 | DUSP6,G6PC3,INPPL1,ITPK1,PGAM5,PPP4C,PPP5C,PTPRN,SGPP2,STY |
| 5.56E-02 | -1.134 | DUSP6,G6PC3,INPPL1,ITPK1,PGAM5,PPP4C,PPP5C,PTPRN,SGPP2,STY |
| 7.58E-02 | 1 | CREB1,FZD1,FZD4,PLCB4,REL |
| 8.70E-02 | NaN | CHUK,NFKBIA,REL,TBK1 |
| 4.62E-02 | -0.258 | CDC26,CHUK,CREB1,DUSP6,MAPK1,MPPE1,NFKBIA,NTN1,PDE4C,PLC |
| 5.34E-02 | -1.414 | DUSP6,G6PC3,INPPL1,ITPK1,PGAM5,PIP4K2A,PPP4C,PPP5C,PTPRN,S[illegible] |
| 7.46E-02 | 2 | CHUK,MAPK1,MAPK8,NFKBIA,REL |
| 7.46E-02 | NaN | CDC26,CDC7,PLK2,PPP2R2D,SMC3 |
| 8.51E-02 | NaN | CREB1,IL17RA,MAPK1,REL |
| 5.93E-02 | 1.633 | IL17RA,MAP3K10,MAPK1,MAPK8,RAP1B,RASD2,RRAS2,TGFBR3 |
| 5.29E-02 | -1.508 | HMOX1,MAPK1,MMP1,MMP17,MMP3,NAA10,RAP1B,RASD2,RRAS2, |
| 7.35E-02 | 0 | MAPK1,PDPK1,RAP1B,RASD2,RRAS2 |
| 5.13E-02 | -1.265 | DUSP6,G6PC3,INPPL1,ITPK1,PGAM5,PIP4K2A,PLCB4,PPP4C,PPP5C,PT |
| 5.00E-01 | NaN | LIAS |
| 5.41E-02 | 0.333 | CHUK,CREB1,H2BC15,IRAK1,MAPK1,MAPK8,NFKBIA,TBK1,USP17L2 (i |
| 7.25E-02 | 1 | MAPK1,MT2A,RAP1B,RASD2,RRAS2 |
| 4.98E-02 | 0.905 | CREB1,HAND2,MAP3K10,MAP3K4,MAPK1,MAPK8,MAPKAPK3,PLCB4 |
| 5.84E-02 | -1.89 | CHD4,HLA-A,HLA-B,HLA-DPA1,HLA-DPB1,ITGB2,NOTCH3,TGFBR3 |
| 6.59E-02 | 0 | MAPK1,RAP1B,RASD2,REL,RRAS2,TCF7L1 |
| 6.14E-02 | 0.378 | ATP6V1G1,MAPK1,MAPK8,NFKBIA,RAP1B,RASD2,RRAS2 |
| 5.80E-02 | 0.378 | ARHGEF16,ARHGEF26,ARPC5,MAP3K10,MAPK1,MAPK8,PDK1,RAP1B |
| 6.09E-02 | -1.342 | ARPC5,BAIAP2,ITGA7,ITGB2,PAK1,PIP4K2A,RHOB |
| 6.09E-02 | -1 | LAMA2,MAPK1,MAPK8,PPP2R2D,RAP1B,RASD2,RRAS2 |
| 1.43E-01 | NaN | IMPDH1,NT5C1B |
| 1.43E-01 | NaN | GALT,UGP2 |
| 4.13E-02 | 0.928 | ADORA2B,CHUK,CREB1,CRHR2,DUSP6,FZD1,FZD4,GPR107,MAP3K10, |
| 9.68E-02 | NaN | AKR1C1/AKR1C2,CYP2E1,PTGR2 |
| 5.98E-02 | 0 | ITGA7,ITGB2,MAPK1,PDPK1,RAP1B,RASD2,RRAS2 |
| 6.38E-02 | -0.816 | ARPC5,EZR,HMOX1,MAPK1,PAK1,VASP |
| 5.67E-02 | 1.342 | KLF2,MAPK1,MAPK8,PLCB4,RAP1B,RASD2,REL,RRAS2 |
| 5.24E-02 | -1.414 | DUSP6,G6PC3,INPPL1,ITPK1,PGAM5,PPP4C,PPP5C,PTPRN,SGPP2,STY |
| 5.93E-02 | 0.816 | INPPL1,MAPK1,MAPK8,PDPK1,RAP1B,RASD2,RRAS2 |
| 4.92E-02 | -1.732 | DKK1,FZD1,FZD4,MAPK1,MMP1,MMP17,MMP3,SMAD1,TCF7L1,TNF[illegible] |
| 6.25E-02 | 0.447 | CHUK,IRAK1,MAPK1,MAPK8,NFKBIA,REL |
| 6.25E-02 | 1.342 | CHUK,MAPK8,NFKBIA,PARP9,REL,TBK1 |
| 1.33E-01 | NaN | GTF3C2,GTF3C6 |
| 5.15E-02 | -1.633 | CHUK,MAPK1,MAPK8,NFKBIA,PLCB4,RAP1B,RASD2,REL,RRAS2,TGFB[illegible] |
| 6.19E-02 | 0 | AK5,IRAK1,MAPK1,MAPK8,PAK1,PRPF4B |
| 5.79E-02 | 1.342 | MAPK1,MAPK8,PAK1,RAP1B,RASD2,REL,RRAS2 |

| | | |
|---|---|---|
| 6.67E-02 | NaN | CAV1,HLA-A,HLA-B,ITGA7,ITGB2 |
| 5.26E-02 | 0.707 | FZD1,FZD4,MAPK1,PLCB4,RAP1B,RASD2,RHOB,RRAS2,WNT16 |
| 6.58E-02 | NaN | CREB1,NFKBIA,UBE2Q2,UBE2Z,VHL |
| 6.06E-02 | 0.447 | MAPK1,RAP1B,RASD2,ROCK2,RRAS2,VEGFC |
| 6.00E-02 | 0 | CREB1,MAPK8,NFKBIA,PPP2R2D,SMC3,TRIM28 |
| 7.27E-02 | 1 | CHUK,MAPK1,NFKBIA,PDPK1 |
| 3.33E-01 | NaN | GSTO1 |
| 3.33E-01 | NaN | PTS |
| 3.33E-01 | NaN | HLCS |
| 3.33E-01 | NaN | APRT |
| 3.33E-01 | NaN | ITPK1 |
| 3.33E-01 | NaN | PTS |
| 1.18E-01 | NaN | KPNA3,RCC1 |
| 1.18E-01 | NaN | INPPL1,ITPK1 |
| 6.33E-02 | 1 | MAPK1,PAK1,RAP1B,RASD2,RRAS2 |
| 6.33E-02 | 0.447 | MAPK1,MAPK8,MMP1,PIAS1,VEGFC |
| 5.51E-02 | 0.816 | CREB1,MAPK1,MAPK8,RAP1B,RASD2,RRAS2,STMN2 |
| 8.33E-02 | NaN | MAPK1,NFKBIA,REL |
| 8.33E-02 | NaN | ISG15,PIAS1,PSMB8 |
| 7.02E-02 | NaN | MAPK1,RAP1B,RASD2,RRAS2 |
| 8.11E-02 | NaN | GSTO1,GSTP1,MGST2 |
| 5.34E-02 | -0.378 | H2BC15,HMOX1,MAPK1,RAP1B,RASD2,RRAS2,SLC3A2 |
| 4.31E-02 | -0.5 | BHLHE40,CDH1,COL11A1,COL15A1,COL4A1,COL4A2,COL4A3,COL7A1 |
| 6.67E-02 | 0 | MAP3K4,MAPK1,MAPK8,REL |
| 5.30E-02 | 1.633 | CREB1,MAPK1,PLCB4,RAP1B,RAPGEF3,RASD2,RRAS2 |
| 1.05E-01 | NaN | IMPDH1,NT5C1B |
| 5.56E-02 | 0 | MAPK1,PDPK1,PPP2R2D,RAP1B,RASD2,RRAS2 |
| 5.26E-02 | 1.342 | CREB1,MAPK1,PLCB4,RAP1B,RASD2,REL,RRAS2 |
| 5.95E-02 | NaN | MAPK1,MAPK8,RAP1B,RASD2,RRAS2 |
| 2.50E-01 | NaN | DLST |
| 2.50E-01 | NaN | HMOX1 |
| 2.50E-01 | NaN | BHMT |
| 5.03E-02 | NaN | ALDH1B1,GSTO1,GSTP1,MAPK1,MAPK8,MGST2,NQO2,REL |
| 5.22E-02 | 1.342 | MAPK1,PAK1,PLCB4,RAP1B,RASD2,ROCK2,RRAS2 |
| 4.81E-02 | -0.333 | HMOX1,MAPK1,MTCL1,RAP1B,RASD2,RRAS2,SMAD1,TPR,XPO5 |
| 4.81E-02 | -1 | IL17RA,MAPK1,MAPK8,MMP3,RAP1B,RASD2,RRAS2,TNFRSF11B,VEG |
| 5.15E-02 | 1.342 | CHUK,MAPK1,MAPK8,NFKBIA,NLRP1,REL,TIRAP |
| 6.35E-02 | NaN | MAPK1,RAP1B,RASD2,RRAS2 |
| 4.76E-02 | 0.333 | ADORA2B,CREB1,CSF1,HLA-DPA1,HLA-DPB1,MAPK1,NFKBIA,REL,TSC[illegible] |
| 5.11E-02 | NaN | ACO1,ATP6V1G1,HMOX1,ISCU,MAPK1,SLC25A37,SMAD1 |
| 5.00E-02 | 0 | INPPL1,MAPK1,MAPK8,PDPK1,RAP1B,RASD2,RRAS2 |

| | | |
|---|---|---|
| 4.79E-02 | 0.816 | MAPK1,MAPK8,RAP1B,RASD2,REL,RHOB,RRAS2,TNFRSF11B |
| 2.00E-01 | NaN | GSTO1 |
| 2.00E-01 | NaN | PGD |
| 2.00E-01 | NaN | GALT |
| 6.06E-02 | NaN | KPNA3,MAPK1,NFKBIA,REL |
| 6.82E-02 | NaN | MAPK1,NTHL1,UNG |
| 5.43E-02 | 0.447 | HAND2,MAPK8,ROCK2,TPM1,VEGFC |
| 4.71E-02 | 0.816 | CHUK,HMOX1,MAPK1,NFKBIA,PLCB4,REL,RHOB,ROCK2 |
| 3.76E-02 | -0.557 | ADORA2B,CDH1,CHUK,CREB1,CRHR2,EZR,FZD1,FZD4,GPR107,IRAK1,[illegible] |
| 8.70E-02 | NaN | ACO1,DLST |
| 4.55E-02 | NaN | CAV1,GJA1,MAPK1,PLCB4,RAP1B,RASD2,RRAS2,SP3,TUBB2A |
| 4.65E-02 | 0.447 | HSPA4L,HSPB9,MAPK1,NR3C2,PDPK1,PIP4K2A,PLCB4,TRAP1 |
| 4.65E-02 | NaN | CHD4,HLA-A,HLA-B,HLA-DPA1,HLA-DPB1,ITGB2,NOTCH3,TGFBR3 |
| 5.80E-02 | NaN | MAPK1,RAP1B,RASD2,RRAS2 |
| 5.32E-02 | 0 | MAPK1,PDPK1,RAP1B,RASD2,RRAS2 |
| 8.33E-02 | NaN | MAPK1,MAPK8 |
| 4.76E-02 | 0.378 | CDH1,CREB1,MAPK1,ROCK2,TCF7L1,TRIB3,VEGFC |
| 5.71E-02 | NaN | CYP2E1,PTGR2,SIRT6,THEM5 |
| 5.71E-02 | NaN | MAPK1,RAP1B,RASD2,RRAS2 |
| 5.26E-02 | 0 | MAPK1,PDPK1,RAP1B,RASD2,RRAS2 |
| 4.21E-02 | 0.302 | ATP6V1G1,BLOC1S3,CREB1,MAPK1,PPP2R2D,RAP1B,RASD2,RRAS2,T[illegible] |
| 4.01E-02 | 0 | COL11A1,COL15A1,COL4A1,COL4A2,COL4A3,COL7A1,COL8A2,ITGB2, |
| 4.92E-02 | -1.633 | HLA-A,HLA-B,HLA-DPA1,HLA-DPB1,ITGB2,NOTCH3 |
| 6.38E-02 | NaN | NGEF,PAK1,ROCK2 |
| 8.00E-02 | NaN | ALDH1B1,CYGB |
| 5.56E-02 | 0 | FZD1,FZD4,TCF7L1,WNT16 |
| 6.25E-02 | NaN | CREB1,FBP2,MAPK1 |
| 5.10E-02 | 1 | CREB1,MAPK1,RAP1B,RASD2,RRAS2 |
| 7.69E-02 | NaN | PIP4K2A,PLCB4 |
| 4.76E-02 | -0.447 | CYP2E1,HMOX1,MAPK1,MAPK8,REL,VEGFC |
| 4.58E-02 | 0.816 | CHUK,MAPK1,MAPK8,NFKBIA,PDPK1,REL,TNFRSF11B |
| 3.91E-02 | -0.832 | CAV1,CREB1,MAPK1,MMP1,MMP17,MMP3,NDUFB7,PAK1,PLCB4,RA |
| 1.43E-01 | NaN | NXN |
| 1.43E-01 | NaN | UGP2 |
| 1.43E-01 | NaN | B3GAT3 |
| 7.41E-02 | NaN | ACP2,NT5C1B |
| 4.52E-02 | 1 | CREB1,MAPK1,MPPE1,NFKBIA,PDE4C,RAP1B,REL |
| 7.14E-02 | NaN | GSTP1,MGST2 |
| 5.19E-02 | NaN | MAPK1,RAP1B,RASD2,RRAS2 |
| 5.13E-02 | NaN | FZD1,FZD4,TCF7L1,WNT16 |
| 1.25E-01 | NaN | CYGB |

| | | |
|---|---|---|
| 5.66E-02 | NaN | SF3A3,XAB2,YJU2 |
| 4.51E-02 | NaN | CSF1,ITGB2,MMP1,MMP3,REL,TNFRSF12A |
| 4.23E-02 | 0.378 | CHUK,HLA-DPA1,HLA-DPB1,IRAK1,JARID2,NFKBIA,PARP9,UQCRC2 |
| 4.67E-02 | NaN | HLA-A,HLA-B,HLA-DPA1,HLA-DPB1,TNFRSF11B |
| 4.19E-02 | NaN | CHUK,IRAK1,MAP3K4,MAPK8,NFKBIA,REL,TIRAP,TNFRSF11B |
| 1.11E-01 | NaN | SGPP2 |
| 4.59E-02 | 1 | MAPK1,RAP1B,RASD2,RRAS2,SLC8A3 |
| 3.80E-02 | -1.069 | CAPN5,CHUK,INHBA,MAPK1,MAPK8,PSMB8,PSMC3,PSMC4,PSMD11, |
| 5.36E-02 | NaN | GPAT2,PLPP3,SIRT6 |
| 3.90E-02 | 1.508 | CREB1,GJA1,MAPK1,MAPK8,OXTR,PLCB4,RAP1B,RASD2,REL,ROCK2,R |
| 1.00E-01 | NaN | HOXB5 |
| 4.65E-02 | 2 | CREB1,FGF1,MAPK1,MAPK8 |
| 5.00E-02 | NaN | CRABP2,PARP9,TNFRSF10D |
| 5.88E-02 | NaN | MAPK1,MAPK8 |
| 3.96E-02 | 0.378 | CDH17,CHUK,FZD1,FZD4,MAPK1,REL,TCF7L1,WNT16 |
| 9.09E-02 | NaN | PGD |
| 9.09E-02 | NaN | BHMT |
| 3.98E-02 | 0 | MAPK1,NFKBIA,PDPK1,RAP1B,RASD2,REL,RRAS2 |
| 4.40E-02 | 1 | GSTP1,MAPK1,MAPK8,MGST2 |
| 3.85E-02 | NaN | AP2S1,AP3S2,ARPC5,CLTA,FGF1,ITGB2,UBB,VEGFC |
| 8.33E-02 | NaN | CSF1 |
| 8.33E-02 | NaN | SCD |
| 5.41E-02 | NaN | ITGB2,MASP1 |
| 4.13E-02 | NaN | CREB1,HLA-A,HLA-B,MASP1,SRY |
| 3.97E-02 | 0 | GJA1,H2BC15,PARP9,SIRT6,SIRT7,SLC7A5 |
| 5.26E-02 | NaN | CNOT7,PPP2R2D |
| 5.26E-02 | NaN | NOTCH3,RBPJ |
| 7.69E-02 | NaN | POLR1B |
| 7.69E-02 | NaN | ACP2 |
| 7.69E-02 | NaN | NT5C1B |
| 4.41E-02 | NaN | ARPC5,CDH1,TUBB2A |
| 5.00E-02 | NaN | AK5,ENTPD3 |
| 7.14E-02 | NaN | XRCC5 |
| 7.14E-02 | NaN | IL17RA |
| 7.14E-02 | NaN | THEM5 |
| 7.14E-02 | NaN | MGST2 |
| 4.08E-02 | NaN | CCNK,DRAM1,MAPK8,PIAS1 |
| 4.88E-02 | NaN | CHMP6,VPS25 |
| 4.04E-02 | 2 | MAPK1,MAPK8,PLCB4,SLC8A3 |
| 4.76E-02 | NaN | CYP2E1,PTGR2 |
| 6.67E-02 | NaN | XRCC5 |

| | | |
|---|---|---|
| 6.67E-02 | NaN | HMOX1 |
| 3.96E-02 | 2 | CREB1,MAPK1,PLCB4,TACR1 |
| 4.65E-02 | NaN | AKR1C1/AKR1C2,NT5C1B |
| 4.65E-02 | NaN | AK5,ENTPD3 |
| 3.61E-02 | 0.816 | HMOX1,MAPK1,MAPK8,PLCB4,RAP1B,RASD2,RRAS2 |
| 6.25E-02 | NaN | ARSB |
| 6.25E-02 | NaN | AKR1C1/AKR1C2 |
| 6.25E-02 | NaN | INPPL1 |
| 6.25E-02 | NaN | NT5C1B |
| 5.88E-02 | NaN | INPPL1 |
| 3.52E-02 | 0.378 | CREB1,MAPK1,NLRP1,OXTR,RAP1B,RASD2,RRAS2 |
| 3.52E-02 | 1.342 | MAPK1,MAPK8,PLCB4,RAP1B,RASD2,REL,RRAS2 |
| 3.65E-02 | -0.447 | HMOX1,RAP1B,RASD2,RRAS2,THRB |
| 4.35E-02 | NaN | MPPE1,PDE4C |
| 4.35E-02 | NaN | MAPK8,REL |
| 4.35E-02 | NaN | NFKBIA,REL |
| 3.45E-02 | 1.414 | ADORA2B,ENTPD3,GJA1,KCNJ2,KCNJ6,PLCB4,PTGR2,ROCK2 |
| 3.90E-02 | NaN | IRAK1,MAPK1,REL |
| 3.53E-02 | 0.816 | CDH1,CREB1,MAPK1,MAPK8,PARP9,SMARCC1 |
| 3.70E-02 | -1 | ISG15,ITGB2,MAPK8,TBK1 |
| 3.60E-02 | -0.447 | FGF1,FZD1,FZD4,SAP30,SMAD1 |
| 3.57E-02 | NaN | CAV1,MAPK1,RAP1B,RASD2,RRAS2 |
| 3.64E-02 | 0 | CDC26,MAD1L1,SMC3,STAG3 |
| 3.80E-02 | NaN | MAPK1,MAPK8,NFKBIA |
| 3.52E-02 | NaN | RAP1B,RASD2,RRAS2,SMARCC1,UBB |
| 3.57E-02 | -2 | COX15,NDUFB7,UQCRC2,VPS9D1 |
| 4.00E-02 | NaN | CHUK,TNFRSF11B |
| 4.00E-02 | NaN | YWHAB,YWHAH |
| 5.00E-02 | NaN | HIBCH |
| 5.00E-02 | NaN | NLRP1 |
| 3.20E-02 | -0.229 | ADORA2B,ARHGEF16,CREB1,CRHR2,FZD1,FZD4,GPR107,MAPK1,NPY[illegible] |
| 3.92E-02 | NaN | CAPN5,MAPK1 |
| 3.17E-02 | 0.853 | ADORA2B,ARPC5,CRHR2,ELMO2,FZD1,FZD4,GPR107,HMOX1,ITGA7,[illegible] |
| 3.85E-02 | NaN | MAPK1,MAPK8 |
| 3.45E-02 | 1 | IL17RA,MAPK1,POSTN,ROCK2 |
| 3.45E-02 | 0 | CAPN5,HECW1,PAK1,VEGFC |
| 4.76E-02 | NaN | ALDH1B1 |
| 4.76E-02 | NaN | ALDH1B1 |
| 4.76E-02 | NaN | MAPK8 |
| 3.19E-02 | 0.905 | CAPN5,COX15,CREB1,FIS1,GSTP1,MAPK8,MGST2,NDUFB7,RAPGEF3,[illegible] |
| 4.55E-02 | NaN | INPPL1 |

| | | |
|---|---|---|
| 4.55E-02 | NaN | CYP2E1 |
| 4.55E-02 | NaN | ALDH1B1 |
| 3.29E-02 | 1 | ARPC5,CREB1,CRHR2,MAPK1,RAP1B |
| 3.33E-02 | NaN | CAV1,MAPK1,SLC7A1,VEGFC |
| 4.35E-02 | NaN | AK5 |
| 4.35E-02 | NaN | CYP2E1 |
| 3.57E-02 | NaN | ALDH1B1,CYP2E1 |
| 3.57E-02 | NaN | MAPK1,MAPK8 |
| 3.23E-02 | 0.447 | CAPN5,CHUK,PGAM5,PPID,TNFRSF11B |
| 3.21E-02 | NaN | AQP11,CAV1,PDPK1,SLC7A1,VEGFC |
| 3.17E-02 | 0.447 | CHUK,MAPK1,MAPK8,NFKBIA,SLC15A3,TBK1 |
| 3.17E-02 | NaN | EZR,ITGB2,MMP1,MMP17,MMP3,TNFRSF11B |
| 3.30E-02 | NaN | HLA-A,HLA-B,REL |
| 3.45E-02 | NaN | CSF1,ITGB2 |
| 3.11E-02 | NaN | ALDH1B1,CYP2E1,GSTO1,GSTP1,IRAK1,MAPK8,MGST2,TNFRSF11B |
| 3.12E-02 | 1.633 | MAPK1,MAPK8,PLCB4,RAP1B,RASD2,RRAS2,SLC8A3 |
| 3.20E-02 | NaN | MAPK1,RAP1B,RASD2,RRAS2 |
| 4.00E-02 | NaN | CYP2E1 |
| 4.00E-02 | NaN | FBP2 |
| 4.00E-02 | NaN | FBP2 |
| 3.17E-02 | 2 | ADORA2B,CREB1,MAPK1,RAPGEF3 |
| 3.39E-02 | NaN | MAPK1,SLC3A2 |
| 3.39E-02 | NaN | B3GAT3,CYP2E1 |
| 3.23E-02 | NaN | MAPK1,NLRP1,TNFRSF11B |
| 3.09E-02 | 0 | ADAM19,CREB1,HMOX1,KCNJ2,MAPK1,PLCB4 |
| 3.09E-02 | -1.342 | ALDH1B1,GSTO1,GSTP1,MAPK1,MGST2,PPP2R2D |
| 3.85E-02 | NaN | REL |
| 3.85E-02 | NaN | CYP2E1 |
| 3.12E-02 | 0 | AKR1C1/AKR1C2,THRB,THRSP,VEGFC |
| 3.70E-02 | NaN | SMAD1 |
| 3.70E-02 | NaN | ALDH1B1 |
| 3.70E-02 | NaN | MAPK8 |
| 3.12E-02 | NaN | CHUK,NFKBIA,REL |
| 3.17E-02 | NaN | B3GAT3,CYP2E1 |
| 3.57E-02 | NaN | GPAT2 |
| 3.33E-02 | NaN | GPAT2 |
| 3.12E-02 | NaN | ALDH1B1 |
| 2.00E-02 | NaN | CNN1,MAPK1,TPM1 |
| 2.86E-02 | NaN | OXTR |
| 7.35E-03 | NaN | SYN3 |
| 2.22E-02 | NaN | PGAM5,TNFRSF11B |

| | | |
|---|---|---|
| 2.61E-02 | NaN | BHLHE40,CREB1,MAPK1,PLCB4,RAP1B,RASD2,RRAS2 |
| 3.03E-02 | 1 | MAPK1,PLCB4,PPP2R2D,RAP1B,RASD2,RRAS2 |
| 1.89E-02 | 2.309 | ATP6V1G1,CHUK,IL17RA,MAPK1,MMP1,MMP17,MMP3,NFKBIA,PDP |
| 1.89E-02 | NaN | NR3C2,VHL |
| 1.00E-02 | NaN | ATP6V1G1,PLCB4 |
| 7.19E-03 | NaN | KCNJ6 |
| 1.54E-02 | 1.342 | CHUK,CREB1,HMOX1,MAPK1,PLCB4 |
| 8.91E-03 | 2 | CHUK,NFKBIA,REL,TNFRSF11B |
| 2.94E-02 | -0.378 | ATP6V1G1,KCNJ6,MAPK1,PIP4K2A,PLCB4,SLC8A3,SMAD1 |
| 2.86E-02 | NaN | PLAU |
| 2.44E-02 | NaN | REL,SCD,TNFRSF11B |
| 2.56E-02 | NaN | IGFBP5,SEMA3B |
| 1.59E-02 | NaN | G6PC3,MAPK8 |
| 1.54E-02 | NaN | SCD |
| 2.22E-02 | NaN | PPP2R2D,REL,TNFRSF11B,VASP |
| 1.25E-02 | NaN | SMARCC1 |
| 2.13E-02 | 1.342 | CHUK,MAPK1,MAPK8,REL,XRCC5 |
| 2.56E-02 | NaN | CREB1,MAPK1,MAPK8,REL |
| 1.35E-02 | 0.378 | CHUK,HLA-A,HLA-B,HLA-DPA1,HLA-DPB1,KPNA3,MAPK1,NFKBIA,OR |
| 1.13E-02 | 0.447 | DOK1,MAPK8,PDPK1,RAP1B,RASD2,RRAS2 |
| 4.01E-03 | NaN | MAPK1,MAPK8 |
| 1.09E-02 | NaN | HLA-A,HLA-B,HLA-DPA1,HLA-DPB1,ORAI1 |
| 4.71E-03 | NaN | HLA-A,HLA-B |
| 1.69E-02 | NaN | FGF1,MMP1 |
| 2.30E-02 | 0 | AP2S1,CLTA,HLA-A,HLA-B,HLA-DPA1,HLA-DPB1,HMOX1,ITGB2,MAPK |
| 2.44E-02 | NaN | AATK,EPHB1,REL |
| 1.06E-02 | NaN | HLA-A,HLA-B,HLA-DPA1,HLA-DPB1,TNFRSF11B |
| 2.86E-02 | NaN | REL |
| 2.13E-02 | 0.447 | ARPC5,CHUK,HLA-A,HLA-B,HLA-DPA1,HLA-DPB1,MAPK8,NFKBIA,PAK |
| 9.47E-03 | 0 | MAPK1,RAP1B,RASD2,REL,RRAS2 |
| 2.02E-02 | 0.577 | CHUK,CREB1,HLA-A,HLA-B,HLA-DPA1,HLA-DPB1,MAPK1,MAPK8,NFK |
| 2.78E-02 | NaN | MAPK1,PLCB4 |
| 1.33E-02 | NaN | PDE4C,PLCB4 |
| 2.63E-02 | NaN | PDPK1 |
| 1.58E-02 | -1 | CHUK,HLA-A,HLA-B,HLA-DPA1,HLA-DPB1,NFKBIA,PDPK1,REL |
| 2.41E-03 | NaN | AP2S1 |
| 1.72E-02 | NaN | PPP2R2D |
| 3.06E-02 | NaN | CHD4,SAP30,TET2 |
| 2.37E-02 | NaN | MAPK1,REL,SRY,TAF9B |
| 2.17E-02 | NaN | JARID2 |
| 2.82E-02 | NaN | MAPK1,PDPK1 |

| | | |
|---|---|---|
| 2.47E-02 | 0.775 | ADORA2B,CREB1,CRHR2,FZD1,FZD4,GPR107,MAPK1,NPY4R/NPY4R2, |
| 2.75E-02 | 1.134 | CHUK,CPE,HLA-A,HLA-B,HLA-DPA1,HLA-DPB1,IRAK1,MAPK1,MAPK8,[illegible] |
| 1.67E-02 | NaN | UNG |
| 8.21E-03 | NaN | HLA-A,HLA-B,HLA-DPA1,HLA-DPB1 |
| 8.73E-03 | NaN | HLA-A,HLA-B,HLA-DPA1,HLA-DPB1 |
| 8.97E-03 | NaN | HLA-A,HLA-B,HLA-DPA1,HLA-DPB1 |
| 1.55E-02 | 0.447 | MAPK1,PDPK1,PLCB4,PPP2R2D,RAP1B,RASD2,RRAS2,YWHAB,YWHA[illegible] |
| 6.41E-03 | NaN | PLCB4,THRB,TUB |
| 2.14E-03 | NaN | HLA-A,HLA-B |
| 2.50E-02 | NaN | MAPK1,PLCB4,RHOB |
| 7.50E-03 | NaN | HLA-A,HLA-B,LSM7,MAPK1,PRPF4B,RAP1B,RASD2,RRAS2 |
| 2.26E-02 | -0.378 | ARPC5,BAIAP2,CDC42EP5,HLA-A,HLA-B,HLA-DPA1,HLA-DPB1,IQGAP2 |
| 2.21E-02 | 0.209 | ADORA2B,ARPC5,CAPN5,CDH1,CRHR2,FZD1,FZD4,GPR107,IL17RA,IT[illegible] |
| 2.48E-02 | NaN | AK5,CREB1,MAPK1,ORAI1,PPP2R2D,SMARCC1 |
| 1.25E-02 | 0.333 | ARHGEF16,CREB1,HMOX1,ITGA7,ITGB2,MAPK1,MARCKS,PLCB4,RAP1 |
| 7.45E-03 | NaN | CHUK,CSF1,HLA-A,HLA-B,HLA-DPA1,HLA-DPB1,REL |
| 2.63E-02 | NaN | MAPK1,PLCB4 |
| 8.18E-03 | NaN | HLA-A,HLA-B,HLA-DPA1,HLA-DPB1 |
| 1.73E-02 | 0.707 | CHUK,MAPK1,MAPK8,NFKBIA,RAP1B,RASD2,REL,RRAS2 |
| 1.33E-02 | NaN | NDUFB7 |
| 1.17E-02 | NaN | HLA-A,HLA-B,HLA-DPA1,HLA-DPB1,MAPK1,PDK1 |
| 2.51E-02 | 0 | CHUK,HLA-A,HLA-B,HLA-DPA1,HLA-DPB1,MAP3K10,MAP3K4,MAPK1, |
| 1.16E-02 | NaN | ISG15 |
| 2.35E-03 | NaN | MAPK1 |
| 1.46E-02 | NaN | HLA-A,HLA-B,HLA-DPA1,HLA-DPB1,MAPK8,NFKBIA,REL |
| 1.69E-02 | 0.707 | CHUK,CREB1,MAPK1,NFKBIA,PDPK1,PLCB4,RAP1B,RASD2,REL,RRAS2 |
| 1.18E-02 | NaN | PPP2R2D |
| 1.79E-02 | NaN | CDC7 |
| 2.00E-02 | NaN | TAF9B |
| 1.61E-02 | NaN | VEGFC |
| 2.69E-02 | NaN | CREB1,KCNJ2,KCNJ6,PLCB4,PPP2R2D |
| 2.24E-03 | NaN | CSF1 |
| 2.13E-02 | NaN | CAPN5 |
| 2.78E-02 | NaN | NTN1,UNC5B |
| 1.11E-02 | NaN | B3GAT3 |
| 2.44E-02 | NaN | B3GAT3 |
| 2.90E-02 | NaN | HMOX1,PLCB4 |
| 2.17E-02 | NaN | CYP2E1 |
| 1.64E-02 | NaN | B3GAT3 |
| 1.59E-02 | NaN | B3GAT3 |
| 2.99E-02 | NaN | B3GAT3,CYP2E1 |

| | | |
|---|---|---|
| 2.78E-02 | NaN | ALDH1B1,B3GAT3 |
| 2.94E-02 | NaN | B3GAT3,CYP2E1 |
| 2.70E-02 | NaN | ALDH1B1 |
| 2.86E-02 | NaN | ALDH1B1 |
| 2.63E-02 | NaN | BHMT |
| 2.38E-02 | NaN | EZR,ITGB2,MMP1,MMP17,MMP3 |
| 1.56E-02 | NaN | AATK,EPHB1,PDE4C,PLCB4 |
| 1.04E-02 | NaN | ITGA7,ITGB2,MAPK8,PAK1,REL,RHOB |
| 1.11E-02 | NaN | MAPK8 |
| 1.20E-02 | 0.447 | MAP3K10,MAP3K4,MAPK8,RAP1B,RASD2,RRAS2 |
| 2.22E-02 | NaN | MPPE1,PDE4C,PPP2R2D,SLC8A3 |
| 1.52E-02 | NaN | AP2S1,UBB |
| 2.60E-02 | -0.258 | COL11A1,COL15A1,COL4A1,COL4A2,COL4A3,COL7A1,COL8A2,CREB1, |
| 2.05E-02 | 0.905 | CHUK,CREB1,INPPL1,MAP3K10,MAP3K4,MAPK1,MAPK8,NFKBIA,PDP |
| 1.85E-02 | NaN | GNGT1 |
| 2.50E-02 | NaN | PPP2R2D,PTS |
| 2.97E-02 | 0.378 | ADORA2B,CREB1,DUSP6,MAPK1,MPPE1,PDE4C,RAPGEF3 |
| 2.50E-02 | NaN | CREB1,IRAK1,MAPKAPK3 |
| 1.93E-02 | 0.632 | CHUK,IRAK1,MAPK8,NFKBIA,RAP1B,RASD2,RRAS2,TBK1,TGFBR3,TIRA |
| 2.60E-02 | 0 | CHUK,DUSP6,HLA-A,HLA-B,HLA-DPA1,HLA-DPB1,ITGB2,MAPK1,MAPK |
| 1.33E-02 | NaN | PLCB4 |
| 8.47E-03 | NaN | PLCB4 |
| 9.90E-03 | NaN | LIPI,ORAI1 |
| 2.44E-02 | NaN | ATP6V1G1,HLA-A,HLA-B,TUBB2A |
| 2.56E-02 | NaN | MAPK1,PDPK1 |
| 4.13E-03 | NaN | IRAK1,REL |
| 1.67E-02 | NaN | CPE |
| 1.11E-02 | NaN | XAB2 |
| 2.68E-02 | 2 | KCNJ6,MAPK1,MAPK8,PLCB4 |
| 2.12E-02 | NaN | HLA-A,HLA-B,HLA-DPA1,HLA-DPB1,MAPK1,MAPK8,PDK1,PPP2R2D,RA |
| 2.33E-02 | -1.604 | CREB1,EZR,HLA-A,HLA-B,HLA-DPA1,HLA-DPB1,MAPK1,ORAI1,PDK1,P |
| 1.52E-02 | -0.632 | INPPL1,IRAK1,ISG15,LILRB3,MAPK1,PDPK1,RAP1B,RASD2,REL,RRAS2, |
| 2.90E-02 | 1 | CREB1,FNDC5,MAPK1,THRB |
| 2.05E-02 | -1 | ALDH1B1,GSTO1,GSTP1,MGST2 |
| 2.55E-02 | 1.134 | CLCN3,CPE,CREB1,MAPK1,PLCB4,RAP1B,SEC61B |
| 2.22E-02 | NaN | TUBB2A |
| 2.27E-02 | 1 | CREB1,MAPK1,RAP1B,SLC8A3,TPM1 |

L21,RPL22,RPL26,RPL26L1,RPL30,RPL35,RPL37,RPL37A,RPL7A,RPL8,RPS12,RPS14,RPS16,RPS2,RPS
S12,RPS14,RPS16,RPS2,RPS20,RPS21,RPS27,RPS3,RPS5,RPS7,RPS8,RPS9,RPSA,RRAS2
A1,COL8A2,FZD1,FZD4,IL17RA,JARID2,MAPK1,MAPK8,MMP1,MMP17,MMP3,NOTCH3,PLAU,RAP
S14,RPS16,RPS2,RPS20,RPS21,RPS27,RPS3,RPS5,RPS7,RPS8,RPS9,RPSA,RRAS2,VEGFC

COL4A3,COL7A1,COL8A2,CREB1,CSF1,MAPK1,MAPK8,MMP1,MMP17,MMP3,NFKBIA,OSCAR,RAP

APK1,MMP1,MMP17,MMP3,NGEF,NRP2,NTF4,NTN1,PAK1,PLCB4,PLXNA2,RAP1B,RASD2,RGS3,RO

LAMA2,MAPK1,MAPK8,MMP1,NFKBIA,RAP1B,RASD2,RRAS2,TGFBR3,TNFRSF11B,VEGFC

1,MMP3,NFKBIA,PLCB4,RAP1B,RASD2,ROCK2,RRAS2,TCF7L1,TNFRSF11B,VEGFC,WNT16
MMP1,NFKBIA,PDK1,RAP1B,RASD2,REL,RHOB,ROCK2,RRAS2,SMAD7,TCF7L1,TGFBR3,TIRAP,TNFRS

K1,PLCB4,RAP1B,RAPGEF3,RASD2,RASGRF2,RBPJ,REL,RHOB,RRAS2,SMAD1,SMAD7,WNT16,ZBTB[illegible]

RT18,KRT34,KRT8,MAPK1,MAPK8,MMP1,MMP3,NDUFB7,NFKBIA,NR3C2,PLAU,RAP1B,RASD2,RR[illegible]

MP1,MMP3,MPPE1,NT5C1B,PDE4C,PDPK1,PLAU,REL,RHOB,RPL7A,SMARCC1,TGFBR3

K4,MAPK1,MAPK8,MAPKAPK3,MPPE1,PDE4C,PDK1,PLCB4,RAP1B,RASD2,REL,ROCK2,RRAS2,TGFB

,MAP3K4,MAPK1,MAPK8,MEGF6,MPPE1,NFKBIA,NPY4R/NPY4R2,OXTR,PAK1,PDE4C,PDPK1,PLCB4

MAPK1,MAPK8,MMP1,MMP17,MMP3,NPY4R/NPY4R2,NTF4,OXTR,PLCB4,ROCK2,S100A3,SMAD1

TGB2,MAPK1,MARCKS,NPY4R/NPY4R2,OXTR,PAK1,PIP4K2A,RAP1B,RAPGEF3,RASD2,ROCK2,RRAS

GA7,ITGB2,MAPK1,MAPK8,NPY4R/NPY4R2,OXTR,PAK1,PDPK1,RAP1B,RASD2,RRAS2,TACR1,TCF7L

ↄCK2,RRAS2,SEMA3B,SEMA7A,SLIT3,TUBB2A,UNC5B,VASP,VEGFC,WNT16
