## Supplementary material for "TBCK Deficiency Alters Ribosomal Function, RNA Splicing, and miRNA Networks: Insights from Multi-Omics Analyses": All_DE_genes_RNAseq_curated

| ESNG ID | Gene ID | baseMean | log2FoldChange | lfcSE | stat |
|---|---|---|---|---|---|
| ENSG00000225937.2 | PCA3 | 393.875735 | 2.366316797 | 0.23283949 | 10.1628671 |
| ENSG00000187210.13 | GCNT1 | 126.712631 | 2.427325537 | 0.25448337 | 9.53824817 |
| ENSG00000272234.1 | AC008945.1 | 36.7362865 | 2.918084885 | 0.32017507 | 9.11402893 |
| ENSG00000100036.12 | SLC35E4 | 129.381544 | 0.974218294 | 0.11418488 | 8.53193755 |
| ENSG00000230928.1 | AL139241.1 | 325.712485 | 3.195447641 | 0.39317911 | 8.12720598 |
| ENSG00000108641.15 | B9D1 | 27.9667317 | 1.625655522 | 0.21502969 | 7.56014458 |
| ENSG00000198865.9 | CCDC152 | 163.15812 | 3.192871467 | 0.4241143 | 7.52832782 |
| ENSG00000222047.8 | C10orf55 | 311.087166 | 2.812464746 | 0.37332912 | 7.53347274 |
| ENSG00000255050.1 | AC067930.5 | 100.098072 | -1.361798465 | 0.18589072 | -7.3258013 |
| ENSG00000122085.16 | MTERF4 | 733.512007 | 1.067736975 | 0.14859448 | 7.18557644 |
| ENSG00000253194.2 | AL365275.1 | 151.501561 | 1.837820539 | 0.25530954 | 7.19840145 |
| ENSG00000271581.1 | AL671883.2 | 1026.35235 | -1.457161618 | 0.20306284 | -7.1759149 |
| ENSG00000231584.8 | FAHD2CP | 21.1929942 | -6.064130433 | 0.85077358 | -7.1277841 |
| ENSG00000170425.3 | ADORA2B | 6.64370414 | 4.248110501 | 0.60789442 | 6.98823731 |
| ENSG00000177042.14 | TMEM80 | 208.630801 | 1.364424213 | 0.19600088 | 6.9613166 |
| ENSG00000261240.1 | AC009065.6 | 281.913221 | 0.83404446 | 0.12078967 | 6.90493204 |
| ENSG00000267193.5 | AC023421.2 | 8.60236232 | 4.892348945 | 0.71665403 | 6.8266538 |
| ENSG00000188015.9 | S100A3 | 34.0883119 | 1.902198278 | 0.28138689 | 6.76008129 |
| ENSG00000233351.1 | AL356124.2 | 169.824385 | 1.305578088 | 0.19587906 | 6.66522543 |
| ENSG00000231609.6 | AC009501.1 | 425.80042 | 1.183407704 | 0.17950181 | 6.59273409 |
| ENSG00000250483.1 | PPM1AP1 | 16.8246506 | 1.681099204 | 0.25606186 | 6.56520733 |
| ENSG00000248774.1 | AC097534.1 | 105.160574 | 1.227968611 | 0.1883144 | 6.52084274 |
| ENSG00000226457.1 | RPL22P3 | 10.7194606 | 4.409609577 | 0.67747065 | 6.50893081 |
| ENSG00000279035.1 | AC022211.4 | 114.386886 | -1.003392935 | 0.15669124 | -6.4036312 |
| ENSG00000097046.12 | CDC7 | 12.4267201 | 5.618466351 | 0.8928112 | 6.2930061 |
| ENSG00000115694.14 | STK25 | 82.6757211 | 0.903943451 | 0.14365787 | 6.2923352 |
| ENSG00000254866.2 | DEFB109D | 4.33101463 | 4.821516158 | 0.77250724 | 6.24138637 |
| ENSG00000231329.8 | AL031772.1 | 151.411532 | 0.838953305 | 0.13485857 | 6.22098628 |
| ENSG00000277443.2 | MARCKS | 90.5561212 | -1.54928799 | 0.251101 | -6.1699793 |
| ENSG00000186063.12 | AIDA | 96.8585688 | 0.775929162 | 0.12687215 | 6.11583518 |
| ENSG00000257607.1 | AC073957.2 | 25.0377757 | 2.383278437 | 0.38947532 | 6.11920269 |
| ENSG00000135424.16 | ITGA7 | 8.59637489 | 2.620514325 | 0.43066211 | 6.08484996 |
| ENSG00000246763.6 | RGMB-AS1 | 605.206922 | 1.504980098 | 0.24732015 | 6.08514956 |
| ENSG00000132763.14 | MMACHC | 2231.78267 | -0.647085633 | 0.10654654 | -6.0732677 |
| ENSG00000170584.10 | NUDCD2 | 66.2909137 | 1.525440333 | 0.25173867 | 6.0596187 |
| ENSG00000148331.11 | ASB6 | 318.50383 | -0.707657465 | 0.11787193 | -6.0036134 |
| ENSG00000257181.1 | AC025423.4 | 404.734404 | 0.935601368 | 0.15644261 | 5.98047664 |
| ENSG00000204706.14 | MAMDC2-AS | 5.88864397 | 4.577985211 | 0.77207156 | 5.92948299 |
| ENSG00000278367.1 | AL356652.1 | 92.2072617 | -1.336413531 | 0.22591943 | -5.915443 |
| ENSG00000120215.9 | MLANA | 20.5397208 | 1.661896245 | 0.28204398 | 5.89233023 |
| ENSG00000237525.6 | AC012668.3 | 15.2899794 | 1.730778022 | 0.29434061 | 5.88018761 |
| ENSG00000264785.1 | AC005722.3 | 8.84260925 | 3.884503874 | 0.66267026 | 5.86189559 |

| | | | | | |
|---|---|---|---|---|---|
| ENSG00000258471.2 | AL355922.2 | 158.941736 | -0.794018145 | 0.13576232 | -5.84859 |
| ENSG00000272356.1 | AL080317.3 | 245.451635 | 1.118457789 | 0.19116566 | 5.85072541 |
| ENSG00000204183.1 | GDF5OS | 60.1261561 | 3.183168746 | 0.54611262 | 5.8287771 |
| ENSG00000225521.1 | AC005237.1 | 45.6653244 | 1.22742066 | 0.21054654 | 5.82968815 |
| ENSG00000135452.9 | TSPAN31 | 674.652832 | -0.720548641 | 0.12413657 | -5.8044833 |
| ENSG00000157734.13 | SNX22 | 8549.88986 | -0.732138437 | 0.12676172 | -5.7757063 |
| ENSG00000273521.1 | AL162274.1 | 15.8657355 | 2.357079987 | 0.4090272 | 5.76264857 |
| ENSG00000230521.1 | HCG4P7 | 30.9447015 | -3.321262509 | 0.57849041 | -5.7412577 |
| ENSG00000262211.1 | AC008914.1 | 987.695079 | 1.093736706 | 0.19057957 | 5.7390028 |
| ENSG00000244676.5 | AL109761.1 | 19.4074911 | 1.282142567 | 0.22374509 | 5.73037181 |
| ENSG00000283117.1 | AC004949.1 | 34.1184263 | -1.563456269 | 0.27297657 | -5.7274375 |
| ENSG00000108511.9 | HOXB6 | 47.3629917 | 3.12042574 | 0.5473553 | 5.70091442 |
| ENSG00000154723.12 | ATP5PF | 53.051444 | 0.938702749 | 0.16477416 | 5.69690506 |
| ENSG00000187325.4 | TAF9B | 119.632396 | 1.382921239 | 0.24311051 | 5.68844694 |
| ENSG00000106772.17 | PRUNE2 | 23.3249801 | 2.061458429 | 0.36311308 | 5.67718025 |
| ENSG00000171497.4 | PPID | 33.5031233 | 1.28525324 | 0.22701357 | 5.66156999 |
| ENSG00000213347.10 | MXD3 | 1286.33105 | -0.589146313 | 0.10416055 | -5.6561365 |
| ENSG00000183250.11 | LINC01547 | 73.5419447 | 2.769653218 | 0.49203018 | 5.62903122 |
| ENSG00000228417.1 | AL360182.2 | 21.7186648 | 2.28919922 | 0.40705979 | 5.6237419 |
| ENSG00000267458.1 | AC092069.1 | 3659.89242 | -0.728148894 | 0.12946151 | -5.6244432 |
| ENSG00000224597.10 | SVIL-AS1 | 224.18514 | 1.422392755 | 0.2535628 | 5.60962706 |
| ENSG00000272821.1 | U62317.3 | 135.713313 | -0.7920977 | 0.14261714 | -5.5540149 |
| ENSG00000256377.5 | AC009509.1 | 39.3493332 | 1.231337933 | 0.2228878 | 5.52447424 |
| ENSG00000259884.1 | AC025259.3 | 178.391331 | -1.624726273 | 0.29469293 | -5.5132855 |
| ENSG00000255435.6 | AP001267.3 | 93.6891869 | 0.716562388 | 0.13082696 | 5.47717666 |
| ENSG00000246273.7 | SBF2-AS1 | 391.581041 | 0.897404274 | 0.16403457 | 5.47082398 |
| ENSG00000254042.1 | AC011365.1 | 1329.23059 | 1.713985613 | 0.3143547 | 5.4523938 |
| ENSG00000214413.7 | BBIP1 | 84.3132884 | 0.975649029 | 0.17915076 | 5.44596638 |
| ENSG00000285868.1 | | 12.4585781 | 2.105481102 | 0.38861715 | 5.41788007 |
| ENSG00000281344.1 | HELLPAR | 64.2960543 | -5.92873238 | 1.09501403 | -5.4142981 |
| ENSG00000174804.3 | FZD4 | 396.67791 | 1.682651225 | 0.31108306 | 5.40900956 |
| ENSG00000236256.9 | DIAPH2-AS1 | 29.3786896 | 1.472148269 | 0.27277615 | 5.39690982 |
| ENSG00000245293.2 | AC096564.1 | 106.896233 | 1.115669931 | 0.20673683 | 5.39657084 |
| ENSG00000255487.1 | AC087362.2 | 26.7275759 | 1.116463376 | 0.20691062 | 5.39587284 |
| ENSG00000260618.1 | AC025917.1 | 102.75427 | 1.012077073 | 0.18755167 | 5.3962573 |
| ENSG00000104848.1 | KCNA7 | 66.1553944 | -2.686191355 | 0.49882821 | -5.3850029 |
| ENSG00000215030.5 | RPL13P12 | 12.7156151 | -4.64877684 | 0.86316566 | -5.3857296 |
| ENSG00000163320.10 | CGGBP1 | 26.4867958 | 1.283531588 | 0.23911642 | 5.36781026 |
| ENSG00000213713.3 | PIGCP1 | 211.133126 | 0.90842391 | 0.1704861 | 5.32843401 |
| ENSG00000267002.3 | AC060780.1 | 26.4664857 | 1.056532397 | 0.19882626 | 5.31384745 |
| ENSG00000140416.20 | TPM1 | 81.9565708 | -1.424731529 | 0.26872675 | -5.3017853 |
| ENSG00000237036.4 | ZEB1-AS1 | 21.9071036 | 2.336799259 | 0.44216213 | 5.28493761 |
| ENSG00000256928.1 | AP000763.4 | 96.4296577 | -1.103012021 | 0.20904882 | -5.276337 |

| | | | | | |
|---|---|---|---|---|---|
| ENSG00000204055.4 | AL158151.1 | 218.339379 | -1.072418309 | 0.20367483 | -5.2653453 |
| ENSG00000267128.1 | RNF157-AS1 | 12.9265815 | 3.24934019 | 0.61771687 | 5.26024196 |
| ENSG00000283703.2 | VSIG10L2 | 207.182575 | 2.437705059 | 0.46352293 | 5.25908193 |
| ENSG00000147687.18 | TATDN1 | 215.923694 | -0.875966095 | 0.16719437 | -5.2392082 |
| ENSG00000143878.9 | RHOB | 11.5213592 | -2.442299085 | 0.46743551 | -5.22489 |
| ENSG00000148303.16 | RPL7A | 43.3273678 | -1.145457971 | 0.21920649 | -5.2254746 |
| ENSG00000284830.1 | AL049557.1 | 6.69344211 | 2.61935682 | 0.50382524 | 5.19893926 |
| ENSG00000188368.9 | PRR19 | 62.1013722 | -1.002925186 | 0.19342226 | -5.1851591 |
| ENSG00000115457.9 | IGFBP2 | 4.89025312 | 4.612952313 | 0.89250238 | 5.16856022 |
| ENSG00000127928.12 | GNGT1 | 709.081647 | -4.249653599 | 0.82210399 | -5.169241 |
| ENSG00000279658.1 | AC110491.2 | 80.4207647 | 1.060407157 | 0.20616647 | 5.14345098 |
| ENSG00000214282.3 | KRT8P14 | 14.7704046 | 1.685430152 | 0.32823001 | 5.13490576 |
| ENSG00000251239.1 | AC004590.1 | 2.45154238 | 4.128591493 | 0.80446614 | 5.13208858 |
| ENSG00000254913.1 | AC239802.1 | 41.6809147 | 1.68283793 | 0.32958282 | 5.10596381 |
| ENSG00000130640.13 | TUBGCP2 | 124.255947 | -0.655694008 | 0.1285804 | -5.0994864 |
| ENSG00000166794.4 | PPIB | 32.0765734 | -1.102718948 | 0.21691674 | -5.0836046 |
| ENSG00000179218.13 | CALR | 165.240762 | -0.930664251 | 0.18336411 | -5.0754985 |
| ENSG00000130762.14 | ARHGEF16 | 23.0967634 | -2.30848717 | 0.4558104 | -5.0645777 |
| ENSG00000230046.1 | BIRC6-AS1 | 36.3482891 | 0.974266414 | 0.19276791 | 5.05409028 |
| ENSG00000249835.2 | VCAN-AS1 | 2937.12386 | 1.067556382 | 0.21115361 | 5.05582812 |
| ENSG00000281021.1 | AL078581.3 | 295.136189 | 0.534264532 | 0.10614401 | 5.03339325 |
| ENSG00000272173.1 | U47924.2 | 591.114352 | -1.007899968 | 0.20043804 | -5.0284863 |
| ENSG00000170909.13 | OSCAR | 14.6539125 | 2.552090235 | 0.50801922 | 5.02360957 |
| ENSG00000257950.3 | P2RX5-TAX1B | 284.514513 | -0.673559932 | 0.1343398 | -5.0138523 |
| ENSG00000231389.7 | HLA-DPA1 | 11.2145709 | -2.494893168 | 0.49929872 | -4.9967946 |
| ENSG00000262655.3 | SPON1 | 267.855716 | 2.051689949 | 0.41245379 | 4.9743511 |
| ENSG00000165443.11 | PHYHIPL | 13.2036022 | 2.550110345 | 0.51320209 | 4.96901782 |
| ENSG00000125779.22 | PANK2 | 125.144694 | 1.096176355 | 0.22200151 | 4.93769777 |
| ENSG00000278156.1 | TSC22D1-AS1 | 265.536969 | 1.083584332 | 0.21953541 | 4.93580662 |
| ENSG00000230513.1 | THAP7-AS1 | 28.9996059 | -1.134766136 | 0.23002957 | -4.9331316 |
| ENSG00000134871.18 | COL4A2 | 73.7092329 | -1.350367837 | 0.27423873 | -4.9240596 |
| ENSG00000246323.2 | AC113382.1 | 97.2522557 | 0.718076205 | 0.14588892 | 4.92207488 |
| ENSG00000284060.1 | AC002472.2 | 10.4399556 | -1.730825073 | 0.35194608 | -4.91787 |
| ENSG00000267765.1 | AC100793.3 | 237.568604 | 0.96665373 | 0.19684804 | 4.91065975 |
| ENSG00000234175.1 | AL355355.2 | 12.9467369 | 1.316224888 | 0.26838923 | 4.90416428 |
| ENSG00000259564.2 | AC009554.1 | 59.4523258 | 0.985170608 | 0.20290069 | 4.85543241 |
| ENSG00000117318.8 | ID3 | 5.90408637 | -2.599575708 | 0.53651589 | -4.8452912 |
| ENSG00000254192.1 | AC011365.2 | 1379.89957 | 1.632226903 | 0.33703348 | 4.84292208 |
| ENSG00000273398.6 | AC017083.3 | 104.31126 | -0.826701715 | 0.17078991 | -4.8404599 |
| ENSG00000145916.18 | RMND5B | 372.22716 | -0.616503701 | 0.12752746 | -4.8342819 |
| ENSG00000255817.1 | AC025576.2 | 63.6206737 | 1.121524612 | 0.23219583 | 4.83008075 |
| ENSG00000126602.10 | TRAP1 | 64.6996374 | 1.161197364 | 0.24051745 | 4.82791318 |
| ENSG00000267302.5 | RNFT1-DT | 13.7227827 | 1.325953515 | 0.27531613 | 4.81611272 |

| | | | | | |
|---|---|---|---|---|---|
| ENSG00000262728.5 | AC123768.3 | 17.6984827 | 1.320847705 | 0.27556343 | 4.79326194 |
| ENSG00000262533.1 | AC090617.3 | 411.396173 | -0.68709704 | 0.14358135 | -4.7854197 |
| ENSG00000092820.17 | EZR | 36.2134534 | -1.275288362 | 0.26730801 | -4.7708573 |
| ENSG00000136536.14 | 7-Mar | 274.887738 | 0.65741479 | 0.13775269 | 4.77242807 |
| ENSG00000272476.1 | AL024507.2 | 182.031942 | 0.628577224 | 0.13173155 | 4.77165294 |
| ENSG00000108298.11 | RPL19 | 58.9696581 | -0.954220457 | 0.20016921 | -4.7670692 |
| ENSG00000225822.4 | UBXN7-AS1 | 22.547778 | 1.038523727 | 0.21810277 | 4.76162565 |
| ENSG00000272320.1 | AL445309.1 | 14.5682239 | 1.923281123 | 0.40535927 | 4.7446334 |
| ENSG00000126768.12 | TIMM17B | 60.5307877 | -0.720521551 | 0.15192426 | -4.7426366 |
| ENSG00000170439.6 | METTL7B | 5.10016373 | 3.054480586 | 0.64476316 | 4.73736833 |
| ENSG00000259125.1 | LRP1-AS | 1005.60143 | 0.708569597 | 0.14959303 | 4.73664837 |
| ENSG00000066923.17 | STAG3 | 200.79437 | 1.018666249 | 0.21526348 | 4.73218329 |
| ENSG00000135678.11 | CPM | 163.244071 | 0.924493236 | 0.19542326 | 4.73072272 |
| ENSG00000279232.2 | AC008522.1 | 9246.12344 | 1.166269951 | 0.24675543 | 4.72642064 |
| ENSG00000283529.1 | AL035685.1 | 14.4110891 | 2.584938388 | 0.54702678 | 4.72543296 |
| ENSG00000184009.11 | ACTG1 | 235.575819 | -0.916592498 | 0.19415822 | -4.7208535 |
| ENSG00000176809.10 | LRRC37A3 | 7.4830392 | -2.157715854 | 0.4588137 | -4.7028147 |
| ENSG00000273489.1 | AC008264.2 | 250.397008 | 0.955477143 | 0.20315265 | 4.70324716 |
| ENSG00000118181.10 | RPS25 | 46.6366389 | -0.90129225 | 0.19226264 | -4.687818 |
| ENSG00000236051.7 | MYCBP2-AS1 | 360.080342 | 0.693423449 | 0.14818169 | 4.67954879 |
| ENSG00000223865.10 | HLA-DPB1 | 2.67340617 | -4.300054333 | 0.91981459 | -4.6749143 |
| ENSG00000285677.1 | | 804.978269 | -0.687591437 | 0.14713993 | -4.6730444 |
| ENSG00000124571.17 | XPO5 | 23.0466067 | 1.364008204 | 0.29257771 | 4.66203737 |
| ENSG00000226715.3 | LINC01709 | 19.1076907 | -1.076026857 | 0.23104004 | -4.6573177 |
| ENSG00000256040.2 | PAPPA-AS1 | 1383.94797 | 1.278299772 | 0.27444829 | 4.6577072 |
| ENSG00000002745.12 | WNT16 | 13.0577513 | 3.079837688 | 0.66193847 | 4.65275524 |
| ENSG00000282843.1 | AL590644.3 | 7.85836192 | -2.211805743 | 0.47578949 | -4.6487066 |
| ENSG00000235237.1 | Z82188.2 | 38.1224174 | -0.960294811 | 0.20664179 | -4.6471473 |
| ENSG00000251260.1 | WDFY3-AS1 | 130.006421 | 0.943623384 | 0.20356776 | 4.63542637 |
| ENSG00000197415.11 | VEPH1 | 252.728327 | 1.750126757 | 0.37862234 | 4.62235476 |
| ENSG00000279569.1 | AC020763.4 | 4401.17853 | 1.019934497 | 0.22078037 | 4.61967933 |
| ENSG00000258520.1 | AL359317.1 | 20.0121328 | 3.177060732 | 0.68818246 | 4.61659646 |
| ENSG00000232358.1 | AL050404.1 | 41.5561696 | -1.797278398 | 0.389692 | -4.6120484 |
| ENSG00000100234.11 | TIMP3 | 98.796517 | -2.011638243 | 0.43734017 | -4.5997107 |
| ENSG00000283375.1 | AC087521.4 | 394.891976 | 0.609797488 | 0.13292859 | 4.58740648 |
| ENSG00000149294.16 | NCAM1 | 6.56165092 | 2.844508486 | 0.62077801 | 4.58216693 |
| ENSG00000172345.13 | STARD5 | 12.3054035 | 1.609880825 | 0.35123911 | 4.58343271 |
| ENSG00000266208.1 | AC080112.1 | 13.7303864 | -1.753058876 | 0.38275601 | -4.580095 |
| ENSG00000253992.1 | AC103726.2 | 2.27235554 | 4.105007642 | 0.89874308 | 4.56749846 |
| ENSG00000279122.1 | AC020763.2 | 2.36244258 | 4.313899134 | 0.94489966 | 4.56545737 |
| ENSG00000226605.1 | AC092567.1 | 167.1919 | 1.20235255 | 0.26416402 | 4.55153783 |
| ENSG00000233885.7 | YEATS2-AS1 | 268.755235 | 0.738308813 | 0.16234392 | 4.54780689 |
| ENSG00000264666.1 | AC020558.1 | 58.67669 | -0.725840181 | 0.15962038 | -4.5472901 |

| | | | | | |
|---|---|---|---|---|---|
| ENSG00000170889.13 | RPS9 | 15.1789441 | -1.212979547 | 0.26704587 | -4.5422142 |
| ENSG00000232832.1 | LMLN-AS1 | 44.399982 | 0.804843181 | 0.17724274 | 4.54090912 |
| ENSG00000127241.16 | MASP1 | 26.4943694 | 1.834290811 | 0.40464906 | 4.53304104 |
| ENSG00000233706.1 | AL353689.2 | 251.80299 | 0.67085911 | 0.14804664 | 4.53140381 |
| ENSG00000149262.16 | INTS4 | 8.8123716 | 1.583590237 | 0.35018428 | 4.52216258 |
| ENSG00000107821.14 | KAZALD1 | 7.00269404 | -2.588379988 | 0.57486434 | -4.5025927 |
| ENSG00000144741.17 | SLC25A26 | 194.954943 | 1.926259216 | 0.42761737 | 4.50463275 |
| ENSG00000271647.1 | KRT8P47 | 6.00305841 | 3.155564742 | 0.70071311 | 4.50336194 |
| ENSG00000143450.16 | OAZ3 | 125.103549 | -0.775485313 | 0.17228402 | -4.5012028 |
| ENSG00000186714.12 | CCDC73 | 292.791026 | -0.625994629 | 0.13921375 | -4.4966437 |
| ENSG00000197915.5 | HRNR | 6.45003249 | -2.246923807 | 0.50044057 | -4.4898914 |
| ENSG00000275327.1 | AL354950.2 | 57.789524 | 2.393533336 | 0.5335191 | 4.48631234 |
| ENSG00000118894.14 | EEF2KMT | 222.162629 | -0.460942282 | 0.10285484 | -4.4814837 |
| ENSG00000270751.1 | FBXW7-AS1 | 47.7328063 | 0.880393052 | 0.19710907 | 4.46652734 |
| ENSG00000267165.1 | CHMP1B-AS1 | 588.304133 | -1.455860147 | 0.32648679 | -4.4591701 |
| ENSG00000253304.1 | TMEM200B | 32.2270089 | 1.055292018 | 0.23703464 | 4.45205814 |
| ENSG00000138443.15 | ABI2 | 3199.76902 | 0.721860997 | 0.16231059 | 4.44740531 |
| ENSG00000198732.10 | SMOC1 | 2.6096443 | 4.223689257 | 0.94956397 | 4.44803024 |
| ENSG00000279333.1 | AC096636.1 | 12.2418195 | -2.91981942 | 0.65672691 | -4.4460176 |
| ENSG00000255629.1 | AC025576.1 | 162.061312 | 0.783968014 | 0.17641109 | 4.44398381 |
| ENSG00000234616.8 | JRK | 14.1095475 | 2.071310251 | 0.46687938 | 4.43649979 |
| ENSG00000254902.1 | ANO1-AS1 | 3.24322809 | -4.605127015 | 1.03788123 | -4.4370463 |
| ENSG00000105974.11 | CAV1 | 23.5808758 | 1.572219634 | 0.35497196 | 4.42913759 |
| ENSG00000259627.1 | AC079328.2 | 5527.26475 | -1.016020607 | 0.2294188 | -4.4286719 |
| ENSG00000166676.15 | TVP23A | 159.265075 | -0.625171525 | 0.14133351 | -4.423378 |
| ENSG00000225921.6 | NOL7 | 297.817327 | 0.590930605 | 0.1336296 | 4.42215335 |
| ENSG00000172613.7 | RAD9A | 265.515419 | -0.563405418 | 0.12760548 | -4.4152132 |
| ENSG00000162391.11 | FAM151A | 44.7235504 | -1.019809607 | 0.23126507 | -4.4097 |
| ENSG00000259188.5 | AC025040.1 | 1.44788727 | 4.271677463 | 0.96935609 | 4.40671647 |
| ENSG00000171790.15 | SLFNL1 | 21.3574234 | -1.937147084 | 0.4402493 | -4.4001139 |
| ENSG00000249252.5 | AC098829.1 | 52.9915348 | 0.986671885 | 0.22451473 | 4.39468657 |
| ENSG00000144355.14 | DLX1 | 5.82043466 | -2.087711907 | 0.47537405 | -4.3917246 |
| ENSG00000135250.16 | SRPK2 | 87.4370064 | 0.803287041 | 0.18296447 | 4.39039906 |
| ENSG00000175029.16 | CTBP2 | 519.442032 | 0.527789198 | 0.1202397 | 4.38947527 |
| ENSG00000196337.11 | CGB7 | 5.72499865 | -2.555613386 | 0.58380421 | -4.3775179 |
| ENSG00000138131.3 | LOXL4 | 10.2998188 | 2.476151807 | 0.5665329 | 4.37071137 |
| ENSG00000285608.1 | | 637.536728 | 0.812281582 | 0.18583279 | 4.37103485 |
| ENSG00000258959.2 | AL118558.1 | 1750.54668 | 0.566859411 | 0.12974908 | 4.36888976 |
| ENSG00000257605.2 | AC073611.1 | 115.626706 | -0.660385305 | 0.15129712 | -4.3648242 |
| ENSG00000166323.12 | C11orf65 | 62.4235384 | 0.946191124 | 0.21690818 | 4.36217352 |
| ENSG00000196923.13 | PDLIM7 | 30.2484606 | -1.186838451 | 0.27260476 | -4.3536967 |
| ENSG00000285051.1 | AC026316.4 | 14.1507494 | 4.536727749 | 1.04297324 | 4.34980265 |
| ENSG00000261716.2 | AC239868.1 | 45.3780979 | 0.81679215 | 0.18784632 | 4.34819345 |

| | | | | | |
|---|---|---|---|---|---|
| ENSG00000263873.1 | AP003396.5 | 7602.23615 | 1.164026537 | 0.26793658 | 4.34441063 |
| ENSG00000138079.13 | SLC3A1 | 337.119506 | 0.831637429 | 0.19166569 | 4.33899996 |
| ENSG00000165458.13 | INPPL1 | 8.90155787 | -1.53890734 | 0.35537009 | -4.3304357 |
| ENSG00000237768.2 | AL731563.2 | 406.364759 | 0.88521499 | 0.20444077 | 4.32993379 |
| ENSG00000236498.1 | AC107081.2 | 255.28547 | -0.543454946 | 0.12563203 | -4.3257676 |
| ENSG00000250186.3 | AC091180.3 | 702.170878 | -0.592135098 | 0.13687039 | -4.326247 |
| ENSG00000181350.11 | LRRC75A | 1364.26551 | -0.888307368 | 0.20546778 | -4.3233414 |
| ENSG00000264443.1 | AL445686.2 | 196.899168 | 0.765726075 | 0.17715883 | 4.32225737 |
| ENSG00000215375.6 | MYL5 | 513.567568 | -0.679496982 | 0.15747986 | -4.3148183 |
| ENSG00000160255.17 | ITGB2 | 12.1125134 | 2.001967473 | 0.46442035 | 4.31067987 |
| ENSG00000236107.8 | AC010127.1 | 52.6778976 | 1.774805608 | 0.41279179 | 4.29951771 |
| ENSG00000228830.1 | AL160408.2 | 722.256068 | 0.624506902 | 0.14543561 | 4.294044 |
| ENSG00000248890.1 | HHIP-AS1 | 12.037326 | -5.261959648 | 1.22751344 | -4.2866819 |
| ENSG00000223390.1 | AL445685.1 | 44.8705216 | -0.86107377 | 0.20098186 | -4.2843357 |
| ENSG00000177082.12 | WDR73 | 108.581237 | 0.897850422 | 0.20973918 | 4.280795 |
| ENSG00000086232.12 | EIF2AK1 | 230.065234 | -0.573607195 | 0.13411057 | -4.2771214 |
| ENSG00000174177.12 | CTU2 | 80.1688928 | 0.542209205 | 0.12708411 | 4.26653805 |
| ENSG00000267023.5 | LRRC37A16P | 665.074821 | 0.571426394 | 0.13390549 | 4.2673859 |
| ENSG00000279605.1 | AC067930.6 | 992.207397 | -0.583762431 | 0.13678422 | -4.2677615 |
| ENSG00000285649.1 | | 51.047398 | -0.665379841 | 0.15602021 | -4.2647028 |
| ENSG00000122729.18 | ACO1 | 6.94673458 | 1.605974176 | 0.37681457 | 4.26197469 |
| ENSG00000107731.12 | UNC5B | 4.10810742 | -3.04681963 | 0.71578723 | -4.2565996 |
| ENSG00000198791.11 | CNOT7 | 48.8616127 | 1.063640553 | 0.2502435 | 4.25042232 |
| ENSG00000205485.13 | AC004980.1 | 81.4878397 | -1.019202187 | 0.24006666 | -4.2454967 |
| ENSG00000228109.1 | MELTF-AS1 | 8.03665089 | -1.713175588 | 0.40343846 | -4.246436 |
| ENSG00000281371.1 | INE2 | 72.8307381 | 0.945340263 | 0.22280656 | 4.24287438 |
| ENSG00000146707.14 | POMZP3 | 26.3297387 | 1.685300683 | 0.39740912 | 4.24071966 |
| ENSG00000260285.1 | AL133367.1 | 22.563613 | 1.610339609 | 0.3800283 | 4.23742026 |
| ENSG00000226430.6 | USP17L7 | 2.40824434 | 4.878257016 | 1.15166117 | 4.23584398 |
| ENSG00000151748.14 | SAV1 | 35.9779319 | 1.687647409 | 0.39944633 | 4.22496663 |
| ENSG00000101665.9 | SMAD7 | 9.21079758 | -1.683201598 | 0.39891892 | -4.2194078 |
| ENSG00000226548.1 | AC016722.1 | 7.72567839 | 2.492198135 | 0.59081037 | 4.21827083 |
| ENSG00000125753.13 | VASP | 17.0000899 | -1.187675446 | 0.28177383 | -4.2149956 |
| ENSG00000006327.13 | TNFRSF12A | 15.2712044 | -1.252345082 | 0.29766154 | -4.2072788 |
| ENSG00000254721.1 | AP000879.1 | 404.24265 | -0.498367462 | 0.11846258 | -4.2069611 |
| ENSG00000269915.1 | AP006621.4 | 112.302869 | 0.549773153 | 0.13075771 | 4.20451816 |
| ENSG00000164022.16 | AIMP1 | 24.8163121 | -1.425389882 | 0.33932948 | -4.2006072 |
| ENSG00000124243.17 | BCAS4 | 27.5128645 | -1.069574307 | 0.25470721 | -4.1992306 |
| ENSG00000139637.13 | C12orf10 | 23.5346904 | -1.094293997 | 0.26071357 | -4.1973036 |
| ENSG00000263235.1 | AC006111.2 | 5.94908183 | 1.742803793 | 0.41521433 | 4.19735941 |
| ENSG00000169609.13 | C15orf40 | 114.384662 | -0.625310633 | 0.14908555 | -4.1943075 |
| ENSG00000107984.9 | DKK1 | 27.8262064 | -1.549937327 | 0.36982132 | -4.1910437 |
| ENSG00000125630.15 | POLR1B | 3.50924945 | 2.601051813 | 0.62045351 | 4.19217843 |

| | | | | | |
|---|---|---|---|---|---|
| ENSG00000156050.8 | FAM161B | 287.745126 | -0.512983688 | 0.12241497 | -4.1905308 |
| ENSG00000092931.11 | MFSD11 | 1192.38992 | -0.536286284 | 0.12803126 | -4.1887135 |
| ENSG00000258317.1 | AC034102.6 | 437.123416 | 0.547554281 | 0.13070058 | 4.18937909 |
| ENSG00000078795.16 | PKD2L2 | 288.901733 | 0.718226586 | 0.17150572 | 4.18777045 |
| ENSG00000243230.1 | AC011005.4 | 12.5551115 | -1.296375746 | 0.3098915 | -4.1833214 |
| ENSG00000121310.16 | ECHDC2 | 3.95926401 | -2.836641424 | 0.67921054 | -4.1763802 |
| ENSG00000267064.1 | UXT-AS1 | 160.90513 | -0.844296213 | 0.20216716 | -4.1762284 |
| ENSG00000235512.1 | TAB3-AS2 | 92.1697258 | 0.887541379 | 0.21257563 | 4.17517931 |
| ENSG00000249889.1 | ALG1L11P | 4.96638794 | 2.453315288 | 0.58786691 | 4.17324952 |
| ENSG00000214140.10 | PRCD | 86.1704991 | -1.504224098 | 0.36084009 | -4.1686723 |
| ENSG00000182934.11 | SRPRA | 46.6877239 | 0.819293707 | 0.1967212 | 4.16474529 |
| ENSG00000144567.10 | RETREG2 | 172.162853 | 0.634530697 | 0.15246882 | 4.16170796 |
| ENSG00000100292.16 | HMOX1 | 22.7477813 | -1.338049834 | 0.32165046 | -4.15995 |
| ENSG00000136942.14 | RPL35 | 18.3830134 | -1.192515244 | 0.28679265 | -4.1581094 |
| ENSG00000285908.1 | | 98.6099112 | -0.957732601 | 0.23041711 | -4.1565168 |
| ENSG00000227487.3 | NCAM1-AS1 | 51.6974165 | 3.014313697 | 0.72630952 | 4.15017786 |
| ENSG00000165801.9 | ARHGEF40 | 57.1511373 | 0.755860831 | 0.18218864 | 4.14878133 |
| ENSG00000237125.9 | HAND2-AS1 | 88.3561043 | 1.615615089 | 0.38954102 | 4.14748382 |
| ENSG00000232675.7 | AL121830.1 | 2.58009334 | 4.972630501 | 1.20358424 | 4.13151846 |
| ENSG00000225407.3 | AC025188.1 | 6.83246074 | -1.613333645 | 0.39063627 | -4.130015 |
| ENSG00000185112.5 | FAM43A | 1.77709894 | -3.419570567 | 0.82865137 | -4.1266698 |
| ENSG00000270084.1 | GAS5-AS1 | 68.1479855 | -0.893996338 | 0.21669524 | -4.1255929 |
| ENSG00000090266.12 | NDUFB2 | 101.487946 | 0.580572596 | 0.14079453 | 4.123545 |
| ENSG00000099904.15 | ZDHHC8 | 12.8180805 | -1.914165537 | 0.46424186 | -4.1232076 |
| ENSG00000260001.6 | TGFBR3L | 5.80334696 | -2.106288619 | 0.51106614 | -4.1213621 |
| ENSG00000285582.1 | | 1.80429366 | 4.097038975 | 0.99516007 | 4.11696481 |
| ENSG00000247416.3 | AP000802.1 | 10.8862528 | 2.930921384 | 0.71426085 | 4.10343276 |
| ENSG00000134480.14 | CCNH | 418.451912 | 0.739144865 | 0.18022468 | 4.10124118 |
| ENSG00000100345.21 | MYH9 | 318.985019 | -0.832303847 | 0.20312386 | -4.0975188 |
| ENSG00000268364.1 | SMC5-AS1 | 4.33881913 | 3.000834534 | 0.73228694 | 4.09789437 |
| ENSG00000225950.8 | NTF4 | 23.7557997 | -2.23459391 | 0.54634799 | -4.0900561 |
| ENSG00000079805.16 | DNM2 | 323.160178 | -0.736151232 | 0.18080723 | -4.0714701 |
| ENSG00000197815.4 | AC122129.1 | 52.2042081 | 0.709215287 | 0.17420701 | 4.07110639 |
| ENSG00000251630.1 | AC119751.6 | 4.31832856 | 3.916146194 | 0.96138293 | 4.07345095 |
| ENSG00000260367.2 | AC109460.1 | 117.057565 | -0.574393177 | 0.14109135 | -4.0710731 |
| ENSG00000137573.13 | SULF1 | 47.385751 | -2.086537746 | 0.51275682 | -4.069254 |
| ENSG00000130176.7 | CNN1 | 4.53938924 | -2.821952375 | 0.69369704 | -4.0679897 |
| ENSG00000130956.13 | HABP4 | 54.1196266 | 0.923346126 | 0.22726563 | 4.06284982 |
| ENSG00000266933.2 | AC005775.1 | 97.0187415 | -1.038125578 | 0.2555679 | -4.0620343 |
| ENSG00000269349.1 | AC022150.3 | 26.6802882 | 1.336306327 | 0.3290455 | 4.06115966 |
| ENSG00000272182.1 | AC135507.1 | 393.100527 | -0.572983898 | 0.14111645 | -4.0603623 |
| ENSG00000234055.1 | AL158151.2 | 120.366431 | 0.660932545 | 0.1629755 | 4.05541044 |
| ENSG00000261762.1 | AC027228.2 | 4.27913385 | -3.453100058 | 0.8529829 | -4.0482641 |

| | | | | | |
|---|---|---|---|---|---|
| ENSG00000126453.9 | BCL2L12 | 74.6361545 | -0.572574401 | 0.1417412 | -4.0395764 |
| ENSG00000263272.1 | AC004148.2 | 61.4880556 | -0.839565648 | 0.20799907 | -4.0363913 |
| ENSG00000255886.1 | AC020611.2 | 81.0950292 | 1.06634479 | 0.26442592 | 4.0326788 |
| ENSG00000164708.5 | PGAM2 | 35.3383501 | 0.908663467 | 0.22581111 | 4.02399805 |
| ENSG00000221916.3 | C19orf73 | 14.039303 | -1.147136437 | 0.28527498 | -4.0211604 |
| ENSG00000109686.17 | SH3D19 | 2726.56965 | -0.923409097 | 0.23020118 | -4.0113135 |
| ENSG00000249087.6 | ZNF436-AS1 | 25.8963689 | 1.329578929 | 0.33140648 | 4.01192802 |
| ENSG00000152700.13 | SAR1B | 129.291551 | 0.49889257 | 0.12447731 | 4.00789975 |
| ENSG00000167157.10 | PRRX2 | 13.6070465 | -1.52344588 | 0.38004531 | -4.0085901 |
| ENSG00000125968.8 | ID1 | 12.0463223 | -1.84668211 | 0.46186984 | -3.9982739 |
| ENSG00000253368.3 | TRNP1 | 10.7889837 | -1.526937125 | 0.38228035 | -3.9942862 |
| ENSG00000157570.11 | TSPAN18 | 53.1115515 | 2.051578683 | 0.51375127 | 3.99333063 |
| ENSG00000151466.11 | SCLT1 | 74.3864219 | 1.084269825 | 0.27170023 | 3.99068416 |
| ENSG00000161970.14 | RPL26 | 15.3188823 | -1.252691631 | 0.31392922 | -3.9903633 |
| ENSG00000239718.1 | HLTF-AS1 | 28.9980951 | 0.841717648 | 0.21110767 | 3.98714861 |
| ENSG00000108474.16 | PIGL | 7.26610324 | -1.99341076 | 0.5001991 | -3.9852346 |
| ENSG00000170222.11 | ADPRM | 7.82596773 | -1.648367709 | 0.41355221 | -3.9858757 |
| ENSG00000034510.5 | TMSB10 | 109.509851 | -0.904525325 | 0.22717318 | -3.9816555 |
| ENSG00000115484.14 | CCT4 | 4.69729223 | -1.998186837 | 0.5017948 | -3.9820796 |
| ENSG00000179454.13 | KLHL28 | 34.5578727 | 0.715023415 | 0.17988519 | 3.97488762 |
| ENSG00000176102.12 | CSTF3 | 73.7678585 | 0.687809782 | 0.17322362 | 3.97064652 |
| ENSG00000154025.15 | SLC5A10 | 285.139198 | -1.11018259 | 0.27989269 | -3.966458 |
| ENSG00000244165.1 | P2RY11 | 208.565296 | -0.56506153 | 0.14244903 | -3.9667629 |
| ENSG00000266043.1 | MIR3649 | 7.0621696 | 2.028386275 | 0.51135333 | 3.96670202 |
| ENSG00000170854.17 | RIOX2 | 236.151439 | 0.817053836 | 0.20617257 | 3.96296099 |
| ENSG00000145362.17 | ANK2 | 8.89246812 | 2.462118429 | 0.62190292 | 3.95900767 |
| ENSG00000170315.13 | UBB | 18.3173281 | -1.107325891 | 0.27989693 | -3.9561916 |
| ENSG00000186812.12 | ZNF397 | 56.7220493 | 0.649483776 | 0.16414786 | 3.95669953 |
| ENSG00000188522.14 | FAM83G | 8.36300217 | -1.500996569 | 0.37928511 | -3.9574361 |
| ENSG00000236671.8 | PRKG1-AS1 | 2268.22872 | -1.253588492 | 0.31705737 | -3.9538222 |
| ENSG00000033011.12 | ALG1 | 86.9327405 | -0.553988416 | 0.14020531 | -3.9512655 |
| ENSG00000267379.1 | AC008569.1 | 356.766339 | 0.748502591 | 0.18940456 | 3.95187198 |
| ENSG00000275807.1 | AC145285.6 | 66.1957867 | -0.59439159 | 0.15040701 | -3.9518874 |
| ENSG00000176700.20 | SCAND2P | 341.618129 | -0.353803084 | 0.08971415 | -3.9436708 |
| ENSG00000257769.1 | AC026401.1 | 51.2067448 | 0.812290033 | 0.20601221 | 3.94292184 |
| ENSG00000101333.16 | PLCB4 | 3.79910426 | 2.039468456 | 0.51788857 | 3.93804491 |
| ENSG00000105825.12 | TFPI2 | 4.72823719 | -4.158189733 | 1.05599069 | -3.9377144 |
| ENSG00000269621.1 | AL589765.7 | 124.250099 | -0.65359372 | 0.16610026 | -3.9349349 |
| ENSG00000115486.11 | GGCX | 601.489082 | -0.336393372 | 0.0856574 | -3.9271958 |
| ENSG00000204577.11 | LILRB3 | 13.098244 | -1.791995513 | 0.45671133 | -3.923694 |
| ENSG00000171988.18 | JMJD1C | 10.5112782 | -1.371853636 | 0.35047478 | -3.914272 |
| ENSG00000230953.2 | AC099677.1 | 8.90700259 | -1.600090361 | 0.40933654 | -3.9089849 |
| ENSG00000133110.14 | POSTN | 15.738218 | 2.152890545 | 0.55089191 | 3.90800898 |

| | | | | | |
|---|---|---|---|---|---|
| ENSG00000130024.14 | PHF10 | 225.505152 | 0.601630588 | 0.15398617 | 3.90704311 |
| ENSG00000125741.4 | OPA3 | 19.6259158 | -0.957792808 | 0.24525131 | -3.9053524 |
| ENSG00000072110.13 | ACTN1 | 46.0727824 | -0.990559865 | 0.25406467 | -3.8988493 |
| ENSG00000184451.5 | CCR10 | 98.53796 | 0.704218133 | 0.18055711 | 3.90025147 |
| ENSG00000197756.9 | RPL37A | 25.2471506 | -1.119519672 | 0.28720787 | -3.8979421 |
| ENSG00000233295.3 | FAM90A20P | 2.28752792 | -3.718111896 | 0.95378256 | -3.8982805 |
| ENSG00000263218.2 | AC127496.6 | 212.001411 | -0.631623527 | 0.16195416 | -3.9000143 |
| ENSG00000283849.1 | AC092053.2 | 12.7200913 | -1.23078689 | 0.3156979 | -3.8986224 |
| ENSG00000128652.11 | HOXD3 | 15.4743005 | -1.230180423 | 0.31580379 | -3.8953947 |
| ENSG00000276578.1 | AC004696.1 | 71.3752667 | -1.044773093 | 0.26820118 | -3.8954829 |
| ENSG00000224383.7 | PRR29 | 2.82807378 | -2.793046804 | 0.71731849 | -3.8937332 |
| ENSG00000231345.3 | BEND3P1 | 12.9643133 | -2.691508367 | 0.69130029 | -3.8933997 |
| ENSG00000237278.2 | RLIMP2 | 42.0007676 | 0.788653649 | 0.2026401 | 3.89189327 |
| ENSG00000254943.1 | AP003501.2 | 3.00657334 | 2.387916109 | 0.61423813 | 3.88760644 |
| ENSG00000125457.14 | MIF4GD | 38.9650518 | -0.715515001 | 0.1841276 | -3.8859737 |
| ENSG00000129048.6 | ACKR4 | 78.043777 | 0.884771709 | 0.2277632 | 3.88461229 |
| ENSG00000146243.13 | IRAK1BP1 | 90.1262006 | 0.817342168 | 0.21047186 | 3.88337983 |
| ENSG00000136854.20 | STXBP1 | 35.1783469 | 0.857740154 | 0.22093693 | 3.88228512 |
| ENSG00000122641.10 | INHBA | 10.450059 | -2.771781237 | 0.71425766 | -3.8806462 |
| ENSG00000104375.16 | STK3 | 8.92112032 | 1.749649814 | 0.45096637 | 3.87977895 |
| ENSG00000164776.9 | PHKG1 | 497.585412 | -0.343403265 | 0.08856035 | -3.8776188 |
| ENSG00000224789.1 | AC012363.1 | 39.0342412 | 0.885276582 | 0.22832064 | 3.87733921 |
| ENSG00000001460.17 | STPG1 | 134.168826 | 0.575047195 | 0.14858404 | 3.87018147 |
| ENSG00000246250.2 | AC087521.2 | 205.89246 | 0.600605826 | 0.15533838 | 3.86643559 |
| ENSG00000152661.8 | GJA1 | 2.9139352 | 2.65236251 | 0.6862602 | 3.86495168 |
| ENSG00000131737.5 | KRT34 | 5.13168039 | 3.509847554 | 0.91069556 | 3.85402952 |
| ENSG00000232284.7 | GNG12-AS1 | 43.4490272 | -1.913636865 | 0.49673866 | -3.8524017 |
| ENSG00000159176.13 | CSRP1 | 14.3226253 | -1.005717042 | 0.26166729 | -3.8434954 |
| ENSG00000254254.5 | AC012349.1 | 132.505398 | -3.349918529 | 0.87304366 | -3.8370573 |
| ENSG00000261312.1 | AC002550.1 | 76.7182077 | -0.590481191 | 0.15388062 | -3.8372681 |
| ENSG00000283208.1 | AC001226.2 | 132.217475 | 0.836551317 | 0.21797182 | 3.83788742 |
| ENSG00000277182.1 | AC006449.5 | 4.94681489 | -2.051841169 | 0.53516981 | -3.8340002 |
| ENSG00000197982.13 | C1orf122 | 54.4684783 | -0.734235905 | 0.19153984 | -3.8333326 |
| ENSG00000250909.1 | AC138956.1 | 414.146148 | 0.597315931 | 0.15584803 | 3.83268186 |
| ENSG00000197457.9 | STMN3 | 3.87119937 | -2.559085635 | 0.6680335 | -3.8307744 |
| ENSG00000240567.1 | LINC02067 | 24.8868536 | -0.862752266 | 0.22529874 | -3.82937 |
| ENSG00000259353.1 | AC090515.4 | 33.3045126 | 1.094439064 | 0.28592656 | 3.82769285 |
| ENSG00000063177.12 | RPL18 | 28.8239801 | -1.463350111 | 0.38272512 | -3.8235016 |
| ENSG00000225610.1 | AC007679.1 | 37.7147434 | 0.908889889 | 0.23816387 | 3.81623742 |
| ENSG00000235927.4 | NEXN-AS1 | 87.0556374 | -0.690824281 | 0.18116452 | -3.8132428 |
| ENSG00000179965.11 | ZNF771 | 77.9788169 | -0.925130137 | 0.24282343 | -3.8098883 |
| ENSG00000255201.1 | AC087623.2 | 247.163117 | 0.522814333 | 0.13739958 | 3.80506497 |
| ENSG00000014919.12 | COX15 | 90.1325328 | -1.172066359 | 0.30862873 | -3.797658 |

| | | | | | |
|---|---|---|---|---|---|
| ENSG00000123243.14 | ITIH5 | 2.38192807 | 4.343913389 | 1.14401137 | 3.79708935 |
| ENSG00000105662.15 | CRTC1 | 6.53496365 | -3.038699178 | 0.80159262 | -3.7908273 |
| ENSG00000227617.8 | CERS6-AS1 | 384.701616 | 0.642486825 | 0.16966203 | 3.78686279 |
| ENSG00000104450.12 | SPAG1 | 128.049699 | 0.525364055 | 0.13885118 | 3.78364843 |
| ENSG00000235426.2 | AL133481.1 | 237.750208 | 0.544069721 | 0.14377481 | 3.78417966 |
| ENSG00000155657.26 | TTN | 510.441916 | 0.777104201 | 0.20555764 | 3.78046863 |
| ENSG00000140988.15 | RPS2 | 58.1564938 | -1.094808206 | 0.28971172 | -3.7789573 |
| ENSG00000102977.14 | ACD | 5.74194405 | -1.939217868 | 0.51338996 | -3.7772805 |
| ENSG00000150967.17 | ABCB9 | 596.579912 | -0.332826058 | 0.08812115 | -3.7769144 |
| ENSG00000234771.3 | SLC25A25-AS | 52.3022875 | -0.750532265 | 0.19870692 | -3.7770817 |
| ENSG00000265618.1 | AC002094.2 | 130.142323 | -0.622692266 | 0.16497079 | -3.7745606 |
| ENSG00000203472.3 | AC009163.1 | 2.44274829 | -3.840446937 | 1.01973829 | -3.7661104 |
| ENSG00000082805.19 | ERC1 | 8.33082926 | 1.476823024 | 0.39226433 | 3.76486701 |
| ENSG00000013016.15 | EHD3 | 2.55802196 | -2.795146845 | 0.74303242 | -3.7618101 |
| ENSG00000164985.14 | PSIP1 | 93.8920765 | 0.980819878 | 0.26080114 | 3.76079591 |
| ENSG00000260192.2 | LINC02240 | 14.3741927 | -1.358875749 | 0.36216798 | -3.7520593 |
| ENSG00000266049.1 | AP001011.1 | 36.3892063 | 0.910866439 | 0.24272502 | 3.75266806 |
| ENSG00000112237.12 | CCNC | 23.1815464 | 0.939774431 | 0.25133987 | 3.73905826 |
| ENSG00000259113.1 | AL118556.1 | 226.413558 | 0.648954938 | 0.17365887 | 3.73695237 |
| ENSG00000218336.8 | TENM3 | 24.63388 | -1.037578469 | 0.27774789 | -3.7356844 |
| ENSG00000263177.1 | MTND1P8 | 2.65794706 | 2.93014127 | 0.78483592 | 3.73344438 |
| ENSG00000252680.1 | RNA5SP449 | 1.65602852 | -3.584442425 | 0.96025461 | -3.7328042 |
| ENSG00000245970.2 | AP003352.1 | 2420.55363 | -0.649483408 | 0.17427289 | -3.7268183 |
| ENSG00000265399.1 | AP005329.2 | 15.0690168 | -1.183319753 | 0.31749374 | -3.7270648 |
| ENSG00000151632.17 | AKR1C2 | 18.0143348 | 1.070232666 | 0.28749112 | 3.72266333 |
| ENSG00000160226.15 | C21orf2 | 89.5941409 | 0.773848345 | 0.20779972 | 3.72401051 |
| ENSG00000236883.1 | AP001615.1 | 27.6217736 | -1.89463803 | 0.50891829 | -3.7228727 |
| ENSG00000271969.1 | U47924.1 | 20.001582 | -0.894405846 | 0.24026616 | -3.7225628 |
| ENSG00000227906.7 | SNAP25-AS1 | 12.6482692 | 1.748977675 | 0.47143416 | 3.7099087 |
| ENSG00000260246.1 | AC000032.1 | 84.8504483 | 2.414983136 | 0.65100259 | 3.70963678 |
| ENSG00000170561.12 | IRX2 | 3.0195968 | -2.299464639 | 0.62086474 | -3.7036483 |
| ENSG00000237298.9 | TTN-AS1 | 7.15545521 | 2.461991048 | 0.66471606 | 3.70382362 |
| ENSG00000150787.7 | PTS | 1.79401242 | -3.757086653 | 1.01468019 | -3.7027299 |
| ENSG00000220323.4 | HIST2H2BD | 28.4258567 | 0.81154636 | 0.21925846 | 3.70132297 |
| ENSG00000259409.1 | BMF-AS1 | 28.3638337 | -1.650866304 | 0.44597756 | -3.7016802 |
| ENSG00000258777.1 | HIF1A-AS1 | 1039.10867 | 0.789989429 | 0.21348163 | 3.70050311 |
| ENSG00000237945.7 | LINC00649 | 202.968175 | -0.665053092 | 0.17979474 | -3.6989574 |
| ENSG00000259006.1 | AC092143.2 | 52.1697736 | -1.080514595 | 0.29211604 | -3.6989225 |
| ENSG00000136918.7 | WDR38 | 158.238135 | -0.957670653 | 0.2590514 | -3.6968365 |
| ENSG00000204291.10 | COL15A1 | 7.07777456 | 1.726082459 | 0.467184 | 3.69465237 |
| ENSG00000246283.2 | AC090510.1 | 140.52504 | 0.60084523 | 0.16262398 | 3.69469022 |
| ENSG00000188846.13 | RPL14 | 21.3269755 | -0.947950567 | 0.2568605 | -3.6905268 |
| ENSG00000115414.18 | FN1 | 1293.48701 | 0.779804321 | 0.21150917 | 3.68685826 |

| | | | | | |
|---|---|---|---|---|---|
| ENSG00000177640.15 | CASC2 | 50.1757202 | 0.770316293 | 0.20906346 | 3.68460504 |
| ENSG00000255920.2 | CCND2-AS1 | 6.9646242 | 3.413550078 | 0.92697352 | 3.68246774 |
| ENSG00000138496.16 | PARP9 | 20.0164822 | 1.103093403 | 0.30004476 | 3.67642952 |
| ENSG00000140400.16 | MAN2C1 | 9.7359985 | -1.195470093 | 0.32517094 | -3.6764358 |
| ENSG00000177971.8 | IMP3 | 218.233175 | 0.558227843 | 0.15187375 | 3.67560444 |
| ENSG00000145703.15 | IQGAP2 | 2.72123811 | -2.939653451 | 0.80028007 | -3.6732809 |
| ENSG00000216331.2 | HIST1H1PS1 | 10.3048292 | 2.628235446 | 0.7159167 | 3.67114702 |
| ENSG00000280414.1 | AC018470.1 | 20.421196 | -5.207887421 | 1.41881797 | -3.6705818 |
| ENSG00000179348.11 | GATA2 | 10.9499681 | -1.62340491 | 0.44252793 | -3.6684801 |
| ENSG00000258908.1 | AL355075.3 | 274.335789 | -1.117502545 | 0.30476066 | -3.6668202 |
| ENSG00000273132.1 | AL355312.3 | 86.8999983 | 0.601284006 | 0.16398093 | 3.66679219 |
| ENSG00000241202.1 | ZIC4-AS1 | 37.731654 | 1.862140887 | 0.50814745 | 3.66456799 |
| ENSG00000033800.13 | PIAS1 | 22.8267411 | -1.076140589 | 0.29388301 | -3.6617992 |
| ENSG00000120075.5 | HOXB5 | 10.5452003 | 2.716536912 | 0.74209242 | 3.660645 |
| ENSG00000272851.1 | AC096772.1 | 226.542311 | 0.767682316 | 0.20971755 | 3.66055351 |
| ENSG00000072832.14 | CRMP1 | 735.228743 | 0.420538204 | 0.11491947 | 3.65941637 |
| ENSG00000128346.10 | C22orf23 | 363.178252 | 0.545848491 | 0.14921577 | 3.6581154 |
| ENSG00000197948.10 | FCHSD1 | 87.1394424 | -0.481577456 | 0.13164709 | -3.6580941 |
| ENSG00000278259.4 | MYO19 | 68.6827213 | -0.60873315 | 0.16635691 | -3.6591998 |
| ENSG00000233893.2 | EZR-AS1 | 150.662701 | -0.924935943 | 0.25290915 | -3.6571866 |
| ENSG00000143631.10 | FLG | 113.786629 | -2.26300763 | 0.61906515 | -3.6555242 |
| ENSG00000188976.10 | NOC2L | 206.342439 | -1.67124685 | 0.45761985 | -3.6520419 |
| ENSG00000265298.1 | AC132812.1 | 90.0037155 | 0.841097013 | 0.23055038 | 3.64821358 |
| ENSG00000186710.11 | CFAP73 | 269.318996 | -0.396254347 | 0.10875766 | -3.6434616 |
| ENSG00000257761.1 | AC078860.1 | 23.4943456 | 2.934881326 | 0.80559571 | 3.64311936 |
| ENSG00000255508.7 | AP002990.1 | 34.8751999 | -0.695334157 | 0.19109055 | -3.6387679 |
| ENSG00000268475.1 | AC011462.2 | 23.5576754 | 0.934561545 | 0.25681506 | 3.63904492 |
| ENSG00000214128.10 | TMEM213 | 59.2994995 | 0.779060421 | 0.21422655 | 3.63661942 |
| ENSG00000254418.1 | SPON1-AS1 | 25.9833351 | -3.252323509 | 0.89472683 | -3.6349905 |
| ENSG00000168763.15 | CNNM3 | 26.3635454 | 0.809562947 | 0.22286365 | 3.6325481 |
| ENSG00000113273.16 | ARSB | 10.021566 | -1.114028952 | 0.30698748 | -3.6289068 |
| ENSG00000268006.1 | PTOV1-AS1 | 270.258896 | 0.423635982 | 0.11678692 | 3.62742673 |
| ENSG00000257663.1 | AC025259.1 | 384.432118 | -0.44129577 | 0.12172079 | -3.6254756 |
| ENSG00000164117.13 | FBXO8 | 15.6043368 | 0.891148903 | 0.24600077 | 3.6225452 |
| ENSG00000255542.1 | AC090625.2 | 3.66385764 | 1.859919012 | 0.51339806 | 3.62276207 |
| ENSG00000131871.14 | SELENOS | 1.65135367 | -3.271898087 | 0.90462292 | -3.616864 |
| ENSG00000108352.12 | RAPGEFL1 | 2.95402166 | 2.398832914 | 0.66356591 | 3.61506349 |
| ENSG00000103145.10 | HCFC1R1 | 42.9782135 | -0.61666874 | 0.17066358 | -3.6133587 |
| ENSG00000104892.16 | KLC3 | 171.430904 | -0.62664592 | 0.17338267 | -3.6142362 |
| ENSG00000168273.7 | SMIM4 | 122.241712 | 0.66520145 | 0.18406979 | 3.61385451 |
| ENSG00000259052.1 | AL157871.6 | 70.4527424 | 0.635982432 | 0.17631447 | 3.60709152 |
| ENSG00000123144.10 | TRIR | 23.765833 | -1.166709501 | 0.32443576 | -3.5961187 |
| ENSG00000129636.12 | ITFG1 | 51.9198344 | 0.771702721 | 0.2148139 | 3.59242457 |

| | | | | | |
|---|---|---|---|---|---|
| ENSG00000229893.2 | AC005091.1 | 153.276273 | 0.563661692 | 0.15692252 | 3.59197455 |
| ENSG00000090061.17 | CCNK | 44.6507178 | -1.029328523 | 0.28668752 | -3.5904198 |
| ENSG00000182993.4 | C12orf60 | 36.4951218 | -3.734898175 | 1.04045225 | -3.5896872 |
| ENSG00000270835.2 | AP001425.1 | 1.88781774 | 3.724865867 | 1.03809354 | 3.58817941 |
| ENSG00000256699.1 | TMEM132D-/ | 13.6288414 | 3.486272207 | 0.97317421 | 3.58237217 |
| ENSG00000256001.1 | AC079949.1 | 9.62591909 | 3.837254539 | 1.07201145 | 3.57949025 |
| ENSG00000257553.1 | AC034102.4 | 473.951445 | -0.406125084 | 0.11345028 | -3.5797627 |
| ENSG00000267342.1 | AC087289.2 | 83.9216803 | -0.612999508 | 0.17127118 | -3.5791165 |
| ENSG00000258425.1 | AC013451.1 | 1614.01658 | 0.734817211 | 0.20547365 | 3.57621139 |
| ENSG00000101082.13 | SLA2 | 163.119429 | -0.48933416 | 0.13693812 | -3.5733962 |
| ENSG00000163516.13 | ANKZF1 | 9.25444452 | 1.08029729 | 0.3023293 | 3.57324708 |
| ENSG00000188992.11 | LIPI | 1.33311306 | 3.918936133 | 1.09644691 | 3.57421422 |
| ENSG00000272325.1 | NUDT3 | 126.287614 | 0.77722472 | 0.21769069 | 3.57031682 |
| ENSG00000115461.4 | IGFBP5 | 26.3380073 | -2.113623187 | 0.59212596 | -3.5695499 |
| ENSG00000213904.8 | LIPE-AS1 | 85.095789 | 0.788503984 | 0.22093921 | 3.56887303 |
| ENSG00000181800.5 | CELF2-AS1 | 6.41591292 | 2.565508466 | 0.71898572 | 3.56823288 |
| ENSG00000064601.18 | CTSA | 49.7479262 | -0.503232438 | 0.14105443 | -3.5676473 |
| ENSG00000099219.13 | ERMP1 | 137.354923 | 0.491578021 | 0.13799682 | 3.56224165 |
| ENSG00000141552.17 | ANAPC11 | 80.0398994 | -0.500702223 | 0.14068639 | -3.5589955 |
| ENSG00000083845.8 | RPS5 | 14.6997167 | -1.34595841 | 0.37824611 | -3.5584197 |
| ENSG00000225670.4 | CADM3-AS1 | 2.61655512 | 4.405425747 | 1.24029318 | 3.55192289 |
| ENSG00000253356.1 | AC084024.3 | 342.306568 | -0.544110922 | 0.15354455 | -3.543668 |
| ENSG00000258034.1 | AC012157.1 | 162.134413 | 0.496428215 | 0.14010523 | 3.54325263 |
| ENSG00000264577.1 | AC010761.1 | 2527.96856 | -0.433744341 | 0.12248757 | -3.5411295 |
| ENSG00000166888.11 | STAT6 | 525.797111 | 0.50250825 | 0.14198237 | 3.53922984 |
| ENSG00000124181.14 | PLCG1 | 201.398918 | 0.466908708 | 0.13195065 | 3.53851021 |
| ENSG00000269926.1 | DDIT4-AS1 | 473.50411 | 1.095971437 | 0.31019073 | 3.53321792 |
| ENSG00000127528.5 | KLF2 | 3.78019837 | -2.107911103 | 0.5972201 | -3.5295381 |
| ENSG00000263326.1 | AC133552.4 | 2.4127407 | 2.154609999 | 0.61035981 | 3.53006535 |
| ENSG00000285921.1 | | 98.1717846 | 0.495604673 | 0.14043517 | 3.52906373 |
| ENSG00000111052.7 | LIN7A | 3.64470379 | 1.906092187 | 0.54044257 | 3.52690975 |
| ENSG00000259589.2 | AC073167.1 | 185.705041 | 0.66416968 | 0.18834714 | 3.52630611 |
| ENSG00000265148.5 | TSPOAP1-AS1 | 374.565005 | -0.493412073 | 0.13997929 | -3.5248935 |
| ENSG00000166140.17 | ZFYVE19 | 25.1282462 | -0.73526219 | 0.2087127 | -3.5228435 |
| ENSG00000121897.14 | LIAS | 441.969717 | -1.133380726 | 0.32209986 | -3.5187247 |
| ENSG00000154277.12 | UCHL1 | 3.19647801 | -2.495571533 | 0.7094702 | -3.5175143 |
| ENSG00000182463.15 | TSHZ2 | 2.18151839 | -3.281987559 | 0.93321583 | -3.516858 |
| ENSG00000111752.10 | PHC1 | 235.778413 | -0.574738519 | 0.16347711 | -3.5157124 |
| ENSG00000187514.16 | PTMA | 48.9692809 | -0.949409942 | 0.27008237 | -3.5152607 |
| ENSG00000106624.10 | AEBP1 | 178.900497 | -0.846394629 | 0.24086057 | -3.514044 |
| ENSG00000182325.10 | FBXL6 | 295.688032 | -0.447178375 | 0.12735371 | -3.5113102 |
| ENSG00000201778.1 | RF00019 | 23.7037282 | 1.110361422 | 0.31625059 | 3.51101767 |
| ENSG00000164587.12 | RPS14 | 21.6434426 | -1.080549958 | 0.30796724 | -3.5086523 |

| | | | | | |
|---|---|---|---|---|---|
| ENSG00000238279.1 | BX470102.1 | 37.4673101 | -0.622255017 | 0.17746279 | -3.5063972 |
| ENSG00000270075.1 | AL162742.2 | 162.512578 | -0.741977431 | 0.21190478 | -3.5014662 |
| ENSG00000161981.10 | SNRNP25 | 44.749881 | -0.878773422 | 0.25102793 | -3.5006997 |
| ENSG00000164105.3 | SAP30 | 3.90230557 | 2.635118047 | 0.75284761 | 3.5002011 |
| ENSG00000285698.1 | | 5.28967822 | 2.102559791 | 0.60077445 | 3.49974903 |
| ENSG00000269313.5 | MAGIX | 2.30763061 | 2.808653181 | 0.80278068 | 3.49865569 |
| ENSG00000254923.1 | AC130366.1 | 2.00498801 | 3.769504221 | 1.07792978 | 3.49698496 |
| ENSG00000260051.1 | AL031600.1 | 16.4244948 | -1.504212354 | 0.43073122 | -3.4922297 |
| ENSG00000078674.17 | PCM1 | 2.56892616 | 2.792349857 | 0.80010414 | 3.48998302 |
| ENSG00000234166.1 | ARHGEF19-AS | 23.8161419 | -1.040879957 | 0.29832424 | -3.4890894 |
| ENSG00000226445.1 | BX322234.1 | 12.6402578 | -1.541428822 | 0.44269375 | -3.4819304 |
| ENSG00000150051.13 | MKX | 6.05040568 | -1.670532648 | 0.48003045 | -3.4800556 |
| ENSG00000258114.1 | AC005871.2 | 390.546683 | 1.243732105 | 0.3577467 | 3.47657189 |
| ENSG00000170379.20 | TCAF2 | 2.93204755 | 2.978872545 | 0.85701956 | 3.4758513 |
| ENSG00000248464.1 | FGF10-AS1 | 2.21557965 | 3.626854341 | 1.04355301 | 3.47548644 |
| ENSG00000223764.2 | AL645608.1 | 22.1102243 | -1.723288026 | 0.49632798 | -3.4720751 |
| ENSG00000280339.1 | AP001528.3 | 3.36249868 | 3.284231179 | 0.94586055 | 3.472215 |
| ENSG00000272418.1 | AC090607.4 | 13.8966336 | -1.083199938 | 0.31236257 | -3.4677648 |
| ENSG00000000460.16 | C1orf112 | 65.0634509 | 0.5612332 | 0.16204232 | 3.46349761 |
| ENSG00000141736.13 | ERBB2 | 258.000723 | -0.574549516 | 0.16602413 | -3.4606386 |
| ENSG00000142657.20 | PGD | 4.3696037 | -1.824457029 | 0.52743572 | -3.4591079 |
| ENSG00000152455.15 | SUV39H2 | 31.5644571 | 0.977450802 | 0.28254784 | 3.45941702 |
| ENSG00000170017.12 | ALCAM | 18.0514367 | -1.170588353 | 0.33846972 | -3.4584728 |
| ENSG00000186166.8 | CCDC84 | 171.843253 | -0.604875553 | 0.17486002 | -3.4591987 |
| ENSG00000204866.8 | IGFL2 | 4.71900634 | 2.380820627 | 0.68854317 | 3.45776523 |
| ENSG00000269275.1 | AC020922.3 | 2.79344025 | -2.962686018 | 0.85679234 | -3.4578811 |
| ENSG00000100316.15 | RPL3 | 46.0277023 | -0.703750566 | 0.20376373 | -3.4537578 |
| ENSG00000139514.12 | SLC7A1 | 15.265848 | -1.348240123 | 0.39052321 | -3.4523944 |
| ENSG00000050438.16 | SLC4A8 | 31.8880905 | 0.866749848 | 0.25136578 | 3.44816166 |
| ENSG00000271976.1 | AC012467.2 | 4.26497402 | 2.510643142 | 0.72837268 | 3.4469211 |
| ENSG00000175137.10 | SH3BP5L | 7.96432103 | -1.435053861 | 0.41643961 | -3.4460071 |
| ENSG00000272088.1 | AL512413.1 | 5.48231568 | -1.640181101 | 0.47607226 | -3.4452356 |
| ENSG00000103855.17 | CD276 | 10.2679503 | -1.475864765 | 0.42872535 | -3.4424481 |
| ENSG00000142453.11 | CARM1 | 389.411085 | -0.502428505 | 0.14593943 | -3.4427193 |
| ENSG00000145216.15 | FIP1L1 | 50.6282715 | 0.927017224 | 0.26925928 | 3.44284229 |
| ENSG00000177106.15 | EPS8L2 | 47.1611072 | 0.642441866 | 0.18665531 | 3.44186226 |
| ENSG00000259322.1 | AC090607.1 | 70.1431952 | -0.58956983 | 0.17131298 | -3.4414778 |
| ENSG00000005075.15 | POLR2J | 12.7128232 | -0.94877271 | 0.27585003 | -3.4394512 |
| ENSG00000236782.7 | AL391650.1 | 159.581519 | -0.898515592 | 0.26135201 | -3.4379517 |
| ENSG00000137135.17 | ARHGEF39 | 321.554795 | -0.453449498 | 0.13196997 | -3.4360051 |
| ENSG00000125631.7 | HTR5BP | 69.2230718 | 0.584860084 | 0.17024655 | 3.43537113 |
| ENSG00000186814.13 | ZSCAN30 | 35.6560854 | 1.045442465 | 0.30439358 | 3.43450887 |
| ENSG00000222043.2 | AC079305.1 | 98.7735837 | 0.433558512 | 0.12627609 | 3.43341729 |

| | | | | | |
|---|---|---|---|---|---|
| ENSG00000084207.16 | GSTP1 | 5.18226531 | -1.967421062 | 0.57399351 | -3.4276016 |
| ENSG00000159917.16 | ZNF235 | 65.9388484 | 0.741900483 | 0.21638993 | 3.42853512 |
| ENSG00000279762.3 | AC005899.8 | 101.724007 | -0.603750623 | 0.17612552 | -3.4279566 |
| ENSG00000247903.1 | AC024896.1 | 83.9457778 | 0.735369683 | 0.21476746 | 3.42402748 |
| ENSG00000232878.3 | DPYD-AS1 | 90.0935878 | 0.92648136 | 0.27066589 | 3.42297051 |
| ENSG00000272918.1 | AC005070.3 | 62.2355688 | 0.841602674 | 0.24583683 | 3.42341984 |
| ENSG00000105248.15 | CCDC94 | 1.71275759 | -2.952431413 | 0.86312673 | -3.4206233 |
| ENSG00000227039.6 | ITGB2-AS1 | 19.8216278 | 1.596467263 | 0.46685635 | 3.41961137 |
| ENSG00000265558.1 | MIR3918 | 8.95173582 | -1.678632422 | 0.49101732 | -3.4186827 |
| ENSG00000250838.1 | AC091133.2 | 1.82531437 | -2.919605779 | 0.85420149 | -3.4179357 |
| ENSG00000140553.17 | UNC45A | 76.0977377 | -0.505686019 | 0.14799204 | -3.4169813 |
| ENSG00000019144.18 | PHLDB1 | 12.9914247 | -1.014540333 | 0.29714828 | -3.4142561 |
| ENSG00000180822.11 | PSMG4 | 272.823466 | 1.18909132 | 0.34814504 | 3.41550552 |
| ENSG00000260017.1 | AC138811.1 | 119.357659 | 0.582055063 | 0.17048068 | 3.41419954 |
| ENSG00000274897.2 | PANO1 | 12.8289374 | -1.252585242 | 0.36678914 | -3.4150009 |
| ENSG00000106034.17 | CPED1 | 2.88924598 | 2.51326392 | 0.73686172 | 3.41076737 |
| ENSG00000143341.11 | HMCN1 | 12.0486306 | 1.889842953 | 0.55456869 | 3.40777074 |
| ENSG00000120438.11 | TCP1 | 130.053557 | -0.938488762 | 0.27548077 | -3.4067306 |
| ENSG00000267265.5 | AC011476.3 | 17.3348422 | 0.88800275 | 0.26076587 | 3.40536419 |
| ENSG00000269888.1 | AC112491.1 | 48.4615139 | -0.826450879 | 0.24289617 | -3.4024862 |
| ENSG00000103023.11 | PRSS54 | 28.8252378 | 1.009653924 | 0.29690565 | 3.40058846 |
| ENSG00000175414.6 | ARL10 | 107.094957 | -0.559839562 | 0.16461347 | -3.4009341 |
| ENSG00000237133.1 | AC020594.1 | 6.15017019 | 1.565913225 | 0.4603701 | 3.40142252 |
| ENSG00000239388.8 | ASB14 | 122.295469 | 0.742714687 | 0.21837015 | 3.40117317 |
| ENSG00000104894.11 | CD37 | 130.905191 | -0.647212233 | 0.19046693 | -3.3980295 |
| ENSG00000237975.6 | FLG-AS1 | 34.0825002 | 1.898984181 | 0.55880956 | 3.39826716 |
| ENSG00000274425.1 | AC114271.1 | 160.791232 | -0.492807735 | 0.14503722 | -3.3978019 |
| ENSG00000012061.15 | ERCC1 | 230.382192 | -0.638960001 | 0.18842517 | -3.3910545 |
| ENSG00000227689.1 | SRP68P2 | 2.86448884 | 3.386253618 | 0.99862403 | 3.39091941 |
| ENSG00000264859.5 | DSG2-AS1 | 4.91573647 | 3.591546809 | 1.05905113 | 3.39128744 |
| ENSG00000241560.5 | ZBTB20-AS1 | 11.5367288 | 1.254018208 | 0.36999135 | 3.38931763 |
| ENSG00000157240.3 | FZD1 | 7.03251286 | -1.443236613 | 0.42608961 | -3.3871669 |
| ENSG00000260350.1 | AC012173.1 | 322.091717 | -0.443903634 | 0.13106429 | -3.3869151 |
| ENSG00000110315.6 | RNF141 | 22.0221146 | 1.50377363 | 0.4441813 | 3.3854951 |
| ENSG00000204623.9 | ZNRD1ASP | 110.289692 | -0.744857734 | 0.2199585 | -3.3863557 |
| ENSG00000255114.1 | AP003392.3 | 575.869805 | -0.444320156 | 0.13123747 | -3.3856197 |
| ENSG00000251003.8 | ZFPM2-AS1 | 1.61204839 | -3.587980934 | 1.06004676 | -3.3847384 |
| ENSG00000246859.2 | STARD4-AS1 | 2011.57718 | 1.171467332 | 0.34622137 | 3.3835789 |
| ENSG00000125434.10 | SLC25A35 | 199.827924 | -0.576072564 | 0.1703607 | -3.3814873 |
| ENSG00000240225.10 | ZNF542P | 7.10432858 | 1.176562024 | 0.34803387 | 3.38059631 |
| ENSG00000164530.14 | PI16 | 3.26696174 | -3.919788011 | 1.15967618 | -3.3800712 |
| ENSG00000065320.8 | NTN1 | 4.37757015 | 1.759823003 | 0.52086106 | 3.37868031 |
| ENSG00000275005.1 | AL354950.1 | 2.87242421 | 3.69103241 | 1.09251407 | 3.37847585 |

| | | | | | |
|---|---|---|---|---|---|
| ENSG00000188343.12 | FAM92A | 99.1537333 | 0.687198205 | 0.20354736 | 3.37610969 |
| ENSG00000263797.1 | AP005210.1 | 2.18902624 | 2.607692534 | 0.77252548 | 3.37554241 |
| ENSG00000259863.1 | SH3RF3-AS1 | 25.6187425 | 0.770345568 | 0.22835433 | 3.37346593 |
| ENSG00000229847.8 | EMX2OS | 27.711872 | 1.051463531 | 0.31180227 | 3.37221257 |
| ENSG00000164761.8 | TNFRSF11B | 7.22511942 | 1.873169468 | 0.55569235 | 3.37087501 |
| ENSG00000233308.1 | OSTN-AS1 | 1.3347601 | 3.985738647 | 1.18300134 | 3.36917509 |
| ENSG00000185013.16 | NT5C1B | 5.03213463 | 1.509268626 | 0.44808416 | 3.36827043 |
| ENSG00000258634.3 | AL160006.1 | 3.26125883 | -3.960296402 | 1.17776457 | -3.3625535 |
| ENSG00000169962.4 | TAS1R3 | 23.040814 | 0.95857293 | 0.2851463 | 3.36168808 |
| ENSG00000082146.12 | STRADB | 170.556724 | 0.685362683 | 0.20395247 | 3.36040385 |
| ENSG00000258757.1 | AL133453.1 | 273.537456 | 0.797914143 | 0.23747663 | 3.35996905 |
| ENSG00000127314.17 | RAP1B | 9.6292882 | -1.916187403 | 0.571044 | -3.3555863 |
| ENSG00000083814.13 | ZNF671 | 2.23357879 | 2.536819781 | 0.75624815 | 3.35448064 |
| ENSG00000103257.8 | SLC7A5 | 25.828853 | -1.524457578 | 0.45448014 | -3.3542887 |
| ENSG00000146374.13 | RSPO3 | 1.86209572 | 3.174505547 | 0.94667635 | 3.35331661 |
| ENSG00000234925.2 | ATP5PDP4 | 5.65483379 | -1.510577832 | 0.45096202 | -3.3496786 |
| ENSG00000277959.1 | AL162274.2 | 5.08355299 | 1.770263782 | 0.52861412 | 3.34887724 |
| ENSG00000104960.15 | PTOV1 | 137.56335 | -1.064722462 | 0.31815346 | -3.3465689 |
| ENSG00000161179.13 | YDJC | 7.28866504 | -1.506347437 | 0.45020253 | -3.3459329 |
| ENSG00000168350.7 | DEGS2 | 105.215863 | 0.769351363 | 0.22996693 | 3.34548694 |
| ENSG00000185869.14 | ZNF829 | 3.38585739 | 2.130987819 | 0.63701845 | 3.34525293 |
| ENSG00000204620.3 | AC115618.1 | 216.136588 | -0.757782185 | 0.22657345 | -3.3445321 |
| ENSG00000240211.1 | AC092849.1 | 7.79291246 | -1.844733305 | 0.55182406 | -3.3429737 |
| ENSG00000258843.1 | AL133485.1 | 44.3809263 | 0.695040978 | 0.2078938 | 3.34325014 |
| ENSG00000203722.7 | RAET1G | 5.37167166 | 2.249796862 | 0.67326036 | 3.34164462 |
| ENSG00000085998.13 | POMGNT1 | 38.360807 | 0.745057224 | 0.22306752 | 3.34005253 |
| ENSG00000261093.1 | AC141586.3 | 413.432696 | 0.323095209 | 0.096782 | 3.33838103 |
| ENSG00000182093.15 | WRB | 3.46939829 | 1.671726187 | 0.50091892 | 3.33731891 |
| ENSG00000175470.19 | PPP2R2D | 4.00085666 | 1.894324715 | 0.56779439 | 3.33628645 |
| ENSG00000115844.10 | DLX2 | 1.90042713 | -2.828918453 | 0.84836224 | -3.3345643 |
| ENSG00000087302.8 | RTRAF | 354.170885 | -1.034495663 | 0.31040324 | -3.3327476 |
| ENSG00000280515.1 | SALRNA2 | 1.94363924 | 2.682591019 | 0.80584671 | 3.3289098 |
| ENSG00000230140.5 | AC016738.2 | 38.7002049 | 0.818968643 | 0.24615397 | 3.32705849 |
| ENSG00000234745.10 | HLA-B | 13.8546822 | -1.219040044 | 0.36642945 | -3.326807 |
| ENSG00000285190.1 | AC018754.1 | 65.5479167 | -0.611366358 | 0.18373324 | -3.3274674 |
| ENSG00000227543.4 | SPAG5-AS1 | 1190.37672 | 0.430668236 | 0.12948887 | 3.3259092 |
| ENSG00000182718.16 | ANXA2 | 63.1695022 | -0.605089626 | 0.18218904 | -3.3212186 |
| ENSG00000173065.13 | FAM222B | 63.5829674 | -0.493494114 | 0.14872873 | -3.318082 |
| ENSG00000261646.1 | AC093849.1 | 1.76474621 | 3.330122231 | 1.0036879 | 3.31788621 |
| ENSG00000279488.1 | AC004623.1 | 109.382463 | -0.381759613 | 0.11509218 | -3.3169899 |
| ENSG00000142552.7 | RCN3 | 51.6435855 | -0.904869447 | 0.2728553 | -3.3162979 |
| ENSG00000162244.11 | RPL29 | 20.9175705 | -0.967042647 | 0.29197038 | -3.3121259 |
| ENSG00000184162.14 | NR2C2AP | 66.5519346 | -0.601549873 | 0.18169475 | -3.3107719 |

| | | | | | |
|---|---|---|---|---|---|
| ENSG00000277290.1 | AC136475.10 | 8.32789475 | -1.689924816 | 0.51050662 | -3.3102897 |
| ENSG00000274523.4 | RCC1L | 91.9874786 | 0.677402184 | 0.20476282 | 3.30822842 |
| ENSG00000257831.1 | AL136418.1 | 140.335644 | 0.584788276 | 0.17717498 | 3.30062564 |
| ENSG00000265840.1 | AC010761.5 | 67.792809 | -0.525435619 | 0.15918556 | -3.3007744 |
| ENSG00000236276.1 | NDP-AS1 | 4.43335581 | 3.015462964 | 0.91449634 | 3.297403 |
| ENSG00000100097.11 | LGALS1 | 40.2682994 | -0.971506475 | 0.29466828 | -3.2969496 |
| ENSG00000135439.11 | AGAP2 | 424.450822 | -0.384510535 | 0.11670624 | -3.294687 |
| ENSG00000167984.17 | NLRC3 | 21.0908323 | 0.69116517 | 0.20974183 | 3.29531391 |
| ENSG00000256092.2 | AC137767.1 | 24.1147924 | 0.800617942 | 0.24300147 | 3.29470405 |
| ENSG00000114790.12 | ARHGEF26 | 1.90051573 | 2.377927658 | 0.72227501 | 3.29227459 |
| ENSG00000223923.1 | AC010136.1 | 95.1007527 | 0.641318354 | 0.19492735 | 3.29003777 |
| ENSG00000267192.1 | AC006116.4 | 2.25354724 | 2.499959831 | 0.75982888 | 3.29016163 |
| ENSG00000042317.16 | SPATA7 | 117.796642 | 0.703422633 | 0.21409827 | 3.28551294 |
| ENSG00000154096.13 | THY1 | 36.1809121 | 0.857064487 | 0.26106853 | 3.28291001 |
| ENSG00000142541.16 | RPL13A | 43.3310679 | -0.771124361 | 0.23525848 | -3.277775 |
| ENSG00000198113.2 | TOR4A | 1.68692942 | -2.939875888 | 0.89739142 | -3.2760241 |
| ENSG00000122861.15 | PLAU | 4.33122977 | 2.425924558 | 0.74103989 | 3.27367609 |
| ENSG00000076344.15 | RGS11 | 2.69943677 | 1.878801055 | 0.57414508 | 3.27234546 |
| ENSG00000225420.1 | AC104134.1 | 194.146488 | 0.626102855 | 0.19146793 | 3.27001422 |
| ENSG00000079308.18 | TNS1 | 27.9101328 | -0.918107502 | 0.28114975 | -3.2655462 |
| ENSG00000161016.17 | RPL8 | 28.8595601 | -1.001957225 | 0.30685632 | -3.2652324 |
| ENSG00000167721.10 | TSR1 | 197.521326 | 0.441751741 | 0.1353019 | 3.26493372 |
| ENSG00000160957.12 | RECQL4 | 27.8593163 | -0.878075376 | 0.26905371 | -3.2635691 |
| ENSG00000269176.2 | AP001160.3 | 52.1563196 | -0.68212271 | 0.20899241 | -3.2638636 |
| ENSG00000232085.1 | AL606534.3 | 172.06009 | 0.631405319 | 0.19353329 | 3.26251524 |
| ENSG00000157542.10 | KCNJ6 | 2.57639152 | 3.412036901 | 1.04628432 | 3.26109914 |
| ENSG00000124766.6 | SOX4 | 5.12335283 | -1.637202628 | 0.50255774 | -3.2577403 |
| ENSG00000256690.1 | AP001160.1 | 89.672377 | -0.436641339 | 0.13401711 | -3.2581015 |
| ENSG00000259661.1 | AC068831.4 | 246.516948 | 0.666245151 | 0.20451814 | 3.2576335 |
| ENSG00000266341.1 | AC004477.2 | 1513.1776 | 0.566801028 | 0.17402168 | 3.25707131 |
| ENSG00000253125.1 | AC055854.1 | 11.5015179 | -2.033681956 | 0.62453876 | -3.2562942 |
| ENSG00000117385.15 | P3H1 | 7.84037786 | -1.219350192 | 0.37491489 | -3.2523387 |
| ENSG00000238018.2 | AC093110.1 | 792.185809 | 0.605014849 | 0.18602314 | 3.25236332 |
| ENSG00000106246.17 | PTCD1 | 498.063981 | -0.365611165 | 0.11268206 | -3.2446262 |
| ENSG00000085788.13 | DDHD2 | 68.4948077 | 0.729554813 | 0.22505527 | 3.2416696 |
| ENSG00000107957.16 | SH3PXD2A | 41.976063 | -0.834835487 | 0.25763845 | -3.2403374 |
| ENSG00000133740.10 | E2F5 | 200.24951 | -0.497094277 | 0.15354863 | -3.2373735 |
| ENSG00000163082.9 | SGPP2 | 8.65055024 | -1.506845916 | 0.46583128 | -3.2347461 |
| ENSG00000170113.15 | NIPA1 | 4.18218038 | -2.182304501 | 0.67454727 | -3.2352136 |
| ENSG00000257494.1 | AC004217.1 | 20.7980865 | 0.67297892 | 0.20796053 | 3.23608962 |
| ENSG00000258428.5 | AL161757.2 | 49.0187104 | 0.669975976 | 0.20708129 | 3.23532841 |
| ENSG00000267787.6 | AC027097.2 | 241.88793 | 0.975987197 | 0.30169635 | 3.23499837 |
| ENSG00000148824.18 | MTG1 | 19.1793454 | 1.066443587 | 0.3297433 | 3.23416302 |

| | | | | | |
|---|---|---|---|---|---|
| ENSG00000149716.12 | ORAOV1 | 8956.85788 | 0.744149089 | 0.23014113 | 3.23344669 |
| ENSG00000228242.6 | AC093495.1 | 363.049234 | 0.502635286 | 0.15546864 | 3.23303325 |
| ENSG00000066855.15 | MTFR1 | 37.6675883 | 0.92762637 | 0.28696393 | 3.2325539 |
| ENSG00000151806.13 | GUF1 | 15.849271 | 1.060102376 | 0.32814923 | 3.23054961 |
| ENSG00000107223.12 | EDF1 | 23.7588255 | -0.908036418 | 0.28114786 | -3.2297468 |
| ENSG00000229358.3 | DPY19L1P1 | 64.7559017 | 0.529655721 | 0.16399527 | 3.22970115 |
| ENSG00000280721.1 | LINC01943 | 346.897202 | -0.537469831 | 0.1664679 | -3.2286696 |
| ENSG00000149260.16 | CAPN5 | 2.85172712 | -2.470449257 | 0.76549115 | -3.2272735 |
| ENSG00000104131.12 | EIF3J | 51.783071 | 0.666112016 | 0.20651061 | 3.22555829 |
| ENSG00000100678.18 | SLC8A3 | 1.94013511 | 4.478760197 | 1.38969638 | 3.2228336 |
| ENSG00000121988.17 | ZRANB3 | 190.433979 | 0.438483008 | 0.13603809 | 3.22323711 |
| ENSG00000259351.1 | AC015914.1 | 33.2277883 | 0.891921305 | 0.27663985 | 3.22412439 |
| ENSG00000274024.1 | AL590282.1 | 4.66265923 | 1.72027567 | 0.53379164 | 3.22274751 |
| ENSG00000285278.1 | AL138885.3 | 61.3588282 | -0.9686626 | 0.30048665 | -3.223646 |
| ENSG00000169764.15 | UGP2 | 10.2430063 | 1.045100702 | 0.32494203 | 3.21626813 |
| ENSG00000170365.9 | SMAD1 | 3.11190243 | -2.448181841 | 0.76131568 | -3.215725 |
| ENSG00000058729.10 | RIOK2 | 1.59881952 | 2.638717839 | 0.82120803 | 3.21321486 |
| ENSG00000114021.11 | NIT2 | 30.5398463 | 0.723430822 | 0.22510092 | 3.21380662 |
| ENSG00000176383.8 | B3GNT4 | 377.252612 | -0.384498531 | 0.11966149 | -3.2132188 |
| ENSG00000237489.4 | C10orf143 | 4.44999042 | 1.9642432 | 0.61126847 | 3.2133887 |
| ENSG00000263069.5 | AC124319.2 | 186.304378 | 0.666033981 | 0.20741275 | 3.21115259 |
| ENSG00000233360.4 | Z83844.2 | 33.9857701 | -0.803978228 | 0.25045291 | -3.2100974 |
| ENSG00000187185.4 | AC092118.1 | 7.91652224 | -1.342334405 | 0.41827396 | -3.2092229 |
| ENSG00000251196.1 | AC106760.1 | 45.1477006 | 1.198417259 | 0.37364739 | 3.20734818 |
| ENSG00000273387.1 | AC005005.3 | 264.571346 | 0.419347092 | 0.13082446 | 3.20541811 |
| ENSG00000130758.7 | MAP3K10 | 7.84675841 | 1.357299526 | 0.42364619 | 3.2038516 |
| ENSG00000145632.14 | PLK2 | 1.490934 | -2.810478008 | 0.8772922 | -3.2035826 |
| ENSG00000279344.1 | AC007342.6 | 289.709238 | 0.440711513 | 0.13756915 | 3.20356361 |
| ENSG00000105372.7 | RPS19 | 31.136532 | -0.945747355 | 0.29527286 | -3.2029607 |
| ENSG00000137124.7 | ALDH1B1 | 2.24552213 | -2.306809419 | 0.72031027 | -3.2025219 |
| ENSG00000179889.18 | PDXDC1 | 467.322131 | 0.440563451 | 0.13758539 | 3.20210927 |
| ENSG00000140992.18 | PDPK1 | 15.1513025 | -1.474634391 | 0.46077194 | -3.2003563 |
| ENSG00000273621.1 | RF02271 | 1.55328515 | -3.930275282 | 1.22845987 | -3.1993518 |
| ENSG00000244151.1 | AC010973.2 | 705.141607 | -0.37565219 | 0.11745911 | -3.1981529 |
| ENSG00000178297.13 | TMPRSS9 | 74.008542 | -0.646676428 | 0.20225943 | -3.1972622 |
| ENSG00000262791.1 | AC130343.1 | 83.5822118 | -0.480567007 | 0.15030293 | -3.197323 |
| ENSG00000070404.9 | FSTL3 | 6.97448961 | -1.214840992 | 0.38020669 | -3.195212 |
| ENSG00000115317.11 | HTRA2 | 337.566396 | -0.80074881 | 0.25105467 | -3.1895396 |
| ENSG00000165916.8 | PSMC3 | 8.83749257 | -1.116184503 | 0.35002823 | -3.1888414 |
| ENSG00000164106.7 | SCRG1 | 2.58695218 | 2.136386782 | 0.67016848 | 3.18783536 |
| ENSG00000123342.15 | MMP19 | 131.90413 | 0.788060993 | 0.24728094 | 3.18690555 |
| ENSG00000163806.15 | SPDYA | 34.9825899 | -0.615736821 | 0.19330328 | -3.1853409 |
| ENSG00000244104.3 | RN7SL659P | 1.89441654 | 3.270383242 | 1.02696418 | 3.18451538 |

| | | | | | |
|---|---|---|---|---|---|
| ENSG00000260465.1 | AC018557.1 | 928.262277 | 0.535285481 | 0.16817407 | 3.18292512 |
| ENSG00000282556.2 | AC068733.3 | 1296.99763 | -0.572039641 | 0.17975094 | -3.1824015 |
| ENSG00000285244.1 | DINOL | 2.36451288 | 2.357322251 | 0.7408463 | 3.1819316 |
| ENSG00000124588.19 | NQO2 | 293.540804 | 1.173822315 | 0.36907281 | 3.1804627 |
| ENSG00000231104.8 | AC022395.1 | 121.004639 | 0.831318813 | 0.26160948 | 3.17770901 |
| ENSG00000165124.17 | SVEP1 | 13.3605094 | 0.900823692 | 0.28358382 | 3.17656944 |
| ENSG00000187634.11 | SAMD11 | 233.891948 | -0.544708627 | 0.17149279 | -3.1762772 |
| ENSG00000058668.14 | ATP2B4 | 37.8527516 | 0.808648015 | 0.25519616 | 3.16873109 |
| ENSG00000168028.13 | RPSA | 3.61087597 | -1.737014221 | 0.54804522 | -3.1694725 |
| ENSG00000215717.5 | TMEM167B | 27.9824743 | 0.662406203 | 0.20902986 | 3.16895491 |
| ENSG00000250794.2 | ALG1L12P | 2.31747682 | 3.177024391 | 1.00361848 | 3.16556985 |
| ENSG00000006015.17 | REX1BD | 62.0669019 | -3.585896777 | 1.13344438 | -3.1637166 |
| ENSG00000154928.17 | EPHB1 | 15.1041406 | 1.555707431 | 0.4919431 | 3.16237273 |
| ENSG00000051523.10 | CYBA | 14.9549832 | -1.409508827 | 0.44582273 | -3.1615903 |
| ENSG00000183287.14 | CCBE1 | 10.0308252 | 1.606194935 | 0.50803533 | 3.1615812 |
| ENSG00000142937.11 | RPS8 | 43.9304658 | -1.138521473 | 0.36018366 | -3.160947 |
| ENSG00000230155.6 | FO393401.1 | 40.0689342 | 0.67365639 | 0.21322299 | 3.15939846 |
| ENSG00000236886.2 | AC007563.2 | 339.270569 | -1.884320981 | 0.59659408 | -3.1584641 |
| ENSG00000169926.10 | KLF13 | 4.12885238 | -1.788133834 | 0.56633093 | -3.157401 |
| ENSG00000271064.1 | AC027644.2 | 11.3297725 | 0.925076379 | 0.29312594 | 3.15590074 |
| ENSG00000167264.17 | DUS2 | 12.3152013 | -0.926596975 | 0.29373314 | -3.1545537 |
| ENSG00000267257.1 | AC105105.1 | 2488.28795 | 0.676312586 | 0.21434508 | 3.15525132 |
| ENSG00000273149.1 | AL138963.3 | 11097.3912 | -0.639124925 | 0.20258856 | -3.1547928 |
| ENSG00000260007.3 | AC107871.1 | 55.555381 | 0.686921165 | 0.21814069 | 3.14898221 |
| ENSG00000112578.9 | BYSL | 2.03058764 | -2.577065216 | 0.81897243 | -3.1467057 |
| ENSG00000266969.1 | AP002449.1 | 48.8476583 | -0.649168276 | 0.20644699 | -3.1444793 |
| ENSG00000272842.1 | AL391834.1 | 36.9202925 | 0.92233851 | 0.29345199 | 3.14306437 |
| ENSG00000182541.17 | LIMK2 | 15.9997893 | 0.988392863 | 0.31464967 | 3.14124866 |
| ENSG00000116171.17 | SCP2 | 14.6709111 | 0.985867604 | 0.31390144 | 3.14069159 |
| ENSG00000121940.15 | CLCC1 | 120.389098 | 1.261255404 | 0.40259169 | 3.13284013 |
| ENSG00000168769.13 | TET2 | 2.96187063 | 1.835701332 | 0.58597463 | 3.13273174 |
| ENSG00000255203.1 | OR7E2P | 2.00895491 | 2.80326996 | 0.89508537 | 3.13184647 |
| ENSG00000259954.1 | IL21R-AS1 | 47.8869687 | -1.311599819 | 0.41886835 | -3.1312937 |
| ENSG00000227766.1 | HCG4P5 | 2002.20722 | -0.662573466 | 0.21181639 | -3.1280557 |
| ENSG00000264486.1 | AC061975.4 | 2.22071883 | 2.331184796 | 0.74550717 | 3.12697836 |
| ENSG00000240889.1 | NDUFB2-AS1 | 245.977405 | -0.654258944 | 0.20928158 | -3.1262137 |
| ENSG00000234149.1 | AC018511.2 | 49.4430951 | 0.806473907 | 0.25801571 | 3.12567758 |
| ENSG00000267112.1 | AC098848.1 | 10.0987492 | -2.490140838 | 0.79718508 | -3.1236671 |
| ENSG00000267576.1 | AC011472.3 | 53.312196 | 0.992481686 | 0.3177238 | 3.12372474 |
| ENSG00000236358.1 | AL355472.3 | 5.25650505 | -1.541948972 | 0.49396202 | -3.1215942 |
| ENSG00000235481.2 | UBE2R2-AS1 | 9.69369357 | -1.567539114 | 0.50236371 | -3.1203272 |
| ENSG00000223443.2 | USP17L2 | 1.4600888 | 3.282549242 | 1.05263443 | 3.11841333 |
| ENSG00000198198.16 | SZT2 | 223.78361 | -0.492981567 | 0.1581462 | -3.117252 |

| | | | | | |
|---|---|---|---|---|---|
| ENSG00000101624.10 | CEP76 | 3.28388266 | 1.875783408 | 0.60190973 | 3.11638657 |
| ENSG00000106785.14 | TRIM14 | 336.476227 | -0.496836357 | 0.15942673 | -3.116393 |
| ENSG00000210195.2 | MT-TT | 6.59777835 | -1.493612437 | 0.47937977 | -3.1157185 |
| ENSG00000075624.14 | ACTB | 382.714834 | -0.591975132 | 0.19015514 | -3.1131167 |
| ENSG00000205542.10 | TMSB4X | 74.0406953 | -0.683297371 | 0.21960292 | -3.1115132 |
| ENSG00000116194.12 | ANGPTL1 | 2.64009991 | 2.364780486 | 0.76027522 | 3.11042688 |
| ENSG00000129625.12 | REEP5 | 21.3295268 | 0.696670739 | 0.22404294 | 3.10954109 |
| ENSG00000267040.6 | AC027097.1 | 261.57708 | 1.009509771 | 0.32486463 | 3.10747824 |
| ENSG00000180448.10 | ARHGAP45 | 118.797683 | 0.723873123 | 0.23298857 | 3.10690406 |
| ENSG00000125148.6 | MT2A | 5.71294438 | -2.057870082 | 0.66245379 | -3.1064357 |
| ENSG00000137962.12 | ARHGAP29 | 1.2735537 | 2.980441821 | 0.95989741 | 3.10495871 |
| ENSG00000196569.12 | LAMA2 | 17.1676764 | 1.075788226 | 0.34642432 | 3.10540617 |
| ENSG00000210112.1 | MT-TM | 10.0117139 | -1.47682103 | 0.47565152 | -3.1048382 |
| ENSG00000085511.19 | MAP3K4 | 22.1039164 | -1.604562778 | 0.51689592 | -3.104228 |
| ENSG00000183765.21 | CHEK2 | 22.8774092 | -0.899835554 | 0.29015356 | -3.1012391 |
| ENSG00000239665.8 | AL157392.3 | 181.354748 | -0.972607389 | 0.31360306 | -3.1013964 |
| ENSG00000245311.2 | ARNTL2-AS1 | 56.2749594 | 0.9765148 | 0.3149192 | 3.10084238 |
| ENSG00000269843.1 | AC008537.2 | 29.8530485 | 0.752239244 | 0.2425756 | 3.10105069 |
| ENSG00000130021.13 | PUDP | 26.0039653 | 1.021295992 | 0.32946917 | 3.09982267 |
| ENSG00000123700.4 | KCNJ2 | 1.88217371 | 2.317071392 | 0.74786439 | 3.09825072 |
| ENSG00000136699.19 | SMPD4 | 658.736852 | -0.655769909 | 0.21166082 | -3.0982111 |
| ENSG00000164404.8 | GDF9 | 312.518957 | -0.818071 | 0.26400165 | -3.0987344 |
| ENSG00000156453.13 | PCDH1 | 1.82108138 | 3.56598887 | 1.1518865 | 3.09578146 |
| ENSG00000258177.1 | AC008149.1 | 377.377158 | 0.715474368 | 0.23114913 | 3.09529335 |
| ENSG00000125459.15 | MSTO1 | 4.02661513 | 1.638901585 | 0.52963818 | 3.09437962 |
| ENSG00000283073.1 | SMUG1-AS1 | 1.74418595 | 2.502275408 | 0.80886845 | 3.09355051 |
| ENSG00000173141.4 | MRPL57 | 1.68008847 | -2.611916423 | 0.84444328 | -3.0930632 |
| ENSG00000255112.2 | CHMP1B | 4.60467465 | -1.960490109 | 0.63422089 | -3.0911787 |
| ENSG00000176915.14 | ANKLE2 | 5.0125171 | -1.370328393 | 0.44363433 | -3.0888691 |
| ENSG00000249996.1 | AC106786.2 | 109.294555 | -0.711286796 | 0.23032087 | -3.0882429 |
| ENSG00000231607.10 | DLEU2 | 23.5418165 | -0.742412403 | 0.24060623 | -3.085591 |
| ENSG00000248801.6 | C8orf34-AS1 | 1.34338215 | 2.806276578 | 0.90937916 | 3.08592577 |
| ENSG00000266002.1 | AC091059.1 | 15.3127949 | 1.242504684 | 0.40260708 | 3.08614711 |
| ENSG00000136235.16 | GPNMB | 6.16956132 | -1.435560872 | 0.46558571 | -3.0833439 |
| ENSG00000257702.3 | LBX2-AS1 | 34.5966182 | -0.775965395 | 0.2516485 | -3.0835287 |
| ENSG00000267383.6 | AC011447.3 | 16.9801965 | -0.821126507 | 0.26633276 | -3.0830849 |
| ENSG00000157227.12 | MMP14 | 132.431655 | -0.905898093 | 0.29389742 | -3.0823615 |
| ENSG00000162931.11 | TRIM17 | 15.3299041 | 0.760378197 | 0.24678592 | 3.0811247 |
| ENSG00000188312.13 | CENPP | 208.067517 | -0.380610416 | 0.12352064 | -3.0813507 |
| ENSG00000210144.1 | MT-TY | 666.983249 | 0.501842819 | 0.16292048 | 3.08029305 |
| ENSG00000100426.6 | ZBED4 | 14.1250409 | 1.050194624 | 0.34104976 | 3.07930029 |
| ENSG00000110237.4 | ARHGEF17 | 22.6853964 | -0.921326317 | 0.29919714 | -3.0793286 |
| ENSG00000112096.17 | SOD2 | 39.5239983 | -0.690821969 | 0.22438101 | -3.0787899 |

| | | | | | |
|---|---|---|---|---|---|
| ENSG00000159761.14 | C16orf86 | 11.2671019 | -1.006485545 | 0.32689416 | -3.078934 |
| ENSG00000198695.2 | MT-ND6 | 20828.1155 | 0.627793783 | 0.20398593 | 3.07763283 |
| ENSG00000166441.12 | RPL27A | 75.7657763 | -0.557685247 | 0.18148968 | -3.0728207 |
| ENSG00000260176.1 | AC141586.2 | 6.66627995 | 1.267116411 | 0.41240985 | 3.07246881 |
| ENSG00000225721.5 | AL592166.1 | 874.531295 | -0.770591779 | 0.25088596 | -3.0714823 |
| ENSG00000213839.4 | TMX2P1 | 122.047758 | 0.456083638 | 0.14851708 | 3.07091707 |
| ENSG00000181409.12 | AATK | 3.67227747 | -2.567497711 | 0.83642329 | -3.0696153 |
| ENSG00000115459.17 | ELMOD3 | 91.128606 | 0.444416419 | 0.14485732 | 3.06795959 |
| ENSG00000126217.20 | MCF2L | 1.44953212 | 3.722662521 | 1.21320993 | 3.06844053 |
| ENSG00000229619.3 | MBNL1-AS1 | 150.09955 | 0.648685448 | 0.21141744 | 3.06826833 |
| ENSG00000227644.2 | HIGD1AP11 | 16.696343 | -1.137601756 | 0.3709808 | -3.0664707 |
| ENSG00000183431.11 | SF3A3 | 2.84411508 | -2.19713183 | 0.7167781 | -3.0652887 |
| ENSG00000244493.1 | SLC9A9-AS2 | 11.2921693 | 1.302164731 | 0.42494252 | 3.06433147 |
| ENSG00000271889.1 | AC016747.2 | 2.59238781 | 1.825256578 | 0.59559474 | 3.06459485 |
| ENSG00000245213.6 | AC105285.1 | 29.0991238 | 0.819045598 | 0.26732497 | 3.06385744 |
| ENSG00000172349.17 | IL16 | 216.524153 | 1.130807406 | 0.36937935 | 3.06137146 |
| ENSG00000134013.15 | LOXL2 | 49.0753274 | -0.744675104 | 0.24355824 | -3.0574827 |
| ENSG00000153885.14 | KCTD15 | 3.25252076 | -1.763050485 | 0.57659522 | -3.0576918 |
| ENSG00000188869.12 | TMC3 | 13.8677289 | 1.097699353 | 0.35904315 | 3.05729092 |
| ENSG00000257042.1 | AC008011.2 | 63.361936 | -2.670077361 | 0.87387427 | -3.055448 |
| ENSG00000215866.7 | LINC01356 | 2.92831247 | -1.846134378 | 0.60430794 | -3.0549563 |
| ENSG00000229950.1 | TFAP2A-AS1 | 8.59200275 | -1.191299237 | 0.39005031 | -3.0542194 |
| ENSG00000108592.16 | FTSJ3 | 528.02067 | -0.267627127 | 0.08765541 | -3.0531728 |
| ENSG00000170881.4 | RNF139 | 2.27593215 | -2.647803438 | 0.86746232 | -3.0523556 |
| ENSG00000124942.13 | AHNAK | 151.29507 | 0.576017879 | 0.18878457 | 3.05119158 |
| ENSG00000149927.17 | DOC2A | 200.195143 | -0.35442357 | 0.11616619 | -3.0510044 |
| ENSG00000109339.21 | MAPK10 | 377.270124 | 0.429463385 | 0.14091494 | 3.04767817 |
| ENSG00000253309.6 | SERPINE3 | 35.0175508 | 0.667806883 | 0.21915179 | 3.04723446 |
| ENSG00000272114.1 | AL136131.3 | 169.750917 | 1.029767728 | 0.33795631 | 3.0470439 |
| ENSG00000285545.1 | | 20.8568203 | 0.769402927 | 0.25245621 | 3.04766883 |
| ENSG00000159267.14 | HLCS | 3.39122891 | -1.884923081 | 0.61886527 | -3.0457729 |
| ENSG00000285932.1 | | 39.8964816 | -0.876929985 | 0.28801067 | -3.0447829 |
| ENSG00000173545.4 | ZNF622 | 2.96618797 | -1.775164115 | 0.583356 | -3.0430202 |
| ENSG00000164199.17 | ADGRV1 | 25.3852713 | 0.628995505 | 0.20694438 | 3.03944237 |
| ENSG00000164107.8 | HAND2 | 23.992361 | 1.212200226 | 0.39901123 | 3.03801031 |
| ENSG00000264044.1 | AC005726.2 | 1690.02125 | 0.502629198 | 0.16543896 | 3.03815494 |
| ENSG00000104529.17 | EEF1D | 75.1550253 | -0.527322285 | 0.17360755 | -3.0374386 |
| ENSG00000198755.10 | RPL10A | 11.042606 | -1.269495264 | 0.41827932 | -3.0350419 |
| ENSG00000013293.5 | SLC7A14 | 1.56304174 | 2.897603871 | 0.95555983 | 3.03236258 |
| ENSG00000269559.2 | AC093677.2 | 10.4272981 | -1.07963717 | 0.3560844 | -3.0319699 |
| ENSG00000140199.11 | SLC12A6 | 217.375084 | -0.527627971 | 0.17404787 | -3.0315107 |
| ENSG00000227440.1 | ATP5MC1P4 | 6.43034188 | 2.103711038 | 0.6941011 | 3.03084237 |
| ENSG00000210140.1 | MT-TC | 12.336802 | -1.273328526 | 0.42097201 | -3.0247344 |

| | | | | | |
|---|---|---|---|---|---|
| ENSG00000164597.13 | COG5 | 480.323658 | 0.351551962 | 0.11643448 | 3.01931153 |
| ENSG00000168569.7 | TMEM223 | 306.10028 | -0.483276305 | 0.1600437 | -3.0196521 |
| ENSG00000162971.10 | TYW5 | 11.3144163 | 0.94700852 | 0.31391851 | 3.01673358 |
| ENSG00000129235.10 | TXNDC17 | 65.2469691 | -0.539154919 | 0.17892121 | -3.013365 |
| ENSG00000248710.1 | AC079594.2 | 9.01265334 | -2.57842305 | 0.856117 | -3.0117648 |
| ENSG00000214262.4 | ANKRD36BP1 | 201.176914 | 0.563604923 | 0.18720325 | 3.01065786 |
| ENSG00000243305.1 | AC026347.1 | 26.8833089 | 0.729024585 | 0.24220077 | 3.01000109 |
| ENSG00000188760.10 | TMEM198 | 41.5917002 | -0.953279686 | 0.31674796 | -3.0095843 |
| ENSG00000226877.8 | AL354733.1 | 6.48314074 | 1.299240637 | 0.43187003 | 3.00840658 |
| ENSG00000235106.9 | BRD3OS | 130.209984 | 0.627740549 | 0.2087063 | 3.00776997 |
| ENSG00000108821.13 | COL1A1 | 1996.75387 | -0.819636251 | 0.27275051 | -3.0050769 |
| ENSG00000248923.1 | MTND5P11 | 42.4534748 | -0.637954461 | 0.21227892 | -3.0052653 |
| ENSG00000155666.11 | KDM8 | 10.6433926 | 0.9512791 | 0.31669164 | 3.00380236 |
| ENSG00000231500.6 | RPS18 | 20.6583058 | -0.792364988 | 0.26379032 | -3.0037682 |
| ENSG00000172757.12 | CFL1 | 100.580457 | -0.434109823 | 0.14455198 | -3.0031398 |
| ENSG00000271806.1 | AL590822.2 | 3.58653255 | -2.009582292 | 0.6694634 | -3.0017807 |
| ENSG00000072071.16 | ADGRL1 | 62.9647381 | 0.471045147 | 0.15696708 | 3.0009168 |
| ENSG00000233822.4 | HIST1H2BN | 5.54699066 | 1.851505447 | 0.61721073 | 2.99979465 |
| ENSG00000075073.14 | TACR2 | 12.0213526 | 0.966755744 | 0.32236881 | 2.99891213 |
| ENSG00000225969.2 | ABHD11-AS1 | 9.68154915 | -1.024322116 | 0.34161961 | -2.9984289 |
| ENSG00000272211.1 | AC114760.2 | 5.66117382 | 1.175579214 | 0.39202555 | 2.99873108 |
| ENSG00000009790.14 | TRAF3IP3 | 37.6534864 | -1.126734957 | 0.3758725 | -2.997652 |
| ENSG00000080947.14 | CROCCP3 | 8.01088541 | 0.988878801 | 0.32999971 | 2.99660508 |
| ENSG00000263683.1 | AC005154.4 | 6.68020697 | 1.208295861 | 0.40321356 | 2.99666476 |
| ENSG00000003509.15 | NDUFAF7 | 188.936142 | 0.553727941 | 0.18484615 | 2.99561527 |
| ENSG00000105193.8 | RPS16 | 9.10927571 | -1.280297797 | 0.42774525 | -2.9931315 |
| ENSG00000178980.14 | SELENOW | 3.31703763 | -1.536606541 | 0.51334146 | -2.993342 |
| ENSG00000265962.1 | GACAT2 | 1.71434066 | 2.796462178 | 0.9344066 | 2.99276801 |
| ENSG00000180104.15 | EXOC3 | 99.4655585 | -0.486481263 | 0.1628454 | -2.9873811 |
| ENSG00000176222.8 | ZNF404 | 8.17286222 | 1.207983626 | 0.40452813 | 2.98615482 |
| ENSG00000168061.14 | SAC3D1 | 2.22008782 | -2.191296967 | 0.73392906 | -2.9857068 |
| ENSG00000148835.10 | TAF5 | 59.0162432 | -0.472576549 | 0.15835051 | -2.9843701 |
| ENSG00000214253.8 | FIS1 | 6.52525877 | -1.34041863 | 0.44934045 | -2.9830803 |
| ENSG00000259652.1 | AC090181.1 | 20.8464936 | 0.756139268 | 0.25349832 | 2.98281771 |
| ENSG00000151665.12 | PIGF | 1427.16806 | 0.653896406 | 0.21932434 | 2.98141286 |
| ENSG00000148848.14 | ADAM12 | 10.6193733 | -1.173920556 | 0.39414224 | -2.9784185 |
| ENSG00000204954.9 | C12orf73 | 58.824923 | 1.579825938 | 0.53036663 | 2.97874312 |
| ENSG00000147140.15 | NONO | 19.2603551 | -0.64322926 | 0.21611378 | -2.9763454 |
| ENSG00000150455.13 | TIRAP | 3.34393475 | -1.73732297 | 0.58372616 | -2.9762637 |
| ENSG00000189060.5 | H1F0 | 40.7666735 | -0.794961202 | 0.26706193 | -2.9766923 |
| ENSG00000089693.10 | MLF2 | 17.0494684 | -0.80766032 | 0.27158818 | -2.9738419 |
| ENSG00000103148.15 | NPRL3 | 413.152148 | -0.569131057 | 0.19141583 | -2.9732706 |
| ENSG00000138074.14 | SLC5A6 | 264.287927 | -0.604296347 | 0.20326475 | -2.972952 |

| | | | | | |
|---|---|---|---|---|---|
| ENSG00000165280.15 | VCP | 13.5498092 | -0.799206795 | 0.26873951 | -2.9739089 |
| ENSG00000175115.11 | PACS1 | 12.123188 | -1.005544608 | 0.33818347 | -2.9733701 |
| ENSG00000259187.1 | AC122108.1 | 7.11104591 | 2.489941306 | 0.83744201 | 2.97327012 |
| ENSG00000280054.1 | AC004241.5 | 11.2241004 | 1.541294309 | 0.51809702 | 2.97491442 |
| ENSG00000105373.18 | NOP53 | 15.0871029 | -1.117458014 | 0.37618669 | -2.9704879 |
| ENSG00000233766.7 | AC098617.1 | 61.6385157 | -2.268038453 | 0.76382917 | -2.9693007 |
| ENSG00000111665.11 | CDCA3 | 30.0424563 | -0.926223306 | 0.31199587 | -2.9687037 |
| ENSG00000111737.11 | RAB35 | 26.3672692 | -0.676999704 | 0.22813006 | -2.9676041 |
| ENSG00000138069.17 | RAB1A | 13.9785724 | 0.725767747 | 0.24464293 | 2.96664096 |
| ENSG00000130255.12 | RPL36 | 71.4319226 | -0.614649743 | 0.20722657 | -2.9660759 |
| ENSG00000185475.10 | TMEM179B | 41.8925098 | -0.434594826 | 0.14653712 | -2.9657662 |
| ENSG00000235865.2 | GSN-AS1 | 553.606385 | 0.900436731 | 0.30387684 | 2.96316344 |
| ENSG00000229117.8 | RPL41 | 19.8560428 | -0.841818898 | 0.28469924 | -2.9568709 |
| ENSG00000149273.14 | RPS3 | 15.4433011 | -1.169572182 | 0.39560424 | -2.9564197 |
| ENSG00000172922.9 | RNASEH2C | 532.889172 | -0.295170674 | 0.09985858 | -2.9558869 |
| ENSG00000166337.9 | TAF10 | 2212.71367 | -0.353839302 | 0.11975374 | -2.9547243 |
| ENSG00000042753.11 | AP2S1 | 10.9319041 | -1.134076803 | 0.38394152 | -2.9537749 |
| ENSG00000168528.11 | SERINC2 | 2.26239399 | -2.378487141 | 0.80520453 | -2.9538919 |
| ENSG00000136048.13 | DRAM1 | 5.9847032 | 1.441549141 | 0.48829363 | 2.9522178 |
| ENSG00000204356.13 | NELFE | 11.4447024 | -0.990759488 | 0.33569258 | -2.9513893 |
| ENSG00000226754.1 | AL606760.1 | 15.995641 | -1.566700288 | 0.53085473 | -2.9512788 |
| ENSG00000231638.1 | LUARIS | 13.8897381 | 1.31399998 | 0.44521347 | 2.95139317 |
| ENSG00000252423.1 | RNU6-229P | 1.35426686 | -2.896625276 | 0.98144078 | -2.951401 |
| ENSG00000090581.9 | GNPTG | 124.588663 | 0.430243628 | 0.14586927 | 2.94951515 |
| ENSG00000280416.1 | AC009084.3 | 189.358347 | 0.478007002 | 0.16207057 | 2.94937575 |
| ENSG00000237009.2 | GLIS3-AS1 | 23.2220843 | -1.040262819 | 0.35281763 | -2.9484434 |
| ENSG00000249695.6 | AC026369.1 | 1.64137023 | -3.179014658 | 1.07809361 | -2.9487371 |
| ENSG00000067369.13 | TP53BP1 | 228.427801 | 0.390045048 | 0.13236734 | 2.94668639 |
| ENSG00000269987.1 | AC004542.2 | 558.646054 | 0.448437283 | 0.15218511 | 2.94665673 |
| ENSG00000261546.1 | AC135782.3 | 33.6684959 | -0.605885036 | 0.20570897 | -2.9453506 |
| ENSG00000151773.12 | CCDC122 | 9.37597402 | 1.183915337 | 0.40207887 | 2.94448531 |
| ENSG00000235652.7 | AL356599.1 | 114.346642 | 0.52991362 | 0.1799668 | 2.94450769 |
| ENSG00000104419.14 | NDRG1 | 13.2222088 | 1.327796134 | 0.45132294 | 2.94200899 |
| ENSG00000278811.4 | LINC00624 | 54.7374231 | -0.675353825 | 0.22957961 | -2.9416977 |
| ENSG00000006118.14 | TMEM132A | 2.71179759 | -1.977443418 | 0.67249921 | -2.9404398 |
| ENSG00000142676.14 | RPL11 | 14.9062379 | -0.808186107 | 0.27496016 | -2.9392844 |
| ENSG00000240661.3 | AC063952.1 | 2900.1155 | 0.537599449 | 0.18295947 | 2.9383527 |
| ENSG00000273212.1 | AC000068.2 | 294.889132 | -0.561835881 | 0.1911866 | -2.9386781 |
| ENSG00000154305.16 | MIA3 | 282.70888 | 0.938422492 | 0.31956595 | 2.93655341 |
| ENSG00000231187.2 | AL356056.2 | 1.84915379 | -3.106643623 | 1.0582641 | -2.9356033 |
| ENSG00000234665.8 | AL512625.3 | 4.74179542 | -1.509684947 | 0.51444221 | -2.9346055 |
| ENSG00000189398.5 | OR7E12P | 25.4885039 | 0.703283901 | 0.23972201 | 2.93374774 |
| ENSG00000267125.2 | AC012615.3 | 156.491727 | -0.452999956 | 0.15441932 | -2.9335704 |

| | | | | | |
|---|---|---|---|---|---|
| ENSG00000092068.19 | SLC7A8 | 17.5678394 | -1.674501549 | 0.57110362 | -2.9320451 |
| ENSG00000128245.14 | YWHAH | 3.33426543 | -2.022602683 | 0.68982375 | -2.9320572 |
| ENSG00000265069.1 | AP002409.1 | 12.0762502 | -1.052190924 | 0.35888756 | -2.9318122 |
| ENSG00000285938.1 | | 9.75960016 | 1.010120701 | 0.34481667 | 2.92944274 |
| ENSG00000168502.17 | MTCL1 | 29.4860526 | 1.417503878 | 0.48432502 | 2.92676162 |
| ENSG00000268970.1 | AC022150.2 | 8.81308652 | 1.034252899 | 0.35342309 | 2.92638748 |
| ENSG00000121764.11 | HCRTR1 | 110.440113 | -0.565622747 | 0.19339351 | -2.9247246 |
| ENSG00000163322.13 | ABRAXAS1 | 157.439352 | -0.522586686 | 0.17871383 | -2.9241535 |
| ENSG00000054965.10 | FAM168A | 31.8007665 | -0.882610878 | 0.30197122 | -2.9228311 |
| ENSG00000182831.11 | C16orf72 | 7.06135952 | -1.167986397 | 0.39980275 | -2.9214066 |
| ENSG00000200714.1 | RF00019 | 1.72312471 | 2.168983773 | 0.74224413 | 2.92219729 |
| ENSG00000260774.1 | AC021087.3 | 3.40688404 | -1.994949124 | 0.68291233 | -2.9212375 |
| ENSG00000263050.1 | AC090617.5 | 12.3001239 | -0.885768776 | 0.30316695 | -2.9217194 |
| ENSG00000275055.1 | AC011468.5 | 10.6118297 | 0.935282983 | 0.32018329 | 2.92108621 |
| ENSG00000084234.17 | APLP2 | 81.4779722 | 0.974119672 | 0.3336006 | 2.92001771 |
| ENSG00000243710.7 | CFAP57 | 111.670341 | -0.585993999 | 0.2006761 | -2.9200986 |
| ENSG00000255250.1 | AP003059.2 | 1.44394511 | 2.851596697 | 0.97642982 | 2.92043179 |
| ENSG00000162032.15 | SPSB3 | 70.4360635 | -0.577261657 | 0.19773535 | -2.9193651 |
| ENSG00000050820.16 | BCAR1 | 16.6822871 | -0.719694055 | 0.24687829 | -2.9151776 |
| ENSG00000095066.11 | HOOK2 | 267.12986 | -0.540397719 | 0.18536089 | -2.9153816 |
| ENSG00000143994.13 | ABHD1 | 30.9433734 | -0.55180705 | 0.18925417 | -2.915693 |
| ENSG00000213742.6 | ZNF337-AS1 | 272.768641 | 0.538321499 | 0.18459384 | 2.91624838 |
| ENSG00000213970.4 | AC006122.1 | 3.09728784 | 2.442439343 | 0.83758593 | 2.9160463 |
| ENSG00000283973.1 | AC099795.1 | 4.989618 | 1.342274479 | 0.46031902 | 2.91596571 |
| ENSG00000145592.13 | RPL37 | 7.2404271 | -1.204171481 | 0.41315669 | -2.9145637 |
| ENSG00000237172.3 | B3GNT9 | 43.8307877 | -0.471953493 | 0.16194307 | -2.9143172 |
| ENSG00000250877.1 | AC095056.1 | 1.34632028 | 3.390476902 | 1.16404492 | 2.91266844 |
| ENSG00000091409.14 | ITGA6 | 17.1063985 | 0.963342894 | 0.33083876 | 2.91181992 |
| ENSG00000116062.14 | MSH6 | 725.679974 | 0.413496887 | 0.1420235 | 2.91146806 |
| ENSG00000240401.8 | AC012358.3 | 200.585466 | 0.615463385 | 0.2113705 | 2.91177519 |
| ENSG00000244040.6 | IL12A-AS1 | 4.80324761 | 1.65152844 | 0.56777175 | 2.90878941 |
| ENSG00000260083.1 | MIR762HG | 385.977673 | -0.411777954 | 0.14173536 | -2.9052592 |
| ENSG00000253406.1 | AC012613.2 | 107.251569 | 0.909513345 | 0.31312682 | 2.90461654 |
| ENSG00000134363.11 | FST | 4.19654358 | 1.594201395 | 0.54927911 | 2.90235214 |
| ENSG00000141753.6 | IGFBP4 | 83.4592363 | -1.036141173 | 0.35692914 | -2.9029324 |
| ENSG00000266872.1 | AC015688.7 | 47.8077169 | 0.66429754 | 0.22888509 | 2.90231896 |
| ENSG00000104852.14 | SNRNP70 | 66.2871352 | -0.809861974 | 0.27912939 | -2.9013855 |
| ENSG00000229400.1 | AL596330.1 | 54.3471606 | 0.937323814 | 0.32306399 | 2.90135654 |
| ENSG00000259773.1 | AC012100.2 | 152.399997 | 0.340922133 | 0.11755725 | 2.90005188 |
| ENSG00000148358.19 | GPR107 | 7.17510122 | -1.046066047 | 0.3607969 | -2.8993211 |
| ENSG00000105664.10 | COMP | 151.095615 | -2.105803414 | 0.72651249 | -2.8985096 |
| ENSG00000258890.6 | CEP95 | 59.9637601 | 0.731165652 | 0.25249581 | 2.89575361 |
| ENSG00000259198.1 | AC020658.2 | 31.8077646 | -0.856845944 | 0.29592357 | -2.8954974 |

| | | | | | |
|---|---|---|---|---|---|
| ENSG00000238278.3 | ALG1L6P | 109.85116 | -0.758683522 | 0.2620887 | -2.8947586 |
| ENSG00000183153.6 | GJD3 | 47.4493142 | -0.807113519 | 0.27893063 | -2.8935994 |
| ENSG00000116251.10 | RPL22 | 2.76536318 | -1.785117125 | 0.61701076 | -2.8931701 |
| ENSG00000272814.1 | AC093732.2 | 80.9538456 | 0.590870033 | 0.20427149 | 2.89257213 |
| ENSG00000176170.13 | SPHK1 | 14.0490932 | -0.947284705 | 0.32757376 | -2.8918211 |
| ENSG00000239552.2 | HOXB-AS2 | 26.9984526 | 1.385742849 | 0.47950771 | 2.88992822 |
| ENSG00000275339.1 | Z99129.1 | 1.82062886 | 2.459037779 | 0.85143302 | 2.88811654 |
| ENSG00000258655.2 | ARHGAP5-AS | 6.14611328 | 1.192732625 | 0.41333148 | 2.88565636 |
| ENSG00000130332.14 | LSM7 | 2.31981363 | -2.095481821 | 0.72638417 | -2.8848121 |
| ENSG00000285704.1 | | 5.69471102 | 1.46536505 | 0.50841043 | 2.88224821 |
| ENSG00000271730.1 | AL390208.1 | 64.5033714 | 0.88358204 | 0.30662854 | 2.881604 |
| ENSG00000163875.15 | MEAF6 | 2.09602782 | -2.528326075 | 0.8788663 | -2.876804 |
| ENSG00000214243.3 | AC004980.2 | 6.53140612 | 1.092838618 | 0.38009641 | 2.87516163 |
| ENSG00000278341.1 | AC138028.6 | 41.4198404 | -0.636027242 | 0.2212495 | -2.8747058 |
| ENSG00000232593.7 | KANTR | 9.2306824 | -1.044681727 | 0.36367401 | -2.8725774 |
| ENSG00000136718.9 | IMP4 | 7.27612877 | -1.002073368 | 0.34911599 | -2.8703164 |
| ENSG00000258749.1 | AL110504.1 | 93.8466695 | -1.009070433 | 0.35168228 | -2.8692672 |
| ENSG00000266957.1 | AC012254.1 | 5.50078984 | 1.318311816 | 0.45940661 | 2.86959695 |
| ENSG00000285728.1 | | 84.5396786 | -0.50269213 | 0.17519025 | -2.869407 |
| ENSG00000101246.19 | ARFRP1 | 6.70749772 | -1.405134096 | 0.49003201 | -2.8674333 |
| ENSG00000228677.1 | TTC3-AS1 | 462.426175 | 0.583415164 | 0.20354792 | 2.86623008 |
| ENSG00000122705.16 | CLTA | 14.265135 | -1.046968612 | 0.36551641 | -2.8643546 |
| ENSG00000145425.9 | RPS3A | 11.2093012 | -0.895468469 | 0.31256792 | -2.8648765 |
| ENSG00000165650.11 | PDZD8 | 4.72930096 | -2.325977049 | 0.81206238 | -2.8642837 |
| ENSG00000184895.7 | SRY | 10.8870309 | 1.596648626 | 0.55732972 | 2.86481872 |
| ENSG00000213341.10 | CHUK | 1.28381315 | 3.045137331 | 1.06305128 | 2.86452534 |
| ENSG00000254829.1 | AP003032.2 | 5.45459631 | -1.478701229 | 0.51603995 | -2.8654782 |
| ENSG00000051620.10 | HEBP2 | 6.5581893 | -1.402670802 | 0.48988285 | -2.863278 |
| ENSG00000235954.6 | TTC28-AS1 | 192.351136 | 0.468457577 | 0.16362254 | 2.86303816 |
| ENSG00000246174.7 | KCTD21-AS1 | 169.476604 | 0.492139855 | 0.1718991 | 2.86295779 |

| pvalue | padj |
|---|---|
| 2.90E-24 | 3.52E-20 |
| 1.45E-21 | 8.81E-18 |
| 7.94E-20 | 3.21E-16 |
| 1.44E-17 | 4.36E-14 |
| 4.39E-16 | 1.07E-12 |
| 4.03E-14 | 7.79E-11 |
| 5.14E-14 | 7.79E-11 |
| 4.94E-14 | 7.79E-11 |
| 2.37E-13 | 3.20E-10 |
| 6.69E-13 | 7.26E-10 |
| 6.09E-13 | 7.26E-10 |
| 7.18E-13 | 7.26E-10 |
| 1.02E-12 | 9.51E-10 |
| 2.78E-12 | 2.41E-09 |
| 3.37E-12 | 2.72E-09 |
| 5.02E-12 | 3.81E-09 |
| 8.69E-12 | 6.20E-09 |
| 1.38E-11 | 9.29E-09 |
| 2.64E-11 | 1.69E-08 |
| 4.32E-11 | 2.62E-08 |
| 5.20E-11 | 3.00E-08 |
| 6.99E-11 | 3.85E-08 |
| 7.57E-11 | 3.99E-08 |
| 1.52E-10 | 7.66E-08 |
| 3.11E-10 | 1.46E-07 |
| 3.13E-10 | 1.46E-07 |
| 4.34E-10 | 1.95E-07 |
| 4.94E-10 | 2.14E-07 |
| 6.83E-10 | 2.86E-07 |
| 9.61E-10 | 3.76E-07 |
| 9.40E-10 | 3.76E-07 |
| 1.17E-09 | 4.28E-07 |
| 1.16E-09 | 4.28E-07 |
| 1.25E-09 | 4.47E-07 |
| 1.36E-09 | 4.73E-07 |
| 1.93E-09 | 6.50E-07 |
| 2.22E-09 | 7.29E-07 |
| 3.04E-09 | 9.69E-07 |
| 3.31E-09 | 1.03E-06 |
| 3.81E-09 | 1.15E-06 |
| 4.10E-09 | 1.21E-06 |
| 4.58E-09 | 1.32E-06 |

| | |
|---|---|
| 4.96E-09 | 1.37E-06 |
| 4.89E-09 | 1.37E-06 |
| 5.58E-09 | 1.47E-06 |
| 5.55E-09 | 1.47E-06 |
| 6.46E-09 | 1.67E-06 |
| 7.66E-09 | 1.94E-06 |
| 8.28E-09 | 2.05E-06 |
| 9.40E-09 | 2.26E-06 |
| 9.52E-09 | 2.26E-06 |
| 1.00E-08 | 2.33E-06 |
| 1.02E-08 | 2.33E-06 |
| 1.19E-08 | 2.68E-06 |
| 1.22E-08 | 2.69E-06 |
| 1.28E-08 | 2.78E-06 |
| 1.37E-08 | 2.91E-06 |
| 1.50E-08 | 3.14E-06 |
| 1.55E-08 | 3.18E-06 |
| 1.81E-08 | 3.65E-06 |
| 1.87E-08 | 3.65E-06 |
| 1.86E-08 | 3.65E-06 |
| 2.03E-08 | 3.90E-06 |
| 2.79E-08 | 5.29E-06 |
| 3.30E-08 | 6.16E-06 |
| 3.52E-08 | 6.47E-06 |
| 4.32E-08 | 7.82E-06 |
| 4.48E-08 | 7.99E-06 |
| 4.97E-08 | 8.73E-06 |
| 5.15E-08 | 8.92E-06 |
| 6.03E-08 | 1.03E-05 |
| 6.15E-08 | 1.04E-05 |
| 6.34E-08 | 1.05E-05 |
| 6.78E-08 | 1.07E-05 |
| 6.79E-08 | 1.07E-05 |
| 6.82E-08 | 1.07E-05 |
| 6.80E-08 | 1.07E-05 |
| 7.24E-08 | 1.11E-05 |
| 7.22E-08 | 1.11E-05 |
| 7.97E-08 | 1.21E-05 |
| 9.91E-08 | 1.48E-05 |
| 1.07E-07 | 1.59E-05 |
| 1.15E-07 | 1.67E-05 |
| 1.26E-07 | 1.81E-05 |
| 1.32E-07 | 1.88E-05 |

| | |
|---|---|
| 1.40E-07 | 1.97E-05 |
| 1.44E-07 | 1.99E-05 |
| 1.45E-07 | 1.99E-05 |
| 1.61E-07 | 2.20E-05 |
| 1.74E-07 | 2.32E-05 |
| 1.74E-07 | 2.32E-05 |
| 2.00E-07 | 2.64E-05 |
| 2.16E-07 | 2.81E-05 |
| 2.36E-07 | 3.01E-05 |
| 2.35E-07 | 3.01E-05 |
| 2.70E-07 | 3.41E-05 |
| 2.82E-07 | 3.53E-05 |
| 2.87E-07 | 3.54E-05 |
| 3.29E-07 | 4.03E-05 |
| 3.41E-07 | 4.13E-05 |
| 3.70E-07 | 4.45E-05 |
| 3.86E-07 | 4.59E-05 |
| 4.09E-07 | 4.82E-05 |
| 4.32E-07 | 4.99E-05 |
| 4.29E-07 | 4.99E-05 |
| 4.82E-07 | 5.51E-05 |
| 4.94E-07 | 5.60E-05 |
| 5.07E-07 | 5.69E-05 |
| 5.34E-07 | 5.93E-05 |
| 5.83E-07 | 6.42E-05 |
| 6.55E-07 | 7.15E-05 |
| 6.73E-07 | 7.28E-05 |
| 7.91E-07 | 8.48E-05 |
| 7.98E-07 | 8.49E-05 |
| 8.09E-07 | 8.53E-05 |
| 8.48E-07 | 8.86E-05 |
| 8.56E-07 | 8.87E-05 |
| 8.75E-07 | 8.99E-05 |
| 9.08E-07 | 9.25E-05 |
| 9.38E-07 | 9.48E-05 |
| 1.20E-06 | 0.00012035 |
| 1.26E-06 | 0.00012563 |
| 1.28E-06 | 0.0001261 |
| 1.30E-06 | 0.00012665 |
| 1.34E-06 | 0.0001296 |
| 1.36E-06 | 0.00013131 |
| 1.38E-06 | 0.0001317 |
| 1.46E-06 | 0.00013864 |

| | |
|---|---|
| 1.64E-06 | 0.00015421 |
| 1.71E-06 | 0.00015912 |
| 1.83E-06 | 0.00016721 |
| 1.82E-06 | 0.00016721 |
| 1.83E-06 | 0.00016721 |
| 1.87E-06 | 0.00016911 |
| 1.92E-06 | 0.00017245 |
| 2.09E-06 | 0.0001862 |
| 2.11E-06 | 0.00018667 |
| 2.17E-06 | 0.0001895 |
| 2.17E-06 | 0.0001895 |
| 2.22E-06 | 0.00019234 |
| 2.24E-06 | 0.00019235 |
| 2.29E-06 | 0.00019467 |
| 2.30E-06 | 0.00019467 |
| 2.35E-06 | 0.00019772 |
| 2.57E-06 | 0.00021307 |
| 2.56E-06 | 0.00021307 |
| 2.76E-06 | 0.00022773 |
| 2.88E-06 | 0.0002355 |
| 2.94E-06 | 0.00023927 |
| 2.97E-06 | 0.00023985 |
| 3.13E-06 | 0.00025137 |
| 3.20E-06 | 0.00025383 |
| 3.20E-06 | 0.00025383 |
| 3.28E-06 | 0.00025783 |
| 3.34E-06 | 0.00026125 |
| 3.37E-06 | 0.00026154 |
| 3.56E-06 | 0.00027505 |
| 3.79E-06 | 0.00029111 |
| 3.84E-06 | 0.00029304 |
| 3.90E-06 | 0.00029556 |
| 3.99E-06 | 0.00030023 |
| 4.23E-06 | 0.0003166 |
| 4.49E-06 | 0.00033378 |
| 4.60E-06 | 0.00033811 |
| 4.57E-06 | 0.00033811 |
| 4.65E-06 | 0.00033942 |
| 4.94E-06 | 0.0003583 |
| 4.98E-06 | 0.00035965 |
| 5.33E-06 | 0.00038202 |
| 5.42E-06 | 0.00038525 |
| 5.43E-06 | 0.00038525 |

| | |
|---|---|
| 5.57E-06 | 0.00039235 |
| 5.60E-06 | 0.00039251 |
| 5.81E-06 | 0.00040508 |
| 5.86E-06 | 0.0004059 |
| 6.12E-06 | 0.00042163 |
| 6.71E-06 | 0.00045464 |
| 6.65E-06 | 0.00045464 |
| 6.69E-06 | 0.00045464 |
| 6.76E-06 | 0.00045508 |
| 6.90E-06 | 0.00046238 |
| 7.13E-06 | 0.00047466 |
| 7.25E-06 | 0.00048006 |
| 7.41E-06 | 0.00048838 |
| 7.95E-06 | 0.00052096 |
| 8.23E-06 | 0.00053626 |
| 8.51E-06 | 0.00055138 |
| 8.69E-06 | 0.00055749 |
| 8.67E-06 | 0.00055749 |
| 8.75E-06 | 0.00055815 |
| 8.83E-06 | 0.0005605 |
| 9.14E-06 | 0.00057433 |
| 9.12E-06 | 0.00057433 |
| 9.46E-06 | 0.00058946 |
| 9.48E-06 | 0.00058946 |
| 9.72E-06 | 0.00060101 |
| 9.77E-06 | 0.00060136 |
| 1.01E-05 | 0.00061785 |
| 1.04E-05 | 0.0006306 |
| 1.05E-05 | 0.00063615 |
| 1.08E-05 | 0.00065256 |
| 1.11E-05 | 0.00066576 |
| 1.12E-05 | 0.00067157 |
| 1.13E-05 | 0.00067194 |
| 1.14E-05 | 0.00067194 |
| 1.20E-05 | 0.00070642 |
| 1.24E-05 | 0.0007218 |
| 1.24E-05 | 0.0007218 |
| 1.25E-05 | 0.00072436 |
| 1.27E-05 | 0.00073445 |
| 1.29E-05 | 0.00073989 |
| 1.34E-05 | 0.00076547 |
| 1.36E-05 | 0.00077553 |
| 1.37E-05 | 0.00077759 |

| | |
|---|---|
| 1.40E-05 | 0.00078743 |
| 1.43E-05 | 0.00080333 |
| 1.49E-05 | 0.00082945 |
| 1.49E-05 | 0.00082945 |
| 1.52E-05 | 0.0008376 |
| 1.52E-05 | 0.0008376 |
| 1.54E-05 | 0.00084303 |
| 1.54E-05 | 0.00084337 |
| 1.60E-05 | 0.00086837 |
| 1.63E-05 | 0.00088083 |
| 1.71E-05 | 0.00092227 |
| 1.75E-05 | 0.00094113 |
| 1.81E-05 | 0.00096857 |
| 1.83E-05 | 0.00097455 |
| 1.86E-05 | 0.00098586 |
| 1.89E-05 | 0.00099791 |
| 1.99E-05 | 0.00103295 |
| 1.98E-05 | 0.00103295 |
| 1.97E-05 | 0.00103295 |
| 2.00E-05 | 0.00103703 |
| 2.03E-05 | 0.0010453 |
| 2.08E-05 | 0.00106621 |
| 2.13E-05 | 0.00109142 |
| 2.18E-05 | 0.00110634 |
| 2.17E-05 | 0.00110634 |
| 2.21E-05 | 0.00111468 |
| 2.23E-05 | 0.00112077 |
| 2.26E-05 | 0.00113266 |
| 2.28E-05 | 0.00113594 |
| 2.39E-05 | 0.00118734 |
| 2.45E-05 | 0.00121203 |
| 2.46E-05 | 0.0012132 |
| 2.50E-05 | 0.00122596 |
| 2.58E-05 | 0.00126015 |
| 2.59E-05 | 0.00126015 |
| 2.62E-05 | 0.00126874 |
| 2.66E-05 | 0.00128571 |
| 2.68E-05 | 0.00128842 |
| 2.70E-05 | 0.0012892 |
| 2.70E-05 | 0.0012892 |
| 2.74E-05 | 0.00130123 |
| 2.78E-05 | 0.0013077 |
| 2.76E-05 | 0.0013077 |

| | |
|---|---|
| 2.78E-05 | 0.0013077 |
| 2.81E-05 | 0.00130807 |
| 2.80E-05 | 0.00130807 |
| 2.82E-05 | 0.00130849 |
| 2.87E-05 | 0.00132928 |
| 2.96E-05 | 0.001361 |
| 2.96E-05 | 0.001361 |
| 2.98E-05 | 0.00136213 |
| 3.00E-05 | 0.00136856 |
| 3.06E-05 | 0.0013911 |
| 3.12E-05 | 0.00140998 |
| 3.16E-05 | 0.00142356 |
| 3.18E-05 | 0.00142924 |
| 3.21E-05 | 0.00143549 |
| 3.23E-05 | 0.00144021 |
| 3.32E-05 | 0.00147526 |
| 3.34E-05 | 0.00147887 |
| 3.36E-05 | 0.00148187 |
| 3.60E-05 | 0.00158291 |
| 3.63E-05 | 0.00158754 |
| 3.68E-05 | 0.00160501 |
| 3.70E-05 | 0.00160676 |
| 3.73E-05 | 0.00161194 |
| 3.74E-05 | 0.00161194 |
| 3.77E-05 | 0.00161915 |
| 3.84E-05 | 0.00164451 |
| 4.07E-05 | 0.00173762 |
| 4.11E-05 | 0.00174801 |
| 4.18E-05 | 0.00176397 |
| 4.17E-05 | 0.00176397 |
| 4.31E-05 | 0.00181537 |
| 4.67E-05 | 0.00194288 |
| 4.68E-05 | 0.00194288 |
| 4.63E-05 | 0.00194288 |
| 4.68E-05 | 0.00194288 |
| 4.72E-05 | 0.00195143 |
| 4.74E-05 | 0.00195537 |
| 4.85E-05 | 0.00199217 |
| 4.86E-05 | 0.00199239 |
| 4.88E-05 | 0.00199313 |
| 4.90E-05 | 0.00199324 |
| 5.00E-05 | 0.00202914 |
| 5.16E-05 | 0.00208511 |

| | |
|---|---|
| 5.35E-05 | 0.00215668 |
| 5.43E-05 | 0.00217892 |
| 5.51E-05 | 0.00220633 |
| 5.72E-05 | 0.00228177 |
| 5.79E-05 | 0.00230187 |
| 6.04E-05 | 0.0023844 |
| 6.02E-05 | 0.0023844 |
| 6.13E-05 | 0.00240346 |
| 6.11E-05 | 0.00240346 |
| 6.38E-05 | 0.00249523 |
| 6.49E-05 | 0.00252944 |
| 6.52E-05 | 0.00253152 |
| 6.59E-05 | 0.00254707 |
| 6.60E-05 | 0.00254707 |
| 6.69E-05 | 0.00257362 |
| 6.74E-05 | 0.00257809 |
| 6.72E-05 | 0.00257809 |
| 6.84E-05 | 0.00260082 |
| 6.83E-05 | 0.00260082 |
| 7.04E-05 | 0.00266753 |
| 7.17E-05 | 0.00270701 |
| 7.29E-05 | 0.00272949 |
| 7.29E-05 | 0.00272949 |
| 7.29E-05 | 0.00272949 |
| 7.40E-05 | 0.00276128 |
| 7.53E-05 | 0.00279877 |
| 7.62E-05 | 0.00280613 |
| 7.60E-05 | 0.00280613 |
| 7.58E-05 | 0.00280613 |
| 7.69E-05 | 0.00282549 |
| 7.77E-05 | 0.00283012 |
| 7.75E-05 | 0.00283012 |
| 7.75E-05 | 0.00283012 |
| 8.02E-05 | 0.00291256 |
| 8.05E-05 | 0.00291295 |
| 8.21E-05 | 0.00295921 |
| 8.23E-05 | 0.00295921 |
| 8.32E-05 | 0.00298481 |
| 8.59E-05 | 0.00307338 |
| 8.72E-05 | 0.00310925 |
| 9.07E-05 | 0.0032237 |
| 9.27E-05 | 0.00328543 |
| 9.31E-05 | 0.00328911 |

| | |
|---|---|
| 9.34E-05 | 0.00329268 |
| 9.41E-05 | 0.00330618 |
| 9.67E-05 | 0.0033507 |
| 9.61E-05 | 0.0033507 |
| 9.70E-05 | 0.0033507 |
| 9.69E-05 | 0.0033507 |
| 9.62E-05 | 0.0033507 |
| 9.67E-05 | 0.0033507 |
| 9.80E-05 | 0.00336693 |
| 9.80E-05 | 0.00336693 |
| 9.87E-05 | 0.00337563 |
| 9.88E-05 | 0.00337563 |
| 9.95E-05 | 0.00338712 |
| 0.00010124 | 0.00343782 |
| 0.00010192 | 0.00345135 |
| 0.00010249 | 0.00346108 |
| 0.00010301 | 0.00346901 |
| 0.00010348 | 0.00347502 |
| 0.00010418 | 0.00348886 |
| 0.00010455 | 0.00349167 |
| 0.00010548 | 0.00350754 |
| 0.00010561 | 0.00350754 |
| 0.00010875 | 0.00360227 |
| 0.00011044 | 0.00364805 |
| 0.00011111 | 0.00366033 |
| 0.00011619 | 0.00381725 |
| 0.00011696 | 0.00383234 |
| 0.00012129 | 0.00396348 |
| 0.00012452 | 0.00403616 |
| 0.00012441 | 0.00403616 |
| 0.0001241 | 0.00403616 |
| 0.00012608 | 0.00407579 |
| 0.00012642 | 0.00407596 |
| 0.00012675 | 0.00407596 |
| 0.00012774 | 0.00409682 |
| 0.00012847 | 0.0041094 |
| 0.00012935 | 0.00412661 |
| 0.00013157 | 0.0041864 |
| 0.0001355 | 0.00430024 |
| 0.00013716 | 0.00434134 |
| 0.00013903 | 0.00438921 |
| 0.00014177 | 0.004464 |
| 0.00014607 | 0.00458622 |

| | |
|---|---|
| 0.00014641 | 0.00458622 |
| 0.00015015 | 0.0046913 |
| 0.00015256 | 0.0047545 |
| 0.00015455 | 0.00479172 |
| 0.00015422 | 0.00479172 |
| 0.00015653 | 0.00484096 |
| 0.00015749 | 0.00485804 |
| 0.00015855 | 0.00486093 |
| 0.00015878 | 0.00486093 |
| 0.00015868 | 0.00486093 |
| 0.00016029 | 0.0048947 |
| 0.00016581 | 0.00505056 |
| 0.00016664 | 0.00506303 |
| 0.00016869 | 0.00511252 |
| 0.00016937 | 0.00512049 |
| 0.00017539 | 0.005276 |
| 0.00017496 | 0.005276 |
| 0.00018471 | 0.0055427 |
| 0.00018626 | 0.00557551 |
| 0.00018721 | 0.00558988 |
| 0.00018888 | 0.00562599 |
| 0.00018936 | 0.00562649 |
| 0.00019391 | 0.00573365 |
| 0.00019372 | 0.00573365 |
| 0.00019713 | 0.00577485 |
| 0.00019608 | 0.00577485 |
| 0.00019697 | 0.00577485 |
| 0.00019721 | 0.00577485 |
| 0.00020733 | 0.00604858 |
| 0.00020756 | 0.00604858 |
| 0.00021252 | 0.00616361 |
| 0.00021237 | 0.00616361 |
| 0.00021329 | 0.00617121 |
| 0.00021448 | 0.00617606 |
| 0.00021418 | 0.00617606 |
| 0.00021517 | 0.00618136 |
| 0.00021649 | 0.00619064 |
| 0.00021652 | 0.00619064 |
| 0.0002183 | 0.00622703 |
| 0.00022019 | 0.00625137 |
| 0.00022016 | 0.00625137 |
| 0.00022379 | 0.0063388 |
| 0.00022704 | 0.00641586 |

| | |
|---|---|
| 0.00022906 | 0.00645782 |
| 0.00023099 | 0.00649712 |
| 0.00023652 | 0.00662205 |
| 0.00023652 | 0.00662205 |
| 0.00023729 | 0.00662818 |
| 0.00023946 | 0.00667339 |
| 0.00024146 | 0.0067134 |
| 0.000242 | 0.0067134 |
| 0.000244 | 0.00675336 |
| 0.00024559 | 0.00676718 |
| 0.00024561 | 0.00676718 |
| 0.00024776 | 0.00681077 |
| 0.00025045 | 0.00686925 |
| 0.00025158 | 0.00687164 |
| 0.00025167 | 0.00687164 |
| 0.00025279 | 0.00687595 |
| 0.00025408 | 0.00687595 |
| 0.0002541 | 0.00687595 |
| 0.000253 | 0.00687595 |
| 0.000255 | 0.00688497 |
| 0.00025666 | 0.00691434 |
| 0.00026016 | 0.00699326 |
| 0.00026407 | 0.00708257 |
| 0.000269 | 0.00719246 |
| 0.00026935 | 0.00719246 |
| 0.00027395 | 0.00728299 |
| 0.00027365 | 0.00728299 |
| 0.00027624 | 0.00732791 |
| 0.00027799 | 0.00735826 |
| 0.00028064 | 0.0074121 |
| 0.00028462 | 0.00750108 |
| 0.00028626 | 0.00752783 |
| 0.00028843 | 0.00756848 |
| 0.00029172 | 0.00762177 |
| 0.00029147 | 0.00762177 |
| 0.00029819 | 0.0077742 |
| 0.00030027 | 0.00781164 |
| 0.00030226 | 0.0078129 |
| 0.00030123 | 0.0078129 |
| 0.00030168 | 0.0078129 |
| 0.00030965 | 0.00798695 |
| 0.000323 | 0.00831366 |
| 0.00032762 | 0.00841131 |

| | |
|---|---|
| 0.00032818 | 0.00841131 |
| 0.00033015 | 0.00844379 |
| 0.00033107 | 0.00844973 |
| 0.000333 | 0.00848088 |
| 0.00034049 | 0.00865354 |
| 0.00034427 | 0.00870729 |
| 0.00034391 | 0.00870729 |
| 0.00034476 | 0.00870729 |
| 0.00034861 | 0.00878627 |
| 0.00035238 | 0.0088313 |
| 0.00035258 | 0.0088313 |
| 0.00035128 | 0.0088313 |
| 0.00035655 | 0.00891227 |
| 0.0003576 | 0.00892001 |
| 0.00035852 | 0.00892472 |
| 0.0003594 | 0.00892822 |
| 0.0003602 | 0.00892989 |
| 0.0003677 | 0.00909724 |
| 0.00037228 | 0.00919165 |
| 0.00037309 | 0.0091931 |
| 0.00038243 | 0.00940398 |
| 0.0003946 | 0.00967939 |
| 0.00039522 | 0.00967939 |
| 0.00039842 | 0.00973795 |
| 0.0004013 | 0.00978856 |
| 0.00040239 | 0.00979557 |
| 0.00041053 | 0.00997375 |
| 0.00041629 | 0.01007109 |
| 0.00041546 | 0.01007109 |
| 0.00041703 | 0.01007109 |
| 0.00042044 | 0.01013319 |
| 0.0004214 | 0.01013617 |
| 0.00042365 | 0.01017019 |
| 0.00042694 | 0.01022892 |
| 0.00043363 | 0.01036855 |
| 0.00043561 | 0.01039545 |
| 0.00043669 | 0.01040071 |
| 0.00043858 | 0.01042252 |
| 0.00043932 | 0.01042252 |
| 0.00044134 | 0.01044992 |
| 0.0004459 | 0.01052849 |
| 0.00044639 | 0.01052849 |
| 0.00045038 | 0.01060193 |

| | |
|---|---|
| 0.00045422 | 0.01067146 |
| 0.00046271 | 0.01084987 |
| 0.00046404 | 0.01085699 |
| 0.00046491 | 0.01085699 |
| 0.0004657 | 0.01085699 |
| 0.00046761 | 0.01088068 |
| 0.00047055 | 0.01092808 |
| 0.00047901 | 0.01110323 |
| 0.00048305 | 0.01117563 |
| 0.00048467 | 0.0111917 |
| 0.00049781 | 0.01147336 |
| 0.00050131 | 0.01153203 |
| 0.00050787 | 0.01166077 |
| 0.00050923 | 0.01166388 |
| 0.00050993 | 0.01166388 |
| 0.00051645 | 0.01176869 |
| 0.00051618 | 0.01176869 |
| 0.00052481 | 0.01193664 |
| 0.0005332 | 0.01210487 |
| 0.0005389 | 0.01220537 |
| 0.00054197 | 0.01220537 |
| 0.00054135 | 0.01220537 |
| 0.00054325 | 0.01220537 |
| 0.00054179 | 0.01220537 |
| 0.00054468 | 0.01220537 |
| 0.00054444 | 0.01220537 |
| 0.00055283 | 0.01236532 |
| 0.00055563 | 0.01240509 |
| 0.00056442 | 0.01257797 |
| 0.00056701 | 0.01261268 |
| 0.00056894 | 0.01263224 |
| 0.00057056 | 0.01264519 |
| 0.00057647 | 0.012706 |
| 0.0005759 | 0.012706 |
| 0.00057563 | 0.012706 |
| 0.00057772 | 0.012706 |
| 0.00057855 | 0.012706 |
| 0.00058289 | 0.01277836 |
| 0.00058613 | 0.01282614 |
| 0.00059036 | 0.01289537 |
| 0.00059174 | 0.01290234 |
| 0.00059363 | 0.01292022 |
| 0.00059602 | 0.0129491 |

| | |
|---|---|
| 0.00060894 | 0.01315893 |
| 0.00060685 | 0.01315893 |
| 0.00060814 | 0.01315893 |
| 0.000617 | 0.0133095 |
| 0.00061941 | 0.01331398 |
| 0.00061838 | 0.01331398 |
| 0.00062478 | 0.01340564 |
| 0.00062711 | 0.01343183 |
| 0.00062925 | 0.01345398 |
| 0.00063098 | 0.0134672 |
| 0.0006332 | 0.01349075 |
| 0.00063956 | 0.01353411 |
| 0.00063664 | 0.01353411 |
| 0.0006397 | 0.01353411 |
| 0.00063782 | 0.01353411 |
| 0.0006478 | 0.01368174 |
| 0.00065496 | 0.01380881 |
| 0.00065746 | 0.01383747 |
| 0.00066076 | 0.0138828 |
| 0.00066776 | 0.01400557 |
| 0.00067241 | 0.01400623 |
| 0.00067156 | 0.01400623 |
| 0.00067036 | 0.01400623 |
| 0.00067097 | 0.01400623 |
| 0.00067873 | 0.0140771 |
| 0.00067814 | 0.0140771 |
| 0.0006793 | 0.0140771 |
| 0.00069624 | 0.01436175 |
| 0.00069659 | 0.01436175 |
| 0.00069565 | 0.01436175 |
| 0.00070067 | 0.01442139 |
| 0.00070618 | 0.01449904 |
| 0.00070683 | 0.01449904 |
| 0.0007105 | 0.01450065 |
| 0.00070828 | 0.01450065 |
| 0.00071018 | 0.01450065 |
| 0.00071246 | 0.01451625 |
| 0.00071548 | 0.01455322 |
| 0.00072095 | 0.0146399 |
| 0.00072329 | 0.01466289 |
| 0.00072467 | 0.01466641 |
| 0.00072835 | 0.01470268 |
| 0.00072889 | 0.01470268 |

| | |
|---|---|
| 0.00073519 | 0.01480508 |
| 0.0007367 | 0.01481104 |
| 0.00074228 | 0.01489849 |
| 0.00074567 | 0.01494172 |
| 0.0007493 | 0.01498968 |
| 0.00075394 | 0.01505759 |
| 0.00075641 | 0.01508224 |
| 0.00077225 | 0.01537275 |
| 0.00077468 | 0.01539573 |
| 0.00077829 | 0.01544122 |
| 0.00077951 | 0.01544122 |
| 0.00079197 | 0.01566239 |
| 0.00079514 | 0.01568486 |
| 0.00079569 | 0.01568486 |
| 0.00079849 | 0.0157145 |
| 0.00080905 | 0.01589653 |
| 0.0008114 | 0.01591678 |
| 0.00081818 | 0.01602255 |
| 0.00082006 | 0.01602255 |
| 0.00082138 | 0.01602255 |
| 0.00082208 | 0.01602255 |
| 0.00082421 | 0.01603846 |
| 0.00082886 | 0.01607719 |
| 0.00082803 | 0.01607719 |
| 0.00083284 | 0.01612855 |
| 0.00083763 | 0.01619543 |
| 0.00084268 | 0.01626723 |
| 0.00084591 | 0.01630357 |
| 0.00084906 | 0.01633827 |
| 0.00085433 | 0.01641372 |
| 0.00085993 | 0.01649513 |
| 0.00087187 | 0.01669769 |
| 0.00087768 | 0.01674484 |
| 0.00087847 | 0.01674484 |
| 0.00087639 | 0.01674484 |
| 0.00088131 | 0.01677249 |
| 0.00089625 | 0.01703021 |
| 0.00090638 | 0.01718083 |
| 0.00090701 | 0.01718083 |
| 0.00090993 | 0.01720916 |
| 0.00091219 | 0.01722496 |
| 0.0009259 | 0.01745672 |
| 0.00093039 | 0.01751416 |

| | |
|---|---|
| 0.00093199 | 0.01751716 |
| 0.00093888 | 0.0176193 |
| 0.0009647 | 0.01804784 |
| 0.00096418 | 0.01804784 |
| 0.00097583 | 0.01822809 |
| 0.00097741 | 0.01822946 |
| 0.00098531 | 0.01829243 |
| 0.00098312 | 0.01829243 |
| 0.00098525 | 0.01829243 |
| 0.00099381 | 0.01842187 |
| 0.00100174 | 0.01851233 |
| 0.0010013 | 0.01851233 |
| 0.00101797 | 0.01878361 |
| 0.00102741 | 0.0189291 |
| 0.00104629 | 0.01924757 |
| 0.0010528 | 0.01933795 |
| 0.00106158 | 0.01946984 |
| 0.00106659 | 0.01953216 |
| 0.00107542 | 0.01966414 |
| 0.00109253 | 0.01993008 |
| 0.00109374 | 0.01993008 |
| 0.0010949 | 0.01993008 |
| 0.00110018 | 0.01996636 |
| 0.00109904 | 0.01996636 |
| 0.00110428 | 0.02001078 |
| 0.00110981 | 0.02008098 |
| 0.00112303 | 0.0202372 |
| 0.0011216 | 0.0202372 |
| 0.00112345 | 0.0202372 |
| 0.00112568 | 0.02024724 |
| 0.00112877 | 0.02027266 |
| 0.0011446 | 0.0204962 |
| 0.0011445 | 0.0204962 |
| 0.00117605 | 0.02102838 |
| 0.00118832 | 0.02121645 |
| 0.00119388 | 0.02128448 |
| 0.00120635 | 0.02147523 |
| 0.00121751 | 0.02151584 |
| 0.00121552 | 0.02151584 |
| 0.00121179 | 0.02151584 |
| 0.00121503 | 0.02151584 |
| 0.00121643 | 0.02151584 |
| 0.00122 | 0.02152844 |

| | |
|---|---|
| 0.00122306 | 0.02155102 |
| 0.00122483 | 0.02155102 |
| 0.00122689 | 0.02155592 |
| 0.00123552 | 0.02167622 |
| 0.001239 | 0.0216779 |
| 0.0012392 | 0.0216779 |
| 0.00124368 | 0.02172489 |
| 0.00124976 | 0.02179976 |
| 0.00125727 | 0.0218993 |
| 0.00126929 | 0.02195758 |
| 0.00126751 | 0.02195758 |
| 0.00126358 | 0.02195758 |
| 0.00126967 | 0.02195758 |
| 0.0012657 | 0.02195758 |
| 0.00129869 | 0.02242745 |
| 0.00130115 | 0.02243798 |
| 0.00131258 | 0.02250695 |
| 0.00130988 | 0.02250695 |
| 0.00131256 | 0.02250695 |
| 0.00131179 | 0.02250695 |
| 0.00132204 | 0.02263709 |
| 0.0013269 | 0.02268831 |
| 0.00133094 | 0.02272537 |
| 0.00133965 | 0.02284184 |
| 0.00134866 | 0.02296328 |
| 0.00135602 | 0.02301472 |
| 0.00135729 | 0.02301472 |
| 0.00135738 | 0.02301472 |
| 0.00136023 | 0.02303075 |
| 0.0013623 | 0.02303369 |
| 0.00136425 | 0.02303458 |
| 0.00137258 | 0.02314292 |
| 0.00137737 | 0.02319147 |
| 0.00138311 | 0.0232558 |
| 0.00138739 | 0.02326321 |
| 0.0013871 | 0.02326321 |
| 0.00139728 | 0.02339675 |
| 0.001425 | 0.0238279 |
| 0.00142844 | 0.02385262 |
| 0.00143342 | 0.02390284 |
| 0.00143804 | 0.02394688 |
| 0.00144584 | 0.02404372 |
| 0.00144997 | 0.02407937 |

| | |
|---|---|
| 0.00145795 | 0.02417889 |
| 0.00146059 | 0.02418956 |
| 0.00146296 | 0.02419578 |
| 0.0014704 | 0.02428565 |
| 0.00148444 | 0.02448411 |
| 0.00149028 | 0.02453852 |
| 0.00149178 | 0.02453852 |
| 0.00153106 | 0.02508248 |
| 0.00152716 | 0.02508248 |
| 0.00152988 | 0.02508248 |
| 0.00154779 | 0.02532243 |
| 0.00155768 | 0.02544987 |
| 0.00156489 | 0.02553321 |
| 0.0015691 | 0.02553398 |
| 0.00156915 | 0.02553398 |
| 0.00157257 | 0.02555533 |
| 0.00158095 | 0.02565713 |
| 0.00158603 | 0.02570511 |
| 0.00159182 | 0.02576458 |
| 0.00160003 | 0.02586293 |
| 0.00160744 | 0.02587911 |
| 0.0016036 | 0.02587911 |
| 0.00160612 | 0.02587911 |
| 0.0016384 | 0.02634263 |
| 0.00165121 | 0.02651341 |
| 0.00166383 | 0.02668065 |
| 0.00167189 | 0.02677455 |
| 0.00168229 | 0.02690556 |
| 0.00168549 | 0.02692127 |
| 0.00173124 | 0.02758941 |
| 0.00173188 | 0.02758941 |
| 0.00173711 | 0.02763641 |
| 0.00174038 | 0.0276522 |
| 0.00175967 | 0.02792206 |
| 0.00176613 | 0.02798795 |
| 0.00177073 | 0.02802421 |
| 0.00177396 | 0.02803873 |
| 0.00178612 | 0.02815758 |
| 0.00178577 | 0.02815758 |
| 0.00179875 | 0.02831975 |
| 0.0018065 | 0.02840497 |
| 0.00181828 | 0.02855306 |
| 0.00182545 | 0.0286287 |

| | |
|---|---|
| 0.00183082 | 0.02863877 |
| 0.00183078 | 0.02863877 |
| 0.00183497 | 0.02866673 |
| 0.00185123 | 0.02888344 |
| 0.00186131 | 0.02900345 |
| 0.00186817 | 0.02907297 |
| 0.00187378 | 0.0291229 |
| 0.00188691 | 0.02928937 |
| 0.00189058 | 0.02930879 |
| 0.00189358 | 0.02931777 |
| 0.00190306 | 0.02936405 |
| 0.00190018 | 0.02936405 |
| 0.00190383 | 0.02936405 |
| 0.00190776 | 0.0293873 |
| 0.00192713 | 0.02957506 |
| 0.0019261 | 0.02957506 |
| 0.00192971 | 0.02957506 |
| 0.00192835 | 0.02957506 |
| 0.00193637 | 0.02963959 |
| 0.00194667 | 0.0296888 |
| 0.00194693 | 0.0296888 |
| 0.00194349 | 0.0296888 |
| 0.00196295 | 0.02989553 |
| 0.00196618 | 0.0299072 |
| 0.00197225 | 0.02996188 |
| 0.00197777 | 0.03000812 |
| 0.00198102 | 0.03001988 |
| 0.00199364 | 0.03017334 |
| 0.0020092 | 0.03037098 |
| 0.00201344 | 0.03039715 |
| 0.00203148 | 0.03055538 |
| 0.00202919 | 0.03055538 |
| 0.00202768 | 0.03055538 |
| 0.00204688 | 0.03069961 |
| 0.00204561 | 0.03069961 |
| 0.00204867 | 0.03069961 |
| 0.00205365 | 0.03073634 |
| 0.0020622 | 0.03078828 |
| 0.00206064 | 0.03078828 |
| 0.00206797 | 0.03083641 |
| 0.00207487 | 0.03084068 |
| 0.00207468 | 0.03084068 |
| 0.00207843 | 0.03084068 |

| | |
|---|---|
| 0.00207743 | 0.03084068 |
| 0.00208652 | 0.03092281 |
| 0.00212046 | 0.03138615 |
| 0.00212296 | 0.03138615 |
| 0.00212999 | 0.0314517 |
| 0.00213402 | 0.03147296 |
| 0.00214335 | 0.03157204 |
| 0.00215526 | 0.03163219 |
| 0.00215179 | 0.03163219 |
| 0.00215303 | 0.03163219 |
| 0.00216602 | 0.03175171 |
| 0.0021746 | 0.03183897 |
| 0.00218157 | 0.03186407 |
| 0.00217965 | 0.03186407 |
| 0.00218503 | 0.0318762 |
| 0.00220326 | 0.03210345 |
| 0.00223205 | 0.03242683 |
| 0.00223049 | 0.03242683 |
| 0.00223347 | 0.03242683 |
| 0.00224725 | 0.03258775 |
| 0.00225093 | 0.03260222 |
| 0.00225647 | 0.03264341 |
| 0.00226435 | 0.03271844 |
| 0.00227053 | 0.03276861 |
| 0.00227935 | 0.03283824 |
| 0.00228077 | 0.03283824 |
| 0.00230617 | 0.0331167 |
| 0.00230957 | 0.0331167 |
| 0.00231104 | 0.0331167 |
| 0.00230624 | 0.0331167 |
| 0.00232083 | 0.03321774 |
| 0.00232848 | 0.03328797 |
| 0.00234217 | 0.03344414 |
| 0.00237017 | 0.03380414 |
| 0.00238146 | 0.03388546 |
| 0.00238032 | 0.03388546 |
| 0.00238598 | 0.03391 |
| 0.00240502 | 0.03414062 |
| 0.00242648 | 0.03440487 |
| 0.00242963 | 0.03440942 |
| 0.00243333 | 0.03442159 |
| 0.00243873 | 0.03445765 |
| 0.00248852 | 0.03512023 |

| | |
|---|---|
| 0.0025335 | 0.03567202 |
| 0.00253065 | 0.03567202 |
| 0.00255514 | 0.03593503 |
| 0.00258368 | 0.03629426 |
| 0.00259734 | 0.03644389 |
| 0.00260682 | 0.03653471 |
| 0.00261247 | 0.03657153 |
| 0.00261605 | 0.03657951 |
| 0.00262622 | 0.03667927 |
| 0.00263172 | 0.03671389 |
| 0.00265514 | 0.03695548 |
| 0.00265349 | 0.03695548 |
| 0.00266628 | 0.03702978 |
| 0.00266658 | 0.03702978 |
| 0.0026721 | 0.03706387 |
| 0.00268405 | 0.0371872 |
| 0.00269168 | 0.03725028 |
| 0.00270162 | 0.03734515 |
| 0.00270945 | 0.03738506 |
| 0.00271375 | 0.03738506 |
| 0.00271106 | 0.03738506 |
| 0.00272068 | 0.03743794 |
| 0.00273004 | 0.03748164 |
| 0.00272951 | 0.03748164 |
| 0.00273892 | 0.03756097 |
| 0.00276131 | 0.03778253 |
| 0.0027594 | 0.03778253 |
| 0.0027646 | 0.03778492 |
| 0.00281379 | 0.0384139 |
| 0.0028251 | 0.0385249 |
| 0.00282924 | 0.03853803 |
| 0.00284163 | 0.03866336 |
| 0.00285363 | 0.03877297 |
| 0.00285608 | 0.03877297 |
| 0.00286922 | 0.03890774 |
| 0.0028974 | 0.03920222 |
| 0.00289433 | 0.03920222 |
| 0.00291706 | 0.03934701 |
| 0.00291784 | 0.03934701 |
| 0.00291376 | 0.03934701 |
| 0.00294097 | 0.03946673 |
| 0.00294645 | 0.03946673 |
| 0.00294951 | 0.03946673 |

| | |
|---|---|
| 0.00294032 | 0.03946673 |
| 0.00294549 | 0.03946673 |
| 0.00294645 | 0.03946673 |
| 0.0029307 | 0.03946673 |
| 0.00297327 | 0.03974086 |
| 0.00298478 | 0.03985081 |
| 0.00299059 | 0.03988438 |
| 0.00300131 | 0.03998334 |
| 0.00301072 | 0.04006476 |
| 0.00301626 | 0.04009087 |
| 0.0030193 | 0.04009087 |
| 0.00304495 | 0.04038721 |
| 0.00310778 | 0.04117557 |
| 0.00311233 | 0.04119084 |
| 0.00311771 | 0.04121707 |
| 0.00312949 | 0.04132761 |
| 0.00313913 | 0.04136485 |
| 0.00313794 | 0.04136485 |
| 0.003155 | 0.04147523 |
| 0.00316348 | 0.04147523 |
| 0.00316461 | 0.04147523 |
| 0.00316344 | 0.04147523 |
| 0.00316336 | 0.04147523 |
| 0.00318273 | 0.04164147 |
| 0.00318417 | 0.04164147 |
| 0.00319379 | 0.04167736 |
| 0.00319075 | 0.04167736 |
| 0.00321199 | 0.04182885 |
| 0.0032123 | 0.04182885 |
| 0.00322589 | 0.04196078 |
| 0.00323492 | 0.04198819 |
| 0.00323469 | 0.04198819 |
| 0.0032609 | 0.04227746 |
| 0.00326418 | 0.04227746 |
| 0.00327747 | 0.0424042 |
| 0.00328971 | 0.04251723 |
| 0.00329961 | 0.04255449 |
| 0.00329615 | 0.04255449 |
| 0.00331882 | 0.04275666 |
| 0.003329 | 0.04284228 |
| 0.00333972 | 0.0429347 |
| 0.00334896 | 0.04298696 |
| 0.00335088 | 0.04298696 |

| | |
|---|---|
| 0.00336738 | 0.04309425 |
| 0.00336725 | 0.04309425 |
| 0.0033699 | 0.04309425 |
| 0.0033957 | 0.04337842 |
| 0.00342511 | 0.04370805 |
| 0.00342924 | 0.04371464 |
| 0.00344761 | 0.04390274 |
| 0.00345394 | 0.0439372 |
| 0.00346865 | 0.04407799 |
| 0.00348455 | 0.04409451 |
| 0.00347571 | 0.04409451 |
| 0.00348644 | 0.04409451 |
| 0.00348105 | 0.04409451 |
| 0.00348813 | 0.04409451 |
| 0.00350011 | 0.044108 |
| 0.00349921 | 0.044108 |
| 0.00349547 | 0.044108 |
| 0.00350745 | 0.04415456 |
| 0.00355486 | 0.0444743 |
| 0.00355254 | 0.0444743 |
| 0.003549 | 0.0444743 |
| 0.00354268 | 0.0444743 |
| 0.00354498 | 0.0444743 |
| 0.00354589 | 0.0444743 |
| 0.00356186 | 0.0445052 |
| 0.00356467 | 0.0445052 |
| 0.00358355 | 0.04469482 |
| 0.0035933 | 0.04472886 |
| 0.00359735 | 0.04472886 |
| 0.00359381 | 0.04472886 |
| 0.00362831 | 0.04506765 |
| 0.00366949 | 0.04553249 |
| 0.00367703 | 0.04557942 |
| 0.00370372 | 0.04577467 |
| 0.00369686 | 0.04577467 |
| 0.00370411 | 0.04577467 |
| 0.00371517 | 0.04582209 |
| 0.00371551 | 0.04582209 |
| 0.00373101 | 0.04596649 |
| 0.00373972 | 0.046027 |
| 0.00374941 | 0.04609947 |
| 0.00378249 | 0.04645 |
| 0.00378558 | 0.04645 |

| | |
|---|---|
| 0.0037945 | 0.04651239 |
| 0.00380854 | 0.04663726 |
| 0.00381375 | 0.04665393 |
| 0.00382101 | 0.04669572 |
| 0.00383016 | 0.04676035 |
| 0.0038533 | 0.04699551 |
| 0.00387556 | 0.04721954 |
| 0.00390598 | 0.04754241 |
| 0.00391647 | 0.04762228 |
| 0.00394849 | 0.04796342 |
| 0.00395657 | 0.04801347 |
| 0.00401725 | 0.04870113 |
| 0.00403821 | 0.04890628 |
| 0.00404404 | 0.04892805 |
| 0.00407138 | 0.04920975 |
| 0.00410061 | 0.04951367 |
| 0.00411424 | 0.04953022 |
| 0.00410995 | 0.04953022 |
| 0.00411242 | 0.04953022 |
| 0.00413816 | 0.04976877 |
| 0.00415392 | 0.04990882 |
| 0.0041786 | 0.04991971 |
| 0.00417172 | 0.04991971 |
| 0.00417953 | 0.04991971 |
| 0.00417248 | 0.04991971 |
| 0.00417635 | 0.04991971 |
| 0.0041638 | 0.04991971 |
| 0.00419282 | 0.04998135 |
| 0.004196 | 0.04998135 |
| 0.00419706 | 0.04998135 |

| ENSG ID | Gene id | baseMean | log2FoldChange | lfcSE | stat |
|---|---|---|---|---|---|
| ENSG00000225937.2 | PCA3 | 393.875735 | 2.366316797 | 0.23283949 | 10.1628671 |
| ENSG00000187210.13 | GCNT1 | 126.712631 | 2.427325537 | 0.25448337 | 9.53824817 |
| ENSG00000272234.1 | AC008945.1 | 36.7362865 | 2.918084885 | 0.32017507 | 9.11402893 |
| ENSG00000100036.12 | SLC35E4 | 129.381544 | 0.974218294 | 0.11418488 | 8.53193755 |
| ENSG00000230928.1 | AL139241.1 | 325.712485 | 3.195447641 | 0.39317911 | 8.12720598 |
| ENSG00000198865.9 | CCDC152 | 163.15812 | 3.192871467 | 0.4241143 | 7.52832782 |
| ENSG00000222047.8 | C10orf55 | 311.087166 | 2.812464746 | 0.37332912 | 7.53347274 |
| ENSG00000108641.15 | B9D1 | 27.9667317 | 1.625655522 | 0.21502969 | 7.56014458 |
| ENSG00000253194.2 | AL365275.1 | 151.501561 | 1.837820539 | 0.25530954 | 7.19840145 |
| ENSG00000122085.16 | MTERF4 | 733.512007 | 1.067736975 | 0.14859448 | 7.18557644 |
| ENSG00000170425.3 | ADORA2B | 6.64370414 | 4.248110501 | 0.60789442 | 6.98823731 |
| ENSG00000177042.14 | TMEM80 | 208.630801 | 1.364424213 | 0.19600088 | 6.9613166 |
| ENSG00000261240.1 | AC009065.6 | 281.913221 | 0.83404446 | 0.12078967 | 6.90493204 |
| ENSG00000267193.5 | AC023421.2 | 8.60236232 | 4.892348945 | 0.71665403 | 6.8266538 |
| ENSG00000188015.9 | S100A3 | 34.0883119 | 1.902198278 | 0.28138689 | 6.76008129 |
| ENSG00000233351.1 | AL356124.2 | 169.824385 | 1.305578088 | 0.19587906 | 6.66522543 |
| ENSG00000231609.6 | AC009501.1 | 425.80042 | 1.183407704 | 0.17950181 | 6.59273409 |
| ENSG00000250483.1 | PPM1AP1 | 16.8246506 | 1.681099204 | 0.25606186 | 6.56520733 |
| ENSG00000248774.1 | AC097534.1 | 105.160574 | 1.227968611 | 0.1883144 | 6.52084274 |
| ENSG00000226457.1 | RPL22P3 | 10.7194606 | 4.409609577 | 0.67747065 | 6.50893081 |
| ENSG00000097046.12 | CDC7 | 12.4267201 | 5.618466351 | 0.8928112 | 6.2930061 |
| ENSG00000115694.14 | STK25 | 82.6757211 | 0.903943451 | 0.14365787 | 6.2923352 |
| ENSG00000254866.2 | DEFB109D | 4.33101463 | 4.821516158 | 0.77250724 | 6.24138637 |
| ENSG00000231329.8 | AL031772.1 | 151.411532 | 0.838953305 | 0.13485857 | 6.22098628 |
| ENSG00000257607.1 | AC073957.2 | 25.0377757 | 2.383278437 | 0.38947532 | 6.11920269 |
| ENSG00000186063.12 | AIDA | 96.8585688 | 0.775929162 | 0.12687215 | 6.11583518 |
| ENSG00000135424.16 | ITGA7 | 8.59637489 | 2.620514325 | 0.43066211 | 6.08484996 |
| ENSG00000246763.6 | RGMB-AS1 | 605.206922 | 1.504980098 | 0.24732015 | 6.08514956 |
| ENSG00000170584.10 | NUDCD2 | 66.2909137 | 1.525440333 | 0.25173867 | 6.0596187 |
| ENSG00000257181.1 | AC025423.4 | 404.734404 | 0.935601368 | 0.15644261 | 5.98047664 |
| ENSG00000204706.14 | MAMDC2-AS: | 5.88864397 | 4.577985211 | 0.77207156 | 5.92948299 |
| ENSG00000120215.9 | MLANA | 20.5397208 | 1.661896245 | 0.28204398 | 5.89233023 |
| ENSG00000237525.6 | AC012668.3 | 15.2899794 | 1.730778022 | 0.29434061 | 5.88018761 |
| ENSG00000264785.1 | AC005722.3 | 8.84260925 | 3.884503874 | 0.66267026 | 5.86189559 |
| ENSG00000272356.1 | AL080317.3 | 245.451635 | 1.118457789 | 0.19116566 | 5.85072541 |
| ENSG00000204183.1 | GDF5OS | 60.1261561 | 3.183168746 | 0.54611262 | 5.8287771 |
| ENSG00000225521.1 | AC005237.1 | 45.6653244 | 1.22742066 | 0.21054654 | 5.82968815 |
| ENSG00000273521.1 | AL162274.1 | 15.8657355 | 2.357079987 | 0.4090272 | 5.76264857 |
| ENSG00000262211.1 | AC008914.1 | 987.695079 | 1.093736706 | 0.19057957 | 5.7390028 |
| ENSG00000244676.5 | AL109761.1 | 19.4074911 | 1.282142567 | 0.22374509 | 5.73037181 |
| ENSG00000108511.9 | HOXB6 | 47.3629917 | 3.12042574 | 0.5473553 | 5.70091442 |
| ENSG00000154723.12 | ATP5PF | 53.051444 | 0.938702749 | 0.16477416 | 5.69690506 |

| | | | | | |
|---|---|---|---|---|---|
| ENSG00000187325.4 | TAF9B | 119.632396 | 1.382921239 | 0.24311051 | 5.68844694 |
| ENSG00000106772.17 | PRUNE2 | 23.3249801 | 2.061458429 | 0.36311308 | 5.67718025 |
| ENSG00000171497.4 | PPID | 33.5031233 | 1.28525324 | 0.22701357 | 5.66156999 |
| ENSG00000183250.11 | LINC01547 | 73.5419447 | 2.769653218 | 0.49203018 | 5.62903122 |
| ENSG00000228417.1 | AL360182.2 | 21.7186648 | 2.28919922 | 0.40705979 | 5.6237419 |
| ENSG00000224597.10 | SVIL-AS1 | 224.18514 | 1.422392755 | 0.2535628 | 5.60962706 |
| ENSG00000256377.5 | AC009509.1 | 39.3493332 | 1.231337933 | 0.2228878 | 5.52447424 |
| ENSG00000255435.6 | AP001267.3 | 93.6891869 | 0.716562388 | 0.13082696 | 5.47717666 |
| ENSG00000246273.7 | SBF2-AS1 | 391.581041 | 0.897404274 | 0.16403457 | 5.47082398 |
| ENSG00000254042.1 | AC011365.1 | 1329.23059 | 1.713985613 | 0.3143547 | 5.4523938 |
| ENSG00000214413.7 | BBIP1 | 84.3132884 | 0.975649029 | 0.17915076 | 5.44596638 |
| ENSG00000285868.1 | | 12.4585781 | 2.105481102 | 0.38861715 | 5.41788007 |
| ENSG00000174804.3 | FZD4 | 396.67791 | 1.682651225 | 0.31108306 | 5.40900956 |
| ENSG00000236256.9 | DIAPH2-AS1 | 29.3786896 | 1.472148269 | 0.27277615 | 5.39690982 |
| ENSG00000255487.1 | AC087362.2 | 26.7275759 | 1.116463376 | 0.20691062 | 5.39587284 |
| ENSG00000245293.2 | AC096564.1 | 106.896233 | 1.115669931 | 0.20673683 | 5.39657084 |
| ENSG00000260618.1 | AC025917.1 | 102.75427 | 1.012077073 | 0.18755167 | 5.3962573 |
| ENSG00000163320.10 | CGGBP1 | 26.4867958 | 1.283531588 | 0.23911642 | 5.36781026 |
| ENSG00000213713.3 | PIGCP1 | 211.133126 | 0.90842391 | 0.1704861 | 5.32843401 |
| ENSG00000267002.3 | AC060780.1 | 26.4664857 | 1.056532397 | 0.19882626 | 5.31384745 |
| ENSG00000237036.4 | ZEB1-AS1 | 21.9071036 | 2.336799259 | 0.44216213 | 5.28493761 |
| ENSG00000267128.1 | RNF157-AS1 | 12.9265815 | 3.24934019 | 0.61771687 | 5.26024196 |
| ENSG00000283703.2 | VSIG10L2 | 207.182575 | 2.437705059 | 0.46352293 | 5.25908193 |
| ENSG00000284830.1 | AL049557.1 | 6.69344211 | 2.61935682 | 0.50382524 | 5.19893926 |
| ENSG00000115457.9 | IGFBP2 | 4.89025312 | 4.612952313 | 0.89250238 | 5.16856022 |
| ENSG00000279658.1 | AC110491.2 | 80.4207647 | 1.060407157 | 0.20616647 | 5.14345098 |
| ENSG00000214282.3 | KRT8P14 | 14.7704046 | 1.685430152 | 0.32823001 | 5.13490576 |
| ENSG00000251239.1 | AC004590.1 | 2.45154238 | 4.128591493 | 0.80446614 | 5.13208858 |
| ENSG00000254913.1 | AC239802.1 | 41.6809147 | 1.68283793 | 0.32958282 | 5.10596381 |
| ENSG00000249835.2 | VCAN-AS1 | 2937.12386 | 1.067556382 | 0.21115361 | 5.05582812 |
| ENSG00000230046.1 | BIRC6-AS1 | 36.3482891 | 0.974266414 | 0.19276791 | 5.05409028 |
| ENSG00000281021.1 | AL078581.3 | 295.136189 | 0.534264532 | 0.10614401 | 5.03339325 |
| ENSG00000170909.13 | OSCAR | 14.6539125 | 2.552090235 | 0.50801922 | 5.02360957 |
| ENSG00000262655.3 | SPON1 | 267.855716 | 2.051689949 | 0.41245379 | 4.9743511 |
| ENSG00000165443.11 | PHYHIPL | 13.2036022 | 2.550110345 | 0.51320209 | 4.96901782 |
| ENSG00000125779.22 | PANK2 | 125.144694 | 1.096176355 | 0.22200151 | 4.93769777 |
| ENSG00000278156.1 | TSC22D1-AS1 | 265.536969 | 1.083584332 | 0.21953541 | 4.93580662 |
| ENSG00000246323.2 | AC113382.1 | 97.2522557 | 0.718076205 | 0.14588892 | 4.92207488 |
| ENSG00000267765.1 | AC100793.3 | 237.568604 | 0.96665373 | 0.19684804 | 4.91065975 |
| ENSG00000234175.1 | AL355355.2 | 12.9467369 | 1.316224888 | 0.26838923 | 4.90416428 |
| ENSG00000259564.2 | AC009554.1 | 59.4523258 | 0.985170608 | 0.20290069 | 4.85543241 |
| ENSG00000254192.1 | AC011365.2 | 1379.89957 | 1.632226903 | 0.33703348 | 4.84292208 |
| ENSG00000255817.1 | AC025576.2 | 63.6206737 | 1.121524612 | 0.23219583 | 4.83008075 |

| | | | | | |
|---|---|---|---|---|---|
| ENSG00000126602.10 | TRAP1 | 64.6996374 | 1.161197364 | 0.24051745 | 4.82791318 |
| ENSG00000267302.5 | RNFT1-DT | 13.7227827 | 1.325953515 | 0.27531613 | 4.81611272 |
| ENSG00000262728.5 | AC123768.3 | 17.6984827 | 1.320847705 | 0.27556343 | 4.79326194 |
| ENSG00000136536.14 | 7-Mar | 274.887738 | 0.65741479 | 0.13775269 | 4.77242807 |
| ENSG00000272476.1 | AL024507.2 | 182.031942 | 0.628577224 | 0.13173155 | 4.77165294 |
| ENSG00000225822.4 | UBXN7-AS1 | 22.547778 | 1.038523727 | 0.21810277 | 4.76162565 |
| ENSG00000272320.1 | AL445309.1 | 14.5682239 | 1.923281123 | 0.40535927 | 4.7446334 |
| ENSG00000170439.6 | METTL7B | 5.10016373 | 3.054480586 | 0.64476316 | 4.73736833 |
| ENSG00000259125.1 | LRP1-AS | 1005.60143 | 0.708569597 | 0.14959303 | 4.73664837 |
| ENSG00000066923.17 | STAG3 | 200.79437 | 1.018666249 | 0.21526348 | 4.73218329 |
| ENSG00000135678.11 | CPM | 163.244071 | 0.924493236 | 0.19542326 | 4.73072272 |
| ENSG00000283529.1 | AL035685.1 | 14.4110891 | 2.584938388 | 0.54702678 | 4.72543296 |
| ENSG00000279232.2 | AC008522.1 | 9246.12344 | 1.166269951 | 0.24675543 | 4.72642064 |
| ENSG00000273489.1 | AC008264.2 | 250.397008 | 0.955477143 | 0.20315265 | 4.70324716 |
| ENSG00000236051.7 | MYCBP2-AS1 | 360.080342 | 0.693423449 | 0.14818169 | 4.67954879 |
| ENSG00000124571.17 | XPO5 | 23.0466067 | 1.364008204 | 0.29257771 | 4.66203737 |
| ENSG00000256040.2 | PAPPA-AS1 | 1383.94797 | 1.278299772 | 0.27444829 | 4.6577072 |
| ENSG00000002745.12 | WNT16 | 13.0577513 | 3.079837688 | 0.66193847 | 4.65275524 |
| ENSG00000251260.1 | WDFY3-AS1 | 130.006421 | 0.943623384 | 0.20356776 | 4.63542637 |
| ENSG00000197415.11 | VEPH1 | 252.728327 | 1.750126757 | 0.37862234 | 4.62235476 |
| ENSG00000279569.1 | AC020763.4 | 4401.17853 | 1.019934497 | 0.22078037 | 4.61967933 |
| ENSG00000258520.1 | AL359317.1 | 20.0121328 | 3.177060732 | 0.68818246 | 4.61659646 |
| ENSG00000283375.1 | AC087521.4 | 394.891976 | 0.609797488 | 0.13292859 | 4.58740648 |
| ENSG00000149294.16 | NCAM1 | 6.56165092 | 2.844508486 | 0.62077801 | 4.58216693 |
| ENSG00000172345.13 | STARD5 | 12.3054035 | 1.609880825 | 0.35123911 | 4.58343271 |
| ENSG00000253992.1 | AC103726.2 | 2.27235554 | 4.105007642 | 0.89874308 | 4.56749846 |
| ENSG00000279122.1 | AC020763.2 | 2.36244258 | 4.313899134 | 0.94489966 | 4.56545737 |
| ENSG00000226605.1 | AC092567.1 | 167.1919 | 1.20235255 | 0.26416402 | 4.55153783 |
| ENSG00000233885.7 | YEATS2-AS1 | 268.755235 | 0.738308813 | 0.16234392 | 4.54780689 |
| ENSG00000232832.1 | LMLN-AS1 | 44.399982 | 0.804843181 | 0.17724274 | 4.54090912 |
| ENSG00000127241.16 | MASP1 | 26.4943694 | 1.834290811 | 0.40464906 | 4.53304104 |
| ENSG00000233706.1 | AL353689.2 | 251.80299 | 0.67085911 | 0.14804664 | 4.53140381 |
| ENSG00000149262.16 | INTS4 | 8.8123716 | 1.583590237 | 0.35018428 | 4.52216258 |
| ENSG00000271647.1 | KRT8P47 | 6.00305841 | 3.155564742 | 0.70071311 | 4.50336194 |
| ENSG00000144741.17 | SLC25A26 | 194.954943 | 1.926259216 | 0.42761737 | 4.50463275 |
| ENSG00000275327.1 | AL354950.2 | 57.789524 | 2.393533336 | 0.5335191 | 4.48631234 |
| ENSG00000270751.1 | FBXW7-AS1 | 47.7328063 | 0.880393052 | 0.19710907 | 4.46652734 |
| ENSG00000253304.1 | TMEM200B | 32.2270089 | 1.055292018 | 0.23703464 | 4.45205814 |
| ENSG00000198732.10 | SMOC1 | 2.6096443 | 4.223689257 | 0.94956397 | 4.44803024 |
| ENSG00000138443.15 | ABI2 | 3199.76902 | 0.721860997 | 0.16231059 | 4.44740531 |
| ENSG00000255629.1 | AC025576.1 | 162.061312 | 0.783968014 | 0.17641109 | 4.44398381 |
| ENSG00000234616.8 | JRK | 14.1095475 | 2.071310251 | 0.46687938 | 4.43649979 |
| ENSG00000105974.11 | CAV1 | 23.5808758 | 1.572219634 | 0.35497196 | 4.42913759 |

| | | | | | |
|---|---|---|---|---|---|
| ENSG00000225921.6 | NOL7 | 297.817327 | 0.590930605 | 0.1336296 | 4.42215335 |
| ENSG00000259188.5 | AC025040.1 | 1.44788727 | 4.271677463 | 0.96935609 | 4.40671647 |
| ENSG00000249252.5 | AC098829.1 | 52.9915348 | 0.986671885 | 0.22451473 | 4.39468657 |
| ENSG00000135250.16 | SRPK2 | 87.4370064 | 0.803287041 | 0.18296447 | 4.39039906 |
| ENSG00000175029.16 | CTBP2 | 519.442032 | 0.527789198 | 0.1202397 | 4.38947527 |
| ENSG00000138131.3 | LOXL4 | 10.2998188 | 2.476151807 | 0.5665329 | 4.37071137 |
| ENSG00000285608.1 | | 637.536728 | 0.812281582 | 0.18583279 | 4.37103485 |
| ENSG00000258959.2 | AL118558.1 | 1750.54668 | 0.566859411 | 0.12974908 | 4.36888976 |
| ENSG00000166323.12 | C11orf65 | 62.4235384 | 0.946191124 | 0.21690818 | 4.36217352 |
| ENSG00000285051.1 | AC026316.4 | 14.1507494 | 4.536727749 | 1.04297324 | 4.34980265 |
| ENSG00000261716.2 | AC239868.1 | 45.3780979 | 0.81679215 | 0.18784632 | 4.34819345 |
| ENSG00000263873.1 | AP003396.5 | 7602.23615 | 1.164026537 | 0.26793658 | 4.34441063 |
| ENSG00000138079.13 | SLC3A1 | 337.119506 | 0.831637429 | 0.19166569 | 4.33899996 |
| ENSG00000237768.2 | AL731563.2 | 406.364759 | 0.88521499 | 0.20444077 | 4.32993379 |
| ENSG00000264443.1 | AL445686.2 | 196.899168 | 0.765726075 | 0.17715883 | 4.32225737 |
| ENSG00000160255.17 | ITGB2 | 12.1125134 | 2.001967473 | 0.46442035 | 4.31067987 |
| ENSG00000236107.8 | AC010127.1 | 52.6778976 | 1.774805608 | 0.41279179 | 4.29951771 |
| ENSG00000228830.1 | AL160408.2 | 722.256068 | 0.624506902 | 0.14543561 | 4.294044 |
| ENSG00000177082.12 | WDR73 | 108.581237 | 0.897850422 | 0.20973918 | 4.280795 |
| ENSG00000267023.5 | LRRC37A16P | 665.074821 | 0.571426394 | 0.13390549 | 4.2673859 |
| ENSG00000174177.12 | CTU2 | 80.1688928 | 0.542209205 | 0.12708411 | 4.26653805 |
| ENSG00000122729.18 | ACO1 | 6.94673458 | 1.605974176 | 0.37681457 | 4.26197469 |
| ENSG00000198791.11 | CNOT7 | 48.8616127 | 1.063640553 | 0.2502435 | 4.25042232 |
| ENSG00000281371.1 | INE2 | 72.8307381 | 0.945340263 | 0.22280656 | 4.24287438 |
| ENSG00000146707.14 | POMZP3 | 26.3297387 | 1.685300683 | 0.39740912 | 4.24071966 |
| ENSG00000260285.1 | AL133367.1 | 22.563613 | 1.610339609 | 0.3800283 | 4.23742026 |
| ENSG00000226430.6 | USP17L7 | 2.40824434 | 4.878257016 | 1.15166117 | 4.23584398 |
| ENSG00000151748.14 | SAV1 | 35.9779319 | 1.687647409 | 0.39944633 | 4.22496663 |
| ENSG00000226548.1 | AC016722.1 | 7.72567839 | 2.492198135 | 0.59081037 | 4.21827083 |
| ENSG00000269915.1 | AP006621.4 | 112.302869 | 0.549773153 | 0.13075771 | 4.20451816 |
| ENSG00000263235.1 | AC006111.2 | 5.94908183 | 1.742803793 | 0.41521433 | 4.19735941 |
| ENSG00000125630.15 | POLR1B | 3.50924945 | 2.601051813 | 0.62045351 | 4.19217843 |
| ENSG00000258317.1 | AC034102.6 | 437.123416 | 0.547554281 | 0.13070058 | 4.18937909 |
| ENSG00000078795.16 | PKD2L2 | 288.901733 | 0.718226586 | 0.17150572 | 4.18777045 |
| ENSG00000235512.1 | TAB3-AS2 | 92.1697258 | 0.887541379 | 0.21257563 | 4.17517931 |
| ENSG00000249889.1 | ALG1L11P | 4.96638794 | 2.453315288 | 0.58786691 | 4.17324952 |
| ENSG00000182934.11 | SRPRA | 46.6877239 | 0.819293707 | 0.1967212 | 4.16474529 |
| ENSG00000144567.10 | RETREG2 | 172.162853 | 0.634530697 | 0.15246882 | 4.16170796 |
| ENSG00000227487.3 | NCAM1-AS1 | 51.6974165 | 3.014313697 | 0.72630952 | 4.15017786 |
| ENSG00000165801.9 | ARHGEF40 | 57.1511373 | 0.755860831 | 0.18218864 | 4.14878133 |
| ENSG00000237125.9 | HAND2-AS1 | 88.3561043 | 1.615615089 | 0.38954102 | 4.14748382 |
| ENSG00000232675.7 | AL121830.1 | 2.58009334 | 4.972630501 | 1.20358424 | 4.13151846 |
| ENSG00000090266.12 | NDUFB2 | 101.487946 | 0.580572596 | 0.14079453 | 4.123545 |

| | | | | | |
|---|---|---|---|---|---|
| ENSG00000285582.1 | | 1.80429366 | 4.097038975 | 0.99516007 | 4.11696481 |
| ENSG00000247416.3 | AP000802.1 | 10.8862528 | 2.930921384 | 0.71426085 | 4.10343276 |
| ENSG00000134480.14 | CCNH | 418.451912 | 0.739144865 | 0.18022468 | 4.10124118 |
| ENSG00000268364.1 | SMC5-AS1 | 4.33881913 | 3.000834534 | 0.73228694 | 4.09789437 |
| ENSG00000251630.1 | AC119751.6 | 4.31832856 | 3.916146194 | 0.96138293 | 4.07345095 |
| ENSG00000197815.4 | AC122129.1 | 52.2042081 | 0.709215287 | 0.17420701 | 4.07110639 |
| ENSG00000130956.13 | HABP4 | 54.1196266 | 0.923346126 | 0.22726563 | 4.06284982 |
| ENSG00000269349.1 | AC022150.3 | 26.6802882 | 1.336306327 | 0.3290455 | 4.06115966 |
| ENSG00000234055.1 | AL158151.2 | 120.366431 | 0.660932545 | 0.1629755 | 4.05541044 |
| ENSG00000255886.1 | AC020611.2 | 81.0950292 | 1.06634479 | 0.26442592 | 4.0326788 |
| ENSG00000164708.5 | PGAM2 | 35.3383501 | 0.908663467 | 0.22581111 | 4.02399805 |
| ENSG00000249087.6 | ZNF436-AS1 | 25.8963689 | 1.329578929 | 0.33140648 | 4.01192802 |
| ENSG00000152700.13 | SAR1B | 129.291551 | 0.49889257 | 0.12447731 | 4.00789975 |
| ENSG00000157570.11 | TSPAN18 | 53.1115515 | 2.051578683 | 0.51375127 | 3.99333063 |
| ENSG00000151466.11 | SCLT1 | 74.3864219 | 1.084269825 | 0.27170023 | 3.99068416 |
| ENSG00000239718.1 | HLTF-AS1 | 28.9980951 | 0.841717648 | 0.21110767 | 3.98714861 |
| ENSG00000179454.13 | KLHL28 | 34.5578727 | 0.715023415 | 0.17988519 | 3.97488762 |
| ENSG00000176102.12 | CSTF3 | 73.7678585 | 0.687809782 | 0.17322362 | 3.97064652 |
| ENSG00000266043.1 | MIR3649 | 7.0621696 | 2.028386275 | 0.51135333 | 3.96670202 |
| ENSG00000170854.17 | RIOX2 | 236.151439 | 0.817053836 | 0.20617257 | 3.96296099 |
| ENSG00000145362.17 | ANK2 | 8.89246812 | 2.462118429 | 0.62190292 | 3.95900767 |
| ENSG00000186812.12 | ZNF397 | 56.7220493 | 0.649483776 | 0.16414786 | 3.95669953 |
| ENSG00000267379.1 | AC008569.1 | 356.766339 | 0.748502591 | 0.18940456 | 3.95187198 |
| ENSG00000257769.1 | AC026401.1 | 51.2067448 | 0.812290033 | 0.20601221 | 3.94292184 |
| ENSG00000101333.16 | PLCB4 | 3.79910426 | 2.039468456 | 0.51788857 | 3.93804491 |
| ENSG00000133110.14 | POSTN | 15.738218 | 2.152890545 | 0.55089191 | 3.90800898 |
| ENSG00000130024.14 | PHF10 | 225.505152 | 0.601630588 | 0.15398617 | 3.90704311 |
| ENSG00000184451.5 | CCR10 | 98.53796 | 0.704218133 | 0.18055711 | 3.90025147 |
| ENSG00000237278.2 | RLIMP2 | 42.0007676 | 0.788653649 | 0.2026401 | 3.89189327 |
| ENSG00000254943.1 | AP003501.2 | 3.00657334 | 2.387916109 | 0.61423813 | 3.88760644 |
| ENSG00000129048.6 | ACKR4 | 78.043777 | 0.884771709 | 0.2277632 | 3.88461229 |
| ENSG00000146243.13 | IRAK1BP1 | 90.1262006 | 0.817342168 | 0.21047186 | 3.88337983 |
| ENSG00000136854.20 | STXBP1 | 35.1783469 | 0.857740154 | 0.22093693 | 3.88228512 |
| ENSG00000104375.16 | STK3 | 8.92112032 | 1.749649814 | 0.45096637 | 3.87977895 |
| ENSG00000224789.1 | AC012363.1 | 39.0342412 | 0.885276582 | 0.22832064 | 3.87733921 |
| ENSG00000001460.17 | STPG1 | 134.168826 | 0.575047195 | 0.14858404 | 3.87018147 |
| ENSG00000246250.2 | AC087521.2 | 205.89246 | 0.600605826 | 0.15533838 | 3.86643559 |
| ENSG00000152661.8 | GJA1 | 2.9139352 | 2.65236251 | 0.6862602 | 3.86495168 |
| ENSG00000131737.5 | KRT34 | 5.13168039 | 3.509847554 | 0.91069556 | 3.85402952 |
| ENSG00000283208.1 | AC001226.2 | 132.217475 | 0.836551317 | 0.21797182 | 3.83788742 |
| ENSG00000250909.1 | AC138956.1 | 414.146148 | 0.597315931 | 0.15584803 | 3.83268186 |
| ENSG00000259353.1 | AC090515.4 | 33.3045126 | 1.094439064 | 0.28592656 | 3.82769285 |
| ENSG00000225610.1 | AC007679.1 | 37.7147434 | 0.908889889 | 0.23816387 | 3.81623742 |

| | | | | | |
|---|---|---|---|---|---|
| ENSG00000255201.1 | AC087623.2 | 247.163117 | 0.522814333 | 0.13739958 | 3.80506497 |
| ENSG00000123243.14 | ITIH5 | 2.38192807 | 4.343913389 | 1.14401137 | 3.79708935 |
| ENSG00000227617.8 | CERS6-AS1 | 384.701616 | 0.642486825 | 0.16966203 | 3.78686279 |
| ENSG00000235426.2 | AL133481.1 | 237.750208 | 0.544069721 | 0.14377481 | 3.78417966 |
| ENSG00000104450.12 | SPAG1 | 128.049699 | 0.525364055 | 0.13885118 | 3.78364843 |
| ENSG00000155657.26 | TTN | 510.441916 | 0.777104201 | 0.20555764 | 3.78046863 |
| ENSG00000082805.19 | ERC1 | 8.33082926 | 1.476823024 | 0.39226433 | 3.76486701 |
| ENSG00000164985.14 | PSIP1 | 93.8920765 | 0.980819878 | 0.26080114 | 3.76079591 |
| ENSG00000266049.1 | AP001011.1 | 36.3892063 | 0.910866439 | 0.24272502 | 3.75266806 |
| ENSG00000112237.12 | CCNC | 23.1815464 | 0.939774431 | 0.25133987 | 3.73905826 |
| ENSG00000259113.1 | AL118556.1 | 226.413558 | 0.648954938 | 0.17365887 | 3.73695237 |
| ENSG00000263177.1 | MTND1P8 | 2.65794706 | 2.93014127 | 0.78483592 | 3.73344438 |
| ENSG00000151632.17 | AKR1C2 | 18.0143348 | 1.070232666 | 0.28749112 | 3.72266333 |
| ENSG00000160226.15 | C21orf2 | 89.5941409 | 0.773848345 | 0.20779972 | 3.72401051 |
| ENSG00000260246.1 | AC000032.1 | 84.8504483 | 2.414983136 | 0.65100259 | 3.70963678 |
| ENSG00000227906.7 | SNAP25-AS1 | 12.6482692 | 1.748977675 | 0.47143416 | 3.7099087 |
| ENSG00000237298.9 | TTN-AS1 | 7.15545521 | 2.461991048 | 0.66471606 | 3.70382362 |
| ENSG00000220323.4 | HIST2H2BD | 28.4258567 | 0.81154636 | 0.21925846 | 3.70132297 |
| ENSG00000258777.1 | HIF1A-AS1 | 1039.10867 | 0.789989429 | 0.21348163 | 3.70050311 |
| ENSG00000204291.10 | COL15A1 | 7.07777456 | 1.726082459 | 0.467184 | 3.69465237 |
| ENSG00000246283.2 | AC090510.1 | 140.52504 | 0.60084523 | 0.16262398 | 3.69469022 |
| ENSG00000115414.18 | FN1 | 1293.48701 | 0.779804321 | 0.21150917 | 3.68685826 |
| ENSG00000177640.15 | CASC2 | 50.1757202 | 0.770316293 | 0.20906346 | 3.68460504 |
| ENSG00000255920.2 | CCND2-AS1 | 6.9646242 | 3.413550078 | 0.92697352 | 3.68246774 |
| ENSG00000138496.16 | PARP9 | 20.0164822 | 1.103093403 | 0.30004476 | 3.67642952 |
| ENSG00000177971.8 | IMP3 | 218.233175 | 0.558227843 | 0.15187375 | 3.67560444 |
| ENSG00000216331.2 | HIST1H1PS1 | 10.3048292 | 2.628235446 | 0.7159167 | 3.67114702 |
| ENSG00000273132.1 | AL355312.3 | 86.8999983 | 0.601284006 | 0.16398093 | 3.66679219 |
| ENSG00000241202.1 | ZIC4-AS1 | 37.731654 | 1.862140887 | 0.50814745 | 3.66456799 |
| ENSG00000120075.5 | HOXB5 | 10.5452003 | 2.716536912 | 0.74209242 | 3.660645 |
| ENSG00000272851.1 | AC096772.1 | 226.542311 | 0.767682316 | 0.20971755 | 3.66055351 |
| ENSG00000128346.10 | C22orf23 | 363.178252 | 0.545848491 | 0.14921577 | 3.6581154 |
| ENSG00000072832.14 | CRMP1 | 735.228743 | 0.420538204 | 0.11491947 | 3.65941637 |
| ENSG00000265298.1 | AC132812.1 | 90.0037155 | 0.841097013 | 0.23055038 | 3.64821358 |
| ENSG00000257761.1 | AC078860.1 | 23.4943456 | 2.934881326 | 0.80559571 | 3.64311936 |
| ENSG00000268475.1 | AC011462.2 | 23.5576754 | 0.934561545 | 0.25681506 | 3.63904492 |
| ENSG00000214128.10 | TMEM213 | 59.2994995 | 0.779060421 | 0.21422655 | 3.63661942 |
| ENSG00000168763.15 | CNNM3 | 26.3635454 | 0.809562947 | 0.22286365 | 3.6325481 |
| ENSG00000268006.1 | PTOV1-AS1 | 270.258896 | 0.423635982 | 0.11678692 | 3.62742673 |
| ENSG00000255542.1 | AC090625.2 | 3.66385764 | 1.859919012 | 0.51339806 | 3.62276207 |
| ENSG00000164117.13 | FBXO8 | 15.6043368 | 0.891148903 | 0.24600077 | 3.6225452 |
| ENSG00000108352.12 | RAPGEFL1 | 2.95402166 | 2.398832914 | 0.66356591 | 3.61506349 |
| ENSG00000168273.7 | SMIM4 | 122.241712 | 0.66520145 | 0.18406979 | 3.61385451 |

| | | | | | |
|---|---|---|---|---|---|
| ENSG00000259052.1 | AL157871.6 | 70.4527424 | 0.635982432 | 0.17631447 | 3.60709152 |
| ENSG00000129636.12 | ITFG1 | 51.9198344 | 0.771702721 | 0.2148139 | 3.59242457 |
| ENSG00000229893.2 | AC005091.1 | 153.276273 | 0.563661692 | 0.15692252 | 3.59197455 |
| ENSG00000270835.2 | AP001425.1 | 1.88781774 | 3.724865867 | 1.03809354 | 3.58817941 |
| ENSG00000256699.1 | TMEM132D-/ | 13.6288414 | 3.486272207 | 0.97317421 | 3.58237217 |
| ENSG00000256001.1 | AC079949.1 | 9.62591909 | 3.837254539 | 1.07201145 | 3.57949025 |
| ENSG00000258425.1 | AC013451.1 | 1614.01658 | 0.734817211 | 0.20547365 | 3.57621139 |
| ENSG00000188992.11 | LIPI | 1.33311306 | 3.918936133 | 1.09644691 | 3.57421422 |
| ENSG00000163516.13 | ANKZF1 | 9.25444452 | 1.08029729 | 0.3023293 | 3.57324708 |
| ENSG00000272325.1 | NUDT3 | 126.287614 | 0.77722472 | 0.21769069 | 3.57031682 |
| ENSG00000213904.8 | LIPE-AS1 | 85.095789 | 0.788503984 | 0.22093921 | 3.56887303 |
| ENSG00000181800.5 | CELF2-AS1 | 6.41591292 | 2.565508466 | 0.71898572 | 3.56823288 |
| ENSG00000099219.13 | ERMP1 | 137.354923 | 0.491578021 | 0.13799682 | 3.56224165 |
| ENSG00000225670.4 | CADM3-AS1 | 2.61655512 | 4.405425747 | 1.24029318 | 3.55192289 |
| ENSG00000258034.1 | AC012157.1 | 162.134413 | 0.496428215 | 0.14010523 | 3.54325263 |
| ENSG00000166888.11 | STAT6 | 525.797111 | 0.50250825 | 0.14198237 | 3.53922984 |
| ENSG00000124181.14 | PLCG1 | 201.398918 | 0.466908708 | 0.13195065 | 3.53851021 |
| ENSG00000269926.1 | DDIT4-AS1 | 473.50411 | 1.095971437 | 0.31019073 | 3.53321792 |
| ENSG00000263326.1 | AC133552.4 | 2.4127407 | 2.154609999 | 0.61035981 | 3.53006535 |
| ENSG00000285921.1 | | 98.1717846 | 0.495604673 | 0.14043517 | 3.52906373 |
| ENSG00000111052.7 | LIN7A | 3.64470379 | 1.906092187 | 0.54044257 | 3.52690975 |
| ENSG00000259589.2 | AC073167.1 | 185.705041 | 0.66416968 | 0.18834714 | 3.52630611 |
| ENSG00000201778.1 | RF00019 | 23.7037282 | 1.110361422 | 0.31625059 | 3.51101767 |
| ENSG00000164105.3 | SAP30 | 3.90230557 | 2.635118047 | 0.75284761 | 3.5002011 |
| ENSG00000285698.1 | | 5.28967822 | 2.102559791 | 0.60077445 | 3.49974903 |
| ENSG00000269313.5 | MAGIX | 2.30763061 | 2.808653181 | 0.80278068 | 3.49865569 |
| ENSG00000254923.1 | AC130366.1 | 2.00498801 | 3.769504221 | 1.07792978 | 3.49698496 |
| ENSG00000078674.17 | PCM1 | 2.56892616 | 2.792349857 | 0.80010414 | 3.48998302 |
| ENSG00000258114.1 | AC005871.2 | 390.546683 | 1.243732105 | 0.3577467 | 3.47657189 |
| ENSG00000248464.1 | FGF10-AS1 | 2.21557965 | 3.626854341 | 1.04355301 | 3.47548644 |
| ENSG00000170379.20 | TCAF2 | 2.93204755 | 2.978872545 | 0.85701956 | 3.4758513 |
| ENSG00000280339.1 | AP001528.3 | 3.36249868 | 3.284231179 | 0.94586055 | 3.472215 |
| ENSG00000000460.16 | C1orf112 | 65.0634509 | 0.5612332 | 0.16204232 | 3.46349761 |
| ENSG00000204866.8 | IGFL2 | 4.71900634 | 2.380820627 | 0.68854317 | 3.45776523 |
| ENSG00000152455.15 | SUV39H2 | 31.5644571 | 0.977450802 | 0.28254784 | 3.45941702 |
| ENSG00000050438.16 | SLC4A8 | 31.8880905 | 0.866749848 | 0.25136578 | 3.44816166 |
| ENSG00000271976.1 | AC012467.2 | 4.26497402 | 2.510643142 | 0.72837268 | 3.4469211 |
| ENSG00000145216.15 | FIP1L1 | 50.6282715 | 0.927017224 | 0.26925928 | 3.44284229 |
| ENSG00000177106.15 | EPS8L2 | 47.1611072 | 0.642441866 | 0.18665531 | 3.44186226 |
| ENSG00000125631.7 | HTR5BP | 69.2230718 | 0.584860084 | 0.17024655 | 3.43537113 |
| ENSG00000186814.13 | ZSCAN30 | 35.6560854 | 1.045442465 | 0.30439358 | 3.43450887 |
| ENSG00000222043.2 | AC079305.1 | 98.7735837 | 0.433558512 | 0.12627609 | 3.43341729 |
| ENSG00000159917.16 | ZNF235 | 65.9388484 | 0.741900483 | 0.21638993 | 3.42853512 |

| | | | | | |
|---|---|---|---|---|---|
| ENSG00000247903.1 | AC024896.1 | 83.9457778 | 0.735369683 | 0.21476746 | 3.42402748 |
| ENSG00000232878.3 | DPYD-AS1 | 90.0935878 | 0.92648136 | 0.27066589 | 3.42297051 |
| ENSG00000272918.1 | AC005070.3 | 62.2355688 | 0.841602674 | 0.24583683 | 3.42341984 |
| ENSG00000227039.6 | ITGB2-AS1 | 19.8216278 | 1.596467263 | 0.46685635 | 3.41961137 |
| ENSG00000180822.11 | PSMG4 | 272.823466 | 1.18909132 | 0.34814504 | 3.41550552 |
| ENSG00000260017.1 | AC138811.1 | 119.357659 | 0.582055063 | 0.17048068 | 3.41419954 |
| ENSG00000106034.17 | CPED1 | 2.88924598 | 2.51326392 | 0.73686172 | 3.41076737 |
| ENSG00000143341.11 | HMCN1 | 12.0486306 | 1.889842953 | 0.55456869 | 3.40777074 |
| ENSG00000267265.5 | AC011476.3 | 17.3348422 | 0.88800275 | 0.26076587 | 3.40536419 |
| ENSG00000237133.1 | AC020594.1 | 6.15017019 | 1.565913225 | 0.4603701 | 3.40142252 |
| ENSG00000103023.11 | PRSS54 | 28.8252378 | 1.009653924 | 0.29690565 | 3.40058846 |
| ENSG00000239388.8 | ASB14 | 122.295469 | 0.742714687 | 0.21837015 | 3.40117317 |
| ENSG00000237975.6 | FLG-AS1 | 34.0825002 | 1.898984181 | 0.55880956 | 3.39826716 |
| ENSG00000264859.5 | DSG2-AS1 | 4.91573647 | 3.591546809 | 1.05905113 | 3.39128744 |
| ENSG00000227689.1 | SRP68P2 | 2.86448884 | 3.386253618 | 0.99862403 | 3.39091941 |
| ENSG00000241560.5 | ZBTB20-AS1 | 11.5367288 | 1.254018208 | 0.36999135 | 3.38931763 |
| ENSG00000110315.6 | RNF141 | 22.0221146 | 1.50377363 | 0.4441813 | 3.3854951 |
| ENSG00000246859.2 | STARD4-AS1 | 2011.57718 | 1.171467332 | 0.34622137 | 3.3835789 |
| ENSG00000240225.10 | ZNF542P | 7.10432858 | 1.176562024 | 0.34803387 | 3.38059631 |
| ENSG00000275005.1 | AL354950.1 | 2.87242421 | 3.69103241 | 1.09251407 | 3.37847585 |
| ENSG00000065320.8 | NTN1 | 4.37757015 | 1.759823003 | 0.52086106 | 3.37868031 |
| ENSG00000188343.12 | FAM92A | 99.1537333 | 0.687198205 | 0.20354736 | 3.37610969 |
| ENSG00000263797.1 | AP005210.1 | 2.18902624 | 2.607692534 | 0.77252548 | 3.37554241 |
| ENSG00000259863.1 | SH3RF3-AS1 | 25.6187425 | 0.770345568 | 0.22835433 | 3.37346593 |
| ENSG00000229847.8 | EMX2OS | 27.711872 | 1.051463531 | 0.31180227 | 3.37221257 |
| ENSG00000164761.8 | TNFRSF11B | 7.22511942 | 1.873169468 | 0.55569235 | 3.37087501 |
| ENSG00000233308.1 | OSTN-AS1 | 1.3347601 | 3.985738647 | 1.18300134 | 3.36917509 |
| ENSG00000185013.16 | NT5C1B | 5.03213463 | 1.509268626 | 0.44808416 | 3.36827043 |
| ENSG00000169962.4 | TAS1R3 | 23.040814 | 0.95857293 | 0.2851463 | 3.36168808 |
| ENSG00000258757.1 | AL133453.1 | 273.537456 | 0.797914143 | 0.23747663 | 3.35996905 |
| ENSG00000082146.12 | STRADB | 170.556724 | 0.685362683 | 0.20395247 | 3.36040385 |
| ENSG00000083814.13 | ZNF671 | 2.23357879 | 2.536819781 | 0.75624815 | 3.35448064 |
| ENSG00000146374.13 | RSPO3 | 1.86209572 | 3.174505547 | 0.94667635 | 3.35331661 |
| ENSG00000277959.1 | AL162274.2 | 5.08355299 | 1.770263782 | 0.52861412 | 3.34887724 |
| ENSG00000185869.14 | ZNF829 | 3.38585739 | 2.130987819 | 0.63701845 | 3.34525293 |
| ENSG00000168350.7 | DEGS2 | 105.215863 | 0.769351363 | 0.22996693 | 3.34548694 |
| ENSG00000258843.1 | AL133485.1 | 44.3809263 | 0.695040978 | 0.2078938 | 3.34325014 |
| ENSG00000203722.7 | RAET1G | 5.37167166 | 2.249796862 | 0.67326036 | 3.34164462 |
| ENSG00000085998.13 | POMGNT1 | 38.360807 | 0.745057224 | 0.22306752 | 3.34005253 |
| ENSG00000261093.1 | AC141586.3 | 413.432696 | 0.323095209 | 0.096782 | 3.33838103 |
| ENSG00000182093.15 | WRB | 3.46939829 | 1.671726187 | 0.50091892 | 3.33731891 |
| ENSG00000175470.19 | PPP2R2D | 4.00085666 | 1.894324715 | 0.56779439 | 3.33628645 |
| ENSG00000280515.1 | SALRNA2 | 1.94363924 | 2.682591019 | 0.80584671 | 3.3289098 |

| | | | | | |
|---|---|---|---|---|---|
| ENSG00000230140.5 | AC016738.2 | 38.7002049 | 0.818968643 | 0.24615397 | 3.32705849 |
| ENSG00000227543.4 | SPAG5-AS1 | 1190.37672 | 0.430668236 | 0.12948887 | 3.3259092 |
| ENSG00000261646.1 | AC093849.1 | 1.76474621 | 3.330122231 | 1.0036879 | 3.31788621 |
| ENSG00000274523.4 | RCC1L | 91.9874786 | 0.677402184 | 0.20476282 | 3.30822842 |
| ENSG00000257831.1 | AL136418.1 | 140.335644 | 0.584788276 | 0.17717498 | 3.30062564 |
| ENSG00000236276.1 | NDP-AS1 | 4.43335581 | 3.015462964 | 0.91449634 | 3.297403 |
| ENSG00000256092.2 | AC137767.1 | 24.1147924 | 0.800617942 | 0.24300147 | 3.29470405 |
| ENSG00000167984.17 | NLRC3 | 21.0908323 | 0.69116517 | 0.20974183 | 3.29531391 |
| ENSG00000114790.12 | ARHGEF26 | 1.90051573 | 2.377927658 | 0.72227501 | 3.29227459 |
| ENSG00000267192.1 | AC006116.4 | 2.25354724 | 2.499959831 | 0.75982888 | 3.29016163 |
| ENSG00000223923.1 | AC010136.1 | 95.1007527 | 0.641318354 | 0.19492735 | 3.29003777 |
| ENSG00000042317.16 | SPATA7 | 117.796642 | 0.703422633 | 0.21409827 | 3.28551294 |
| ENSG00000154096.13 | THY1 | 36.1809121 | 0.857064487 | 0.26106853 | 3.28291001 |
| ENSG00000122861.15 | PLAU | 4.33122977 | 2.425924558 | 0.74103989 | 3.27367609 |
| ENSG00000076344.15 | RGS11 | 2.69943677 | 1.878801055 | 0.57414508 | 3.27234546 |
| ENSG00000225420.1 | AC104134.1 | 194.146488 | 0.626102855 | 0.19146793 | 3.27001422 |
| ENSG00000167721.10 | TSR1 | 197.521326 | 0.441751741 | 0.1353019 | 3.26493372 |
| ENSG00000232085.1 | AL606534.3 | 172.06009 | 0.631405319 | 0.19353329 | 3.26251524 |
| ENSG00000157542.10 | KCNJ6 | 2.57639152 | 3.412036901 | 1.04628432 | 3.26109914 |
| ENSG00000259661.1 | AC068831.4 | 246.516948 | 0.666245151 | 0.20451814 | 3.2576335 |
| ENSG00000266341.1 | AC004477.2 | 1513.1776 | 0.566801028 | 0.17402168 | 3.25707131 |
| ENSG00000238018.2 | AC093110.1 | 792.185809 | 0.605014849 | 0.18602314 | 3.25236332 |
| ENSG00000085788.13 | DDHD2 | 68.4948077 | 0.729554813 | 0.22505527 | 3.2416696 |
| ENSG00000267787.6 | AC027097.2 | 241.88793 | 0.975987197 | 0.30169635 | 3.23499837 |
| ENSG00000257494.1 | AC004217.1 | 20.7980865 | 0.67297892 | 0.20796053 | 3.23608962 |
| ENSG00000258428.5 | AL161757.2 | 49.0187104 | 0.669975976 | 0.20708129 | 3.23532841 |
| ENSG00000148824.18 | MTG1 | 19.1793454 | 1.066443587 | 0.3297433 | 3.23416302 |
| ENSG00000149716.12 | ORAOV1 | 8956.85788 | 0.744149089 | 0.23014113 | 3.23344669 |
| ENSG00000228242.6 | AC093495.1 | 363.049234 | 0.502635286 | 0.15546864 | 3.23303325 |
| ENSG00000066855.15 | MTFR1 | 37.6675883 | 0.92762637 | 0.28696393 | 3.2325539 |
| ENSG00000151806.13 | GUF1 | 15.849271 | 1.060102376 | 0.32814923 | 3.23054961 |
| ENSG00000229358.3 | DPY19L1P1 | 64.7559017 | 0.529655721 | 0.16399527 | 3.22970115 |
| ENSG00000104131.12 | EIF3J | 51.783071 | 0.666112016 | 0.20651061 | 3.22555829 |
| ENSG00000100678.18 | SLC8A3 | 1.94013511 | 4.478760197 | 1.38969638 | 3.2228336 |
| ENSG00000274024.1 | AL590282.1 | 4.66265923 | 1.72027567 | 0.53379164 | 3.22274751 |
| ENSG00000259351.1 | AC015914.1 | 33.2277883 | 0.891921305 | 0.27663985 | 3.22412439 |
| ENSG00000121988.17 | ZRANB3 | 190.433979 | 0.438483008 | 0.13603809 | 3.22323711 |
| ENSG00000169764.15 | UGP2 | 10.2430063 | 1.045100702 | 0.32494203 | 3.21626813 |
| ENSG00000058729.10 | RIOK2 | 1.59881952 | 2.638717839 | 0.82120803 | 3.21321486 |
| ENSG00000237489.4 | C10orf143 | 4.44999042 | 1.9642432 | 0.61126847 | 3.2133887 |
| ENSG00000114021.11 | NIT2 | 30.5398463 | 0.723430822 | 0.22510092 | 3.21380662 |
| ENSG00000263069.5 | AC124319.2 | 186.304378 | 0.666033981 | 0.20741275 | 3.21115259 |
| ENSG00000251196.1 | AC106760.1 | 45.1477006 | 1.198417259 | 0.37364739 | 3.20734818 |

| | | | | | |
|---|---|---|---|---|---|
| ENSG00000273387.1 | AC005005.3 | 264.571346 | 0.419347092 | 0.13082446 | 3.20541811 |
| ENSG00000130758.7 | MAP3K10 | 7.84675841 | 1.357299526 | 0.42364619 | 3.2038516 |
| ENSG00000279344.1 | AC007342.6 | 289.709238 | 0.440711513 | 0.13756915 | 3.20356361 |
| ENSG00000179889.18 | PDXDC1 | 467.322131 | 0.440563451 | 0.13758539 | 3.20210927 |
| ENSG00000164106.7 | SCRG1 | 2.58695218 | 2.136386782 | 0.67016848 | 3.18783536 |
| ENSG00000123342.15 | MMP19 | 131.90413 | 0.788060993 | 0.24728094 | 3.18690555 |
| ENSG00000244104.3 | RN7SL659P | 1.89441654 | 3.270383242 | 1.02696418 | 3.18451538 |
| ENSG00000260465.1 | AC018557.1 | 928.262277 | 0.535285481 | 0.16817407 | 3.18292512 |
| ENSG00000285244.1 | DINOL | 2.36451288 | 2.357322251 | 0.7408463 | 3.1819316 |
| ENSG00000124588.19 | NQO2 | 293.540804 | 1.173822315 | 0.36907281 | 3.1804627 |
| ENSG00000231104.8 | AC022395.1 | 121.004639 | 0.831318813 | 0.26160948 | 3.17770901 |
| ENSG00000165124.17 | SVEP1 | 13.3605094 | 0.900823692 | 0.28358382 | 3.17656944 |
| ENSG00000058668.14 | ATP2B4 | 37.8527516 | 0.808648015 | 0.25519616 | 3.16873109 |
| ENSG00000215717.5 | TMEM167B | 27.9824743 | 0.662406203 | 0.20902986 | 3.16895491 |
| ENSG00000250794.2 | ALG1L12P | 2.31747682 | 3.177024391 | 1.00361848 | 3.16556985 |
| ENSG00000154928.17 | EPHB1 | 15.1041406 | 1.555707431 | 0.4919431 | 3.16237273 |
| ENSG00000183287.14 | CCBE1 | 10.0308252 | 1.606194935 | 0.50803533 | 3.1615812 |
| ENSG00000230155.6 | FO393401.1 | 40.0689342 | 0.67365639 | 0.21322299 | 3.15939846 |
| ENSG00000271064.1 | AC027644.2 | 11.3297725 | 0.925076379 | 0.29312594 | 3.15590074 |
| ENSG00000267257.1 | AC105105.1 | 2488.28795 | 0.676312586 | 0.21434508 | 3.15525132 |
| ENSG00000260007.3 | AC107871.1 | 55.555381 | 0.686921165 | 0.21814069 | 3.14898221 |
| ENSG00000272842.1 | AL391834.1 | 36.9202925 | 0.92233851 | 0.29345199 | 3.14306437 |
| ENSG00000182541.17 | LIMK2 | 15.9997893 | 0.988392863 | 0.31464967 | 3.14124866 |
| ENSG00000116171.17 | SCP2 | 14.6709111 | 0.985867604 | 0.31390144 | 3.14069159 |
| ENSG00000168769.13 | TET2 | 2.96187063 | 1.835701332 | 0.58597463 | 3.13273174 |
| ENSG00000121940.15 | CLCC1 | 120.389098 | 1.261255404 | 0.40259169 | 3.13284013 |
| ENSG00000255203.1 | OR7E2P | 2.00895491 | 2.80326996 | 0.89508537 | 3.13184647 |
| ENSG00000264486.1 | AC061975.4 | 2.22071883 | 2.331184796 | 0.74550717 | 3.12697836 |
| ENSG00000234149.1 | AC018511.2 | 49.4430951 | 0.806473907 | 0.25801571 | 3.12567758 |
| ENSG00000267576.1 | AC011472.3 | 53.312196 | 0.992481686 | 0.3177238 | 3.12372474 |
| ENSG00000223443.2 | USP17L2 | 1.4600888 | 3.282549242 | 1.05263443 | 3.11841333 |
| ENSG00000101624.10 | CEP76 | 3.28388266 | 1.875783408 | 0.60190973 | 3.11638657 |
| ENSG00000116194.12 | ANGPTL1 | 2.64009991 | 2.364780486 | 0.76027522 | 3.11042688 |
| ENSG00000129625.12 | REEP5 | 21.3295268 | 0.696670739 | 0.22404294 | 3.10954109 |
| ENSG00000267040.6 | AC027097.1 | 261.57708 | 1.009509771 | 0.32486463 | 3.10747824 |
| ENSG00000180448.10 | ARHGAP45 | 118.797683 | 0.723873123 | 0.23298857 | 3.10690406 |
| ENSG00000137962.12 | ARHGAP29 | 1.2735537 | 2.980441821 | 0.95989741 | 3.10495871 |
| ENSG00000196569.12 | LAMA2 | 17.1676764 | 1.075788226 | 0.34642432 | 3.10540617 |
| ENSG00000245311.2 | ARNTL2-AS1 | 56.2749594 | 0.9765148 | 0.3149192 | 3.10084238 |
| ENSG00000269843.1 | AC008537.2 | 29.8530485 | 0.752239244 | 0.2425756 | 3.10105069 |
| ENSG00000130021.13 | PUDP | 26.0039653 | 1.021295992 | 0.32946917 | 3.09982267 |
| ENSG00000123700.4 | KCNJ2 | 1.88217371 | 2.317071392 | 0.74786439 | 3.09825072 |
| ENSG00000156453.13 | PCDH1 | 1.82108138 | 3.56598887 | 1.1518865 | 3.09578146 |

| | | | | | |
|---|---|---|---|---|---|
| ENSG00000258177.1 | AC008149.1 | 377.377158 | 0.715474368 | 0.23114913 | 3.09529335 |
| ENSG00000125459.15 | MSTO1 | 4.02661513 | 1.638901585 | 0.52963818 | 3.09437962 |
| ENSG00000283073.1 | SMUG1-AS1 | 1.74418595 | 2.502275408 | 0.80886845 | 3.09355051 |
| ENSG00000248801.6 | C8orf34-AS1 | 1.34338215 | 2.806276578 | 0.90937916 | 3.08592577 |
| ENSG00000266002.1 | AC091059.1 | 15.3127949 | 1.242504684 | 0.40260708 | 3.08614711 |
| ENSG00000162931.11 | TRIM17 | 15.3299041 | 0.760378197 | 0.24678592 | 3.0811247 |
| ENSG00000210144.1 | MT-TY | 666.983249 | 0.501842819 | 0.16292048 | 3.08029305 |
| ENSG00000100426.6 | ZBED4 | 14.1250409 | 1.050194624 | 0.34104976 | 3.07930029 |
| ENSG00000198695.2 | MT-ND6 | 20828.1155 | 0.627793783 | 0.20398593 | 3.07763283 |
| ENSG00000260176.1 | AC141586.2 | 6.66627995 | 1.267116411 | 0.41240985 | 3.07246881 |
| ENSG00000213839.4 | TMX2P1 | 122.047758 | 0.456083638 | 0.14851708 | 3.07091707 |
| ENSG00000126217.20 | MCF2L | 1.44953212 | 3.722662521 | 1.21320993 | 3.06844053 |
| ENSG00000229619.3 | MBNL1-AS1 | 150.09955 | 0.648685448 | 0.21141744 | 3.06826833 |
| ENSG00000115459.17 | ELMOD3 | 91.128606 | 0.444416419 | 0.14485732 | 3.06795959 |
| ENSG00000271889.1 | AC016747.2 | 2.59238781 | 1.825256578 | 0.59559474 | 3.06459485 |
| ENSG00000244493.1 | SLC9A9-AS2 | 11.2921693 | 1.302164731 | 0.42494252 | 3.06433147 |
| ENSG00000245213.6 | AC105285.1 | 29.0991238 | 0.819045598 | 0.26732497 | 3.06385744 |
| ENSG00000172349.17 | IL16 | 216.524153 | 1.130807406 | 0.36937935 | 3.06137146 |
| ENSG00000188869.12 | TMC3 | 13.8677289 | 1.097699353 | 0.35904315 | 3.05729092 |
| ENSG00000124942.13 | AHNAK | 151.29507 | 0.576017879 | 0.18878457 | 3.05119158 |
| ENSG00000272114.1 | AL136131.3 | 169.750917 | 1.029767728 | 0.33795631 | 3.0470439 |
| ENSG00000285545.1 | | 20.8568203 | 0.769402927 | 0.25245621 | 3.04766883 |
| ENSG00000253309.6 | SERPINE3 | 35.0175508 | 0.667806883 | 0.21915179 | 3.04723446 |
| ENSG00000109339.21 | MAPK10 | 377.270124 | 0.429463385 | 0.14091494 | 3.04767817 |
| ENSG00000164199.17 | ADGRV1 | 25.3852713 | 0.628995505 | 0.20694438 | 3.03944237 |
| ENSG00000164107.8 | HAND2 | 23.992361 | 1.212200226 | 0.39901123 | 3.03801031 |
| ENSG00000264044.1 | AC005726.2 | 1690.02125 | 0.502629198 | 0.16543896 | 3.03815494 |
| ENSG00000013293.5 | SLC7A14 | 1.56304174 | 2.897603871 | 0.95555983 | 3.03236258 |
| ENSG00000227440.1 | ATP5MC1P4 | 6.43034188 | 2.103711038 | 0.6941011 | 3.03084237 |
| ENSG00000164597.13 | COG5 | 480.323658 | 0.351551962 | 0.11643448 | 3.01931153 |
| ENSG00000162971.10 | TYW5 | 11.3144163 | 0.94700852 | 0.31391851 | 3.01673358 |
| ENSG00000214262.4 | ANKRD36BP1 | 201.176914 | 0.563604923 | 0.18720325 | 3.01065786 |
| ENSG00000243305.1 | AC026347.1 | 26.8833089 | 0.729024585 | 0.24220077 | 3.01000109 |
| ENSG00000226877.8 | AL354733.1 | 6.48314074 | 1.299240637 | 0.43187003 | 3.00840658 |
| ENSG00000235106.9 | BRD3OS | 130.209984 | 0.627740549 | 0.2087063 | 3.00776997 |
| ENSG00000155666.11 | KDM8 | 10.6433926 | 0.9512791 | 0.31669164 | 3.00380236 |
| ENSG00000072071.16 | ADGRL1 | 62.9647381 | 0.471045147 | 0.15696708 | 3.0009168 |
| ENSG00000233822.4 | HIST1H2BN | 5.54699066 | 1.851505447 | 0.61721073 | 2.99979465 |
| ENSG00000272211.1 | AC114760.2 | 5.66117382 | 1.175579214 | 0.39202555 | 2.99873108 |
| ENSG00000075073.14 | TACR2 | 12.0213526 | 0.966755744 | 0.32236881 | 2.99891213 |
| ENSG00000263683.1 | AC005154.4 | 6.68020697 | 1.208295861 | 0.40321356 | 2.99666476 |
| ENSG00000080947.14 | CROCCP3 | 8.01088541 | 0.988878801 | 0.32999971 | 2.99660508 |
| ENSG00000003509.15 | NDUFAF7 | 188.936142 | 0.553727941 | 0.18484615 | 2.99561527 |

| | | | | | |
|---|---|---|---|---|---|
| ENSG00000265962.1 | GACAT2 | 1.71434066 | 2.796462178 | 0.9344066 | 2.99276801 |
| ENSG00000176222.8 | ZNF404 | 8.17286222 | 1.207983626 | 0.40452813 | 2.98615482 |
| ENSG00000259652.1 | AC090181.1 | 20.8464936 | 0.756139268 | 0.25349832 | 2.98281771 |
| ENSG00000151665.12 | PIGF | 1427.16806 | 0.653896406 | 0.21932434 | 2.98141286 |
| ENSG00000204954.9 | C12orf73 | 58.824923 | 1.579825938 | 0.53036663 | 2.97874312 |
| ENSG00000259187.1 | AC122108.1 | 7.11104591 | 2.489941306 | 0.83744201 | 2.97327012 |
| ENSG00000280054.1 | AC004241.5 | 11.2241004 | 1.541294309 | 0.51809702 | 2.97491442 |
| ENSG00000138069.17 | RAB1A | 13.9785724 | 0.725767747 | 0.24464293 | 2.96664096 |
| ENSG00000235865.2 | GSN-AS1 | 553.606385 | 0.900436731 | 0.30387684 | 2.96316344 |
| ENSG00000136048.13 | DRAM1 | 5.9847032 | 1.441549141 | 0.48829363 | 2.9522178 |
| ENSG00000231638.1 | LUARIS | 13.8897381 | 1.31399998 | 0.44521347 | 2.95139317 |
| ENSG00000280416.1 | AC009084.3 | 189.358347 | 0.478007002 | 0.16207057 | 2.94937575 |
| ENSG00000090581.9 | GNPTG | 124.588663 | 0.430243628 | 0.14586927 | 2.94951515 |
| ENSG00000269987.1 | AC004542.2 | 558.646054 | 0.448437283 | 0.15218511 | 2.94665673 |
| ENSG00000067369.13 | TP53BP1 | 228.427801 | 0.390045048 | 0.13236734 | 2.94668639 |
| ENSG00000151773.12 | CCDC122 | 9.37597402 | 1.183915337 | 0.40207887 | 2.94448531 |
| ENSG00000235652.7 | AL356599.1 | 114.346642 | 0.52991362 | 0.1799668 | 2.94450769 |
| ENSG00000104419.14 | NDRG1 | 13.2222088 | 1.327796134 | 0.45132294 | 2.94200899 |
| ENSG00000240661.3 | AC063952.1 | 2900.1155 | 0.537599449 | 0.18295947 | 2.9383527 |
| ENSG00000154305.16 | MIA3 | 282.70888 | 0.938422492 | 0.31956595 | 2.93655341 |
| ENSG00000189398.5 | OR7E12P | 25.4885039 | 0.703283901 | 0.23972201 | 2.93374774 |
| ENSG00000285938.1 | | 9.75960016 | 1.010120701 | 0.34481667 | 2.92944274 |
| ENSG00000168502.17 | MTCL1 | 29.4860526 | 1.417503878 | 0.48432502 | 2.92676162 |
| ENSG00000268970.1 | AC022150.2 | 8.81308652 | 1.034252899 | 0.35342309 | 2.92638748 |
| ENSG00000200714.1 | RF00019 | 1.72312471 | 2.168983773 | 0.74224413 | 2.92219729 |
| ENSG00000275055.1 | AC011468.5 | 10.6118297 | 0.935282983 | 0.32018329 | 2.92108621 |
| ENSG00000255250.1 | AP003059.2 | 1.44394511 | 2.851596697 | 0.97642982 | 2.92043179 |
| ENSG00000084234.17 | APLP2 | 81.4779722 | 0.974119672 | 0.3336006 | 2.92001771 |
| ENSG00000213970.4 | AC006122.1 | 3.09728784 | 2.442439343 | 0.83758593 | 2.9160463 |
| ENSG00000283973.1 | AC099795.1 | 4.989618 | 1.342274479 | 0.46031902 | 2.91596571 |
| ENSG00000213742.6 | ZNF337-AS1 | 272.768641 | 0.538321499 | 0.18459384 | 2.91624838 |
| ENSG00000250877.1 | AC095056.1 | 1.34632028 | 3.390476902 | 1.16404492 | 2.91266844 |
| ENSG00000091409.14 | ITGA6 | 17.1063985 | 0.963342894 | 0.33083876 | 2.91181992 |
| ENSG00000240401.8 | AC012358.3 | 200.585466 | 0.615463385 | 0.2113705 | 2.91177519 |
| ENSG00000116062.14 | MSH6 | 725.679974 | 0.413496887 | 0.1420235 | 2.91146806 |
| ENSG00000244040.6 | IL12A-AS1 | 4.80324761 | 1.65152844 | 0.56777175 | 2.90878941 |
| ENSG00000253406.1 | AC012613.2 | 107.251569 | 0.909513345 | 0.31312682 | 2.90461654 |
| ENSG00000134363.11 | FST | 4.19654358 | 1.594201395 | 0.54927911 | 2.90235214 |
| ENSG00000266872.1 | AC015688.7 | 47.8077169 | 0.66429754 | 0.22888509 | 2.90231896 |
| ENSG00000229400.1 | AL596330.1 | 54.3471606 | 0.937323814 | 0.32306399 | 2.90135654 |
| ENSG00000259773.1 | AC012100.2 | 152.399997 | 0.340922133 | 0.11755725 | 2.90005188 |
| ENSG00000258890.6 | CEP95 | 59.9637601 | 0.731165652 | 0.25249581 | 2.89575361 |
| ENSG00000272814.1 | AC093732.2 | 80.9538456 | 0.590870033 | 0.20427149 | 2.89257213 |

| | | | | | |
|---|---|---|---|---|---|
| ENSG00000239552.2 | HOXB-AS2 | 26.9984526 | 1.385742849 | 0.47950771 | 2.88992822 |
| ENSG00000275339.1 | Z99129.1 | 1.82062886 | 2.459037779 | 0.85143302 | 2.88811654 |
| ENSG00000258655.2 | ARHGAP5-AS | 6.14611328 | 1.192732625 | 0.41333148 | 2.88565636 |
| ENSG00000285704.1 | | 5.69471102 | 1.46536505 | 0.50841043 | 2.88224821 |
| ENSG00000271730.1 | AL390208.1 | 64.5033714 | 0.88358204 | 0.30662854 | 2.881604 |
| ENSG00000214243.3 | AC004980.2 | 6.53140612 | 1.092838618 | 0.38009641 | 2.87516163 |
| ENSG00000266957.1 | AC012254.1 | 5.50078984 | 1.318311816 | 0.45940661 | 2.86959695 |
| ENSG00000228677.1 | TTC3-AS1 | 462.426175 | 0.583415164 | 0.20354792 | 2.86623008 |
| ENSG00000213341.10 | CHUK | 1.28381315 | 3.045137331 | 1.06305128 | 2.86452534 |
| ENSG00000184895.7 | SRY | 10.8870309 | 1.596648626 | 0.55732972 | 2.86481872 |
| ENSG00000246174.7 | KCTD21-AS1 | 169.476604 | 0.492139855 | 0.1718991 | 2.86295779 |
| ENSG00000235954.6 | TTC28-AS1 | 192.351136 | 0.468457577 | 0.16362254 | 2.86303816 |

| pvalue | padj |
|---|---|
| 2.90E-24 | 3.52E-20 |
| 1.45E-21 | 8.81E-18 |
| 7.94E-20 | 3.21E-16 |
| 1.44E-17 | 4.36E-14 |
| 4.39E-16 | 1.07E-12 |
| 5.14E-14 | 7.79E-11 |
| 4.94E-14 | 7.79E-11 |
| 4.03E-14 | 7.79E-11 |
| 6.09E-13 | 7.26E-10 |
| 6.69E-13 | 7.26E-10 |
| 2.78E-12 | 2.41E-09 |
| 3.37E-12 | 2.72E-09 |
| 5.02E-12 | 3.81E-09 |
| 8.69E-12 | 6.20E-09 |
| 1.38E-11 | 9.29E-09 |
| 2.64E-11 | 1.69E-08 |
| 4.32E-11 | 2.62E-08 |
| 5.20E-11 | 3.00E-08 |
| 6.99E-11 | 3.85E-08 |
| 7.57E-11 | 3.99E-08 |
| 3.11E-10 | 1.46E-07 |
| 3.13E-10 | 1.46E-07 |
| 4.34E-10 | 1.95E-07 |
| 4.94E-10 | 2.14E-07 |
| 9.40E-10 | 3.76E-07 |
| 9.61E-10 | 3.76E-07 |
| 1.17E-09 | 4.28E-07 |
| 1.16E-09 | 4.28E-07 |
| 1.36E-09 | 4.73E-07 |
| 2.22E-09 | 7.29E-07 |
| 3.04E-09 | 9.69E-07 |
| 3.81E-09 | 1.15E-06 |
| 4.10E-09 | 1.21E-06 |
| 4.58E-09 | 1.32E-06 |
| 4.89E-09 | 1.37E-06 |
| 5.58E-09 | 1.47E-06 |
| 5.55E-09 | 1.47E-06 |
| 8.28E-09 | 2.05E-06 |
| 9.52E-09 | 2.26E-06 |
| 1.00E-08 | 2.33E-06 |
| 1.19E-08 | 2.68E-06 |
| 1.22E-08 | 2.69E-06 |

| | |
|---|---|
| 1.28E-08 | 2.78E-06 |
| 1.37E-08 | 2.91E-06 |
| 1.50E-08 | 3.14E-06 |
| 1.81E-08 | 3.65E-06 |
| 1.87E-08 | 3.65E-06 |
| 2.03E-08 | 3.90E-06 |
| 3.30E-08 | 6.16E-06 |
| 4.32E-08 | 7.82E-06 |
| 4.48E-08 | 7.99E-06 |
| 4.97E-08 | 8.73E-06 |
| 5.15E-08 | 8.92E-06 |
| 6.03E-08 | 1.03E-05 |
| 6.34E-08 | 1.05E-05 |
| 6.78E-08 | 1.07E-05 |
| 6.82E-08 | 1.07E-05 |
| 6.79E-08 | 1.07E-05 |
| 6.80E-08 | 1.07E-05 |
| 7.97E-08 | 1.21E-05 |
| 9.91E-08 | 1.48E-05 |
| 1.07E-07 | 1.59E-05 |
| 1.26E-07 | 1.81E-05 |
| 1.44E-07 | 1.99E-05 |
| 1.45E-07 | 1.99E-05 |
| 2.00E-07 | 2.64E-05 |
| 2.36E-07 | 3.01E-05 |
| 2.70E-07 | 3.41E-05 |
| 2.82E-07 | 3.53E-05 |
| 2.87E-07 | 3.54E-05 |
| 3.29E-07 | 4.03E-05 |
| 4.29E-07 | 4.99E-05 |
| 4.32E-07 | 4.99E-05 |
| 4.82E-07 | 5.51E-05 |
| 5.07E-07 | 5.69E-05 |
| 6.55E-07 | 7.15E-05 |
| 6.73E-07 | 7.28E-05 |
| 7.91E-07 | 8.48E-05 |
| 7.98E-07 | 8.49E-05 |
| 8.56E-07 | 8.87E-05 |
| 9.08E-07 | 9.25E-05 |
| 9.38E-07 | 9.48E-05 |
| 1.20E-06 | 0.00012035 |
| 1.28E-06 | 0.0001261 |
| 1.36E-06 | 0.00013131 |

| | |
|---|---|
| 1.38E-06 | 0.0001317 |
| 1.46E-06 | 0.00013864 |
| 1.64E-06 | 0.00015421 |
| 1.82E-06 | 0.00016721 |
| 1.83E-06 | 0.00016721 |
| 1.92E-06 | 0.00017245 |
| 2.09E-06 | 0.0001862 |
| 2.17E-06 | 0.0001895 |
| 2.17E-06 | 0.0001895 |
| 2.22E-06 | 0.00019234 |
| 2.24E-06 | 0.00019235 |
| 2.30E-06 | 0.00019467 |
| 2.29E-06 | 0.00019467 |
| 2.56E-06 | 0.00021307 |
| 2.88E-06 | 0.0002355 |
| 3.13E-06 | 0.00025137 |
| 3.20E-06 | 0.00025383 |
| 3.28E-06 | 0.00025783 |
| 3.56E-06 | 0.00027505 |
| 3.79E-06 | 0.00029111 |
| 3.84E-06 | 0.00029304 |
| 3.90E-06 | 0.00029556 |
| 4.49E-06 | 0.00033378 |
| 4.60E-06 | 0.00033811 |
| 4.57E-06 | 0.00033811 |
| 4.94E-06 | 0.0003583 |
| 4.98E-06 | 0.00035965 |
| 5.33E-06 | 0.00038202 |
| 5.42E-06 | 0.00038525 |
| 5.60E-06 | 0.00039251 |
| 5.81E-06 | 0.00040508 |
| 5.86E-06 | 0.0004059 |
| 6.12E-06 | 0.00042163 |
| 6.69E-06 | 0.00045464 |
| 6.65E-06 | 0.00045464 |
| 7.25E-06 | 0.00048006 |
| 7.95E-06 | 0.00052096 |
| 8.51E-06 | 0.00055138 |
| 8.67E-06 | 0.00055749 |
| 8.69E-06 | 0.00055749 |
| 8.83E-06 | 0.0005605 |
| 9.14E-06 | 0.00057433 |
| 9.46E-06 | 0.00058946 |

| | |
|---|---|
| 9.77E-06 | 0.00060136 |
| 1.05E-05 | 0.00063615 |
| 1.11E-05 | 0.00066576 |
| 1.13E-05 | 0.00067194 |
| 1.14E-05 | 0.00067194 |
| 1.24E-05 | 0.0007218 |
| 1.24E-05 | 0.0007218 |
| 1.25E-05 | 0.00072436 |
| 1.29E-05 | 0.00073989 |
| 1.36E-05 | 0.00077553 |
| 1.37E-05 | 0.00077759 |
| 1.40E-05 | 0.00078743 |
| 1.43E-05 | 0.00080333 |
| 1.49E-05 | 0.00082945 |
| 1.54E-05 | 0.00084337 |
| 1.63E-05 | 0.00088083 |
| 1.71E-05 | 0.00092227 |
| 1.75E-05 | 0.00094113 |
| 1.86E-05 | 0.00098586 |
| 1.98E-05 | 0.00103295 |
| 1.99E-05 | 0.00103295 |
| 2.03E-05 | 0.0010453 |
| 2.13E-05 | 0.00109142 |
| 2.21E-05 | 0.00111468 |
| 2.23E-05 | 0.00112077 |
| 2.26E-05 | 0.00113266 |
| 2.28E-05 | 0.00113594 |
| 2.39E-05 | 0.00118734 |
| 2.46E-05 | 0.0012132 |
| 2.62E-05 | 0.00126874 |
| 2.70E-05 | 0.0012892 |
| 2.76E-05 | 0.0013077 |
| 2.80E-05 | 0.00130807 |
| 2.82E-05 | 0.00130849 |
| 2.98E-05 | 0.00136213 |
| 3.00E-05 | 0.00136856 |
| 3.12E-05 | 0.00140998 |
| 3.16E-05 | 0.00142356 |
| 3.32E-05 | 0.00147526 |
| 3.34E-05 | 0.00147887 |
| 3.36E-05 | 0.00148187 |
| 3.60E-05 | 0.00158291 |
| 3.73E-05 | 0.00161194 |

| | |
|---|---|
| 3.84E-05 | 0.00164451 |
| 4.07E-05 | 0.00173762 |
| 4.11E-05 | 0.00174801 |
| 4.17E-05 | 0.00176397 |
| 4.63E-05 | 0.00194288 |
| 4.68E-05 | 0.00194288 |
| 4.85E-05 | 0.00199217 |
| 4.88E-05 | 0.00199313 |
| 5.00E-05 | 0.00202914 |
| 5.51E-05 | 0.00220633 |
| 5.72E-05 | 0.00228177 |
| 6.02E-05 | 0.0023844 |
| 6.13E-05 | 0.00240346 |
| 6.52E-05 | 0.00253152 |
| 6.59E-05 | 0.00254707 |
| 6.69E-05 | 0.00257362 |
| 7.04E-05 | 0.00266753 |
| 7.17E-05 | 0.00270701 |
| 7.29E-05 | 0.00272949 |
| 7.40E-05 | 0.00276128 |
| 7.53E-05 | 0.00279877 |
| 7.60E-05 | 0.00280613 |
| 7.75E-05 | 0.00283012 |
| 8.05E-05 | 0.00291295 |
| 8.21E-05 | 0.00295921 |
| 9.31E-05 | 0.00328911 |
| 9.34E-05 | 0.00329268 |
| 9.61E-05 | 0.0033507 |
| 9.95E-05 | 0.00338712 |
| 0.00010124 | 0.00343782 |
| 0.00010249 | 0.00346108 |
| 0.00010301 | 0.00346901 |
| 0.00010348 | 0.00347502 |
| 0.00010455 | 0.00349167 |
| 0.00010561 | 0.00350754 |
| 0.00010875 | 0.00360227 |
| 0.00011044 | 0.00364805 |
| 0.00011111 | 0.00366033 |
| 0.00011619 | 0.00381725 |
| 0.0001241 | 0.00403616 |
| 0.00012675 | 0.00407596 |
| 0.00012935 | 0.00412661 |
| 0.0001355 | 0.00430024 |

| | |
|---|---|
| 0.00014177 | 0.004464 |
| 0.00014641 | 0.00458622 |
| 0.00015256 | 0.0047545 |
| 0.00015422 | 0.00479172 |
| 0.00015455 | 0.00479172 |
| 0.00015653 | 0.00484096 |
| 0.00016664 | 0.00506303 |
| 0.00016937 | 0.00512049 |
| 0.00017496 | 0.005276 |
| 0.00018471 | 0.0055427 |
| 0.00018626 | 0.00557551 |
| 0.00018888 | 0.00562599 |
| 0.00019713 | 0.00577485 |
| 0.00019608 | 0.00577485 |
| 0.00020756 | 0.00604858 |
| 0.00020733 | 0.00604858 |
| 0.00021237 | 0.00616361 |
| 0.00021448 | 0.00617606 |
| 0.00021517 | 0.00618136 |
| 0.00022019 | 0.00625137 |
| 0.00022016 | 0.00625137 |
| 0.00022704 | 0.00641586 |
| 0.00022906 | 0.00645782 |
| 0.00023099 | 0.00649712 |
| 0.00023652 | 0.00662205 |
| 0.00023729 | 0.00662818 |
| 0.00024146 | 0.0067134 |
| 0.00024561 | 0.00676718 |
| 0.00024776 | 0.00681077 |
| 0.00025158 | 0.00687164 |
| 0.00025167 | 0.00687164 |
| 0.00025408 | 0.00687595 |
| 0.00025279 | 0.00687595 |
| 0.00026407 | 0.00708257 |
| 0.00026935 | 0.00719246 |
| 0.00027365 | 0.00728299 |
| 0.00027624 | 0.00732791 |
| 0.00028064 | 0.0074121 |
| 0.00028626 | 0.00752783 |
| 0.00029147 | 0.00762177 |
| 0.00029172 | 0.00762177 |
| 0.00030027 | 0.00781164 |
| 0.00030168 | 0.0078129 |

| | |
|---|---|
| 0.00030965 | 0.00798695 |
| 0.00032762 | 0.00841131 |
| 0.00032818 | 0.00841131 |
| 0.000333 | 0.00848088 |
| 0.00034049 | 0.00865354 |
| 0.00034427 | 0.00870729 |
| 0.00034861 | 0.00878627 |
| 0.00035128 | 0.0088313 |
| 0.00035258 | 0.0088313 |
| 0.00035655 | 0.00891227 |
| 0.00035852 | 0.00892472 |
| 0.0003594 | 0.00892822 |
| 0.0003677 | 0.00909724 |
| 0.00038243 | 0.00940398 |
| 0.00039522 | 0.00967939 |
| 0.0004013 | 0.00978856 |
| 0.00040239 | 0.00979557 |
| 0.00041053 | 0.00997375 |
| 0.00041546 | 0.01007109 |
| 0.00041703 | 0.01007109 |
| 0.00042044 | 0.01013319 |
| 0.0004214 | 0.01013617 |
| 0.00044639 | 0.01052849 |
| 0.00046491 | 0.01085699 |
| 0.0004657 | 0.01085699 |
| 0.00046761 | 0.01088068 |
| 0.00047055 | 0.01092808 |
| 0.00048305 | 0.01117563 |
| 0.00050787 | 0.01166077 |
| 0.00050993 | 0.01166388 |
| 0.00050923 | 0.01166388 |
| 0.00051618 | 0.01176869 |
| 0.0005332 | 0.01210487 |
| 0.00054468 | 0.01220537 |
| 0.00054135 | 0.01220537 |
| 0.00056442 | 0.01257797 |
| 0.00056701 | 0.01261268 |
| 0.00057563 | 0.012706 |
| 0.00057772 | 0.012706 |
| 0.00059174 | 0.01290234 |
| 0.00059363 | 0.01292022 |
| 0.00059602 | 0.0129491 |
| 0.00060685 | 0.01315893 |

| | |
|---|---|
| 0.000617 | 0.0133095 |
| 0.00061941 | 0.01331398 |
| 0.00061838 | 0.01331398 |
| 0.00062711 | 0.01343183 |
| 0.00063664 | 0.01353411 |
| 0.0006397 | 0.01353411 |
| 0.0006478 | 0.01368174 |
| 0.00065496 | 0.01380881 |
| 0.00066076 | 0.0138828 |
| 0.00067036 | 0.01400623 |
| 0.00067241 | 0.01400623 |
| 0.00067097 | 0.01400623 |
| 0.00067814 | 0.0140771 |
| 0.00069565 | 0.01436175 |
| 0.00069659 | 0.01436175 |
| 0.00070067 | 0.01442139 |
| 0.0007105 | 0.01450065 |
| 0.00071548 | 0.01455322 |
| 0.00072329 | 0.01466289 |
| 0.00072889 | 0.01470268 |
| 0.00072835 | 0.01470268 |
| 0.00073519 | 0.01480508 |
| 0.0007367 | 0.01481104 |
| 0.00074228 | 0.01489849 |
| 0.00074567 | 0.01494172 |
| 0.0007493 | 0.01498968 |
| 0.00075394 | 0.01505759 |
| 0.00075641 | 0.01508224 |
| 0.00077468 | 0.01539573 |
| 0.00077951 | 0.01544122 |
| 0.00077829 | 0.01544122 |
| 0.00079514 | 0.01568486 |
| 0.00079849 | 0.0157145 |
| 0.0008114 | 0.01591678 |
| 0.00082208 | 0.01602255 |
| 0.00082138 | 0.01602255 |
| 0.00082803 | 0.01607719 |
| 0.00083284 | 0.01612855 |
| 0.00083763 | 0.01619543 |
| 0.00084268 | 0.01626723 |
| 0.00084591 | 0.01630357 |
| 0.00084906 | 0.01633827 |
| 0.00087187 | 0.01669769 |

| | |
|---|---|
| 0.00087768 | 0.01674484 |
| 0.00088131 | 0.01677249 |
| 0.00090701 | 0.01718083 |
| 0.00093888 | 0.0176193 |
| 0.0009647 | 0.01804784 |
| 0.00097583 | 0.01822809 |
| 0.00098525 | 0.01829243 |
| 0.00098312 | 0.01829243 |
| 0.00099381 | 0.01842187 |
| 0.0010013 | 0.01851233 |
| 0.00100174 | 0.01851233 |
| 0.00101797 | 0.01878361 |
| 0.00102741 | 0.0189291 |
| 0.00106158 | 0.01946984 |
| 0.00106659 | 0.01953216 |
| 0.00107542 | 0.01966414 |
| 0.0010949 | 0.01993008 |
| 0.00110428 | 0.02001078 |
| 0.00110981 | 0.02008098 |
| 0.00112345 | 0.0202372 |
| 0.00112568 | 0.02024724 |
| 0.0011445 | 0.0204962 |
| 0.00118832 | 0.02121645 |
| 0.00121643 | 0.02151584 |
| 0.00121179 | 0.02151584 |
| 0.00121503 | 0.02151584 |
| 0.00122 | 0.02152844 |
| 0.00122306 | 0.02155102 |
| 0.00122483 | 0.02155102 |
| 0.00122689 | 0.02155592 |
| 0.00123552 | 0.02167622 |
| 0.0012392 | 0.0216779 |
| 0.00125727 | 0.0218993 |
| 0.00126929 | 0.02195758 |
| 0.00126967 | 0.02195758 |
| 0.00126358 | 0.02195758 |
| 0.00126751 | 0.02195758 |
| 0.00129869 | 0.02242745 |
| 0.00131258 | 0.02250695 |
| 0.00131179 | 0.02250695 |
| 0.00130988 | 0.02250695 |
| 0.00132204 | 0.02263709 |
| 0.00133965 | 0.02284184 |

| | |
|---|---|
| 0.00134866 | 0.02296328 |
| 0.00135602 | 0.02301472 |
| 0.00135738 | 0.02301472 |
| 0.00136425 | 0.02303458 |
| 0.00143342 | 0.02390284 |
| 0.00143804 | 0.02394688 |
| 0.00144997 | 0.02407937 |
| 0.00145795 | 0.02417889 |
| 0.00146296 | 0.02419578 |
| 0.0014704 | 0.02428565 |
| 0.00148444 | 0.02448411 |
| 0.00149028 | 0.02453852 |
| 0.00153106 | 0.02508248 |
| 0.00152988 | 0.02508248 |
| 0.00154779 | 0.02532243 |
| 0.00156489 | 0.02553321 |
| 0.00156915 | 0.02553398 |
| 0.00158095 | 0.02565713 |
| 0.00160003 | 0.02586293 |
| 0.0016036 | 0.02587911 |
| 0.0016384 | 0.02634263 |
| 0.00167189 | 0.02677455 |
| 0.00168229 | 0.02690556 |
| 0.00168549 | 0.02692127 |
| 0.00173188 | 0.02758941 |
| 0.00173124 | 0.02758941 |
| 0.00173711 | 0.02763641 |
| 0.00176613 | 0.02798795 |
| 0.00177396 | 0.02803873 |
| 0.00178577 | 0.02815758 |
| 0.00181828 | 0.02855306 |
| 0.00183082 | 0.02863877 |
| 0.00186817 | 0.02907297 |
| 0.00187378 | 0.0291229 |
| 0.00188691 | 0.02928937 |
| 0.00189058 | 0.02930879 |
| 0.00190306 | 0.02936405 |
| 0.00190018 | 0.02936405 |
| 0.00192971 | 0.02957506 |
| 0.00192835 | 0.02957506 |
| 0.00193637 | 0.02963959 |
| 0.00194667 | 0.0296888 |
| 0.00196295 | 0.02989553 |

| | |
|---|---|
| 0.00196618 | 0.0299072 |
| 0.00197225 | 0.02996188 |
| 0.00197777 | 0.03000812 |
| 0.00202919 | 0.03055538 |
| 0.00202768 | 0.03055538 |
| 0.0020622 | 0.03078828 |
| 0.00206797 | 0.03083641 |
| 0.00207487 | 0.03084068 |
| 0.00208652 | 0.03092281 |
| 0.00212296 | 0.03138615 |
| 0.00213402 | 0.03147296 |
| 0.00215179 | 0.03163219 |
| 0.00215303 | 0.03163219 |
| 0.00215526 | 0.03163219 |
| 0.00217965 | 0.03186407 |
| 0.00218157 | 0.03186407 |
| 0.00218503 | 0.0318762 |
| 0.00220326 | 0.03210345 |
| 0.00223347 | 0.03242683 |
| 0.00227935 | 0.03283824 |
| 0.00231104 | 0.0331167 |
| 0.00230624 | 0.0331167 |
| 0.00230957 | 0.0331167 |
| 0.00230617 | 0.0331167 |
| 0.00237017 | 0.03380414 |
| 0.00238146 | 0.03388546 |
| 0.00238032 | 0.03388546 |
| 0.00242648 | 0.03440487 |
| 0.00243873 | 0.03445765 |
| 0.0025335 | 0.03567202 |
| 0.00255514 | 0.03593503 |
| 0.00260682 | 0.03653471 |
| 0.00261247 | 0.03657153 |
| 0.00262622 | 0.03667927 |
| 0.00263172 | 0.03671389 |
| 0.00266628 | 0.03702978 |
| 0.00269168 | 0.03725028 |
| 0.00270162 | 0.03734515 |
| 0.00271106 | 0.03738506 |
| 0.00270945 | 0.03738506 |
| 0.00272951 | 0.03748164 |
| 0.00273004 | 0.03748164 |
| 0.00273892 | 0.03756097 |

| | |
|---|---|
| 0.0027646 | 0.03778492 |
| 0.0028251 | 0.0385249 |
| 0.00285608 | 0.03877297 |
| 0.00286922 | 0.03890774 |
| 0.00289433 | 0.03920222 |
| 0.00294645 | 0.03946673 |
| 0.0029307 | 0.03946673 |
| 0.00301072 | 0.04006476 |
| 0.00304495 | 0.04038721 |
| 0.003155 | 0.04147523 |
| 0.00316344 | 0.04147523 |
| 0.00318417 | 0.04164147 |
| 0.00318273 | 0.04164147 |
| 0.0032123 | 0.04182885 |
| 0.00321199 | 0.04182885 |
| 0.00323492 | 0.04198819 |
| 0.00323469 | 0.04198819 |
| 0.0032609 | 0.04227746 |
| 0.00329961 | 0.04255449 |
| 0.00331882 | 0.04275666 |
| 0.00334896 | 0.04298696 |
| 0.0033957 | 0.04337842 |
| 0.00342511 | 0.04370805 |
| 0.00342924 | 0.04371464 |
| 0.00347571 | 0.04409451 |
| 0.00348813 | 0.04409451 |
| 0.00349547 | 0.044108 |
| 0.00350011 | 0.044108 |
| 0.00354498 | 0.0444743 |
| 0.00354589 | 0.0444743 |
| 0.00354268 | 0.0444743 |
| 0.00358355 | 0.04469482 |
| 0.0035933 | 0.04472886 |
| 0.00359381 | 0.04472886 |
| 0.00359735 | 0.04472886 |
| 0.00362831 | 0.04506765 |
| 0.00367703 | 0.04557942 |
| 0.00370372 | 0.04577467 |
| 0.00370411 | 0.04577467 |
| 0.00371551 | 0.04582209 |
| 0.00373101 | 0.04596649 |
| 0.00378249 | 0.04645 |
| 0.00382101 | 0.04669572 |

| | |
|---|---|
| 0.0038533 | 0.04699551 |
| 0.00387556 | 0.04721954 |
| 0.00390598 | 0.04754241 |
| 0.00394849 | 0.04796342 |
| 0.00395657 | 0.04801347 |
| 0.00403821 | 0.04890628 |
| 0.00410995 | 0.04953022 |
| 0.00415392 | 0.04990882 |
| 0.00417635 | 0.04991971 |
| 0.00417248 | 0.04991971 |
| 0.00419706 | 0.04998135 |
| 0.004196 | 0.04998135 |

| Gene id | baseMean | log2FoldChange | lfcSE | stat | pvalue |
|---|---|---|---|---|---|
| ENSG00000255050.1 | 100.098072 | -1.361798465 | 0.18589072 | -7.3258013 | 2.37E-13 |
| ENSG00000271581.1 | 1026.35235 | -1.457161618 | 0.20306284 | -7.1759149 | 7.18E-13 |
| ENSG00000231584.8 | 21.1929942 | -6.064130433 | 0.85077358 | -7.1277841 | 1.02E-12 |
| ENSG00000279035.1 | 114.386886 | -1.003392935 | 0.15669124 | -6.4036312 | 1.52E-10 |
| ENSG00000277443.2 | 90.5561212 | -1.54928799 | 0.251101 | -6.1699793 | 6.83E-10 |
| ENSG00000132763.14 | 2231.78267 | -0.647085633 | 0.10654654 | -6.0732677 | 1.25E-09 |
| ENSG00000148331.11 | 318.50383 | -0.707657465 | 0.11787193 | -6.0036134 | 1.93E-09 |
| ENSG00000278367.1 | 92.2072617 | -1.336413531 | 0.22591943 | -5.915443 | 3.31E-09 |
| ENSG00000258471.2 | 158.941736 | -0.794018145 | 0.13576232 | -5.84859 | 4.96E-09 |
| ENSG00000135452.9 | 674.652832 | -0.720548641 | 0.12413657 | -5.8044833 | 6.46E-09 |
| ENSG00000157734.13 | 8549.88986 | -0.732138437 | 0.12676172 | -5.7757063 | 7.66E-09 |
| ENSG00000230521.1 | 30.9447015 | -3.321262509 | 0.57849041 | -5.7412577 | 9.40E-09 |
| ENSG00000283117.1 | 34.1184263 | -1.563456269 | 0.27297657 | -5.7274375 | 1.02E-08 |
| ENSG00000213347.10 | 1286.33105 | -0.589146313 | 0.10416055 | -5.6561365 | 1.55E-08 |
| ENSG00000267458.1 | 3659.89242 | -0.728148894 | 0.12946151 | -5.6244432 | 1.86E-08 |
| ENSG00000272821.1 | 135.713313 | -0.7920977 | 0.14261714 | -5.5540149 | 2.79E-08 |
| ENSG00000259884.1 | 178.391331 | -1.624726273 | 0.29469293 | -5.5132855 | 3.52E-08 |
| ENSG00000281344.1 | 64.2960543 | -5.92873238 | 1.09501403 | -5.4142981 | 6.15E-08 |
| ENSG00000215030.5 | 12.7156151 | -4.64877684 | 0.86316566 | -5.3857296 | 7.22E-08 |
| ENSG00000104848.1 | 66.1553944 | -2.686191355 | 0.49882821 | -5.3850029 | 7.24E-08 |
| ENSG00000140416.20 | 81.9565708 | -1.424731529 | 0.26872675 | -5.3017853 | 1.15E-07 |
| ENSG00000256928.1 | 96.4296577 | -1.103012021 | 0.20904882 | -5.276337 | 1.32E-07 |
| ENSG00000204055.4 | 218.339379 | -1.072418309 | 0.20367483 | -5.2653453 | 1.40E-07 |
| ENSG00000147687.18 | 215.923694 | -0.875966095 | 0.16719437 | -5.2392082 | 1.61E-07 |
| ENSG00000143878.9 | 11.5213592 | -2.442299085 | 0.46743551 | -5.22489 | 1.74E-07 |
| ENSG00000148303.16 | 43.3273678 | -1.145457971 | 0.21920649 | -5.2254746 | 1.74E-07 |
| ENSG00000188368.9 | 62.1013722 | -1.002925186 | 0.19342226 | -5.1851591 | 2.16E-07 |
| ENSG00000127928.12 | 709.081647 | -4.249653599 | 0.82210399 | -5.169241 | 2.35E-07 |
| ENSG00000130640.13 | 124.255947 | -0.655694008 | 0.1285804 | -5.0994864 | 3.41E-07 |
| ENSG00000166794.4 | 32.0765734 | -1.102718948 | 0.21691674 | -5.0836046 | 3.70E-07 |
| ENSG00000179218.13 | 165.240762 | -0.930664251 | 0.18336411 | -5.0754985 | 3.86E-07 |
| ENSG00000130762.14 | 23.0967634 | -2.30848717 | 0.4558104 | -5.0645777 | 4.09E-07 |
| ENSG00000272173.1 | 591.114352 | -1.007899968 | 0.20043804 | -5.0284863 | 4.94E-07 |
| ENSG00000257950.3 | 284.514513 | -0.673559932 | 0.1343398 | -5.0138523 | 5.34E-07 |
| ENSG00000231389.7 | 11.2145709 | -2.494893168 | 0.49929872 | -4.9967946 | 5.83E-07 |
| ENSG00000230513.1 | 28.9996059 | -1.134766136 | 0.23002957 | -4.9331316 | 8.09E-07 |
| ENSG00000134871.18 | 73.7092329 | -1.350367837 | 0.27423873 | -4.9240596 | 8.48E-07 |
| ENSG00000284060.1 | 10.4399556 | -1.730825073 | 0.35194608 | -4.91787 | 8.75E-07 |
| ENSG00000117318.8 | 5.90408637 | -2.599575708 | 0.53651589 | -4.8452912 | 1.26E-06 |
| ENSG00000273398.6 | 104.31126 | -0.826701715 | 0.17078991 | -4.8404599 | 1.30E-06 |
| ENSG00000145916.18 | 372.22716 | -0.616503701 | 0.12752746 | -4.8342819 | 1.34E-06 |
| ENSG00000262533.1 | 411.396173 | -0.68709704 | 0.14358135 | -4.7854197 | 1.71E-06 |

| | | | | | |
|---|---|---|---|---|---|
| ENSG00000092820.17 | 36.2134534 | -1.275288362 | 0.26730801 | -4.7708573 | 1.83E-06 |
| ENSG00000108298.11 | 58.9696581 | -0.954220457 | 0.20016921 | -4.7670692 | 1.87E-06 |
| ENSG00000126768.12 | 60.5307877 | -0.720521551 | 0.15192426 | -4.7426366 | 2.11E-06 |
| ENSG00000184009.11 | 235.575819 | -0.916592498 | 0.19415822 | -4.7208535 | 2.35E-06 |
| ENSG00000176809.10 | 7.4830392 | -2.157715854 | 0.4588137 | -4.7028147 | 2.57E-06 |
| ENSG00000118181.10 | 46.6366389 | -0.90129225 | 0.19226264 | -4.687818 | 2.76E-06 |
| ENSG00000223865.10 | 2.67340617 | -4.300054333 | 0.91981459 | -4.6749143 | 2.94E-06 |
| ENSG00000285677.1 | 804.978269 | -0.687591437 | 0.14713993 | -4.6730444 | 2.97E-06 |
| ENSG00000226715.3 | 19.1076907 | -1.076026857 | 0.23104004 | -4.6573177 | 3.20E-06 |
| ENSG00000282843.1 | 7.85836192 | -2.211805743 | 0.47578949 | -4.6487066 | 3.34E-06 |
| ENSG00000235237.1 | 38.1224174 | -0.960294811 | 0.20664179 | -4.6471473 | 3.37E-06 |
| ENSG00000232358.1 | 41.5561696 | -1.797278398 | 0.389692 | -4.6120484 | 3.99E-06 |
| ENSG00000100234.11 | 98.796517 | -2.011638243 | 0.43734017 | -4.5997107 | 4.23E-06 |
| ENSG00000266208.1 | 13.7303864 | -1.753058876 | 0.38275601 | -4.580095 | 4.65E-06 |
| ENSG00000264666.1 | 58.67669 | -0.725840181 | 0.15962038 | -4.5472901 | 5.43E-06 |
| ENSG00000170889.13 | 15.1789441 | -1.212979547 | 0.26704587 | -4.5422142 | 5.57E-06 |
| ENSG00000107821.14 | 7.00269404 | -2.588379988 | 0.57486434 | -4.5025927 | 6.71E-06 |
| ENSG00000143450.16 | 125.103549 | -0.775485313 | 0.17228402 | -4.5012028 | 6.76E-06 |
| ENSG00000186714.12 | 292.791026 | -0.625994629 | 0.13921375 | -4.4966437 | 6.90E-06 |
| ENSG00000197915.5 | 6.45003249 | -2.246923807 | 0.50044057 | -4.4898914 | 7.13E-06 |
| ENSG00000118894.14 | 222.162629 | -0.460942282 | 0.10285484 | -4.4814837 | 7.41E-06 |
| ENSG00000267165.1 | 588.304133 | -1.455860147 | 0.32648679 | -4.4591701 | 8.23E-06 |
| ENSG00000279333.1 | 12.2418195 | -2.91981942 | 0.65672691 | -4.4460176 | 8.75E-06 |
| ENSG00000254902.1 | 3.24322809 | -4.605127015 | 1.03788123 | -4.4370463 | 9.12E-06 |
| ENSG00000259627.1 | 5527.26475 | -1.016020607 | 0.2294188 | -4.4286719 | 9.48E-06 |
| ENSG00000166676.15 | 159.265075 | -0.625171525 | 0.14133351 | -4.423378 | 9.72E-06 |
| ENSG00000172613.7 | 265.515419 | -0.563405418 | 0.12760548 | -4.4152132 | 1.01E-05 |
| ENSG00000162391.11 | 44.7235504 | -1.019809607 | 0.23126507 | -4.4097 | 1.04E-05 |
| ENSG00000171790.15 | 21.3574234 | -1.937147084 | 0.4402493 | -4.4001139 | 1.08E-05 |
| ENSG00000144355.14 | 5.82043466 | -2.087711907 | 0.47537405 | -4.3917246 | 1.12E-05 |
| ENSG00000196337.11 | 5.72499865 | -2.555613386 | 0.58380421 | -4.3775179 | 1.20E-05 |
| ENSG00000257605.2 | 115.626706 | -0.660385305 | 0.15129712 | -4.3648242 | 1.27E-05 |
| ENSG00000196923.13 | 30.2484606 | -1.186838451 | 0.27260476 | -4.3536967 | 1.34E-05 |
| ENSG00000165458.13 | 8.90155787 | -1.53890734 | 0.35537009 | -4.3304357 | 1.49E-05 |
| ENSG00000250186.3 | 702.170878 | -0.592135098 | 0.13687039 | -4.326247 | 1.52E-05 |
| ENSG00000236498.1 | 255.28547 | -0.543454946 | 0.12563203 | -4.3257676 | 1.52E-05 |
| ENSG00000181350.11 | 1364.26551 | -0.888307368 | 0.20546778 | -4.3233414 | 1.54E-05 |
| ENSG00000215375.6 | 513.567568 | -0.679496982 | 0.15747986 | -4.3148183 | 1.60E-05 |
| ENSG00000248890.1 | 12.037326 | -5.261959648 | 1.22751344 | -4.2866819 | 1.81E-05 |
| ENSG00000223390.1 | 44.8705216 | -0.86107377 | 0.20098186 | -4.2843357 | 1.83E-05 |
| ENSG00000086232.12 | 230.065234 | -0.573607195 | 0.13411057 | -4.2771214 | 1.89E-05 |
| ENSG00000279605.1 | 992.207397 | -0.583762431 | 0.13678422 | -4.2677615 | 1.97E-05 |
| ENSG00000285649.1 | 51.047398 | -0.665379841 | 0.15602021 | -4.2647028 | 2.00E-05 |

| | | | | | |
|---|---|---|---|---|---|
| ENSG00000107731.12 | 4.10810742 | -3.04681963 | 0.71578723 | -4.2565996 | 2.08E-05 |
| ENSG00000228109.1 | 8.03665089 | -1.713175588 | 0.40343846 | -4.246436 | 2.17E-05 |
| ENSG00000205485.13 | 81.4878397 | -1.019202187 | 0.24006666 | -4.2454967 | 2.18E-05 |
| ENSG00000101665.9 | 9.21079758 | -1.683201598 | 0.39891892 | -4.2194078 | 2.45E-05 |
| ENSG00000125753.13 | 17.0000899 | -1.187675446 | 0.28177383 | -4.2149956 | 2.50E-05 |
| ENSG00000006327.13 | 15.2712044 | -1.252345082 | 0.29766154 | -4.2072788 | 2.58E-05 |
| ENSG00000254721.1 | 404.24265 | -0.498367462 | 0.11846258 | -4.2069611 | 2.59E-05 |
| ENSG00000164022.16 | 24.8163121 | -1.425389882 | 0.33932948 | -4.2006072 | 2.66E-05 |
| ENSG00000124243.17 | 27.5128645 | -1.069574307 | 0.25470721 | -4.1992306 | 2.68E-05 |
| ENSG00000139637.13 | 23.5346904 | -1.094293997 | 0.26071357 | -4.1973036 | 2.70E-05 |
| ENSG00000169609.13 | 114.384662 | -0.625310633 | 0.14908555 | -4.1943075 | 2.74E-05 |
| ENSG00000107984.9 | 27.8262064 | -1.549937327 | 0.36982132 | -4.1910437 | 2.78E-05 |
| ENSG00000156050.8 | 287.745126 | -0.512983688 | 0.12241497 | -4.1905308 | 2.78E-05 |
| ENSG00000092931.11 | 1192.38992 | -0.536286284 | 0.12803126 | -4.1887135 | 2.81E-05 |
| ENSG00000243230.1 | 12.5551115 | -1.296375746 | 0.3098915 | -4.1833214 | 2.87E-05 |
| ENSG00000121310.16 | 3.95926401 | -2.836641424 | 0.67921054 | -4.1763802 | 2.96E-05 |
| ENSG00000267064.1 | 160.90513 | -0.844296213 | 0.20216716 | -4.1762284 | 2.96E-05 |
| ENSG00000214140.10 | 86.1704991 | -1.504224098 | 0.36084009 | -4.1686723 | 3.06E-05 |
| ENSG00000100292.16 | 22.7477813 | -1.338049834 | 0.32165046 | -4.15995 | 3.18E-05 |
| ENSG00000136942.14 | 18.3830134 | -1.192515244 | 0.28679265 | -4.1581094 | 3.21E-05 |
| ENSG00000285908.1 | 98.6099112 | -0.957732601 | 0.23041711 | -4.1565168 | 3.23E-05 |
| ENSG00000225407.3 | 6.83246074 | -1.613333645 | 0.39063627 | -4.130015 | 3.63E-05 |
| ENSG00000185112.5 | 1.77709894 | -3.419570567 | 0.82865137 | -4.1266698 | 3.68E-05 |
| ENSG00000270084.1 | 68.1479855 | -0.893996338 | 0.21669524 | -4.1255929 | 3.70E-05 |
| ENSG00000099904.15 | 12.8180805 | -1.914165537 | 0.46424186 | -4.1232076 | 3.74E-05 |
| ENSG00000260001.6 | 5.80334696 | -2.106288619 | 0.51106614 | -4.1213621 | 3.77E-05 |
| ENSG00000100345.21 | 318.985019 | -0.832303847 | 0.20312386 | -4.0975188 | 4.18E-05 |
| ENSG00000225950.8 | 23.7557997 | -2.23459391 | 0.54634799 | -4.0900561 | 4.31E-05 |
| ENSG00000079805.16 | 323.160178 | -0.736151232 | 0.18080723 | -4.0714701 | 4.67E-05 |
| ENSG00000260367.2 | 117.057565 | -0.574393177 | 0.14109135 | -4.0710731 | 4.68E-05 |
| ENSG00000137573.13 | 47.385751 | -2.086537746 | 0.51275682 | -4.069254 | 4.72E-05 |
| ENSG00000130176.7 | 4.53938924 | -2.821952375 | 0.69369704 | -4.0679897 | 4.74E-05 |
| ENSG00000266933.2 | 97.0187415 | -1.038125578 | 0.2555679 | -4.0620343 | 4.86E-05 |
| ENSG00000272182.1 | 393.100527 | -0.572983898 | 0.14111645 | -4.0603623 | 4.90E-05 |
| ENSG00000261762.1 | 4.27913385 | -3.453100058 | 0.8529829 | -4.0482641 | 5.16E-05 |
| ENSG00000126453.9 | 74.6361545 | -0.572574401 | 0.1417412 | -4.0395764 | 5.35E-05 |
| ENSG00000263272.1 | 61.4880556 | -0.839565648 | 0.20799907 | -4.0363913 | 5.43E-05 |
| ENSG00000221916.3 | 14.039303 | -1.147136437 | 0.28527498 | -4.0211604 | 5.79E-05 |
| ENSG00000109686.17 | 2726.56965 | -0.923409097 | 0.23020118 | -4.0113135 | 6.04E-05 |
| ENSG00000167157.10 | 13.6070465 | -1.52344588 | 0.38004531 | -4.0085901 | 6.11E-05 |
| ENSG00000125968.8 | 12.0463223 | -1.84668211 | 0.46186984 | -3.9982739 | 6.38E-05 |
| ENSG00000253368.3 | 10.7889837 | -1.526937125 | 0.38228035 | -3.9942862 | 6.49E-05 |
| ENSG00000161970.14 | 15.3188823 | -1.252691631 | 0.31392922 | -3.9903633 | 6.60E-05 |

| | | | | | |
|---|---|---|---|---|---|
| ENSG00000108474.16 | 7.26610324 | -1.99341076 | 0.5001991 | -3.9852346 | 6.74E-05 |
| ENSG00000170222.11 | 7.82596773 | -1.648367709 | 0.41355221 | -3.9858757 | 6.72E-05 |
| ENSG00000115484.14 | 4.69729223 | -1.998186837 | 0.5017948 | -3.9820796 | 6.83E-05 |
| ENSG00000034510.5 | 109.509851 | -0.904525325 | 0.22717318 | -3.9816555 | 6.84E-05 |
| ENSG00000154025.15 | 285.139198 | -1.11018259 | 0.27989269 | -3.966458 | 7.29E-05 |
| ENSG00000244165.1 | 208.565296 | -0.56506153 | 0.14244903 | -3.9667629 | 7.29E-05 |
| ENSG00000188522.14 | 8.36300217 | -1.500996569 | 0.37928511 | -3.9574361 | 7.58E-05 |
| ENSG00000170315.13 | 18.3173281 | -1.107325891 | 0.27989693 | -3.9561916 | 7.62E-05 |
| ENSG00000236671.8 | 2268.22872 | -1.253588492 | 0.31705737 | -3.9538222 | 7.69E-05 |
| ENSG00000275807.1 | 66.1957867 | -0.59439159 | 0.15040701 | -3.9518874 | 7.75E-05 |
| ENSG00000033011.12 | 86.9327405 | -0.553988416 | 0.14020531 | -3.9512655 | 7.77E-05 |
| ENSG00000176700.20 | 341.618129 | -0.353803084 | 0.08971415 | -3.9436708 | 8.02E-05 |
| ENSG00000105825.12 | 4.72823719 | -4.158189733 | 1.05599069 | -3.9377144 | 8.23E-05 |
| ENSG00000269621.1 | 124.250099 | -0.65359372 | 0.16610026 | -3.9349349 | 8.32E-05 |
| ENSG00000115486.11 | 601.489082 | -0.336393372 | 0.0856574 | -3.9271958 | 8.59E-05 |
| ENSG00000204577.11 | 13.098244 | -1.791995513 | 0.45671133 | -3.923694 | 8.72E-05 |
| ENSG00000171988.18 | 10.5112782 | -1.371853636 | 0.35047478 | -3.914272 | 9.07E-05 |
| ENSG00000230953.2 | 8.90700259 | -1.600090361 | 0.40933654 | -3.9089849 | 9.27E-05 |
| ENSG00000125741.4 | 19.6259158 | -0.957792808 | 0.24525131 | -3.9053524 | 9.41E-05 |
| ENSG00000233295.3 | 2.28752792 | -3.718111896 | 0.95378256 | -3.8982805 | 9.69E-05 |
| ENSG00000283849.1 | 12.7200913 | -1.23078689 | 0.3156979 | -3.8986224 | 9.67E-05 |
| ENSG00000197756.9 | 25.2471506 | -1.119519672 | 0.28720787 | -3.8979421 | 9.70E-05 |
| ENSG00000072110.13 | 46.0727824 | -0.990559865 | 0.25406467 | -3.8988493 | 9.67E-05 |
| ENSG00000263218.2 | 212.001411 | -0.631623527 | 0.16195416 | -3.9000143 | 9.62E-05 |
| ENSG00000128652.11 | 15.4743005 | -1.230180423 | 0.31580379 | -3.8953947 | 9.80E-05 |
| ENSG00000276578.1 | 71.3752667 | -1.044773093 | 0.26820118 | -3.8954829 | 9.80E-05 |
| ENSG00000224383.7 | 2.82807378 | -2.793046804 | 0.71731849 | -3.8937332 | 9.87E-05 |
| ENSG00000231345.3 | 12.9643133 | -2.691508367 | 0.69130029 | -3.8933997 | 9.88E-05 |
| ENSG00000125457.14 | 38.9650518 | -0.715515001 | 0.1841276 | -3.8859737 | 0.00010192 |
| ENSG00000122641.10 | 10.450059 | -2.771781237 | 0.71425766 | -3.8806462 | 0.00010418 |
| ENSG00000164776.9 | 497.585412 | -0.343403265 | 0.08856035 | -3.8776188 | 0.00010548 |
| ENSG00000232284.7 | 43.4490272 | -1.913636865 | 0.49673866 | -3.8524017 | 0.00011696 |
| ENSG00000159176.13 | 14.3226253 | -1.005717042 | 0.26166729 | -3.8434954 | 0.00012129 |
| ENSG00000254254.5 | 132.505398 | -3.349918529 | 0.87304366 | -3.8370573 | 0.00012452 |
| ENSG00000261312.1 | 76.7182077 | -0.590481191 | 0.15388062 | -3.8372681 | 0.00012441 |
| ENSG00000277182.1 | 4.94681489 | -2.051841169 | 0.53516981 | -3.8340002 | 0.00012608 |
| ENSG00000197982.13 | 54.4684783 | -0.734235905 | 0.19153984 | -3.8333326 | 0.00012642 |
| ENSG00000197457.9 | 3.87119937 | -2.559085635 | 0.6680335 | -3.8307744 | 0.00012774 |
| ENSG00000240567.1 | 24.8868536 | -0.862752266 | 0.22529874 | -3.82937 | 0.00012847 |
| ENSG00000063177.12 | 28.8239801 | -1.463350111 | 0.38272512 | -3.8235016 | 0.00013157 |
| ENSG00000235927.4 | 87.0556374 | -0.690824281 | 0.18116452 | -3.8132428 | 0.00013716 |
| ENSG00000179965.11 | 77.9788169 | -0.925130137 | 0.24282343 | -3.8098883 | 0.00013903 |
| ENSG00000014919.12 | 90.1325328 | -1.172066359 | 0.30862873 | -3.797658 | 0.00014607 |

| | | | | | |
|---|---|---|---|---|---|
| ENSG00000105662.15 | 6.53496365 | -3.038699178 | 0.80159262 | -3.7908273 | 0.00015015 |
| ENSG00000140988.15 | 58.1564938 | -1.094808206 | 0.28971172 | -3.7789573 | 0.00015749 |
| ENSG00000102977.14 | 5.74194405 | -1.939217868 | 0.51338996 | -3.7772805 | 0.00015855 |
| ENSG00000234771.3 | 52.3022875 | -0.750532265 | 0.19870692 | -3.7770817 | 0.00015868 |
| ENSG00000150967.17 | 596.579912 | -0.332826058 | 0.08812115 | -3.7769144 | 0.00015878 |
| ENSG00000265618.1 | 130.142323 | -0.622692266 | 0.16497079 | -3.7745606 | 0.00016029 |
| ENSG00000203472.3 | 2.44274829 | -3.840446937 | 1.01973829 | -3.7661104 | 0.00016581 |
| ENSG00000013016.15 | 2.55802196 | -2.795146845 | 0.74303242 | -3.7618101 | 0.00016869 |
| ENSG00000260192.2 | 14.3741927 | -1.358875749 | 0.36216798 | -3.7520593 | 0.00017539 |
| ENSG00000218336.8 | 24.63388 | -1.037578469 | 0.27774789 | -3.7356844 | 0.00018721 |
| ENSG00000252680.1 | 1.65602852 | -3.584442425 | 0.96025461 | -3.7328042 | 0.00018936 |
| ENSG00000265399.1 | 15.0690168 | -1.183319753 | 0.31749374 | -3.7270648 | 0.00019372 |
| ENSG00000245970.2 | 2420.55363 | -0.649483408 | 0.17427289 | -3.7268183 | 0.00019391 |
| ENSG00000236883.1 | 27.6217736 | -1.89463803 | 0.50891829 | -3.7228727 | 0.00019697 |
| ENSG00000271969.1 | 20.001582 | -0.894405846 | 0.24026616 | -3.7225628 | 0.00019721 |
| ENSG00000170561.12 | 3.0195968 | -2.299464639 | 0.62086474 | -3.7036483 | 0.00021252 |
| ENSG00000150787.7 | 1.79401242 | -3.757086653 | 1.01468019 | -3.7027299 | 0.00021329 |
| ENSG00000259409.1 | 28.3638337 | -1.650866304 | 0.44597756 | -3.7016802 | 0.00021418 |
| ENSG00000259006.1 | 52.1697736 | -1.080514595 | 0.29211604 | -3.6989225 | 0.00021652 |
| ENSG00000237945.7 | 202.968175 | -0.665053092 | 0.17979474 | -3.6989574 | 0.00021649 |
| ENSG00000136918.7 | 158.238135 | -0.957670653 | 0.2590514 | -3.6968365 | 0.0002183 |
| ENSG00000188846.13 | 21.3269755 | -0.947950567 | 0.2568605 | -3.6905268 | 0.00022379 |
| ENSG00000140400.16 | 9.7359985 | -1.195470093 | 0.32517094 | -3.6764358 | 0.00023652 |
| ENSG00000145703.15 | 2.72123811 | -2.939653451 | 0.80028007 | -3.6732809 | 0.00023946 |
| ENSG00000280414.1 | 20.421196 | -5.207887421 | 1.41881797 | -3.6705818 | 0.000242 |
| ENSG00000179348.11 | 10.9499681 | -1.62340491 | 0.44252793 | -3.6684801 | 0.000244 |
| ENSG00000258908.1 | 274.335789 | -1.117502545 | 0.30476066 | -3.6668202 | 0.00024559 |
| ENSG00000033800.13 | 22.8267411 | -1.076140589 | 0.29388301 | -3.6617992 | 0.00025045 |
| ENSG00000278259.4 | 68.6827213 | -0.60873315 | 0.16635691 | -3.6591998 | 0.000253 |
| ENSG00000197948.10 | 87.1394424 | -0.481577456 | 0.13164709 | -3.6580941 | 0.0002541 |
| ENSG00000233893.2 | 150.662701 | -0.924935943 | 0.25290915 | -3.6571866 | 0.000255 |
| ENSG00000143631.10 | 113.786629 | -2.26300763 | 0.61906515 | -3.6555242 | 0.00025666 |
| ENSG00000188976.10 | 206.342439 | -1.67124685 | 0.45761985 | -3.6520419 | 0.00026016 |
| ENSG00000186710.11 | 269.318996 | -0.396254347 | 0.10875766 | -3.6434616 | 0.000269 |
| ENSG00000255508.7 | 34.8751999 | -0.695334157 | 0.19109055 | -3.6387679 | 0.00027395 |
| ENSG00000254418.1 | 25.9833351 | -3.252323509 | 0.89472683 | -3.6349905 | 0.00027799 |
| ENSG00000113273.16 | 10.021566 | -1.114028952 | 0.30698748 | -3.6289068 | 0.00028462 |
| ENSG00000257663.1 | 384.432118 | -0.44129577 | 0.12172079 | -3.6254756 | 0.00028843 |
| ENSG00000131871.14 | 1.65135367 | -3.271898087 | 0.90462292 | -3.616864 | 0.00029819 |
| ENSG00000104892.16 | 171.430904 | -0.62664592 | 0.17338267 | -3.6142362 | 0.00030123 |
| ENSG00000103145.10 | 42.9782135 | -0.61666874 | 0.17066358 | -3.6133587 | 0.00030226 |
| ENSG00000123144.10 | 23.765833 | -1.166709501 | 0.32443576 | -3.5961187 | 0.000323 |
| ENSG00000090061.17 | 44.6507178 | -1.029328523 | 0.28668752 | -3.5904198 | 0.00033015 |

| | | | | | |
|---|---|---|---|---|---|
| ENSG00000182993.4 | 36.4951218 | -3.734898175 | 1.04045225 | -3.5896872 | 0.00033107 |
| ENSG00000267342.1 | 83.9216803 | -0.612999508 | 0.17127118 | -3.5791165 | 0.00034476 |
| ENSG00000257553.1 | 473.951445 | -0.406125084 | 0.11345028 | -3.5797627 | 0.00034391 |
| ENSG00000101082.13 | 163.119429 | -0.48933416 | 0.13693812 | -3.5733962 | 0.00035238 |
| ENSG00000115461.4 | 26.3380073 | -2.113623187 | 0.59212596 | -3.5695499 | 0.0003576 |
| ENSG00000064601.18 | 49.7479262 | -0.503232438 | 0.14105443 | -3.5676473 | 0.0003602 |
| ENSG00000141552.17 | 80.0398994 | -0.500702223 | 0.14068639 | -3.5589955 | 0.00037228 |
| ENSG00000083845.8 | 14.6997167 | -1.34595841 | 0.37824611 | -3.5584197 | 0.00037309 |
| ENSG00000253356.1 | 342.306568 | -0.544110922 | 0.15354455 | -3.543668 | 0.0003946 |
| ENSG00000264577.1 | 2527.96856 | -0.433744341 | 0.12248757 | -3.5411295 | 0.00039842 |
| ENSG00000127528.5 | 3.78019837 | -2.107911103 | 0.5972201 | -3.5295381 | 0.00041629 |
| ENSG00000265148.5 | 374.565005 | -0.493412073 | 0.13997929 | -3.5248935 | 0.00042365 |
| ENSG00000166140.17 | 25.1282462 | -0.73526219 | 0.2087127 | -3.5228435 | 0.00042694 |
| ENSG00000121897.14 | 441.969717 | -1.133380726 | 0.32209986 | -3.5187247 | 0.00043363 |
| ENSG00000154277.12 | 3.19647801 | -2.495571533 | 0.7094702 | -3.5175143 | 0.00043561 |
| ENSG00000182463.15 | 2.18151839 | -3.281987559 | 0.93321583 | -3.516858 | 0.00043669 |
| ENSG00000187514.16 | 48.9692809 | -0.949409942 | 0.27008237 | -3.5152607 | 0.00043932 |
| ENSG00000111752.10 | 235.778413 | -0.574738519 | 0.16347711 | -3.5157124 | 0.00043858 |
| ENSG00000106624.10 | 178.900497 | -0.846394629 | 0.24086057 | -3.514044 | 0.00044134 |
| ENSG00000182325.10 | 295.688032 | -0.447178375 | 0.12735371 | -3.5113102 | 0.0004459 |
| ENSG00000164587.12 | 21.6434426 | -1.080549958 | 0.30796724 | -3.5086523 | 0.00045038 |
| ENSG00000238279.1 | 37.4673101 | -0.622255017 | 0.17746279 | -3.5063972 | 0.00045422 |
| ENSG00000270075.1 | 162.512578 | -0.741977431 | 0.21190478 | -3.5014662 | 0.00046271 |
| ENSG00000161981.10 | 44.749881 | -0.878773422 | 0.25102793 | -3.5006997 | 0.00046404 |
| ENSG00000260051.1 | 16.4244948 | -1.504212354 | 0.43073122 | -3.4922297 | 0.00047901 |
| ENSG00000234166.1 | 23.8161419 | -1.040879957 | 0.29832424 | -3.4890894 | 0.00048467 |
| ENSG00000226445.1 | 12.6402578 | -1.541428822 | 0.44269375 | -3.4819304 | 0.00049781 |
| ENSG00000150051.13 | 6.05040568 | -1.670532648 | 0.48003045 | -3.4800556 | 0.00050131 |
| ENSG00000223764.2 | 22.1102243 | -1.723288026 | 0.49632798 | -3.4720751 | 0.00051645 |
| ENSG00000272418.1 | 13.8966336 | -1.083199938 | 0.31236257 | -3.4677648 | 0.00052481 |
| ENSG00000269275.1 | 2.79344025 | -2.962686018 | 0.85679234 | -3.4578811 | 0.00054444 |
| ENSG00000142657.20 | 4.3696037 | -1.824457029 | 0.52743572 | -3.4591079 | 0.00054197 |
| ENSG00000170017.12 | 18.0514367 | -1.170588353 | 0.33846972 | -3.4584728 | 0.00054325 |
| ENSG00000186166.8 | 171.843253 | -0.604875553 | 0.17486002 | -3.4591987 | 0.00054179 |
| ENSG00000141736.13 | 258.000723 | -0.574549516 | 0.16602413 | -3.4606386 | 0.0005389 |
| ENSG00000100316.15 | 46.0277023 | -0.703750566 | 0.20376373 | -3.4537578 | 0.00055283 |
| ENSG00000139514.12 | 15.265848 | -1.348240123 | 0.39052321 | -3.4523944 | 0.00055563 |
| ENSG00000175137.10 | 7.96432103 | -1.435053861 | 0.41643961 | -3.4460071 | 0.00056894 |
| ENSG00000272088.1 | 5.48231568 | -1.640181101 | 0.47607226 | -3.4452356 | 0.00057056 |
| ENSG00000103855.17 | 10.2679503 | -1.475864765 | 0.42872535 | -3.4424481 | 0.00057647 |
| ENSG00000259322.1 | 70.1431952 | -0.58956983 | 0.17131298 | -3.4414778 | 0.00057855 |
| ENSG00000142453.11 | 389.411085 | -0.502428505 | 0.14593943 | -3.4427193 | 0.0005759 |
| ENSG00000005075.15 | 12.7128232 | -0.94877271 | 0.27585003 | -3.4394512 | 0.00058289 |

| | | | | | |
|---|---|---|---|---|---|
| ENSG00000236782.7 | 159.581519 | -0.898515592 | 0.26135201 | -3.4379517 | 0.00058613 |
| ENSG00000137135.17 | 321.554795 | -0.453449498 | 0.13196997 | -3.4360051 | 0.00059036 |
| ENSG00000084207.16 | 5.18226531 | -1.967421062 | 0.57399351 | -3.4276016 | 0.00060894 |
| ENSG00000279762.3 | 101.724007 | -0.603750623 | 0.17612552 | -3.4279566 | 0.00060814 |
| ENSG00000105248.15 | 1.71275759 | -2.952431413 | 0.86312673 | -3.4206233 | 0.00062478 |
| ENSG00000265558.1 | 8.95173582 | -1.678632422 | 0.49101732 | -3.4186827 | 0.00062925 |
| ENSG00000250838.1 | 1.82531437 | -2.919605779 | 0.85420149 | -3.4179357 | 0.00063098 |
| ENSG00000140553.17 | 76.0977377 | -0.505686019 | 0.14799204 | -3.4169813 | 0.0006332 |
| ENSG00000274897.2 | 12.8289374 | -1.252585242 | 0.36678914 | -3.4150009 | 0.00063782 |
| ENSG00000019144.18 | 12.9914247 | -1.014540333 | 0.29714828 | -3.4142561 | 0.00063956 |
| ENSG00000120438.11 | 130.053557 | -0.938488762 | 0.27548077 | -3.4067306 | 0.00065746 |
| ENSG00000269888.1 | 48.4615139 | -0.826450879 | 0.24289617 | -3.4024862 | 0.00066776 |
| ENSG00000175414.6 | 107.094957 | -0.559839562 | 0.16461347 | -3.4009341 | 0.00067156 |
| ENSG00000104894.11 | 130.905191 | -0.647212233 | 0.19046693 | -3.3980295 | 0.00067873 |
| ENSG00000274425.1 | 160.791232 | -0.492807735 | 0.14503722 | -3.3978019 | 0.0006793 |
| ENSG00000012061.15 | 230.382192 | -0.638960001 | 0.18842517 | -3.3910545 | 0.00069624 |
| ENSG00000157240.3 | 7.03251286 | -1.443236613 | 0.42608961 | -3.3871669 | 0.00070618 |
| ENSG00000260350.1 | 322.091717 | -0.443903634 | 0.13106429 | -3.3869151 | 0.00070683 |
| ENSG00000204623.9 | 110.289692 | -0.744857734 | 0.2199585 | -3.3863557 | 0.00070828 |
| ENSG00000255114.1 | 575.869805 | -0.444320156 | 0.13123747 | -3.3856197 | 0.00071018 |
| ENSG00000251003.8 | 1.61204839 | -3.587980934 | 1.06004676 | -3.3847384 | 0.00071246 |
| ENSG00000125434.10 | 199.827924 | -0.576072564 | 0.1703607 | -3.3814873 | 0.00072095 |
| ENSG00000164530.14 | 3.26696174 | -3.919788011 | 1.15967618 | -3.3800712 | 0.00072467 |
| ENSG00000258634.3 | 3.26125883 | -3.960296402 | 1.17776457 | -3.3625535 | 0.00077225 |
| ENSG00000127314.17 | 9.6292882 | -1.916187403 | 0.571044 | -3.3555863 | 0.00079197 |
| ENSG00000103257.8 | 25.828853 | -1.524457578 | 0.45448014 | -3.3542887 | 0.00079569 |
| ENSG00000234925.2 | 5.65483379 | -1.510577832 | 0.45096202 | -3.3496786 | 0.00080905 |
| ENSG00000161179.13 | 7.28866504 | -1.506347437 | 0.45020253 | -3.3459329 | 0.00082006 |
| ENSG00000104960.15 | 137.56335 | -1.064722462 | 0.31815346 | -3.3465689 | 0.00081818 |
| ENSG00000204620.3 | 216.136588 | -0.757782185 | 0.22657345 | -3.3445321 | 0.00082421 |
| ENSG00000240211.1 | 7.79291246 | -1.844733305 | 0.55182406 | -3.3429737 | 0.00082886 |
| ENSG00000115844.10 | 1.90042713 | -2.828918453 | 0.84836224 | -3.3345643 | 0.00085433 |
| ENSG00000087302.8 | 354.170885 | -1.034495663 | 0.31040324 | -3.3327476 | 0.00085993 |
| ENSG00000234745.10 | 13.8546822 | -1.219040044 | 0.36642945 | -3.326807 | 0.00087847 |
| ENSG00000285190.1 | 65.5479167 | -0.611366358 | 0.18373324 | -3.3274674 | 0.00087639 |
| ENSG00000182718.16 | 63.1695022 | -0.605089626 | 0.18218904 | -3.3212186 | 0.00089625 |
| ENSG00000173065.13 | 63.5829674 | -0.493494114 | 0.14872873 | -3.318082 | 0.00090638 |
| ENSG00000279488.1 | 109.382463 | -0.381759613 | 0.11509218 | -3.3169899 | 0.00090993 |
| ENSG00000142552.7 | 51.6435855 | -0.904869447 | 0.2728553 | -3.3162979 | 0.00091219 |
| ENSG00000162244.11 | 20.9175705 | -0.967042647 | 0.29197038 | -3.3121259 | 0.0009259 |
| ENSG00000184162.14 | 66.5519346 | -0.601549873 | 0.18169475 | -3.3107719 | 0.00093039 |
| ENSG00000277290.1 | 8.32789475 | -1.689924816 | 0.51050662 | -3.3102897 | 0.00093199 |
| ENSG00000265840.1 | 67.792809 | -0.525435619 | 0.15918556 | -3.3007744 | 0.00096418 |

| | | | | | |
|---|---|---|---|---|---|
| ENSG00000100097.11 | 40.2682994 | -0.971506475 | 0.29466828 | -3.2969496 | 0.00097741 |
| ENSG00000135439.11 | 424.450822 | -0.384510535 | 0.11670624 | -3.294687 | 0.00098531 |
| ENSG00000142541.16 | 43.3310679 | -0.771124361 | 0.23525848 | -3.277775 | 0.00104629 |
| ENSG00000198113.2 | 1.68692942 | -2.939875888 | 0.89739142 | -3.2760241 | 0.0010528 |
| ENSG00000161016.17 | 28.8595601 | -1.001957225 | 0.30685632 | -3.2652324 | 0.00109374 |
| ENSG00000079308.18 | 27.9101328 | -0.918107502 | 0.28114975 | -3.2655462 | 0.00109253 |
| ENSG00000160957.12 | 27.8593163 | -0.878075376 | 0.26905371 | -3.2635691 | 0.00110018 |
| ENSG00000269176.2 | 52.1563196 | -0.68212271 | 0.20899241 | -3.2638636 | 0.00109904 |
| ENSG00000124766.6 | 5.12335283 | -1.637202628 | 0.50255774 | -3.2577403 | 0.00112303 |
| ENSG00000256690.1 | 89.672377 | -0.436641339 | 0.13401711 | -3.2581015 | 0.0011216 |
| ENSG00000253125.1 | 11.5015179 | -2.033681956 | 0.62453876 | -3.2562942 | 0.00112877 |
| ENSG00000117385.15 | 7.84037786 | -1.219350192 | 0.37491489 | -3.2523387 | 0.0011446 |
| ENSG00000106246.17 | 498.063981 | -0.365611165 | 0.11268206 | -3.2446262 | 0.00117605 |
| ENSG00000107957.16 | 41.976063 | -0.834835487 | 0.25763845 | -3.2403374 | 0.00119388 |
| ENSG00000133740.10 | 200.24951 | -0.497094277 | 0.15354863 | -3.2373735 | 0.00120635 |
| ENSG00000170113.15 | 4.18218038 | -2.182304501 | 0.67454727 | -3.2352136 | 0.00121552 |
| ENSG00000163082.9 | 8.65055024 | -1.506845916 | 0.46583128 | -3.2347461 | 0.00121751 |
| ENSG00000107223.12 | 23.7588255 | -0.908036418 | 0.28114786 | -3.2297468 | 0.001239 |
| ENSG00000280721.1 | 346.897202 | -0.537469831 | 0.1664679 | -3.2286696 | 0.00124368 |
| ENSG00000149260.16 | 2.85172712 | -2.470449257 | 0.76549115 | -3.2272735 | 0.00124976 |
| ENSG00000285278.1 | 61.3588282 | -0.9686626 | 0.30048665 | -3.223646 | 0.0012657 |
| ENSG00000170365.9 | 3.11190243 | -2.448181841 | 0.76131568 | -3.215725 | 0.00130115 |
| ENSG00000176383.8 | 377.252612 | -0.384498531 | 0.11966149 | -3.2132188 | 0.00131256 |
| ENSG00000233360.4 | 33.9857701 | -0.803978228 | 0.25045291 | -3.2100974 | 0.0013269 |
| ENSG00000187185.4 | 7.91652224 | -1.342334405 | 0.41827396 | -3.2092229 | 0.00133094 |
| ENSG00000145632.14 | 1.490934 | -2.810478008 | 0.8772922 | -3.2035826 | 0.00135729 |
| ENSG00000105372.7 | 31.136532 | -0.945747355 | 0.29527286 | -3.2029607 | 0.00136023 |
| ENSG00000137124.7 | 2.24552213 | -2.306809419 | 0.72031027 | -3.2025219 | 0.0013623 |
| ENSG00000140992.18 | 15.1513025 | -1.474634391 | 0.46077194 | -3.2003563 | 0.00137258 |
| ENSG00000273621.1 | 1.55328515 | -3.930275282 | 1.22845987 | -3.1993518 | 0.00137737 |
| ENSG00000244151.1 | 705.141607 | -0.37565219 | 0.11745911 | -3.1981529 | 0.00138311 |
| ENSG00000178297.13 | 74.008542 | -0.646676428 | 0.20225943 | -3.1972622 | 0.00138739 |
| ENSG00000262791.1 | 83.5822118 | -0.480567007 | 0.15030293 | -3.197323 | 0.0013871 |
| ENSG00000070404.9 | 6.97448961 | -1.214840992 | 0.38020669 | -3.195212 | 0.00139728 |
| ENSG00000115317.11 | 337.566396 | -0.80074881 | 0.25105467 | -3.1895396 | 0.001425 |
| ENSG00000165916.8 | 8.83749257 | -1.116184503 | 0.35002823 | -3.1888414 | 0.00142844 |
| ENSG00000163806.15 | 34.9825899 | -0.615736821 | 0.19330328 | -3.1853409 | 0.00144584 |
| ENSG00000282556.2 | 1296.99763 | -0.572039641 | 0.17975094 | -3.1824015 | 0.00146059 |
| ENSG00000187634.11 | 233.891948 | -0.544708627 | 0.17149279 | -3.1762772 | 0.00149178 |
| ENSG00000168028.13 | 3.61087597 | -1.737014221 | 0.54804522 | -3.1694725 | 0.00152716 |
| ENSG00000006015.17 | 62.0669019 | -3.585896777 | 1.13344438 | -3.1637166 | 0.00155768 |
| ENSG00000051523.10 | 14.9549832 | -1.409508827 | 0.44582273 | -3.1615903 | 0.0015691 |
| ENSG00000142937.11 | 43.9304658 | -1.138521473 | 0.36018366 | -3.160947 | 0.00157257 |

| | | | | | |
|---|---|---|---|---|---|
| ENSG00000236886.2 | 339.270569 | -1.884320981 | 0.59659408 | -3.1584641 | 0.00158603 |
| ENSG00000169926.10 | 4.12885238 | -1.788133834 | 0.56633093 | -3.157401 | 0.00159182 |
| ENSG00000167264.17 | 12.3152013 | -0.926596975 | 0.29373314 | -3.1545537 | 0.00160744 |
| ENSG00000273149.1 | 11097.3912 | -0.639124925 | 0.20258856 | -3.1547928 | 0.00160612 |
| ENSG00000112578.9 | 2.03058764 | -2.577065216 | 0.81897243 | -3.1467057 | 0.00165121 |
| ENSG00000266969.1 | 48.8476583 | -0.649168276 | 0.20644699 | -3.1444793 | 0.00166383 |
| ENSG00000259954.1 | 47.8869687 | -1.311599819 | 0.41886835 | -3.1312937 | 0.00174038 |
| ENSG00000227766.1 | 2002.20722 | -0.662573466 | 0.21181639 | -3.1280557 | 0.00175967 |
| ENSG00000240889.1 | 245.977405 | -0.654258944 | 0.20928158 | -3.1262137 | 0.00177073 |
| ENSG00000267112.1 | 10.0987492 | -2.490140838 | 0.79718508 | -3.1236671 | 0.00178612 |
| ENSG00000236358.1 | 5.25650505 | -1.541948972 | 0.49396202 | -3.1215942 | 0.00179875 |
| ENSG00000235481.2 | 9.69369357 | -1.567539114 | 0.50236371 | -3.1203272 | 0.0018065 |
| ENSG00000198198.16 | 223.78361 | -0.492981567 | 0.1581462 | -3.117252 | 0.00182545 |
| ENSG00000106785.14 | 336.476227 | -0.496836357 | 0.15942673 | -3.116393 | 0.00183078 |
| ENSG00000210195.2 | 6.59777835 | -1.493612437 | 0.47937977 | -3.1157185 | 0.00183497 |
| ENSG00000075624.14 | 382.714834 | -0.591975132 | 0.19015514 | -3.1131167 | 0.00185123 |
| ENSG00000205542.10 | 74.0406953 | -0.683297371 | 0.21960292 | -3.1115132 | 0.00186131 |
| ENSG00000125148.6 | 5.71294438 | -2.057870082 | 0.66245379 | -3.1064357 | 0.00189358 |
| ENSG00000210112.1 | 10.0117139 | -1.47682103 | 0.47565152 | -3.1048382 | 0.00190383 |
| ENSG00000085511.19 | 22.1039164 | -1.604562778 | 0.51689592 | -3.104228 | 0.00190776 |
| ENSG00000239665.8 | 181.354748 | -0.972607389 | 0.31360306 | -3.1013964 | 0.0019261 |
| ENSG00000183765.21 | 22.8774092 | -0.899835554 | 0.29015356 | -3.1012391 | 0.00192713 |
| ENSG00000164404.8 | 312.518957 | -0.818071 | 0.26400165 | -3.0987344 | 0.00194349 |
| ENSG00000136699.19 | 658.736852 | -0.655769909 | 0.21166082 | -3.0982111 | 0.00194693 |
| ENSG00000173141.4 | 1.68008847 | -2.611916423 | 0.84444328 | -3.0930632 | 0.00198102 |
| ENSG00000255112.2 | 4.60467465 | -1.960490109 | 0.63422089 | -3.0911787 | 0.00199364 |
| ENSG00000176915.14 | 5.0125171 | -1.370328393 | 0.44363433 | -3.0888691 | 0.0020092 |
| ENSG00000249996.1 | 109.294555 | -0.711286796 | 0.23032087 | -3.0882429 | 0.00201344 |
| ENSG00000231607.10 | 23.5418165 | -0.742412403 | 0.24060623 | -3.085591 | 0.00203148 |
| ENSG00000136235.16 | 6.16956132 | -1.435560872 | 0.46558571 | -3.0833439 | 0.00204688 |
| ENSG00000267383.6 | 16.9801965 | -0.821126507 | 0.26633276 | -3.0830849 | 0.00204867 |
| ENSG00000257702.3 | 34.5966182 | -0.775965395 | 0.2516485 | -3.0835287 | 0.00204561 |
| ENSG00000157227.12 | 132.431655 | -0.905898093 | 0.29389742 | -3.0823615 | 0.00205365 |
| ENSG00000188312.13 | 208.067517 | -0.380610416 | 0.12352064 | -3.0813507 | 0.00206064 |
| ENSG00000159761.14 | 11.2671019 | -1.006485545 | 0.32689416 | -3.078934 | 0.00207743 |
| ENSG00000110237.4 | 22.6853964 | -0.921326317 | 0.29919714 | -3.0793286 | 0.00207468 |
| ENSG00000112096.17 | 39.5239983 | -0.690821969 | 0.22438101 | -3.0787899 | 0.00207843 |
| ENSG00000166441.12 | 75.7657763 | -0.557685247 | 0.18148968 | -3.0728207 | 0.00212046 |
| ENSG00000225721.5 | 874.531295 | -0.770591779 | 0.25088596 | -3.0714823 | 0.00212999 |
| ENSG00000181409.12 | 3.67227747 | -2.567497711 | 0.83642329 | -3.0696153 | 0.00214335 |
| ENSG00000227644.2 | 16.696343 | -1.137601756 | 0.3709808 | -3.0664707 | 0.00216602 |
| ENSG00000183431.11 | 2.84411508 | -2.19713183 | 0.7167781 | -3.0652887 | 0.0021746 |
| ENSG00000153885.14 | 3.25252076 | -1.763050485 | 0.57659522 | -3.0576918 | 0.00223049 |

| | | | | | |
|---|---|---|---|---|---|
| ENSG00000134013.15 | 49.0753274 | -0.744675104 | 0.24355824 | -3.0574827 | 0.00223205 |
| ENSG00000257042.1 | 63.361936 | -2.670077361 | 0.87387427 | -3.055448 | 0.00224725 |
| ENSG00000215866.7 | 2.92831247 | -1.846134378 | 0.60430794 | -3.0549563 | 0.00225093 |
| ENSG00000229950.1 | 8.59200275 | -1.191299237 | 0.39005031 | -3.0542194 | 0.00225647 |
| ENSG00000108592.16 | 528.02067 | -0.267627127 | 0.08765541 | -3.0531728 | 0.00226435 |
| ENSG00000170881.4 | 2.27593215 | -2.647803438 | 0.86746232 | -3.0523556 | 0.00227053 |
| ENSG00000149927.17 | 200.195143 | -0.35442357 | 0.11616619 | -3.0510044 | 0.00228077 |
| ENSG00000159267.14 | 3.39122891 | -1.884923081 | 0.61886527 | -3.0457729 | 0.00232083 |
| ENSG00000285932.1 | 39.8964816 | -0.876929985 | 0.28801067 | -3.0447829 | 0.00232848 |
| ENSG00000173545.4 | 2.96618797 | -1.775164115 | 0.583356 | -3.0430202 | 0.00234217 |
| ENSG00000104529.17 | 75.1550253 | -0.527322285 | 0.17360755 | -3.0374386 | 0.00238598 |
| ENSG00000198755.10 | 11.042606 | -1.269495264 | 0.41827932 | -3.0350419 | 0.00240502 |
| ENSG00000269559.2 | 10.4272981 | -1.07963717 | 0.3560844 | -3.0319699 | 0.00242963 |
| ENSG00000140199.11 | 217.375084 | -0.527627971 | 0.17404787 | -3.0315107 | 0.00243333 |
| ENSG00000210140.1 | 12.336802 | -1.273328526 | 0.42097201 | -3.0247344 | 0.00248852 |
| ENSG00000168569.7 | 306.10028 | -0.483276305 | 0.1600437 | -3.0196521 | 0.00253065 |
| ENSG00000129235.10 | 65.2469691 | -0.539154919 | 0.17892121 | -3.013365 | 0.00258368 |
| ENSG00000248710.1 | 9.01265334 | -2.57842305 | 0.856117 | -3.0117648 | 0.00259734 |
| ENSG00000188760.10 | 41.5917002 | -0.953279686 | 0.31674796 | -3.0095843 | 0.00261605 |
| ENSG00000108821.13 | 1996.75387 | -0.819636251 | 0.27275051 | -3.0050769 | 0.00265514 |
| ENSG00000248923.1 | 42.4534748 | -0.637954461 | 0.21227892 | -3.0052653 | 0.00265349 |
| ENSG00000231500.6 | 20.6583058 | -0.792364988 | 0.26379032 | -3.0037682 | 0.00266658 |
| ENSG00000172757.12 | 100.580457 | -0.434109823 | 0.14455198 | -3.0031398 | 0.0026721 |
| ENSG00000271806.1 | 3.58653255 | -2.009582292 | 0.6694634 | -3.0017807 | 0.00268405 |
| ENSG00000225969.2 | 9.68154915 | -1.024322116 | 0.34161961 | -2.9984289 | 0.00271375 |
| ENSG00000009790.14 | 37.6534864 | -1.126734957 | 0.3758725 | -2.997652 | 0.00272068 |
| ENSG00000178980.14 | 3.31703763 | -1.536606541 | 0.51334146 | -2.993342 | 0.0027594 |
| ENSG00000105193.8 | 9.10927571 | -1.280297797 | 0.42774525 | -2.9931315 | 0.00276131 |
| ENSG00000180104.15 | 99.4655585 | -0.486481263 | 0.1628454 | -2.9873811 | 0.00281379 |
| ENSG00000168061.14 | 2.22008782 | -2.191296967 | 0.73392906 | -2.9857068 | 0.00282924 |
| ENSG00000148835.10 | 59.0162432 | -0.472576549 | 0.15835051 | -2.9843701 | 0.00284163 |
| ENSG00000214253.8 | 6.52525877 | -1.34041863 | 0.44934045 | -2.9830803 | 0.00285363 |
| ENSG00000148848.14 | 10.6193733 | -1.173920556 | 0.39414224 | -2.9784185 | 0.0028974 |
| ENSG00000150455.13 | 3.34393475 | -1.73732297 | 0.58372616 | -2.9762637 | 0.00291784 |
| ENSG00000189060.5 | 40.7666735 | -0.794961202 | 0.26706193 | -2.9766923 | 0.00291376 |
| ENSG00000147140.15 | 19.2603551 | -0.64322926 | 0.21611378 | -2.9763454 | 0.00291706 |
| ENSG00000175115.11 | 12.123188 | -1.005544608 | 0.33818347 | -2.9733701 | 0.00294549 |
| ENSG00000089693.10 | 17.0494684 | -0.80766032 | 0.27158818 | -2.9738419 | 0.00294097 |
| ENSG00000165280.15 | 13.5498092 | -0.799206795 | 0.26873951 | -2.9739089 | 0.00294032 |
| ENSG00000138074.14 | 264.287927 | -0.604296347 | 0.20326475 | -2.972952 | 0.00294951 |
| ENSG00000103148.15 | 413.152148 | -0.569131057 | 0.19141583 | -2.9732706 | 0.00294645 |
| ENSG00000105373.18 | 15.0871029 | -1.117458014 | 0.37618669 | -2.9704879 | 0.00297327 |
| ENSG00000233766.7 | 61.6385157 | -2.268038453 | 0.76382917 | -2.9693007 | 0.00298478 |

| | | | | | |
|---|---|---|---|---|---|
| ENSG00000111665.11 | 30.0424563 | -0.926223306 | 0.31199587 | -2.9687037 | 0.00299059 |
| ENSG00000111737.11 | 26.3672692 | -0.676999704 | 0.22813006 | -2.9676041 | 0.00300131 |
| ENSG00000130255.12 | 71.4319226 | -0.614649743 | 0.20722657 | -2.9660759 | 0.00301626 |
| ENSG00000185475.10 | 41.8925098 | -0.434594826 | 0.14653712 | -2.9657662 | 0.0030193 |
| ENSG00000229117.8 | 19.8560428 | -0.841818898 | 0.28469924 | -2.9568709 | 0.00310778 |
| ENSG00000149273.14 | 15.4433011 | -1.169572182 | 0.39560424 | -2.9564197 | 0.00311233 |
| ENSG00000172922.9 | 532.889172 | -0.295170674 | 0.09985858 | -2.9558869 | 0.00311771 |
| ENSG00000166337.9 | 2212.71367 | -0.353839302 | 0.11975374 | -2.9547243 | 0.00312949 |
| ENSG00000168528.11 | 2.26239399 | -2.378487141 | 0.80520453 | -2.9538919 | 0.00313794 |
| ENSG00000042753.11 | 10.9319041 | -1.134076803 | 0.38394152 | -2.9537749 | 0.00313913 |
| ENSG00000252423.1 | 1.35426686 | -2.896625276 | 0.98144078 | -2.951401 | 0.00316336 |
| ENSG00000226754.1 | 15.995641 | -1.566700288 | 0.53085473 | -2.9512788 | 0.00316461 |
| ENSG00000204356.13 | 11.4447024 | -0.990759488 | 0.33569258 | -2.9513893 | 0.00316348 |
| ENSG00000249695.6 | 1.64137023 | -3.179014658 | 1.07809361 | -2.9487371 | 0.00319075 |
| ENSG00000237009.2 | 23.2220843 | -1.040262819 | 0.35281763 | -2.9484434 | 0.00319379 |
| ENSG00000261546.1 | 33.6684959 | -0.605885036 | 0.20570897 | -2.9453506 | 0.00322589 |
| ENSG00000278811.4 | 54.7374231 | -0.675353825 | 0.22957961 | -2.9416977 | 0.00326418 |
| ENSG00000006118.14 | 2.71179759 | -1.977443418 | 0.67249921 | -2.9404398 | 0.00327747 |
| ENSG00000142676.14 | 14.9062379 | -0.808186107 | 0.27496016 | -2.9392844 | 0.00328971 |
| ENSG00000273212.1 | 294.889132 | -0.561835881 | 0.1911866 | -2.9386781 | 0.00329615 |
| ENSG00000231187.2 | 1.84915379 | -3.106643623 | 1.0582641 | -2.9356033 | 0.003329 |
| ENSG00000234665.8 | 4.74179542 | -1.509684947 | 0.51444221 | -2.9346055 | 0.00333972 |
| ENSG00000267125.2 | 156.491727 | -0.452999956 | 0.15441932 | -2.9335704 | 0.00335088 |
| ENSG00000128245.14 | 3.33426543 | -2.022602683 | 0.68982375 | -2.9320572 | 0.00336725 |
| ENSG00000092068.19 | 17.5678394 | -1.674501549 | 0.57110362 | -2.9320451 | 0.00336738 |
| ENSG00000265069.1 | 12.0762502 | -1.052190924 | 0.35888756 | -2.9318122 | 0.0033699 |
| ENSG00000121764.11 | 110.440113 | -0.565622747 | 0.19339351 | -2.9247246 | 0.00344761 |
| ENSG00000163322.13 | 157.439352 | -0.522586686 | 0.17871383 | -2.9241535 | 0.00345394 |
| ENSG00000054965.10 | 31.8007665 | -0.882610878 | 0.30197122 | -2.9228311 | 0.00346865 |
| ENSG00000260774.1 | 3.40688404 | -1.994949124 | 0.68291233 | -2.9212375 | 0.00348644 |
| ENSG00000182831.11 | 7.06135952 | -1.167986397 | 0.39980275 | -2.9214066 | 0.00348455 |
| ENSG00000263050.1 | 12.3001239 | -0.885768776 | 0.30316695 | -2.9217194 | 0.00348105 |
| ENSG00000243710.7 | 111.670341 | -0.585993999 | 0.2006761 | -2.9200986 | 0.00349921 |
| ENSG00000162032.15 | 70.4360635 | -0.577261657 | 0.19773535 | -2.9193651 | 0.00350745 |
| ENSG00000050820.16 | 16.6822871 | -0.719694055 | 0.24687829 | -2.9151776 | 0.00355486 |
| ENSG00000143994.13 | 30.9433734 | -0.55180705 | 0.18925417 | -2.915693 | 0.003549 |
| ENSG00000095066.11 | 267.12986 | -0.540397719 | 0.18536089 | -2.9153816 | 0.00355254 |
| ENSG00000145592.13 | 7.2404271 | -1.204171481 | 0.41315669 | -2.9145637 | 0.00356186 |
| ENSG00000237172.3 | 43.8307877 | -0.471953493 | 0.16194307 | -2.9143172 | 0.00356467 |
| ENSG00000260083.1 | 385.977673 | -0.411777954 | 0.14173536 | -2.9052592 | 0.00366949 |
| ENSG00000141753.6 | 83.4592363 | -1.036141173 | 0.35692914 | -2.9029324 | 0.00369686 |
| ENSG00000104852.14 | 66.2871352 | -0.809861974 | 0.27912939 | -2.9013855 | 0.00371517 |
| ENSG00000148358.19 | 7.17510122 | -1.046066047 | 0.3607969 | -2.8993211 | 0.00373972 |

| | | | | | |
|---|---|---|---|---|---|
| ENSG00000105664.10 | 151.095615 | -2.105803414 | 0.72651249 | -2.8985096 | 0.00374941 |
| ENSG00000259198.1 | 31.8077646 | -0.856845944 | 0.29592357 | -2.8954974 | 0.00378558 |
| ENSG00000238278.3 | 109.85116 | -0.758683522 | 0.2620887 | -2.8947586 | 0.0037945 |
| ENSG00000183153.6 | 47.4493142 | -0.807113519 | 0.27893063 | -2.8935994 | 0.00380854 |
| ENSG00000116251.10 | 2.76536318 | -1.785117125 | 0.61701076 | -2.8931701 | 0.00381375 |
| ENSG00000176170.13 | 14.0490932 | -0.947284705 | 0.32757376 | -2.8918211 | 0.00383016 |
| ENSG00000130332.14 | 2.31981363 | -2.095481821 | 0.72638417 | -2.8848121 | 0.00391647 |
| ENSG00000163875.15 | 2.09602782 | -2.528326075 | 0.8788663 | -2.876804 | 0.00401725 |
| ENSG00000278341.1 | 41.4198404 | -0.636027242 | 0.2212495 | -2.8747058 | 0.00404404 |
| ENSG00000232593.7 | 9.2306824 | -1.044681727 | 0.36367401 | -2.8725774 | 0.00407138 |
| ENSG00000136718.9 | 7.27612877 | -1.002073368 | 0.34911599 | -2.8703164 | 0.00410061 |
| ENSG00000258749.1 | 93.8466695 | -1.009070433 | 0.35168228 | -2.8692672 | 0.00411424 |
| ENSG00000285728.1 | 84.5396786 | -0.50269213 | 0.17519025 | -2.869407 | 0.00411242 |
| ENSG00000101246.19 | 6.70749772 | -1.405134096 | 0.49003201 | -2.8674333 | 0.00413816 |
| ENSG00000165650.11 | 4.72930096 | -2.325977049 | 0.81206238 | -2.8642837 | 0.00417953 |
| ENSG00000254829.1 | 5.45459631 | -1.478701229 | 0.51603995 | -2.8654782 | 0.0041638 |
| ENSG00000122705.16 | 14.265135 | -1.046968612 | 0.36551641 | -2.8643546 | 0.0041786 |
| ENSG00000145425.9 | 11.2093012 | -0.895468469 | 0.31256792 | -2.8648765 | 0.00417172 |
| ENSG00000051620.10 | 6.5581893 | -1.402670802 | 0.48988285 | -2.863278 | 0.00419282 |

| padj |
|---|
| 3.20E-10 |
| 7.26E-10 |
| 9.51E-10 |
| 7.66E-08 |
| 2.86E-07 |
| 4.47E-07 |
| 6.50E-07 |
| 1.03E-06 |
| 1.37E-06 |
| 1.67E-06 |
| 1.94E-06 |
| 2.26E-06 |
| 2.33E-06 |
| 3.18E-06 |
| 3.65E-06 |
| 5.29E-06 |
| 6.47E-06 |
| 1.04E-05 |
| 1.11E-05 |
| 1.11E-05 |
| 1.67E-05 |
| 1.88E-05 |
| 1.97E-05 |
| 2.20E-05 |
| 2.32E-05 |
| 2.32E-05 |
| 2.81E-05 |
| 3.01E-05 |
| 4.13E-05 |
| 4.45E-05 |
| 4.59E-05 |
| 4.82E-05 |
| 5.60E-05 |
| 5.93E-05 |
| 6.42E-05 |
| 8.53E-05 |
| 8.86E-05 |
| 8.99E-05 |
| 0.00012563 |
| 0.00012665 |
| 0.0001296 |
| 0.00015912 |

| |
|---|
| 0.00016721 |
| 0.00016911 |
| 0.00018667 |
| 0.00019772 |
| 0.00021307 |
| 0.00022773 |
| 0.00023927 |
| 0.00023985 |
| 0.00025383 |
| 0.00026125 |
| 0.00026154 |
| 0.00030023 |
| 0.0003166 |
| 0.00033942 |
| 0.00038525 |
| 0.00039235 |
| 0.00045464 |
| 0.00045508 |
| 0.00046238 |
| 0.00047466 |
| 0.00048838 |
| 0.00053626 |
| 0.00055815 |
| 0.00057433 |
| 0.00058946 |
| 0.00060101 |
| 0.00061785 |
| 0.0006306 |
| 0.00065256 |
| 0.00067157 |
| 0.00070642 |
| 0.00073445 |
| 0.00076547 |
| 0.00082945 |
| 0.0008376 |
| 0.0008376 |
| 0.00084303 |
| 0.00086837 |
| 0.00096857 |
| 0.00097455 |
| 0.00099791 |
| 0.00103295 |
| 0.00103703 |

| |
|---|
| 0.00106621 |
| 0.00110634 |
| 0.00110634 |
| 0.00121203 |
| 0.00122596 |
| 0.00126015 |
| 0.00126015 |
| 0.00128571 |
| 0.00128842 |
| 0.0012892 |
| 0.00130123 |
| 0.0013077 |
| 0.0013077 |
| 0.00130807 |
| 0.00132928 |
| 0.001361 |
| 0.001361 |
| 0.0013911 |
| 0.00142924 |
| 0.00143549 |
| 0.00144021 |
| 0.00158754 |
| 0.00160501 |
| 0.00160676 |
| 0.00161194 |
| 0.00161915 |
| 0.00176397 |
| 0.00181537 |
| 0.00194288 |
| 0.00194288 |
| 0.00195143 |
| 0.00195537 |
| 0.00199239 |
| 0.00199324 |
| 0.00208511 |
| 0.00215668 |
| 0.00217892 |
| 0.00230187 |
| 0.0023844 |
| 0.00240346 |
| 0.00249523 |
| 0.00252944 |
| 0.00254707 |

0.00257809
0.00257809
0.00260082
0.00260082
0.00272949
0.00272949
0.00280613
0.00280613
0.00282549
0.00283012
0.00283012
0.00291256
0.00295921
0.00298481
0.00307338
0.00310925
0.0032237
0.00328543
0.00330618
0.0033507
0.0033507
0.0033507
0.0033507
0.0033507
0.00336693
0.00336693
0.00337563
0.00337563
0.00345135
0.00348886
0.00350754
0.00383234
0.00396348
0.00403616
0.00403616
0.00407579
0.00407596
0.00409682
0.0041094
0.0041864
0.00434134
0.00438921
0.00458622

| |
|---|
| 0.0046913 |
| 0.00485804 |
| 0.00486093 |
| 0.00486093 |
| 0.00486093 |
| 0.0048947 |
| 0.00505056 |
| 0.00511252 |
| 0.005276 |
| 0.00558988 |
| 0.00562649 |
| 0.00573365 |
| 0.00573365 |
| 0.00577485 |
| 0.00577485 |
| 0.00616361 |
| 0.00617121 |
| 0.00617606 |
| 0.00619064 |
| 0.00619064 |
| 0.00622703 |
| 0.0063388 |
| 0.00662205 |
| 0.00667339 |
| 0.0067134 |
| 0.00675336 |
| 0.00676718 |
| 0.00686925 |
| 0.00687595 |
| 0.00687595 |
| 0.00688497 |
| 0.00691434 |
| 0.00699326 |
| 0.00719246 |
| 0.00728299 |
| 0.00735826 |
| 0.00750108 |
| 0.00756848 |
| 0.0077742 |
| 0.0078129 |
| 0.0078129 |
| 0.00831366 |
| 0.00844379 |

| |
|---|
| 0.00844973 |
| 0.00870729 |
| 0.00870729 |
| 0.0088313 |
| 0.00892001 |
| 0.00892989 |
| 0.00919165 |
| 0.0091931 |
| 0.00967939 |
| 0.00973795 |
| 0.01007109 |
| 0.01017019 |
| 0.01022892 |
| 0.01036855 |
| 0.01039545 |
| 0.01040071 |
| 0.01042252 |
| 0.01042252 |
| 0.01044992 |
| 0.01052849 |
| 0.01060193 |
| 0.01067146 |
| 0.01084987 |
| 0.01085699 |
| 0.01110323 |
| 0.0111917 |
| 0.01147336 |
| 0.01153203 |
| 0.01176869 |
| 0.01193664 |
| 0.01220537 |
| 0.01220537 |
| 0.01220537 |
| 0.01220537 |
| 0.01220537 |
| 0.01236532 |
| 0.01240509 |
| 0.01263224 |
| 0.01264519 |
| 0.012706 |
| 0.012706 |
| 0.012706 |
| 0.01277836 |

| |
|---|
| 0.01282614 |
| 0.01289537 |
| 0.01315893 |
| 0.01315893 |
| 0.01340564 |
| 0.01345398 |
| 0.0134672 |
| 0.01349075 |
| 0.01353411 |
| 0.01353411 |
| 0.01383747 |
| 0.01400557 |
| 0.01400623 |
| 0.0140771 |
| 0.0140771 |
| 0.01436175 |
| 0.01449904 |
| 0.01449904 |
| 0.01450065 |
| 0.01450065 |
| 0.01451625 |
| 0.0146399 |
| 0.01466641 |
| 0.01537275 |
| 0.01566239 |
| 0.01568486 |
| 0.01589653 |
| 0.01602255 |
| 0.01602255 |
| 0.01603846 |
| 0.01607719 |
| 0.01641372 |
| 0.01649513 |
| 0.01674484 |
| 0.01674484 |
| 0.01703021 |
| 0.01718083 |
| 0.01720916 |
| 0.01722496 |
| 0.01745672 |
| 0.01751416 |
| 0.01751716 |
| 0.01804784 |

| |
|---|
| 0.01822946 |
| 0.01829243 |
| 0.01924757 |
| 0.01933795 |
| 0.01993008 |
| 0.01993008 |
| 0.01996636 |
| 0.01996636 |
| 0.0202372 |
| 0.0202372 |
| 0.02027266 |
| 0.0204962 |
| 0.02102838 |
| 0.02128448 |
| 0.02147523 |
| 0.02151584 |
| 0.02151584 |
| 0.0216779 |
| 0.02172489 |
| 0.02179976 |
| 0.02195758 |
| 0.02243798 |
| 0.02250695 |
| 0.02268831 |
| 0.02272537 |
| 0.02301472 |
| 0.02303075 |
| 0.02303369 |
| 0.02314292 |
| 0.02319147 |
| 0.0232558 |
| 0.02326321 |
| 0.02326321 |
| 0.02339675 |
| 0.0238279 |
| 0.02385262 |
| 0.02404372 |
| 0.02418956 |
| 0.02453852 |
| 0.02508248 |
| 0.02544987 |
| 0.02553398 |
| 0.02555533 |

| |
|---|
| 0.02570511 |
| 0.02576458 |
| 0.02587911 |
| 0.02587911 |
| 0.02651341 |
| 0.02668065 |
| 0.0276522 |
| 0.02792206 |
| 0.02802421 |
| 0.02815758 |
| 0.02831975 |
| 0.02840497 |
| 0.0286287 |
| 0.02863877 |
| 0.02866673 |
| 0.02888344 |
| 0.02900345 |
| 0.02931777 |
| 0.02936405 |
| 0.0293873 |
| 0.02957506 |
| 0.02957506 |
| 0.0296888 |
| 0.0296888 |
| 0.03001988 |
| 0.03017334 |
| 0.03037098 |
| 0.03039715 |
| 0.03055538 |
| 0.03069961 |
| 0.03069961 |
| 0.03069961 |
| 0.03073634 |
| 0.03078828 |
| 0.03084068 |
| 0.03084068 |
| 0.03084068 |
| 0.03138615 |
| 0.0314517 |
| 0.03157204 |
| 0.03175171 |
| 0.03183897 |
| 0.03242683 |

| |
|---|
| 0.03242683 |
| 0.03258775 |
| 0.03260222 |
| 0.03264341 |
| 0.03271844 |
| 0.03276861 |
| 0.03283824 |
| 0.03321774 |
| 0.03328797 |
| 0.03344414 |
| 0.03391 |
| 0.03414062 |
| 0.03440942 |
| 0.03442159 |
| 0.03512023 |
| 0.03567202 |
| 0.03629426 |
| 0.03644389 |
| 0.03657951 |
| 0.03695548 |
| 0.03695548 |
| 0.03702978 |
| 0.03706387 |
| 0.0371872 |
| 0.03738506 |
| 0.03743794 |
| 0.03778253 |
| 0.03778253 |
| 0.0384139 |
| 0.03853803 |
| 0.03866336 |
| 0.03877297 |
| 0.03920222 |
| 0.03934701 |
| 0.03934701 |
| 0.03934701 |
| 0.03946673 |
| 0.03946673 |
| 0.03946673 |
| 0.03946673 |
| 0.03946673 |
| 0.03974086 |
| 0.03985081 |

0.03988438
0.03998334
0.04009087
0.04009087
0.04117557
0.04119084
0.04121707
0.04132761
0.04136485
0.04136485
0.04147523
0.04147523
0.04147523
0.04167736
0.04167736
0.04196078
0.04227746
0.0424042
0.04251723
0.04255449
0.04284228
0.0429347
0.04298696
0.04309425
0.04309425
0.04309425
0.04390274
0.0439372
0.04407799
0.04409451
0.04409451
0.04409451
0.044108
0.04415456
0.0444743
0.0444743
0.0444743
0.0445052
0.0445052
0.04553249
0.04577467
0.04582209
0.046027

| |
|---|
| 0.04609947 |
| 0.04645 |
| 0.04651239 |
| 0.04663726 |
| 0.04665393 |
| 0.04676035 |
| 0.04762228 |
| 0.04870113 |
| 0.04892805 |
| 0.04920975 |
| 0.04951367 |
| 0.04953022 |
| 0.04953022 |
| 0.04976877 |
| 0.04991971 |
| 0.04991971 |
| 0.04991971 |
| 0.04991971 |
| 0.04998135 |

| Gene id | baseMean | og2FoldChang | lfcSE | stat | pvalue | padj |
|---|---|---|---|---|---|---|
| ENSG000001 | 733.512007 | 1.06773697 | 0.14859448 | 7.18557644 | 6.69E-13 | 7.26E-10 |
| ENSG000001 | 46.6366389 | -0.9012922 | 0.19226264 | -4.687818 | 2.76E-06 | 0.00022773 |
| ENSG000001 | 15.1789441 | -1.2129795 | 0.26704587 | -4.5422142 | 5.57E-06 | 0.00039235 |
| ENSG000001 | 18.3830134 | -1.1925152 | 0.28679265 | -4.1581094 | 3.21E-05 | 0.00143549 |
| ENSG000001 | 15.3188823 | -1.2526916 | 0.31392922 | -3.9903633 | 6.60E-05 | 0.00254707 |
| ENSG000001 | 236.151439 | 0.81705384 | 0.20617257 | 3.96296099 | 7.40E-05 | 0.00276128 |
| ENSG000001 | 21.3269755 | -0.9479506 | 0.2568605 | -3.6905268 | 0.00022379 | 0.0063388 |
| ENSG000001 | 218.233175 | 0.55822784 | 0.15187375 | 3.67560444 | 0.00023729 | 0.00662818 |
| ENSG000001 | 206.342439 | -1.6712468 | 0.45761985 | -3.6520419 | 0.00026016 | 0.00699326 |
| ENSG000000 | 14.6997167 | -1.3459584 | 0.37824611 | -3.5584197 | 0.00037309 | 0.0091931 |
| ENSG000001 | 21.6434426 | -1.08055 | 0.30796724 | -3.5086523 | 0.00045038 | 0.01060193 |
| ENSG000001 | 197.521326 | 0.44175174 | 0.1353019 | 3.26493372 | 0.0010949 | 0.01993008 |
| ENSG000001 | 8956.85788 | 0.74414909 | 0.23014113 | 3.23344669 | 0.00122306 | 0.02155102 |
| ENSG000000 | 1.59881952 | 2.63871784 | 0.82120803 | 3.21321486 | 0.00131258 | 0.02250695 |
| ENSG000001 | 31.136532 | -0.9457474 | 0.29527286 | -3.2029607 | 0.00136023 | 0.02303075 |
| ENSG000001 | 3.61087597 | -1.7370142 | 0.54804522 | -3.1694725 | 0.00152716 | 0.02508248 |
| ENSG000001 | 43.9304658 | -1.1385215 | 0.36018366 | -3.160947 | 0.00157257 | 0.02555533 |
| ENSG000001 | 2.03058764 | -2.5770652 | 0.81897243 | -3.1467057 | 0.00165121 | 0.02651341 |
| ENSG000001 | 528.02067 | -0.2676271 | 0.08765541 | -3.0531728 | 0.00226435 | 0.03271844 |
| ENSG000001 | 2.96618797 | -1.7751641 | 0.583356 | -3.0430202 | 0.00234217 | 0.03344414 |
| ENSG000001 | 9.10927571 | -1.2802978 | 0.42774525 | -2.9931315 | 0.00276131 | 0.03778253 |
| ENSG000001 | 15.0871029 | -1.117458 | 0.37618669 | -2.9704879 | 0.00297327 | 0.03974086 |
| ENSG000001 | 14.9062379 | -0.8081861 | 0.27496016 | -2.9392844 | 0.00328971 | 0.04251723 |
| ENSG000001 | 7.27612877 | -1.0020734 | 0.34911599 | -2.8703164 | 0.00410061 | 0.04951367 |
| ENSG000001 | 18.8803752 | -1.0259471 | 0.35938387 | -2.8547389 | 0.00430722 | 0.05089324 |
| ENSG000001 | 2.8863428 | 1.99070876 | 0.71046382 | 2.80198472 | 0.00507893 | 0.05684495 |
| ENSG000002 | 95.8575744 | -0.5939246 | 0.21229338 | -2.7976597 | 0.00514743 | 0.05718686 |
| ENSG000001 | 294.524879 | 0.46126686 | 0.16576735 | 2.78261595 | 0.00539226 | 0.05883921 |
| ENSG000002 | 2.46379519 | -1.8289478 | 0.66044012 | -2.7692863 | 0.00561792 | 0.06048499 |
| ENSG000001 | 16.6787496 | -0.802871 | 0.29439796 | -2.7271622 | 0.00638816 | 0.06550654 |
| ENSG000001 | 120.167705 | -0.3623861 | 0.13611458 | -2.6623604 | 0.00775947 | 0.07513427 |
| ENSG000001 | 24.6825369 | -0.5852753 | 0.22055273 | -2.6536753 | 0.00796204 | 0.0763034 |
| ENSG000001 | 19.8944968 | -0.7297882 | 0.28612738 | -2.550571 | 0.01075466 | 0.09413629 |
| ENSG000001 | 2.17548067 | -1.8962914 | 0.74549053 | -2.5436828 | 0.01096907 | 0.09557521 |
| ENSG000001 | 2.33860248 | -1.8130001 | 0.72081663 | -2.5152029 | 0.01189639 | 0.10085309 |
| ENSG000001 | 6.16954763 | -1.0566809 | 0.42539461 | -2.4840015 | 0.01299153 | 0.10699475 |
| ENSG000001 | 7.44518733 | -0.9886564 | 0.40345427 | -2.4504794 | 0.01426661 | 0.11344562 |
| ENSG000001 | 288.209361 | 0.52463046 | 0.2192 | 2.39338712 | 0.01669362 | 0.1256218 |
| ENSG000000 | 21.0621217 | -0.8926831 | 0.37356989 | -2.3896014 | 0.01686667 | 0.12676667 |
| ENSG000001 | 3.28543129 | -1.4789988 | 0.6454836 | -2.2913035 | 0.02194587 | 0.15042531 |
| ENSG000001 | 1.88097893 | -1.7139882 | 0.76096792 | -2.252379 | 0.02429833 | 0.16074537 |
| ENSG000000 | 1.31355368 | 2.09992156 | 0.93772339 | 2.23938272 | 0.02513102 | 0.16450508 |

| | | | | | | |
|---|---|---|---|---|---|---|
| ENSG000001( | 5.82998102 | -1.0436509 | 0.46825886 | -2.2287904 | 0.02582785 | 0.16619482 |
| ENSG000001: | 353.848973 | 0.44903008 | 0.20204084 | 2.22247191 | 0.02625143 | 0.16798741 |
| ENSG000001: | 159.474289 | -0.2654803 | 0.11949171 | -2.2217467 | 0.02630043 | 0.16798741 |
| ENSG000000: | 3.59398832 | -1.2421676 | 0.56442599 | -2.2007626 | 0.02775283 | 0.1736952 |
| ENSG000001: | 4.04094017 | -0.9935258 | 0.45141132 | -2.2009325 | 0.0277408 | 0.1736952 |
| ENSG000001! | 5203.21253 | 0.55553378 | 0.25803791 | 2.15291542 | 0.03132533 | 0.18570021 |
| ENSG000001 | 6.67327186 | 0.89879063 | 0.41773819 | 2.15156441 | 0.03143168 | 0.18596693 |
| ENSG000001( | 3.42882709 | -1.095653 | 0.51802517 | -2.1150574 | 0.03442505 | 0.19602391 |
| ENSG000001: | 14.7850445 | -0.5767257 | 0.2735904 | -2.1079895 | 0.0350319 | 0.19795079 |
| ENSG000001: | 288.465248 | 0.27616665 | 0.13119584 | 2.10499538 | 0.03529171 | 0.198996 |
| ENSG000000: | 2.74324542 | -1.2699693 | 0.60497522 | -2.0992088 | 0.0357985 | 0.20046233 |
| ENSG000001 | 15.2985983 | -0.6417166 | 0.30569664 | -2.0991941 | 0.0357998 | 0.20046233 |
| ENSG000001! | 1.86353098 | 2.04676145 | 0.98324967 | 2.08162943 | 0.03737633 | 0.20633934 |
| ENSG000000: | 50.0441908 | 0.56092883 | 0.27694352 | 2.0254268 | 0.04282356 | 0.22209902 |
| ENSG000001: | 3.75528464 | 1.4029198 | 0.70035536 | 2.00315424 | 0.04516074 | 0.23013182 |
| ENSG000000: | 161.333082 | 0.44236775 | 0.22357322 | 1.97862583 | 0.04785815 | 0.23778047 |
| ENSG000001: | 3.49455205 | -1.0175995 | 0.52471749 | -1.9393283 | 0.05246137 | 0.24904783 |
| ENSG000001: | 17.6423251 | -0.7087306 | 0.36703562 | -1.9309586 | 0.05348818 | 0.2508461 |
| ENSG000001( | 41.7504397 | 0.35741082 | 0.18568934 | 1.9247783 | 0.0542571 | 0.25278971 |
| ENSG000000: | 543.948742 | -0.4431453 | 0.23556106 | -1.8812331 | 0.05994022 | 0.26703846 |
| ENSG000001! | 32.6338963 | -0.3206924 | 0.171673 | -1.8680422 | 0.06175619 | 0.27115909 |
| ENSG000001( | 3.87394914 | -1.1171037 | 0.60086954 | -1.8591452 | 0.06300656 | 0.27367557 |
| ENSG000001( | 1.5278073 | -1.4512443 | 0.78927905 | -1.838696 | 0.06595991 | 0.28156056 |
| ENSG000002( | 68.5240569 | -0.4598109 | 0.25251168 | -1.8209492 | 0.06861458 | 0.28729103 |
| ENSG000001: | 3437.14055 | 0.6863671 | 0.37979714 | 1.80719398 | 0.07073204 | 0.29195932 |
| ENSG000001 | 3.42625479 | -1.1485808 | 0.6493688 | -1.768765 | 0.0769331 | 0.30669515 |
| ENSG000001( | 4.02346981 | -0.9849852 | 0.57173643 | -1.7227959 | 0.08492544 | 0.32274331 |
| ENSG000001: | 10.8146812 | -0.6544344 | 0.38275792 | -1.7097867 | 0.08730533 | 0.32656666 |
| ENSG000000! | 6.18182957 | -0.6260399 | 0.36703525 | -1.7056669 | 0.08807011 | 0.32790969 |
| ENSG000001 | 14.6322186 | -0.4387548 | 0.26064732 | -1.6833274 | 0.09231174 | 0.33688495 |
| ENSG000001( | 1.39216442 | -1.2836937 | 0.77695164 | -1.6522183 | 0.09849005 | 0.34813734 |
| ENSG000001: | 7.01972152 | -0.6940556 | 0.42148986 | -1.6466721 | 0.09962545 | 0.34987235 |
| ENSG000001( | 2.68850412 | 0.95731503 | 0.58707623 | 1.63064859 | 0.10296449 | 0.35684349 |
| ENSG000001: | 41.6943463 | -0.3453745 | 0.21211509 | -1.6282411 | 0.10347378 | 0.35809667 |
| ENSG000001( | 4.92252044 | -0.6769128 | 0.41780375 | -1.6201693 | 0.10519592 | 0.36075808 |
| ENSG000001! | 90.0773096 | 1.1049117 | 0.69675986 | 1.58578552 | 0.11278797 | 0.37478256 |
| ENSG000001! | 2.52429985 | -0.8837447 | 0.57828749 | -1.5282099 | 0.12646043 | 0.39955167 |
| ENSG000001( | 6.43126115 | 0.64829461 | 0.43133532 | 1.50299449 | 0.13284047 | 0.4080124 |
| ENSG000000: | 38.6884247 | 0.31874554 | 0.21473208 | 1.48438712 | 0.13770625 | 0.41486403 |
| ENSG000002: | 1.97436916 | -1.3931639 | 0.94846212 | -1.4688661 | 0.1418691 | 0.42074421 |
| ENSG000001: | 10.2228006 | -1.4046512 | 1.01993781 | -1.377193 | 0.1684526 | 0.46068339 |
| ENSG000000( | 1.73521341 | -1.0857982 | 0.79171977 | -1.3714426 | 1.70E-01 | 0.46304322 |
| ENSG000001( | 3.46834234 | -0.6770275 | 0.49743347 | -1.3610414 | 0.17350062 | 0.46834737 |

| | | | | | | |
|---|---|---|---|---|---|---|
| ENSG0000001 | 5.22624476 | -0.5859663 | 0.43085855 | -1.3599968 | 0.17383092 | 0.46861292 |
| ENSG0000001 | 31.0392252 | -0.2609852 | 0.19220071 | -1.3578782 | 0.17450234 | 0.46943012 |
| ENSG0000000 | 1.30154369 | -1.194629 | 0.88513605 | -1.3496558 | 0.17712642 | 0.47180683 |
| ENSG0000001 | 1.41498299 | -1.0895441 | 0.81378033 | -1.3388675 | 0.1806138 | 0.47540855 |
| ENSG0000001 | 96.8016308 | -0.2121341 | 0.15873721 | -1.3363857 | 0.18142324 | 0.47628036 |
| ENSG0000001 | 1.30607988 | -1.3205821 | 0.99071289 | -1.3329615 | 0.18254445 | 0.47776045 |
| ENSG0000001 | 32.5561584 | 0.25827579 | 0.19400986 | 1.33125081 | 0.1831065 | 0.47859286 |
| ENSG0000001 | 4.20631622 | -0.6487381 | 0.48812456 | -1.3290422 | 0.18383406 | 0.47943954 |
| ENSG0000001 | 1.55800491 | -0.9958086 | 0.75429467 | -1.3201851 | 0.18677321 | 0.48167816 |
| ENSG0000001 | 2.11311147 | 0.94247122 | 0.72638179 | 1.2974874 | 0.19446354 | 0.49017583 |
| ENSG0000001 | 18.1419426 | -0.4366311 | 0.33972042 | -1.285266 | 0.19869933 | 0.4952251 |
| ENSG0000000 | 25.0884293 | 0.34788043 | 0.27231454 | 1.2774949 | 0.20142758 | 0.49723204 |
| ENSG0000002 | 5.02801001 | -0.5461814 | 0.42998381 | -1.2702372 | 0.20400016 | 0.50140991 |
| ENSG0000001 | 16.4938168 | -0.5040394 | 0.40048818 | -1.2585624 | 0.20818844 | 0.50648176 |
| ENSG0000001 | 1.39156963 | -1.0935829 | 0.86898317 | -1.2584627 | 0.20822446 | 0.50648176 |
| ENSG0000000 | 4.08047785 | -0.673527 | 0.55033322 | -1.2238531 | 2.21E-01 | 0.52337813 |
| ENSG0000001 | 8.01905488 | -0.4273694 | 0.35076806 | -1.2183819 | 0.22307889 | 0.5253263 |
| ENSG0000001 | 6.45148559 | -0.4488831 | 0.36879046 | -1.2171767 | 0.223537 | 0.5259975 |
| ENSG0000001 | 1.8045453 | -0.8465459 | 0.69569445 | -1.2168359 | 0.22366668 | 0.52609839 |
| ENSG0000001 | 22.940677 | -0.2787654 | 0.23030029 | -1.2104433 | 0.22610884 | 0.52886696 |
| ENSG0000001 | 1.40419932 | -1.2491655 | 1.04018629 | -1.2009055 | 0.22978785 | 0.53264209 |
| ENSG0000002 | 20.8089858 | 0.28910476 | 0.2415186 | 1.19702893 | 0.23129528 | 0.53458024 |
| ENSG0000000 | 2.94966366 | -0.7441522 | 0.62947958 | -1.1821704 | 0.23713808 | 0.54099076 |
| ENSG0000001 | 1.90186926 | -0.7813467 | 0.66680194 | -1.1717822 | 0.24128449 | 0.5446827 |
| ENSG0000000 | 10.2660887 | -0.3319996 | 0.28840888 | -1.1511422 | 0.24967375 | 0.55509922 |
| ENSG0000001 | 11.1465464 | -0.4107005 | 0.35888233 | -1.1443878 | 0.25246287 | 0.55860692 |
| ENSG0000001 | 1.74990055 | 0.78990064 | 0.69084887 | 1.1433769 | 0.25288214 | 0.55922841 |
| ENSG0000001 | 11.7023263 | -0.3137243 | 0.27596801 | -1.136814 | 0.25561604 | 0.56183833 |
| ENSG0000001 | 7.79463138 | -0.41836 | 0.37257381 | -1.1228916 | 0.26148352 | 0.5667983 |
| ENSG0000001 | 27.9084772 | 0.25293976 | 0.22573799 | 1.12050152 | 0.26250011 | 0.56786024 |
| ENSG0000001 | 2.57973844 | -0.6514199 | 0.58638906 | -1.1109004 | 0.2666112 | 0.57243234 |
| ENSG0000000 | 159.416837 | -0.1477045 | 0.13653429 | -1.0818123 | 0.27933595 | 0.58771082 |
| ENSG0000002 | 3.13344704 | 0.63228019 | 0.59466471 | 1.06325494 | 0.28766635 | 0.59564687 |
| ENSG0000001 | 16.9952928 | -0.2680984 | 0.25944962 | -1.0333349 | 0.30144718 | 0.60775723 |
| ENSG0000001 | 7.65286371 | -0.3516947 | 0.34966393 | -1.0058079 | 0.314508 | 0.62108556 |
| ENSG0000001 | 6.97398514 | -0.3817433 | 0.38116062 | -1.0015287 | 0.31657128 | 0.62352455 |
| ENSG0000001 | 14.8432747 | -0.3594997 | 0.36088008 | -0.9961751 | 0.31916508 | 0.62598905 |
| ENSG0000001 | 53.3670185 | -0.157809 | 0.16015753 | -0.9853359 | 0.32445913 | 0.63061923 |
| ENSG0000002 | 2.38037409 | -0.7215999 | 0.74241228 | -0.9719666 | 0.33106718 | 0.63595745 |
| ENSG0000001 | 1.80064613 | -0.7266952 | 0.74822935 | -0.9712199 | 0.3314388 | 0.63631007 |
| ENSG0000001 | 12.2926 | -0.2615357 | 0.26926749 | -0.9712858 | 0.33140599 | 0.63631007 |
| ENSG0000001 | 10.2344617 | -0.2716697 | 0.28178453 | -0.9641043 | 0.33499362 | 0.63936836 |
| ENSG0000001 | 5.51490527 | -0.3902385 | 0.40715892 | -0.9584426 | 0.3378396 | 0.64124464 |

| | | | | | | |
|---|---|---|---|---|---|---|
| ENSG000001 | 6.67612223 | 0.35416327 | 0.37529276 | 0.94369866 | 0.34532366 | 0.649069 |
| ENSG000001 | 17.1103339 | 0.27202506 | 0.30393572 | 0.8950085 | 0.37078254 | 0.67039475 |
| ENSG000001 | 1.68168138 | -0.7088932 | 0.79578534 | -0.8908096 | 0.3730313 | 0.67215495 |
| ENSG000001 | 2.29437346 | 0.611435 | 0.68768624 | 0.88911914 | 0.37393905 | 0.67269077 |
| ENSG000001 | 1.29135001 | -0.8025393 | 0.91059341 | -0.8813367 | 0.37813563 | 0.67650562 |
| ENSG000001 | 3.73632289 | -0.4626488 | 0.53421466 | -0.8660353 | 0.38647079 | 0.68167982 |
| ENSG000001 | 17.8832596 | -0.2084471 | 0.24454259 | -0.8523959 | 0.39399438 | 0.68784474 |
| ENSG000002 | 16.1089297 | 0.24693803 | 0.2925355 | 0.84413012 | 0.3985967 | 0.69219134 |
| ENSG000002 | 2.63552853 | -0.6328395 | 0.76689325 | -0.8251989 | 0.40925866 | 0.70126399 |
| ENSG000001 | 5.66649796 | 0.31147493 | 0.39210094 | 0.79437434 | 0.42697754 | 0.71459873 |
| ENSG000001 | 7.72141875 | -0.2813958 | 0.35601411 | -0.7904064 | 0.42929046 | 0.71655284 |
| ENSG000001 | 5.44371296 | 0.32385532 | 0.41507817 | 0.78022729 | 0.4352571 | 0.71949357 |
| ENSG000001 | 35.5512311 | -0.2143929 | 0.27469521 | -0.7804758 | 0.43511089 | 0.71949357 |
| ENSG000002 | 122.229866 | -0.1695453 | 0.21799668 | -0.7777424 | 0.43672089 | 0.72058558 |
| ENSG000000 | 3.62433638 | -0.3682151 | 0.48138917 | -0.7649011 | 4.44E-01 | 0.72479564 |
| ENSG000000 | 6.64375869 | -0.3200115 | 0.41813442 | -0.7653315 | 0.44407414 | 0.72479564 |
| ENSG000001 | 93.6077383 | -0.1408149 | 0.18685247 | -0.7536153 | 0.45108025 | 0.72832561 |
| ENSG000001 | 1.90833356 | -0.6131018 | 0.83671763 | -0.7327464 | 0.46371312 | 0.73735495 |
| ENSG000001 | 2.38756217 | 0.47923552 | 0.66358778 | 0.72218859 | 0.47017854 | 0.74149378 |
| ENSG000001 | 3.02714484 | 0.40732236 | 0.56827628 | 0.71676819 | 0.47351714 | 0.74400728 |
| ENSG000001 | 1.28547553 | 0.61572751 | 0.86254731 | 0.71384781 | 0.47532129 | 0.7448282 |
| ENSG000000 | 2.181605 | -0.4736958 | 0.66702359 | -0.7101634 | 4.78E-01 | 0.74632361 |
| ENSG000000 | 375.191485 | 0.11424296 | 0.16206285 | 0.70492994 | 0.48085383 | 0.74937537 |
| ENSG000000 | 6.47268023 | -0.2607388 | 0.37141017 | -0.702024 | 0.4826642 | 0.75055646 |
| ENSG000001 | 6.24340871 | 0.28340929 | 0.4062629 | 0.69760072 | 0.48542693 | 0.75243967 |
| ENSG000001 | 94.4341822 | 0.11231127 | 0.16142441 | 0.69575148 | 0.48658448 | 0.75317462 |
| ENSG000001 | 58.3385948 | -0.1571972 | 0.22944998 | -0.6851046 | 0.49327794 | 0.75739601 |
| ENSG000001 | 4.46371736 | -0.3125167 | 0.45734981 | -0.683321 | 0.494404 | 0.75767674 |
| ENSG000001 | 22.659981 | -0.1380022 | 0.20414325 | -0.6760069 | 0.49903628 | 0.76034007 |
| ENSG000001 | 9.40506366 | -0.3379615 | 0.49998452 | -0.675944 | 0.49907623 | 0.76034007 |
| ENSG000000 | 2.26268535 | 0.43994593 | 0.68620692 | 0.6411272 | 0.52144004 | 0.77364063 |
| ENSG000001 | 15.8637167 | -0.2919191 | 0.45690044 | -0.6389119 | 0.52288027 | 0.77442088 |
| ENSG000001 | 1.56337135 | -0.4755992 | 0.76927199 | -0.6182459 | 0.53641328 | 0.78345899 |
| ENSG000001 | 18.5985072 | -0.2123546 | 0.35084235 | -0.6052707 | 0.54499913 | 0.78861595 |
| ENSG000001 | 2.13096508 | 0.35921748 | 0.60071396 | 0.59798425 | 0.54985044 | 0.79194926 |
| ENSG000001 | 59.4482714 | 0.08691906 | 0.14563175 | 0.59684143 | 0.55061326 | 0.79243775 |
| ENSG000001 | 2.54656061 | -0.3590836 | 0.61005967 | -0.588604 | 0.55612692 | 0.7962592 |
| ENSG000001 | 1.55231222 | -0.5235846 | 0.90377841 | -0.5793285 | 0.56236752 | 0.80122006 |
| ENSG000001 | 3.47886306 | 0.27768394 | 0.49079674 | 0.56578196 | 0.57154201 | 0.80833712 |
| ENSG000001 | 1.62554068 | -0.5830804 | 1.07845825 | -0.5406611 | 0.58874122 | 0.81796318 |
| ENSG000001 | 3.10465378 | 0.29673126 | 0.55577183 | 0.53390843 | 0.59340489 | 0.81950051 |
| ENSG000001 | 1.54042135 | -0.4958241 | 0.94924886 | -0.5223331 | 0.60143839 | 0.82446083 |
| ENSG000001 | 2.9401637 | 0.26040812 | 0.50971794 | 0.51088671 | 0.60943039 | 0.82919468 |

| | | | | | | |
|---|---|---|---|---|---|---|
| ENSG0000014 | 7.56073703 | -0.181849 | 0.3613983 | -0.5031818 | 0.61483646 | 0.83139403 |
| ENSG0000014 | 7.22617521 | -0.1805554 | 0.36649496 | -0.4926544 | 0.62225679 | 0.83575294 |
| ENSG0000011 | 4.92753541 | -0.2021061 | 0.4329403 | -0.4668221 | 0.6406272 | 0.84462463 |
| ENSG0000011 | 1.53438382 | -0.3655494 | 0.80942995 | -0.4516134 | 0.65154749 | 0.85142937 |
| ENSG000000! | 3.13522165 | -0.2479298 | 0.55474973 | -0.4469218 | 0.6549315 | 0.85386925 |
| ENSG0000011 | 10.3986187 | -0.1445273 | 0.3362089 | -0.4298734 | 0.6672877 | 0.86079232 |
| ENSG000001! | 1.30325122 | 0.36197481 | 0.88625073 | 0.40843386 | 0.68295518 | 0.86841468 |
| ENSG0000011 | 2.88732332 | -0.2265786 | 0.57328484 | -0.3952286 | 0.69267419 | 0.87335301 |
| ENSG0000002 | 4.91255681 | -0.1796912 | 0.46276562 | -0.3882984 | 6.98E-01 | 0.87607408 |
| ENSG000001( | 1.77943563 | 0.25328915 | 0.66375476 | 0.3816005 | 0.70275772 | 0.87893653 |
| ENSG0000014 | 8.34798175 | -0.1326166 | 0.36591947 | -0.3624203 | 0.71703798 | 0.8865826 |
| ENSG000001! | 1.58095108 | -0.3108604 | 0.98016042 | -0.3171526 | 0.7511278 | 0.90516127 |
| ENSG0000014 | 594.90805 | -0.0919986 | 0.34546365 | -0.2663048 | 0.7900045 | 0.9222171 |
| ENSG0000017 | 27.1637785 | -0.0667358 | 0.27797395 | -0.2400792 | 0.81026885 | 0.93282476 |
| ENSG0000011 | 384.088742 | 0.04570736 | 0.19827301 | 0.23052741 | 0.81768197 | 0.93675662 |
| ENSG0000011 | 2.09403167 | 0.13877145 | 0.61955517 | 0.22398562 | 0.82276849 | 0.93834564 |
| ENSG0000011 | 3.97902086 | 0.0970437 | 0.50497007 | 0.19217714 | 0.84760345 | 0.94799806 |
| ENSG0000018 | 6.71920157 | -0.08442 | 0.46421835 | -0.181854 | 0.85569728 | 0.95031429 |
| ENSG0000014 | 27.7308196 | 0.04564668 | 0.27000376 | 0.16905942 | 0.8657499 | 0.95430861 |
| ENSG0000011 | 81.9677135 | 0.0349692 | 0.21579068 | 0.1620515 | 0.8712653 | 0.95567977 |
| ENSG000002! | 4.50231203 | -0.0701144 | 0.46383451 | -0.1511626 | 0.87984744 | 0.95717479 |
| ENSG000000( | 1.56298138 | 0.11406559 | 0.77208384 | 0.14773731 | 0.88255009 | 0.95844797 |
| ENSG0000014 | 2.6354562 | -0.0863335 | 0.62615478 | -0.1378788 | 0.8903362 | 0.96224889 |
| ENSG0000017 | 8.53661211 | 0.05348967 | 0.38845603 | 0.13769814 | 0.89047899 | 0.96230108 |
| ENSG0000011 | 75.7935643 | 0.03852105 | 0.29803105 | 0.12925178 | 0.89715842 | 0.96440799 |
| ENSG0000011 | 1.45746384 | 0.12692872 | 1.02075772 | 0.12434755 | 0.9010401 | 0.96543952 |
| ENSG0000011 | 2.26636057 | 0.07601291 | 0.63344431 | 0.11999936 | 0.90448365 | 0.96645877 |
| ENSG000002! | 5.36292037 | 0.0486235 | 0.4601617 | 0.10566612 | 0.91584726 | 0.97080376 |
| ENSG000001( | 32.2981357 | -0.0281091 | 0.27397709 | -0.1025966 | 0.9182831 | 0.97126972 |
| ENSG000001( | 1.83935963 | 0.07000924 | 0.75277229 | 0.09300189 | 0.92590206 | 0.97358112 |
| ENSG0000011 | 23.209451 | -0.0220541 | 0.27699868 | -0.0796179 | 0.93654114 | 0.97632198 |
| ENSG0000011 | 1.69683506 | -0.0439954 | 0.7689477 | -0.057215 | 0.95437389 | 0.98363059 |
| ENSG0000017 | 8.52232142 | -0.0150083 | 0.39017765 | -0.0384654 | 0.96931665 | 0.99059375 |
| ENSG0000014 | 2.28154999 | 0.02590446 | 0.73524311 | 0.03523251 | 0.97189434 | 0.99106699 |
| ENSG0000014 | 3.86051011 | -0.0107047 | 0.48125206 | -0.0222433 | 0.98225384 | 0.99316457 |
| ENSG0000017 | 5.88574631 | -0.0092403 | 0.3843905 | -0.0240389 | 0.98082158 | 0.99316457 |
| ENSG0000011 | 1.70946642 | -0.0092187 | 0.83438857 | -0.0110485 | 0.99118478 | 0.99731839 |
| ENSG000000! | 0.6036222 | -2.0593953 | 1.13519594 | -1.814132 | 0.06965741 | NA |
| ENSG000000( | 1.13646217 | 0.99365696 | 0.8962263 | 1.10871212 | 0.26755439 | NA |
| ENSG000000( | 0.85805431 | 0.61755286 | 0.99760202 | 0.6190373 | 0.53589179 | NA |
| ENSG0000007 | 0.58108083 | -0.5755676 | 1.15328883 | -0.4990663 | 0.61773269 | NA |
| ENSG0000007 | 0.72515461 | -0.86809 | 1.04269363 | -0.8325456 | 0.40510107 | NA |
| ENSG0000007 | 0.97643922 | -0.468184 | 0.94023062 | -0.4979459 | 0.61852218 | NA |

| | | | | | | |
|---|---|---|---|---|---|---|
| ENSG0000008 | 0.17074281 | 0.08569952 | 2.31928159 | 0.03695089 | 0.97052416 | NA |
| ENSG0000008 | 0.11402337 | 0.83985718 | 3.04669342 | 0.27566186 | 0.78280779 | NA |
| ENSG0000008 | 0.46997227 | -0.6042168 | 1.35687585 | -0.4453 | 0.65610297 | NA |
| ENSG0000010 | 0.89859789 | -1.5612998 | 0.96239243 | -1.622311 | 0.10473678 | NA |
| ENSG0000010 | 0.30935535 | -1.0559063 | 1.36912316 | -0.7712281 | 0.44057172 | NA |
| ENSG0000010 | 1.10349769 | 0.3529629 | 0.8163889 | 0.43234652 | 0.66548958 | NA |
| ENSG0000010 | 0.35318851 | 1.47557261 | 1.53711687 | 0.95996123 | 0.33707473 | NA |
| ENSG0000010 | 1.26266987 | -0.9493801 | 0.93109291 | -1.0196405 | 0.30789898 | NA |
| ENSG0000010 | 0.00492421 | 0.29059237 | 3.04687218 | 0.09537399 | 0.92401777 | NA |
| ENSG0000010 | 0.01477263 | 0.11637786 | 3.04654239 | 0.03819998 | 0.96952824 | NA |
| ENSG0000010 | 0.38665268 | -0.0828177 | 1.76734265 | -0.04686 | 0.96262478 | NA |
| ENSG0000010 | 0.7110017 | -0.1437493 | 1.08882541 | -0.1320223 | 0.89496664 | NA |
| ENSG0000010 | 0.67625805 | 0.52344385 | 1.10817705 | 0.47234678 | 0.63667929 | NA |
| ENSG0000011 | 0.79762087 | 1.09308016 | 1.11862514 | 0.97716395 | 0.32848799 | NA |
| ENSG0000011 | 0.48216699 | -1.5650465 | 1.25678686 | -1.245276 | 0.21303033 | NA |
| ENSG0000011 | 0.77418505 | -0.6114785 | 1.0159767 | -0.6018628 | 0.5472655 | NA |
| ENSG0000011 | 0.25718231 | -0.8124713 | 1.87531244 | -0.4332458 | 0.6648362 | NA |
| ENSG0000011 | 1.19052892 | 1.12433256 | 0.8569701 | 1.31198574 | 0.18952494 | NA |
| ENSG0000011 | 0.26085127 | 1.29653245 | 2.01011825 | 0.64500307 | 0.51892519 | NA |
| ENSG0000011 | 0.78322585 | -2.9519761 | 1.0336991 | -2.8557402 | 0.00429366 | NA |
| ENSG0000011 | 0.2897579 | 0.05417567 | 1.78180048 | 0.03040502 | 0.97574404 | NA |
| ENSG0000011 | 0.39708474 | 0.58413749 | 1.57207409 | 0.37157122 | 0.71021212 | NA |
| ENSG0000011 | 0.11479014 | -0.2352577 | 3.04598255 | -0.0772354 | 0.93843629 | NA |
| ENSG0000011 | 0.87406364 | -0.5405805 | 0.9022221 | -0.5991657 | 0.54906241 | NA |
| ENSG0000011 | 0.13045495 | 0.23083956 | 2.51348241 | 0.09184053 | 0.92682474 | NA |
| ENSG0000011 | 0.55793791 | -0.3487587 | 1.52267079 | -0.2290441 | 0.81883467 | NA |
| ENSG0000011 | 1.21557469 | 0.07767618 | 0.96590171 | 0.08041831 | 0.93590457 | NA |
| ENSG0000011 | 0.09271673 | 0.36235122 | 3.04641237 | 0.11894359 | 0.90532005 | NA |
| ENSG0000011 | 0.23724706 | 0.40688093 | 1.83579955 | 0.22163691 | 0.82459655 | NA |
| ENSG0000011 | 0.65003767 | -0.6733187 | 1.54003094 | -0.4372111 | 0.66195824 | NA |
| ENSG0000011 | 0.96567315 | -1.1057747 | 1.03875877 | -1.0645154 | 0.2870953 | NA |
| ENSG0000011 | 0.01477263 | 0.11637786 | 3.04654239 | 0.03819998 | 0.96952824 | NA |
| ENSG0000011 | 0.46259576 | 0.06801986 | 1.26122806 | 0.05393145 | 0.95698978 | NA |
| ENSG0000011 | 0.00492421 | 0.29059237 | 3.04687218 | 0.09537399 | 0.92401777 | NA |
| ENSG0000011 | 0.25371783 | -1.4014611 | 1.88923068 | -0.7418158 | 0.45819897 | NA |
| ENSG0000011 | 0.20848103 | -0.0884262 | 2.31291911 | -0.0382314 | 0.96950317 | NA |
| ENSG0000011 | 0.45639843 | 0.244604 | 1.17936006 | 0.20740401 | 0.83569435 | NA |
| ENSG0000011 | 1.14469496 | 0.30639119 | 0.87415582 | 0.35049951 | 0.72596386 | NA |
| ENSG0000014 | 0.83805849 | -0.8856928 | 1.12093191 | -0.7901397 | 0.42944618 | NA |
| ENSG0000014 | 1.02182416 | -1.7138245 | 0.93505171 | -1.8328661 | 0.06682249 | NA |
| ENSG0000014 | 0.34280984 | 0.39219983 | 1.57648661 | 0.24878095 | 0.80353023 | NA |
| ENSG0000014 | 1.03892727 | -1.5358666 | 0.91772265 | -1.6735629 | 0.09421654 | NA |
| ENSG0000014 | 0.03446947 | -0.1484891 | 3.04610873 | -0.0487471 | 0.96112081 | NA |

| | | | | | | |
|---|---|---|---|---|---|---|
| ENSG0000014 | 0.28674078 | -0.8608114 | 1.64761223 | -0.52246 | 0.60135012 | NA |
| ENSG0000014 | 0.58347887 | 0.3364064 | 1.17861533 | 0.2854251 | 0.77531848 | NA |
| ENSG0000014 | 0.06352272 | 0.61118818 | 3.04687218 | 0.20059528 | 0.84101505 | NA |
| ENSG0000015 | 1.14873079 | -0.0474015 | 1.08084891 | -0.0438558 | 0.96501935 | NA |
| ENSG0000015 | 0.39198229 | 0.31668499 | 1.35253427 | 0.23414193 | 0.81487479 | NA |
| ENSG0000015 | 0.26185448 | -1.4647466 | 1.79175817 | -0.8174912 | 0.41364777 | NA |
| ENSG0000016 | 0.27011385 | -0.1697993 | 2.19746375 | -0.0772706 | 0.93840831 | NA |
| ENSG0000016 | 0.1168766 | -0.4822492 | 2.60524132 | -0.1851073 | 0.85314487 | NA |
| ENSG0000016 | 0 | NA | NA | NA | NA | NA |
| ENSG0000016 | 0.59122315 | -0.6248081 | 1.11693388 | -0.5593957 | 0.57589167 | NA |
| ENSG0000016 | 0.4628743 | -1.2274296 | 1.44128497 | -0.8516218 | 0.39442406 | NA |
| ENSG0000016 | 0.05208748 | 0.71566443 | 3.04708895 | 0.23486824 | 0.814311 | NA |
| ENSG0000016 | 0.14375618 | 0.02141902 | 2.44053712 | 0.00877635 | 0.99299757 | NA |
| ENSG0000017 | 0.24447832 | 0.72742975 | 1.86308548 | 0.39044357 | 0.69620857 | NA |
| ENSG0000017 | 0.27434412 | 0.20082908 | 1.62012686 | 0.12395886 | 0.90134785 | NA |
| ENSG0000017 | 0.46646308 | 0.98259318 | 1.72485883 | 0.56966585 | 0.56890436 | NA |
| ENSG0000017 | 0.83182334 | -0.9931977 | 0.98958023 | -1.0036556 | 0.31554466 | NA |
| ENSG0000017 | 1.22306097 | -1.1604828 | 0.90596245 | -1.2809392 | 0.20021501 | NA |
| ENSG0000017 | 0.66717386 | 0.37931534 | 1.2091115 | 0.31371411 | 0.75373818 | NA |
| ENSG0000017 | 1.26125991 | -0.839765 | 0.8128979 | -1.033051 | 0.30158002 | NA |
| ENSG0000017 | 0.57491258 | 0.02060269 | 1.35372558 | 0.01521925 | 0.98785726 | NA |
| ENSG0000018 | 0.71025839 | -1.3318087 | 1.05393945 | -1.2636482 | 0.20635632 | NA |
| ENSG0000018 | 0 | NA | NA | NA | NA | NA |
| ENSG0000018 | 0.15188313 | -0.2109443 | 2.03882691 | -0.1034636 | 0.91759507 | NA |
| ENSG0000018 | 0.42361119 | 0.95695198 | 1.25565903 | 0.76211133 | 0.44599356 | NA |
| ENSG0000018 | 0.16029699 | -1.0141658 | 2.40720941 | -0.4213035 | 0.67353345 | NA |
| ENSG0000018 | 0.09135804 | -0.4264876 | 3.04572908 | -0.1400281 | 0.8886378 | NA |
| ENSG0000018 | 0.0840626 | -0.3534751 | 3.04582204 | -0.1160525 | 0.90761097 | NA |
| ENSG0000019 | 0.06728348 | -0.2352574 | 3.04598255 | -0.0772353 | 0.93843635 | NA |
| ENSG0000019 | 0.42972057 | -0.2968834 | 1.59647791 | -0.1859615 | 0.85247497 | NA |
| ENSG0000019 | 0.40602563 | 0.00253565 | 1.37059394 | 0.00185004 | 0.99852388 | NA |
| ENSG000002[illegible] | 0 | NA | NA | NA | NA | NA |
| ENSG0000024 | 0.16582426 | -0.209743 | 1.99595879 | -0.1050838 | 0.9163093 | NA |
| ENSG0000025 | 0 | NA | NA | NA | NA | NA |
| ENSG0000025 | 0 | NA | NA | NA | NA | NA |
| ENSG0000027 | 0.30469479 | 0.46664263 | 1.42077504 | 0.32844231 | 0.74257726 | NA |
| ENSG0000027 | 0 | NA | NA | NA | NA | NA |
| ENSG0000027 | 0 | NA | NA | NA | NA | NA |
| ENSG0000027 | 0.60218874 | -2.0025179 | 1.44551418 | -1.3853326 | 0.16595079 | NA |
| ENSG0000027 | 0.17964869 | 0.99881998 | 2.50541705 | 0.39866416 | 0.69014068 | NA |

| ensg | gene | |
|---|---|---|
| ENSG000001 | MTERF4 | |
| ENSG000001 | RPS25 | |
| ENSG000001 | RPS9 | |
| ENSG000001 | RPL35 | |
| ENSG000001 | RPL26 | |
| ENSG000001 | RIOX2 | |
| ENSG000001 | RPL14 | |
| ENSG000001 | IMP3 | |
| ENSG000001 | NOC2L | |
| ENSG000000 | RPS5 | |
| ENSG000001 | RPS14 | |
| ENSG000001 | TSR1 | |
| ENSG000001 | LTO1 | |
| ENSG000000 | RIOK2 | |
| ENSG000001 | RPS19 | |
| ENSG000001 | RPSA | |
| ENSG000001 | RPS8 | |
| ENSG000001 | BYSL | |
| ENSG000001 | FTSJ3 | |
| ENSG000001 | ZNF622 | |
| ENSG000001 | RPS16 | |
| ENSG000001 | NOP53 | |
| ENSG000001 | RPL11 | |
| ENSG000001 | IMP4 | p-val <0.05 |
| ENSG000001 | RPS27 | |
| ENSG000001 | NSUN5 | |
| ENSG000002 | RPS28 | |
| ENSG000001 | NUP88 | |
| ENSG000002 | RPLP0P6 | |
| ENSG000001 | DDX51 | |
| ENSG000001 | TRMT112 | |
| ENSG000001 | RPS24 | |
| ENSG000001 | SART1 | |
| ENSG000001 | SIRT7 | |
| ENSG000001 | RRP7A | |
| ENSG000001 | RPS21 | |
| ENSG000001 | RPL27 | |
| ENSG000001 | UTP23 | |
| ENSG000000 | RPLP0 | |
| ENSG000001 | RPS7 | |
| ENSG000001 | GTPBP4 | |
| ENSG000000 | DDX18 | |

| | |
|---|---|
| ENSG000001 | DDX21 |
| ENSG000001 | HEATR1 |
| ENSG000001 | WDR55 |
| ENSG000000 | XRCC5 |
| ENSG000001 | RRP36 |
| ENSG000001 | SRFBP1 |
| ENSG000001 | EIF2A |
| ENSG000001 | FBL |
| ENSG000001 | DDX54 |
| ENSG000001 | EMG1 |
| ENSG000000 | RPL26L1 |
| ENSG000001 | METTL25B |
| ENSG000001 | NOL8 |
| ENSG000000 | DIS3 |
| ENSG000001 | MRPL44 |
| ENSG000000 | DIMT1 |
| ENSG000001 | DHX30 |
| ENSG000001 | RRP7BP |
| ENSG000001 | EXOSC3 |
| ENSG000000 | MTREX |
| ENSG000001 | MAK16 |
| ENSG000001 | ERCC2 |
| ENSG000001 | C1QBP |
| ENSG000002 | BOP1 |
| ENSG000001 | KRR1 |
| ENSG000001 | EIF4A3 |
| ENSG000001 | NOLC1 |
| ENSG000001 | DDX28 |
| ENSG000000 | USP36 |
| ENSG000001 | ISG20L2 |
| ENSG000001 | GRWD1 |
| ENSG000001 | NPM1 |
| ENSG000001 | TSC1 |
| ENSG000001 | CUL4A |
| ENSG000001 | DDX49 |
| ENSG000001 | MALSU1 |
| ENSG000001 | DHX37 |
| ENSG000001 | SNU13 |
| ENSG000000 | NOP14 |
| ENSG000002 | EXOSC6 |
| ENSG000001 | ISG20 |
| ENSG000000 | RPUSD1 |
| ENSG000001 | NOP56 |

| | |
|---|---|
| ENSG000001 | PA2G4 |
| ENSG000001 | PIH1D1 |
| ENSG000000 | WDR18 |
| ENSG000001 | SLX9 |
| ENSG000001 | DCAF13 |
| ENSG000001 | NAT10 |
| ENSG000001 | NHP2 |
| ENSG000001 | PWP1 |
| ENSG000001 | URB1 |
| ENSG000001 | TENT4B |
| ENSG000001 | RPS6 |
| ENSG000000 | XPO1 |
| ENSG000002 | WDR46 |
| ENSG000001 | RPS15 |
| ENSG000001 | NOL6 |
| ENSG000000 | TSR3 |
| ENSG000001 | TRAF7 |
| ENSG000001 | WDR74 |
| ENSG000001 | RIOK1 |
| ENSG000001 | DDX17 |
| ENSG000001 | SPATA5L1 |
| ENSG000002 | MRPL20 |
| ENSG000000 | RRP12 |
| ENSG000001 | ABT1 |
| ENSG000000 | RPL6 |
| ENSG000001 | PDCD11 |
| ENSG000001 | BMS1 |
| ENSG000001 | MYBBP1A |
| ENSG000001 | PES1 |
| ENSG000001 | EXOSC8 |
| ENSG000001 | REXO4 |
| ENSG000000 | BUD23 |
| ENSG000002 | DDX3X |
| ENSG000001 | DDX31 |
| ENSG000001 | MDN1 |
| ENSG000001 | MRPS2 |
| ENSG000001 | MRM3 |
| ENSG000001 | KRI1 |
| ENSG000002 | EIF6 |
| ENSG000001 | ERI3 |
| ENSG000001 | MRPS7 |
| ENSG000001 | RPL23A |
| ENSG000001 | NMD3 |

| | |
|---|---|
| ENSG000001 | RPS27L |
| ENSG000001 | UTP15 |
| ENSG000001 | UTP3 |
| ENSG000001 | ERI2 |
| ENSG000001 | DDX10 |
| ENSG000001 | EBNA1BP2 |
| ENSG000001 | ABCE1 |
| ENSG000002 | NSUN5P1 |
| ENSG000002 | AATF |
| ENSG000001 | DDX27 |
| ENSG000001 | RPL38 |
| ENSG000001 | NOB1 |
| ENSG000001 | RPP38 |
| ENSG000002 | ZNHIT3 |
| ENSG000000 | UTP18 |
| ENSG000000 | LSG1 |
| ENSG000001 | GTF3A |
| ENSG000001 | KAT2B |
| ENSG000001 | WDR43 |
| ENSG000001 | DDX56 |
| ENSG000001 | TFB2M |
| ENSG000000 | LAS1L |
| ENSG000000 | NOP16 |
| ENSG000000 | XRN2 |
| ENSG000001 | RPP40 |
| ENSG000001 | NSUN5P2 |
| ENSG000001 | NIP7 |
| ENSG000001 | TBL3 |
| ENSG000001 | TSR2 |
| ENSG000001 | RBIS |
| ENSG000000 | WBP11 |
| ENSG000001 | NOP9 |
| ENSG000001 | CUL4B |
| ENSG000001 | PIH1D2 |
| ENSG000001 | YBEY |
| ENSG000001 | CINP |
| ENSG000001 | MRPS11 |
| ENSG000001 | NSUN3 |
| ENSG000001 | GLUL |
| ENSG000001 | RAN |
| ENSG000001 | NGRN |
| ENSG000001 | UTP14A |
| ENSG000001 | YTHDF2 |

| | |
|---|---|
| ENSG000001 | RPL7 |
| ENSG000001 | NOM1 |
| ENSG000001 | GNL2 |
| ENSG000001 | UTP25 |
| ENSG000000 | MRTO4 |
| ENSG000001 | UTP20 |
| ENSG000001 | SDAD1 |
| ENSG000001 | DKC1 |
| ENSG000000 | TFB1M |
| ENSG000001 | PIN4 |
| ENSG000001 | SURF6 |
| ENSG000001 | NUDT16 |
| ENSG000001 | RPL10 |
| ENSG000001 | GEMIN4 |
| ENSG000001 | METTL5 |
| ENSG000001 | METTL16 |
| ENSG000001 | DROSHA |
| ENSG000001 | RPL35A |
| ENSG000001 | UTP4 |
| ENSG000001 | BRIX1 |
| ENSG000002 | MPV17L2 |
| ENSG000000 | DHX29 |
| ENSG000001 | RPL7L1 |
| ENSG000001 | EXOSC10 |
| ENSG000001 | EXOSC9 |
| ENSG000001 | MPHOSPH10 |
| ENSG000001 | SBDS |
| ENSG000002 | PRKDC |
| ENSG000001 | NSA2 |
| ENSG000001 | ERI1 |
| ENSG000001 | RPL5 |
| ENSG000001 | RPF1 |
| ENSG000001 | EXOSC4 |
| ENSG000001 | LYAR |
| ENSG000001 | PELP1 |
| ENSG000001 | RRS1 |
| ENSG000001 | ZNHIT6 |
| ENSG000000 | NOP58 |
| ENSG000000 | WDR3 |
| ENSG000000 | RRP15 |
| ENSG000000 | NLE1 |
| ENSG000000 | EXOSC7 |
| ENSG000000 | EXOSC5 |

| | |
|---|---|
| ENSG0000008 | MRPL22 |
| ENSG0000008 | RRN3 |
| ENSG0000008 | ESF1 |
| ENSG0000010 | MTG2 |
| ENSG0000010 | RBFA |
| ENSG0000010 | RIOK3 |
| ENSG0000010 | SUV39H1 |
| ENSG0000010 | POP4 |
| ENSG0000010 | GTPBP10 |
| ENSG0000010 | NPM3 |
| ENSG0000010 | UTP6 |
| ENSG0000010 | GAR1 |
| ENSG0000010 | FRG1 |
| ENSG0000011 | NOP2 |
| ENSG0000011 | PAK1IP1 |
| ENSG0000011 | RRP9 |
| ENSG0000011 | WDR75 |
| ENSG0000011 | NOL10 |
| ENSG0000011 | C1orf109 |
| ENSG0000011 | NSUN4 |
| ENSG0000011 | FASTKD2 |
| ENSG0000011 | FCF1 |
| ENSG0000011 | RCL1 |
| ENSG0000011 | MRM2 |
| ENSG0000011 | NGDN |
| ENSG0000011 | GNL3L |
| ENSG0000011 | EXOSC2 |
| ENSG0000011 | PPAN |
| ENSG0000011 | NOL11 |
| ENSG0000011 | RRP8 |
| ENSG0000011 | ERAL1 |
| ENSG0000011 | WDR36 |
| ENSG0000011 | LTV1 |
| ENSG0000011 | MPHOSPH6 |
| ENSG0000011 | URB2 |
| ENSG0000011 | RNASEL |
| ENSG0000011 | RSL24D1 |
| ENSG0000011 | WDR12 |
| ENSG0000014 | EFL1 |
| ENSG0000014 | ZNF593 |
| ENSG0000014 | NVL |
| ENSG0000014 | SDE2 |
| ENSG0000014 | SHQ1 |

| | |
|---|---|
| ENSG0000014 | SPATA5 |
| ENSG0000014 | NAF1 |
| ENSG0000014 | RPP30 |
| ENSG0000015 | HEATR3 |
| ENSG0000015 | NIFK |
| ENSG0000015 | MTERF3 |
| ENSG0000016 | NOL9 |
| ENSG0000016 | LSM6 |
| ENSG0000016 | RPL10L |
| ENSG0000016 | METTL17 |
| ENSG0000016 | RPUSD2 |
| ENSG0000016 | ZCCHC4 |
| ENSG0000016 | METTL15 |
| ENSG0000017 | TRMT61B |
| ENSG0000017 | EXOSC1 |
| ENSG0000017 | CHD7 |
| ENSG0000017 | MRPL36 |
| ENSG0000017 | METTL18 |
| ENSG0000017 | POP7 |
| ENSG0000017 | METTL15P1 |
| ENSG0000017 | RPP25 |
| ENSG0000018 | NOP10 |
| ENSG0000018 | VCX |
| ENSG0000018 | RPS17 |
| ENSG0000018 | UTP11 |
| ENSG0000018 | NOC4L |
| ENSG0000018 | FBLL1 |
| ENSG0000018 | TRMT2B |
| ENSG0000019 | C1D |
| ENSG0000019 | RPF2 |
| ENSG0000019 | TMA16 |
| ENSG0000021 | DDX47 |
| ENSG0000024 | PWP2 |
| ENSG0000025 | UTP14C |
| ENSG0000025 | FDXACB1 |
| ENSG0000027 | GTF2H5 |
| ENSG0000027 | ZNF658 |
| ENSG0000027 | RMRP |
| ENSG0000027 | DDX52 |
| ENSG0000027 | MRM1 |

| Gene id | baseMean | og2FoldChang | lfcSE | stat | pvalue | padj |
|---|---|---|---|---|---|---|
| ENSG000001 | 82.6757211 | 0.90394345 | 0.14365787 | 6.2923352 | 3.13E-10 | 1.46E-07 |
| ENSG000001 | 396.67791 | 1.68265122 | 0.31108306 | 5.40900956 | 6.34E-08 | 1.05E-05 |
| ENSG000001 | 709.081647 | -4.2496536 | 0.82210399 | -5.169241 | 2.35E-07 | 3.01E-05 |
| ENSG000001 | 6.56165092 | 2.84450849 | 0.62077801 | 4.58216693 | 4.60E-06 | 0.00033811 |
| ENSG000001 | 3199.76902 | 0.721861 | 0.16231059 | 4.44740531 | 8.69E-06 | 0.00055749 |
| ENSG000001 | 4.10810742 | -3.0468196 | 0.71578723 | -4.2565996 | 2.08E-05 | 0.00106621 |
| ENSG000001 | 17.0000899 | -1.1876754 | 0.28177383 | -4.2149956 | 2.50E-05 | 0.00122596 |
| ENSG000001 | 27.8262064 | -1.5499373 | 0.36982132 | -4.1910437 | 2.78E-05 | 0.0013077 |
| ENSG000001 | 57.1511373 | 0.75586083 | 0.18218864 | 4.14878133 | 3.34E-05 | 0.00147887 |
| ENSG000002 | 23.7557997 | -2.2345939 | 0.54634799 | -4.0900561 | 4.31E-05 | 0.00181537 |
| ENSG000000 | 323.160178 | -0.7361512 | 0.18080723 | -4.0714701 | 4.67E-05 | 0.00194288 |
| ENSG000001 | 12.0463223 | -1.8466821 | 0.46186984 | -3.9982739 | 6.38E-05 | 0.00249523 |
| ENSG000001 | 74.3864219 | 1.08426982 | 0.27170023 | 3.99068416 | 6.59E-05 | 0.00254707 |
| ENSG000001 | 18.3173281 | -1.1073259 | 0.27989693 | -3.9561916 | 7.62E-05 | 0.00280613 |
| ENSG000001 | 35.1783469 | 0.85774015 | 0.22093693 | 3.88228512 | 0.00010348 | 0.00347502 |
| ENSG000001 | 3.87119937 | -2.5590856 | 0.6680335 | -3.8307744 | 0.00012774 | 0.00409682 |
| ENSG000001 | 6.53496365 | -3.0386992 | 0.80159262 | -3.7908273 | 0.00015015 | 0.0046913 |
| ENSG000002 | 24.63388 | -1.0375785 | 0.27774789 | -3.7356844 | 0.00018721 | 0.00558988 |
| ENSG000001 | 1293.48701 | 0.77980432 | 0.21150917 | 3.68685826 | 0.00022704 | 0.00641586 |
| ENSG000001 | 10.9499681 | -1.6234049 | 0.44252793 | -3.6684801 | 0.000244 | 0.00675336 |
| ENSG000000 | 735.228743 | 0.4205382 | 0.11491947 | 3.65941637 | 0.00025279 | 0.00687595 |
| ENSG000001 | 10.021566 | -1.114029 | 0.30698748 | -3.6289068 | 0.00028462 | 0.00750108 |
| ENSG000001 | 3.19647801 | -2.4955715 | 0.7094702 | -3.5175143 | 0.00043561 | 0.01039545 |
| ENSG000001 | 258.000723 | -0.5745495 | 0.16602413 | -3.4606386 | 0.0005389 | 0.01220537 |
| ENSG000001 | 18.0514367 | -1.1705884 | 0.33846972 | -3.4584728 | 0.00054325 | 0.01220537 |
| ENSG000001 | 389.411085 | -0.5024285 | 0.14593943 | -3.4427193 | 0.0005759 | 0.012706 |
| ENSG000001 | 7.03251286 | -1.4432366 | 0.42608961 | -3.3871669 | 0.00070618 | 0.01449904 |
| ENSG000000 | 4.37757015 | 1.759823 | 0.52086106 | 3.37868031 | 0.00072835 | 0.01470268 |
| ENSG000001 | 36.1809121 | 0.85706449 | 0.26106853 | 3.28291001 | 0.00102741 | 0.0189291 |
| ENSG000001 | 7.84037786 | -1.2193502 | 0.37491489 | -3.2523387 | 0.0011446 | 0.0204962 |
| ENSG000001 | 337.566396 | -0.8007488 | 0.25105467 | -3.1895396 | 0.001425 | 0.0238279 |
| ENSG000001 | 15.1041406 | 1.55570743 | 0.4919431 | 3.16237273 | 0.00156489 | 0.02553321 |
| ENSG000001 | 223.78361 | -0.4929816 | 0.1581462 | -3.117252 | 0.00182545 | 0.0286287 |
| ENSG000001 | 25.3852713 | 0.62899551 | 0.20694438 | 3.03944237 | 0.00237017 | 0.03380414 |
| ENSG000001 | 23.992361 | 1.21220023 | 0.39901123 | 3.03801031 | 0.00238146 | 0.03388546 |
| ENSG000001 | 100.580457 | -0.4341098 | 0.14455198 | -3.0031398 | 0.0026721 | 0.03706387 |
| ENSG000001 | 26.3672692 | -0.6769997 | 0.22813006 | -2.9676041 | 0.00300131 | 0.03998334 |
| ENSG000001 | 3.33426543 | -2.0226027 | 0.68982375 | -2.9320572 | 0.00336725 | 0.04309425 |
| ENSG000000 | 81.4779722 | 0.97411967 | 0.3336006 | 2.92001771 | 0.00350011 | 0.044108 |
| ENSG000000 | 17.1063985 | 0.96334289 | 0.33083876 | 2.91181992 | 0.0035933 | 0.04472886 |
| ENSG000001 | 13.958955 | 0.84548514 | 0.29651888 | 2.85137038 | 0.00435312 | 0.05128563 |
| ENSG000000 | 2.69286508 | -1.8800728 | 0.66401064 | -2.8313895 | 0.00463462 | 0.05345913 |

| | | | | | | |
|---|---|---|---|---|---|---|
| ENSG000001 | 41.5539647 | 1.54139351 | 0.55100481 | 2.79742299 | 0.0051512 | 0.05718686 |
| ENSG000001 | 15.3205922 | -1.0036202 | 0.36432866 | -2.754711 | 0.0058744 | 0.06204145 |
| ENSG000001 | 13.8708681 | -0.9299884 | 0.33959016 | -2.7385611 | 0.00617087 | 0.06404004 |
| ENSG000001 | 10.4921493 | -0.8857311 | 0.32474618 | -2.7274567 | 0.00638246 | 0.06550654 |
| ENSG000001 | 10.4483529 | 0.94444814 | 0.34955586 | 2.70185183 | 0.00689545 | 0.0691427 |
| ENSG000000 | 9.13610076 | -1.6336737 | 0.6103909 | -2.6764384 | 0.00744092 | 0.07310074 |
| ENSG000001 | 81.0347971 | 0.83902377 | 0.31430246 | 2.66947884 | 0.00759691 | 0.07398466 |
| ENSG000001 | 5.90413139 | 1.06213678 | 0.39908566 | 2.66142561 | 0.00778105 | 0.07522305 |
| ENSG000001 | 6.47099402 | -1.2250701 | 0.47133009 | -2.5991765 | 0.00934477 | 0.08537051 |
| ENSG000001 | 5.67622071 | -1.1621195 | 0.45020687 | -2.5813012 | 0.00984287 | 0.08838733 |
| ENSG000001 | 1.60088693 | 3.77002474 | 1.48800233 | 2.5336148 | 0.01128928 | 0.09740917 |
| ENSG000001 | 2.40958923 | -1.5649215 | 0.61984737 | -2.5246885 | 0.01158009 | 0.09896192 |
| ENSG000002 | 4.76756535 | -1.3350617 | 0.53530362 | -2.4940271 | 0.01263029 | 0.10505775 |
| ENSG000000 | 49.1945586 | 0.62383974 | 0.25218398 | 2.47374846 | 0.01337038 | 0.1088577 |
| ENSG000001 | 1.82448808 | 1.62377888 | 0.65846822 | 2.46599431 | 0.01366335 | 0.11020679 |
| ENSG000001 | 10.0112331 | -0.8250043 | 0.33522908 | -2.4610166 | 0.0138544 | 0.1112297 |
| ENSG000001 | 2.3437222 | -1.9083867 | 0.77657366 | -2.4574446 | 0.01399294 | 0.1121934 |
| ENSG000001 | 26.050095 | -0.7627555 | 0.31070846 | -2.4548913 | 0.01409272 | 0.11247269 |
| ENSG000001 | 835.699215 | 0.57952316 | 0.23957764 | 2.41893672 | 0.01556595 | 0.12004198 |
| ENSG000001 | 3.62277805 | -1.9839228 | 0.82234225 | -2.4125269 | 0.01584237 | 0.12156395 |
| ENSG000001 | 5.94723235 | -0.9382788 | 0.39650223 | -2.3663898 | 0.01796252 | 0.13149738 |
| ENSG000001 | 148.463546 | 0.42218758 | 0.18218698 | 2.31733121 | 0.0204857 | 0.14314011 |
| ENSG000001 | 15.6590998 | 0.67124738 | 0.29042773 | 2.31123718 | 0.02081976 | 0.14494456 |
| ENSG000001 | 6.27592296 | -0.9047517 | 0.39239412 | -2.305722 | 0.02112617 | 0.14638666 |
| ENSG000001 | 3.80369715 | 1.56902423 | 0.69022129 | 2.27321912 | 0.02301298 | 0.15516482 |
| ENSG000001 | 7.98447701 | -0.8092067 | 0.35707819 | -2.266189 | 0.02343981 | 0.15708173 |
| ENSG000001 | 3.11457593 | -1.248335 | 0.55148272 | -2.2635976 | 0.02359888 | 0.15788586 |
| ENSG000001 | 370.942936 | 0.39849322 | 0.17685153 | 2.25326421 | 0.02424249 | 0.16066377 |
| ENSG000002 | 8.39167659 | 0.71601056 | 0.31878899 | 2.24603293 | 0.0247019 | 0.16254846 |
| ENSG000001 | 3.63594926 | 1.03575155 | 0.46295837 | 2.23724553 | 0.0252703 | 0.16494066 |
| ENSG000001 | 8.52369927 | 0.80081845 | 0.35822173 | 2.23553847 | 0.02538202 | 0.16516705 |
| ENSG000000 | 1.63311517 | 1.80218212 | 0.81649399 | 2.20722032 | 2.73E-02 | 1.72E-01 |
| ENSG000001 | 61.5361176 | -0.5685231 | 0.25866053 | -2.1979509 | 0.02795261 | 0.17422595 |
| ENSG000001 | 7.47766792 | -0.9722554 | 0.44313281 | -2.1940498 | 0.02823183 | 0.175254 |
| ENSG000000 | 3.67801364 | -1.2078694 | 0.55360617 | -2.1818207 | 0.02912277 | 0.17804103 |
| ENSG000001 | 21.0064333 | -0.5102091 | 0.23555916 | -2.165949 | 0.03031508 | 0.18182847 |
| ENSG000001 | 5891.32317 | -0.3019048 | 0.13965186 | -2.1618387 | 0.03063061 | 0.18291532 |
| ENSG000001 | 15.0605417 | -0.6827923 | 0.31579347 | -2.1621482 | 0.03060675 | 0.18291532 |
| ENSG000001 | 2.08960849 | -1.4451313 | 0.67120286 | -2.153047 | 0.03131498 | 0.18570021 |
| ENSG000001 | 2.25974232 | -1.4134048 | 0.66290463 | -2.1321389 | 0.03299543 | 0.19218913 |
| ENSG000001 | 32.1307829 | -0.6459246 | 0.30331548 | -2.1295471 | 0.03320902 | 0.19290509 |
| ENSG000001 | 3.22953543 | -1.2462197 | 0.58575689 | -2.1275375 | 0.03337545 | 0.19314576 |
| ENSG000001 | 4.01557578 | -1.4333403 | 0.67452345 | -2.1249673 | 0.03358934 | 0.19374081 |

| | | | | | | |
|---|---|---|---|---|---|---|
| ENSG000002 | 11.4900674 | -0.8688522 | 0.40986166 | -2.119867 | 0.03401726 | 0.19498407 |
| ENSG000000 | 1.53786732 | 1.64883099 | 0.77885803 | 2.11698529 | 0.0342611 | 0.19564167 |
| ENSG000000 | 2.4401884 | -1.4376547 | 0.6802233 | -2.1135041 | 0.03455765 | 0.19630897 |
| ENSG000001 | 36.2999943 | -0.5457388 | 0.25902495 | -2.1068965 | 0.03512655 | 0.19834149 |
| ENSG000001 | 9.66148983 | -0.740016 | 0.35460118 | -2.0868966 | 0.03689748 | 0.20467145 |
| ENSG000001 | 4.47209139 | -0.9724804 | 0.46986836 | -2.0696869 | 0.03848168 | 0.20916262 |
| ENSG000001 | 1.40621664 | 2.17989706 | 1.05547252 | 2.0653281 | 0.03889197 | 0.21048468 |
| ENSG000000 | 3.23066825 | 1.09119352 | 0.53491443 | 2.03994034 | 0.04135627 | 0.21788876 |
| ENSG000001 | 1.44790413 | -1.5550214 | 0.76397523 | -2.0354344 | 0.04180719 | 0.21936737 |
| ENSG000001 | 6.19618755 | 0.80409759 | 0.39745146 | 2.02313408 | 0.04305933 | 0.2227948 |
| ENSG000000 | 1.4509663 | -2.0080366 | 1.01532328 | -1.9777313 | 0.04795903 | 0.23779071 |
| ENSG000000 | 13.5615213 | -0.7772211 | 0.39390355 | -1.9731254 | 4.85E-02 | 0.23889701 |
| ENSG000001 | 23.0242627 | 0.59127282 | 0.30031167 | 1.96886399 | 0.04896871 | 0.24024593 |
| ENSG000001 | 7.96468281 | 0.78827771 | 0.40553559 | 1.94379419 | 0.05192028 | 0.24780055 |
| ENSG000001 | 13.2232 | -0.5153722 | 0.26590579 | -1.938176 | 0.05260176 | 0.24919542 |
| ENSG000001 | 2.81638662 | 1.47435646 | 0.76344124 | 1.93119834 | 0.05345853 | 0.25080409 |
| ENSG000001 | 14.7296882 | -0.7777945 | 0.40451208 | -1.9227966 | 0.0545056 | 0.2531691 |
| ENSG000001 | 1.62092646 | 2.13703686 | 1.11642355 | 1.91418111 | 0.05559702 | 0.2558856 |
| ENSG000001 | 1.5676871 | -2.235937 | 1.17613967 | -1.9010812 | 0.05729138 | 0.26067083 |
| ENSG000000 | 84.9541985 | -0.3653875 | 0.19252389 | -1.8978814 | 0.05771171 | 0.26164511 |
| ENSG000001 | 7.41737077 | 0.72602261 | 0.38415914 | 1.88990065 | 0.05877125 | 0.2646671 |
| ENSG000001 | 443.493361 | -0.4131349 | 0.22007632 | -1.8772345 | 0.06048597 | 0.26800854 |
| ENSG000001 | 9.47206041 | 0.59772868 | 0.31969188 | 1.86970244 | 0.06152515 | 0.2705366 |
| ENSG000001 | 7.67991786 | -0.7407339 | 0.39650492 | -1.8681583 | 0.06174001 | 0.27115909 |
| ENSG000001 | 18.0500428 | 0.78645607 | 0.42110441 | 1.86760349 | 0.06181736 | 0.27132942 |
| ENSG000001 | 2.81051806 | -1.0827485 | 0.58569969 | -1.848641 | 0.06450967 | 0.27761828 |
| ENSG000001 | 19.0334819 | -0.6504127 | 0.3522736 | -1.8463283 | 0.06484454 | 0.27826917 |
| ENSG000001 | 1.69081544 | 1.29492705 | 0.70738112 | 1.83059317 | 0.06716129 | 0.28421691 |
| ENSG000001 | 9.41822638 | -0.5793665 | 0.3179724 | -1.8220654 | 0.06844506 | 0.287114 |
| ENSG000001 | 14.7443225 | -0.5406723 | 0.29801936 | -1.8142187 | 0.06964406 | 0.28997368 |
| ENSG000001 | 9.94187206 | 0.58266491 | 0.32238122 | 1.80737857 | 0.07070327 | 0.29194177 |
| ENSG000001 | 18.1782699 | -0.4608628 | 0.25557851 | -1.8032142 | 0.07135458 | 0.29392851 |
| ENSG000001 | 6.54392342 | -0.8405851 | 0.46682923 | -1.8006265 | 0.07176176 | 0.29540504 |
| ENSG000001 | 94.121428 | 0.3183749 | 0.17841357 | 1.78447693 | 0.0743462 | 0.30093455 |
| ENSG000001 | 17.9821422 | 0.78380461 | 0.44176911 | 1.7742404 | 0.07602339 | 0.30477236 |
| ENSG000000 | 3.46976749 | -1.0016069 | 0.56616569 | -1.7691055 | 0.07687627 | 0.30658577 |
| ENSG000001 | 6.20185695 | -0.9583803 | 0.54256909 | -1.7663746 | 0.07733302 | 0.30758143 |
| ENSG000000 | 6.99961764 | -0.6380518 | 0.36155766 | -1.7647304 | 0.07760908 | 0.30817389 |
| ENSG000001 | 2.39755227 | -1.0821511 | 0.61399225 | -1.7624833 | 0.07798768 | 0.30856548 |
| ENSG000001 | 1.42434064 | -1.4821014 | 0.84739173 | -1.7490157 | 0.08028831 | 0.3131709 |
| ENSG000002 | 8.24368887 | -0.6800082 | 0.39043654 | -1.7416612 | 0.08156775 | 0.31562266 |
| ENSG000001 | 1.36225501 | -2.0712418 | 1.19163859 | -1.7381459 | 0.0821851 | 0.31724061 |
| ENSG000001 | 1.63389791 | 1.23629093 | 0.71153655 | 1.73749462 | 0.0822999 | 0.31724061 |

| | | | | | | |
|---|---|---|---|---|---|---|
| ENSG000001( | 12.4736847 | -0.7968391 | 0.45903759 | -1.7358907 | 0.08258316 | 0.31782718 |
| ENSG000001: | 4.6961788 | 0.97334194 | 0.56173913 | 1.73272945 | 0.08314377 | 0.31927523 |
| ENSG000002( | 1.90347791 | -1.1994559 | 0.69325543 | -1.7301788 | 0.08359833 | 0.32031053 |
| ENSG000002' | 470.605094 | -0.2624134 | 0.15166662 | -1.7301985 | 0.08359481 | 0.32031053 |
| ENSG000001' | 8.04105613 | -0.7097099 | 0.41113309 | -1.726229 | 0.08430622 | 0.32149869 |
| ENSG000001( | 1.91727365 | -2.2826039 | 1.32734962 | -1.7196704 | 0.08549237 | 0.32372339 |
| ENSG000001: | 2.86302969 | 1.04626631 | 0.60865273 | 1.71898728 | 0.08561669 | 0.32372339 |
| ENSG000001/ | 20.3926527 | 0.56888534 | 0.33083875 | 1.7195245 | 0.08551891 | 0.32372339 |
| ENSG000002( | 4.6528018 | -0.8711065 | 0.50703478 | -1.718041 | 0.08578914 | 0.32408148 |
| ENSG000001: | 19.293148 | -0.3980411 | 0.23258341 | -1.7113906 | 0.08700902 | 0.32592321 |
| ENSG000000: | 7.07107706 | -0.6594041 | 0.3856005 | -1.7100708 | 0.08725279 | 0.32647084 |
| ENSG000001( | 2.40787755 | -1.0874094 | 0.64599407 | -1.6833117 | 0.09231476 | 0.33688495 |
| ENSG000001: | 33.7937448 | -0.3989975 | 0.23801964 | -1.6763218 | 0.09367518 | 0.34000725 |
| ENSG000001/ | 40.6351815 | -0.3312106 | 0.19932117 | -1.6616928 | 0.09657438 | 0.34553192 |
| ENSG000001( | 37.483628 | 1.58740309 | 0.96342162 | 1.64767228 | 0.09941994 | 0.34945645 |
| ENSG000001( | 2.68850412 | 0.95731503 | 0.58707623 | 1.63064859 | 0.10296449 | 0.35684349 |
| ENSG000001: | 4.93919975 | -0.7742649 | 0.47647661 | -1.6249799 | 0.10416685 | 0.35916231 |
| ENSG000001: | 13.6905342 | -0.6156492 | 0.37948653 | -1.6223215 | 0.10473452 | 0.36029983 |
| ENSG000000! | 25.0074418 | -0.4261924 | 0.2650136 | -1.6081905 | 0.10779346 | 0.36553289 |
| ENSG000001/ | 213.631608 | 0.24667574 | 0.15351111 | 1.60689185 | 0.1080781 | 0.3660885 |
| ENSG000001( | 1.83678355 | 1.09154077 | 0.68133679 | 1.60205758 | 0.10914288 | 0.36790539 |
| ENSG000001: | 3.56137604 | -0.8136518 | 0.50895325 | -1.5986768 | 0.10989243 | 0.36912693 |
| ENSG000001! | 2.32258812 | -1.1948614 | 0.75218917 | -1.5885118 | 0.11217065 | 0.37417127 |
| ENSG000001! | 19.6125674 | 0.44218161 | 0.27888086 | 1.58555736 | 0.11283975 | 0.37478256 |
| ENSG000000: | 36.474435 | -0.3907017 | 0.24736771 | -1.5794368 | 0.1142359 | 0.3780731 |
| ENSG000001: | 1.28211691 | 1.44628425 | 0.91636019 | 1.57829233 | 0.11449847 | 0.37853026 |
| ENSG000001( | 13.2336402 | -0.9561579 | 0.61279761 | -1.560316 | 0.11868522 | 0.38584632 |
| ENSG000001: | 8.96333543 | 0.53580938 | 0.34752904 | 1.54176867 | 0.12312982 | 0.39364524 |
| ENSG000001: | 26.7081403 | -0.4081132 | 0.26574895 | -1.5357094 | 0.12460968 | 0.39637419 |
| ENSG000001' | 2.57643126 | -0.9385223 | 0.61551195 | -1.5247832 | 0.12731318 | 0.40029462 |
| ENSG000001! | 12.5214239 | -0.4310861 | 0.28521329 | -1.5114518 | 0.13067338 | 0.40432704 |
| ENSG000001/ | 12.984337 | -0.5143202 | 0.34057109 | -1.5101698 | 0.13100011 | 0.40481352 |
| ENSG000000: | 2.07272649 | -1.1720892 | 0.7815094 | -1.4997762 | 0.13367237 | 0.40952999 |
| ENSG000001: | 3.74930028 | -0.7448961 | 0.49682049 | -1.4993265 | 0.13378894 | 0.40966879 |
| ENSG000001/ | 4.93651232 | 0.63306062 | 0.42258298 | 1.49807411 | 0.134114 | 0.40990205 |
| ENSG000001' | 5.45228764 | 0.68095942 | 0.45642738 | 1.49193378 | 0.13571651 | 0.41173454 |
| ENSG000001( | 4.17397415 | -0.7256414 | 0.49289576 | -1.4722005 | 0.14096675 | 0.41991863 |
| ENSG000001( | 3.75705221 | 0.77084019 | 0.52572206 | 1.46625041 | 0.14258008 | 0.42220281 |
| ENSG000001! | 1.59254567 | 1.20075415 | 0.81912633 | 1.46589616 | 0.14267658 | 0.42238538 |
| ENSG000001( | 1.87028445 | -1.0438771 | 0.7131409 | -1.4637739 | 0.14325573 | 0.4235182 |
| ENSG000001( | 2.91469932 | -0.8510138 | 0.59092741 | -1.4401325 | 0.14982992 | 0.43379723 |
| ENSG000000( | 4.13262055 | 0.80372862 | 0.55888772 | 1.43808603 | 0.15040965 | 0.43449854 |
| ENSG000001: | 2.47585966 | 0.98644784 | 0.68676482 | 1.4363692 | 0.15089731 | 0.43451974 |

| | | | | | | |
|---|---|---|---|---|---|---|
| ENSG000001 | 5.65500499 | -0.6009496 | 0.41807166 | -1.4374321 | 0.15059526 | 0.43451974 |
| ENSG000002 | 3.50931131 | -1.0223098 | 0.72592653 | -1.4082828 | 0.15904735 | 0.44803527 |
| ENSG000001 | 2.71436153 | -0.8572483 | 0.60962374 | -1.4061924 | 0.159667 | 0.44884877 |
| ENSG000001 | 2.36622334 | -0.902365 | 0.64217915 | -1.4051609 | 0.15997344 | 0.44893168 |
| ENSG000000 | 19.9712664 | 0.41863158 | 0.29993044 | 1.39576226 | 0.1627861 | 0.45283522 |
| ENSG000001 | 2.66788176 | 1.09518997 | 0.78730305 | 1.39106533 | 0.16420562 | 0.45520966 |
| ENSG000001 | 4.98788483 | -0.6577028 | 0.47582907 | -1.382225 | 0.16690263 | 0.45965687 |
| ENSG000001 | 1.45526539 | 1.20831613 | 0.87738602 | 1.37717731 | 0.16845743 | 0.46068339 |
| ENSG000000 | 18.6022771 | 0.49420593 | 0.36014188 | 1.37225343 | 0.16998456 | 0.4628829 |
| ENSG000001 | 11.0909328 | -0.4364394 | 0.31852286 | -1.3701981 | 0.17062508 | 0.46393592 |
| ENSG000001 | 1.90900375 | -1.096402 | 0.80114155 | -1.3685497 | 0.17114009 | 0.46446585 |
| ENSG000001 | 5.79373739 | 0.60246165 | 0.44115311 | 1.36565204 | 0.1720482 | 0.46581305 |
| ENSG000001 | 1.32184128 | -1.3121815 | 0.96479672 | -1.3600601 | 0.17381092 | 0.46861292 |
| ENSG000000 | 18.5090826 | -0.3148018 | 0.23564197 | -1.3359327 | 0.18157127 | 0.47651578 |
| ENSG000002 | 8.77430235 | -0.6008924 | 0.45280551 | -1.3270429 | 0.18449448 | 0.47996279 |
| ENSG000000 | 22.5638681 | 0.34241994 | 0.25820227 | 1.32616939 | 0.1847836 | 0.47998964 |
| ENSG000001 | 1.82802565 | -1.0125068 | 0.76427016 | -1.3248022 | 0.18523676 | 0.48024493 |
| ENSG000001 | 1.52886403 | -1.2094848 | 0.9156193 | -1.3209473 | 0.18651895 | 0.48167816 |
| ENSG000000 | 1.58962568 | 1.0778584 | 0.81818777 | 1.31737291 | 0.18771366 | 0.48264107 |
| ENSG000001 | 84.8414251 | -0.1717539 | 0.13068315 | -1.3142774 | 0.18875286 | 0.48346807 |
| ENSG000001 | 1.39614728 | -1.0751867 | 0.82076611 | -1.3099794 | 0.19020281 | 0.48576883 |
| ENSG000001 | 6.16034562 | -0.6753758 | 0.51697191 | -1.3064071 | 0.19141415 | 0.48674983 |
| ENSG000001 | 269.569242 | 0.21385474 | 0.16411138 | 1.3031073 | 0.19253813 | 0.48770158 |
| ENSG000001 | 3.1670673 | -0.7209063 | 0.55317561 | -1.3032142 | 0.19250166 | 0.48770158 |
| ENSG000000 | 10.8506592 | 0.37121125 | 0.28640725 | 1.29609583 | 0.19494247 | 0.49041037 |
| ENSG000002 | 5.35106607 | -0.6045686 | 0.46898499 | -1.2891002 | 0.19736325 | 0.49414182 |
| ENSG000000 | 5.04059592 | 0.61186732 | 0.47484562 | 1.28856051 | 0.19755092 | 0.49420343 |
| ENSG000001 | 1557.17196 | 0.27217859 | 0.21220096 | 1.28264546 | 0.19961632 | 0.49605691 |
| ENSG000001 | 16.3247781 | -0.3847734 | 0.30033246 | -1.2811582 | 0.20013809 | 0.49605691 |
| ENSG000001 | 2.11107154 | -0.8876683 | 0.693468 | -1.2800422 | 0.20053031 | 0.49643229 |
| ENSG000001 | 9.92071991 | -0.4383703 | 0.3429732 | -1.2781473 | 0.2011975 | 0.49723204 |
| ENSG000001 | 4.28153749 | -1.0578146 | 0.83778176 | -1.2626374 | 0.20671952 | 0.50474536 |
| ENSG000001 | 2.81713877 | -0.7248028 | 0.57418388 | -1.2623182 | 0.2068343 | 0.50482226 |
| ENSG000001 | 2.67944151 | -0.7599836 | 0.60556834 | -1.2549924 | 0.20948153 | 0.50716765 |
| ENSG000001 | 2.19007992 | 0.82068931 | 0.65617938 | 1.25070876 | 0.21104075 | 0.50893868 |
| ENSG000000 | 140.519466 | -0.3823926 | 0.30620934 | -1.2487948 | 0.21174014 | 0.50998868 |
| ENSG000001 | 7.20266107 | 0.47752129 | 0.38303421 | 1.24668052 | 0.21251466 | 0.51084366 |
| ENSG000001 | 7.61908219 | -0.4583395 | 0.37232615 | -1.231016 | 0.21831687 | 0.5196653 |
| ENSG000001 | 4.35513537 | 0.67150389 | 0.54772301 | 1.22599174 | 0.22020178 | 0.52200728 |
| ENSG000001 | 3.97149722 | -0.726223 | 0.60500062 | -1.2003673 | 0.22999672 | 0.53285785 |
| ENSG000000 | 3.4542565 | -0.685259 | 0.57218799 | -1.1976117 | 0.23106822 | 0.53458024 |
| ENSG000001 | 4.28931571 | 0.6091031 | 0.50988943 | 1.19457879 | 0.23225164 | 0.53579194 |
| ENSG000001 | 29.8266858 | 0.31963251 | 0.26820754 | 1.19173575 | 0.23336488 | 0.53682778 |

| | | | | | | |
|---|---|---|---|---|---|---|
| ENSG000001 | 5.92576489 | -0.6337538 | 0.53351241 | -1.1878895 | 0.23487694 | 0.53934491 |
| ENSG000001 | 3.85113298 | 0.58592905 | 0.49483514 | 1.18408942 | 0.23637767 | 0.54027271 |
| ENSG000000 | 28.2319493 | -0.3703406 | 0.3133053 | -1.1820438 | 0.23718834 | 0.54100361 |
| ENSG000001 | 43.6005739 | 0.34356206 | 0.29213883 | 1.17602328 | 0.23958557 | 0.54350597 |
| ENSG000001 | 2.35325397 | 0.85535813 | 0.72827024 | 1.1745065 | 0.24019221 | 0.54409567 |
| ENSG000001 | 22.1988033 | -0.4459469 | 0.38472073 | -1.1591446 | 0.24639726 | 0.55071423 |
| ENSG000001 | 5.10754074 | -0.5493314 | 0.47860987 | -1.1477645 | 0.25106581 | 0.55714275 |
| ENSG000001 | 22.1052198 | 0.30116132 | 0.26273066 | 1.14627399 | 0.2516818 | 0.55784772 |
| ENSG000000 | 2.93612619 | 0.68051122 | 0.59418223 | 1.14529043 | 0.25208887 | 0.55798771 |
| ENSG000001 | 2.18928865 | 0.70130284 | 0.61725513 | 1.13616365 | 0.25588808 | 0.5618498 |
| ENSG000001 | 31.4556448 | -0.4061679 | 0.36090823 | -1.125405 | 0.26041744 | 0.56620936 |
| ENSG000000 | 1.84973202 | -0.9988794 | 0.88976634 | -1.1226311 | 0.2615942 | 0.5667983 |
| ENSG000001 | 170.847048 | 0.13872245 | 0.12357966 | 1.12253466 | 0.26163518 | 0.5667983 |
| ENSG000000 | 30.0311202 | 0.2418716 | 0.21595106 | 1.12002969 | 0.26270111 | 0.56809232 |
| ENSG000001 | 9.46781621 | -0.573367 | 0.51859738 | -1.105611 | 0.26889491 | 0.57451761 |
| ENSG000000 | 6.95844833 | -0.6157537 | 0.56265528 | -1.0943712 | 0.27379223 | 0.58058128 |
| ENSG000001 | 1.27076053 | 0.95890835 | 0.88236356 | 1.08674971 | 0.27714744 | 0.58542951 |
| ENSG000002 | 1.26477387 | 0.84946691 | 0.78875182 | 1.07697617 | 0.28149091 | 0.58968625 |
| ENSG000001 | 10.9949666 | -0.4281 | 0.40040562 | -1.0691659 | 0.28499492 | 0.5928266 |
| ENSG000001 | 1.75373891 | 1.14937051 | 1.0786673 | 1.06554681 | 0.28662855 | 0.59449067 |
| ENSG000001 | 273.650874 | -0.1623855 | 0.15280537 | -1.0626947 | 0.28792044 | 0.59601092 |
| ENSG000002 | 1.59391003 | -0.9388367 | 0.88386249 | -1.0621977 | 0.28814596 | 0.59601092 |
| ENSG000000 | 9.89115047 | -0.3474298 | 0.32782255 | -1.0598106 | 0.28923079 | 0.59652005 |
| ENSG000001 | 2.72247038 | -1.0292703 | 0.97118117 | -1.0598128 | 0.28922976 | 0.59652005 |
| ENSG000000 | 8.59078563 | 0.36073125 | 0.34363775 | 1.04974279 | 0.29383638 | 0.60009745 |
| ENSG000001 | 8.86646017 | 0.35457055 | 0.33862527 | 1.04708829 | 0.29505886 | 0.6014795 |
| ENSG000001 | 9.10410486 | -0.465916 | 0.44948886 | -1.0365463 | 0.29994734 | 0.60584165 |
| ENSG000002 | 201.309503 | 0.188276 | 0.18184629 | 1.03535793 | 0.30050179 | 0.60645633 |
| ENSG000001 | 30.3387535 | -0.3967202 | 0.38358387 | -1.0342462 | 0.30102107 | 0.60715204 |
| ENSG000000 | 2.802714 | -0.6868016 | 0.66669384 | -1.0301605 | 0.30293469 | 0.61004605 |
| ENSG000000 | 2.48898617 | -0.6385658 | 0.62337745 | -1.0243646 | 0.30566308 | 0.61298709 |
| ENSG000001 | 2.78018142 | 0.55028737 | 0.54450808 | 1.01061378 | 0.31220132 | 0.61836547 |
| ENSG000001 | 32.7706006 | -0.1922081 | 0.19072099 | -1.0077972 | 0.31355181 | 0.61989378 |
| ENSG000001 | 10.5395239 | -0.2891717 | 0.28735059 | -1.0063374 | 0.31425329 | 0.62107803 |
| ENSG000000 | 109.261258 | -0.1969223 | 0.19722924 | -0.9984439 | 0.31806417 | 0.625132 |
| ENSG000001 | 10.1173821 | 0.32259154 | 0.32340104 | 0.99749691 | 0.31852337 | 0.62557046 |
| ENSG000001 | 4.90616028 | -0.557075 | 0.55849078 | -0.997465 | 0.31853885 | 0.62557046 |
| ENSG000001 | 9.38410581 | -0.3201015 | 0.32112053 | -0.9968265 | 0.31884872 | 0.625849 |
| ENSG000001 | 1.37673554 | -0.8289908 | 0.83792998 | -0.9893319 | 0.32250079 | 0.62896992 |
| ENSG000001 | 1.77051046 | 0.81679147 | 0.83083569 | 0.98309627 | 0.32556006 | 0.63158339 |
| ENSG000001 | 3.08901302 | 0.64377485 | 0.65574518 | 0.98174545 | 0.32622527 | 0.63190127 |
| ENSG000000 | 2.28498843 | -0.7446685 | 0.76394219 | -0.9747708 | 3.30E-01 | 0.63489078 |
| ENSG000000 | 4.82842616 | 0.62190955 | 0.63841823 | 0.97414127 | 0.3299864 | 0.63507619 |

| | | | | | | |
|---|---|---|---|---|---|---|
| ENSG000001 | 2.901361 | 0.77017309 | 0.79584636 | 0.96774092 | 0.33317378 | 0.63814061 |
| ENSG000001 | 4.02177464 | 0.82618748 | 0.85850858 | 0.96235203 | 0.3358728 | 0.63977137 |
| ENSG000001 | 6.12568223 | 0.3987726 | 0.41454851 | 0.96194437 | 0.33607755 | 0.63977137 |
| ENSG000001 | 8.54837935 | 0.35873604 | 0.37364018 | 0.96011097 | 0.33699937 | 0.64068937 |
| ENSG000001 | 2.89611342 | -0.5622406 | 0.58904728 | -0.9544914 | 0.33983494 | 0.64372172 |
| ENSG000000 | 196.454081 | -0.188574 | 0.19844701 | -0.9502485 | 0.34198598 | 0.64616192 |
| ENSG000001 | 3.46033747 | -0.6545305 | 0.69418371 | -0.9428779 | 0.34574334 | 0.649069 |
| ENSG000001 | 1.27153096 | 0.98832974 | 1.05365962 | 0.93799717 | 0.34824586 | 0.65120848 |
| ENSG000001 | 13.0952733 | 0.27249047 | 0.290633 | 0.93757581 | 0.34846245 | 0.65135096 |
| ENSG000000 | 3.65869111 | -0.4640283 | 0.49692949 | -0.9337911 | 0.35041166 | 0.65345494 |
| ENSG000000 | 25.108003 | 0.27164609 | 0.29481459 | 0.92141329 | 0.35683469 | 0.65849126 |
| ENSG000001 | 2.32713809 | -0.6159523 | 0.67179506 | -0.9168753 | 0.35920801 | 0.65979981 |
| ENSG000001 | 6.58827284 | 0.39491644 | 0.43338292 | 0.91124135 | 0.36216822 | 0.66298781 |
| ENSG000001 | 20.7275745 | -0.2846356 | 0.31241755 | -0.9110745 | 0.36225615 | 0.66298781 |
| ENSG000001 | 1.90910458 | 0.78306571 | 0.85930232 | 0.91128081 | 0.36214743 | 0.66298781 |
| ENSG000001 | 13.1865592 | -0.2683917 | 0.29605916 | -0.9065477 | 0.36464605 | 0.66525268 |
| ENSG000001 | 28.3926337 | -0.2560231 | 0.28516064 | -0.8978205 | 0.36928128 | 0.66975165 |
| ENSG000001 | 3.13655084 | 0.4528562 | 0.50480089 | 0.89709864 | 0.36966628 | 0.66981505 |
| ENSG000001 | 3.02688642 | -0.4718635 | 0.52681679 | -0.8956881 | 0.37041937 | 0.67019425 |
| ENSG000001 | 2.63192844 | -0.5952651 | 0.66575856 | -0.8941155 | 0.37126008 | 0.6704582 |
| ENSG000001 | 4.21726763 | 0.47762912 | 0.53529964 | 0.89226497 | 0.37225092 | 0.67164726 |
| ENSG000000 | 137.996076 | -0.2322271 | 0.26067416 | -0.8908711 | 0.37299834 | 0.67215495 |
| ENSG000001 | 1.33172728 | -0.9011085 | 1.01609683 | -0.8868333 | 0.37516868 | 0.67340389 |
| ENSG000000 | 45.359038 | 0.18227428 | 0.20563334 | 0.8864043 | 0.3753997 | 0.6734785 |
| ENSG000001 | 4.57545967 | -0.5067092 | 0.57214704 | -0.8856276 | 0.37581825 | 0.67386967 |
| ENSG000001 | 2.51958072 | -0.5777121 | 0.65354079 | -0.8839725 | 0.37671104 | 0.6750923 |
| ENSG000001 | 10.6218894 | -0.315115 | 0.3632998 | -0.867369 | 0.38573986 | 0.68138194 |
| ENSG000001 | 4.41778681 | -0.3881603 | 0.45029888 | -0.8620058 | 0.3886843 | 0.68305399 |
| ENSG000001 | 5.70879456 | 0.33790201 | 0.39171693 | 0.86261783 | 0.38834762 | 0.68305399 |
| ENSG000001 | 1.30806429 | -0.7692777 | 0.89643123 | -0.8581558 | 0.39080644 | 0.68464545 |
| ENSG000001 | 17.6694894 | 0.19932963 | 0.23224952 | 0.85825637 | 0.39075092 | 0.68464545 |
| ENSG000000 | 5.2905509 | 0.32080443 | 0.37570943 | 0.85386314 | 0.39318083 | 0.68686248 |
| ENSG000001 | 5.19832777 | 0.37212531 | 0.43721228 | 0.85113189 | 0.39469609 | 0.68808015 |
| ENSG000001 | 4.18916344 | -0.418159 | 0.49956682 | -0.8370432 | 0.4025683 | 0.69659755 |
| ENSG000000 | 4.81952338 | 0.39454498 | 0.47211455 | 0.83569755 | 4.03E-01 | 0.69672317 |
| ENSG000001 | 2.59367904 | 0.5374252 | 0.64855768 | 0.82864672 | 0.40730434 | 0.69979458 |
| ENSG000001 | 2.18229479 | -0.4973222 | 0.62316931 | -0.7980532 | 0.42483964 | 0.71323836 |
| ENSG000001 | 2.69739639 | -0.4889849 | 0.61410502 | -0.7962562 | 0.42588313 | 0.71364215 |
| ENSG000001 | 19.5159532 | -0.2181444 | 0.27466147 | -0.7942299 | 0.4270616 | 0.71459873 |
| ENSG000001 | 6.9328857 | -0.3811472 | 0.48038215 | -0.7934249 | 0.42753031 | 0.71518559 |
| ENSG000000 | 7.59325096 | -0.2812717 | 0.35675222 | -0.7884232 | 4.30E-01 | 0.71723626 |
| ENSG000001 | 2.26285604 | 0.536543 | 0.68616766 | 0.78194155 | 0.43424892 | 0.71917412 |
| ENSG000001 | 7.0905918 | 0.3290519 | 0.42056472 | 0.78240492 | 0.43397665 | 0.71917412 |

| | | | | | | |
|---|---|---|---|---|---|---|
| ENSG000001( | 6.44782715 | -0.3295907 | 0.42598084 | -0.773722 | 0.43909523 | 0.72178325 |
| ENSG000001: | 6.97596946 | -0.3581404 | 0.46468949 | -0.770709 | 0.44087946 | 0.72323852 |
| ENSG000000: | 4.63583914 | 0.4704806 | 0.61114117 | 0.76983947 | 0.44139512 | 0.72334736 |
| ENSG000001( | 6.78897502 | 0.3158925 | 0.41418484 | 0.76268485 | 0.44565137 | 0.72528279 |
| ENSG000001( | 8.18968777 | -0.3030391 | 0.39830674 | -0.7608183 | 0.44676558 | 0.7260295 |
| ENSG000001( | 5.82962923 | 0.31844002 | 0.41896329 | 0.76006665 | 0.44721474 | 0.72606344 |
| ENSG000001( | 1.83324878 | -0.5088047 | 0.67724349 | -0.7512877 | 0.45247956 | 0.72911951 |
| ENSG000001( | 21.8538601 | -0.1506918 | 0.20157866 | -0.7475582 | 0.45472666 | 0.73044273 |
| ENSG000001. | 26.6207041 | 0.17608101 | 0.23577275 | 0.74682512 | 0.45516913 | 0.73073377 |
| ENSG000001( | 3.30887085 | 0.40703165 | 0.54818528 | 0.74250745 | 0.45777994 | 0.73254357 |
| ENSG000001( | 1.49306792 | 0.60365078 | 0.81318323 | 0.74233058 | 0.45788707 | 0.73257903 |
| ENSG000001: | 12.1824716 | -0.2850817 | 0.38415157 | -0.7421074 | 0.45802229 | 0.73257903 |
| ENSG000000! | 15.2383601 | -0.213726 | 0.28911531 | -0.7392414 | 0.4597604 | 0.73347485 |
| ENSG000001( | 3.29240622 | 0.40810136 | 0.55859074 | 0.73059098 | 0.46502902 | 0.73832422 |
| ENSG000001: | 13.6958784 | -0.2210185 | 0.30654378 | -0.7210014 | 0.47090867 | 0.74246661 |
| ENSG000001( | 2.09669124 | 0.45979781 | 0.63804165 | 0.72063917 | 0.47113155 | 0.74262485 |
| ENSG000000( | 45.0462964 | 0.26603761 | 0.36982862 | 0.71935376 | 4.72E-01 | 7.43E-01 |
| ENSG000001: | 62.2956209 | -0.1730553 | 0.24047104 | -0.7196515 | 0.4717396 | 0.74290641 |
| ENSG000001: | 3.02714484 | 0.40732236 | 0.56827628 | 0.71676819 | 0.47351714 | 0.74400728 |
| ENSG000002: | 641.542796 | -0.2151185 | 0.30474768 | -0.7058906 | 0.48025616 | 0.74863641 |
| ENSG000001! | 9.77162568 | 0.23397012 | 0.33309437 | 0.7024139 | 0.48242108 | 0.75047051 |
| ENSG000001( | 3.2738685 | 0.38358095 | 0.54820923 | 0.69969809 | 0.48411587 | 0.75175313 |
| ENSG000001' | 4.28351599 | -0.3757226 | 0.53779349 | -0.6986373 | 0.48477874 | 0.75201186 |
| ENSG000001( | 10.7123185 | -0.265437 | 0.38274796 | -0.6935035 | 0.48799366 | 0.75429646 |
| ENSG000001( | 9.40095759 | -0.2842379 | 0.41125764 | -0.6911432 | 0.48947557 | 0.75562362 |
| ENSG000000! | 1.72172798 | 0.48845078 | 0.71208654 | 0.685943 | 0.49274905 | 0.7571831 |
| ENSG000000. | 12.3874286 | 0.18458315 | 0.27040781 | 0.68261028 | 0.49485314 | 0.75805163 |
| ENSG000001: | 3.71083143 | 0.36819848 | 0.55013824 | 0.66928356 | 0.50331461 | 0.76275237 |
| ENSG000001! | 1.82562411 | 0.42731893 | 0.64301208 | 0.66455817 | 0.50633313 | 0.76422766 |
| ENSG000001! | 22.6316679 | 0.21514049 | 0.32532082 | 0.66131792 | 0.50840845 | 0.76597351 |
| ENSG000001: | 17.1914444 | -0.1930046 | 0.2966049 | -0.6507129 | 0.51523185 | 0.76909443 |
| ENSG000001: | 3.85727234 | -0.3841934 | 0.58992848 | -0.6512542 | 0.51488243 | 0.76909443 |
| ENSG000001' | 1.35233326 | -0.6669493 | 1.02930162 | -0.647963 | 0.5170089 | 0.77008219 |
| ENSG000001' | 1.71670927 | -0.41659 | 0.64765428 | -0.643229 | 0.52007553 | 0.7729405 |
| ENSG000001. | 1.8603751 | 0.5298995 | 0.82576192 | 0.64170978 | 0.52106164 | 0.77332448 |
| ENSG000001! | 28.7973158 | -0.2435004 | 0.38595363 | -0.6309059 | 0.52810204 | 0.77803877 |
| ENSG000001. | 19.5449689 | -0.1966069 | 0.31361743 | -0.6269005 | 0.53072444 | 0.77978092 |
| ENSG000000' | 2.15585686 | 0.47799647 | 0.76571813 | 0.62424598 | 0.53246605 | 0.78093427 |
| ENSG000000! | 17.8135367 | -0.1481877 | 0.23971521 | -0.6181823 | 0.53645516 | 0.78345899 |
| ENSG000001! | 1.56337135 | -0.4755992 | 0.76927199 | -0.6182459 | 0.53641328 | 0.78345899 |
| ENSG000001! | 2.35100031 | 0.3692493 | 0.5973579 | 0.61813746 | 0.53648473 | 0.78345899 |
| ENSG000001( | 6.79105215 | -0.3149957 | 0.51016277 | -0.6174417 | 0.53694345 | 0.7835716 |
| ENSG000002: | 10.7573323 | 0.23810241 | 0.38886385 | 0.61230275 | 0.54033747 | 0.78615991 |

| | | | | | | |
|---|---|---|---|---|---|---|
| ENSG000001 | 2.02197413 | 0.40775789 | 0.67031337 | 0.60830935 | 0.54298232 | 0.78688966 |
| ENSG000001 | 93.1231491 | 0.14141202 | 0.23231302 | 0.60871329 | 0.54271449 | 0.78688966 |
| ENSG000001 | 8.65984279 | 0.20046996 | 0.32963248 | 0.60816204 | 0.54308001 | 0.78688966 |
| ENSG000001 | 22.9135684 | 0.13814216 | 0.22728189 | 0.60780099 | 0.54331948 | 0.78703095 |
| ENSG000000 | 84.3029963 | -0.2704069 | 0.44664067 | -0.6054238 | 0.54489741 | 0.78861595 |
| ENSG000001 | 2.27207749 | 0.40705935 | 0.6787996 | 0.5996753 | 0.54872265 | 0.79107679 |
| ENSG000001 | 1.81857056 | -0.484153 | 0.8114536 | -0.5966491 | 0.55074171 | 0.79247973 |
| ENSG000000 | 2.15200617 | -0.5750015 | 0.97130763 | -0.591987 | 0.55385926 | 0.79432578 |
| ENSG000001 | 2.35977786 | 0.43541338 | 0.73689582 | 0.59087508 | 0.55460412 | 0.79511185 |
| ENSG000001 | 1.60883025 | -0.421075 | 0.71641845 | -0.5877501 | 0.55670004 | 0.79630427 |
| ENSG000001 | 8.90063333 | 0.20886396 | 0.3556006 | 0.58735549 | 0.55696498 | 0.79636343 |
| ENSG000001 | 7.7283947 | 0.22209192 | 0.39067996 | 0.56847534 | 0.56971224 | 0.80750866 |
| ENSG000000 | 4.49261572 | 0.23504694 | 0.41440324 | 0.56719377 | 0.57058254 | 0.80780387 |
| ENSG000001 | 21.9088811 | 0.14131444 | 0.24971067 | 0.56591272 | 0.57145312 | 0.80833712 |
| ENSG000001 | 1.86128448 | -0.575334 | 1.01748319 | -0.5654482 | 0.57176896 | 0.80843889 |
| ENSG000001 | 18.1719591 | 0.18275809 | 0.32474878 | 0.5627676 | 0.57359315 | 0.80969092 |
| ENSG000001 | 2.21710562 | -0.369694 | 0.66324587 | -0.5574011 | 0.5772534 | 0.81212058 |
| ENSG000001 | 13.6254706 | 0.28074402 | 0.50423009 | 0.55677761 | 0.57767938 | 0.81243702 |
| ENSG000002 | 1.35752193 | -0.523347 | 0.94080464 | -0.5562759 | 0.57802223 | 0.81263638 |
| ENSG000001 | 9.80165991 | 0.18786649 | 0.34612984 | 0.54276307 | 0.58729294 | 0.81704755 |
| ENSG000001 | 7.74991919 | 0.17685303 | 0.33321336 | 0.53075013 | 0.59559194 | 0.82129471 |
| ENSG000000 | 1.34437545 | -0.5165719 | 0.97399032 | -0.5303666 | 0.5958578 | 0.82141603 |
| ENSG000002 | 1.94838477 | -0.3698956 | 0.69930681 | -0.528946 | 0.59684288 | 0.82193869 |
| ENSG000000 | 75.4230916 | 0.07680995 | 0.14717624 | 0.52189096 | 0.60174625 | 0.82475634 |
| ENSG000001 | 8.11466683 | -0.1752139 | 0.33900963 | -0.5168404 | 0.60526758 | 0.8274985 |
| ENSG000001 | 11.5882191 | -0.225463 | 0.4392364 | -0.5133068 | 0.60773673 | 0.82866368 |
| ENSG000000 | 1.407638 | 0.42817806 | 0.85013927 | 0.50365637 | 0.61450287 | 0.83125105 |
| ENSG000000 | 2.7751214 | 0.27040477 | 0.53683866 | 0.5036984 | 0.61447333 | 0.83125105 |
| ENSG000001 | 20.0020201 | 0.11028033 | 0.218418 | 0.50490495 | 0.6136256 | 0.83125105 |
| ENSG000001 | 2.79057901 | -0.2943835 | 0.58419687 | -0.5039115 | 0.61432358 | 0.83125105 |
| ENSG000000 | 68.4116014 | 0.12064701 | 0.24160389 | 0.49935874 | 0.61752668 | 0.8327825 |
| ENSG000001 | 3.00757692 | -0.3812145 | 0.76571736 | -0.4978528 | 0.61858778 | 0.83377654 |
| ENSG000000 | 4.23111085 | -0.2373954 | 0.48029347 | -0.4942715 | 0.62111446 | 0.83476 |
| ENSG000001 | 13.7927687 | -0.17663 | 0.359213 | -0.4917137 | 0.62292174 | 0.83591768 |
| ENSG000001 | 13.0827628 | -0.1671486 | 0.34294713 | -0.487389 | 0.62598271 | 0.83751061 |
| ENSG000001 | 7.38453158 | 0.17339802 | 0.35783697 | 0.48457268 | 0.6279795 | 0.8381587 |
| ENSG000001 | 7.81412916 | 0.18920708 | 0.39671055 | 0.47693988 | 0.63340493 | 0.84132442 |
| ENSG000001 | 3.30398272 | 0.27898576 | 0.58881997 | 0.47380485 | 0.63563908 | 0.84281445 |
| ENSG000001 | 9.91851344 | -0.1712608 | 0.36668557 | -0.4670508 | 0.64046355 | 0.84459259 |
| ENSG000000 | 178.09952 | -0.1401619 | 0.30326296 | -0.4621795 | 6.44E-01 | 0.84735019 |
| ENSG000001 | 30.506813 | 0.21051184 | 0.45668131 | 0.46096006 | 0.64482726 | 0.847611 |
| ENSG000001 | 4.97008468 | -0.2012413 | 0.43809414 | -0.4593562 | 0.64597836 | 0.84789905 |
| ENSG000001 | 1.64112394 | 0.37297244 | 0.81229138 | 0.4591609 | 0.64611863 | 0.84799135 |

| | | | | | | |
|---|---|---|---|---|---|---|
| ENSG0000010 | 4.76163934 | -0.1780724 | 0.39630187 | -0.4493353 | 0.65318976 | 0.85247654 |
| ENSG0000018 | 8.73464535 | -0.1929032 | 0.43555154 | -0.4428942 | 0.65784227 | 0.85556483 |
| ENSG0000010 | 65.0617964 | -0.0796108 | 0.1828764 | -0.435326 | 0.66332584 | 0.85876753 |
| ENSG0000017 | 7.13436287 | -0.1595521 | 0.36901214 | -0.4323763 | 0.66546793 | 0.85979619 |
| ENSG0000007 | 7.66191306 | 0.15902298 | 0.37135843 | 0.42821965 | 0.66849121 | 0.86159036 |
| ENSG0000000 | 4.41878755 | -0.2054824 | 0.48143365 | -0.4268136 | 0.66951512 | 0.86241787 |
| ENSG0000019 | 116.194726 | 0.09380614 | 0.2217996 | 0.42293193 | 0.67234492 | 0.86398532 |
| ENSG0000010 | 1.53331134 | -0.3123727 | 0.74276364 | -0.4205547 | 0.6740803 | 0.8651304 |
| ENSG0000010 | 4.19438681 | -0.2232942 | 0.53403667 | -0.4181251 | 0.67585562 | 0.86602014 |
| ENSG0000019 | 3.27081224 | 0.23718896 | 0.56967928 | 0.41635525 | 0.67715007 | 0.8664862 |
| ENSG0000014 | 1.47180283 | 0.31377874 | 0.76025997 | 0.41272558 | 0.67980768 | 0.86706958 |
| ENSG0000017 | 126.351094 | -0.0917717 | 0.22265333 | -0.412173 | 0.68021261 | 0.86706958 |
| ENSG0000021 | 1.84129316 | -0.3135591 | 0.76240719 | -0.4112751 | 0.68087083 | 0.86740196 |
| ENSG0000010 | 6.41851576 | -0.1618419 | 0.3945907 | -0.4101513 | 0.68169495 | 0.86778718 |
| ENSG0000011 | 73.6694195 | -0.0726046 | 0.17838286 | -0.4070154 | 0.68399664 | 0.86898936 |
| ENSG0000001 | 1.3038181 | 0.31585594 | 0.79132155 | 0.39914993 | 0.68978273 | 0.87206951 |
| ENSG0000000 | 2.64930735 | 0.21296075 | 0.54715488 | 0.38921476 | 0.69711729 | 0.87585553 |
| ENSG0000010 | 2.06371821 | -0.2622708 | 0.67478501 | -0.3886731 | 0.69751799 | 0.87590492 |
| ENSG0000015 | 1.60474197 | 0.25721694 | 0.66657487 | 0.38587855 | 0.69958662 | 0.87716253 |
| ENSG0000019 | 43.39974 | -0.1088194 | 0.28243378 | -0.3852917 | 0.70002131 | 0.87718184 |
| ENSG0000011 | 50.766361 | -0.1160929 | 0.30775078 | -0.3772302 | 0.70600252 | 0.88008931 |
| ENSG0000010 | 7.64887607 | 0.13826392 | 0.36774694 | 0.37597572 | 0.70693495 | 0.88070829 |
| ENSG0000000 | 4.57528335 | -0.1604339 | 0.4288078 | -0.3741395 | 0.70830052 | 0.88169707 |
| ENSG0000010 | 3.8727638 | 0.16620797 | 0.44639354 | 0.37233508 | 0.70964339 | 0.88205012 |
| ENSG0000009 | 3.98080969 | 0.25028193 | 0.67302024 | 0.37187875 | 0.70998313 | 0.88214876 |
| ENSG0000011 | 8.99815658 | -0.1474706 | 0.39727278 | -0.3712073 | 0.71048312 | 0.88231785 |
| ENSG0000017 | 1.68198746 | 0.28055751 | 0.75951162 | 0.36939199 | 0.71183557 | 0.88300241 |
| ENSG0000011 | 42.3198939 | -0.07613 | 0.20841406 | -0.3652825 | 0.71490058 | 0.88508371 |
| ENSG0000017 | 10.1875454 | 0.11808364 | 0.32390252 | 0.36456537 | 0.71543587 | 0.88527484 |
| ENSG0000018 | 1.62941031 | -0.2652995 | 0.7286441 | -0.3641002 | 0.71578316 | 0.88554335 |
| ENSG0000011 | 76.6326839 | 0.09544573 | 0.26585427 | 0.35901524 | 0.71958369 | 0.88816057 |
| ENSG0000015 | 20.7728251 | 0.07812707 | 0.22100559 | 0.35350722 | 0.72370822 | 0.89047638 |
| ENSG0000014 | 6.98146208 | 0.15828245 | 0.44863117 | 0.35281197 | 0.72422941 | 0.89063026 |
| ENSG0000020 | 2.83667403 | 0.19155505 | 0.54815987 | 0.34945106 | 0.7267507 | 0.89237563 |
| ENSG0000018 | 109.829838 | -0.0580116 | 0.16647651 | -0.3484669 | 0.72748954 | 0.8927377 |
| ENSG0000010 | 2.40437411 | -0.2381299 | 0.69015385 | -0.3450388 | 0.73006518 | 0.893998 |
| ENSG0000010 | 24.858631 | 0.1069176 | 0.3106969 | 0.34412188 | 0.73075463 | 0.89430026 |
| ENSG0000019 | 3.26344429 | -0.1853182 | 0.54044741 | -0.3428977 | 0.73167541 | 0.89476124 |
| ENSG0000011 | 1.74502315 | -0.2318289 | 0.68005171 | -0.3408989 | 0.73317968 | 0.89556399 |
| ENSG0000017 | 18.0944572 | 0.0769565 | 0.22645994 | 0.33982389 | 0.73398916 | 0.89556399 |
| ENSG0000018 | 3.86971226 | -0.2133085 | 0.6284609 | -0.3394141 | 0.73429777 | 0.89563918 |
| ENSG0000014 | 1.9493213 | 0.24133102 | 0.71338632 | 0.33828938 | 0.73514513 | 0.89575729 |
| ENSG0000009 | 17.6798133 | 0.09567115 | 0.28363554 | 0.3373031 | 0.73588842 | 0.89590482 |

| | | | | | | |
|---|---|---|---|---|---|---|
| ENSG000000 | 2.12023105 | 0.26235409 | 0.77900665 | 0.3367803 | 0.73628253 | 0.89602537 |
| ENSG000001 | 1.65782144 | -0.2513118 | 0.78459989 | -0.3203057 | 0.74873661 | 0.90398704 |
| ENSG000001 | 53.312168 | -0.0658227 | 0.20698457 | -0.3180076 | 0.75047915 | 0.90509936 |
| ENSG000002 | 1.4918724 | -0.3449118 | 1.08923703 | -0.3166545 | 0.75150577 | 0.90544966 |
| ENSG000001 | 283.800268 | 0.04170842 | 0.13210457 | 0.31572276 | 0.75221295 | 0.90583864 |
| ENSG000001 | 2.59161561 | 0.21028278 | 0.66887026 | 0.31438501 | 0.75322864 | 0.90653142 |
| ENSG000001 | 2.82199478 | 0.18481547 | 0.59369682 | 0.31129604 | 0.75557558 | 0.90733256 |
| ENSG000001 | 1.89707979 | 0.24879106 | 0.79924782 | 0.3112815 | 0.75558663 | 0.90733256 |
| ENSG000001 | 5.9262382 | 0.13756071 | 0.45938436 | 0.29944578 | 0.76459993 | 0.91175293 |
| ENSG000001 | 1.53353327 | 0.22117928 | 0.75429534 | 0.29322637 | 0.76934913 | 0.91361007 |
| ENSG000001 | 1.42621456 | 0.21518279 | 0.7350064 | 0.29276314 | 0.7697032 | 0.91373991 |
| ENSG000001 | 7.40444921 | -0.1029952 | 0.35473638 | -0.2903428 | 0.77155399 | 0.91486067 |
| ENSG000001 | 1.87635717 | -0.2381592 | 0.8388915 | -0.2838975 | 0.77648891 | 0.91605441 |
| ENSG000001 | 6.33418522 | 0.11491368 | 0.40736705 | 0.2820888 | 0.77787542 | 0.91653064 |
| ENSG000001 | 2.44450607 | -0.2051777 | 0.72884013 | -0.2815126 | 0.77831726 | 0.91674608 |
| ENSG000001 | 10.9195618 | 0.1009259 | 0.36096965 | 0.27959664 | 0.77978698 | 0.91753447 |
| ENSG000001 | 4.54583823 | -0.1903295 | 0.69505147 | -0.2738352 | 0.78421129 | 0.91962872 |
| ENSG000000 | 3.37817212 | 0.15215868 | 0.57913681 | 0.26273357 | 0.79275594 | 0.92446259 |
| ENSG000000 | 7.56800941 | -0.108172 | 0.41231421 | -0.2623534 | 0.79304903 | 0.92446259 |
| ENSG000001 | 5.73064043 | -0.0959288 | 0.36640337 | -0.2618119 | 0.79346646 | 0.92455075 |
| ENSG000001 | 12.923043 | 0.11208948 | 0.43257887 | 0.25911918 | 0.79554329 | 0.92565229 |
| ENSG000001 | 2.08408263 | -0.1766658 | 0.68295354 | -0.2586791 | 0.79588287 | 0.92576429 |
| ENSG000001 | 4.1057392 | -0.1259028 | 0.50841263 | -0.247639 | 0.8044137 | 0.93034668 |
| ENSG000000 | 2.60384933 | -0.1847818 | 0.75084337 | -0.246099 | 0.80560557 | 0.93087872 |
| ENSG000001 | 5.55928403 | -0.115312 | 0.47377451 | -0.24339 | 0.80770333 | 0.93175254 |
| ENSG000001 | 6.28455546 | -0.1009309 | 0.42154145 | -0.2394328 | 0.81076998 | 0.93296186 |
| ENSG000001 | 6.59884735 | -0.0857192 | 0.35812414 | -0.239356 | 0.81082951 | 0.93296186 |
| ENSG000002 | 1.9106335 | -0.1577078 | 0.68237269 | -0.2311168 | 0.81722406 | 0.9366746 |
| ENSG000001 | 5.1613333 | 0.12308297 | 0.54175551 | 0.22719283 | 0.8202738 | 0.9380189 |
| ENSG000000 | 12.2087813 | 0.07046176 | 0.32468935 | 0.21701285 | 0.82819833 | 0.94036231 |
| ENSG000001 | 4.52097698 | 0.11976508 | 0.55161043 | 0.21711896 | 0.82811564 | 0.94036231 |
| ENSG000000 | 7.07943091 | -0.0899252 | 0.41962028 | -0.2143015 | 0.83031198 | 0.94170382 |
| ENSG000001 | 6.01499207 | -0.0879398 | 0.41095275 | -0.2139901 | 0.83055475 | 0.94180294 |
| ENSG000001 | 4.02580336 | -0.1135238 | 0.53404266 | -0.2125744 | 0.83165893 | 0.94205491 |
| ENSG000001 | 76.5856976 | -0.0409225 | 0.19441267 | -0.2104927 | 0.83328311 | 0.94283316 |
| ENSG000000 | 2.05776707 | -0.1278148 | 0.61141656 | -0.209047 | 0.83441152 | 0.94306315 |
| ENSG000001 | 11.8384168 | 0.10886385 | 0.5193659 | 0.20960917 | 0.83397272 | 0.94306315 |
| ENSG000001 | 67.9090953 | -0.0545298 | 0.27235522 | -0.2002159 | 0.84131175 | 0.94533528 |
| ENSG000001 | 5.26663919 | -0.0898159 | 0.45000684 | -0.1995878 | 0.84180297 | 0.94571192 |
| ENSG000000 | 4.24898224 | -0.0951614 | 0.49354944 | -0.1928102 | 0.84710763 | 0.94799806 |
| ENSG000001 | 1.99262975 | -0.1309263 | 0.72206018 | -0.1813232 | 0.85611386 | 0.95034048 |
| ENSG000001 | 3.65574718 | -0.1020297 | 0.57441539 | -0.1776235 | 0.85901868 | 0.9518219 |
| ENSG000000 | 15.6695894 | -0.0600609 | 0.34355162 | -0.1748234 | 0.86121843 | 0.9522704 |

| | | | | | | |
|---|---|---|---|---|---|---|
| ENSG0000004 | 21.8229121 | -0.0600924 | 0.34424515 | -0.1745628 | 0.86142319 | 0.9522704 |
| ENSG0000011 | 3.84303804 | -0.0823671 | 0.47365463 | -0.1738969 | 0.86194649 | 0.95255959 |
| ENSG0000015 | 10.7963727 | -0.0552281 | 0.32505324 | -0.1699047 | 0.86508507 | 0.95418828 |
| ENSG0000018 | 5.63668613 | 0.079505 | 0.46667082 | 0.17036634 | 0.86472204 | 0.95418828 |
| ENSG0000015 | 1.55658457 | -0.1365774 | 0.82113876 | -0.1663269 | 0.86789973 | 0.9547634 |
| ENSG0000019 | 39.9156012 | 0.03116228 | 0.18768201 | 0.16603763 | 0.86812734 | 0.9547634 |
| ENSG0000000 | 1.51138844 | 0.14277982 | 0.89919302 | 0.15878662 | 0.87383699 | 0.95567977 |
| ENSG0000018 | 8.84591706 | -0.0637227 | 0.39586255 | -0.1609718 | 0.87211559 | 0.95567977 |
| ENSG0000018 | 3.0968758 | -0.0923437 | 0.61921246 | -0.1491308 | 0.88145041 | 0.95801207 |
| ENSG0000018 | 4.19963185 | 0.07025818 | 0.47522523 | 0.14784185 | 0.88246758 | 0.95844423 |
| ENSG0000004 | 52.8198003 | -0.0359849 | 0.24807713 | -0.1450551 | 0.88466737 | 0.95928645 |
| ENSG0000010 | 19.3064843 | -0.0398295 | 0.28622217 | -0.139156 | 0.88932691 | 0.96175826 |
| ENSG0000010 | 22.7809971 | 0.0379848 | 0.27637414 | 0.13743976 | 0.8906832 | 0.96230108 |
| ENSG0000010 | 7.23007182 | -0.0522682 | 0.38362822 | -0.136247 | 0.89162602 | 0.9625067 |
| ENSG0000010 | 10.8236284 | 0.06082174 | 0.44620761 | 0.13630817 | 0.89157767 | 0.9625067 |
| ENSG0000010 | 4.27510195 | 0.07220626 | 0.56417124 | 0.12798642 | 0.89815972 | 0.96467433 |
| ENSG0000019 | 23.5162459 | -0.0282739 | 0.22743233 | -0.1243179 | 0.90106358 | 0.96543952 |
| ENSG0000011 | 2.10587316 | 0.07277001 | 0.59491259 | 0.12232051 | 0.90264519 | 0.96616649 |
| ENSG0000011 | 2.247239 | -0.0811338 | 0.68459135 | -0.1185142 | 0.90566027 | 0.96645877 |
| ENSG0000015 | 847.15542 | 0.01405671 | 0.11734912 | 0.1197854 | 0.90465315 | 0.96645877 |
| ENSG0000015 | 1.77968549 | -0.0897807 | 0.76859113 | -0.116812 | 0.90700902 | 0.96656736 |
| ENSG0000011 | 2.0008451 | 0.08372005 | 0.75192823 | 0.11134048 | 0.91134636 | 0.9690599 |
| ENSG0000011 | 3.97500118 | -0.0579333 | 0.5462917 | -0.1060482 | 0.9155441 | 0.97080376 |
| ENSG0000011 | 12.1638148 | 0.0355518 | 0.33874662 | 0.104951 | 0.91641469 | 0.97097975 |
| ENSG0000011 | 11.0345625 | -0.0406317 | 0.40188723 | -0.1011022 | 0.91946931 | 0.97178664 |
| ENSG0000018 | 2.40081129 | -0.0606711 | 0.61672342 | -0.0983764 | 0.92163339 | 0.97210904 |
| ENSG0000011 | 7.58392354 | 0.03923361 | 0.40102831 | 0.09783252 | 0.92206528 | 0.9722708 |
| ENSG0000014 | 2.6184496 | -0.0696393 | 0.7216307 | -0.0965027 | 0.92312134 | 0.97229384 |
| ENSG0000001 | 9.03435725 | -0.0406141 | 0.44051847 | -0.092196 | 0.92654228 | 0.97364008 |
| ENSG0000014 | 11.2436609 | -0.0343816 | 0.37281696 | -0.0922211 | 0.92652234 | 0.97364008 |
| ENSG0000010 | 16.2974661 | -0.0215436 | 0.23659913 | -0.0910552 | 0.92744873 | 0.97364008 |
| ENSG0000000 | 4.36917547 | 0.04432501 | 0.53817002 | 0.08236247 | 0.93435848 | 0.97591183 |
| ENSG0000010 | 1.79659455 | 0.05656198 | 0.70718462 | 0.07998191 | 0.93625164 | 0.97632198 |
| ENSG0000011 | 36.590825 | -0.0201092 | 0.25380181 | -0.0792319 | 0.93684819 | 0.97632198 |
| ENSG0000010 | 3.50495767 | -0.040673 | 0.54012193 | -0.0753033 | 0.93997341 | 0.97771751 |
| ENSG0000010 | 13.4705173 | -0.0329616 | 0.44896696 | -0.0734166 | 0.9414746 | 0.97888355 |
| ENSG0000004 | 53.3055206 | 0.03401718 | 0.46732032 | 0.072792 | 0.94197164 | 0.97895604 |
| ENSG0000000 | 1.7273428 | 0.05237128 | 0.73205596 | 0.07154 | 0.94296799 | 0.97920441 |
| ENSG0000017 | 8.65756638 | -0.029683 | 0.41421811 | -0.0716604 | 0.9428722 | 0.97920441 |
| ENSG0000011 | 13.2373483 | -0.0244926 | 0.35013647 | -0.0699517 | 0.94423211 | 0.97970951 |
| ENSG0000008 | 1.71634142 | 0.04444112 | 0.64190689 | 0.06923297 | 0.94480418 | 0.97983407 |
| ENSG0000017 | 12.1761696 | -0.0227121 | 0.34334868 | -0.0661489 | 0.94725928 | 0.98125475 |
| ENSG0000014 | 17.1621827 | -0.0217046 | 0.34245727 | -0.0633791 | 0.94946462 | 0.98249834 |

| | | | | | | |
|---|---|---|---|---|---|---|
| ENSG000001 | 3.09481028 | -0.0316446 | 0.52360128 | -0.0604364 | 0.95180809 | 0.98327021 |
| ENSG000001 | 70.113631 | -0.0173454 | 0.29202365 | -0.0593972 | 0.95263575 | 0.98327021 |
| ENSG000001 | 8.26689221 | -0.0203223 | 0.33299656 | -0.0610284 | 0.95133658 | 0.98327021 |
| ENSG000000 | 57.9483901 | 0.02408886 | 0.41728538 | 0.05772754 | 9.54E-01 | 0.98363059 |
| ENSG000001 | 1.62813781 | 0.05354336 | 0.93670532 | 0.05716138 | 0.95441665 | 0.98363059 |
| ENSG000001 | 2.36608804 | -0.0410777 | 0.76014198 | -0.0540395 | 0.95690372 | 0.98449906 |
| ENSG000001 | 3.47191903 | -0.0233756 | 0.6212467 | -0.0376269 | 0.96998513 | 0.99091007 |
| ENSG000001 | 2.13334066 | -0.0258187 | 0.68509789 | -0.0376861 | 0.96993797 | 0.99091007 |
| ENSG000000 | 2.26157757 | 0.02592284 | 0.70195196 | 0.03692964 | 0.9705411 | 0.99106046 |
| ENSG000001 | 2.12066918 | 0.0277096 | 0.76993604 | 0.03598948 | 0.97129075 | 0.99106699 |
| ENSG000001 | 1.30674508 | -0.0290308 | 0.8313506 | -0.03492 | 0.97214351 | 0.99106699 |
| ENSG000001 | 7.97799249 | 0.0170687 | 0.51679749 | 0.03302784 | 0.97365239 | 0.99122138 |
| ENSG000001 | 19.7223577 | 0.01436301 | 0.4890911 | 0.02936674 | 0.9765721 | 0.9923208 |
| ENSG000001 | 1.95154975 | 0.01812099 | 0.61948615 | 0.02925164 | 0.97666389 | 0.9923208 |
| ENSG000001 | 3.81676855 | -0.0132946 | 0.59188262 | -0.0224615 | 0.98207979 | 0.99316457 |
| ENSG000001 | 13.6637433 | -0.0066007 | 0.29109819 | -0.0226753 | 0.98190929 | 0.99316457 |
| ENSG000002 | 2.68488528 | 0.01161801 | 0.51121868 | 0.02272612 | 0.98186874 | 0.99316457 |
| ENSG000000 | 8.94342396 | 0.0065676 | 0.35853613 | 0.01831783 | 9.85E-01 | 0.99482229 |
| ENSG000001 | 6.27919865 | -0.0082513 | 0.5101797 | -0.0161732 | 0.9870962 | 0.99587584 |
| ENSG000001 | 14.3062544 | 0.00392755 | 0.32722343 | 0.01200266 | 0.99042349 | 0.99708554 |
| ENSG000001 | 2.43219048 | 0.00751391 | 0.69238693 | 0.01085218 | 0.99134138 | 0.99731839 |
| ENSG000001 | 5.34900279 | 0.00567718 | 0.50197623 | 0.01130967 | 0.99097638 | 0.99731839 |
| ENSG000001 | 2.02195235 | -0.0023487 | 0.70361079 | -0.0033381 | 0.99733657 | 0.99881959 |
| ENSG000001 | 1.62706984 | 0.00160763 | 0.95743204 | 0.00167911 | 0.99866027 | 0.99940221 |
| ENSG000000 | 1.15683144 | 0.96994616 | 0.85232252 | 1.13800367 | 2.55E-01 | NA |
| ENSG000000 | 1.05073509 | 1.61858922 | 0.93542475 | 1.73032542 | 8.36E-02 | NA |
| ENSG000000 | 0 | NA | NA | NA | NA | NA |
| ENSG000000 | 0.65461564 | -0.5390498 | 1.18279486 | -0.4557424 | 6.49E-01 | NA |
| ENSG000000 | 0 | NA | NA | NA | NA | NA |
| ENSG000000 | 0 | NA | NA | NA | NA | NA |
| ENSG000000 | 0.91847241 | -1.6667979 | 1.15554328 | -1.4424366 | 0.14917925 | NA |
| ENSG000000 | 0 | NA | NA | NA | NA | NA |
| ENSG000000 | 0 | NA | NA | NA | NA | NA |
| ENSG000000 | 1.19860032 | 0.91880568 | 0.82734487 | 1.11054739 | 0.26676322 | NA |
| ENSG000000 | 0.00984842 | 0.19866539 | 3.04669343 | 0.06520688 | 0.94800928 | NA |
| ENSG000000 | 0.22790065 | 0.19820861 | 1.66955141 | 0.11871968 | 0.90549744 | NA |
| ENSG000000 | 0 | NA | NA | NA | NA | NA |
| ENSG000000 | 0 | NA | NA | NA | NA | NA |
| ENSG000000 | 0.33019986 | -0.3919726 | 1.7630694 | -0.222324 | 0.8240617 | NA |
| ENSG000000 | 0.20536481 | 0.74391209 | 2.44721022 | 0.30398373 | 0.76114029 | NA |
| ENSG000000 | 0 | NA | NA | NA | NA | NA |
| ENSG000000 | 0 | NA | NA | NA | NA | NA |
| ENSG000000 | 0 | NA | NA | NA | NA | NA |

| | | | | | | |
|---|---|---|---|---|---|---|
| ENSG0000001 | 0 | NA | NA | NA | NA | NA |
| ENSG0000001 | 0.18906859 | 0.84022181 | 2.81986977 | 0.29796476 | 0.76573006 | NA |
| ENSG0000002 | 0.28435067 | -0.8136162 | 1.78656379 | -0.4554084 | 0.64881545 | NA |
| ENSG0000002 | 0.01969684 | 0.0417549 | 3.04641238 | 0.01370625 | 0.98906434 | NA |
| ENSG0000003 | 0.24196645 | 0.33522165 | 1.81705401 | 0.18448634 | 0.85363194 | NA |
| ENSG0000003 | 0.7739332 | -0.6788341 | 1.09285889 | -0.6211544 | 0.53449805 | NA |
| ENSG0000003 | 0 | NA | NA | NA | NA | NA |
| ENSG0000004 | 0.38311506 | 0.48019495 | 1.35173566 | 0.35524324 | 0.72240738 | NA |
| ENSG0000004 | 0.07157543 | 0.71566443 | 3.04708895 | 0.23486824 | 0.814311 | NA |
| ENSG0000004 | 0.09190453 | -0.5036858 | 3.04563564 | -0.1653795 | 0.86864525 | NA |
| ENSG0000004 | 0.06235927 | -0.1687738 | 3.04607858 | -0.0554069 | 0.9558143 | NA |
| ENSG0000004 | 0.20003502 | -0.4264876 | 3.04572908 | -0.1400281 | 0.88863781 | NA |
| ENSG0000004 | 0.70303984 | -0.0559761 | 1.20185939 | -0.0465746 | 0.96285226 | NA |
| ENSG0000004 | 0 | NA | NA | NA | NA | NA |
| ENSG0000004 | 0.11504242 | 0.2294839 | 2.60225812 | 0.08818645 | 0.92972849 | NA |
| ENSG0000005 | 0 | NA | NA | NA | NA | NA |
| ENSG0000005 | 0.84169487 | -0.5800956 | 1.14915243 | -0.504803 | 0.61369721 | NA |
| ENSG0000005 | 0.92456381 | 0.64439949 | 1.21664133 | 0.52965445 | 0.59635154 | NA |
| ENSG0000005 | 0 | NA | NA | NA | NA | NA |
| ENSG0000005 | 0.00492421 | 0.29059237 | 3.04687218 | 0.09537399 | 0.92401777 | NA |
| ENSG0000005 | 0.00984842 | 0.19866539 | 3.04669343 | 0.06520688 | 0.94800928 | NA |
| ENSG0000006 | 0.9067598 | -1.6778214 | 1.24735446 | -1.345104 | 0.17859167 | NA |
| ENSG0000006 | 0 | NA | NA | NA | NA | NA |
| ENSG0000006 | 0.78791549 | -0.9125678 | 1.08973538 | -0.8374215 | 0.4023557 | NA |
| ENSG0000006 | 0.23200192 | -0.0861573 | 2.25600609 | -0.0381902 | 0.96953606 | NA |
| ENSG0000006 | 1.20649266 | 0.51407564 | 0.87511258 | 0.58743943 | 0.55690862 | NA |
| ENSG0000006 | 0 | NA | NA | NA | NA | NA |
| ENSG0000006 | 0 | NA | NA | NA | NA | NA |
| ENSG0000006 | 1.14872869 | 1.18931502 | 1.01387038 | 1.17304445 | 0.24077798 | NA |
| ENSG0000006 | 0.17505744 | 0.63994735 | 2.37197556 | 0.26979509 | 0.7873179 | NA |
| ENSG0000006 | 0.39706894 | 0.92646495 | 1.60918068 | 0.57573705 | 0.56479293 | NA |
| ENSG0000006 | 0.4630558 | 0.79339371 | 1.72900915 | 0.4588719 | 0.64632616 | NA |
| ENSG0000007 | 0 | NA | NA | NA | NA | NA |
| ENSG0000007 | 0.06954201 | 0.61118818 | 3.04687218 | 0.20059528 | 0.84101505 | NA |
| ENSG0000007 | 0.14224343 | 0.67944133 | 2.6140449 | 0.25991953 | 0.79492584 | NA |
| ENSG0000007 | 0 | NA | NA | NA | NA | NA |
| ENSG0000007 | 0.05944154 | -0.0155188 | 3.04631688 | -0.0050943 | 0.99593536 | NA |
| ENSG0000007 | 0.65692839 | 0.00086167 | 1.23000623 | 0.00070054 | 0.99944105 | NA |
| ENSG0000007 | 0.09135804 | -0.4264876 | 3.04572908 | -0.1400281 | 0.8886378 | NA |
| ENSG0000007 | 1.05447091 | 0.15590754 | 0.91631663 | 0.17014592 | 0.86489538 | NA |
| ENSG0000007 | 0.73161064 | -1.8247895 | 1.04272038 | -1.7500277 | 0.08011354 | NA |
| ENSG0000007 | 0 | NA | NA | NA | NA | NA |
| ENSG0000007 | 0.19748872 | -1.1622095 | 2.29168451 | -0.507142 | 0.61205518 | NA |

| | | | | | | |
|---|---|---|---|---|---|---|
| ENSG0000007 | 0.00984842 | 0.19866539 | 3.04669343 | 0.06520688 | 0.94800928 | NA |
| ENSG0000007 | 0.87349749 | 0.70648838 | 1.05391795 | 0.67034477 | 0.50263804 | NA |
| ENSG0000007 | 0 | NA | NA | NA | NA | NA |
| ENSG0000007 | 0 | NA | NA | NA | NA | NA |
| ENSG0000007 | 0 | NA | NA | NA | NA | NA |
| ENSG0000007 | 0.14373984 | 1.03626011 | 3.04708895 | 0.340082 | 0.73379477 | NA |
| ENSG0000007 | 0.18579189 | -0.6474103 | 2.26585871 | -0.285724 | 0.77508949 | NA |
| ENSG0000008 | 0 | NA | NA | NA | NA | NA |
| ENSG0000008 | 1.24213674 | -0.4494385 | 0.87726878 | -0.5123156 | 0.60843014 | NA |
| ENSG0000008 | 0.59445952 | -2.4503039 | 1.33262799 | -1.8387006 | 0.06595924 | NA |
| ENSG0000008 | 1.10689447 | 0.97682039 | 0.892319 | 1.09469864 | 0.27364869 | NA |
| ENSG0000008 | 0 | NA | NA | NA | NA | NA |
| ENSG0000008 | 0 | NA | NA | NA | NA | NA |
| ENSG0000008 | 1.11957119 | 0.70617405 | 1.14480268 | 0.6168522 | 0.53733222 | NA |
| ENSG0000008 | 0.03446947 | -0.1484891 | 3.04610873 | -0.0487471 | 0.96112081 | NA |
| ENSG0000008 | 0.05743506 | -0.0960214 | 3.04618866 | -0.0315218 | 0.97485339 | NA |
| ENSG0000008 | 0 | NA | NA | NA | NA | NA |
| ENSG0000008 | 1.15717123 | 0.89217793 | 0.81034896 | 1.10097991 | 0.2709054 | NA |
| ENSG0000008 | 1.08944588 | -0.0101892 | 1.03204232 | -0.0098729 | 0.99212271 | NA |
| ENSG0000008 | 0.56296697 | 0.21940955 | 1.17172415 | 0.18725358 | 0.85146181 | NA |
| ENSG0000008 | 0.20855507 | 0.23150985 | 2.33051221 | 0.09933861 | 0.92086942 | NA |
| ENSG0000008 | 0.14905163 | 0.30591525 | 2.98698654 | 0.10241601 | 0.91842647 | NA |
| ENSG0000009 | 0 | NA | NA | NA | NA | NA |
| ENSG0000009 | 0.07157543 | 0.71566443 | 3.04708895 | 0.23486824 | 0.814311 | NA |
| ENSG0000009 | 0.13196966 | 0.7575698 | 3.04654238 | 0.24866544 | 0.80361958 | NA |
| ENSG0000009 | 0.66777728 | -1.9171165 | 1.09375496 | -1.7527843 | 0.07963904 | NA |
| ENSG0000009 | 0 | NA | NA | NA | NA | NA |
| ENSG0000009 | 0.55036894 | -0.9881466 | 1.25335177 | -0.7884032 | 0.43046087 | NA |
| ENSG0000009 | 0.1408654 | 0.1553788 | 2.44665881 | 0.06350653 | 0.94936316 | NA |
| ENSG0000009 | 0 | NA | NA | NA | NA | NA |
| ENSG0000009 | 0.13546114 | -0.4264876 | 3.04572908 | -0.1400281 | 0.8886378 | NA |
| ENSG00000010 | 0 | NA | NA | NA | NA | NA |
| ENSG00000010 | 0.49415596 | -0.7587834 | 1.21438879 | -0.6248274 | 0.53208433 | NA |
| ENSG00000010 | 0.36847292 | 0.12379919 | 1.37820705 | 0.08982626 | 0.92842528 | NA |
| ENSG00000010 | 0.20315681 | -0.899061 | 2.81693655 | -0.3191627 | 0.74960316 | NA |
| ENSG00000010 | 0.87030708 | -0.7347436 | 1.14552678 | -0.6414024 | 0.52126129 | NA |
| ENSG00000010 | 0.28343762 | 1.71005718 | 2.0958575 | 0.81592245 | 0.41454449 | NA |
| ENSG00000010 | 0 | NA | NA | NA | NA | NA |
| ENSG00000010 | 0 | NA | NA | NA | NA | NA |
| ENSG00000010 | 1.07458305 | -1.3093105 | 0.87688019 | -1.4931463 | 0.13539888 | NA |
| ENSG00000010 | 0 | NA | NA | NA | NA | NA |
| ENSG00000010 | 0.89581206 | -1.1795772 | 0.963529 | -1.224226 | 0.22086699 | NA |
| ENSG00000010 | 0.08163274 | 0.23083524 | 3.04619839 | 0.07577814 | 0.93959561 | NA |

| | | | | | | |
|---|---|---|---|---|---|---|
| ENSG0000010 | 0.39951104 | -0.6135294 | 2.07343761 | -0.2958996 | 0.76730674 | NA |
| ENSG0000010 | 0 | NA | NA | NA | NA | NA |
| ENSG0000010 | 0.59582056 | -0.2755498 | 1.35938458 | -0.2027019 | 0.83936804 | NA |
| ENSG0000010 | 0 | NA | NA | NA | NA | NA |
| ENSG0000010 | 0.05944154 | -0.0155188 | 3.04631688 | -0.0050943 | 0.99593536 | NA |
| ENSG0000010 | 0 | NA | NA | NA | NA | NA |
| ENSG0000010 | 0 | NA | NA | NA | NA | NA |
| ENSG0000010 | 0.19773843 | -0.3268145 | 2.26778722 | -0.1441116 | 0.88541233 | NA |
| ENSG0000010 | 0 | NA | NA | NA | NA | NA |
| ENSG0000010 | 0.73035918 | 0.00684113 | 1.14415535 | 0.0059792 | 0.99522932 | NA |
| ENSG0000010 | 0.12655369 | 0.84073792 | 2.57666783 | 0.32628882 | 0.74420584 | NA |
| ENSG0000010 | 0 | NA | NA | NA | NA | NA |
| ENSG0000010 | 0.62332043 | -0.7219941 | 1.37544173 | -0.524918 | 0.59964019 | NA |
| ENSG0000010 | 0 | NA | NA | NA | NA | NA |
| ENSG0000010 | 0 | NA | NA | NA | NA | NA |
| ENSG0000010 | 0.22205105 | 1.37590549 | 2.18932263 | 0.62846173 | 0.52970151 | NA |
| ENSG0000010 | 0 | NA | NA | NA | NA | NA |
| ENSG0000010 | 0.35457079 | -0.5664188 | 1.28835213 | -0.439646 | 0.66019354 | NA |
| ENSG0000010 | 0.00492421 | 0.29059237 | 3.04687218 | 0.09537399 | 0.92401777 | NA |
| ENSG0000010 | 0 | NA | NA | NA | NA | NA |
| ENSG0000010 | 0 | NA | NA | NA | NA | NA |
| ENSG0000010 | 0 | NA | NA | NA | NA | NA |
| ENSG0000010 | 0.05859851 | 0.71566443 | 3.04708895 | 0.23486824 | 0.814311 | NA |
| ENSG0000010 | 0.65058961 | 0.83875626 | 1.61161511 | 0.52044452 | 0.60275379 | NA |
| ENSG0000010 | 0 | NA | NA | NA | NA | NA |
| ENSG0000010 | 0.29791677 | 1.77429898 | 2.61997911 | 0.67721875 | 0.49826717 | NA |
| ENSG0000010 | 0.2979179 | -1.539866 | 2.20611591 | -0.6979987 | 0.48517801 | NA |
| ENSG0000010 | 0.09135804 | -0.4264876 | 3.04572908 | -0.1400281 | 0.8886378 | NA |
| ENSG0000010 | 0.00492421 | 0.29059237 | 3.04687218 | 0.09537399 | 0.92401777 | NA |
| ENSG0000010 | 0 | NA | NA | NA | NA | NA |
| ENSG0000010 | 0 | NA | NA | NA | NA | NA |
| ENSG0000010 | 0 | NA | NA | NA | NA | NA |
| ENSG0000011 | 0 | NA | NA | NA | NA | NA |
| ENSG0000011 | 0 | NA | NA | NA | NA | NA |
| ENSG0000011 | 0.9427043 | -3.2412766 | 1.16228963 | -2.7886996 | 0.00529201 | NA |
| ENSG0000011 | 0.72351285 | -0.8920947 | 1.08657496 | -0.8210153 | 0.41163754 | NA |
| ENSG0000011 | 0.40721442 | 0.0973184 | 1.41225279 | 0.06891004 | 0.94506123 | NA |
| ENSG0000011 | 0.25428132 | -1.2524969 | 2.1273312 | -0.5887644 | 0.55601934 | NA |
| ENSG0000011 | 0.2582469 | 0.72747106 | 1.83997789 | 0.39536946 | 0.69257025 | NA |
| ENSG0000011 | 0 | NA | NA | NA | NA | NA |
| ENSG0000011 | 0 | NA | NA | NA | NA | NA |
| ENSG0000011 | 0.70877369 | -0.3186418 | 1.09891507 | -0.2899603 | 0.77184657 | NA |
| ENSG0000011 | 1.03612497 | 0.5149965 | 0.81410556 | 0.6325918 | 0.52700024 | NA |

| | | | | | | |
|---|---|---|---|---|---|---|
| ENSG000001 | 0.99111571 | 1.76216487 | 0.97477107 | 1.80777305 | 0.07064183 | NA |
| ENSG000001 | 0 | NA | NA | NA | NA | NA |
| ENSG000001 | 0.8422975 | -1.1443786 | 1.20253421 | -0.9516391 | 0.34128004 | NA |
| ENSG000001 | 0.08961008 | 0.71566443 | 3.04708895 | 0.23486824 | 0.814311 | NA |
| ENSG000001 | 0.49699554 | 0.22756249 | 1.60177566 | 0.14206889 | 0.88702559 | NA |
| ENSG000001 | 0.52796303 | 0.06073778 | 1.30656617 | 0.04648657 | 0.96292244 | NA |
| ENSG000001 | 0 | NA | NA | NA | NA | NA |
| ENSG000001 | 0.25200605 | -1.4077338 | 1.84552204 | -0.7627835 | 0.44559252 | NA |
| ENSG000001 | 0.02462105 | -0.0266108 | 3.0462988 | -0.0087355 | 0.9930302 | NA |
| ENSG000001 | 0 | NA | NA | NA | NA | NA |
| ENSG000001 | 0.47043286 | 1.61013942 | 1.33805806 | 1.20334048 | 0.22884459 | NA |
| ENSG000001 | 0.15980498 | -0.2102568 | 2.01387021 | -0.1044043 | 0.91684848 | NA |
| ENSG000001 | 0.05944154 | -0.0155188 | 3.04631688 | -0.0050943 | 0.99593536 | NA |
| ENSG000001 | 1.06937801 | -0.2143609 | 0.88694837 | -0.2416836 | 0.80902531 | NA |
| ENSG000001 | 0.08698032 | -0.4565907 | 3.04569207 | -0.1499136 | 0.88083278 | NA |
| ENSG000001 | 1.03371679 | -1.18853 | 0.95009849 | -1.2509546 | 0.21095105 | NA |
| ENSG000001 | 0.10135333 | 1.03626011 | 3.04708895 | 0.340082 | 0.73379477 | NA |
| ENSG000001 | 0 | NA | NA | NA | NA | NA |
| ENSG000001 | 0.05251085 | -0.0155188 | 3.04631688 | -0.0050943 | 0.99593536 | NA |
| ENSG000001 | 0.56819504 | -0.9786571 | 1.12802001 | -0.8675884 | 0.3856197 | NA |
| ENSG000001 | 0 | NA | NA | NA | NA | NA |
| ENSG000001 | 0 | NA | NA | NA | NA | NA |
| ENSG000001 | 0.4622672 | 0.17075411 | 1.40941937 | 0.1211521 | 0.90357057 | NA |
| ENSG000001 | 1.21636776 | 0.3561292 | 0.82326071 | 0.43258374 | 0.66531721 | NA |
| ENSG000001 | 0.63754853 | -0.1391286 | 1.19212406 | -0.1167065 | 0.90709267 | NA |
| ENSG000001 | 0 | NA | NA | NA | NA | NA |
| ENSG000001 | 0.92600175 | -0.5144731 | 1.10219198 | -0.4667727 | 0.64066254 | NA |
| ENSG000001 | 0 | NA | NA | NA | NA | NA |
| ENSG000001 | 0.09453429 | 0.61118818 | 3.04687218 | 0.20059528 | 0.84101505 | NA |
| ENSG000001 | 0.16846445 | 0.15642373 | 2.33417949 | 0.06701444 | 0.94657021 | NA |
| ENSG000001 | 0.42298383 | -1.5018996 | 1.72794227 | -0.8691839 | 0.38474654 | NA |
| ENSG000001 | 0.27093315 | 1.7160506 | 2.11861133 | 0.8099884 | 0.41794684 | NA |
| ENSG000001 | 0.45326568 | -1.1465334 | 1.43494294 | -0.7990097 | 0.42428476 | NA |
| ENSG000001 | 0.00492421 | 0.29059237 | 3.04687218 | 0.09537399 | 0.92401777 | NA |
| ENSG000001 | 0.53993638 | 0.17677472 | 1.25867228 | 0.14044539 | 0.8883081 | NA |
| ENSG000001 | 0 | NA | NA | NA | NA | NA |
| ENSG000001 | 0.29631728 | -1.1993564 | 1.85754456 | -0.6456676 | 0.51849463 | NA |
| ENSG000001 | 0.8174413 | -0.2278793 | 1.2142968 | -0.1876636 | 0.85114037 | NA |
| ENSG000001 | 0 | NA | NA | NA | NA | NA |
| ENSG000001 | 0 | NA | NA | NA | NA | NA |
| ENSG000001 | 0 | NA | NA | NA | NA | NA |
| ENSG000001 | 0 | NA | NA | NA | NA | NA |
| ENSG000001 | 0.13561954 | 0.23120352 | 2.49137491 | 0.09280158 | 0.9260612 | NA |

| | | | | | | |
|---|---|---|---|---|---|---|
| ENSG000001: | 0.38400678 | -0.5333548 | 1.30836117 | -0.407651 | 0.6835299 | NA |
| ENSG000001: | 0 | NA | NA | NA | NA | NA |
| ENSG000001: | 0 | NA | NA | NA | NA | NA |
| ENSG000001: | 1.18307637 | -0.7665174 | 0.93880579 | -0.8164813 | 0.41422491 | NA |
| ENSG000001: | 0 | NA | NA | NA | NA | NA |
| ENSG000001: | 0 | NA | NA | NA | NA | NA |
| ENSG000001: | 0 | NA | NA | NA | NA | NA |
| ENSG000001: | 0.09135804 | -0.4264876 | 3.04572908 | -0.1400281 | 0.8886378 | NA |
| ENSG000001: | 0.00984842 | 0.19866539 | 3.04669343 | 0.06520688 | 0.94800928 | NA |
| ENSG000001: | 0.17991226 | 1.3600127 | 2.82364967 | 0.48165065 | 0.63005414 | NA |
| ENSG000001: | 0.54381767 | -0.7366773 | 1.17099636 | -0.629103 | 0.52928161 | NA |
| ENSG000001: | 0.51664845 | 0.76282209 | 1.27808462 | 0.59684788 | 0.55060895 | NA |
| ENSG000001: | 0.26137856 | 0.72807253 | 1.59489036 | 0.45650319 | 0.64802818 | NA |
| ENSG000001: | 0.47880784 | 1.02793504 | 1.24807267 | 0.82361793 | 0.41015668 | NA |
| ENSG000001: | 0 | NA | NA | NA | NA | NA |
| ENSG000001: | 0.2757571 | 1.07076114 | 1.89485761 | 0.56508792 | 0.57201397 | NA |
| ENSG000001: | 0.36892165 | 0.04967017 | 1.40075989 | 0.03545945 | 0.97171338 | NA |
| ENSG000001: | 0 | NA | NA | NA | NA | NA |
| ENSG000001: | 0 | NA | NA | NA | NA | NA |
| ENSG000001: | 0.59945481 | -1.5639008 | 1.09995355 | -1.421788 | 0.15508779 | NA |
| ENSG000001: | 0 | NA | NA | NA | NA | NA |
| ENSG000001: | 0.24015115 | -1.3541443 | 1.9060789 | -0.7104345 | 0.47743472 | NA |
| ENSG000001: | 0 | NA | NA | NA | NA | NA |
| ENSG000001: | 0.00492421 | 0.29059237 | 3.04687218 | 0.09537399 | 0.92401777 | NA |
| ENSG000001: | 0.36082489 | 0.50587997 | 1.40921845 | 0.3589791 | 0.71961072 | NA |
| ENSG000001: | 1.09190572 | -0.2384019 | 1.06348342 | -0.2241707 | 0.82262444 | NA |
| ENSG000001: | 0.37570872 | -1.348725 | 1.32916456 | -1.0147164 | 0.31024105 | NA |
| ENSG000001: | 0.00492421 | 0.29059237 | 3.04687218 | 0.09537399 | 0.92401777 | NA |
| ENSG000001: | 0 | NA | NA | NA | NA | NA |
| ENSG000001: | 0 | NA | NA | NA | NA | NA |
| ENSG000001: | 0.10120647 | -0.5421949 | 2.67950986 | -0.2023485 | 0.83964425 | NA |
| ENSG000001: | 0.14154881 | 0.3050978 | 3.04478504 | 0.1002034 | 0.92018285 | NA |
| ENSG000001: | 0.26153034 | 0.98909791 | 1.97863323 | 0.49988947 | 0.61715291 | NA |
| ENSG000001: | 0.26809263 | 0.89180705 | 1.83730604 | 0.4853884 | 0.62740086 | NA |
| ENSG000001: | 0.11011821 | 0.30507684 | 3.04631687 | 0.10014613 | 0.92022831 | NA |
| ENSG000001: | 0.47024912 | 0.86040214 | 1.33289532 | 0.64551366 | 0.51859437 | NA |
| ENSG000001: | 0.33645122 | 0.86298616 | 1.84979248 | 0.46653134 | 0.64083522 | NA |
| ENSG000001: | 1.07839475 | -1.0331283 | 0.9104773 | -1.1347107 | 0.25649656 | NA |
| ENSG000001: | 0 | NA | NA | NA | NA | NA |
| ENSG000001: | 0 | NA | NA | NA | NA | NA |
| ENSG000001: | 0.0829503 | -0.0155188 | 3.04631688 | -0.0050943 | 0.99593536 | NA |
| ENSG000001: | 1.12990263 | 0.46100529 | 0.83125086 | 0.55459225 | 0.57917359 | NA |
| ENSG000001: | 0.9858859 | 0.00260446 | 1.35308833 | 0.00192482 | 0.99846421 | NA |

| | | | | | | |
|---|---|---|---|---|---|---|
| ENSG0000001: | 0 | NA | NA | NA | NA | NA |
| ENSG0000001: | 0 | NA | NA | NA | NA | NA |
| ENSG0000001: | 0 | NA | NA | NA | NA | NA |
| ENSG0000001: | 1.2323897 | 0.08080329 | 1.15402485 | 0.07001867 | 0.9441788 | NA |
| ENSG0000001: | 0 | NA | NA | NA | NA | NA |
| ENSG0000001: | 0.00984842 | 0.19866539 | 3.04669343 | 0.06520688 | 0.94800928 | NA |
| ENSG0000001: | 1.24614742 | 0.61419934 | 0.81054946 | 0.75775677 | 0.44859661 | NA |
| ENSG0000001: | 0.05067667 | 0.71566443 | 3.04708895 | 0.23486824 | 0.814311 | NA |
| ENSG0000001: | 0 | NA | NA | NA | NA | NA |
| ENSG0000001: | 0.3693425 | -1.482006 | 2.11352357 | -0.7012016 | 0.48317724 | NA |
| ENSG0000001: | 0.38222493 | -0.633127 | 1.4972191 | -0.4228686 | 0.6723911 | NA |
| ENSG0000001: | 0.27698269 | 0.37096455 | 1.86257229 | 0.19916786 | 0.84213144 | NA |
| ENSG0000001: | 0.33630265 | 0.5873165 | 1.75424961 | 0.33479643 | 0.73777866 | NA |
| ENSG0000001: | 0.01969684 | 0.0417549 | 3.04641238 | 0.01370625 | 0.98906434 | NA |
| ENSG0000001: | 1.18304896 | -1.7564125 | 1.01577852 | -1.7291294 | 0.08378595 | NA |
| ENSG0000001: | 0.77469409 | 0.08210241 | 1.12465068 | 0.07300258 | 0.94180406 | NA |
| ENSG0000001: | 0.35368033 | -0.1583003 | 1.45827902 | -0.1085528 | 0.91355719 | NA |
| ENSG0000001: | 0.06461779 | 0.71566443 | 3.04708895 | 0.23486824 | 0.814311 | NA |
| ENSG0000001: | 0.81604307 | 2.15658535 | 1.33750459 | 1.61239473 | 0.10687607 | NA |
| ENSG0000001: | 0 | NA | NA | NA | NA | NA |
| ENSG0000001: | 0 | NA | NA | NA | NA | NA |
| ENSG0000001: | 0 | NA | NA | NA | NA | NA |
| ENSG0000001: | 0.54322925 | -0.6575334 | 1.24214625 | -0.5293526 | 0.59656087 | NA |
| ENSG0000001: | 0.93195553 | 0.82088512 | 0.85823796 | 0.9564773 | 0.33883115 | NA |
| ENSG0000001: | 0 | NA | NA | NA | NA | NA |
| ENSG0000001: | 0.00492421 | 0.29059237 | 3.04687218 | 0.09537399 | 0.92401777 | NA |
| ENSG0000001: | 0.5468226 | 0.16068917 | 1.36425809 | 0.11778502 | 0.906238 | NA |
| ENSG0000001: | 0.14137525 | -0.03788 | 2.45493009 | -0.0154302 | 0.98768899 | NA |
| ENSG0000001: | 0.00492421 | 0.29059237 | 3.04687218 | 0.09537399 | 0.92401777 | NA |
| ENSG0000001: | 0.9614855 | 0.77036823 | 1.06605093 | 0.72263736 | 0.46990271 | NA |
| ENSG0000001: | 0.06461779 | 0.71566443 | 3.04708895 | 0.23486824 | 0.814311 | NA |
| ENSG0000001: | 0.31014925 | 1.36741038 | 2.45923818 | 0.55603007 | 0.5781903 | NA |
| ENSG0000001: | 0.27439859 | 1.06871634 | 1.61036896 | 0.66364689 | 0.50691634 | NA |
| ENSG0000001: | 0.60171431 | 0.02679816 | 1.13934345 | 0.0235207 | 0.98123493 | NA |
| ENSG0000001: | 0.14756809 | 0.30559924 | 3.00889432 | 0.1015653 | 0.91910173 | NA |
| ENSG0000001: | 0 | NA | NA | NA | NA | NA |
| ENSG0000001: | 0.81018679 | -1.957489 | 1.07775033 | -1.8162732 | 0.06932848 | NA |
| ENSG0000001: | 0.05859851 | 0.71566443 | 3.04708895 | 0.23486824 | 0.814311 | NA |
| ENSG0000001: | 1.03618057 | -0.5648456 | 0.84211639 | -0.6707453 | 0.50238283 | NA |
| ENSG0000001: | 0.1570292 | 0.23271476 | 2.40632706 | 0.09670953 | 0.92295707 | NA |
| ENSG0000001: | 0 | NA | NA | NA | NA | NA |
| ENSG0000001: | 0 | NA | NA | NA | NA | NA |
| ENSG0000001 | 0.09453429 | 0.61118818 | 3.04687218 | 0.20059528 | 0.84101505 | NA |

| | | | | | | |
|---|---|---|---|---|---|---|
| ENSG000001[illegible] | 0.27289168 | 1.23936868 | 1.880867 | 0.65893478 | 0.50993765 | NA |
| ENSG000001[illegible] | 0 | NA | NA | NA | NA | NA |
| ENSG000001[illegible] | 0.10276414 | 1.03626011 | 3.04708895 | 0.340082 | 0.73379477 | NA |
| ENSG000001[illegible] | 0 | NA | NA | NA | NA | NA |
| ENSG000001[illegible] | 1.16886012 | 0.03086055 | 0.83054212 | 0.03715712 | 0.97035973 | NA |
| ENSG000001[illegible] | 1.19571176 | -1.1529526 | 0.84647536 | -1.3620628 | 0.17317806 | NA |
| ENSG000001[illegible] | 0 | NA | NA | NA | NA | NA |
| ENSG000001[illegible] | 0.20823536 | 0.10486235 | 1.92834815 | 0.05437936 | 0.95663292 | NA |
| ENSG000001[illegible] | 0.50485869 | 1.25976574 | 1.35938219 | 0.92671932 | 0.35407228 | NA |
| ENSG000001[illegible] | 0.51030604 | -0.2379368 | 1.54805164 | -0.1537008 | 0.87784566 | NA |
| ENSG000001[illegible] | 0 | NA | NA | NA | NA | NA |
| ENSG000001[illegible] | 0 | NA | NA | NA | NA | NA |
| ENSG000001[illegible] | 0 | NA | NA | NA | NA | NA |
| ENSG000001[illegible] | 0.32258161 | 0.23942608 | 2.14700946 | 0.11151608 | 0.91120711 | NA |
| ENSG000001[illegible] | 0 | NA | NA | NA | NA | NA |
| ENSG000001[illegible] | 0 | NA | NA | NA | NA | NA |
| ENSG000001[illegible] | 0.05251085 | -0.0155188 | 3.04631688 | -0.0050943 | 0.99593536 | NA |
| ENSG000001[illegible] | 0.42308145 | 1.36007352 | 1.50931442 | 0.90112007 | 0.36752448 | NA |
| ENSG000001[illegible] | 0 | NA | NA | NA | NA | NA |
| ENSG000001[illegible] | 0.90311498 | -0.366242 | 1.04643271 | -0.349991 | 0.72634545 | NA |
| ENSG000001[illegible] | 0 | NA | NA | NA | NA | NA |
| ENSG000001[illegible] | 0 | NA | NA | NA | NA | NA |
| ENSG000001[illegible] | 0.05067667 | 0.71566443 | 3.04708895 | 0.23486824 | 0.814311 | NA |
| ENSG000001[illegible] | 0.49403678 | 0.16642624 | 1.39916511 | 0.11894682 | 0.90531749 | NA |
| ENSG000001[illegible] | 0.10120647 | -0.5421949 | 2.67950986 | -0.2023485 | 0.83964425 | NA |
| ENSG000001[illegible] | 0 | NA | NA | NA | NA | NA |
| ENSG000001[illegible] | 0.96378477 | 0.88251293 | 1.01147614 | 0.8725 | 0.38293567 | NA |
| ENSG000001[illegible] | 0.33180118 | -1.6188944 | 3.02784505 | -0.5346688 | 0.59287887 | NA |
| ENSG000001[illegible] | 0 | NA | NA | NA | NA | NA |
| ENSG000001[illegible] | 0 | NA | NA | NA | NA | NA |
| ENSG000001[illegible] | 0 | NA | NA | NA | NA | NA |
| ENSG000001[illegible] | 0 | NA | NA | NA | NA | NA |
| ENSG000001[illegible] | 1.07783824 | 0.5839253 | 0.94684403 | 0.61670696 | 0.53742803 | NA |
| ENSG000001[illegible] | 0 | NA | NA | NA | NA | NA |
| ENSG000001[illegible] | 0.34325914 | -0.1266631 | 1.49770753 | -0.0845713 | 0.93260219 | NA |
| ENSG000001[illegible] | 0 | NA | NA | NA | NA | NA |
| ENSG000001[illegible] | 0.54578954 | -0.214454 | 1.27828078 | -0.1677675 | 0.86676619 | NA |
| ENSG000001[illegible] | 0 | NA | NA | NA | NA | NA |
| ENSG000001[illegible] | 0 | NA | NA | NA | NA | NA |
| ENSG000001[illegible] | 0 | NA | NA | NA | NA | NA |
| ENSG000001[illegible] | 0.32721134 | 0.18676088 | 1.81190536 | 0.1030743 | 0.917904 | NA |
| ENSG000001[illegible] | 0 | NA | NA | NA | NA | NA |
| ENSG000001[illegible] | 0.23522694 | -1.3282229 | 2.28705536 | -0.5807568 | 0.56140438 | NA |

| | | | | | | |
|---|---|---|---|---|---|---|
| ENSG0000015 | 1.14539016 | -1.5056738 | 0.90491447 | -1.6638852 | 0.09613537 | NA |
| ENSG0000015 | 0 | NA | NA | NA | NA | NA |
| ENSG0000015 | 0 | NA | NA | NA | NA | NA |
| ENSG0000015 | 0 | NA | NA | NA | NA | NA |
| ENSG0000015 | 0.56750183 | -0.2658728 | 1.11620888 | -0.2381927 | 0.81173163 | NA |
| ENSG0000015 | 0.59694379 | -1.6363886 | 1.09247787 | -1.4978689 | 0.13416731 | NA |
| ENSG0000015 | 0 | NA | NA | NA | NA | NA |
| ENSG0000015 | 0.06607753 | -0.0155188 | 3.04631688 | -0.0050943 | 0.99593536 | NA |
| ENSG0000015 | 0.00984842 | 0.19866539 | 3.04669343 | 0.06520688 | 0.94800928 | NA |
| ENSG0000015 | 0.06686011 | 0.43697383 | 3.04654239 | 0.14343271 | 0.88594845 | NA |
| ENSG0000015 | 0.28851196 | 0.72816839 | 1.56666736 | 0.46478813 | 0.64208319 | NA |
| ENSG0000015 | 0.65168423 | -0.5818926 | 1.35828697 | -0.4284018 | 0.66835862 | NA |
| ENSG0000015 | 0.32525812 | -0.6011169 | 2.20933245 | -0.2720808 | 0.7855599 | NA |
| ENSG0000015 | 0.55233523 | -0.070031 | 1.34573982 | -0.052039 | 0.95849761 | NA |
| ENSG0000015 | 0.20667768 | -1.1993893 | 1.87833338 | -0.6385391 | 0.52312282 | NA |
| ENSG0000015 | 0.1939928 | 0.75698296 | 2.85338935 | 0.26529256 | 0.7907841 | NA |
| ENSG0000015 | 0 | NA | NA | NA | NA | NA |
| ENSG0000015 | 0 | NA | NA | NA | NA | NA |
| ENSG0000015 | 0.07679413 | 0.61118818 | 3.04687218 | 0.20059528 | 0.84101505 | NA |
| ENSG0000015 | 0 | NA | NA | NA | NA | NA |
| ENSG0000015 | 1.02778338 | -2.1629091 | 1.0079853 | -2.1457745 | 0.03189098 | NA |
| ENSG0000015 | 0.58415996 | -1.8081226 | 1.40958193 | -1.2827368 | 0.1995843 | NA |
| ENSG0000015 | 0 | NA | NA | NA | NA | NA |
| ENSG0000015 | 0.15485162 | 1.35817061 | 2.94711035 | 0.46084824 | 0.64490749 | NA |
| ENSG0000015 | 0.14951189 | -0.1942751 | 2.44650616 | -0.0794092 | 0.93670714 | NA |
| ENSG0000015 | 0 | NA | NA | NA | NA | NA |
| ENSG0000015 | 0.55976769 | -0.9502489 | 1.11242616 | -0.8542131 | 0.39298695 | NA |
| ENSG0000015 | 0 | NA | NA | NA | NA | NA |
| ENSG0000015 | 0.37112966 | -0.8291341 | 1.36559587 | -0.6071592 | 0.54374524 | NA |
| ENSG0000015 | 0.1767129 | 0.27467078 | 2.50455297 | 0.10966859 | 0.91267222 | NA |
| ENSG0000015 | 0 | NA | NA | NA | NA | NA |
| ENSG0000016 | 0.27575285 | -1.3630472 | 1.59061536 | -0.8569307 | 0.39148318 | NA |
| ENSG0000016 | 0 | NA | NA | NA | NA | NA |
| ENSG0000016 | 0.36130513 | 0.4733714 | 1.29688855 | 0.36500546 | 0.71510733 | NA |
| ENSG0000016 | 0.7374338 | 0.10934703 | 1.11188876 | 0.09834349 | 0.92165954 | NA |
| ENSG0000016 | 1.09645608 | -0.5876056 | 1.31064391 | -0.4483335 | 0.65391252 | NA |
| ENSG0000016 | 1.0964988 | -0.2920623 | 0.85378314 | -0.3420802 | 0.73229054 | NA |
| ENSG0000016 | 0 | NA | NA | NA | NA | NA |
| ENSG0000016 | 0.22400103 | 0.40639847 | 2.20049736 | 0.18468483 | 0.85347625 | NA |
| ENSG0000016 | 0 | NA | NA | NA | NA | NA |
| ENSG0000016 | 0.63811834 | 1.55293431 | 1.26812743 | 1.22458854 | 0.22073029 | NA |
| ENSG0000016 | 0 | NA | NA | NA | NA | NA |
| ENSG0000016 | 0 | NA | NA | NA | NA | NA |

| | | | | | | |
|---|---|---|---|---|---|---|
| ENSG000001( | 0.05208748 | 0.71566443 | 3.04708895 | 0.23486824 | 0.814311 | NA |
| ENSG000001( | 0 | NA | NA | NA | NA | NA |
| ENSG000001( | 0 | NA | NA | NA | NA | NA |
| ENSG000001( | 0.06461779 | 0.71566443 | 3.04708895 | 0.23486824 | 0.814311 | NA |
| ENSG000001( | 0.36945613 | -1.2943534 | 1.51888375 | -0.8521741 | 0.39411747 | NA |
| ENSG000001( | 0 | NA | NA | NA | NA | NA |
| ENSG000001( | 0.05944154 | -0.0155188 | 3.04631688 | -0.0050943 | 0.99593536 | NA |
| ENSG000001( | 0.32560475 | -0.8569865 | 1.41142164 | -0.6071796 | 0.5437317 | NA |
| ENSG000001( | 0.76851901 | 0.16494026 | 1.35673391 | 0.12157156 | 0.90323834 | NA |
| ENSG000001( | 0.00984842 | 0.19866539 | 3.04669343 | 0.06520688 | 0.94800928 | NA |
| ENSG000001( | 0 | NA | NA | NA | NA | NA |
| ENSG000001( | 0.62158592 | -1.1837135 | 1.07223631 | -1.103967 | 0.26960746 | NA |
| ENSG000001( | 0.05251085 | -0.0155188 | 3.04631688 | -0.0050943 | 0.99593536 | NA |
| ENSG000001( | 0.6126728 | -0.3848043 | 1.18152379 | -0.3256848 | 0.74466286 | NA |
| ENSG000001( | 0.1613944 | 0.67859138 | 2.5366322 | 0.26751666 | 0.78907139 | NA |
| ENSG000001( | 1.19067423 | -1.2572641 | 0.82986919 | -1.5150148 | 0.12976864 | NA |
| ENSG000001( | 0 | NA | NA | NA | NA | NA |
| ENSG000001( | 0.10417495 | 1.03626011 | 3.04708895 | 0.340082 | 0.73379477 | NA |
| ENSG000001( | 0.45814646 | 1.86883772 | 1.5180673 | 1.23106381 | 0.218299 | NA |
| ENSG000001( | 0 | NA | NA | NA | NA | NA |
| ENSG000001( | 0 | NA | NA | NA | NA | NA |
| ENSG000001( | 1.19459568 | 0.00202232 | 0.79400137 | 0.002547 | 0.99796779 | NA |
| ENSG000001( | 0.11928691 | 0.71566443 | 3.04708895 | 0.23486824 | 0.814311 | NA |
| ENSG000001( | 0 | NA | NA | NA | NA | NA |
| ENSG000001( | 0.32152411 | 0.63705732 | 2.46808752 | 0.2581178 | 0.796316 | NA |
| ENSG000001( | 0 | NA | NA | NA | NA | NA |
| ENSG000001( | 0.32259735 | -0.7976754 | 1.65175983 | -0.4829245 | 0.62914932 | NA |
| ENSG000001( | 0.82635697 | -0.4642628 | 1.04459123 | -0.4444445 | 0.65672127 | NA |
| ENSG000001( | 0 | NA | NA | NA | NA | NA |
| ENSG000001( | 0 | NA | NA | NA | NA | NA |
| ENSG000001( | 0 | NA | NA | NA | NA | NA |
| ENSG000001( | 0.07100174 | -0.0960214 | 3.04618866 | -0.0315218 | 0.97485339 | NA |
| ENSG000001( | 0.32966559 | 1.05535296 | 1.79883967 | 0.5866854 | 0.55741502 | NA |
| ENSG000001( | 0 | NA | NA | NA | NA | NA |
| ENSG000001( | 0.00492421 | 0.29059237 | 3.04687218 | 0.09537399 | 0.92401777 | NA |
| ENSG000001( | 0.05701169 | 0.61118818 | 3.04687218 | 0.20059528 | 0.84101505 | NA |
| ENSG000001( | 0.20855507 | 0.23150985 | 2.33051221 | 0.09933861 | 0.92086942 | NA |
| ENSG000001( | 0.16148 | 1.03860824 | 2.87568052 | 0.36116955 | 0.7179727 | NA |
| ENSG000001( | 0.16582426 | -0.209743 | 1.99595879 | -0.1050838 | 0.9163093 | NA |
| ENSG000001( | 0 | NA | NA | NA | NA | NA |
| ENSG000001( | 0.13046843 | 1.03626011 | 3.04708895 | 0.340082 | 0.73379477 | NA |
| ENSG000001( | 0 | NA | NA | NA | NA | NA |
| ENSG000001( | 0.01477263 | 0.11637786 | 3.04654239 | 0.03819998 | 0.96952824 | NA |

| | | | | | | |
|---|---|---|---|---|---|---|
| ENSG0000010 | 0 | NA | NA | NA | NA | NA |
| ENSG0000010 | 0 | NA | NA | NA | NA | NA |
| ENSG0000010 | 0 | NA | NA | NA | NA | NA |
| ENSG0000010 | 0.64956618 | -0.1815445 | 1.25618505 | -0.1445205 | 0.88508945 | NA |
| ENSG0000010 | 0.26593565 | -0.9062066 | 1.81804377 | -0.4984515 | 0.61816585 | NA |
| ENSG0000010 | 0 | NA | NA | NA | NA | NA |
| ENSG0000010 | 0.10120647 | -0.5421949 | 2.67950986 | -0.2023485 | 0.83964425 | NA |
| ENSG0000010 | 0.26731577 | -0.4265629 | 2.10008601 | -0.2031169 | 0.83904367 | NA |
| ENSG0000010 | 0.44866945 | -0.4778921 | 1.55202302 | -0.3079156 | 0.75814655 | NA |
| ENSG0000010 | 0.17922017 | 1.03626011 | 3.04708895 | 0.340082 | 0.73379477 | NA |
| ENSG0000001 | 0.58475736 | -0.7131268 | 1.18786726 | -0.6003422 | 0.54827823 | NA |
| ENSG0000001 | 0 | NA | NA | NA | NA | NA |
| ENSG0000001 | 0.48996987 | -1.2242269 | 1.49129229 | -0.8209168 | 0.41169364 | NA |
| ENSG0000001 | 0.31924106 | 0.8728566 | 1.89572916 | 0.46043318 | 0.64520532 | NA |
| ENSG0000001 | 0 | NA | NA | NA | NA | NA |
| ENSG0000001 | 0.39906853 | -0.9778373 | 1.23653172 | -0.7907903 | 0.42906638 | NA |
| ENSG0000001 | 0 | NA | NA | NA | NA | NA |
| ENSG0000001 | 0.42128991 | -0.4308327 | 2.07906333 | -0.2072244 | 0.8358346 | NA |
| ENSG0000001 | 0.46111361 | -0.5371436 | 1.35006521 | -0.3978649 | 0.69072974 | NA |
| ENSG0000001 | 0.09135804 | -0.4264876 | 3.04572908 | -0.1400281 | 0.8886378 | NA |
| ENSG0000001 | 1.11428553 | 2.24828972 | 0.91127336 | 2.46719571 | 0.01361759 | NA |
| ENSG0000001 | 0 | NA | NA | NA | NA | NA |
| ENSG0000001 | 0.30505672 | -1.2772294 | 2.60677678 | -0.489965 | 0.62415865 | NA |
| ENSG0000001 | 0 | NA | NA | NA | NA | NA |
| ENSG0000001 | 0.01477263 | 0.11637786 | 3.04654239 | 0.03819998 | 0.96952824 | NA |
| ENSG0000001 | 0.06352272 | 0.61118818 | 3.04687218 | 0.20059528 | 0.84101505 | NA |
| ENSG0000001 | 0 | NA | NA | NA | NA | NA |
| ENSG0000001 | 0.37360913 | -1.8992669 | 1.74730605 | -1.0869686 | 0.27705067 | NA |
| ENSG0000001 | 0.43943093 | 1.24564258 | 1.23801475 | 1.00616134 | 0.31433797 | NA |
| ENSG0000001 | 0 | NA | NA | NA | NA | NA |
| ENSG0000001 | 0.74041086 | -0.3206827 | 1.04414876 | -0.3071236 | 0.7587493 | NA |
| ENSG0000001 | 0 | NA | NA | NA | NA | NA |
| ENSG0000001 | 0 | NA | NA | NA | NA | NA |
| ENSG0000001 | 0 | NA | NA | NA | NA | NA |
| ENSG0000001 | 0.19812913 | 1.1637379 | 1.97360168 | 0.58965186 | 0.55542408 | NA |
| ENSG0000001 | 0 | NA | NA | NA | NA | NA |
| ENSG0000001 | 0.18579189 | -0.6474103 | 2.26585871 | -0.285724 | 0.77508949 | NA |
| ENSG0000001 | 0.05944154 | -0.0155188 | 3.04631688 | -0.0050943 | 0.99593536 | NA |
| ENSG0000001 | 0 | NA | NA | NA | NA | NA |
| ENSG0000001 | 0.43266259 | 0.75829637 | 1.80011204 | 0.42124954 | 0.67357287 | NA |
| ENSG0000001 | 0.77781219 | 1.17046023 | 1.08785064 | 1.07593836 | 0.28195483 | NA |
| ENSG0000001 | 0.05067667 | 0.71566443 | 3.04708895 | 0.23486824 | 0.814311 | NA |
| ENSG0000001 | 1.15270196 | -1.0372336 | 0.92690614 | -1.1190276 | 0.26312835 | NA |

| | | | | | | |
|---|---|---|---|---|---|---|
| ENSG00000017 | 0 | NA | NA | NA | NA | NA |
| ENSG00000017 | 0.05067667 | 0.71566443 | 3.04708895 | 0.23486824 | 0.814311 | NA |
| ENSG00000017 | 0 | NA | NA | NA | NA | NA |
| ENSG00000017 | 1.0995353 | -1.0508073 | 0.84871554 | -1.2381148 | 0.21567348 | NA |
| ENSG00000017 | 0 | NA | NA | NA | NA | NA |
| ENSG00000017 | 0.06461779 | 0.71566443 | 3.04708895 | 0.23486824 | 0.814311 | NA |
| ENSG00000017 | 0.19052164 | -0.6952687 | 2.75333233 | -0.252519 | 0.80063994 | NA |
| ENSG00000017 | 0 | NA | NA | NA | NA | NA |
| ENSG00000017 | 0.05859851 | 0.71566443 | 3.04708895 | 0.23486824 | 0.814311 | NA |
| ENSG00000017 | 0.14639103 | 0.75567352 | 2.53677343 | 0.29788767 | 0.7657889 | NA |
| ENSG00000017 | 0.25683111 | -0.5121256 | 2.71163047 | -0.1888626 | 0.8502005 | NA |
| ENSG00000017 | 0 | NA | NA | NA | NA | NA |
| ENSG00000017 | 0 | NA | NA | NA | NA | NA |
| ENSG00000017 | 0 | NA | NA | NA | NA | NA |
| ENSG00000018 | 0 | NA | NA | NA | NA | NA |
| ENSG00000018 | 0.29139307 | -1.1759445 | 2.2070411 | -0.532815 | 0.59416167 | NA |
| ENSG00000018 | 0.71844709 | 1.84765405 | 1.32526666 | 1.3941753 | 0.16326467 | NA |
| ENSG00000018 | 0.16079488 | 0.62694332 | 2.95786617 | 0.21195798 | 0.83213982 | NA |
| ENSG00000018 | 0.08961008 | 0.71566443 | 3.04708895 | 0.23486824 | 0.814311 | NA |
| ENSG00000018 | 0.61709061 | 0.89867655 | 1.22935045 | 0.73101739 | 0.46476853 | NA |
| ENSG00000018 | 0.94531844 | 0.03852443 | 0.88985858 | 0.04329275 | 0.96546817 | NA |
| ENSG00000018 | 0.40011792 | 0.58537002 | 1.79330268 | 0.32642009 | 0.74410654 | NA |
| ENSG00000018 | 0 | NA | NA | NA | NA | NA |
| ENSG00000018 | 0.85688312 | 2.79746692 | 1.37050247 | 2.04119801 | 0.04123115 | NA |
| ENSG00000018 | 0 | NA | NA | NA | NA | NA |
| ENSG00000018 | 0.09135804 | -0.4264876 | 3.04572908 | -0.1400281 | 0.8886378 | NA |
| ENSG00000018 | 0 | NA | NA | NA | NA | NA |
| ENSG00000018 | 0.17341416 | -1.0491205 | 1.96063591 | -0.5350919 | 0.59258628 | NA |
| ENSG00000018 | 0.82311129 | 0.55528153 | 1.04666658 | 0.5305238 | 0.59574881 | NA |
| ENSG00000018 | 0.47900132 | -1.0344052 | 1.47615939 | -0.7007409 | 0.48346474 | NA |
| ENSG00000018 | 0.24492197 | 0.96482177 | 2.38306858 | 0.4048653 | 0.68557653 | NA |
| ENSG00000018 | 0.03446947 | -0.1484891 | 3.04610873 | -0.0487471 | 0.96112081 | NA |
| ENSG00000018 | 0 | NA | NA | NA | NA | NA |
| ENSG00000018 | 0 | NA | NA | NA | NA | NA |
| ENSG00000018 | 0.22088022 | 1.61786428 | 2.71897911 | 0.59502637 | 0.55182584 | NA |
| ENSG00000018 | 1.09654973 | -0.7404114 | 0.99333623 | -0.7453785 | 0.45604295 | NA |
| ENSG00000018 | 0.00984842 | 0.19866539 | 3.04669343 | 0.06520688 | 0.94800928 | NA |
| ENSG00000018 | 0.30057116 | 0.60389527 | 1.64205505 | 0.36776798 | 0.71304624 | NA |
| ENSG00000018 | 0.07100174 | -0.0960214 | 3.04618866 | -0.0315218 | 0.97485339 | NA |
| ENSG00000018 | 0 | NA | NA | NA | NA | NA |
| ENSG00000018 | 0.54410882 | 0.05273674 | 1.48440464 | 0.0355272 | 0.97165936 | NA |
| ENSG00000018 | 0 | NA | NA | NA | NA | NA |
| ENSG00000018 | 0.34216346 | 1.8064198 | 1.8348529 | 0.98450388 | 0.32486782 | NA |

| | | | | | | |
|---|---|---|---|---|---|---|
| ENSG0000018 | 0 | NA | NA | NA | NA | NA |
| ENSG0000018 | 0.68429271 | 0.21767934 | 1.26859287 | 0.17159118 | 0.86375894 | NA |
| ENSG0000018 | 0.00492421 | 0.29059237 | 3.04687218 | 0.09537399 | 0.92401777 | NA |
| ENSG0000018 | 0.16963949 | -0.1041831 | 2.93592601 | -0.0354856 | 0.97169252 | NA |
| ENSG0000018 | 0.29316878 | -1.3304056 | 2.54613959 | -0.5225187 | 0.60130922 | NA |
| ENSG0000018 | 0.62276317 | 1.23431237 | 1.11432378 | 1.10767839 | 0.26800073 | NA |
| ENSG0000018 | 0.59898494 | -0.367066 | 1.25436086 | -0.2926319 | 0.76980352 | NA |
| ENSG0000018 | 0.09135804 | -0.4264876 | 3.04572908 | -0.1400281 | 0.8886378 | NA |
| ENSG0000018 | 0 | NA | NA | NA | NA | NA |
| ENSG0000018 | 0.10417495 | 1.03626011 | 3.04708895 | 0.340082 | 0.73379477 | NA |
| ENSG0000018 | 0 | NA | NA | NA | NA | NA |
| ENSG0000018 | 0.93360932 | 0.48047972 | 1.22397288 | 0.39255749 | 0.69464634 | NA |
| ENSG0000018 | 0 | NA | NA | NA | NA | NA |
| ENSG0000018 | 1.17282356 | -0.3246394 | 1.25462902 | -0.2587533 | 0.79582561 | NA |
| ENSG0000018 | 0 | NA | NA | NA | NA | NA |
| ENSG0000018 | 0.82535073 | -0.5276146 | 0.97263142 | -0.542461 | 0.58750095 | NA |
| ENSG0000018 | 0.69148778 | 2.39430787 | 1.29445614 | 1.84966318 | 0.06436211 | NA |
| ENSG0000018 | 0 | NA | NA | NA | NA | NA |
| ENSG0000018 | 0.01477263 | 0.11637786 | 3.04654239 | 0.03819998 | 0.96952824 | NA |
| ENSG0000018 | 0.10276414 | 1.03626011 | 3.04708895 | 0.340082 | 0.73379477 | NA |
| ENSG0000018 | 0.15329394 | -0.2108209 | 2.03426 | -0.1036352 | 0.91745888 | NA |
| ENSG0000019 | 0.61614633 | -0.0066504 | 1.19865506 | -0.0055482 | 0.99557319 | NA |
| ENSG0000019 | 0.05701169 | 0.61118818 | 3.04687218 | 0.20059528 | 0.84101505 | NA |
| ENSG0000019 | 0 | NA | NA | NA | NA | NA |
| ENSG0000019 | 0.10001751 | -0.0155189 | 3.04631688 | -0.0050943 | 0.99593535 | NA |
| ENSG0000019 | 0.10547122 | -0.5036858 | 3.04563564 | -0.1653795 | 0.86864525 | NA |
| ENSG0000019 | 0 | NA | NA | NA | NA | NA |
| ENSG0000019 | 0 | NA | NA | NA | NA | NA |
| ENSG0000019 | 0.19714969 | 0.56637552 | 1.94203679 | 0.29163996 | 0.77056192 | NA |
| ENSG0000019 | 0.05859851 | 0.71566443 | 3.04708895 | 0.23486824 | 0.814311 | NA |
| ENSG0000019 | 0 | NA | NA | NA | NA | NA |
| ENSG0000019 | 0.22381948 | -0.4804046 | 1.97616341 | -0.2430996 | 0.80792821 | NA |
| ENSG0000019 | 0.43974255 | 2.39343901 | 1.96427727 | 1.21848328 | 0.22304038 | NA |
| ENSG0000019 | 0.63541884 | -0.4307329 | 1.13352975 | -0.3799925 | 0.70395095 | NA |
| ENSG0000019 | 0.64235911 | -2.6280125 | 1.59684864 | -1.6457493 | 0.09981539 | NA |
| ENSG0000019 | 0 | NA | NA | NA | NA | NA |
| ENSG0000019 | 0.18961549 | 0.6413497 | 2.3298559 | 0.27527441 | 0.78310542 | NA |
| ENSG0000019 | 0.11110936 | 0.30507684 | 3.04631687 | 0.10014613 | 0.92022831 | NA |
| ENSG0000019 | 0.14031686 | 0.84091495 | 2.5080774 | 0.3352827 | 0.73741185 | NA |
| ENSG0000019 | 0 | NA | NA | NA | NA | NA |
| ENSG0000019 | 0 | NA | NA | NA | NA | NA |
| ENSG0000019 | 0.36738201 | -1.8487639 | 1.79776134 | -1.02837 | 0.3037758 | NA |
| ENSG0000019 | 0 | NA | NA | NA | NA | NA |

| | | | | | | |
|---|---|---|---|---|---|---|
| ENSG0000020 | 0 | NA | NA | NA | NA | NA |
| ENSG0000020 | 0 | NA | NA | NA | NA | NA |
| ENSG0000020 | 0 | NA | NA | NA | NA | NA |
| ENSG0000020 | 0.53251179 | -1.3419452 | 1.12098403 | -1.1971136 | 0.23126227 | NA |
| ENSG0000020 | 0 | NA | NA | NA | NA | NA |
| ENSG0000020 | 0 | NA | NA | NA | NA | NA |
| ENSG0000020 | 0 | NA | NA | NA | NA | NA |
| ENSG0000020 | 0 | NA | NA | NA | NA | NA |
| ENSG0000020 | 0.19763559 | 0.17715574 | 1.98082048 | 0.08943554 | 0.92873578 | NA |
| ENSG0000020 | 0 | NA | NA | NA | NA | NA |
| ENSG0000020 | 0 | NA | NA | NA | NA | NA |
| ENSG0000021 | 0 | NA | NA | NA | NA | NA |
| ENSG0000021 | 0.10318752 | 0.30507684 | 3.04631687 | 0.10014613 | 0.92022831 | NA |
| ENSG0000021 | 1.24936948 | -0.3430482 | 0.81499689 | -0.4209197 | 0.67381374 | NA |
| ENSG0000021 | 0.34300146 | 1.98722742 | 1.76641755 | 1.12500435 | 0.26058719 | NA |
| ENSG0000021 | 0 | NA | NA | NA | NA | NA |
| ENSG0000021 | 0 | NA | NA | NA | NA | NA |
| ENSG0000021 | 0 | NA | NA | NA | NA | NA |
| ENSG0000022 | 1.21390625 | -0.6270177 | 1.07391437 | -0.5838619 | 0.55931323 | NA |
| ENSG0000023 | 0 | NA | NA | NA | NA | NA |
| ENSG0000023 | 0.05944154 | -0.0155188 | 3.04631688 | -0.0050943 | 0.99593536 | NA |
| ENSG0000024 | 1.08803136 | 1.25866368 | 1.14049861 | 1.10360826 | 0.26976311 | NA |
| ENSG0000024 | 0 | NA | NA | NA | NA | NA |
| ENSG0000024 | 0 | NA | NA | NA | NA | NA |
| ENSG0000024 | 0.15980498 | -0.2102568 | 2.01387021 | -0.1044043 | 0.91684848 | NA |
| ENSG0000024 | 0 | NA | NA | NA | NA | NA |
| ENSG0000025 | 0 | NA | NA | NA | NA | NA |
| ENSG0000025 | 0.63376946 | -0.4757447 | 1.1131113 | -0.4274008 | 0.66908739 | NA |
| ENSG0000025 | 0.05944154 | -0.0155188 | 3.04631688 | -0.0050943 | 0.99593536 | NA |
| ENSG0000027 | 0.18845204 | -0.1429887 | 1.97138989 | -0.0725319 | 0.94217862 | NA |
| ENSG0000027 | 0.15568762 | 0.30640095 | 2.95432761 | 0.10371258 | 0.91739744 | NA |
| ENSG0000027 | 0 | NA | NA | NA | NA | NA |
| ENSG0000027 | 0.20151267 | -0.3769624 | 2.25890984 | -0.166878 | 0.86746602 | NA |
| ENSG0000027 | 0.26459647 | -0.2024686 | 1.79416149 | -0.1128486 | 0.9101506 | NA |
| ENSG0000028 | 0 | NA | NA | NA | NA | NA |

| ensg | gene | |
|---|---|---|
| ENSG000001: | STK25 | |
| ENSG000001' | FZD4 | |
| ENSG000001: | GNGT1 | |
| ENSG000001. | NCAM1 | |
| ENSG000001: | ABI2 | |
| ENSG000001( | UNC5B | |
| ENSG000001: | VASP | |
| ENSG000001( | DKK1 | |
| ENSG000001( | ARHGEF40 | |
| ENSG000002: | NTF4 | |
| ENSG000000' | DNM2 | |
| ENSG000001: | ID1 | |
| ENSG000001! | SCLT1 | |
| ENSG000001' | UBB | |
| ENSG000001: | STXBP1 | |
| ENSG000001! | STMN3 | |
| ENSG000001( | CRTC1 | |
| ENSG000002: | TENM3 | |
| ENSG000001: | FN1 | |
| ENSG000001' | GATA2 | |
| ENSG000000' | CRMP1 | |
| ENSG000001: | ARSB | |
| ENSG000001! | UCHL1 | |
| ENSG000001. | ERBB2 | |
| ENSG000001' | ALCAM | |
| ENSG000001. | CARM1 | |
| ENSG000001! | FZD1 | |
| ENSG000000( | NTN1 | |
| ENSG000001! | THY1 | |
| ENSG000001: | P3H1 | |
| ENSG000001: | HTRA2 | |
| ENSG000001! | EPHB1 | |
| ENSG000001! | SZT2 | |
| ENSG000001( | ADGRV1 | |
| ENSG000001( | HAND2 | |
| ENSG000001' | CFL1 | |
| ENSG000001: | RAB35 | |
| ENSG000001: | YWHAH | |
| ENSG000000: | APLP2 | |
| ENSG000000! | ITGA6 | p-val <0.05 |
| ENSG000001! | RAB6B | |
| ENSG000000' | PLXNA2 | |

| | |
|---|---|
| ENSG000001 | SLIT3 |
| ENSG000001 | SEMA7A |
| ENSG000001 | SPOCK1 |
| ENSG000001 | PQBP1 |
| ENSG000001 | IFT20 |
| ENSG000000 | NOTCH3 |
| ENSG000001 | MAP1B |
| ENSG000001 | CREB1 |
| ENSG000001 | SSNA1 |
| ENSG000001 | ADARB1 |
| ENSG000001 | STMN2 |
| ENSG000001 | PDLIM5 |
| ENSG000002 | TWF2 |
| ENSG000000 | MAP4K4 |
| ENSG000001 | THRB |
| ENSG000001 | SRF |
| ENSG000001 | NRP2 |
| ENSG000001 | TOPORS |
| ENSG000001 | PAQR3 |
| ENSG000001 | HDGFL3 |
| ENSG000001 | SIPA1L1 |
| ENSG000001 | NDEL1 |
| ENSG000001 | NFASC |
| ENSG000001 | LIMK1 |
| ENSG000001 | ITSN2 |
| ENSG000001 | NEXN |
| ENSG000001 | TTL |
| ENSG000001 | SAMD14 |
| ENSG000002 | SRGAP2 |
| ENSG000001 | FLRT1 |
| ENSG000001 | DDR2 |
| ENSG000000 | HECW1 |
| ENSG000001 | ACAP3 |
| ENSG000001 | ITM2C |
| ENSG000000 | RPGRIP1 |
| ENSG000001 | MINK1 |
| ENSG000001 | PPT1 |
| ENSG000001 | PTPRG |
| ENSG000001 | PBX3 |
| ENSG000001 | BAIAP2 |
| ENSG000001 | S100A6 |
| ENSG000001 | BLOC1S3 |
| ENSG000001 | CRABP2 |

| | |
|---|---|
| ENSG000002 | FOXD1 |
| ENSG000000 | NGEF |
| ENSG000000 | SEMA3B |
| ENSG000001 | ROM1 |
| ENSG000001 | TUBA1A |
| ENSG000001 | B2M |
| ENSG000001 | PAK1 |
| ENSG000000 | CDH1 |
| ENSG000001 | UNK |
| ENSG000001 | NREP |
| ENSG000000 | CYFIP2 |
| ENSG000000 | PLXND1 |
| ENSG000001 | FEZ2 |
| ENSG000001 | PTPRF |
| ENSG000001 | ARF6 |
| ENSG000001 | VPS54 |
| ENSG000001 | PTK7 |
| ENSG000001 | OSTN |
| ENSG000001 | ULK4 |
| ENSG000000 | CTTN |
| ENSG000001 | CCDC88A |
| ENSG000001 | FLNA |
| ENSG000001 | MFSD8 |
| ENSG000001 | SF3A2 |
| ENSG000001 | ADM |
| ENSG000001 | YTHDF1 |
| ENSG000001 | ZMIZ1 |
| ENSG000001 | GPRIN1 |
| ENSG000001 | XBP1 |
| ENSG000001 | KIRREL3 |
| ENSG000001 | GRIN3A |
| ENSG000001 | TIAM2 |
| ENSG000001 | VAPA |
| ENSG000001 | SIAH1 |
| ENSG000001 | PALLD |
| ENSG000000 | GLI2 |
| ENSG000001 | TSPO |
| ENSG000000 | PAFAH1B1 |
| ENSG000001 | PARD3 |
| ENSG000001 | ATXN10 |
| ENSG000002 | GDI1 |
| ENSG000001 | FBXO7 |
| ENSG000001 | RAP1A |

| | |
|---|---|
| ENSG000001( | DVL3 |
| ENSG000001: | DOCK7 |
| ENSG000002( | DDR1 |
| ENSG000002' | CYFIP1 |
| ENSG000001' | GAK |
| ENSG000001( | CDH23 |
| ENSG000001: | LRP4 |
| ENSG000001‹ | RAB13 |
| ENSG000002( | AGER |
| ENSG000001{ | RNF220 |
| ENSG000000: | RUFY3 |
| ENSG000001( | MICALL2 |
| ENSG000001{ | PRKG1 |
| ENSG000001‹ | SCYL1 |
| ENSG000001( | TNXB |
| ENSG000001( | TSC1 |
| ENSG000001: | MTCH1 |
| ENSG000001: | PLXNA1 |
| ENSG000000! | HSP90AB1 |
| ENSG000001‹ | RETREG3 |
| ENSG000001( | EZH2 |
| ENSG000001: | PRMT1 |
| ENSG000001! | PLXNB3 |
| ENSG000001! | SMURF1 |
| ENSG000000: | TRIO |
| ENSG000001: | MTR |
| ENSG000001( | SFRP1 |
| ENSG000001: | NOTCH2 |
| ENSG000001: | DBNL |
| ENSG000001' | ANAPC2 |
| ENSG000001! | DLG5 |
| ENSG000001‹ | TPBG |
| ENSG000000: | LYPLA2 |
| ENSG000001: | PPP3CA |
| ENSG000001‹ | SLIT2 |
| ENSG000001' | SRGAP2C |
| ENSG000001( | EN1 |
| ENSG000001( | KANK1 |
| ENSG000001! | NPR2 |
| ENSG000001( | DIP2A |
| ENSG000001( | PTPN9 |
| ENSG000000( | ARHGAP44 |
| ENSG000001: | HOXD9 |

| | |
|---|---|
| ENSG000001 | APBB1 |
| ENSG000002 | TUBB3 |
| ENSG000001 | KLF7 |
| ENSG000001 | KIFBP |
| ENSG000000 | HSP90AA1 |
| ENSG000001 | OMG |
| ENSG000001 | CREB3L2 |
| ENSG000001 | APLP1 |
| ENSG000000 | GPM6B |
| ENSG000001 | IFT140 |
| ENSG000001 | VASH2 |
| ENSG000001 | ASAP1 |
| ENSG000001 | TRPV2 |
| ENSG000000 | CTNNA1 |
| ENSG000002 | TCTN1 |
| ENSG000000 | NRDC |
| ENSG000001 | KATNB1 |
| ENSG000001 | CIB1 |
| ENSG000000 | PSEN1 |
| ENSG000001 | ATF5 |
| ENSG000001 | NOTCH1 |
| ENSG000001 | AUTS2 |
| ENSG000001 | CLN5 |
| ENSG000001 | HS6ST1 |
| ENSG000000 | MAP3K13 |
| ENSG000002 | CUX1 |
| ENSG000000 | GSK3B |
| ENSG000001 | STK24 |
| ENSG000001 | STK11 |
| ENSG000001 | GBA1 |
| ENSG000001 | C12orf57 |
| ENSG000001 | LHX9 |
| ENSG000001 | ZPR1 |
| ENSG000001 | PRKD1 |
| ENSG000001 | HOXA2 |
| ENSG000000 | MMP2 |
| ENSG000001 | METRN |
| ENSG000001 | IGF1R |
| ENSG000001 | FZD2 |
| ENSG000001 | MDK |
| ENSG000000 | TOP2B |
| ENSG000001 | SMAD4 |
| ENSG000001 | NBL1 |

| | |
|---|---|
| ENSG0000001 | NRN1 |
| ENSG0000001 | VLDLR |
| ENSG0000000 | PSD |
| ENSG0000001 | MAP1A |
| ENSG0000001 | DNM3 |
| ENSG0000001 | ENAH |
| ENSG0000001 | RAC1 |
| ENSG0000001 | ECE1 |
| ENSG0000000 | SEMA3C |
| ENSG0000001 | ZNF212 |
| ENSG0000001 | TBCD |
| ENSG0000000 | NEO1 |
| ENSG0000001 | CERS2 |
| ENSG0000000 | WNK1 |
| ENSG0000001 | NTM |
| ENSG0000000 | PHGDH |
| ENSG0000001 | NTN3 |
| ENSG0000002 | NCKIPSD |
| ENSG0000001 | MAP1S |
| ENSG0000001 | HOXD10 |
| ENSG0000001 | AFG3L2 |
| ENSG0000002 | NR2E3 |
| ENSG0000000 | PAX6 |
| ENSG0000001 | TMEM108 |
| ENSG0000000 | MTMR2 |
| ENSG0000001 | PHACTR1 |
| ENSG0000001 | MAP2K2 |
| ENSG0000002 | TBCE |
| ENSG0000001 | DVL1 |
| ENSG0000000 | NCK2 |
| ENSG0000000 | RAB10 |
| ENSG0000001 | PTEN |
| ENSG0000001 | AVIL |
| ENSG0000001 | NPTN |
| ENSG0000000 | PRKCZ |
| ENSG0000001 | BBS4 |
| ENSG0000001 | CDH2 |
| ENSG0000001 | TAOK2 |
| ENSG0000001 | KCNB1 |
| ENSG0000001 | SOS1 |
| ENSG0000001 | TMEM30A |
| ENSG0000000 | DVL2 |
| ENSG0000000 | SYT1 |

| | |
|---|---|
| ENSG000001 | KIAA1755 |
| ENSG000001 | MYOT |
| ENSG000001 | SYT2 |
| ENSG000001 | GHRL |
| ENSG000001 | CDH4 |
| ENSG000000 | GBA2 |
| ENSG000001 | WNT5A |
| ENSG000001 | WEE1 |
| ENSG000001 | RUNX1 |
| ENSG000000 | MARK2 |
| ENSG000000 | ITGA3 |
| ENSG000001 | SHANK1 |
| ENSG000001 | TSC22D4 |
| ENSG000001 | FLRT2 |
| ENSG000001 | ROR1 |
| ENSG000001 | AKT1 |
| ENSG000001 | CDNF |
| ENSG000001 | WDR5 |
| ENSG000001 | BECN1 |
| ENSG000001 | NPHP4 |
| ENSG000001 | ITGA4 |
| ENSG000000 | RHOA |
| ENSG000001 | CPEB3 |
| ENSG000000 | FARP2 |
| ENSG000001 | RAB8A |
| ENSG000001 | EFNB2 |
| ENSG000001 | RERE |
| ENSG000001 | ADNP |
| ENSG000001 | VPS13B |
| ENSG000001 | RAB11A |
| ENSG000001 | ATL1 |
| ENSG000000 | NCKAP1 |
| ENSG000001 | PPP3CB |
| ENSG000001 | OGDH |
| ENSG000000 | MYCBP2 |
| ENSG000001 | THOC2 |
| ENSG000001 | SS18L1 |
| ENSG000001 | UBE3A |
| ENSG000001 | SOD1 |
| ENSG000001 | PTPN11 |
| ENSG000000 | FKBP4 |
| ENSG000001 | DICER1 |
| ENSG000001 | SHOX2 |

| | |
|---|---|
| ENSG000001( | SIN3A |
| ENSG000001: | SCARB2 |
| ENSG000000: | TBC1D23 |
| ENSG000001( | MECP2 |
| ENSG000001( | YWHAZ |
| ENSG000001( | WASL |
| ENSG000001( | ANKRD27 |
| ENSG000001( | GSK3A |
| ENSG000001‹ | CRB2 |
| ENSG000001( | TAOK1 |
| ENSG000001( | SETX |
| ENSG000001: | PLXNC1 |
| ENSG000000! | LAMB1 |
| ENSG000001( | CRK |
| ENSG000001: | PLXNA3 |
| ENSG000001( | NDRG4 |
| ENSG000000( | SCYL3 |
| ENSG000001: | RTN4 |
| ENSG000001: | DDX56 |
| ENSG000000: | VIM |
| ENSG000001: | IST1 |
| ENSG000001( | NEFH |
| ENSG000001ʹ | TANC2 |
| ENSG000001( | CXCL12 |
| ENSG000001( | SDC2 |
| ENSG000000! | EFNB1 |
| ENSG000000‹ | CUL7 |
| ENSG000001: | SEMA5A |
| ENSG000001! | LPAR1 |
| ENSG000001! | CEP290 |
| ENSG000001: | MACF1 |
| ENSG000001: | RNF6 |
| ENSG000001ʹ | FBXO45 |
| ENSG000001ʹ | NEGR1 |
| ENSG000001‹ | TNFRSF21 |
| ENSG000001: | SEMA4D |
| ENSG000001‹ | CDH11 |
| ENSG000000ʹ | RIMS1 |
| ENSG000000! | NRP1 |
| ENSG000001! | CUL4B |
| ENSG000001! | SYNGAP1 |
| ENSG000001( | CTHRC1 |
| ENSG000002: | DHFR |

| | |
|---|---|
| ENSG000001 | KIDINS220 |
| ENSG000001 | ITGB1 |
| ENSG000001 | ARHGAP35 |
| ENSG000001 | GFAP |
| ENSG000000 | SEMA4G |
| ENSG000001 | RAPGEF2 |
| ENSG000001 | FES |
| ENSG000000 | NTN4 |
| ENSG000001 | STX3 |
| ENSG000001 | TSKU |
| ENSG000001 | PCDH12 |
| ENSG000001 | EHD1 |
| ENSG000000 | MYO9A |
| ENSG000001 | NR2F1 |
| ENSG000001 | UNC5C |
| ENSG000001 | CNTNAP1 |
| ENSG000001 | PTK2 |
| ENSG000001 | VEGFA |
| ENSG000002 | ITGA1 |
| ENSG000001 | ABI1 |
| ENSG000001 | EP300 |
| ENSG000000 | WHRN |
| ENSG000002 | PRAG1 |
| ENSG000000 | VCL |
| ENSG000001 | BDNF |
| ENSG000001 | MAP6 |
| ENSG000000 | MUL1 |
| ENSG000000 | CRKL |
| ENSG000001 | SERPINF1 |
| ENSG000001 | CSF1R |
| ENSG000000 | PALS1 |
| ENSG000001 | PLEKHG4B |
| ENSG000000 | NCDN |
| ENSG000001 | PRRX1 |
| ENSG000001 | PLEKHG4 |
| ENSG000001 | GIT1 |
| ENSG000001 | EXT1 |
| ENSG000001 | PGRMC1 |
| ENSG000001 | OBSL1 |
| ENSG000000 | SARM1 |
| ENSG000001 | TENM2 |
| ENSG000001 | NLGN2 |
| ENSG000001 | PRICKLE1 |

| | |
|---|---|
| ENSG000001( | NCS1 |
| ENSG000001{ | AGRN |
| ENSG000001( | SPG11 |
| ENSG000001' | PTPRM |
| ENSG000000' | PICALM |
| ENSG000000( | SPAG9 |
| ENSG000001! | LHFPL5 |
| ENSG000001( | EEF2K |
| ENSG000001( | BAG5 |
| ENSG000001! | HDAC2 |
| ENSG000001‹ | LRP12 |
| ENSG000001' | RAPH1 |
| ENSG000002: | HEXA |
| ENSG000001( | MEGF8 |
| ENSG000001: | CAMK1 |
| ENSG000000: | SLC4A7 |
| ENSG000000( | ROCK1 |
| ENSG000001( | NEURL1 |
| ENSG000001! | DIXDC1 |
| ENSG000001! | PLXNB2 |
| ENSG000001: | DBN1 |
| ENSG000001( | TRIP11 |
| ENSG000000( | SYN1 |
| ENSG000001( | FBXO31 |
| ENSG000000! | PDE6C |
| ENSG000001: | MAPK8IP3 |
| ENSG000001' | ENC1 |
| ENSG000001: | LLGL1 |
| ENSG000001' | ULK1 |
| ENSG000001{ | SHTN1 |
| ENSG000001: | NIBAN2 |
| ENSG000001! | BRAF |
| ENSG000001‹ | IQSEC1 |
| ENSG000002( | BMPR2 |
| ENSG000001{ | PDZD7 |
| ENSG000001( | GDNF |
| ENSG000001( | PPP1R9B |
| ENSG000001! | ZNF335 |
| ENSG000001: | NUMB |
| ENSG000001' | RAB6A |
| ENSG000001{ | NOG |
| ENSG000001‹ | CAMK2G |
| ENSG000000! | ABL1 |

| | |
|---|---|
| ENSG000000 | EDN1 |
| ENSG000001 | SDC4 |
| ENSG000001 | GNAT2 |
| ENSG000002 | FOXO6 |
| ENSG000001 | CNP |
| ENSG000001 | CDC20 |
| ENSG000001 | NR2F6 |
| ENSG000001 | ADCY6 |
| ENSG000001 | USP9X |
| ENSG000001 | HERC1 |
| ENSG000001 | SPTBN4 |
| ENSG000001 | BBS1 |
| ENSG000001 | NDN |
| ENSG000001 | DAB2IP |
| ENSG000001 | PPFIA2 |
| ENSG000001 | PMP22 |
| ENSG000001 | NTF3 |
| ENSG000000 | IFRD1 |
| ENSG000000 | EPB41L3 |
| ENSG000001 | PAK4 |
| ENSG000001 | BOC |
| ENSG000001 | SEMA4C |
| ENSG000001 | GORASP1 |
| ENSG000000 | MUSK |
| ENSG000001 | MAGI2 |
| ENSG000001 | CDK16 |
| ENSG000001 | ARF4 |
| ENSG000002 | ERCC6 |
| ENSG000001 | TAOK3 |
| ENSG000000 | ATP8B1 |
| ENSG000001 | ABITRAM |
| ENSG000000 | SEMA6A |
| ENSG000001 | NSMF |
| ENSG000001 | SNX3 |
| ENSG000001 | LHX4 |
| ENSG000000 | SPAST |
| ENSG000001 | TOR1A |
| ENSG000001 | LAMB2 |
| ENSG000001 | HMGB1 |
| ENSG000000 | FYN |
| ENSG000001 | HECW2 |
| ENSG000001 | SLC9A3R1 |
| ENSG000000 | GRN |

| | |
|---|---|
| ENSG000000 | HSPA5 |
| ENSG000001 | SPART |
| ENSG000001 | DAB2 |
| ENSG000001 | DENND5A |
| ENSG000001 | B4GALT5 |
| ENSG000001 | TRAPPC4 |
| ENSG000000 | PPP2R5B |
| ENSG000001 | SCRIB |
| ENSG000001 | ZDHHC17 |
| ENSG000001 | PTCH1 |
| ENSG000000 | MAP4 |
| ENSG000001 | MICALL1 |
| ENSG000001 | TRIOBP |
| ENSG000001 | NUMBL |
| ENSG000001 | PTPRS |
| ENSG000001 | ROBO1 |
| ENSG000001 | SLC9A6 |
| ENSG000001 | CAMSAP1 |
| ENSG000001 | DLG4 |
| ENSG000001 | FARP1 |
| ENSG000001 | TWF1 |
| ENSG000001 | UPF3B |
| ENSG000001 | STMN1 |
| ENSG000001 | CAPRIN1 |
| ENSG000001 | FLOT1 |
| ENSG000001 | PBX1 |
| ENSG000001 | NFE2L2 |
| ENSG000001 | MANF |
| ENSG000000 | LRRC7 |
| ENSG000001 | IQGAP1 |
| ENSG000001 | MAP2K1 |
| ENSG000000 | MEF2A |
| ENSG000001 | NIN |
| ENSG000001 | EIF2AK4 |
| ENSG000001 | NFATC4 |
| ENSG000001 | CTNNB1 |
| ENSG000000 | TNC |
| ENSG000000 | MAPK6 |
| ENSG000001 | KIF5B |
| ENSG000001 | DPYSL3 |
| ENSG000000 | MAPKAPK5 |
| ENSG000001 | DAG1 |
| ENSG000001 | GOLGA4 |

| | |
|---|---|
| ENSG000001( | NAGLU |
| ENSG000001⁄ | APP |
| ENSG000001! | LRP8 |
| ENSG000000( | KDM1A |
| ENSG000001⁄ | FEZ1 |
| ENSG000001⁄ | GDF7 |
| ENSG000001⁄ | ABL2 |
| ENSG000001⁄ | WDPCP |
| ENSG000000' | CBFA2T2 |
| ENSG000001: | RB1 |
| ENSG000001' | FBXW8 |
| ENSG000001: | KLF4 |
| ENSG000001' | JUN |
| ENSG000001: | PAK2 |
| ENSG000001! | MOV10 |
| ENSG000001' | SVBP |
| ENSG000002( | PBX2 |
| ENSG000000( | CFLAR |
| ENSG000001( | RAPGEF1 |
| ENSG000001( | UGDH |
| ENSG000001( | GLI3 |
| ENSG000001! | INPP5F |
| ENSG000001! | MBP |
| ENSG000001( | SLC25A46 |
| ENSG000000( | SEMA3F |
| ENSG000000( | ALS2 |
| ENSG000000( | CD38 |
| ENSG000000( | ARHGAP33 |
| ENSG000000( | ARX |
| ENSG000000( | CX3CL1 |
| ENSG000000( | ETV1 |
| ENSG000000( | USH1C |
| ENSG000000( | CDKL3 |
| ENSG000000( | GAS7 |
| ENSG000000( | MYLIP |
| ENSG000000( | CDKL5 |
| ENSG000000( | MAPK8IP2 |
| ENSG000000( | TENM1 |
| ENSG000000: | SEMA3G |
| ENSG000000: | MKS1 |
| ENSG000000: | ANOS1 |
| ENSG000000: | STMN4 |
| ENSG000000: | ISL1 |

| | |
|---|---|
| ENSG000000 | CNTN1 |
| ENSG000000 | HGF |
| ENSG000000 | RUNX3 |
| ENSG000000 | NRXN3 |
| ENSG000000 | IFT88 |
| ENSG000000 | UBA6 |
| ENSG000000 | MYOC |
| ENSG000000 | STAU2 |
| ENSG000000 | RTN4R |
| ENSG000000 | MYO16 |
| ENSG000000 | EPHA3 |
| ENSG000000 | PREX2 |
| ENSG000000 | DTNBP1 |
| ENSG000000 | XK |
| ENSG000000 | NEDD4L |
| ENSG000000 | FSTL4 |
| ENSG000000 | KCNQ1 |
| ENSG000000 | PRDM1 |
| ENSG000000 | RASGRF1 |
| ENSG000000 | CAMK2B |
| ENSG000000 | DGKG |
| ENSG000000 | LZTS1 |
| ENSG000000 | LTK |
| ENSG000000 | SEZ6 |
| ENSG000000 | NGFR |
| ENSG000000 | ANKS1A |
| ENSG000000 | PRKCQ |
| ENSG000000 | CTNNA2 |
| ENSG000000 | DIP2B |
| ENSG000000 | FGFR2 |
| ENSG000000 | PLA2G10 |
| ENSG000000 | NEDD4 |
| ENSG000000 | CNGB1 |
| ENSG000000 | CAMK2A |
| ENSG000000 | CDC42 |
| ENSG000000 | EPHA8 |
| ENSG000000 | TRPC5 |
| ENSG000000 | FRY |
| ENSG000000 | SCARF1 |
| ENSG000000 | SEMA3A |
| ENSG000000 | FRYL |
| ENSG000000 | PAX2 |
| ENSG000000 | CAMSAP3 |

| | |
|---|---|
| ENSG00000007 | ACTL6B |
| ENSG00000007 | USP33 |
| ENSG00000007 | PAK3 |
| ENSG00000007 | SPAG6 |
| ENSG00000007 | MAP2 |
| ENSG00000007 | SLC1A3 |
| ENSG00000007 | OPHN1 |
| ENSG00000008 | EPHA6 |
| ENSG00000008 | RAB21 |
| ENSG00000008 | CHRNA3 |
| ENSG00000008 | MEF2C |
| ENSG00000008 | LRP2 |
| ENSG00000008 | SEMA5B |
| ENSG00000008 | ULK2 |
| ENSG00000008 | WDR47 |
| ENSG00000008 | IGSF9 |
| ENSG00000008 | MT3 |
| ENSG00000008 | AURKA |
| ENSG00000008 | LZTS3 |
| ENSG00000008 | SLC23A2 |
| ENSG00000008 | DZANK1 |
| ENSG00000008 | ARHGAP4 |
| ENSG00000009 | POU4F3 |
| ENSG00000009 | NRCAM |
| ENSG00000009 | TGFB2 |
| ENSG00000009 | HDAC6 |
| ENSG00000009 | CRTAC1 |
| ENSG00000009 | JAK2 |
| ENSG00000009 | STX1B |
| ENSG00000009 | EFNA2 |
| ENSG00000009 | CECR2 |
| ENSG00000010 | PLA2G3 |
| ENSG00000010 | PICK1 |
| ENSG00000010 | IFT27 |
| ENSG00000010 | ALKBH1 |
| ENSG00000010 | RPS6KA5 |
| ENSG00000010 | CPNE6 |
| ENSG00000010 | VSX1 |
| ENSG00000010 | BMP7 |
| ENSG00000010 | PTK6 |
| ENSG00000010 | MCF2 |
| ENSG00000010 | FMR1 |
| ENSG00000010 | ZDHHC15 |

| | |
|---|---|
| ENSG000001( | RPGRIP1L |
| ENSG000001( | SYT17 |
| ENSG000001( | BLOC1S6 |
| ENSG000001( | RP1 |
| ENSG000001( | FZD3 |
| ENSG000001( | NOVA2 |
| ENSG000001( | CACNG7 |
| ENSG000001( | RAB3A |
| ENSG000001( | MAG |
| ENSG000001( | SCN1B |
| ENSG000001( | PBX4 |
| ENSG000001( | DLX5 |
| ENSG000001( | PTN |
| ENSG000001( | EVX1 |
| ENSG000001( | COBL |
| ENSG000001( | EPHB6 |
| ENSG000001( | PTPRZ1 |
| ENSG000001( | TMEM106B |
| ENSG000001( | LHX2 |
| ENSG000001( | LHX6 |
| ENSG000001( | LHX3 |
| ENSG000001( | SH3GL2 |
| ENSG000001( | GATA3 |
| ENSG000001( | PITX3 |
| ENSG000001( | LGI1 |
| ENSG000001( | WNT3 |
| ENSG000001( | ABI3 |
| ENSG000001( | RND2 |
| ENSG000001( | EFNB3 |
| ENSG000001( | PHOX2B |
| ENSG000001( | AREG |
| ENSG000001( | IL2 |
| ENSG000001: | FOLR1 |
| ENSG000001: | APOA4 |
| ENSG000001: | NECTIN1 |
| ENSG000001: | CAPRIN2 |
| ENSG000001: | SLC11A2 |
| ENSG000001: | TRPV4 |
| ENSG000001: | CUX2 |
| ENSG000001: | KCNA1 |
| ENSG000001: | RASAL1 |
| ENSG000001: | TDP2 |
| ENSG000001: | UST |

| | |
|---|---|
| ENSG000001 | TULP1 |
| ENSG000001 | BMP5 |
| ENSG000001 | WASF1 |
| ENSG000001 | NR2E1 |
| ENSG000001 | FIG4 |
| ENSG000001 | PRPH2 |
| ENSG000001 | UNC5A |
| ENSG000001 | HES1 |
| ENSG000001 | GNAT1 |
| ENSG000001 | CSPG5 |
| ENSG000001 | DGUOK |
| ENSG000001 | TLX2 |
| ENSG000001 | EFHD1 |
| ENSG000001 | STRN |
| ENSG000001 | VAX2 |
| ENSG000001 | EPHA4 |
| ENSG000001 | MARK1 |
| ENSG000001 | TNR |
| ENSG000001 | RGS2 |
| ENSG000001 | RAB29 |
| ENSG000001 | ARTN |
| ENSG000001 | PLPPR5 |
| ENSG000001 | PLPPR4 |
| ENSG000001 | CAMSAP2 |
| ENSG000001 | B4GALT6 |
| ENSG000001 | CNR1 |
| ENSG000001 | SGK1 |
| ENSG000001 | OLFM3 |
| ENSG000001 | SPP1 |
| ENSG000001 | TRIM67 |
| ENSG000001 | TRIM32 |
| ENSG000001 | ONECUT2 |
| ENSG000001 | FKBP1B |
| ENSG000001 | BCL11A |
| ENSG000001 | KCNIP2 |
| ENSG000001 | TNN |
| ENSG000001 | PLS1 |
| ENSG000001 | PTK2B |
| ENSG000001 | SLITRK3 |
| ENSG000001 | CXCR4 |
| ENSG000001 | NPY |
| ENSG000001 | NEUROG3 |
| ENSG000001 | EGR2 |

| | |
|---|---|
| ENSG000001 | ATF1 |
| ENSG000001 | NEUROD4 |
| ENSG000001 | NCKAP1L |
| ENSG000001 | PLP1 |
| ENSG000001 | SLC12A5 |
| ENSG000001 | PARD6B |
| ENSG000001 | EDN3 |
| ENSG000001 | PACSIN1 |
| ENSG000001 | CPNE5 |
| ENSG000001 | RAB17 |
| ENSG000001 | RAP2A |
| ENSG000001 | C3 |
| ENSG000001 | FLRT3 |
| ENSG000001 | ZC4H2 |
| ENSG000001 | EDN2 |
| ENSG000001 | BCL11B |
| ENSG000001 | STYXL1 |
| ENSG000001 | ADORA2A |
| ENSG000001 | LIF |
| ENSG000001 | SMO |
| ENSG000001 | FEZF1 |
| ENSG000001 | CHN1 |
| ENSG000001 | ATP1B2 |
| ENSG000001 | KLK8 |
| ENSG000001 | FGF13 |
| ENSG000001 | CDKN1C |
| ENSG000001 | APOE |
| ENSG000001 | KIF1A |
| ENSG000001 | RTN4IP1 |
| ENSG000001 | EPO |
| ENSG000001 | UNC13A |
| ENSG000001 | SULT4A1 |
| ENSG000001 | OLFM1 |
| ENSG000001 | MNX1 |
| ENSG000001 | PRDM12 |
| ENSG000001 | C1QL1 |
| ENSG000001 | SNAP25 |
| ENSG000001 | BTBD3 |
| ENSG000001 | SYT4 |
| ENSG000001 | ATP8A2 |
| ENSG000001 | LGR6 |
| ENSG000001 | DCLK1 |
| ENSG000001 | EPHB2 |

| | |
|---|---|
| ENSG000001 | CNTN6 |
| ENSG000001 | CHL1 |
| ENSG000001 | TSPAN2 |
| ENSG000001 | NGF |
| ENSG000001 | CRB1 |
| ENSG000001 | AGTPBP1 |
| ENSG000001 | BICDL1 |
| ENSG000001 | ADGRB3 |
| ENSG000001 | EPHA7 |
| ENSG000001 | BLOC1S1 |
| ENSG000001 | AHI1 |
| ENSG000001 | SEMA4F |
| ENSG000001 | EMX1 |
| ENSG000001 | AGT |
| ENSG000001 | DOCK10 |
| ENSG000001 | SCYL2 |
| ENSG000001 | NEK3 |
| ENSG000001 | EDNRB |
| ENSG000001 | IL6 |
| ENSG000001 | NKX2-8 |
| ENSG000001 | NKX2-1 |
| ENSG000001 | TBR1 |
| ENSG000001 | SKIL |
| ENSG000001 | PLAA |
| ENSG000001 | KIAA0319 |
| ENSG000001 | TUBB2B |
| ENSG000001 | ATAT1 |
| ENSG000001 | MGARP |
| ENSG000001 | MYO7A |
| ENSG000001 | PARP6 |
| ENSG000001 | ITPKA |
| ENSG000001 | PAK6 |
| ENSG000001 | SEMA6D |
| ENSG000001 | ARL3 |
| ENSG000001 | KIF20B |
| ENSG000001 | ZNF365 |
| ENSG000001 | MYPN |
| ENSG000001 | BMPR1B |
| ENSG000001 | SEC24B |
| ENSG000001 | LLPH |
| ENSG000001 | ASCL1 |
| ENSG000001 | SRRM4 |
| ENSG000001 | NTRK3 |

| | |
|---|---|
| ENSG000001 | ST8SIA2 |
| ENSG000001 | ADCYAP1 |
| ENSG000001 | RNF157 |
| ENSG000001 | RNF165 |
| ENSG000001 | ACP4 |
| ENSG000001 | LMO4 |
| ENSG000001 | BARHL2 |
| ENSG000001 | CELSR2 |
| ENSG000001 | SEMA6C |
| ENSG000001 | SNAPIN |
| ENSG000001 | EFNA3 |
| ENSG000001 | SLC4A10 |
| ENSG000001 | NYAP2 |
| ENSG000001 | CPNE9 |
| ENSG000001 | CNTN4 |
| ENSG000001 | LRTM1 |
| ENSG000001 | EPHA5 |
| ENSG000001 | SFRP2 |
| ENSG000001 | GABRB2 |
| ENSG000001 | FBXO38 |
| ENSG000001 | GFRA3 |
| ENSG000001 | DCDC2 |
| ENSG000001 | SDK1 |
| ENSG000001 | NFIB |
| ENSG000001 | C9orf72 |
| ENSG000001 | NTRK2 |
| ENSG000001 | UGCG |
| ENSG000001 | HMCN2 |
| ENSG000001 | SLC39A12 |
| ENSG000001 | CDHR1 |
| ENSG000001 | VAX1 |
| ENSG000001 | LRRC4C |
| ENSG000001 | TENM4 |
| ENSG000001 | DRD2 |
| ENSG000001 | TBX6 |
| ENSG000001 | CNTN5 |
| ENSG000001 | CTF1 |
| ENSG000001 | KLHL1 |
| ENSG000001 | GPM6A |
| ENSG000001 | ANK3 |
| ENSG000001 | PTPRO |
| ENSG000001 | POU4F2 |
| ENSG000001 | EDNRA |

| | |
|---|---|
| ENSG000001! | GFRA1 |
| ENSG000001! | POU4F1 |
| ENSG000001! | GRID2 |
| ENSG000001! | RIT2 |
| ENSG000001! | UHMK1 |
| ENSG000001! | NR4A2 |
| ENSG000001! | FEZF2 |
| ENSG000001! | ADGRF1 |
| ENSG000001! | PTPRD |
| ENSG000001! | LGI4 |
| ENSG000001! | SEMA3D |
| ENSG000001! | IMPACT |
| ENSG000001! | ROBO4 |
| ENSG000001! | ROBO3 |
| ENSG000001! | TNIK |
| ENSG000001! | WNT3A |
| ENSG000001! | C21orf91 |
| ENSG000001! | CHODL |
| ENSG000001! | NCAM2 |
| ENSG000001! | WNT7A |
| ENSG000001! | ZFYVE27 |
| ENSG000001! | GRIP1 |
| ENSG000001! | KIF5A |
| ENSG000001! | CFAP418 |
| ENSG000001! | TIAM1 |
| ENSG000001! | DPYSL5 |
| ENSG000001! | NCK1 |
| ENSG000001! | PPP1R9A |
| ENSG000001! | IGF2BP1 |
| ENSG000001! | BTG2 |
| ENSG000001! | ISL2 |
| ENSG000001( | KALRN |
| ENSG000001( | S100B |
| ENSG000001( | RDH13 |
| ENSG000001( | BRSK1 |
| ENSG000001( | CHRNB2 |
| ENSG000001( | TBC1D24 |
| ENSG000001( | ELAVL4 |
| ENSG000001( | DRAXIN |
| ENSG000001( | LHX8 |
| ENSG000001( | NTNG1 |
| ENSG000001( | GFI1 |
| ENSG000001( | LMX1A |

| | |
|---|---|
| ENSG000001( | DISC1 |
| ENSG000001( | S100A9 |
| ENSG000001( | GABRB1 |
| ENSG000001( | CCKAR |
| ENSG000001( | TRIM46 |
| ENSG000001( | FEV |
| ENSG000001( | IHH |
| ENSG000001( | SERPINI1 |
| ENSG000001( | PRKCI |
| ENSG000001( | NKX6-1 |
| ENSG000001( | CLRN1 |
| ENSG000001( | RYK |
| ENSG000001( | UCN |
| ENSG000001( | PLXNB1 |
| ENSG000001( | CASP3 |
| ENSG000001( | UQCRQ |
| ENSG000001( | HCN1 |
| ENSG000001( | SHH |
| ENSG000001( | ADCY1 |
| ENSG000001( | EN2 |
| ENSG000001( | CSMD3 |
| ENSG000001( | CDK5 |
| ENSG000001( | GBX1 |
| ENSG000001( | TMC1 |
| ENSG000001( | ATP7A |
| ENSG000001( | SLITRK5 |
| ENSG000001( | FAT3 |
| ENSG000001( | TTC8 |
| ENSG000001( | OTX2 |
| ENSG000001( | DRGX |
| ENSG000001( | FRMD7 |
| ENSG000001( | HPRT1 |
| ENSG000001( | RET |
| ENSG000001( | CLMN |
| ENSG000001( | LRTM2 |
| ENSG000001( | NYAP1 |
| ENSG000001( | ISLR2 |
| ENSG000001( | ATCAY |
| ENSG000001( | SEMA6B |
| ENSG000001( | KLK6 |
| ENSG000001( | KIF5C |
| ENSG000001( | SCN11A |
| ENSG000001( | MFSD2A |

| | |
|---|---|
| ENSG000001 | GBX2 |
| ENSG000001 | SPRY3 |
| ENSG000001 | SPRY3 |
| ENSG000001 | ROR2 |
| ENSG000001 | EFNA1 |
| ENSG000001 | IL1RAPL1 |
| ENSG000001 | MINAR1 |
| ENSG000001 | RAC3 |
| ENSG000001 | NLGN1 |
| ENSG000001 | CTNND2 |
| ENSG000001 | B3GNT2 |
| ENSG000001 | SEMA3E |
| ENSG000001 | DCLK2 |
| ENSG000001 | ZNF804A |
| ENSG000001 | EMB |
| ENSG000001 | ZNF296 |
| ENSG000001 | GPR37 |
| ENSG000001 | OR10A4 |
| ENSG000001 | ALK |
| ENSG000001 | NRTN |
| ENSG000001 | NPTX1 |
| ENSG000001 | LPAR3 |
| ENSG000001 | NEUROD2 |
| ENSG000001 | DSCAM |
| ENSG000001 | BCL2 |
| ENSG000001 | KNDC1 |
| ENSG000001 | FOXB1 |
| ENSG000001 | GAP43 |
| ENSG000001 | MBOAT1 |
| ENSG000001 | ATOH1 |
| ENSG000001 | TTC36 |
| ENSG000001 | FUT9 |
| ENSG000001 | RND1 |
| ENSG000001 | C1QA |
| ENSG000001 | NDNF |
| ENSG000001 | INSM1 |
| ENSG000001 | DAB1 |
| ENSG000001 | CNTNAP2 |
| ENSG000001 | UGT8 |
| ENSG000001 | BRSK2 |
| ENSG000001 | B4GAT1 |
| ENSG000001 | LEP |
| ENSG000001 | DHX36 |

| | |
|---|---|
| ENSG0000017 | CHRNA7 |
| ENSG0000017 | CABP4 |
| ENSG0000017 | AMIGO3 |
| ENSG0000017 | TPRN |
| ENSG0000017 | FOXG1 |
| ENSG0000017 | RIMS2 |
| ENSG0000017 | CDK5R1 |
| ENSG0000017 | DSCAML1 |
| ENSG0000017 | NHLH2 |
| ENSG0000017 | SLITRK1 |
| ENSG0000017 | PCARE |
| ENSG0000017 | SLITRK4 |
| ENSG0000017 | ATOH7 |
| ENSG0000017 | NRXN1 |
| ENSG0000018 | TH |
| ENSG0000018 | TMEM132E |
| ENSG0000018 | AMIGO1 |
| ENSG0000018 | ADGRB1 |
| ENSG0000018 | USH1G |
| ENSG0000018 | RGMA |
| ENSG0000018 | EPHB3 |
| ENSG0000018 | GALR2 |
| ENSG0000018 | SOX1 |
| ENSG0000018 | CAMK1D |
| ENSG0000018 | EPHA10 |
| ENSG0000018 | RP1L1 |
| ENSG0000018 | EFHC2 |
| ENSG0000018 | OPCML |
| ENSG0000018 | KREMEN1 |
| ENSG0000018 | CNTN2 |
| ENSG0000018 | EFNA5 |
| ENSG0000018 | POU3F2 |
| ENSG0000018 | SLITRK6 |
| ENSG0000018 | DRD1 |
| ENSG0000018 | ROBO2 |
| ENSG0000018 | SEMA4B |
| ENSG0000018 | INPP5J |
| ENSG0000018 | PRMT3 |
| ENSG0000018 | PRKN |
| ENSG0000018 | GPRIN3 |
| ENSG0000018 | RTN4RL1 |
| ENSG0000018 | SLITRK2 |
| ENSG0000018 | PLK5 |

| | |
|---|---|
| ENSG00000018 | AGBL4 |
| ENSG00000018 | BLOC1S4 |
| ENSG00000018 | GABRA5 |
| ENSG00000018 | GLDN |
| ENSG00000018 | MYT1L |
| ENSG00000018 | FSCN2 |
| ENSG00000018 | MAPT |
| ENSG00000018 | RTN4RL2 |
| ENSG00000018 | CCK |
| ENSG00000018 | SLIT1 |
| ENSG00000018 | DCC |
| ENSG00000018 | SECISBP2 |
| ENSG00000018 | NRN1L |
| ENSG00000018 | WNT7B |
| ENSG00000018 | BLOC1S5 |
| ENSG00000018 | NANOS1 |
| ENSG00000018 | LRRK2 |
| ENSG00000018 | DHFRP1 |
| ENSG00000018 | RELN |
| ENSG00000018 | APOD |
| ENSG00000018 | ALKAL2 |
| ENSG00000019 | BLOC1S2 |
| ENSG00000019 | SEMA4A |
| ENSG00000019 | GRM7 |
| ENSG00000019 | NLGN3 |
| ENSG00000019 | NTNG2 |
| ENSG00000019 | ALKAL1 |
| ENSG00000019 | DIO3 |
| ENSG00000019 | KIF13B |
| ENSG00000019 | HES5 |
| ENSG00000019 | KEL |
| ENSG00000019 | NTRK1 |
| ENSG00000019 | ARC |
| ENSG00000019 | LRIG2 |
| ENSG00000019 | TOX |
| ENSG00000019 | CD3E |
| ENSG00000019 | GPRASP3 |
| ENSG00000019 | L1CAM |
| ENSG00000019 | DMD |
| ENSG00000019 | RORB |
| ENSG00000019 | MIR219A1 |
| ENSG00000019 | MIR210 |
| ENSG00000019 | MIR133B |

| | |
|---|---|
| ENSG000002 | KHDC3L |
| ENSG000002 | NEU4 |
| ENSG000002 | GPRIN2 |
| ENSG000002 | PJVK |
| ENSG000002 | SLC44A4 |
| ENSG000002 | LST1 |
| ENSG000002 | GRXCR2 |
| ENSG000002 | MIR200C |
| ENSG000002 | MIR222 |
| ENSG000002 | MIR221 |
| ENSG000002 | MIR431 |
| ENSG000002 | SYT3 |
| ENSG000002 | GPC2 |
| ENSG000002 | CPNE1 |
| ENSG000002 | CRPPA |
| ENSG000002 | SYT14P1 |
| ENSG000002 | GRXCR1 |
| ENSG000002 | SKOR2 |
| ENSG000002 | PLXNA4 |
| ENSG000002 | STMND1 |
| ENSG000002 | MFRP |
| ENSG000002 | ARHGEF25 |
| ENSG000002 | CNTF |
| ENSG000002 | STRC |
| ENSG000002 | EFNA4 |
| ENSG000002 | CLRN2 |
| ENSG000002 | TUNAR |
| ENSG000002 | SHANK3 |
| ENSG000002 | LYN |
| ENSG000002 | MICOS10-NBL1 |
| ENSG000002 | ARMCX5-GPRASP2 |
| ENSG000002 | LHX1 |
| ENSG000002 | SRCIN1 |
| ENSG000002 | NEFL |
| ENSG000002 | TBCE |
