## Supplementary material for "TBCK Deficiency Alters Ribosomal Function, RNA Splicing, and miRNA Networks: Insights from Multi-Omics Analyses": splicing_maser-filtered-rMATs-events

| GeneID | geneSymbol | PValue | FDR | IncLevelDifference | PSI_1 | PSI_2 |
|---|---|---|---|---|---|---|
| ENSG000001 | SLC3A2 | 4.17E-12 | 6.94E-09 | -0.484 | 0.163,0.328,0 | 0.823,0.919,1 |
| ENSG000002 | FAM86DP | 0 | 0 | -0.435 | 0.03,0,0,0,0,0 | 0.4,0.75,0.34 |
| ENSG000001 | ADAM15 | 0 | 0 | -0.417 | 0.339,0.52,0. | 0.995,1,0.995 |
| ENSG000001 | POSTN | 8.08E-09 | 5.5523E-06 | -0.369 | 0.149,0.181,0 | 0.978,0.865,0 |
| ENSG000001 | PER1 | 1.59E-12 | 2.94E-09 | -0.323 | 0.846,0.682,1 | 1,1,1,0.852,0. |
| ENSG000001 | DUSP18 | 1.79E-07 | 6.5862E-05 | -0.318 | 0.032,0.022,0 | 0.385,0.268,0 |
| ENSG000001 | C19orf12 | 0.00073644 | 0.04752408 | -0.29 | 0.571,0.243,0 | 0.586,1,0.488 |
| ENSG000002 | GLI4 | 0.00058465 | 0.04012557 | -0.27 | 0.3,0.68,0.86 | 0.833,0.8,1,1, |
| ENSG000001 | SNX19 | 0 | 0 | -0.26 | 0.72,0.636,0. | 1,1,1,1,1,1, |
| ENSG000001 | GPATCH11 | 2.68E-08 | 1.4315E-05 | -0.255 | 1,0.824,0.765 | 0.742,0.92,1, |
| ENSG000001 | MIR4435-2H | 6.697E-05 | 0.00779121 | -0.254 | 0.2,0.385,0.7 | 0.304,0.867,1 |
| ENSG000001 | NAGK | 2.488E-05 | 0.00353174 | -0.25 | 1,0.79,0.715, | 1,1,1,1,1,1,0. |
| ENSG000002 | TCEA3 | 4.12E-10 | 4.51E-07 | -0.25 | 0.648,0.187,1 | 1,1,1,0.871,1, |
| ENSG000001 | PRPF4B | 1.6901E-05 | 0.00259376 | -0.247 | 0.516,0.959,0 | 0.739,0.929,0 |
| ENSG000002 | LINC00963 | 0.00065358 | 0.04359748 | -0.241 | 0.6,0.077,0.3 | 0.391,0.778,1 |
| ENSG000002 | WASH8P | 0.00024925 | 0.02102727 | -0.237 | 0.333,0.5,1,0. | 1,1,0.692,1,0. |
| ENSG000001 | SLFN11 | 3.195E-06 | 0.00067346 | -0.237 | 0.726,0.879,0 | 0.785,1,0.726 |
| ENSG000001 | SLFN11 | 7.54E-07 | 0.00020829 | -0.233 | 0.716,0.898,0 | 0.811,1,0.841 |
| ENSG000001 | IL17RA | 0.0002682 | 0.02213508 | -0.232 | 0.431,0.565,0 | 0.297,0.456,0 |
| ENSG000001 | ZBTB8OS | 6.73E-08 | 3.0116E-05 | -0.231 | 0.5,0.478,0.6 | 1,0.273,0.333 |
| ENSG000001 | TTC23 | 6.2207E-05 | 0.00738155 | -0.225 | 0.714,0.4,0.7 | 0.778,1,0.8,1, |
| ENSG000001 | ABCA5 | 1.6207E-05 | 0.00251048 | -0.223 | 1,0.857,0.846 | 1,1,1,1,1,1, |
| ENSG000001 | HACL1 | 0.00021188 | 0.018474 | -0.219 | 0,1,0.435,0.5 | 1,1,0.919,0.7 |
| ENSG000001 | ANXA2 | 3.83E-09 | 2.9491E-06 | -0.209 | 1,0.778,0.704 | 0.867,1,0.852 |
| ENSG000001 | ANXA2 | 3.81E-09 | 2.9491E-06 | -0.208 | 1,0.778,0.704 | 0.867,1,0.852 |
| ENSG000001 | CEP164 | 0.00012462 | 0.01249179 | -0.207 | 0.35,0.34,0.4 | 0.609,0.656,0 |
| ENSG000001 | PKIG | 3.48E-08 | 1.7752E-05 | -0.207 | 0.182,0.328,0 | 0.602,0.721,0 |
| ENSG000001 | CDK20 | 1.36E-07 | 5.159E-05 | -0.204 | 0.636,0.538,0 | 1,0.75,0.429, |
| ENSG000002 | AC090114.3 | 1.86E-07 | 6.7903E-05 | -0.203 | 0.747,0.957,0 | 0.915,0.689,0 |
| ENSG000001 | IL6 | 1.91E-11 | 2.88E-08 | -0.201 | 0.294,1,0.818 | 1,1,1,1,1,0. |
| ENSG000001 | SUGCT | 3.80E-08 | 1.9254E-05 | -0.2 | 0.5,0.4,0.647, | 0.538,0.333,1 |
| ENSG000002 | MEG3 | 0.00048652 | 0.03488797 | -0.199 | 0.818,0.824,0 | 1,1,1,1,1,1, |
| ENSG000001 | EXO5 | 6.52E-09 | 4.6381E-06 | -0.198 | 0.5,0.7,1,0.66 | 1,0.818,1,0.7 |
| ENSG000001 | ZNF568 | 1.78E-11 | 2.74E-08 | -0.194 | 0.667,0.733,0 | 1,0.889,0.923 |
| ENSG000001 | NR2C2AP | 0.00024434 | 0.02067632 | -0.19 | 0.556,0.578,0 | 0.84,0.95,0.8 |
| ENSG000001 | ABI3BP | 1.40E-08 | 8.3824E-06 | -0.19 | 0.73,0.881,0. | 1,1,1,1,0.742, |
| ENSG000000 | ZFX | 3.15E-09 | 2.5929E-06 | -0.189 | 1,0.444,1,0.5 | 1,1,0.81,1,0.8 |
| ENSG000002 | ZSWIM7 | 2.85E-10 | 3.36E-07 | -0.185 | 0.412,0.9,0.6 | 1,1,1,0.933,1, |
| ENSG000001 | MRPL55 | 1.48E-10 | 1.93E-07 | -0.185 | 0.714,0.818,1 | 1,1,0.8,1,1,1, |
| ENSG000001 | PUS3 | 9.73E-13 | 1.88E-09 | -0.185 | 1,0.562,0.76, | 1,0.789,1,1,1, |
| ENSG000001 | ZBTB8OS | 8.46E-10 | 8.18E-07 | -0.182 | 0.814,0.703,0 | 1,0.742,0.884 |
| ENSG000001 | MOSPD1 | 0.00043881 | 0.03227343 | -0.181 | 0.529,0.469,0 | 0.762,0.872,0 |

| | | | | | | |
|---|---|---|---|---|---|---|
| ENSG000000 | UFD1 | 0.00033861 | 0.02652212 | -0.178 | 0.471,0.597,0 | 0.693,0.947,0 |
| ENSG000002 | GOLGA2P1 | 3.37E-09 | 2.7183E-06 | -0.172 | 0.692,0.656,0 | 1,1,1,1,1,1,0. |
| ENSG000001 | APTX | 5.7912E-05 | 0.00697136 | -0.171 | 0.552,0.664,0 | 1,0.638,0.876 |
| ENSG000001 | OBSL1 | 2.1549E-06 | 0.00049245 | -0.169 | 0.524,0.75,0. | 0.778,1,0.789 |
| ENSG000001 | GORASP1 | 2.2267E-05 | 0.00324383 | -0.167 | 0.926,0.622,0 | 0.771,0.636,1 |
| ENSG000001 | TFDP2 | 0.00033612 | 0.02637404 | -0.165 | 1,0.783,0.753 | 0.825,0.865,0 |
| ENSG000001 | CCDC125 | 4.8898E-05 | 0.00610538 | -0.165 | 1,0.716,1,0.3 | 1,1,0.807,0.6 |
| ENSG000001 | PRR16 | 9.9446E-06 | 0.00170912 | -0.162 | 1,1,0.857,1,0. | 1,0.867,1,0.7 |
| ENSG000002 | LINC00963 | 3.7909E-05 | 0.00497369 | -0.161 | 0.833,0.507,0 | 0.65,0.917,1, |
| ENSG000000 | AK2 | 9.5585E-05 | 0.0100883 | -0.158 | 0.333,0.879,0 | 0.879,1,0.677 |
| ENSG000001 | KDM8 | 0.00075598 | 0.04826792 | -0.157 | 1,0.879,1,1,0. | 1,1,1,1,1,1,1, |
| ENSG000000 | IP6K2 | 0.0002386 | 0.02036963 | -0.157 | 0.278,0.077,0 | 0.292,0.515,0 |
| ENSG000001 | ATRIP | 6.4449E-05 | 0.00756951 | -0.157 | 0.782,0.886,0 | 1,0.945,1,1,0. |
| ENSG000001 | PDCD10 | 2.0474E-05 | 0.00304112 | -0.157 | 1,0.813,0.581 | 0.761,0.802,0 |
| ENSG000001 | ZNF561 | 0.00010848 | 0.01118894 | -0.155 | 0.789,1,0.75, | 0.862,0.714,0 |
| ENSG000001 | RPS24 | 1.15E-12 | 2.18E-09 | -0.152 | 0.606,0.484,0 | 0.859,0.83,0. |
| ENSG000000 | NDST1 | 0.00054474 | 0.03804587 | -0.151 | 0.111,0.235,0 | 0.228,0.244,0 |
| ENSG000001 | DUS2 | 1.58E-08 | 9.2803E-06 | -0.151 | 0.625,1,0.924 | 0.81,1,1,1,0.8 |
| ENSG000001 | PAQR3 | 8.4018E-06 | 0.00149652 | -0.15 | 1,0.685,1,1,0. | 0.658,0.676,1 |
| ENSG000001 | HRAS | 1.55E-07 | 5.7916E-05 | -0.149 | 1,0.547,0.885 | 1,1,0.789,1,1, |
| ENSG000001 | BTRC | 4.1473E-05 | 0.00534857 | -0.148 | 0.545,0.837,0 | 0.891,1,0.719 |
| ENSG000001 | PLEKHA8 | 2.5499E-06 | 0.00056561 | -0.148 | 0.778,0.333,1 | 1,0.75,1,0.7,1 |
| ENSG000001 | DAB2 | 3.71E-09 | 2.912E-06 | -0.148 | 1,1,0.6,1,1,1, | 0.961,0.952,0 |
| ENSG000001 | POU6F1 | 4.71E-08 | 2.2528E-05 | -0.147 | 1,0.87,0.885, | 0.698,1,1,1,1, |
| ENSG000001 | SLC25A36 | 0.00012378 | 0.01243513 | -0.146 | 0.75,0.619,1, | 0.826,0.778,1 |
| ENSG000001 | METTL2B | 5.34E-07 | 0.00016163 | -0.145 | 0.458,0.527,0 | 1,1,1,0.936,0. |
| ENSG000001 | ZFYVE19 | 0.00052257 | 0.03691025 | -0.144 | 0.541,0.487,0 | 0.744,0.868,0 |
| ENSG000000 | STPG1 | 7.9138E-05 | 0.0088718 | -0.143 | 1,1,1,0.697,0. | 1,1,1,1,1,1,0. |
| ENSG000001 | HACL1 | 0.00018901 | 0.0170154 | -0.142 | 0.529,1,0.663 | 1,1,0.91,0.81 |
| ENSG000001 | A4GALT | 3.6754E-05 | 0.00484508 | -0.14 | 0.389,0.696,1 | 0.817,1,1,1,1, |
| ENSG000001 | ABI3BP | 1.36E-08 | 8.2276E-06 | -0.138 | 0.899,0.925,0 | 1,1,1,1,0.793, |
| ENSG000000 | ARSD | 4.081E-05 | 0.00528765 | -0.134 | 0.818,0.75,0. | 0.81,1,0.913, |
| ENSG000001 | LPGAT1 | 7.4988E-06 | 0.00137697 | -0.134 | 0.412,0.484,0 | 0.862,0.75,0. |
| ENSG000001 | POSTN | 5.41E-11 | 7.62E-08 | -0.134 | 0.214,0.192,0 | 0.278,0.283,0 |
| ENSG000001 | XPO4 | 4.3255E-05 | 0.00552316 | -0.132 | 1,0.574,0.675 | 0.734,1,1,0.8 |
| ENSG000001 | APTX | 4.4773E-06 | 0.00089959 | -0.132 | 0.81,0.808,0. | 1,0.821,0.877 |
| ENSG000001 | ACP1 | 0.00065574 | 0.0437068 | -0.131 | 0.523,0.637,0 | 0.669,0.59,0. |
| ENSG000001 | SUGT1 | 1.1144E-05 | 0.00187642 | -0.131 | 0.338,0.479,0 | 0.562,0.582,0 |
| ENSG000001 | GPS1 | 1.36E-07 | 5.159E-05 | -0.131 | 0.889,1,1,0.9 | 0.917,0.875,0 |
| ENSG000001 | GPS1 | 1.23E-07 | 4.7915E-05 | -0.131 | 0.889,1,1,0.9 | 0.917,0.875,0 |
| ENSG000001 | IL6 | 7.33E-12 | 1.20E-08 | -0.131 | 0.538,1,0.833 | 1,1,1,1,1,1,0. |
| ENSG000001 | GPS1 | 1.07E-06 | 0.00027555 | -0.13 | 0.896,1,1,0.9 | 0.922,0.883,0 |
| ENSG000001 | MFSD3 | 0.0003242 | 0.02565921 | -0.129 | 0.846,0.846,0 | 0.773,1,1,1,0. |

| | | | | | | |
|---|---|---|---|---|---|---|
| ENSG000002 | THAP9-AS1 | 3.06E-07 | 0.00010309 | -0.129 | 0.937,0.917,0 | 1,0.871,0.877 |
| ENSG000001 | TMEM173 | 1.04E-08 | 6.601E-06 | -0.128 | 0.762,0.64,0.( | 0.904,0.92,0.8 |
| ENSG000000 | ADAMTS6 | 0.00028512 | 0.02325182 | -0.126 | 1,0.692,1,0.5, | 1,1,1,0.6,0.9, |
| ENSG000001 | ZBTB24 | 1.62E-08 | 9.4216E-06 | -0.125 | 0.846,1,1,1,1, | 1,1,0.733,1,1, |
| ENSG000001 | TRPT1 | 1.14E-14 | 2.72E-11 | -0.125 | 1,0.833,1,0.74 | 1,1,1,1,1,1,1, |
| ENSG000001 | HARS2 | 0.00070704 | 0.04627316 | -0.124 | 0.902,0.908,0 | 0.86,0.944,1,( |
| ENSG000001 | GAK | 0.00058296 | 0.04004306 | -0.124 | 1,1,1,0.657,0. | 1,1,1,1,1,1,1, |
| ENSG000001 | SPIN1 | 0.00039411 | 0.02972165 | -0.124 | 0.111,0.138,0 | 0.214,0.406,0 |
| ENSG000002 | LINC00963 | 0.00019853 | 0.0176631 | -0.124 | 0.879,0.532,0 | 0.674,0.939,1 |
| ENSG000001 | CD151 | 0.00017621 | 0.01612504 | -0.124 | 0.257,0.234,0 | 0.407,0.585,0 |
| ENSG000000 | P4HA2 | 2.5835E-05 | 0.0036239 | -0.124 | 0.245,0.18,0.1 | 0.35,0.386,0.3 |
| ENSG000001 | MFSD8 | 4.38E-07 | 0.00013633 | -0.124 | 0.692,1,0.778 | 1,0.704,1,1,0. |
| ENSG000001 | RPP40 | 2.27E-07 | 8.0676E-05 | -0.124 | 0.64,0.698,1, | 1,1,1,0.762,0. |
| ENSG000001 | ZDHHC17 | 8.25E-08 | 3.4822E-05 | -0.124 | 0.905,0.895,0 | 0.957,0.962,1 |
| ENSG000001 | RNF216P1 | 0.00055209 | 0.03836583 | -0.123 | 0.714,0.684,1 | 0.846,0.818,1 |
| ENSG000001 | DMAC2 | 0.00019588 | 0.01746664 | -0.123 | 0.839,0.756,0 | 0.886,0.788,0 |
| ENSG000000 | P4HA2 | 5.875E-05 | 0.00704169 | -0.123 | 0.261,0.209,0 | 0.391,0.415,0 |
| ENSG000001 | SLFN11 | 2.6934E-06 | 0.00058803 | -0.123 | 0.604,0.901,0 | 0.753,1,0.802 |
| ENSG000002 | LINC00963 | 2.371E-05 | 0.00339457 | -0.122 | 0.897,0.6,0.7 | 0.759,0.941,1 |
| ENSG000000 | ARHGEF1 | 4.3546E-06 | 0.00087706 | -0.122 | 0.807,0.578,0 | 0.827,0.899,0 |
| ENSG000001 | BIN1 | 0.00070055 | 0.04596804 | -0.121 | 0.95,0.558,0.( | 0.857,0.761,0 |
| ENSG000001 | CCDC126 | 2.0886E-05 | 0.00307598 | -0.121 | 1,0.861,0.59,( | 1,1,0.684,1,1, |
| ENSG000000 | IP6K2 | 0.00056295 | 0.03895807 | -0.12 | 0.212,0.048,0 | 0.227,0.429,0 |
| ENSG000000 | UFD1 | 9.0065E-06 | 0.00158054 | -0.119 | 0.686,0.767,0 | 0.881,0.941,0 |
| ENSG000001 | COL6A3 | 9.5085E-05 | 0.01007564 | -0.117 | 0.343,0.172,0 | 0.51,0.495,0.3 |
| ENSG000001 | APIP | 0 | 0 | -0.117 | 0.738,0.75,1,( | 1,1,1,1,1,1,1, |
| ENSG000001 | AUH | 5.593E-06 | 0.00107743 | -0.116 | 0.788,0.724,0 | 0.895,0.89,0.8 |
| ENSG000001 | SLFN11 | 3.1726E-06 | 0.00067323 | -0.116 | 0.667,0.905,0 | 0.778,1,0.8,0. |
| ENSG000002 | MEG3 | 4.92E-07 | 0.00015034 | -0.116 | 0.812,0.92,1,( | 1,1,0.871,1,0. |
| ENSG000001 | PPRC1 | 1.23E-07 | 4.7915E-05 | -0.116 | 0.8,0.73,0.84( | 0.913,1,1,0.73 |
| ENSG000001 | PHYHD1 | 0.00041669 | 0.03114178 | -0.114 | 0.693,0.621,0 | 1,1,1,0.621, |
| ENSG000001 | MRPL55 | 1.87E-09 | 1.6956E-06 | -0.114 | 0.818,0.964,1 | 1,1,0.929,1,1, |
| ENSG000001 | 2-Sep | 7.74E-09 | 5.363E-06 | -0.113 | 0.875,0.783,0 | 1,1,0.897,1,1, |
| ENSG000001 | TFDP1 | 0.00020683 | 0.01812918 | -0.112 | 0.673,0.862,0 | 0.829,0.818,0 |
| ENSG000002 | WASH8P | 0.00037727 | 0.02863359 | -0.111 | 0.6,0.882,1,0. | 1,1,0.889,1,0. |
| ENSG000002 | PRR3 | 0.00019939 | 0.01772006 | -0.111 | 0.692,1,0.176 | 0.625,0.714,1 |
| ENSG000002 | NBPF14 | 8.6173E-05 | 0.0094565 | -0.111 | 0.909,0.6,1,1, | 0.882,0.958,1 |
| ENSG000001 | SDR39U1 | 2.8428E-06 | 0.00061581 | -0.111 | 0.579,0.68,1, | 1,1,0.92,0.9,1 |
| ENSG000000 | ALDH3A2 | 4.75E-08 | 2.2569E-05 | -0.111 | 0.2,0.152,0.2, | 0.12,0.189,0.2 |
| ENSG000001 | INTS14 | 7.35E-10 | 7.28E-07 | -0.111 | 0.667,0.583,0 | 0.917,1,1,1,0. |
| ENSG000001 | ATP6V0A2 | 2.02E-10 | 2.47E-07 | -0.111 | 1,0.796,0.875 | 0.833,1,1,1,0. |
| ENSG000001 | POSTN | 0 | 0 | -0.111 | 0.027,0.023,0 | 0.271,0.248,0 |
| ENSG000001 | MLX | 5.8046E-06 | 0.00110236 | -0.11 | 0.772,0.52,0.8 | 0.807,0.872,0 |

| | | | | | | |
|---|---|---|---|---|---|---|
| ENSG000001 | GTF2H2C | 3.74E-07 | 0.00011919 | -0.11 | 0.879,0.823,0 | 0.726,1,1,1, |
| ENSG000001 | PLEKHB2 | 2.66E-07 | 9.2647E-05 | -0.11 | 0.867,0.714,1 | 0.92,0.882,1, |
| ENSG000002 | GANC | 9.94E-08 | 4.0322E-05 | -0.11 | 0.556,0.621,1 | 0.867,0.806,1 |
| ENSG000001 | UAP1L1 | 3.4746E-05 | 0.00464668 | -0.109 | 0.9,0.846,0.8 | 1,1,1,0.917,1, |
| ENSG000002 | CENPS-COR | 6.5276E-06 | 0.00122093 | -0.109 | 1,0.896,0.627 | 1,0.771,1,1,1, |
| ENSG000001 | RNF213 | 0.00037789 | 0.02865415 | -0.108 | 0.091,0.04,0. | 0.292,0.245,0 |
| ENSG000001 | SEC13 | 1.58E-05 | 0.00246112 | -0.108 | 1,0.691,0.528 | 0.651,1,1,1,1, |
| ENSG000002 | LCMT1 | 2.94E-07 | 9.9895E-05 | -0.108 | 0.939,0.782,0 | 0.8,0.864,0.9 |
| ENSG000001 | SLC7A6 | 0.00053388 | 0.0375715 | -0.107 | 1,0.739,1,1,0. | 1,1,1,0.909,1, |
| ENSG000001 | ZNF561 | 0.00029839 | 0.02414446 | -0.106 | 0.892,1,0.824 | 0.9,0.754,0.8 |
| ENSG000000 | PDPR | 8.9547E-05 | 0.00970143 | -0.106 | 0.882,0.857,0 | 0.882,1,0.8,1, |
| ENSG000001 | HACL1 | 1.8904E-05 | 0.00285379 | -0.106 | 0.714,0.857,0 | 0.944,1,1,1,0. |
| ENSG000001 | ZBTB8OS | 9.23E-11 | 1.26E-07 | -0.106 | 0.863,0.797,0 | 1,0.742,0.961 |
| ENSG000001 | PKD1P1 | 0.00023406 | 0.02011299 | -0.105 | 0.556,0.5,0.6, | 0.818,1,0.529 |
| ENSG000000 | SPEN | 9.2354E-05 | 0.00988818 | -0.105 | 0.733,0.778,0 | 1,0.882,0.935 |
| ENSG000001 | PABPC1L | 1.482E-05 | 0.00233038 | -0.105 | 0.877,0.884,0 | 0.881,0.901,1 |
| ENSG000000 | NLE1 | 1.2001E-06 | 0.00030528 | -0.105 | 0.714,0.7,0.8 | 0.857,1,1,0.6 |
| ENSG000001 | AVL9 | 9.2365E-05 | 0.00988818 | -0.104 | 0.709,0.63,0. | 1,0.908,1,0.9 |
| ENSG000001 | TXNRD3 | 4.40E-08 | 2.1288E-05 | -0.104 | 0.833,0.789,0 | 1,1,0.931,1,0. |
| ENSG000001 | MIR4435-2H | 0.00020156 | 0.0178366 | -0.103 | 0.643,0.73,0. | 0.739,0.946,1 |
| ENSG000000 | MAP2K3 | 6.734E-06 | 0.00125034 | -0.103 | 1,0.667,1,0.6 | 1,0.765,0.905 |
| ENSG000001 | MGA | 0.00044408 | 0.03254544 | -0.102 | 1,1,0.833,0.5 | 1,0.9,1,0.879, |
| ENSG000001 | CUL9 | 0.00071754 | 0.04673952 | -0.101 | 0.9,0.545,0.7 | 0.786,0.86,1, |
| ENSG000001 | CLSTN3 | 0.00070701 | 0.04627316 | -0.101 | 0.733,0.688,0 | 1,1,1,1,0.923, |
| ENSG000001 | TAPT1 | 6.3391E-06 | 0.00119029 | -0.101 | 1,1,0.636,0.8 | 1,1,1,0.882,1, |
| ENSG000001 | SLC16A4 | 1.46E-08 | 8.6583E-06 | -0.101 | 0.913,0.814,0 | 1,1,1,1,1,1,1, |
| ENSG000001 | YPEL5 | 1.1082E-05 | 0.00186984 | 0.101 | 0.029,0.025,0 | 0.034,0.012,0 |
| ENSG000001 | COL6A3 | 0.00030224 | 0.02436165 | 0.102 | 0.915,0.893,0 | 0.678,0.553,0 |
| ENSG000001 | MTO1 | 0.00024287 | 0.02059344 | 0.103 | 0.917,0.93,0. | 0.977,0.932,0 |
| ENSG000000 | PLXND1 | 2.1108E-06 | 0.00048672 | 0.105 | 0.968,0.85,0. | 0.603,0.729,0 |
| ENSG000001 | RBPJ | 0.00013874 | 0.01356128 | 0.108 | 0.913,1,1,1,0. | 0.741,0.808,0 |
| ENSG000001 | TRA2B | 0.00060404 | 0.04098326 | 0.109 | 0.229,0.157,0 | 0.124,0.136,0 |
| ENSG000001 | C9orf85 | 6.7088E-06 | 0.00124843 | 0.113 | 0.789,0.95,0. | 0.6,0.804,0.8 |
| ENSG000001 | SLC35E2B | 2.83E-09 | 2.3761E-06 | 0.114 | 1,1,1,1,0.943, | 0.694,0.76,0. |
| ENSG000001 | KIF3A | 2.6087E-05 | 0.00364703 | 0.115 | 1,0.959,1,0.9 | 0.904,0.762,0 |
| ENSG000001 | WARS | 1.7095E-06 | 0.00041416 | 0.116 | 0,0,0,0.398,0. | 0,0.009,0,0,0, |
| ENSG000000 | FAM13B | 2.57E-09 | 2.196E-06 | 0.116 | 0.034,0.293,0 | 0.04,0.07,0.0 |
| ENSG000001 | ADAL | 0.00012865 | 0.01283095 | 0.118 | 0.648,0.886,0 | 0.607,0.921,0 |
| ENSG000001 | EPB41 | 6.19E-07 | 0.00018005 | 0.119 | 1,1,1,0.847,1, | 0.605,0.501,0 |
| ENSG000002 | HLA-B | 3.28E-09 | 2.6763E-06 | 0.119 | 0.989,0.997,1 | 0.98,0.97,0.9 |
| ENSG000001 | SYTL3 | 0.00014983 | 0.014323 | 0.12 | 1,0.789,1,1,1, | 0.2,1,1,0.6,1, |
| ENSG000001 | LPIN3 | 1.2777E-05 | 0.0021088 | 0.12 | 1,1,1,1,1,0.77 | 1,1,1,0.857,0. |
| ENSG000002 | LCMT1 | 8.30E-08 | 3.4881E-05 | 0.12 | 0.161,0.139,0 | 0.043,0.093,0 |

| | | | | | | | |
|---|---|---|---|---|---|---|---|
| ENSG00000( | OCEL1 | 0.00042327 | 0.03152079 | | 0.123 | 0.755,0.971,0 | 0.686,0.535,0 |
| ENSG000001 | PIN4 | 0 | 0 | | 0.123 | 1,1,0.982,1,0. | 0.855,0.742,0 |
| ENSG000001 | SLC39A1 | 0 | 0 | | 0.123 | 1,1,1,1,1,1,0.9 | 0.843,0.866,0 |
| ENSG00000( | FRYL | 5.3413E-05 | 0.00654344 | | 0.124 | 0.636,0.917,0 | 0.733,0.8,0.79 |
| ENSG000001 | FRAS1 | 4.5817E-05 | 0.00577445 | | 0.126 | 0.98,0.84,1,0. | 0.633,0.852,0 |
| ENSG000001 | WARS | 5.54E-07 | 0.00016636 | | 0.127 | 0,0,0,0.387,0. | 0,0.009,0,0,0, |
| ENSG000001 | WARS | 5.28E-07 | 0.00016042 | | 0.127 | 0,0,0,0.464,0. | 0,0.006,0,0,0, |
| ENSG000001 | SENP7 | 0.00058912 | 0.04031375 | | 0.128 | 0.903,0.824,0 | 0.824,0.78,0.: |
| ENSG000001 | PREPL | 1.97E-09 | 1.7581E-06 | | 0.132 | 0.357,0.154,0 | 0.158,0.098,0 |
| ENSG00000( | COL16A1 | 0.00036451 | 0.02791623 | | 0.134 | 0.476,0.511,0 | 0.463,0.524,0 |
| ENSG000001 | PTPN13 | 0.0005885 | 0.04031375 | | 0.135 | 0.614,1,0.851 | 0.793,0.793,1 |
| ENSG000001 | URB2 | 0.00011626 | 0.0117653 | | 0.138 | 1,1,0.636,1,1, | 1,0.667,1,0.54 |
| ENSG000001 | TBC1D5 | 6.00E-07 | 0.00017691 | | 0.138 | 0.808,0.691,0 | 0.788,0.62,0.( |
| ENSG000001 | PREPL | 2.86E-11 | 4.18E-08 | | 0.138 | 0.357,0.154,0 | 0.094,0.098,0 |
| ENSG000001 | ZBTB1 | 3.523E-05 | 0.00470384 | | 0.139 | 0.194,0.19,0. | 0.1,0.212,0.0 |
| ENSG000001 | RBPJ | 2.2806E-05 | 0.0033107 | | 0.139 | 0.84,0.714,0.( | 0.658,0.554,0 |
| ENSG000001 | GUF1 | 0.00018238 | 0.0165797 | | 0.141 | 1,1,1,1,1,1,1, | 0.377,0.515,0 |
| ENSG000001 | TMEM263 | 8.2714E-05 | 0.0091494 | | 0.141 | 0.747,0.786,0 | 0.765,0.633,0 |
| ENSG000002 | AC090114.3 | 4.05E-08 | 2.0073E-05 | | 0.142 | 0.722,0.889,1 | 0.928,0.948,0 |
| ENSG000001 | ITGB3BP | 9.03E-09 | 5.964E-06 | | 0.142 | 1,0.855,1,1,1, | 0.712,0.825,0 |
| ENSG000001 | DTWD1 | 3.51E-07 | 0.00011506 | | 0.144 | 0.226,0.352,0 | 0.205,0.158,0 |
| ENSG000001 | ILF3 | 5.0372E-05 | 0.00625374 | | 0.146 | 0.498,0.492,0 | 0.154,0.252,0 |
| ENSG000001 | ZNF45 | 9.0897E-05 | 0.0097687 | | 0.147 | 0.733,1,0.6,1, | 0.714,0.667,0 |
| ENSG000002 | ATXN2 | 0.000418 | 0.03121189 | | 0.149 | 0.542,0.591,0 | 0.561,0.423,0 |
| ENSG00000( | PITPNM2 | 0.0002515 | 0.021167 | | 0.153 | 0.897,0.957,0 | 0.727,0.714,0 |
| ENSG000001 | FAM210A | 7.0385E-05 | 0.0080867 | | 0.153 | 1,0.75,1,1,1,1 | 1,0.455,0.939 |
| ENSG000002 | AC012651.1 | 6.33E-08 | 2.8476E-05 | | 0.155 | 1,1,1,1,1,1,1, | 0.672,0.619,0 |
| ENSG000001 | TBCK | 5.3299E-05 | 0.00653909 | | 0.16 | 0.898,1,1,0.9 | 0.445,0.785,0 |
| ENSG000001 | FAM210A | 5.5593E-05 | 0.00676073 | | 0.162 | 1,0.846,1,1,1, | 1,0.471,0.926 |
| ENSG000001 | PPFIBP1 | 1.30E-07 | 5.0241E-05 | | 0.165 | 0.553,0.834,0 | 0.48,0.447,0.4 |
| ENSG000001 | ASH1L | 8.6629E-05 | 0.00948159 | | 0.167 | 1,1,1,1,1,1,1, | 0.778,0.714,0 |
| ENSG000001 | LPIN3 | 4.071E-05 | 0.00528295 | | 0.167 | 1,1,1,1,1,0.77 | 1,1,1,0.74,0.8 |
| ENSG00000( | NSFL1C | 1.9752E-05 | 0.00296574 | | 0.17 | 0.571,0.373,0 | 0.227,0.281,0 |
| ENSG000002 | LINC01881 | 1.19E-07 | 4.6883E-05 | | 0.172 | 1,1,0.684,1,1, | 1,1,1,0.538,0. |
| ENSG000001 | SH3PXD2A | 0.00031567 | 0.02512853 | | 0.175 | 0.648,0.446,0 | 0.155,0.408,0 |
| ENSG000001 | ITGB3BP | 7.32E-08 | 3.1409E-05 | | 0.176 | 1,0.733,1,1,1, | 0.535,0.706,0 |
| ENSG00000( | YTHDC2 | 2.2022E-05 | 0.00322512 | | 0.177 | 0.714,0.889,0 | 0.419,0.818,0 |
| ENSG000001 | ARHGAP12 | 1.8224E-06 | 0.00043548 | | 0.179 | 0.569,0.79,0.4 | 0.548,0.339,0 |
| ENSG000001 | PREPL | 0.0006712 | 0.04438148 | | 0.182 | 0.357,0.161,0 | 0.2,0.211,0.1 |
| ENSG000001 | BTN2A2 | 5.1044E-06 | 0.00100139 | | 0.182 | 1,1,1,1,1,1,1, | 0.882,0.368,0 |
| ENSG000001 | RFX8 | 0.00019338 | 0.01729668 | | 0.183 | 0.88,0.865,0.8 | 0.676,0.834,0 |
| ENSG000001 | PREPL | 0.00047711 | 0.0343613 | | 0.185 | 0.357,0.161,0 | 0.172,0.211,0 |
| ENSG000001 | PREPL | 0.00072392 | 0.04703032 | | 0.194 | 0.294,0.185,0 | 0.238,0.364,0 |

| | | | | | | |
|---|---|---|---|---|---|---|
| ENSG000001 | ZSCAN32 | 7.6911E-06 | 0.00139686 | 0.202 | 1,1,1,0.556,1, | 0.394,1,0.545 |
| ENSG000002 | AMACR | 7.9678E-05 | 0.00889633 | 0.205 | 0.909,0.871,0 | 0.517,0.418,0 |
| ENSG000000 | ZFX | 0.00021575 | 0.01875134 | 0.208 | 1,1,1,0.795,0. | 0.727,0.455,0 |
| ENSG000002 | AC090114.3 | 2.3036E-05 | 0.0033209 | 0.21 | 0.652,0.909,1 | 0.952,0.915,0 |
| ENSG000002 | CASTOR3 | 0.00040418 | 0.03028881 | 0.214 | 0.616,1,1,0.8 | 0.86,0.746,0. |
| ENSG000001 | CLIP1 | 2.81E-07 | 9.646E-05 | 0.233 | 1,1,1,1,0.734, | 0.75,1,0.79,0. |
| ENSG000001 | BTN2A2 | 6.73E-07 | 0.00019293 | 0.234 | 1,1,1,1,1,1,1, | 0.846,0.25,0. |
| ENSG000001 | ZCCHC10 | 0.00060625 | 0.04109919 | 0.237 | 0.376,0.557,0 | 0.215,0.044,0 |
| ENSG000001 | PIN4 | 0 | 0 | 0.237 | 1,1,0.959,1,0. | 0.738,0.478,0 |
| ENSG000000 | CIRBP | 2.217E-05 | 0.00324099 | 0.244 | 0.236,0.277,0 | 0.172,0.176,0 |
| ENSG000001 | RPGRIP1L | 4.3292E-05 | 0.00552316 | 0.245 | 0.565,0.793,0 | 0.1,0.257,0.1 |
| ENSG000001 | ZNF827 | 3.447E-05 | 0.0046246 | 0.245 | 1,0.923,0.883 | 0.334,0.805,0 |
| ENSG000000 | CIRBP | 1.4665E-05 | 0.0023173 | 0.248 | 0.236,0.306,0 | 0.172,0.208,0 |
| ENSG000001 | ZNF142 | 0.00046322 | 0.03359368 | 0.254 | 0.6,1,0.615,0. | 0.312,0.362,0 |
| ENSG000001 | ZNF142 | 0.00012835 | 0.01281684 | 0.261 | 0.6,1,0.63,0.5 | 0.389,0.375,0 |
| ENSG000001 | SEPT2 | 0.00046502 | 0.0336599 | 0.275 | 1,0.613,0.9,0. | 0.097,0.22,0. |
| ENSG000001 | COPB2 | 9.7402E-06 | 0.00168093 | 0.299 | 0.875,0.8,1,1, | 0.688,0.65,0. |
| ENSG000001 | BICD1 | 0.00014236 | 0.01382913 | 0.306 | 1,0.694,0.862 | 0.221,0.74,0. |
| ENSG000001 | GSE1 | 1.2085E-05 | 0.00201851 | 0.312 | 0.846,0.83,0. | 0.677,0.429,0 |
| ENSG000001 | ENDOV | 0.00035733 | 0.02752037 | 0.314 | 0.8,1,1,0.75,1 | 0.714,1,0.556 |
| ENSG000001 | SEPT2 | 1.6962E-05 | 0.00259388 | 0.314 | 1,0.769,0.943 | 0.152,0.256,0 |
| ENSG000001 | SEPT2 | 5.4381E-06 | 0.00105443 | 0.361 | 1,0.714,0.913 | 0.176,0.158,0 |
| ENSG000002 | MIATNB | 4.86E-07 | 0.0001491 | 0.364 | 0.333,0.619,0 | 0.435,0.714,0 |
| ENSG000001 | SEPT2 | 1.4686E-06 | 0.00036575 | 0.433 | 1,0.647,0.882 | 0.125,0.059,0 |
| ENSG000002 | SNHG3 | 0 | 0 | 0.486 | 1,1,1,1,1,1,1, | 0.389,0.429,0 |
| ENSG000002 | FAM86DP | 2.27E-12 | 3.94E-09 | 0.607 | 0,1,1,1,1,1,0. | 0,0.167,0.12, |

| Chr | Strand | event_type |
|---|---|---|
| chr11 | + | SE |
| chr3 | - | SE |
| chr1 | + | SE |
| chr13 | - | SE |
| chr17 | - | SE |
| chr22 | - | SE |
| chr19 | - | SE |
| chr8 | + | SE |
| chr11 | - | SE |
| chr2 | + | SE |
| chr2 | - | SE |
| chr2 | + | SE |
| chr1 | - | SE |
| chr6 | + | SE |
| chr9 | + | SE |
| chr12 | - | SE |
| chr17 | - | SE |
| chr17 | - | SE |
| chr22 | + | SE |
| chr1 | - | SE |
| chr15 | - | SE |
| chr17 | - | SE |
| chr3 | - | SE |
| chr15 | - | SE |
| chr15 | - | SE |
| chr11 | + | SE |
| chr20 | + | SE |
| chr9 | - | SE |
| chr7 | + | SE |
| chr7 | + | SE |
| chr7 | + | SE |
| chr14 | + | SE |
| chr1 | + | SE |
| chr19 | + | SE |
| chr19 | - | SE |
| chr3 | - | SE |
| chrX | + | SE |
| chr17 | - | SE |
| chr1 | - | SE |
| chr11 | - | SE |
| chr1 | - | SE |
| chrX | - | SE |

| | | |
|---|---|---|
| chr22 | - | SE |
| chr15 | - | SE |
| chr9 | - | SE |
| chr2 | - | SE |
| chr3 | - | SE |
| chr3 | - | SE |
| chr5 | - | SE |
| chr5 | + | SE |
| chr9 | + | SE |
| chr1 | - | SE |
| chr16 | + | SE |
| chr3 | - | SE |
| chr3 | + | SE |
| chr3 | - | SE |
| chr19 | - | SE |
| chr10 | + | SE |
| chr5 | + | SE |
| chr16 | + | SE |
| chr4 | - | SE |
| chr11 | - | SE |
| chr10 | + | SE |
| chr7 | + | SE |
| chr5 | - | SE |
| chr12 | - | SE |
| chr3 | + | SE |
| chr7 | + | SE |
| chr15 | + | SE |
| chr1 | - | SE |
| chr3 | - | SE |
| chr22 | - | SE |
| chr3 | - | SE |
| chrX | - | SE |
| chr1 | - | SE |
| chr13 | - | SE |
| chr13 | - | SE |
| chr9 | - | SE |
| chr2 | + | SE |
| chr13 | + | SE |
| chr17 | + | SE |
| chr17 | + | SE |
| chr7 | + | SE |
| chr17 | + | SE |
| chr8 | + | SE |

| | | |
|---|---|---|
| chr4 | - | SE |
| chr5 | - | SE |
| chr5 | - | SE |
| chr6 | - | SE |
| chr11 | - | SE |
| chr5 | + | SE |
| chr4 | - | SE |
| chr9 | + | SE |
| chr9 | + | SE |
| chr11 | + | SE |
| chr5 | - | SE |
| chr4 | - | SE |
| chr6 | - | SE |
| chr12 | + | SE |
| chr7 | + | SE |
| chr19 | - | SE |
| chr5 | - | SE |
| chr17 | - | SE |
| chr9 | + | SE |
| chr19 | + | SE |
| chr2 | - | SE |
| chr7 | + | SE |
| chr3 | - | SE |
| chr22 | - | SE |
| chr2 | - | SE |
| chr11 | - | SE |
| chr9 | - | SE |
| chr17 | - | SE |
| chr14 | + | SE |
| chr10 | + | SE |
| chr9 | + | SE |
| chr1 | - | SE |
| chr2 | + | SE |
| chr13 | + | SE |
| chr12 | - | SE |
| chr6 | + | SE |
| chr1 | - | SE |
| chr14 | - | SE |
| chr17 | + | SE |
| chr15 | - | SE |
| chr12 | + | SE |
| chr13 | - | SE |
| chr17 | + | SE |

| | | |
|---|---|---|
| chr5 | + | SE |
| chr2 | + | SE |
| chr15 | + | SE |
| chr9 | + | SE |
| chr1 | + | SE |
| chr17 | + | SE |
| chr3 | - | SE |
| chr16 | + | SE |
| chr16 | + | SE |
| chr19 | - | SE |
| chr16 | + | SE |
| chr3 | - | SE |
| chr1 | - | SE |
| chr16 | + | SE |
| chr1 | + | SE |
| chr20 | + | SE |
| chr17 | - | SE |
| chr7 | + | SE |
| chr3 | - | SE |
| chr2 | - | SE |
| chr17 | + | SE |
| chr15 | + | SE |
| chr6 | + | SE |
| chr12 | + | SE |
| chr4 | - | SE |
| chr1 | - | SE |
| chr2 | + | SE |
| chr2 | - | SE |
| chr6 | + | SE |
| chr3 | - | SE |
| chr4 | + | SE |
| chr3 | - | SE |
| chr9 | + | SE |
| chr1 | - | SE |
| chr5 | - | SE |
| chr14 | - | SE |
| chr5 | - | SE |
| chr15 | + | SE |
| chr1 | + | SE |
| chr6 | - | SE |
| chr6 | + | SE |
| chr20 | + | SE |
| chr16 | + | SE |

| | | |
|---|---|---|
| chr19 | + | SE |
| chrX | + | SE |
| chr1 | - | SE |
| chr4 | - | SE |
| chr4 | + | SE |
| chr14 | - | SE |
| chr14 | - | SE |
| chr3 | - | SE |
| chr2 | - | SE |
| chr1 | - | SE |
| chr4 | + | SE |
| chr1 | + | SE |
| chr3 | - | SE |
| chr2 | - | SE |
| chr14 | + | SE |
| chr4 | + | SE |
| chr4 | + | SE |
| chr12 | + | SE |
| chr7 | + | SE |
| chr1 | - | SE |
| chr15 | + | SE |
| chr19 | + | SE |
| chr19 | - | SE |
| chr12 | - | SE |
| chr12 | - | SE |
| chr18 | - | SE |
| chr15 | + | SE |
| chr4 | - | SE |
| chr18 | - | SE |
| chr12 | + | SE |
| chr1 | - | SE |
| chr20 | + | SE |
| chr20 | - | SE |
| chr2 | + | SE |
| chr10 | - | SE |
| chr1 | - | SE |
| chr5 | + | SE |
| chr10 | - | SE |
| chr2 | - | SE |
| chr6 | + | SE |
| chr2 | - | SE |
| chr2 | - | SE |
| chr2 | - | SE |

| | | |
|---|---|---|
| chr16 | - | SE |
| chr5 | - | SE |
| chrX | + | SE |
| chr7 | + | SE |
| chr7 | - | SE |
| chr12 | - | SE |
| chr6 | + | SE |
| chr5 | - | SE |
| chrX | + | SE |
| chr19 | + | SE |
| chr16 | - | SE |
| chr4 | - | SE |
| chr19 | + | SE |
| chr2 | - | SE |
| chr2 | - | SE |
| chr2 | + | SE |
| chr3 | - | SE |
| chr12 | + | SE |
| chr16 | + | SE |
| chr17 | + | SE |
| chr2 | + | SE |
| chr2 | + | SE |
| chr22 | + | SE |
| chr2 | + | SE |
| chr1 | + | SE |
| chr3 | - | SE |

| ID | GeneID | geneSymbol | PValue | FDR | IncLevelDiff | PSI_1 |
|---|---|---|---|---|---|---|
| 51 | ENSG000002 | SPIN3 | 1.1433E-06 | 0.00010887 | -0.146 | 0.714,0.286,0 |
| 253 | ENSG00000( | RTEL1-TNFI | 1.21E-07 | 1.7362E-05 | -0.177 | 0.905,0.755,0 |
| 258 | ENSG000001 | SLC17A9 | 0.00377378 | 0.04740625 | 0.191 | 0.767,0.818,0 |
| 259 | ENSG000001 | SLC17A9 | 0.00051162 | 0.01112253 | 0.248 | 0.609,0.66,0.( |
| 419 | ENSG000001 | DMAC2 | 3.9944E-05 | 0.00142986 | -0.119 | 0.147,0.145,0 |
| 566 | ENSG000002 | QTRT1 | 0.00223985 | 0.03396156 | 0.119 | 0.325,0.594,0 |
| 693 | ENSG000001 | BAIAP2L2 | 0.00393487 | 0.04878625 | -0.138 | 0.567,0.317,0 |
| 808 | ENSG000001 | DDX5 | 2.66E-11 | 1.58E-08 | -0.125 | 0.388,0.325,0 |
| 856 | ENSG000001 | CDK5RAP3 | 0.00285052 | 0.04003348 | 0.107 | 0.399,0.493,0 |
| 1014 | ENSG000002 | ZSWIM7 | 0.0001278 | 0.00370996 | 0.211 | 0.769,0.692,0 |
| 1091 | ENSG000001 | SGSM2 | 0.00048229 | 0.01058147 | 0.133 | 0.506,0.435,0 |
| 1252 | ENSG00000( | TNRC6A | 0.00261495 | 0.03772666 | 0.149 | 0.952,0.885,0 |
| 1267 | ENSG000001 | CCDC159 | 0.00267733 | 0.03827858 | 0.235 | 0.92,1,0.778,( |
| 1428 | ENSG000001 | WDR73 | 0.00344435 | 0.04502765 | -0.175 | 0.185,0.215,0 |
| 1448 | ENSG000002 | AL162258.1 | 2.0844E-06 | 0.0001654 | -0.136 | 0.807,1,0.918 |
| 2009 | ENSG00000( | RHBDF1 | 4.2258E-06 | 0.00028234 | -0.104 | 0.636,0.961,1 |
| 2288 | ENSG000001 | COL4A3BP | 1.0135E-06 | 9.8479E-05 | 0.231 | 1,1,0.412,1,1, |
| 2320 | ENSG000001 | EIF3G | 8.51E-08 | 1.307E-05 | -0.168 | 0.25,0.842,0.8 |
| 2498 | ENSG000001 | GUK1 | 0.00101324 | 0.01855392 | 0.109 | 0.97,0.917,0.8 |
| 2836 | ENSG000001 | INVS | 0.00014286 | 0.00407265 | -0.118 | 0.852,0.773,0 |
| 3171 | ENSG000001 | CAPRIN2 | 0.00037376 | 0.00876592 | 0.186 | 0.706,0.558,0 |
| 3394 | ENSG000001 | PRRT3 | 0.00012918 | 0.00372735 | -0.166 | 0.5,0.714,0.8 |
| 3455 | ENSG000001 | WDR55 | 9.6391E-05 | 0.00297878 | 0.118 | 0.251,0.221,0 |
| 3605 | ENSG000001 | NKTR | 0.00119236 | 0.02129968 | 0.188 | 0.619,0.684,0 |
| 3720 | ENSG000001 | NPRL2 | 1.17E-09 | 4.64E-07 | -0.216 | 1,0.667,0.613 |
| 3799 | ENSG000001 | SOD2 | 0.00120345 | 0.02129968 | -0.105 | 0.698,0.869,0 |
| 3812 | ENSG000001 | GORASP1 | 0.00013979 | 0.00400918 | 0.106 | 0.143,0.391,0 |
| 4347 | ENSG000001 | HARS2 | 1.76E-09 | 6.28E-07 | -0.15 | 0.807,0.67,1,( |
| 4565 | ENSG000002 | NSUN5P1 | 0.00363636 | 0.04641475 | -0.116 | 0.63,0.941,0.8 |
| 4764 | ENSG000002 | HLA-C | 3.06E-08 | 5.5947E-06 | -0.264 | 1,1,1,1,1,1,0. |
| 4867 | ENSG00000( | ARFGEF1 | 4.7706E-06 | 0.00030284 | 0.134 | 0.765,0.9,1,1, |
| 4898 | ENSG000001 | DDB2 | 8.04E-07 | 8.1492E-05 | 0.109 | 0.161,0.365,0 |
| 4961 | ENSG000001 | PIDD1 | 6.9032E-06 | 0.00041083 | 0.305 | 0.514,0.724,0 |
| 4982 | ENSG000002 | EGFL8 | 0.00230776 | 0.03444273 | 0.265 | 0.565,1,0.618 |
| 5093 | ENSG000001 | MINK1 | 4.1737E-05 | 0.00148289 | -0.125 | 0.857,0.913,0 |
| 5097 | ENSG000001 | EML3 | 8.4181E-06 | 0.00046067 | 0.118 | 0.325,0.233,0 |
| 5244 | ENSG000002 | DDAH2 | 5.5664E-06 | 0.0003487 | -0.111 | 0.565,0.905,0 |
| 5286 | ENSG000001 | CDK16 | 1.6629E-05 | 0.00078388 | -0.129 | 0.895,0.949,0 |
| 5379 | ENSG000001 | CARS | 1.2114E-06 | 0.00011019 | -0.118 | 0.651,0.84,0.9 |
| 5455 | ENSG000001 | JUP | 0.00027864 | 0.00687349 | 0.318 | 0.5,0.6,0.467, |
| 5908 | ENSG00000( | DGKA | 2.9669E-05 | 0.00113915 | -0.118 | 1,1,0.622,0.7 |
| 5950 | ENSG000001 | DDIT3 | 0.00016384 | 0.00448293 | 0.182 | 0.314,0.247,0 |

| | | | | | | |
|---|---|---|---|---|---|---|
| 5952 | ENSG000001 | DDIT3 | 5.338E-05 | 0.00178974 | -0.114 | 1,0.911,1,0.7 |
| 6049 | ENSG00000( | CCNK | 9.8518E-05 | 0.0030067 | 0.105 | 0.826,1,1,1,1, |
| 6405 | ENSG000001 | MAN2C1 | 3.1969E-06 | 0.00023061 | 0.18 | 0.4,0.36,0.37 |
| 6406 | ENSG000001 | MAN2C1 | 0.00030725 | 0.00748389 | 0.183 | 0.647,0.62,0. |

| PSI_2 | Chr | Strand | event_type |
|---|---|---|---|
| 1,0.833,0.846 | chrX | - | RI |
| 0.84,0.949,0. | chr20 | + | RI |
| 0.417,0.684,0 | chr20 | + | RI |
| 0.333,0.211,0 | chr20 | + | RI |
| 0.197,0.181,0 | chr19 | - | RI |
| 0.273,0.307,0 | chr19 | + | RI |
| 1,1,1,0.561,0. | chr22 | - | RI |
| 0.312,0.369,0 | chr17 | - | RI |
| 0.421,0.285,0 | chr17 | + | RI |
| 0.455,0.6,0.4 | chr17 | - | RI |
| 0.298,0.16,0. | chr17 | + | RI |
| 0.8,0.704,1,0. | chr16 | + | RI |
| 0.556,0.778,0 | chr19 | + | RI |
| 0.273,0.571,0 | chr15 | - | RI |
| 1,1,1,1,1,1,0. | chr1 | - | RI |
| 0.967,0.929,0 | chr16 | - | RI |
| 0.7,0.909,0.6 | chr5 | - | RI |
| 0.833,1,0.81, | chr19 | - | RI |
| 0.846,0.927,0 | chr1 | + | RI |
| 0.956,0.958,0 | chr9 | + | RI |
| 0.259,0.333,0 | chr12 | - | RI |
| 0.867,0.733,0 | chr3 | - | RI |
| 0.143,0.092,0 | chr5 | + | RI |
| 0.226,0.517,0 | chr3 | + | RI |
| 1,1,0.833,1,0. | chr3 | - | RI |
| 0.812,0.904,0 | chr6 | - | RI |
| 0.132,0.081,0 | chr3 | - | RI |
| 0.899,1,1,1,1, | chr5 | + | RI |
| 0.733,1,1,1,1, | chr7 | + | RI |
| 1,1,1,1,0.75,1 | chr6 | - | RI |
| 0.758,0.643,0 | chr8 | - | RI |
| 0.09,0.115,0. | chr11 | + | RI |
| 0.161,0.097,0 | chr11 | - | RI |
| 0.536,0.491,0 | chr6 | + | RI |
| 1,0.636,0.8,1, | chr17 | + | RI |
| 0.107,0.018,0 | chr11 | - | RI |
| 0.929,1,0.625 | chr6 | - | RI |
| 0.793,1,1,0.8 | chrX | + | RI |
| 0.919,1,1,0.9 | chr11 | - | RI |
| 0.304,0.333,0 | chr17 | - | RI |
| 1,1,1,0.882,1, | chr12 | + | RI |
| 0.224,0.207,0 | chr12 | - | RI |

| | | | |
|---|---|---|---|
| 1,1,1,1,1,1,1, | chr12 | - | RI |
| 0.75,0.871,1, | chr14 | + | RI |
| 0.047,0.105,0 | chr15 | - | RI |
| 0.364,0.179,0 | chr15 | - | RI |

| ID | GeneID | geneSymbol | PValue | FDR | IncLevelDiff | PSI_1 |
|---|---|---|---|---|---|---|
| 591 | ENSG000001 | PLCB4 | 6.2874E-05 | 0.00653284 | -0.111 | 0.058,0,0.268 |
| 1476 | ENSG00000( | GRN | 5.03E-11 | 4.12E-08 | -0.127 | 0.937,0.673,0 |
| 1554 | ENSG00000( | GSDMB | 0.00010167 | 0.00907403 | -0.162 | 0.559,1,0.718 |
| 2625 | ENSG000001 | METTL2B | 6.4806E-05 | 0.00658262 | -0.235 | 0.237,0.218,0 |
| 3861 | ENSG000001 | RPL9 | 1.11E-16 | 2.21E-13 | -0.147 | 0.76,0.671,0.' |
| 5007 | ENSG000001 | TMEM87B | 8.7988E-06 | 0.00131727 | 0.187 | 0.553,0.797,0 |
| 5076 | ENSG000001 | GOLGA4 | 0.0001432 | 0.01179734 | -0.101 | 0.721,0.814,0 |
| 5620 | ENSG000002 | NBPF9 | 2.496E-06 | 0.00049645 | 0.353 | 0,0.75,0.741,( |
| 6078 | ENSG00000( | TMEM161A | 0.00019899 | 0.0155647 | -0.139 | 0,0.864,0.778 |
| 7748 | ENSG000002 | FAM86DP | 5.67E-14 | 7.18E-11 | 0.594 | 0,1,1,1,1,0.: |

| PSI_2 | Chr | Strand | event_type |
|---|---|---|---|
| 0.398,0.215,0 | chr20 | + | MXE |
| 0.973,0.896,0 | chr17 | + | MXE |
| 1,1,1,1,1,1,0.[illegible] | chr17 | - | MXE |
| 0.511,0.516,0 | chr7 | + | MXE |
| 0.854,0.881,0 | chr4 | - | MXE |
| 0.632,0.478,0 | chr2 | + | MXE |
| 0.862,0.953,0 | chr3 | + | MXE |
| 0.429,0.2,0.5[illegible] | chr1 | - | MXE |
| 0.868,1,0.861 | chr19 | - | MXE |
| 0.111,0.333,0 | chr3 | - | MXE |

| ID | GeneID | geneSymbol | PValue | FDR | IncLevelDiff | PSI_1 |
|---|---|---|---|---|---|---|
| 1273 | ENSG000001 | NPAS1 | 0.00020173 | 0.01303031 | 0.159 | 0.421,0.219,0 |
| 1374 | ENSG000001 | MLX | 4.2734E-05 | 0.00424402 | -0.166 | 0.573,0.323,0 |
| 2256 | ENSG000001 | PTK7 | 0.000427 | 0.0218871 | -0.156 | 0.919,0.46,0.[illegible] |
| 2335 | ENSG000001 | DDX39B | 0.0002209 | 0.01354897 | 0.126 | 0.548,0.362,0 |
| 2483 | ENSG000001 | DDX41 | 0.0010357 | 0.04134977 | 0.225 | 1,0.869,1,1,0. |
| 2484 | ENSG000001 | DDX41 | 0.00092798 | 0.03840014 | 0.231 | 1,0.862,1,1,0. |
| 2631 | ENSG000001 | PHF21A | 8.42E-07 | 0.00018591 | -0.218 | 1,0.744,1,0.4[illegible] |
| 2977 | ENSG000001 | MFSD8 | 9.96E-07 | 0.00020297 | -0.122 | 0.636,1,0.625 |
| 3140 | ENSG000001 | SLC25A37 | 1.12E-07 | 4.2283E-05 | -0.103 | 0.71,0.943,0.[illegible] |
| 3284 | ENSG000001 | APTX | 2.1921E-05 | 0.00259941 | -0.141 | 0.75,0.867,0.[illegible] |

| PSI_2 | Chr | Strand | event_type |
|---|---|---|---|
| 0.185,0.101,0 | chr19 | + | A3SS |
| 0.683,0.806,0 | chr17 | + | A3SS |
| 1,1,0.694,0.8[illegible] | chr6 | + | A3SS |
| 0.26,0.493,0.[illegible] | chr6 | - | A3SS |
| 0.889,0.869,0 | chr5 | - | A3SS |
| 0.882,0.867,0 | chr5 | - | A3SS |
| 1,1,1,1,1,0.72 | chr11 | - | A3SS |
| 1,0.579,1,1,0. | chr4 | - | A3SS |
| 0.648,1,1,0.8[illegible] | chr8 | + | A3SS |
| 1,0.765,0.9,0. | chr9 | - | A3SS |

| ID | GeneID | geneSymbol | PValue | FDR | IncLevelDiff | PSI_1 |
|---|---|---|---|---|---|---|
| 802 | ENSG000001 | ZNF652 | 4.61E-09 | 1.5883E-06 | -0.178 | 0.857,0.625,0 |
| 902 | ENSG000001 | SLFN11 | 0.00047735 | 0.02282021 | -0.117 | 0.733,0.836,1 |
| 1273 | ENSG000001 | UBE2I | 4.55E-10 | 2.69E-07 | 0.254 | 1,1,1,1,1,1,1, |
| 1669 | ENSG000001 | ZBTB8OS | 3.57E-09 | 1.3175E-06 | -0.207 | 0.696,0.707,0 |
| 1682 | ENSG000001 | CAPN10 | 0.00046952 | 0.02273939 | 0.115 | 1,1,1,1,1,1,1, |
| 2096 | ENSG000001 | NDUFS5 | 3.09E-07 | 5.1449E-05 | 0.132 | 0.353,0.423,0 |
| 2861 | ENSG000001 | NCK1 | 0.00082141 | 0.03420108 | -0.161 | 0.253,0.277,0 |
| 2978 | ENSG000001 | GSKIP | 2.15E-08 | 6.5399E-06 | -0.131 | 0.565,0.417,0 |
| 3331 | ENSG000001 | ZNF75D | 1.07E-07 | 2.5126E-05 | 0.239 | 1,1,1,1,1,1,1, |
| 4468 | ENSG000001 | PARP2 | 2.89E-15 | 1.49E-11 | 0.493 | 0.438,0.556,0 |

| PSI_2 | Chr | Strand | event_type |
|---|---|---|---|
| 0.917,1,0.833 | chr17 | - | A5SS |
| 0.61,1,0.941, | chr17 | - | A5SS |
| 0.535,0.463,0 | chr16 | + | A5SS |
| 1,0.632,0.842 | chr1 | - | A5SS |
| 0.89,1,0.783, | chr2 | + | A5SS |
| 0.196,0.211,0 | chr1 | + | A5SS |
| 0.224,0.332,0 | chr3 | + | A5SS |
| 1,1,1,0.971,1, | chr14 | + | A5SS |
| 0.625,1,0.778 | chrX | - | A5SS |
| 0.411,0.556,0 | chr14 | + | A5SS |

| GeneID | geneSymbol | PValue | FDR | IncLevelDifference |
|---|---|---|---|---|
| ENSG00000168003.16 | SLC3A2 | 4.17E-12 | 6.94E-09 | -0.484 |
| ENSG00000244026.6 | FAM86DP | 0 | 0 | -0.435 |
| ENSG00000143537.13 | ADAM15 | 0 | 0 | -0.417 |
| ENSG00000133110.14 | POSTN | 8.08E-09 | 5.55228E-06 | -0.369 |
| ENSG00000179094.15 | PER1 | 1.59E-12 | 2.94E-09 | -0.323 |
| ENSG00000167065.13 | DUSP18 | 1.79E-07 | 6.5862E-05 | -0.318 |
| ENSG00000131943.17 | C19orf12 | 0.00073644 | 0.047524083 | -0.29 |
| ENSG00000250571.6 | GLI4 | 0.000584648 | 0.040125567 | -0.27 |
| ENSG00000120451.10 | SNX19 | 0 | 0 | -0.26 |
| ENSG00000152133.14 | GPATCH11 | 2.68E-08 | 1.43145E-05 | -0.255 |
| ENSG00000172965.15 | MIR4435-2HG | 6.69702E-05 | 0.007791209 | -0.254 |
| ENSG00000124357.12 | NAGK | 2.48804E-05 | 0.003531737 | -0.25 |
| ENSG00000204219.10 | TCEA3 | 4.12E-10 | 4.51E-07 | -0.25 |
| ENSG00000112739.16 | PRPF4B | 1.69005E-05 | 0.002593763 | -0.247 |
| ENSG00000204054.13 | LINC00963 | 0.00065358 | 0.043597481 | -0.241 |
| ENSG00000226210.3 | WASH8P | 0.000249247 | 0.021027272 | -0.237 |
| ENSG00000172716.16 | SLFN11 | 3.19503E-06 | 0.00067346 | -0.237 |
| ENSG00000172716.16 | SLFN11 | 7.54E-07 | 0.000208286 | -0.233 |
| ENSG00000177663.13 | IL17RA | 0.000268198 | 0.022135082 | -0.232 |
| ENSG00000176261.15 | ZBTB8OS | 6.73E-08 | 3.01164E-05 | -0.231 |
| ENSG00000103852.12 | TTC23 | 6.22066E-05 | 0.007381554 | -0.225 |
| ENSG00000154265.15 | ABCA5 | 1.6207E-05 | 0.002510484 | -0.223 |
| ENSG00000131373.14 | HACL1 | 0.000211875 | 0.018474 | -0.219 |
| ENSG00000182718.16 | ANXA2 | 3.83E-09 | 2.94911E-06 | -0.209 |
| ENSG00000182718.16 | ANXA2 | 3.81E-09 | 2.94911E-06 | -0.208 |
| ENSG00000110274.15 | CEP164 | 0.000124619 | 0.012491785 | -0.207 |
| ENSG00000168734.13 | PKIG | 3.48E-08 | 1.77525E-05 | -0.207 |
| ENSG00000156345.17 | CDK20 | 1.36E-07 | 5.15904E-05 | -0.204 |
| ENSG00000280828.1 | AC090114.3 | 1.86E-07 | 6.79032E-05 | -0.203 |
| ENSG00000136244.11 | IL6 | 1.91E-11 | 2.88E-08 | -0.201 |
| ENSG00000175600.15 | SUGCT | 3.80E-08 | 1.92537E-05 | -0.2 |
| ENSG00000214548.16 | MEG3 | 0.000486524 | 0.034887966 | -0.199 |
| ENSG00000164002.11 | EXO5 | 6.52E-09 | 4.63813E-06 | -0.198 |
| ENSG00000198453.12 | ZNF568 | 1.78E-11 | 2.74E-08 | -0.194 |
| ENSG00000184162.14 | NR2C2AP | 0.000244342 | 0.020676324 | -0.19 |
| ENSG00000154175.16 | ABI3BP | 1.40E-08 | 8.3824E-06 | -0.19 |
| ENSG00000005889.15 | ZFX | 3.15E-09 | 2.59292E-06 | -0.189 |
| ENSG00000214941.7 | ZSWIM7 | 2.85E-10 | 3.36E-07 | -0.185 |
| ENSG00000162910.18 | MRPL55 | 1.48E-10 | 1.93E-07 | -0.185 |
| ENSG00000110060.8 | PUS3 | 9.73E-13 | 1.88E-09 | -0.185 |
| ENSG00000176261.15 | ZBTB8OS | 8.46E-10 | 8.18E-07 | -0.182 |
| ENSG00000101928.12 | MOSPD1 | 0.000438812 | 0.032273428 | -0.181 |

| | | | | |
|---|---|---|---|---|
| ENSG00000070010.18 | UFD1 | 0.000338613 | 0.026522123 | -0.178 |
| ENSG00000255769.7 | GOLGA2P10 | 3.37E-09 | 2.71828E-06 | -0.172 |
| ENSG00000137074.18 | APTX | 5.79117E-05 | 0.006971362 | -0.171 |
| ENSG00000124006.14 | OBSL1 | 2.15493E-06 | 0.000492449 | -0.169 |
| ENSG00000114745.13 | GORASP1 | 2.22671E-05 | 0.003243827 | -0.167 |
| ENSG00000114126.17 | TFDP2 | 0.00033612 | 0.026374043 | -0.165 |
| ENSG00000183323.12 | CCDC125 | 4.88981E-05 | 0.006105378 | -0.165 |
| ENSG00000184838.14 | PRR16 | 9.94462E-06 | 0.001709119 | -0.162 |
| ENSG00000204054.13 | LINC00963 | 3.79087E-05 | 0.004973691 | -0.161 |
| ENSG00000004455.16 | AK2 | 9.5585E-05 | 0.010088295 | -0.158 |
| ENSG00000155666.11 | KDM8 | 0.000755981 | 0.048267922 | -0.157 |
| ENSG00000068745.14 | IP6K2 | 0.000238598 | 0.020369629 | -0.157 |
| ENSG00000164053.20 | ATRIP | 6.44486E-05 | 0.007569513 | -0.157 |
| ENSG00000114209.14 | PDCD10 | 2.04735E-05 | 0.003041121 | -0.157 |
| ENSG00000171469.10 | ZNF561 | 0.000108481 | 0.011188939 | -0.155 |
| ENSG00000138326.19 | RPS24 | 1.15E-12 | 2.18E-09 | -0.152 |
| ENSG00000070614.14 | NDST1 | 0.000544741 | 0.03804587 | -0.151 |
| ENSG00000167264.17 | DUS2 | 1.58E-08 | 9.28029E-06 | -0.151 |
| ENSG00000163291.14 | PAQR3 | 8.40177E-06 | 0.001496522 | -0.15 |
| ENSG00000174775.16 | HRAS | 1.55E-07 | 5.79161E-05 | -0.149 |
| ENSG00000166167.17 | BTRC | 4.14732E-05 | 0.005348566 | -0.148 |
| ENSG00000106086.19 | PLEKHA8 | 2.54988E-06 | 0.00056561 | -0.148 |
| ENSG00000153071.14 | DAB2 | 3.71E-09 | 2.91201E-06 | -0.148 |
| ENSG00000184271.17 | POU6F1 | 4.71E-08 | 2.25276E-05 | -0.147 |
| ENSG00000114120.12 | SLC25A36 | 0.00012378 | 0.012435125 | -0.146 |
| ENSG00000165055.15 | METTL2B | 5.34E-07 | 0.000161629 | -0.145 |
| ENSG00000166140.17 | ZFYVE19 | 0.000522572 | 0.036910248 | -0.144 |
| ENSG00000001460.17 | STPG1 | 7.91383E-05 | 0.008871804 | -0.143 |
| ENSG00000131373.14 | HACL1 | 0.00018901 | 0.017015405 | -0.142 |
| ENSG00000128274.16 | A4GALT | 3.67537E-05 | 0.004845079 | -0.14 |
| ENSG00000154175.16 | ABI3BP | 1.36E-08 | 8.22758E-06 | -0.138 |
| ENSG00000006756.15 | ARSD | 4.08102E-05 | 0.00528765 | -0.134 |
| ENSG00000123684.12 | LPGAT1 | 7.4988E-06 | 0.001376965 | -0.134 |
| ENSG00000133110.14 | POSTN | 5.41E-11 | 7.62E-08 | -0.134 |
| ENSG00000132953.16 | XPO4 | 4.32548E-05 | 0.005523155 | -0.132 |
| ENSG00000137074.18 | APTX | 4.47729E-06 | 0.000899589 | -0.132 |
| ENSG00000143727.15 | ACP1 | 0.000655744 | 0.043706804 | -0.131 |
| ENSG00000165416.14 | SUGT1 | 1.11436E-05 | 0.001876417 | -0.131 |
| ENSG00000169727.12 | GPS1 | 1.36E-07 | 5.15904E-05 | -0.131 |
| ENSG00000169727.12 | GPS1 | 1.23E-07 | 4.79154E-05 | -0.131 |
| ENSG00000136244.11 | IL6 | 7.33E-12 | 1.20E-08 | -0.131 |
| ENSG00000169727.12 | GPS1 | 1.06997E-06 | 0.000275549 | -0.13 |
| ENSG00000167700.8 | MFSD3 | 0.000324203 | 0.025659212 | -0.129 |

| | | | | |
|---|---|---|---|---|
| ENSG00000251022.6 | THAP9-AS1 | 3.06E-07 | 0.000103094 | -0.129 |
| ENSG00000184584.12 | TMEM173 | 1.04E-08 | 6.60098E-06 | -0.128 |
| ENSG00000049192.14 | ADAMTS6 | 0.00028512 | 0.023251825 | -0.126 |
| ENSG00000112365.4 | ZBTB24 | 1.62E-08 | 9.42164E-06 | -0.125 |
| ENSG00000149743.13 | TRPT1 | 1.14E-14 | 2.72E-11 | -0.125 |
| ENSG00000112855.15 | HARS2 | 0.000707042 | 0.046273161 | -0.124 |
| ENSG00000178950.16 | GAK | 0.000582964 | 0.040043055 | -0.124 |
| ENSG00000106723.16 | SPIN1 | 0.000394111 | 0.029721651 | -0.124 |
| ENSG00000204054.13 | LINC00963 | 0.000198532 | 0.017663103 | -0.124 |
| ENSG00000177697.18 | CD151 | 0.000176212 | 0.016125038 | -0.124 |
| ENSG00000072682.18 | P4HA2 | 2.58346E-05 | 0.003623895 | -0.124 |
| ENSG00000164073.10 | MFSD8 | 4.38E-07 | 0.000136325 | -0.124 |
| ENSG00000124787.13 | RPP40 | 2.27E-07 | 8.06764E-05 | -0.124 |
| ENSG00000186908.14 | ZDHHC17 | 8.25E-08 | 3.48224E-05 | -0.124 |
| ENSG00000196204.11 | RNF216P1 | 0.000552089 | 0.038365831 | -0.123 |
| ENSG00000105341.18 | DMAC2 | 0.000195885 | 0.017466644 | -0.123 |
| ENSG00000072682.18 | P4HA2 | 5.87499E-05 | 0.007041686 | -0.123 |
| ENSG00000172716.16 | SLFN11 | 2.69335E-06 | 0.000588028 | -0.123 |
| ENSG00000204054.13 | LINC00963 | 2.371E-05 | 0.00339457 | -0.122 |
| ENSG00000076928.17 | ARHGEF1 | 4.35463E-06 | 0.000877063 | -0.122 |
| ENSG00000136717.14 | BIN1 | 0.000700545 | 0.045968038 | -0.121 |
| ENSG00000169193.11 | CCDC126 | 2.08855E-05 | 0.003075976 | -0.121 |
| ENSG00000068745.14 | IP6K2 | 0.000562954 | 0.038958069 | -0.12 |
| ENSG00000070010.18 | UFD1 | 9.00646E-06 | 0.001580538 | -0.119 |
| ENSG00000163359.15 | COL6A3 | 9.50852E-05 | 0.010075644 | -0.117 |
| ENSG00000149089.12 | APIP | 0 | 0 | -0.117 |
| ENSG00000148090.11 | AUH | 5.59303E-06 | 0.001077431 | -0.116 |
| ENSG00000172716.16 | SLFN11 | 3.17263E-06 | 0.00067323 | -0.116 |
| ENSG00000214548.16 | MEG3 | 4.92E-07 | 0.000150344 | -0.116 |
| ENSG00000148840.10 | PPRC1 | 1.23E-07 | 4.79154E-05 | -0.116 |
| ENSG00000175287.18 | PHYHD1 | 0.000416686 | 0.031141783 | -0.114 |
| ENSG00000162910.18 | MRPL55 | 1.87E-09 | 1.69558E-06 | -0.114 |
| ENSG00000168385.17 | SEPT2 | 7.74E-09 | 5.36302E-06 | -0.113 |
| ENSG00000198176.12 | TFDP1 | 0.000206831 | 0.018129177 | -0.112 |
| ENSG00000226210.3 | WASH8P | 0.000377274 | 0.028633587 | -0.111 |
| ENSG00000204576.11 | PRR3 | 0.000199394 | 0.017720063 | -0.111 |
| ENSG00000270629.5 | NBPF14 | 8.61728E-05 | 0.009456496 | -0.111 |
| ENSG00000100445.17 | SDR39U1 | 2.84283E-06 | 0.000615813 | -0.111 |
| ENSG00000072210.18 | ALDH3A2 | 4.75E-08 | 2.25692E-05 | -0.111 |
| ENSG00000138614.14 | INTS14 | 7.35E-10 | 7.28E-07 | -0.111 |
| ENSG00000185344.13 | ATP6V0A2 | 2.02E-10 | 2.47E-07 | -0.111 |
| ENSG00000133110.14 | POSTN | 0 | 0 | -0.111 |
| ENSG00000108788.11 | MLX | 5.80457E-06 | 0.001102364 | -0.11 |

| | | | | |
|---|---|---|---|---|
| ENSG00000183474.15 | GTF2H2C | 3.74E-07 | 0.000119194 | -0.11 |
| ENSG00000115762.16 | PLEKHB2 | 2.66E-07 | 9.26466E-05 | -0.11 |
| ENSG00000214013.9 | GANC | 9.94E-08 | 4.0322E-05 | -0.11 |
| ENSG00000197355.10 | UAP1L1 | 3.47459E-05 | 0.00464668 | -0.109 |
| ENSG00000251503.8 | CENPS-CORT | 6.52761E-06 | 0.001220929 | -0.109 |
| ENSG00000173821.19 | RNF213 | 0.00037789 | 0.028654153 | -0.108 |
| ENSG00000157020.17 | SEC13 | 1.57996E-05 | 0.002461125 | -0.108 |
| ENSG00000205629.11 | LCMT1 | 2.94E-07 | 9.98949E-05 | -0.108 |
| ENSG00000103064.14 | SLC7A6 | 0.000533884 | 0.037571497 | -0.107 |
| ENSG00000171469.10 | ZNF561 | 0.000298388 | 0.024144458 | -0.106 |
| ENSG00000090857.13 | PDPR | 8.95475E-05 | 0.009701435 | -0.106 |
| ENSG00000131373.14 | HACL1 | 1.89036E-05 | 0.00285379 | -0.106 |
| ENSG00000176261.15 | ZBTB8OS | 9.23E-11 | 1.26E-07 | -0.106 |
| ENSG00000183889.12 | PKD1P1 | 0.000234058 | 0.020112987 | -0.105 |
| ENSG00000065526.10 | SPEN | 9.23538E-05 | 0.00988818 | -0.105 |
| ENSG00000101104.12 | PABPC1L | 1.48202E-05 | 0.002330385 | -0.105 |
| ENSG00000073536.17 | NLE1 | 1.2001E-06 | 0.000305281 | -0.105 |
| ENSG00000105778.18 | AVL9 | 9.23651E-05 | 0.00988818 | -0.104 |
| ENSG00000197763.15 | TXNRD3 | 4.40E-08 | 2.12877E-05 | -0.104 |
| ENSG00000172965.15 | MIR4435-2HG | 0.000201563 | 0.017836597 | -0.103 |
| ENSG00000034152.18 | MAP2K3 | 6.73403E-06 | 0.001250335 | -0.103 |
| ENSG00000174197.16 | MGA | 0.000444075 | 0.032545439 | -0.102 |
| ENSG00000112659.13 | CUL9 | 0.000717539 | 0.04673952 | -0.101 |
| ENSG00000139182.14 | CLSTN3 | 0.000707007 | 0.046273161 | -0.101 |
| ENSG00000169762.16 | TAPT1 | 6.33911E-06 | 0.001190293 | -0.101 |
| ENSG00000168679.17 | SLC16A4 | 1.46E-08 | 8.6583E-06 | -0.101 |
| ENSG00000119801.12 | YPEL5 | 1.10821E-05 | 0.001869842 | 0.101 |
| ENSG00000163359.15 | COL6A3 | 0.000302244 | 0.024361649 | 0.102 |
| ENSG00000135297.15 | MTO1 | 0.000242867 | 0.020593437 | 0.103 |
| ENSG00000004399.12 | PLXND1 | 2.11077E-06 | 0.000486718 | 0.105 |
| ENSG00000168214.20 | RBPJ | 0.00013874 | 0.01356128 | 0.108 |
| ENSG00000136527.17 | TRA2B | 0.000604043 | 0.040983264 | 0.109 |
| ENSG00000155621.14 | C9orf85 | 6.70877E-06 | 0.001248431 | 0.113 |
| ENSG00000189339.11 | SLC35E2B | 2.83E-09 | 2.37613E-06 | 0.114 |
| ENSG00000131437.15 | KIF3A | 2.60872E-05 | 0.003647028 | 0.115 |
| ENSG00000140105.17 | WARS | 1.7095E-06 | 0.000414161 | 0.116 |
| ENSG00000031003.10 | FAM13B | 2.57E-09 | 2.19598E-06 | 0.116 |
| ENSG00000168803.15 | ADAL | 0.000128646 | 0.012830951 | 0.118 |
| ENSG00000159023.21 | EPB41 | 6.19E-07 | 0.000180049 | 0.119 |
| ENSG00000234745.10 | HLA-B | 3.28E-09 | 2.67631E-06 | 0.119 |
| ENSG00000164674.15 | SYTL3 | 0.000149827 | 0.014322995 | 0.12 |
| ENSG00000132793.11 | LPIN3 | 1.27772E-05 | 0.002108799 | 0.12 |
| ENSG00000205629.11 | LCMT1 | 8.30E-08 | 3.48806E-05 | 0.12 |

| | | | | |
|---|---|---|---|---|
| ENSG00000099330.8 | OCEL1 | 0.000423273 | 0.031520793 | 0.123 |
| ENSG00000102309.12 | PIN4 | 0 | 0 | 0.123 |
| ENSG00000143570.17 | SLC39A1 | 0 | 0 | 0.123 |
| ENSG00000075539.14 | FRYL | 5.34129E-05 | 0.006543437 | 0.124 |
| ENSG00000138759.18 | FRAS1 | 4.58169E-05 | 0.00577445 | 0.126 |
| ENSG00000140105.17 | WARS | 5.54E-07 | 0.000166358 | 0.127 |
| ENSG00000140105.17 | WARS | 5.28E-07 | 0.00016042 | 0.127 |
| ENSG00000138468.15 | SENP7 | 0.000589123 | 0.040313748 | 0.128 |
| ENSG00000138078.15 | PREPL | 1.97E-09 | 1.75805E-06 | 0.132 |
| ENSG00000084636.17 | COL16A1 | 0.000364514 | 0.02791623 | 0.134 |
| ENSG00000163629.12 | PTPN13 | 0.000588502 | 0.040313748 | 0.135 |
| ENSG00000135763.9 | URB2 | 0.000116264 | 0.011765295 | 0.138 |
| ENSG00000131374.14 | TBC1D5 | 6.00E-07 | 0.000176907 | 0.138 |
| ENSG00000138078.15 | PREPL | 2.86E-11 | 4.18E-08 | 0.138 |
| ENSG00000126804.13 | ZBTB1 | 3.52299E-05 | 0.004703842 | 0.139 |
| ENSG00000168214.20 | RBPJ | 2.28058E-05 | 0.003310702 | 0.139 |
| ENSG00000151806.13 | GUF1 | 0.000182376 | 0.016579696 | 0.141 |
| ENSG00000151135.9 | TMEM263 | 8.27144E-05 | 0.009149401 | 0.141 |
| ENSG00000280828.1 | AC090114.3 | 4.05E-08 | 2.00731E-05 | 0.142 |
| ENSG00000142856.16 | ITGB3BP | 9.03E-09 | 5.96395E-06 | 0.142 |
| ENSG00000104047.14 | DTWD1 | 3.51E-07 | 0.000115059 | 0.144 |
| ENSG00000129351.17 | ILF3 | 5.03716E-05 | 0.006253745 | 0.146 |
| ENSG00000124459.11 | ZNF45 | 9.08968E-05 | 0.009768705 | 0.147 |
| ENSG00000204842.15 | ATXN2 | 0.000418 | 0.031211885 | 0.149 |
| ENSG00000090975.12 | PITPNM2 | 0.000251497 | 0.021166998 | 0.153 |
| ENSG00000177150.12 | FAM210A | 7.0385E-05 | 0.008086696 | 0.153 |
| ENSG00000258461.5 | AC012651.1 | 6.33E-08 | 2.84761E-05 | 0.155 |
| ENSG00000145348.16 | TBCK | 5.32988E-05 | 0.006539086 | 0.16 |
| ENSG00000177150.12 | FAM210A | 5.55931E-05 | 0.006760735 | 0.162 |
| ENSG00000110841.13 | PPFIBP1 | 1.30E-07 | 5.02405E-05 | 0.165 |
| ENSG00000116539.12 | ASH1L | 8.66294E-05 | 0.009481587 | 0.167 |
| ENSG00000132793.11 | LPIN3 | 4.07104E-05 | 0.005282954 | 0.167 |
| ENSG00000088833.17 | NSFL1C | 1.97521E-05 | 0.002965741 | 0.17 |
| ENSG00000220804.8 | LINC01881 | 1.19E-07 | 4.68834E-05 | 0.172 |
| ENSG00000107957.16 | SH3PXD2A | 0.000315673 | 0.025128531 | 0.175 |
| ENSG00000142856.16 | ITGB3BP | 7.32E-08 | 3.14088E-05 | 0.176 |
| ENSG00000047188.15 | YTHDC2 | 2.20224E-05 | 0.00322512 | 0.177 |
| ENSG00000165322.17 | ARHGAP12 | 1.82243E-06 | 0.000435478 | 0.179 |
| ENSG00000138078.15 | PREPL | 0.000671202 | 0.044381483 | 0.182 |
| ENSG00000124508.16 | BTN2A2 | 5.10436E-06 | 0.001001393 | 0.182 |
| ENSG00000196460.13 | RFX8 | 0.000193382 | 0.017296678 | 0.183 |
| ENSG00000138078.15 | PREPL | 0.000477114 | 0.034361305 | 0.185 |
| ENSG00000138078.15 | PREPL | 0.000723922 | 0.047030324 | 0.194 |

| | | | | |
|---|---|---|---|---|
| ENSG00000140987.19 | ZSCAN32 | 7.69111E-06 | 0.00139686 | 0.202 |
| ENSG00000242110.7 | AMACR | 7.96779E-05 | 0.008896326 | 0.205 |
| ENSG00000005889.15 | ZFX | 0.000215749 | 0.01875134 | 0.208 |
| ENSG00000280828.1 | AC090114.3 | 2.30357E-05 | 0.003320898 | 0.21 |
| ENSG00000239521.8 | CASTOR3 | 0.000404181 | 0.030288812 | 0.214 |
| ENSG00000130779.20 | CLIP1 | 2.81E-07 | 9.646E-05 | 0.233 |
| ENSG00000124508.16 | BTN2A2 | 6.73E-07 | 0.000192926 | 0.234 |
| ENSG00000155329.11 | ZCCHC10 | 0.000606245 | 0.041099192 | 0.237 |
| ENSG00000102309.12 | PIN4 | 0 | 0 | 0.237 |
| ENSG00000099622.13 | CIRBP | 2.21697E-05 | 0.00324099 | 0.244 |
| ENSG00000103494.13 | RPGRIP1L | 4.32918E-05 | 0.005523155 | 0.245 |
| ENSG00000151612.15 | ZNF827 | 3.44697E-05 | 0.004624604 | 0.245 |
| ENSG00000099622.13 | CIRBP | 1.46652E-05 | 0.002317296 | 0.248 |
| ENSG00000115568.15 | ZNF142 | 0.000463225 | 0.033593683 | 0.254 |
| ENSG00000115568.15 | ZNF142 | 0.00012835 | 0.012816844 | 0.261 |
| ENSG00000168385.17 | SEPT2 | 0.000465016 | 0.033659902 | 0.275 |
| ENSG00000184432.9 | COPB2 | 9.74018E-06 | 0.001680928 | 0.299 |
| ENSG00000151746.13 | BICD1 | 0.000142359 | 0.013829127 | 0.306 |
| ENSG00000131149.18 | GSE1 | 1.20846E-05 | 0.002018512 | 0.312 |
| ENSG00000173818.16 | ENDOV | 0.000357329 | 0.02752037 | 0.314 |
| ENSG00000168385.17 | SEPT2 | 1.69624E-05 | 0.00259388 | 0.314 |
| ENSG00000168385.17 | SEPT2 | 5.4381E-06 | 0.001054434 | 0.361 |
| ENSG00000244625.5 | MIATNB | 4.86E-07 | 0.000149103 | 0.364 |
| ENSG00000168385.17 | SEPT2 | 1.4686E-06 | 0.000365751 | 0.433 |
| ENSG00000242125.3 | SNHG3 | 0 | 0 | 0.486 |
| ENSG00000244026.6 | FAM86DP | 2.27E-12 | 3.94E-09 | 0.607 |
| ENSG00000204271.12 | SPIN3 | 1.14333E-06 | 0.000108868 | -0.146 |
| ENSG00000026036.22 | RTEL1-TNFRSF6[illegible] | 1.21E-07 | 1.73621E-05 | -0.177 |
| ENSG00000101194.17 | SLC17A9 | 0.003773781 | 0.047406254 | 0.191 |
| ENSG00000101194.17 | SLC17A9 | 0.000511622 | 0.011122526 | 0.248 |
| ENSG00000105341.18 | DMAC2 | 3.99437E-05 | 0.001429864 | -0.119 |
| ENSG00000213339.8 | QTRT1 | 0.002239851 | 0.033961561 | 0.119 |
| ENSG00000128298.16 | BAIAP2L2 | 0.003934871 | 0.048786247 | -0.138 |
| ENSG00000108654.14 | DDX5 | 2.66E-11 | 1.58E-08 | -0.125 |
| ENSG00000108465.14 | CDK5RAP3 | 0.002850525 | 0.040033477 | 0.107 |
| ENSG00000214941.7 | ZSWIM7 | 0.000127795 | 0.003709955 | 0.211 |
| ENSG00000141258.12 | SGSM2 | 0.000482289 | 0.010581469 | 0.133 |
| ENSG00000090905.18 | TNRC6A | 0.002614955 | 0.037726663 | 0.149 |
| ENSG00000183401.11 | CCDC159 | 0.00267733 | 0.038278584 | 0.235 |
| ENSG00000177082.12 | WDR73 | 0.003444347 | 0.045027649 | -0.175 |
| ENSG00000271853.5 | AL162258.1 | 2.08442E-06 | 0.000165398 | -0.136 |
| ENSG00000007384.15 | RHBDF1 | 4.22583E-06 | 0.000282342 | -0.104 |
| ENSG00000113163.16 | COL4A3BP | 1.01355E-06 | 9.84794E-05 | 0.231 |

| | | | | |
|---|---|---|---|---|
| ENSG00000130811.11 | EIF3G | 8.51E-08 | 1.307E-05 | -0.168 |
| ENSG00000143774.16 | GUK1 | 0.001013236 | 0.018553916 | 0.109 |
| ENSG00000119509.12 | INVS | 0.000142855 | 0.004072653 | -0.118 |
| ENSG00000110888.17 | CAPRIN2 | 0.000373762 | 0.008765922 | 0.186 |
| ENSG00000163704.11 | PRRT3 | 0.000129177 | 0.003727352 | -0.166 |
| ENSG00000120314.18 | WDR55 | 9.63911E-05 | 0.002978783 | 0.118 |
| ENSG00000114857.17 | NKTR | 0.001192358 | 0.021299676 | 0.188 |
| ENSG00000114388.12 | NPRL2 | 1.17E-09 | 4.64E-07 | -0.216 |
| ENSG00000112096.17 | SOD2 | 0.001203447 | 0.021299676 | -0.105 |
| ENSG00000114745.13 | GORASP1 | 0.000139787 | 0.00400918 | 0.106 |
| ENSG00000112855.15 | HARS2 | 1.76E-09 | 6.28E-07 | -0.15 |
| ENSG00000223705.9 | NSUN5P1 | 0.003636358 | 0.046414753 | -0.116 |
| ENSG00000204525.16 | HLA-C | 3.06E-08 | 5.59467E-06 | -0.264 |
| ENSG00000066777.8 | ARFGEF1 | 4.77063E-06 | 0.00030284 | 0.134 |
| ENSG00000134574.11 | DDB2 | 8.04E-07 | 8.14918E-05 | 0.109 |
| ENSG00000177595.17 | PIDD1 | 6.90321E-06 | 0.000410827 | 0.305 |
| ENSG00000241404.6 | EGFL8 | 0.002307757 | 0.03444273 | 0.265 |
| ENSG00000141503.15 | MINK1 | 4.17365E-05 | 0.001482893 | -0.125 |
| ENSG00000149499.11 | EML3 | 8.41807E-06 | 0.000460671 | 0.118 |
| ENSG00000213722.8 | DDAH2 | 5.56636E-06 | 0.000348703 | -0.111 |
| ENSG00000102225.15 | CDK16 | 1.66292E-05 | 0.000783877 | -0.129 |
| ENSG00000110619.17 | CARS | 1.2114E-06 | 0.000110189 | -0.118 |
| ENSG00000173801.16 | JUP | 0.000278635 | 0.006873488 | 0.318 |
| ENSG00000065357.19 | DGKA | 2.96692E-05 | 0.001139153 | -0.118 |
| ENSG00000175197.12 | DDIT3 | 0.000163838 | 0.004482934 | 0.182 |
| ENSG00000175197.12 | DDIT3 | 5.33801E-05 | 0.001789738 | -0.114 |
| ENSG00000090061.17 | CCNK | 9.85182E-05 | 0.003006699 | 0.105 |
| ENSG00000140400.16 | MAN2C1 | 3.19692E-06 | 0.000230614 | 0.18 |
| ENSG00000140400.16 | MAN2C1 | 0.000307245 | 0.007483891 | 0.183 |
| ENSG00000101333.16 | PLCB4 | 6.28744E-05 | 0.006532838 | -0.111 |
| ENSG00000030582.17 | GRN | 5.03E-11 | 4.12E-08 | -0.127 |
| ENSG00000073605.18 | GSDMB | 0.00010167 | 0.009074033 | -0.162 |
| ENSG00000165055.15 | METTL2B | 6.48061E-05 | 0.006582617 | -0.235 |
| ENSG00000163682.16 | RPL9 | 1.11E-16 | 2.21E-13 | -0.147 |
| ENSG00000153214.10 | TMEM87B | 8.79885E-06 | 0.001317273 | 0.187 |
| ENSG00000144674.16 | GOLGA4 | 0.000143198 | 0.011797336 | -0.101 |
| ENSG00000269713.7 | NBPF9 | 2.49597E-06 | 0.000496449 | 0.353 |
| ENSG00000064545.14 | TMEM161A | 0.000198988 | 0.015564695 | -0.139 |
| ENSG00000244026.6 | FAM86DP | 5.67E-14 | 7.18E-11 | 0.594 |
| ENSG00000130751.9 | NPAS1 | 0.000201728 | 0.013030312 | 0.159 |
| ENSG00000108788.11 | MLX | 4.2734E-05 | 0.004244018 | -0.166 |
| ENSG00000112655.15 | PTK7 | 0.000426998 | 0.021887099 | -0.156 |
| ENSG00000198563.13 | DDX39B | 0.000220898 | 0.013548966 | 0.126 |

| | | | | |
|---|---|---|---|---|
| ENSG00000183258.11 | DDX41 | 0.001035696 | 0.041349766 | 0.225 |
| ENSG00000183258.11 | DDX41 | 0.000927983 | 0.038400142 | 0.231 |
| ENSG00000135365.15 | PHF21A | 8.42E-07 | 0.00018591 | -0.218 |
| ENSG00000164073.10 | MFSD8 | 9.96E-07 | 0.000202968 | -0.122 |
| ENSG00000147454.13 | SLC25A37 | 1.12E-07 | 4.22827E-05 | -0.103 |
| ENSG00000137074.18 | APTX | 2.19208E-05 | 0.002599409 | -0.141 |
| ENSG00000198740.8 | ZNF652 | 4.61E-09 | 1.5883E-06 | -0.178 |
| ENSG00000172716.16 | SLFN11 | 0.000477355 | 0.022820211 | -0.117 |
| ENSG00000103275.19 | UBE2I | 4.55E-10 | 2.69E-07 | 0.254 |
| ENSG00000176261.15 | ZBTB8OS | 3.57E-09 | 1.31755E-06 | -0.207 |
| ENSG00000142330.19 | CAPN10 | 0.000469518 | 0.02273939 | 0.115 |
| ENSG00000168653.10 | NDUFS5 | 3.09E-07 | 5.14486E-05 | 0.132 |
| ENSG00000158092.6 | NCK1 | 0.000821409 | 0.03420108 | -0.161 |
| ENSG00000100744.14 | GSKIP | 2.15E-08 | 6.53988E-06 | -0.131 |
| ENSG00000186376.14 | ZNF75D | 1.07E-07 | 2.51261E-05 | 0.239 |
| ENSG00000129484.13 | PARP2 | 2.89E-15 | 1.49E-11 | 0.493 |

**PSI_1**

0.163,0.328,0.509,0,0,1,0.414,0.293,0.451,0,0
0.03,0,0,0,0,0,0.105,0.03,0.074,0.083,0.042
0.339,0.52,0.494,1,0.982,0.92,0.425,0.278,0.438,0.407,0.541
0.149,0.181,0.172,0.311,0.386,0.45,0.318,0.31,0.283,0.121,0.049
0.846,0.682,1,0.1,0.103,0.127,0.818,0.857,0.862,0.556,0.931
0.032,0.022,0.111,0.286,0.385,0.429,0,0,0.148,0.214,0.455
0.571,0.243,0.391,0.231,0.733,1,0.412,0.25,0.38,0.156,0.333
0.3,0.68,0.862,0.538,0.852,0.667,0.714,0.429,0.366,0,0.25
0.72,0.636,0.742,1,1,1,0.721,0.636,0.74,0.505,0.43
1,0.824,0.765,0.375,0.667,0.304,0.684,0.714,0.837,0.758,0.5
0.2,0.385,0.778,0.412,0.5,0.143,1,1,0.653,1,0.714
1,0.79,0.715,0.834,0.557,1,1,0,0.774,0.334,1
0.648,0.187,1,0.936,0.804,0.598,1,0.551,0.896,0.479,0.839
0.516,0.959,0.96,0.569,0.464,0.578,0.22,0.693,0.719,0.452,0.85
0.6,0.077,0.368,0.5,0.793,0.727,0.6,0,0.276,0.25,0.625
0.333,0.5,1,0.478,0.385,0.333,1,0.5,0.69,0.714,0.733
0.726,0.879,0.57,0.768,0.57,1,0.249,0.249,0.846,0.525,0.665
0.716,0.898,0.694,0.847,0.602,1,0.202,0.558,0.879,0.558,0.602
0.431,0.565,0.386,0.448,0.421,0.653,0.458,0.405,0.559,0.292,0.398
0.5,0.478,0.636,0.6,0.6,0.647,0.778,0.6,0.538,0.44,0.714
0.714,0.4,0.714,0.556,0.412,1,0.231,1,0.756,0.9,0.5
1,0.857,0.846,0.125,0.385,0.556,1,1,1,0.778,1
0,1,0.435,0.562,0.672,1,1,0.339,0.84,1,1
1,0.778,0.704,1,1,0.333,0.538,0.538,0.702,0.867,0.455
1,0.778,0.704,1,1,0.333,0.538,0.538,0.706,0.867,0.455
0.35,0.34,0.41,0.296,0.255,0.513,0.512,0.33,0.524,0.386,0.709
0.182,0.328,0.315,0.299,0.376,0.462,0.409,0.31,0.255,0.3,0.567
0.636,0.538,0.583,0.75,0,1,0.571,1,0.774,1,0.273
0.747,0.957,0.611,0.741,0.665,0.38,0.576,0.605,0.665,0.846,0.728
0.294,1,0.818,0.711,0.746,0.707,1,0.5,0.9,1,0.943
0.5,0.4,0.647,0.667,0.565,0.833,0.867,0.692,0.735,0.778,0.429
0.818,0.824,0.7,1,0.429,1,1,1,0.916,1,0
0.5,0.7,1,0.667,0.474,1,0.692,0.4,0.628,0.769,0.364
0.667,0.733,0.875,0.917,0.846,0.882,0.8,0.429,0.734,0.652,1
0.556,0.578,0.586,0.871,0.846,1,0.931,0.286,0.89,0.553,1
0.73,0.881,0.703,0.67,0.844,1,0.329,1,0.833,0.859,0.628
1,0.444,1,0.571,0.667,0.667,1,1,0.761,0.429,1
0.412,0.9,0.625,0.667,0.857,1,1,1,0.731,0.438,0.619
0.714,0.818,1,0.833,1,1,0.579,0.556,0.582,0.5,0.444
1,0.562,0.76,1,0.871,0.714,0.852,0.389,0.714,0.765,1
0.814,0.703,0.78,0.884,0.891,0.56,0.793,0.814,0.723,0.539,0.877
0.529,0.469,0.389,0.628,0.759,0.524,0.72,0.803,0.769,0.574,0.783

0.471,0.597,0.795,0.826,0.658,0.349,0.738,0.744,0.624,0.671,0.829
0.692,0.656,0.768,0.726,1,0,1,0.9,0.812,1,0.76
0.552,0.664,0.696,0.585,1,0.659,0.565,0.679,0.771,0.597,0.653
0.524,0.75,0.655,0.818,0.812,0.6,0.923,1,0.803,0.6,0.474
0.926,0.622,0.4,0.905,0.786,0.778,0.839,0.778,0.804,0.689,0.526
1,0.783,0.753,0.575,0.844,0.575,0.759,0.119,0.879,1,1
1,0.716,1,0.321,0.557,0.883,1,1,0.55,0.468,1
1,1,0.857,1,0.778,0.385,0.852,0.333,0.811,0.75,0.778
0.833,0.507,0.692,0.773,0.857,0.793,0.81,0.667,0.705,0.591,0.85
0.333,0.879,0.44,1,1,1,0.688,1,0.792,0.652,0.692
1,0.879,1,1,0.58,0.609,0.7,0.851,0.869,1,0.784
0.278,0.077,0.365,0.154,0.103,0.333,0.128,0.057,0.241,0.15,0.6
0.782,0.886,0.795,0.795,0.815,0.356,1,1,0.922,0.782,0.721
1,0.813,0.581,0.79,0.912,1,0.41,0.464,0.685,0.776,0.607
0.789,1,0.75,0.778,0.814,0.727,0.536,0.769,0.847,0.592,0.622
0.606,0.484,0.613,0.839,0.863,0.892,0.802,0.613,0.795,0.671,0.755
0.111,0.235,0.143,0.135,0.333,0,0.183,0.286,0.202,0.133,0.1
0.625,1,0.924,0.893,1,0.526,0.932,0.921,0.893,0.787,0.419
1,0.685,1,1,0.794,0.325,0.562,1,0.899,0.312,0.719
1,0.547,0.885,0.858,0.562,1,1,0.622,0.881,0.794,0.832
0.545,0.837,0.865,1,0.683,0.473,0.86,0.891,0.913,0.649,0.625
0.778,0.333,1,1,1,0.667,1,0.429,0.615,0.636,1
1,1,0.6,1,1,1,0.556,1,0.874,0.462,0.488
1,0.87,0.885,1,0.755,1,0.738,0.339,0.88,0.658,0.606
0.75,0.619,1,0.533,0.778,0.714,0.667,1,0.83,0.833,0.5
0.458,0.527,0.353,0.92,1,1,1,1,0.994,0.942,1
0.541,0.487,0.604,0.75,0.887,1,0.882,0.714,0.771,0.672,0.706
1,1,1,0.697,0.837,0.833,0.533,0.412,0.484,1,1
0.529,1,0.663,0.797,0.584,1,1,0.123,0.829,1,1
0.389,0.696,1,0.875,1,0.761,1,1,0.96,0.56,1
0.899,0.925,0.754,0.748,0.857,1,0.438,1,0.858,0.931,0.71
0.818,0.75,0.667,0.565,0.81,1,0.8,1,0.687,0.862,0.579
0.412,0.484,0.913,1,1,1,0.455,0.636,0.874,1,0.818
0.214,0.192,0.202,0.179,0.172,0.184,0.275,0.249,0.256,0.23,0.212
1,0.574,0.675,0.645,0.861,0.7,1,1,0.959,0.74,0.861
0.81,0.808,0.841,0.866,1,0.718,0.619,0.683,0.819,0.647,0.76
0.523,0.637,0.53,0.502,0.684,0.401,0.371,0.266,0.306,0.563,0.351
0.338,0.479,0.402,0.331,0.349,0.413,0.373,0.394,0.33,0.384,0.275
0.889,1,1,0.913,0.263,1,0.652,1,0.873,0.714,0.895
0.889,1,1,0.913,0.263,1,0.652,1,0.872,0.714,0.889
0.538,1,0.833,0.824,0.835,0.786,1,0.714,0.942,1,0.964
0.896,1,1,0.902,0.279,1,0.637,1,0.878,0.73,0.907
0.846,0.846,0.878,0.938,0.75,0.4,0.829,0.75,0.902,0.915,0.897

0.937,0.917,0.657,0.705,0.827,0.838,0.714,0.908,0.925,0.92,0.6
0.762,0.64,0.655,0.733,0.791,0.731,0.926,0.855,0.921,0.698,0.701
1,0.692,1,0.5,0.6,1,1,0.455,0.952,0.517,1
0.846,1,1,1,0.4,0.512,0.76,0.927,0.826,0.867
1,0.833,1,0.745,1,1,0.754,0.716,0.627,1,0.945
0.902,0.908,0.71,0.5,0.647,0.571,0.784,1,0.889,0.754,0.811
1,1,1,0.657,0.615,0.364,1,1,1,1,1
0.111,0.138,0.093,0.302,0.184,0.293,0.14,0.275,0.162,0.165,0.303
0.879,0.532,0.793,0.808,0.885,0.818,0.86,0.778,0.746,0.617,0.867
0.257,0.234,0.228,0.383,0.414,0.341,0.349,0.389,0.308,0.312,0.49
0.245,0.18,0.216,0.453,0.384,0.392,0.248,0.27,0.25,0.249,0.318
0.692,1,0.778,1,0.704,0.778,1,0.333,0.908,0.789,1
0.64,0.698,1,1,0.703,1,0.71,0.695,0.885,0.727,0.922
0.905,0.895,0.806,0.925,0.927,0.571,0.704,0.721,0.941,0.771,1
0.714,0.684,1,0.778,1,1,1,0.25,0.808,0.538,1
0.839,0.756,0.766,1,0.88,1,0.67,0.488,0.64,0.72,0.798
0.261,0.209,0.253,0.476,0.418,0.401,0.286,0.296,0.283,0.289,0.357
0.604,0.901,0.767,0.82,0.89,1,0.767,0.717,0.901,0.616,0.687
0.897,0.6,0.778,0.867,0.915,0.854,0.893,0.692,0.805,0.705,0.914
0.807,0.578,0.818,0.792,0.873,0.797,0.722,0.658,0.754,0.598,0.744
0.95,0.558,0.623,0.685,0.67,0.611,0.706,0.778,0.694,0.59,0.607
1,0.861,0.59,0.866,0.764,1,0.575,0.565,0.817,0.704,1
0.212,0.048,0.245,0.043,0.037,0.333,0.081,0.057,0.15,0.261,0.455
0.686,0.767,0.829,0.864,0.769,0.632,0.833,0.788,0.777,0.736,0.837
0.343,0.172,0.32,0.457,0.506,0.567,0.261,0.18,0.295,0.136,0.34
0.738,0.75,1,0.714,0.776,1,0.885,1,0.855,1,1
0.788,0.724,0.9,0.661,0.764,0.954,0.901,0.772,0.897,0.788,0.871
0.667,0.905,0.765,0.818,0.889,1,0.778,0.704,0.901,0.647,0.684
0.812,0.92,1,0.795,0.938,0.923,0.846,0.867,0.769,0.707,0.5
0.8,0.73,0.846,0.556,0.647,1,0.957,1,0.88,0.882,1
0.693,0.621,0.86,0.732,0.868,1,1,0.45,0.966,1,1
0.818,0.964,1,0.915,1,1,0.863,0.75,0.876,0.695,0.671
0.875,0.783,0.655,0.939,0.714,0.8,1,1,0.931,0.933,1
0.673,0.862,0.717,0.603,0.727,0.538,0.779,0.625,0.749,0.804,0.735
0.6,0.882,1,0.613,0.714,0.579,1,0.818,0.86,0.833,0.826
0.692,1,0.176,1,0.733,0.6,1,0.429,0.655,0.667,0.444
0.909,0.6,1,1,0.826,1,0.643,0.739,0.867,0.488,0.907
0.579,0.68,1,1,0.895,1,1,1,0.908,0.6,0.9
0.2,0.152,0.2,0.111,0.097,0.082,0.039,0.091,0.076,0.091,0.292
0.667,0.583,0.882,1,1,1,0.905,1,0.871,0.643,0.92
1,0.796,0.875,0.923,0.931,1,0.684,0.771,0.881,0.659,0.684
0.027,0.023,0.014,0.037,0.046,0.039,0.04,0.049,0.039,0.023,0.012
0.772,0.52,0.868,0.691,0.816,0.89,0.803,0.694,0.805,0.544,0.802

0.879,0.823,0.799,0.749,0.57,0.768,1,1,0.855,0.823,0.909
0.867,0.714,1,0.636,1,1,1,0.818,0.856,0.615,0.714
0.556,0.621,1,0.857,0.846,1,0.867,0.929,0.864,0.814,0.793
0.9,0.846,0.897,1,0.81,0.333,0.975,0.758,0.884,1,0.952
1,0.896,0.627,0.599,0.789,1,0.749,0.839,0.872,0.599,1
0.091,0.04,0.167,0.096,0.059,0.111,0.175,0.323,0.282,0.191,0.195
1,0.691,0.528,0.599,0.857,1,0.789,0.599,0.853,0.839,1
0.939,0.782,0.799,0.787,0.885,0.793,0.787,0.73,0.834,0.891,0.765
1,0.739,1,1,0.667,0.273,1,1,0.968,0.947,1
0.892,1,0.824,0.833,0.877,0.8,0.644,0.829,0.897,0.672,0.725
0.882,0.857,0.647,0.905,1,0.778,0.846,0.733,0.88,0.818,0.833
0.714,0.857,0.833,1,0.733,0.6,1,1,0.956,1,0.864
0.863,0.797,0.884,0.923,0.936,0.768,0.863,0.913,0.828,0.718,0.92
0.556,0.5,0.6,1,0.667,1,1,1,0.929,0.385,1
0.733,0.778,0.727,0.913,0.882,1,0.721,0.789,0.875,0.739,0.895
0.877,0.884,0.926,1,0.774,0.533,1,0.696,0.836,0.932,0.632
0.714,0.7,0.818,1,0.455,1,1,1,0.907,0.583,0.739
0.709,0.63,0.93,1,0.694,0.847,0.901,1,0.875,0.933,0.765
0.833,0.789,0.806,0.882,0.926,0.857,0.907,0.7,0.93,0.961,0.898
0.643,0.73,0.903,0.684,0.819,0.72,1,1,0.892,1,0.907
1,0.667,1,0.667,0.879,1,0.765,0.538,0.739,1,0.846
1,1,0.833,0.583,0.882,0.765,1,0.667,0.94,0.867,1
0.9,0.545,0.714,0.875,1,1,0.943,0.368,0.904,0.833,1
0.733,0.688,0.83,0.857,0.862,0.857,1,1,0.973,1,0.867
1,1,0.636,0.889,0.833,0.833,1,1,0.762,0.619,1
0.913,0.814,0.695,0.778,0.826,1,1,1,0.92,0.871,0.913
0.029,0.025,0.034,0.106,0.042,0.042,0.414,0.446,0.377,0.064,0.016
0.915,0.893,0.912,0.817,0.812,0.736,0.94,0.871,0.929,0.917,0.942
0.917,0.93,0.95,1,1,1,0.933,1,0.93,0.836,0.903
0.968,0.85,0.934,0.899,0.899,1,0.943,1,0.955,0.943,0.893
0.913,1,1,1,0.852,1,1,1,0.852,0.951,0.909
0.229,0.157,0.208,0.418,0.399,0.468,0.08,0.16,0.085,0.082,0.054
0.789,0.95,0.943,1,0.953,0.944,1,1,0.933,1,0.913
1,1,1,1,0.943,1,1,1,0.968,0.857,1
1,0.959,1,0.956,0.921,0.963,0.834,1,0.964,0.873,1
0,0,0,0.398,0.358,0.407,0.277,0.37,0.259,0.179,0.24
0.034,0.293,0.132,0.214,0.184,0.236,0.301,0.172,0.141,0.238,0.053
0.648,0.886,0.824,1,0.898,0.525,0.84,0.913,0.889,1,0.892
1,1,1,0.847,1,0.71,1,1,0.926,0.889,1
0.989,0.997,1,1,1,0.991,0.969,0.973,0.975,1,1
1,0.789,1,1,1,0.742,0.615,0.538,0.565,0.75
1,1,1,1,1,0.778,1,0.6,0.985,1,1
0.161,0.139,0.164,0.115,0.267,0.4,0.2,0.184,0.129,0.053,0.014

0.755,0.971,0.833,0.8,0.887,1,0.745,0.889,0.815,0.89,0.906
1,1,0.982,1,0.97,0.981,1,1,0.998,0.849,0.854
1,1,1,1,1,1,0.992,0.977,0.999,1,1
0.636,0.917,0.829,1,0.875,1,1,1,0.912,1,0.789
0.98,0.84,1,0.907,0.891,1,0.885,1,0.945,0.736,0.86
0,0,0,0.387,0.376,0.429,0.263,0.339,0.247,0.164,0.252
0,0,0,0.464,0.373,0.439,0.404,0.408,0.357,0.29,0.313
0.903,0.824,0.897,1,1,0.87,1,0.571,0.842,0.897,0.8
0.357,0.154,0.14,0.205,0.385,0.302,0.222,0.238,0.139,0.206,0.186
0.476,0.511,0.475,0.533,0.526,0.707,0.465,0.53,0.501,0.561,0.411
0.614,1,0.851,0.741,0.939,1,1,1,0.868,0.827,1
1,1,0.636,1,1,0.733,1,1,0.981,1,1
0.808,0.691,0.835,0.719,0.777,0.79,0.737,0.849,0.818,0.888,0.869
0.357,0.154,0.149,0.195,0.404,0.302,0.192,0.238,0.144,0.167,0.2
0.194,0.19,0.196,0.625,0.246,0.447,0.13,0.304,0.104,0.086,0.227
0.84,0.714,0.684,1,1,0.733,0.833,0.949,0.886,0.76,0.778
1,1,1,1,1,1,1,1,1,1,1
0.747,0.786,0.667,0.738,0.803,0.812,0.653,0.828,0.669,0.641,0.665
0.722,0.889,1,1,1,0.946,1,0.918,0.989,0.76,0.954
1,0.855,1,1,1,0.779,0.909,1,0.94,1,0.943
0.226,0.352,0.487,0.481,0.395,0.405,0.264,0.171,0.217,0.316,0.166
0.498,0.492,0.401,0.237,0.407,0.651,0.257,0.475,0.321,0.396,0.273
0.733,1,0.6,1,0.667,1,0.692,1,0.797,1,1
0.542,0.591,0.579,0.638,0.565,0.551,0.582,0.844,0.665,0.73,0.779
0.897,0.957,0.886,0.895,0.818,1,0.769,1,0.883,1,0.8
1,0.75,1,1,1,1,1,1,0.966,1,1
1,1,1,1,1,1,1,1,0.994,0.865,1
0.898,1,1,0.948,0.986,1,1,1,0.97,1,0.941
1,0.846,1,1,1,1,1,1,0.963,1,1
0.553,0.834,0.463,0.528,0.669,0.576,0.582,0.716,0.559,0.684,0.575
1,1,1,1,1,1,1,1,0.936,1,1
1,1,1,1,1,0.773,1,0.46,0.973,1,1
0.571,0.373,0.377,0.393,0.662,0.518,0.347,0.469,0.471,0.448,0.279
1,1,0.684,1,1,1,1,1,0.903,0.8,1
0.648,0.446,0.714,0.408,0.408,0.256,0.588,0.256,0.539,0.535,0.326
1,0.733,1,1,1,0.6,0.714,1,0.914,1,0.846
0.714,0.889,0.913,1,0.895,1,1,0.769,0.894,1,1
0.569,0.79,0.498,0.692,0.533,0.54,0.543,0.711,0.625,0.733,0.656
0.357,0.161,0.371,0.538,0.507,0.704,0.417,0.625,0.256,0.243,0.36
1,1,1,1,1,1,1,1,0.94,1,0.917
0.88,0.865,0.87,1,0.785,1,0.901,0.904,0.823,0.941,0.776
0.357,0.161,0.371,0.538,0.507,0.704,0.417,0.625,0.258,0.243,0.36
0.294,0.185,0.419,0.538,0.593,0.514,0.556,0.417,0.311,0.417,0.692

1,1,1,0.556,1,1,0.917,1,0.896,1,0.882
0.909,0.871,0.832,0.742,0.792,1,0.823,0.95,0.862,0.816,0.931
1,1,1,0.795,0.818,0.875,1,1,0.817,0.455,1
0.652,0.909,1,1,1,0.849,1,0.843,0.985,0.657,0.849
0.616,1,1,0.874,0.74,1,0.728,1,0.753,1,1
1,1,1,1,0.734,1,1,0.871,0.909,0.892,1
1,1,1,1,1,1,1,1,0.891,1,0.867
0.376,0.557,0.256,0.287,0.445,0.231,0.546,0.751,0.583,0.546,0.311
1,1,0.959,1,0.943,0.958,1,1,0.996,0.714,0.704
0.236,0.277,0.522,0.662,0.78,0.73,0.208,0.467,0.268,0.381,0.161
0.565,0.793,0.526,0.702,0.591,0.321,0.455,0.314,0.234,0.154,0.375
1,0.923,0.883,1,1,1,1,1,0.96,0.913,1
0.236,0.306,0.522,0.667,0.783,0.73,0.208,0.467,0.297,0.409,0.188
0.6,1,0.615,0.412,0.895,0.765,0.647,0.333,0.68,0.8,0.714
0.6,1,0.63,0.524,0.905,0.81,0.739,0.4,0.747,0.833,0.714
1,0.613,0.9,0.429,1,0.778,0.478,0.5,0.574,0.188,0.185
0.875,0.8,1,1,0.5,1,1,1,0.634,0.765,1
1,0.694,0.862,1,1,1,1,1,0.875,1,1
0.846,0.83,0.944,1,0.913,0.714,1,0.905,0.809,0.857,0.786
0.8,1,1,0.75,1,0.895,1,1,0.813,0.8,1
1,0.769,0.943,0.543,1,0.833,0.556,0.556,0.703,0.366,0.312
1,0.714,0.913,0.467,1,0.8,0.294,0.385,0.62,0.316,0.185
0.333,0.619,0.517,0.455,0.6,0.857,0.592,0.857,0.631,0.538,0.625
1,0.647,0.882,0.333,1,0.6,0.333,0.273,0.509,0.257,0.185
1,1,1,1,1,1,1,1,1,1,1
0,1,1,1,1,1,0.333,0.778,0.433,0.273,0.81
0.714,0.286,0.895,0.524,1,0.8,1,0.5,0.77,1,0.556
0.905,0.755,0.821,0.455,0.6,0.714,0.806,0.895,0.865,0.947,0.727
0.767,0.818,0.919,1,1,1,0.525,0.862,0.799,0.885,0.784
0.609,0.66,0.61,0.667,0.571,1,0.213,0.556,0.398,0.492,0.321
0.147,0.145,0.102,0.067,0.16,0.057,0.159,0.294,0.166,0.118,0.182
0.325,0.594,0.526,0.46,0.429,0.405,0.271,0.44,0.246,0.471,0.231
0.567,0.317,0.468,1,1,1,0.684,0.379,0.4,0.902,1
0.388,0.325,0.378,0.179,0.203,0.207,0.181,0.253,0.266,0.269,0.187
0.399,0.493,0.382,0.377,0.322,0.417,0.134,0.459,0.223,0.391,0.137
0.769,0.692,0.705,0.8,0.63,0.571,0.767,0.684,0.504,0.686,0.72
0.506,0.435,0.222,0.353,0.236,0.353,0.192,0.201,0.175,0.371,0.045
0.952,0.885,0.854,0.889,0.944,1,1,1,0.837,0.946,0.889
0.92,1,0.778,0.622,1,0.75,0.44,0.5,0.774,0.667,0.806
0.185,0.215,0.379,0.059,0.217,0.053,0.31,0.125,0.207,0.269,0.04
0.807,1,0.918,0.844,0.702,0.466,1,0.724,0.891,0.733,0.825
0.636,0.961,1,1,1,1,0.882,0.571,0.881,0.879,0.867
1,1,0.412,1,1,1,0.833,0.333,0.89,1,1

0.25,0.842,0.8,0.833,0.875,1,0.68,0.385,0.701,0.857,1
0.97,0.917,0.867,0.82,0.971,1,0.915,0.963,0.865,0.967,0.911
0.852,0.773,0.6,0.8,0.724,0.81,0.579,0.733,0.57,0.818,0.55
0.706,0.558,0.47,0.487,0.541,0.371,0.208,0.852,0.403,0.552,0.385
0.5,0.714,0.833,0.429,0.273,1,0.778,1,0.811,1,0.818
0.251,0.221,0.347,0.432,0.281,0.151,0.124,0.199,0.188,0.175,0.155
0.619,0.684,0.731,0.487,0.667,0.786,0.36,0.657,0.41,0.762,0.111
1,0.667,0.613,0.538,1,0.2,0.8,1,0.797,0.333,0.714
0.698,0.869,0.958,0.691,0.901,0.839,0.631,0.391,0.621,0.739,0.754
0.143,0.391,0.364,0.333,0.2,0.3,0.137,0.185,0.123,0.193,0.128
0.807,0.67,1,0.945,1,0.742,0.473,0.84,0.963,0.624,1
0.63,0.941,0.811,1,0.769,1,0.5,0.778,0.625,0.647,0.733
1,1,1,1,1,1,0.113,0.037,0.122,0.75,0.714
0.765,0.9,1,1,0.913,0.75,1,1,0.985,0.636,0.846
0.161,0.365,0.253,0.383,0.236,0.127,0.123,0,0.123,0.333,0.059
0.514,0.724,0.793,0.895,0.667,0.818,0.286,0.474,0.412,0.333,0.091
0.565,1,0.618,0.464,0.698,0.296,0.574,0.776,0.597,1,0.536
0.857,0.913,0.892,0.8,0.857,0.111,0.818,1,0.688,1,1
0.325,0.233,0.324,0.204,0.12,0.467,0.111,0.179,0.138,0.319,0.109
0.565,0.905,0.789,0.818,0.667,0.556,0.75,1,0.812,0.684,1
0.895,0.949,0.92,0.714,0.81,0.3,0.647,0.733,0.927,0.8,0.875
0.651,0.84,0.905,0.782,0.694,0.843,0.8,0.857,0.959,0.875,0.767
0.5,0.6,0.467,0.636,0.684,1,1,0.556,0.458,0.8,0.429
1,1,0.622,0.778,0.643,0.6,1,1,0.953,0.739,1
0.314,0.247,0.168,0.543,0.415,0.671,0.255,0.481,0.223,0.692,0.233
1,0.911,1,0.778,0.641,0.657,0.778,1,0.984,1,1
0.826,1,1,1,1,1,0.879,1,0.907,0.846,1
0.4,0.36,0.372,0.436,0.381,0.462,0.045,0.105,0.1,0.513,0.315
0.647,0.62,0.575,0.457,0.429,0.509,0.089,0.561,0.33,0.491,0.222
0.058,0,0.268,0.091,0.069,0.092,0.089,0.035,0.247,0.357,0.174
0.937,0.673,0.727,0.873,1,1,0.954,0.837,0.86,0.851,0.673
0.559,1,0.718,0.851,0.92,0.718,0.388,0.458,0.851,1,1
0.237,0.218,0.306,0.199,0.257,0.23,0.257,0.502,0.323,0.262,0.62
0.76,0.671,0.729,0.658,0.622,0.592,0.894,0.9,0.869,0.702,0.678
0.553,0.797,0.598,0.63,0.465,0.606,0.545,0.487,0.58,0.751,0.432
0.721,0.814,0.717,0.671,0.906,0.865,0.808,0.858,0.785,0.808,0.688
0,0.75,0.741,0.333,0.571,1,0.167,0.571,0.643,1,0.778
0,0.864,0.778,0.766,0.732,1,0.724,1,0.851,1,0.793
0,1,1,1,1,1,0.385,0.833,0.582,0.538,0.812
0.421,0.219,0.289,0.143,0.481,0.312,0.237,0.386,0.166,0.244,0.538
0.573,0.323,0.755,0.505,0.667,0.783,0.657,0.56,0.693,0.373,0.627
0.919,0.46,0.763,0.57,1,0,0.752,0.202,0.667,0.828,0.752
0.548,0.362,0.542,0.567,0.631,0.488,0.598,0.589,0.498,0.513,0.587

1,0.869,1,1,0.59,1,0.913,1,0.852,1,1

1,0.862,1,1,0.6,1,0.905,1,0.855,1,1

1,0.744,1,0.414,0.91,0.726,0.613,0.76,0.836,1,0.261

0.636,1,0.625,1,0.636,0.778,1,0.391,0.877,0.778,1

0.71,0.943,0.883,0.902,1,0.208,1,0.589,0.833,1,0.913

0.75,0.867,0.742,0.862,1,0.692,0.536,0.692,0.799,0.649,0.765

0.857,0.625,0.6,0.4,0.889,1,0.688,0.923,0.738,0.75,1

0.733,0.836,1,1,0.556,1,0.538,0.778,0.888,0.84,1

1,1,1,1,1,1,1,1,1,1,1

0.696,0.707,0.696,0.792,0.859,0.768,0.832,0.604,0.676,0.495,0.821

1,1,1,1,1,1,1,1,0.93,0.947,1

0.353,0.423,0.384,0.424,0.432,0.399,0.461,0.396,0.412,0.415,0.422

0.253,0.277,0.151,0.303,0.13,0.034,0,0,0.199,0.204,0.061

0.565,0.417,0.558,1,1,1,1,0.959,1,1,1

1,1,1,1,1,1,1,1,0.873,1,1

0.438,0.556,0.541,1,0.962,1,0.627,0.259,0.55,0.788,0.643

| PSI_2 | Chr | Strand | event_type |
|---|---|---|---|
| 0.823,0.919,1,0.341,0.675,1,0.806,0.675,0.7 | chr11 | + | SE |
| 0.4,0.75,0.349,0.562,0.421,0.28,0.354,0.524,0.569 | chr3 | - | SE |
| 0.995,1,0.995,1,0.988,1,0.992,0.982,0.991 | chr1 | + | SE |
| 0.978,0.865,0.84,0.604,0.64,0.665,0.314,0.327,0.323 | chr13 | - | SE |
| 1,1,1,0.852,0.727,1,0.957,1,1 | chr17 | - | SE |
| 0.385,0.268,0.526,0.36,0.667,0.231,0.793,0.6,0.739 | chr22 | - | SE |
| 0.586,1,0.488,0.75,0.467,0.636,0.68,0.846,1 | chr19 | - | SE |
| 0.833,0.8,1,1,0.778,1,0.478,0.667,0.5 | chr8 | + | SE |
| 1,1,1,1,1,1,1,1,0.989 | chr11 | - | SE |
| 0.742,0.92,1,1,1,1,0.833,0.879,1 | chr2 | + | SE |
| 0.304,0.867,1,1,0.867,1,0.8,1,1 | chr2 | - | SE |
| 1,1,1,1,1,1,0.801,1,1 | chr2 | + | SE |
| 1,1,1,0.871,1,1,0.961,0.959,0.954 | chr1 | - | SE |
| 0.739,0.929,0.859,1,0.9,1,0.731,0.88,0.894 | chr6 | + | SE |
| 0.391,0.778,1,0.667,1,0,0.52,1,0.75 | chr9 | + | SE |
| 1,1,0.692,1,0.714,1,0.583,0.846,0.75 | chr12 | - | SE |
| 0.785,1,0.726,0.623,1,1,1,1,0.768 | chr17 | - | SE |
| 0.811,1,0.841,0.78,1,1,1,1,0.847 | chr17 | - | SE |
| 0.297,0.456,0.349,1,1,1,0.683,0.714,0.691 | chr22 | + | SE |
| 1,0.273,0.333,1,1,1,1,1,0.818 | chr1 | - | SE |
| 0.778,1,0.8,1,0.889,1,0.875,0.81,0.75 | chr15 | - | SE |
| 1,1,1,1,1,1,1,1,1 | chr17 | - | SE |
| 1,1,0.919,0.755,1,1,1,0.719,1 | chr3 | - | SE |
| 0.867,1,0.852,1,1,1,0.636,1,1 | chr15 | - | SE |
| 0.867,1,0.852,1,1,1,0.636,1,1 | chr15 | - | SE |
| 0.609,0.656,0.761,0.537,0.683,0.428,0.705,0.669,0.6 | chr11 | + | SE |
| 0.602,0.721,0.651,0.483,0.495,0.595,0.49,0.583,0.355 | chr20 | + | SE |
| 1,0.75,0.429,1,1,1,0.667,0.818,1 | chr9 | - | SE |
| 0.915,0.689,0.869,0.952,0.964,0.849,0.905,1,0.836 | chr7 | + | SE |
| 1,1,1,1,1,1,0.933,0.926,1 | chr7 | + | SE |
| 0.538,0.333,1,1,1,1,1,1,0.75 | chr7 | + | SE |
| 1,1,1,1,1,1,1,0.895,1 | chr14 | + | SE |
| 1,0.818,1,0.714,1,0.5,0.875,1,0.765 | chr1 | + | SE |
| 1,0.889,0.923,1,1,1,0.913,1,1 | chr19 | + | SE |
| 0.84,0.95,0.86,0.905,0.895,1,0.889,1,1 | chr19 | - | SE |
| 1,1,1,1,0.742,1,0.969,0.933,1 | chr3 | - | SE |
| 1,1,0.81,1,0.875,1,1,1,1 | chrX | + | SE |
| 1,1,1,0.933,1,0.667,1,0.818,1 | chr17 | - | SE |
| 1,1,0.8,1,1,1,0.429,1,1 | chr1 | - | SE |
| 1,0.789,1,1,1,1,0.938,1,1 | chr11 | - | SE |
| 1,0.742,0.884,1,1,1,1,1,0.868 | chr1 | - | SE |
| 0.762,0.872,0.849,0.667,0.463,1,0.815,1,0.886 | chrX | - | SE |

| | | | |
|---|---|---|---|
| 0.693,0.947,0.736,0.953,1,0.778,0.857,0.84,0.77 | chr22 | - | SE |
| 1,1,1,1,1,1,0.346,1,1 | chr15 | - | SE |
| 1,0.638,0.876,0.87,0.779,1,0.934,0.779,0.738 | chr9 | - | SE |
| 0.778,1,0.789,1,0.867,1,1,0.882,0.714 | chr2 | - | SE |
| 0.771,0.636,1,1,0.939,1,0.957,1,0.786 | chr3 | - | SE |
| 0.825,0.865,0.859,0.815,1,1,1,1,0.904 | chr3 | - | SE |
| 1,1,0.807,0.626,1,1,1,1,1 | chr5 | - | SE |
| 1,0.867,1,0.714,1,1,0.867,1,1 | chr5 | + | SE |
| 0.65,0.917,1,0.917,1,0.875,0.769,1,0.926 | chr9 | + | SE |
| 0.879,1,0.677,0.8,1,1,1,1,1 | chr1 | - | SE |
| 1,1,1,1,1,1,1,1,1 | chr16 | + | SE |
| 0.292,0.515,0.263,0.366,0.429,0.154,0.551,0.45,0.429 | chr3 | - | SE |
| 1,0.945,1,1,0.847,1,1,0.869,1 | chr3 | + | SE |
| 0.761,0.802,0.874,0.679,0.874,1,1,1,1 | chr3 | - | SE |
| 0.862,0.714,0.829,0.8,1,1,0.963,0.959,1 | chr19 | - | SE |
| 0.859,0.83,0.841,0.85,0.866,0.84,0.925,0.922,0.928 | chr10 | + | SE |
| 0.228,0.244,0.433,0.2,0.263,0.417,0.364,0.333,0.4 | chr5 | + | SE |
| 0.81,1,1,1,0.847,1,1,1,1 | chr16 | + | SE |
| 0.658,0.676,1,1,1,1,1,1,0.806 | chr4 | - | SE |
| 1,1,0.789,1,1,1,1,1,0.898 | chr11 | - | SE |
| 0.891,1,0.719,0.885,1,0.782,0.95,0.925,1 | chr10 | + | SE |
| 1,0.75,1,0.7,1,0.8,1,1,1 | chr7 | + | SE |
| 0.961,0.952,0.951,1,1,0.818,1,1,1 | chr5 | - | SE |
| 0.698,1,1,1,1,1,0.928,0.837,1 | chr12 | - | SE |
| 0.826,0.778,1,1,0.76,0.882,1,0.8,1 | chr3 | + | SE |
| 1,1,1,0.936,0.926,1,1,1,0.964 | chr7 | + | SE |
| 0.744,0.868,0.722,0.821,0.872,0.957,0.939,0.964,0.964 | chr15 | + | SE |
| 1,1,1,1,1,1,0.857,0.83,0.8 | chr1 | - | SE |
| 1,1,0.91,0.818,1,1,1,0.529,1 | chr3 | - | SE |
| 0.817,1,1,1,1,1,1,1,1 | chr22 | - | SE |
| 1,1,1,1,0.793,1,0.974,0.938,1 | chr3 | - | SE |
| 0.81,1,0.913,1,1,1,0.941,0.765,0.76 | chrX | - | SE |
| 0.862,0.75,0.826,1,1,1,0.8,1,1 | chr1 | - | SE |
| 0.278,0.283,0.304,0.466,0.486,0.471,0.284,0.28,0.291 | chr13 | - | SE |
| 0.734,1,1,0.898,1,1,0.933,1,1 | chr13 | - | SE |
| 1,0.821,0.877,0.913,0.86,1,0.952,0.913,0.868 | chr9 | - | SE |
| 0.669,0.59,0.681,0.555,0.584,0.487,0.562,0.65,0.604 | chr2 | + | SE |
| 0.562,0.582,0.501,0.478,0.461,0.595,0.431,0.538,0.363 | chr13 | + | SE |
| 0.917,0.875,0.909,1,1,1,1,1,1 | chr17 | + | SE |
| 0.917,0.875,0.909,1,1,1,1,1,1 | chr17 | + | SE |
| 1,1,1,1,1,1,0.946,0.951,1 | chr7 | + | SE |
| 0.922,0.883,0.915,1,1,1,1,1,1 | chr17 | + | SE |
| 0.773,1,1,1,0.935,1,0.885,0.955,0.937 | chr8 | + | SE |

| | | | |
|---|---|---|---|
| 1,0.871,0.877,0.871,1,1,0.863,1,1 | chr4 | - | SE |
| 0.904,0.92,0.852,0.828,0.891,0.892,0.881,0.954,0.911 | chr5 | - | SE |
| 1,1,1,0.6,0.9,1,1,0.765,1 | chr5 | - | SE |
| 1,1,0.733,1,1,1,0.941,0.931,1 | chr6 | - | SE |
| 1,1,1,1,1,1,1,1,1 | chr11 | - | SE |
| 0.86,0.944,1,0.895,0.724,1,0.872,0.867,0.885 | chr5 | + | SE |
| 1,1,1,1,1,1,1,1,1 | chr4 | - | SE |
| 0.214,0.406,0.153,0.286,0.405,0.5,0.288,0.344,0.288 | chr9 | + | SE |
| 0.674,0.939,1,0.931,1,0.882,0.778,1,0.933 | chr9 | + | SE |
| 0.407,0.585,0.462,0.382,0.535,0.448,0.395,0.446,0.491 | chr11 | + | SE |
| 0.35,0.386,0.386,0.565,0.562,0.494,0.344,0.357,0.297 | chr5 | - | SE |
| 1,0.704,1,1,0.875,1,1,1,0.882 | chr4 | - | SE |
| 1,1,1,0.762,0.904,1,1,0.794,1 | chr6 | - | SE |
| 0.957,0.962,1,0.918,1,0.923,0.971,0.882,1 | chr12 | + | SE |
| 0.846,0.818,1,0.778,1,1,1,0.846,1 | chr7 | + | SE |
| 0.886,0.788,0.934,1,1,1,0.884,0.848,0.772 | chr19 | - | SE |
| 0.391,0.415,0.433,0.598,0.592,0.514,0.366,0.357,0.325 | chr5 | - | SE |
| 0.753,1,0.802,0.753,1,1,1,1,0.896 | chr17 | - | SE |
| 0.759,0.941,1,0.962,1,0.938,0.842,1,0.958 | chr9 | + | SE |
| 0.827,0.899,0.914,0.951,0.852,0.969,0.719,0.754,0.871 | chr19 | + | SE |
| 0.857,0.761,0.855,0.951,0.854,0.538,0.778,0.814,0.796 | chr2 | - | SE |
| 1,1,0.684,1,1,1,0.907,1,0.654 | chr7 | + | SE |
| 0.227,0.429,0.188,0.212,0.368,0.083,0.5,0.29,0.36 | chr3 | - | SE |
| 0.881,0.941,0.876,0.968,1,0.833,0.86,0.855,0.83 | chr22 | - | SE |
| 0.51,0.495,0.536,0.434,0.367,0.369,0.405,0.451,0.409 | chr2 | - | SE |
| 1,1,1,1,1,1,1,1,1 | chr11 | - | SE |
| 0.895,0.89,0.84,1,0.802,1,1,1,1 | chr9 | - | SE |
| 0.778,1,0.8,0.75,1,1,1,1,0.882 | chr17 | - | SE |
| 1,1,0.871,1,0.889,1,0.886,1,0.826 | chr14 | + | SE |
| 0.913,1,1,0.739,1,1,1,1,1 | chr10 | + | SE |
| 1,1,1,1,0.621,1,1,0.925,1 | chr9 | + | SE |
| 1,1,0.929,1,1,1,0.915,1,1 | chr1 | - | SE |
| 1,1,0.897,1,1,1,1,1,1 | chr2 | + | SE |
| 0.829,0.818,0.834,0.843,0.872,0.867,0.806,0.754,0.781 | chr13 | + | SE |
| 1,1,0.889,1,0.889,1,0.655,0.846,0.857 | chr12 | - | SE |
| 0.625,0.714,1,1,0.5,1,0.429,1,0.778 | chr6 | + | SE |
| 0.882,0.958,1,1,0.688,1,0.909,0.909,1 | chr1 | - | SE |
| 1,1,0.92,0.9,1,1,1,1,1 | chr14 | - | SE |
| 0.12,0.189,0.27,0.257,0.273,0.297,0.188,0.296,0.28 | chr17 | + | SE |
| 0.917,1,1,1,0.833,1,1,1,1 | chr15 | - | SE |
| 0.833,1,1,1,0.833,1,0.867,1,1 | chr12 | + | SE |
| 0.271,0.248,0.227,0.135,0.137,0.115,0.048,0.049,0.054 | chr13 | - | SE |
| 0.807,0.872,0.869,0.879,0.815,0.695,0.962,0.878,0.925 | chr17 | + | SE |

| | | | |
|---|---|---|---|
| 0.726,1,1,1,1,1,1,1,0.768 | chr5 | + | SE |
| 0.92,0.882,1,0.786,1,1,1,0.947,1 | chr2 | + | SE |
| 0.867,0.806,1,1,1,1,0.8,1,1 | chr15 | + | SE |
| 1,1,1,0.917,1,1,0.833,0.882,1 | chr9 | + | SE |
| 1,0.771,1,1,1,0.691,0.857,1,1 | chr1 | + | SE |
| 0.292,0.245,0.229,0.25,0.294,0.257,0.327,0.324,0.167 | chr17 | + | SE |
| 0.651,1,1,1,1,1,0.691,1,0.789 | chr3 | - | SE |
| 0.8,0.864,0.937,0.968,0.95,1,0.962,0.844,1 | chr16 | + | SE |
| 1,1,1,0.909,1,1,1,1,0.9 | chr16 | + | SE |
| 0.9,0.754,0.875,0.848,1,1,0.973,0.965,1 | chr19 | - | SE |
| 0.882,1,0.8,1,0.84,1,0.944,1,1 | chr16 | + | SE |
| 0.944,1,1,1,0.917,1,1,0.913,1 | chr3 | - | SE |
| 1,0.742,0.961,1,1,1,1,1,0.953 | chr1 | - | SE |
| 0.818,1,0.529,1,1,1,1,0.667,1 | chr16 | + | SE |
| 1,0.882,0.935,0.886,1,1,0.84,0.812,1 | chr1 | + | SE |
| 0.881,0.901,1,1,0.863,1,1,0.741,1 | chr20 | + | SE |
| 0.857,1,1,0.692,1,1,0.692,1,1 | chr17 | - | SE |
| 1,0.908,1,0.958,0.886,1,0.911,0.962,0.907 | chr7 | + | SE |
| 1,1,0.931,1,0.765,1,1,1,1 | chr3 | - | SE |
| 0.739,0.946,1,1,0.939,1,0.907,1,1 | chr2 | - | SE |
| 1,0.765,0.905,1,1,1,0.946,0.818,0.941 | chr17 | + | SE |
| 1,0.9,1,0.879,1,1,0.943,1,1 | chr15 | + | SE |
| 0.786,0.86,1,1,1,1,0.692,1,1 | chr6 | + | SE |
| 1,1,1,1,0.923,1,0.895,1,1 | chr12 | + | SE |
| 1,1,1,0.882,1,1,0.857,1,1 | chr4 | - | SE |
| 1,1,1,1,1,1,1,0.931,0.938 | chr1 | - | SE |
| 0.034,0.012,0.039,0.109,0.036,0.049,0.034,0.031,0.05 | chr2 | + | SE |
| 0.678,0.553,0.5,0.765,0.858,0.817,0.947,0.953,0.935 | chr2 | - | SE |
| 0.977,0.932,0.964,0.872,0.931,0.846,0.729,0.684,0.649 | chr6 | + | SE |
| 0.603,0.729,0.82,0.941,0.923,0.702,0.984,0.905,0.861 | chr3 | - | SE |
| 0.741,0.808,0.864,0.938,0.892,0.852,0.76,0.795,0.952 | chr4 | + | SE |
| 0.124,0.136,0.111,0.105,0.127,0.067,0.05,0.133,0.085 | chr3 | - | SE |
| 0.6,0.804,0.824,0.824,0.935,0.778,0.833,1,0.917 | chr9 | + | SE |
| 0.694,0.76,0.927,0.952,0.946,0.765,0.913,0.829,1 | chr1 | - | SE |
| 0.904,0.762,0.654,0.902,0.542,0.904,1,0.937,0.922 | chr5 | - | SE |
| 0,0.009,0,0,0,0.026,0.274,0.284,0.398 | chr14 | - | SE |
| 0.04,0.07,0.03,0.116,0.098,0,0.092,0.075,0.066 | chr5 | - | SE |
| 0.607,0.921,0.795,0.659,0.734,0.782,0.869,0.576,0.614 | chr15 | + | SE |
| 0.605,0.501,0.931,0.672,0.795,0.91,1,1,1 | chr1 | + | SE |
| 0.98,0.97,0.984,0.996,0.995,1,0.666,0.547,0.705 | chr6 | - | SE |
| 0.2,1,1,0.6,1,0,1,0.818,0.667 | chr6 | + | SE |
| 1,1,1,0.857,0.846,1,0.36,0.76,0.579 | chr20 | + | SE |
| 0.043,0.093,0.091,0,0,0,0.042,0.103,0.04 | chr16 | + | SE |

| | | | |
|---|---|---|---|
| 0.686,0.535,0.844,0.838,0.733,0.765,0.728,0.763,0.77 | chr19 | + | SE |
| 0.855,0.742,0.871,0.775,0.938,0.831,0.883,0.876,0.823 | chrX | + | SE |
| 0.843,0.866,0.819,0.721,0.802,0.818,1,1,1 | chr1 | - | SE |
| 0.733,0.8,0.795,0.744,0.875,0.727,0.76,1,0.6 | chr4 | - | SE |
| 0.633,0.852,0.547,0.801,0.778,0.673,0.893,0.946,0.96 | chr4 | + | SE |
| 0,0.009,0,0,0,0.026,0.261,0.264,0.309 | chr14 | - | SE |
| 0,0.006,0,0,0,0.014,0.5,0.461,0.373 | chr14 | - | SE |
| 0.824,0.78,0.571,0.743,0.769,1,0.71,0.455,0.857 | chr3 | - | SE |
| 0.158,0.098,0.069,0.111,0.18,0.118,0.026,0.062,0.06 | chr2 | - | SE |
| 0.463,0.524,0.484,0.444,0.515,0.389,0.245,0.207,0.186 | chr1 | - | SE |
| 0.793,0.793,1,0.669,0.851,0.805,0.609,0.498,0.817 | chr4 | + | SE |
| 1,0.667,1,0.545,0.818,0.286,0.909,1,1 | chr1 | + | SE |
| 0.788,0.62,0.624,0.708,0.706,0.514,0.596,0.623,0.759 | chr3 | - | SE |
| 0.094,0.098,0.069,0.111,0.194,0.118,0.026,0.032,0.06 | chr2 | - | SE |
| 0.1,0.212,0.033,0.061,0.154,0.176,0.111,0.111,0.037 | chr14 | + | SE |
| 0.658,0.554,0.649,0.646,0.872,0.605,0.825,0.609,0.837 | chr4 | + | SE |
| 0.377,0.515,0.837,1,1,1,1,1,1 | chr4 | + | SE |
| 0.765,0.633,0.63,0.474,0.444,0.516,0.638,0.553,0.631 | chr12 | + | SE |
| 0.928,0.948,0.946,0.557,0.717,0.795,0.675,0.757,0.722 | chr7 | + | SE |
| 0.712,0.825,0.755,0.702,1,1,0.693,0.884,0.683 | chr1 | - | SE |
| 0.205,0.158,0.123,0.154,0.227,0.213,0.216,0.155,0.1 | chr15 | + | SE |
| 0.154,0.252,0.319,0.3,0.19,0.307,0.222,0.257,0.293 | chr19 | + | SE |
| 0.714,0.667,0.636,0.556,0.846,1,0.5,0.524,1 | chr19 | - | SE |
| 0.561,0.423,0.344,0.428,0.588,0.505,0.399,0.552,0.636 | chr12 | - | SE |
| 0.727,0.714,0.857,0.882,0.556,0.571,0.773,0.92,0.727 | chr12 | - | SE |
| 1,0.455,0.939,1,1,0.833,1,0.667,0.5 | chr18 | - | SE |
| 0.672,0.619,0.891,0.832,0.672,0.804,1,1,1 | chr15 | + | SE |
| 0.445,0.785,0.745,1,1,0.523,0.919,1,0.933 | chr4 | - | SE |
| 1,0.471,0.926,1,1,0.818,1,0.667,0.5 | chr18 | - | SE |
| 0.48,0.447,0.469,0.382,0.489,0.453,0.429,0.506,0.378 | chr12 | + | SE |
| 0.778,0.714,0.7,1,0.667,0.667,0.92,1,1 | chr1 | - | SE |
| 1,1,1,0.74,0.85,1,0.124,0.57,0.561 | chr20 | + | SE |
| 0.227,0.281,0.355,0.202,0.156,0.179,0.395,0.366,0.324 | chr20 | - | SE |
| 1,1,1,0.538,0.81,1,0.733,0.417,0.455 | chr2 | + | SE |
| 0.155,0.408,0.035,0.515,0.441,0.256,0.183,0.284,0.341 | chr10 | - | SE |
| 0.535,0.706,0.795,0.545,1,1,0.538,0.75,0.571 | chr1 | - | SE |
| 0.419,0.818,0.733,0.714,0.846,1,0.68,0.857,0.579 | chr5 | + | SE |
| 0.548,0.339,0.418,0.364,0.548,0.422,0.479,0.517,0.389 | chr10 | - | SE |
| 0.2,0.211,0.137,0.25,0.579,0.286,0.118,0.154,0.143 | chr2 | - | SE |
| 0.882,0.368,0.529,1,0.875,1,0.875,0.9,0.818 | chr6 | + | SE |
| 0.676,0.834,0.584,0.73,0.762,0.859,0.765,0.619,0.494 | chr2 | - | SE |
| 0.172,0.211,0.137,0.25,0.579,0.286,0.118,0.154,0.143 | chr2 | - | SE |
| 0.238,0.364,0.235,0.25,0.314,0.167,0.2,0.154,0.368 | chr2 | - | SE |

| | | | |
|---|---|---|---|
| 0.394,1,0.545,1,0.2,1,0.636,0.793,1 | chr16 | - | SE |
| 0.517,0.418,0.576,0.979,0.882,0.879,0.576,0.625,0.5 | chr5 | - | SE |
| 0.727,0.455,0.7,0.652,0.478,0.333,0.875,1,0.895 | chrX | + | SE |
| 0.952,0.915,0.948,0.567,0.505,0.671,0.384,0.737,0.405 | chr7 | + | SE |
| 0.86,0.746,0.609,0.629,0.797,0.348,0.752,0.658,0.616 | chr7 | - | SE |
| 0.75,1,0.79,0.725,0.84,0.53,0.653,0.131,1 | chr12 | - | SE |
| 0.846,0.25,0.467,1,0.75,1,0.833,0.857,0.692 | chr6 | + | SE |
| 0.215,0.044,0.475,0.118,0.413,0,0.287,0,0.311 | chr5 | - | SE |
| 0.738,0.478,0.753,0.567,0.857,0.611,0.794,0.765,0.714 | chrX | + | SE |
| 0.172,0.176,0.326,0.2,0.29,0.111,0.133,0.061,0.176 | chr19 | + | SE |
| 0.1,0.257,0.116,0.6,0.2,0.217,0.16,0.137,0.119 | chr16 | - | SE |
| 0.334,0.805,0.273,1,1,0.429,1,0.693,1 | chr4 | - | SE |
| 0.172,0.208,0.333,0.2,0.312,0.111,0.133,0.061,0.176 | chr19 | + | SE |
| 0.312,0.362,0.28,0.462,0.2,0.818,0.667,0.353,0.36 | chr2 | - | SE |
| 0.389,0.375,0.333,0.44,0.259,0.818,0.75,0.421,0.333 | chr2 | - | SE |
| 0.097,0.22,0.407,0.1,0.286,0.6,0.273,0.667,0.308 | chr2 | + | SE |
| 0.688,0.65,0.474,1,0.429,0.583,0.471,0.571,0.273 | chr3 | - | SE |
| 0.221,0.74,0.532,0.82,1,0.362,0.221,0.888,1 | chr12 | + | SE |
| 0.677,0.429,0.852,0.36,0.176,0.333,0.647,0.739,0.838 | chr16 | + | SE |
| 0.714,1,0.556,1,0.429,0,0.5,1,0.2 | chr17 | + | SE |
| 0.152,0.256,0.556,0.1,0.333,0.6,0.36,0.667,0.357 | chr2 | + | SE |
| 0.176,0.158,0.429,0,0.333,0.5,0.2,0.333,0.1 | chr2 | + | SE |
| 0.435,0.714,0.524,0,0.067,0.263,0.143,0,0 | chr22 | + | SE |
| 0.125,0.059,0.36,0.053,0.091,0,0.238,0,0.1 | chr2 | + | SE |
| 0.389,0.429,0.591,0.061,0.116,0.04,1,1,1 | chr1 | + | SE |
| 0,0.167,0.12,0.091,0.118,0,0.026,0.158,0.094 | chr3 | - | SE |
| 1,0.833,0.846,0.857,0.833,1,1,0.857,0.667 | chrX | - | RI |
| 0.84,0.949,0.889,0.931,1,1,1,1,0.931 | chr20 | + | RI |
| 0.417,0.684,0.806,0.615,0.636,0.714,0.5,0.562,1 | chr20 | + | RI |
| 0.333,0.211,0.344,0.467,0.455,0.2,0.222,0.294,0.231 | chr20 | + | RI |
| 0.197,0.181,0.111,0.2,0.259,0.385,0.322,0.347,0.374 | chr19 | - | RI |
| 0.273,0.307,0.225,0.286,0.381,0.259,0.302,0.312,0.179 | chr19 | + | RI |
| 1,1,1,0.561,0.585,0.413,1,1,1 | chr22 | - | RI |
| 0.312,0.369,0.351,0.362,0.365,0.35,0.473,0.438,0.428 | chr17 | - | RI |
| 0.421,0.285,0.257,0.161,0.25,0.115,0.179,0.207,0.213 | chr17 | + | RI |
| 0.455,0.6,0.459,0.469,0.448,0.4,0.333,0.44,0.654 | chr17 | - | RI |
| 0.298,0.16,0.17,0.143,0.116,0.127,0.069,0.175,0.071 | chr17 | + | RI |
| 0.8,0.704,1,0.739,0.568,0.913,0.579,0.92,0.778 | chr16 | + | RI |
| 0.556,0.778,0.81,0.714,0.143,0.222,0.535,0.419,0.463 | chr19 | + | RI |
| 0.273,0.571,0.636,0.349,0.333,0.2,0.385,0.25,0.263 | chr15 | - | RI |
| 1,1,1,1,1,1,0.887,0.929,0.702 | chr1 | - | RI |
| 0.967,0.929,0.96,1,1,1,1,1,1 | chr16 | - | RI |
| 0.7,0.909,0.636,0.5,0.25,0.222,0.76,0.895,0.8 | chr5 | - | RI |

| | | | |
|---|---|---|---|
| 0.833,1,0.81,0.846,1,1,1,1,0.75 | chr19 | - | RI |
| 0.846,0.927,0.893,0.875,0.602,0.929,0.818,0.713,0.735 | chr1 | + | RI |
| 0.956,0.958,0.821,0.871,0.81,0.636,0.815,0.706,0.878 | chr9 | + | RI |
| 0.259,0.333,0.412,0.213,0.444,0.333,0.37,0.159,0.333 | chr12 | - | RI |
| 0.867,0.733,0.852,1,1,1,1,1,0.714 | chr3 | - | RI |
| 0.143,0.092,0.262,0.143,0.1,0.062,0.086,0.073,0.039 | chr5 | + | RI |
| 0.226,0.517,0.356,0.418,0.5,0.286,0.283,0.462,0.394 | chr3 | + | RI |
| 1,1,0.833,1,0.667,1,1,0.714,1 | chr3 | - | RI |
| 0.812,0.904,0.964,0.809,0.74,0.719,0.884,0.865,0.873 | chr6 | - | RI |
| 0.132,0.081,0.053,0.029,0.094,0.238,0.149,0.185,0.125 | chr3 | - | RI |
| 0.899,1,1,1,1,1,0.954,0.913,1 | chr5 | + | RI |
| 0.733,1,1,1,1,1,0.31,1,0.905 | chr7 | + | RI |
| 1,1,1,1,0.75,1,0.957,1,1 | chr6 | - | RI |
| 0.758,0.643,0.771,1,0.727,0.294,0.913,0.778,0.926 | chr8 | - | RI |
| 0.09,0.115,0.057,0.085,0.072,0.088,0.209,0.061,0.012 | chr11 | + | RI |
| 0.161,0.097,0.125,0.1,0.385,0.2,0.3,0.25,0.556 | chr11 | - | RI |
| 0.536,0.491,0.634,0.292,0.464,0.536,0.224,0.104,0.162 | chr6 | + | RI |
| 1,0.636,0.8,1,1,1,1,1,1 | chr17 | + | RI |
| 0.107,0.018,0.237,0.137,0.158,0.171,0.08,0.034,0.065 | chr11 | - | RI |
| 0.929,1,0.625,1,1,0.636,1,1,0.8 | chr6 | - | RI |
| 0.793,1,1,0.871,0.818,0.692,1,1,1 | chrX | + | RI |
| 0.919,1,1,0.915,0.887,0.947,0.879,1,0.857 | chr11 | - | RI |
| 0.304,0.333,0.412,0.289,0.5,0.143,0.143,0.417,0.429 | chr17 | - | RI |
| 1,1,1,0.882,1,0.818,1,1,1 | chr12 | + | RI |
| 0.224,0.207,0.18,0.212,0.35,0.165,0.128,0.13,0.241 | chr12 | - | RI |
| 1,1,1,1,1,1,1,1,1 | chr12 | - | RI |
| 0.75,0.871,1,0.733,0.935,0.643,1,0.8,0.882 | chr14 | + | RI |
| 0.047,0.105,0.065,0.04,0.094,0.051,0.297,0.238,0.296 | chr15 | - | RI |
| 0.364,0.179,0.333,0.233,0.253,0.254,0.347,0.259,0.169 | chr15 | - | RI |
| 0.398,0.215,0.312,0.196,0.186,0.203,0.281,0.215,0.205 | chr20 | + | MXE |
| 0.973,0.896,0.972,1,1,1,0.983,1,1 | chr17 | + | MXE |
| 1,1,1,1,1,1,0.826,1,0.559 | chr17 | - | MXE |
| 0.511,0.516,0.213,0.532,0.512,0.651,0.511,0.651,0.81 | chr7 | + | MXE |
| 0.854,0.881,0.847,0.905,0.891,0.882,0.893,0.87,0.91 | chr4 | - | MXE |
| 0.632,0.478,0.457,0.325,0.301,0.4,0.315,0.304,0.373 | chr2 | + | MXE |
| 0.862,0.953,0.853,0.858,0.93,0.792,0.875,0.9,0.955 | chr3 | + | MXE |
| 0.429,0.2,0.538,0.154,0.15,0.182,0.2,0.067,0.263 | chr1 | - | MXE |
| 0.868,1,0.861,0.766,0.766,1,1,1,0.954 | chr19 | - | MXE |
| 0.111,0.333,0.118,0.1,0.158,0,0.258,0.12,0.121 | chr3 | - | MXE |
| 0.185,0.101,0.294,0.057,0.134,0.097,0.163,0.169,0.18 | chr19 | + | A3SS |
| 0.683,0.806,0.797,0.811,0.699,0.5,0.931,0.773,0.829 | chr17 | + | A3SS |
| 1,1,0.694,0.858,0.654,1,0.831,0.275,0.752 | chr6 | + | A3SS |
| 0.26,0.493,0.264,0.467,0.462,0.538,0.343,0.517,0.37 | chr6 | - | A3SS |

| | | | |
|---|---|---|---|
| 0.889,0.869,0.758,0.453,0.57,1,0.356,0.659,0.786 | chr5 | - | A3SS |
| 0.882,0.867,0.769,0.467,0.565,1,0.333,0.636,0.769 | chr5 | - | A3SS |
| 1,1,1,1,1,0.726,1,1,1 | chr11 | - | A3SS |
| 1,0.579,1,1,0.818,1,1,1,0.833 | chr4 | - | A3SS |
| 0.648,1,1,0.871,0.754,1,1,1,1 | chr8 | + | A3SS |
| 1,0.765,0.9,0.892,0.9,1,0.931,0.889,0.829 | chr9 | - | A3SS |
| 0.917,1,0.833,1,1,1,0.778,1,1 | chr17 | - | A5SS |
| 0.61,1,0.941,1,1,1,1,1,1 | chr17 | - | A5SS |
| 0.535,0.463,0.674,0.216,0.828,1,1,1,1 | chr16 | + | A5SS |
| 1,0.632,0.842,1,1,1,1,1,0.894 | chr1 | - | A5SS |
| 0.89,1,0.783,1,0.763,0.612,0.935,0.878,1 | chr2 | + | A5SS |
| 0.196,0.211,0.229,0.21,0.191,0.204,0.434,0.428,0.41 | chr1 | + | A5SS |
| 0.224,0.332,0.297,0.343,0.472,0.138,0.26,0.323,0.383 | chr3 | + | A5SS |
| 1,1,1,0.971,1,1,0.983,1,1 | chr14 | + | A5SS |
| 0.625,1,0.778,0.778,1,0.429,1,0.636,0.5 | chrX | - | A5SS |
| 0.411,0.556,0.609,0.015,0,0,0,0,0 | chr14 | + | A5SS |
