## Supplementary material for "TBCK Deficiency Alters Ribosomal Function, RNA Splicing, and miRNA Networks: Insights from Multi-Omics Analyses": proteomics_analysis_protein_list

| Protein | log2fc | neg_log10_ |
|---|---|---|
| LOC102724159 | 0.420454 | NA |
| CS | 0.133566 | NA |
| MPDU1 | 0.124221 | NA |
| PDIA4 | 0.082241 | NA |
| ACTN4 | -0.10745 | NA |
| SSB | -0.10745 | NA |
| PAFAH1B1 | -0.12167 | NA |
| ALDH4A1 | -0.24107 | 13.9183 |
| RCN1 | 0.189514 | 12.75681 |
| UFM1 | -0.12877 | 12.3299 |
| CLCC1 | -0.29248 | 12.32416 |
| cDNA, FLJ93684, highly similar to Homo sapiens mab-21-like 1 (C. elegans) (M | -1.91478 | 12.25583 |
| GLRX | -0.63691 | 11.74171 |
| ACOT7 | -0.39719 | 11.42609 |
| cDNA FLJ75877, highly similar to Homo sapiens 5'-nucleotidase, cytosolic II (N | -0.15763 | 10.43476 |
| TPP2 | -0.07417 | 9.924228 |
| YTHDF3 | 0.156468 | 9.664133 |
| TMEM167A | -0.30542 | 9.411599 |
| FHOD1 | 0.328849 | 9.292728 |
| GRHPR | -0.1027 | 9.199955 |
| p90 OS=Homo sapiens | 0.861313 | 9.183812 |
| Ubiquitin-specific protease 7 isoform (Fragment) OS=Homo sapiens | 0.115186 | 8.996142 |
| APOBEC3F | -0.48798 | 8.978019 |
| MSRB3 | -0.13114 | 8.945888 |
| SYNJ2 | 0.625085 | 8.919913 |
| Solute carrier family 2 (Facilitated glucose transporter), member 1 variant (Fr | -0.66115 | 8.696248 |
| TEX10 | -0.20604 | 8.566039 |
| RAB11B | -0.12759 | 8.481172 |
| cDNA FLJ54730, highly similar to cAMP-dependent protein kinase, beta-2-cat | -0.29079 | 8.371686 |
| NANS | -0.14414 | 8.370938 |
| Isoform 3 of NTF2-related export protein 2 OS=Homo sapie | 0.286609 | 8.36568 |
| LRRFIP1 | 0.236805 | 8.206659 |
| FUCA2 | -0.41565 | 8.196289 |
| ITPR2 | -0.37221 | 8.09517 |
| ATG2B | -0.22532 | 8.013008 |
| BAX | 0.231699 | 7.893629 |
| WDR36 | 0.144159 | 7.88701 |
| CECR5 | 0.365544 | 7.822896 |
| ALG2 | -0.38265 | 7.815259 |
| PTDSS2 | 0.189057 | 7.787113 |
| RPL9 | -0.07298 | 7.672427 |
| RBM12B | 0.180847 | 7.632192 |
| ZGPAT | 0.27607 | 7.583548 |
| RPL11 | 0.025292 | 7.576652 |
| SLTM | 0.16186 | 7.569385 |
| USE1 | -0.13773 | 7.565119 |

| | | |
|---|---|---|
| HSDL2 | 0.177915 | 7.560433 |
| ASNSD1 | 0.182084 | 7.52747 |
| ARF5 | 0.527775 | 7.521518 |
| CCS | -0.17665 | 7.507843 |
| Isoform 2 of B-cell receptor-associated protein 29 OS=Homo sapiens | -0.14836 | 7.504815 |
| NSUN2 | 0.1948 | 7.504609 |
| FLJ10357 | 0.228858 | 7.467976 |
| VAV2 | -0.37221 | 7.455076 |
| DRG2 | -0.17665 | 7.45398 |
| H2AFY | -0.1854 | 7.420419 |
| ZNF428 | 0.309391 | 7.358087 |
| TMSB10 | 0.609639 | 7.35149 |
| ZC3H7B | -0.17716 | 7.325852 |
| BYSL | 0.054785 | 7.307049 |
| Lipase OS=Homo sapiens | -0.54744 | 7.305643 |
| VPS52 | 0.297541 | 7.2602 |
| EIF5 | 0.175442 | 7.259339 |
| TMSB4X | 1.027834 | 7.257123 |
| HEL-S-22 | -0.1706 | 7.252249 |
| hCG_1773630 | 0.820831 | 7.196651 |
| DDAH1 | -0.23053 | 7.181047 |
| TWF2 | -0.13114 | 7.153132 |
| INPP4B | -0.4718 | 7.129321 |
| MTDH | 0.1948 | 7.125027 |
| cDNA FLJ56092, highly similar to Pentatricopeptide repeat protein 1 OS=Hom | -0.12995 | 7.10825 |
| RHOQ | -0.16302 | 7.081958 |
| Phosphomannomutase OS=Homo sapiens | -0.1027 | 7.065177 |
| 6-phosphogluconate dehydrogenase, decarboxylating OS=Homo sapiens | -0.25394 | 7.062299 |
| DGKA | -0.23938 | 7.040489 |
| ITFG1 | -0.13705 | 7.020983 |
| CLUH | -0.07655 | 7.010402 |
| PTMS | 0.460289 | 6.961426 |
| KIF1BP | 0.289953 | 6.952438 |
| DOPEY2 | 0.354845 | 6.941503 |
| PPP4R3A | -0.14768 | 6.917706 |
| MIA3 | 0.171735 | 6.890126 |
| Amino acid transporter (Fragment) OS=Homo sapiens | 0.395174 | 6.853851 |
| ERLIN2 | -0.20252 | 6.752762 |
| DNAJC3 | -0.12285 | 6.713433 |
| CDK11B | 0.284084 | 6.683457 |
| STAT3 | -0.18187 | 6.678239 |
| CMBL | -0.1901 | 6.643073 |
| Isoform 2 of Neutral alpha-glucosidase AB OS=Homo sapien | 0.153234 | 6.60832 |
| CTPS1 | -0.22598 | 6.602457 |
| PWP1 | 0.191079 | 6.59776 |
| GALM | -0.28846 | 6.594445 |
| HEL-211 | 0.667786 | 6.585317 |

| | | |
|---|---|---|
| SLC25A11 | -0.17548 | 6.579601 |
| TMEM165 | 0.013237 | 6.576618 |
| ARHGAP35 | 0.292927 | 6.575934 |
| PPIG | 0.278591 | 6.559464 |
| GK | -0.23704 | 6.552543 |
| RHBDD3 | 0.722253 | 6.460996 |
| PDCD10 | -0.2265 | 6.459454 |
| SERINC1 | -0.11999 | 6.458035 |
| TAOK1 | 0.144621 | 6.455675 |
| PCYOX1 | -0.14059 | 6.433252 |
| Chondroitin sulfate proteoglycan 2 (Versican) variant (Fragment) OS=Homo s | 1.007705 | 6.386196 |
| PIGU | -0.22012 | 6.357026 |
| MOXD1 | -0.52494 | 6.355613 |
| SLC12A4 | -0.22481 | 6.355363 |
| TMEM87B | -0.08677 | 6.334035 |
| HEATR5A | -0.23704 | 6.323638 |
| NELFA | -0.22115 | 6.315947 |
| MYL6 | -0.14937 | 6.283419 |
| SRPK1 | -0.12098 | 6.280844 |
| TNS3 | -0.46602 | 6.271189 |
| ICAM5 | 0.213898 | 6.26821 |
| TUFM | 0.126672 | 6.265644 |
| PGM2 | -0.28029 | 6.247005 |
| POLR2B | 0.132803 | 6.238269 |
| DDX1 | -0.0558 | 6.23367 |
| FCHSD1 | -0.41517 | 6.212643 |
| Isoform 3 of Plasminogen activator inhibitor 1 RNA-binding protein OS=Homo | 0.194016 | 6.193383 |
| DHTKD1 | 0.487941 | 6.183132 |
| SCRN3 | -0.25693 | 6.182958 |
| POTEKP | -0.20773 | 6.178854 |
| DDX10 | 0.305586 | 6.142482 |
| Isoform 2 of Small EDRK-rich factor 2 OS=Homo sapien | 0.787242 | 6.097444 |
| UTP14A | 0.230106 | 6.078302 |
| OCRL | -0.15122 | 6.067057 |
| PPP1R8 | 0.308947 | 6.059565 |
| CSRP1 | -0.51455 | 6.05821 |
| CLPP | 0.235902 | 5.993587 |
| TCF24 | -0.30361 | 5.977779 |
| TCF25 | 0.139242 | 5.966493 |
| KIF13A | -0.13164 | 5.943541 |
| CYB5B | -0.19833 | 5.934244 |
| WASF1 | -0.50415 | 5.914626 |
| EHD3 | -0.37043 | 5.895815 |
| Isoform 2 of Ras-related protein Rab-5C OS=Homo sapien | -0.28729 | 5.885674 |
| cDNA FLJ56357, highly similar to Homo sapiens apolipoprotein A-I binding pr | -0.0706 | 5.884552 |
| CAPN1 | -0.20538 | 5.88287 |
| PSMA6 | -0.09083 | 5.879881 |

| | | |
|---|---|---|
| DYNLRB1 | -0.24575 | 5.866764 |
| IPO7 | -0.04506 | 5.844665 |
| LRRC59 | 0.202705 | 5.840756 |
| Isoform 2 of Drebrin-like protein OS=Homo sapie | 0.199435 | 5.831143 |
| PHF3 | 0.381831 | 5.79285 |
| OSGEP | -0.37453 | 5.767794 |
| cDNA FLJ16777 fis, clone BRHIP2029567, highly similar to Cell division cycle 5 | 0.120082 | 5.761965 |
| METTL15 | 0.448064 | 5.76118 |
| SLC30A9 | 0.55187 | 5.749849 |
| ETF1 | -0.10695 | 5.748187 |
| RAC2 | 0.51137 | 5.744827 |
| UQCRQ | 0.176678 | 5.725074 |
| cDNA FLJ76981, highly similar to Homo sapiens golgi autoantigen, golgin subf | 0.196953 | 5.701449 |
| NEDD4 | -0.27731 | 5.683782 |
| Isoform A of Protein CutA OS=Homo sapie | -0.12809 | 5.666399 |
| OAS3 | -0.2516 | 5.633305 |
| ZAK | 0.586004 | 5.628116 |
| Isoform 2 of Negative elongation factor E OS=Homo sapien | 0.287872 | 5.615804 |
| RPL22 | -0.12167 | 5.615629 |
| Isoform 3 of Double-strand break repair protein MRE11A OS=Homo sapiens | 0.145851 | 5.61522 |
| RBM3 | 0.525106 | 5.592481 |
| RBCK1 | 0.260163 | 5.580476 |
| MIF | -0.29079 | 5.577301 |
| cDNA FLJ56561 OS=Homo sapiens | 0.074472 | 5.575282 |
| Isoform 3 of GRAM domain-containing protein 3 OS=Homo sapiens | -0.14128 | 5.572849 |
| VAMP3 | -0.1583 | 5.563907 |
| HIP1 | 0.348883 | 5.561711 |
| FDPS | 0.04657 | 5.55472 |
| ITPRIP | 0.648027 | 5.552739 |
| CPD | -0.20706 | 5.551166 |
| YWHAQ | 0.159855 | 5.538568 |
| EIF2AK4 | -0.23587 | 5.537353 |
| hCG_2014768 | 0.113206 | 5.534561 |
| CLTC | -0.14886 | 5.526297 |
| CDC73 | 0.20892 | 5.524755 |
| TOR1B | -0.22532 | 5.510003 |
| CDK4 | -0.33969 | 5.473062 |
| YBX3 | 0.251378 | 5.470671 |
| ITSN1 | 0.176678 | 5.470032 |
| SNRPB2 | 0.386135 | 5.461922 |
| ATM | 0.465969 | 5.46031 |
| cDNA, FLJ94965, highly similar to Homo sapiens leucyl/cystinyl aminopeptida | -0.2089 | 5.460037 |
| Apolipoprotein B mRNA editing enzyme, catalytic polypeptide-like 3C variant | -0.19598 | 5.455788 |
| PRMT3 | 0.166335 | 5.440895 |
| hCG_1821276 | 0.153234 | 5.430381 |
| SP3 | 0.448064 | 5.429677 |
| OTUD4 | 0.319108 | 5.413058 |

| | | |
|---|---|---|
| AAMDC | -0.16353 | 5.402187 |
| C19orf66 | -0.20252 | 5.397704 |
| FLOT2 | -0.1583 | 5.373897 |
| Isoform 1 of Arf-GAP with SH3 domain, ANK repeat and PH domain-containin | -0.17077 | 5.373521 |
| GOLGA2 | 0.102964 | 5.367334 |
| BECN1 | 0.192776 | 5.364676 |
| ACSL3 | 0.145851 | 5.35913 |
| Isoform 5 of Oxysterol-binding protein-related protein 6 OS=Homo sapiens | 1.151753 | 5.358937 |
| NCSTN | -0.1642 | 5.345428 |
| RARS2 | 0.272739 | 5.342961 |
| Isoform 3 of Malate dehydrogenase, cytoplasmic OS=Homo sapie | -0.12877 | 5.335319 |
| ACTR1B | -0.30245 | 5.322735 |
| Isoform 2 of Lysine-specific histone demethylase 1A OS=Homo sapien | 0.430212 | 5.296201 |
| GNPNAT1 | 0.158162 | 5.287193 |
| ABCE1 | 0.048991 | 5.282629 |
| LANCL1 | 0.383556 | 5.280865 |
| LIX1L | 0.481766 | 5.273337 |
| ATG9A | -0.15595 | 5.273262 |
| RPS5 | 0.165561 | 5.269618 |
| Isoform 5 of Splicing factor 1 OS=Homo sapi | 0.110294 | 5.242889 |
| Isoform 6 of Pleckstrin homology domain-containing family A member 5 OS= | 0.304319 | 5.239034 |
| Isoform 2 of Electron transfer flavoprotein subunit beta OS=Homo sapie | -0.03478 | 5.220036 |
| ARFGEF1 | -0.13282 | 5.203279 |
| Isoform 2 of Coiled-coil domain-containing protein 80 OS=Homo sapiens | 0.718033 | 5.199268 |
| KPNA4 | -0.1943 | 5.191442 |
| BOD1L1 | 0.611832 | 5.183799 |
| Annexin OS=Homo sapiens | -0.04315 | 5.181465 |
| Isoform 4 of Pseudouridine-5'-phosphatase OS=Homo sapie | -0.32986 | 5.177117 |
| FAF2 | -0.09152 | 5.165506 |
| cDNA FLJ76888, highly similar to Homo sapiens RNA binding motif protein 6 ( | 0.300518 | 5.164874 |
| UBA5 | -0.06822 | 5.162299 |
| STIM1 | -0.30531 | 5.144338 |
| HEATR3 | -0.33272 | 5.1377 |
| CYBA | 0.299252 | 5.136423 |
| SH3PXD2B | 0.404231 | 5.130844 |
| LUM | -0.49375 | 5.127776 |
| Isoform 3 of Nucleolar protein 3 OS=Homo sapie | -0.17598 | 5.116495 |
| Isoform 5 of Radixin OS=Homo sapi | 0.135557 | 5.112084 |
| HSPG2 | 0.350602 | 5.111135 |
| CAPN5 | -0.12472 | 5.11051 |
| cDNA, FLJ95644, highly similar to Homo sapiens solute carrier family 20 (pho: | -0.17127 | 5.108615 |
| Isoform 2 of Integrin alpha-11 OS=Homo sapiens | 0.734542 | 5.104 |
| MOCS2 | -0.14414 | 5.095216 |
| HECTD1 | 0.267704 | 5.093883 |
| NRGN | -0.37105 | 5.075246 |
| NQO2 | 0.333115 | 5.073427 |
| FAM49B | -0.05987 | 5.063828 |

| | | |
|---|---|---|
| FBXO6 | -0.65653 | 5.059205 |
| UEVLD | -0.18473 | 5.055526 |
| Isoform 2 of Probable hydrolase PNKD OS=Homo sapie | 0.267256 | 5.04639 |
| MYD88 | -0.25225 | 5.029476 |
| TGFB1I1 | -0.14059 | 5.028593 |
| NGDN | 0.365544 | 5.025962 |
| YPEL5 | 0.382267 | 5.022961 |
| Isoform 2 of Splicing regulatory glutamine/lysine-rich protein 1 OS=Homo sap | -0.28846 | 5.020866 |
| SASH1 | 0.237604 | 5.020601 |
| MRPS9 | 0.402936 | 5.017701 |
| PBEF1 | -0.3269 | 5.006895 |
| cDNA FLJ77534, highly similar to Homo sapiens membrane protein, palmitoy | -0.20186 | 5.005068 |
| CPNE3 | -0.17363 | 4.991925 |
| RAB1A | -0.13114 | 4.985448 |
| cDNA FLJ76392, highly similar to Homo sapiens mitochondrial ribosomal prot | 0.37112 | 4.971705 |
| USP39 | -0.16892 | 4.922563 |
| EPRS | 0.126207 | 4.916486 |
| MEN1 | 0.939086 | 4.910961 |
| ZCCHC8 | 0.431516 | 4.91041 |
| MAN1A1 | 0.459405 | 4.908272 |
| PPIL1 | -0.16538 | 4.907517 |
| cDNA FLJ51840, highly similar to Alpha-actinin-2 OS=Homo sapiens | -0.19833 | 4.882551 |
| LXN | -0.46657 | 4.878457 |
| AIM1 | 0.215143 | 4.876417 |
| Isoform Long of Ubiquitin fusion degradation protein 1 homolog OS=Homo sa | -0.16134 | 4.873091 |
| ADAMTSL1 | 0.433692 | 4.851552 |
| UQCRFS1 | 0.138477 | 4.8382 |
| Isoform 2 of HAUS augmin-like complex subunit 7 OS=Homo sapien | 1.638481 | 4.835115 |
| WDR6 | -0.16841 | 4.824466 |
| OSBPL3 | -0.36989 | 4.821646 |
| cDNA, FLJ96161 OS=Homo sapiens | 0.196497 | 4.818992 |
| SNRPD2 | 0.140471 | 4.815615 |
| Toll interacting protein variant (Fragment) OS=Homo sapiens | -0.06394 | 4.811545 |
| BOP1 | -0.09983 | 4.800734 |
| PVRL3 | 0.09611 | 4.80064 |
| Isoform 2 of Leupaxin OS=Homo sapie | 0.376265 | 4.794375 |
| cDNA FLJ55996, highly similar to Conserved oligomeric Golgi complex compo | -0.13705 | 4.778596 |
| RBM25 | 0.139705 | 4.772806 |
| Isoform C of AP-1 complex subunit beta-1 OS=Homo sapien | -0.05629 | 4.769738 |
| NUP58 | 0.31066 | 4.765932 |
| PRPF6 | 0.178373 | 4.754332 |
| RNASEH2C | 1.676539 | 4.753804 |
| Isoform 3 of WAS/WASL-interacting protein family member 1 OS=Homo sapi | 0.568435 | 4.744599 |
| NAV1 | 0.452422 | 4.741279 |
| GOT2 | -0.11861 | 4.737737 |
| TMEM85 | 0.178373 | 4.736803 |
| MTCH2 | 0.135257 | 4.731309 |

| | | |
|---|---|---|
| BCL10 | 0.10785 | 4.728607 |
| PROCR | 0.223866 | 4.722365 |
| GALNT6 | 0.124221 | 4.699928 |
| SELM | -0.23001 | 4.699635 |
| LAP3 | -0.20471 | 4.696955 |
| PAWR | 0.238855 | 4.69627 |
| Isoform 3 of Fermitin family homolog 2 OS=Homo sapiens | -0.13182 | 4.676775 |
| PLIN3 | -0.0939 | 4.671825 |
| Isoform sGi2 of Guanine nucleotide-binding protein G(i) subunit alpha-2 OS=I | -0.146 | 4.666458 |
| PDCD11 | 0.144621 | 4.65443 |
| IFIT1 | -0.48335 | 4.652174 |
| Isoform 6 of Serine/threonine-protein kinase WNK1 OS=Homo sapie | 0.244311 | 4.63276 |
| ABRACL | -0.17665 | 4.628103 |
| TAGLN3 | 0.227156 | 4.625212 |
| MYL9 | -0.42144 | 4.621582 |
| LOXL2 | -0.15359 | 4.62022 |
| Isoform Heart of ATP synthase subunit gamma, mitochondrial OS=Homo sapi | -0.02951 | 4.619145 |
| PIK3IP1 | -0.46542 | 4.618688 |
| CORO1B | -0.13823 | 4.603831 |
| cDNA FLJ10711 fis, clone NT2RP3000917, highly similar to 5'-3' exoribonucle | 0.081024 | 4.598412 |
| Isoform 2 of Neuromodulin OS=Homo sapien | 1.692761 | 4.596275 |
| PLA2G15 | -0.1524 | 4.594038 |
| cDNA, FLJ92608, highly similar to Homo sapiens aldehyde dehydrogenase 1 f | 0.703009 | 4.585813 |
| Isoform 3 of Polypyrimidine tract-binding protein 1 OS=Homo sapien | 0.065698 | 4.5785 |
| RPL13 | -0.12285 | 4.576016 |
| FIP1L1 | 0.080554 | 4.574232 |
| ESYT1 | -0.13755 | 4.567024 |
| RDH11 | -0.1259 | 4.554441 |
| YBX1 | 0.236354 | 4.5511 |
| Mitogen-activated protein kinase kinase kinase 7 interacting protein 1 isofor | -0.30361 | 4.547742 |
| PDS5A | 0.550527 | 4.543481 |
| GIGYF2 | 0.085893 | 4.542921 |
| UNC45A | -0.04864 | 4.534183 |
| Isoform 3 of 5'-AMP-activated protein kinase subunit gamma-1 OS=Homo sa | -0.36177 | 4.532914 |
| TRA1 | 0.179152 | 4.52667 |
| VDAC1 | -0.1583 | 4.526134 |
| Isoform 5 of Prostaglandin G/H synthase 1 OS=Homo sapien | -0.19128 | 4.522288 |
| Hexokinase OS=Homo sapiens | -0.13991 | 4.514491 |
| PHLDB2 | -0.14178 | 4.51437 |
| MACF1 | -0.08369 | 4.512482 |
| PRPF39 | 0.190296 | 4.508815 |
| ENO1 | 0.198979 | 4.503068 |
| Isoform 5 of MMS19 nucleotide excision repair protein homolog OS=Homo s | -0.22063 | 4.502758 |
| URLC2 | 0.225908 | 4.500198 |
| HEL-S-299 | 0.421101 | 4.499997 |
| TXNDC5 | 0.082988 | 4.496172 |
| KDM3B | 0.206433 | 4.485168 |

| | | |
|---|---|---|
| Isoform 3 of Collagen type IV alpha-3-binding protein OS=Homo sapiens G | 0.379254 | 4.482322 |
| KIAA0368 | -0.12877 | 4.480007 |
| LRBA | 0.191993 | 4.459525 |
| DHRS7 | -0.17312 | 4.459448 |
| FGF1 | -0.27913 | 4.454993 |
| PSMA4 | -0.09152 | 4.447747 |
| Isoform 3 of Ataxin-2-like protein OS=Homo sapiens | 0.090765 | 4.42977 |
| Isoform 2 of TATA element modulatory factor OS=Homo sapie | 0.368113 | 4.425387 |
| Flotillin 1 variant (Fragment) OS=Homo sapiens | -0.17598 | 4.425191 |
| PFKP | 0.529109 | 4.419624 |
| HEL-S-44 | -0.13468 | 4.415808 |
| PKM | -0.11337 | 4.411205 |
| cDNA, FLJ95265, highly similar to Homo sapiens acetyl-Coenzyme A acyltrans | 0.253886 | 4.401381 |
| ALDH18A1 | 0.231355 | 4.389577 |
| cDNA FLJ37346 fis, clone BRAMY2021310, highly similar to Transcriptional re | 0.236354 | 4.385643 |
| Heat shock 70kDa protein 1A variant (Fragment) OS=Homo sapiens | 0.302607 | 4.37012 |
| cDNA FLJ56153, highly similar to Homo sapiens transforming growth factor b | 0.18534 | 4.359593 |
| AFTPH | 0.253886 | 4.358089 |
| RABEPK | 0.142163 | 4.349572 |
| FXR2 | 0.294191 | 4.349004 |
| cDNA FLJ76913, highly similar to Homo sapiens F-box protein 7 (FBXO7), mRN | -0.18893 | 4.348685 |
| FAM180A | 1.300548 | 4.348342 |
| ZC3H13 | 1.060656 | 4.340177 |
| PDIA5 | 0.215143 | 4.339898 |
| Isoform 2 of Nuclear protein localization protein 4 homolog OS=Homo sapier | -0.07347 | 4.339867 |
| cDNA FLJ60299, highly similar to Rab GDP dissociation inhibitor beta OS=Hon | -0.06703 | 4.338936 |
| QDPR | -0.21411 | 4.337493 |
| cDNA, FLJ94534, highly similar to Homo sapiens capping protein (actin filame | 0.961037 | 4.325553 |
| SGPL1 | 0.176678 | 4.323516 |
| cDNA FLJ61541, highly similar to Homo sapiens PDZ and LIM domain 5 (PDLIN | -0.52494 | 4.319986 |
| Isoform 3 of Actin-related protein 2/3 complex subunit 4 OS=Homo sapien | -0.19767 | 4.311677 |
| Isoform 2 of Heat shock protein HSP 90-alpha OS=Homo sapiens G | -0.1901 | 4.307624 |
| NDUFS6 | 0.102964 | 4.298651 |
| PARD3 | 0.421969 | 4.298643 |
| OVCA2 | -0.39077 | 4.298607 |
| KCTD17 | 0.388716 | 4.292601 |
| GOT1 | -0.04793 | 4.287865 |
| CHURC1-FNTB | -0.09439 | 4.280598 |
| RNH1 | 0.378839 | 4.26885 |
| MATR3 | 0.158162 | 4.268378 |
| Isoform 2 of COP9 signalosome complex subunit 1 OS=Homo sapie | 0.060373 | 4.261579 |
| AQR | 1.535609 | 4.252541 |
| NUP188 | 0.242608 | 4.252046 |
| H6PD | -0.21528 | 4.244428 |
| C5orf51 | -0.21007 | 4.244393 |
| CAPZA2 | -0.18422 | 4.237917 |
| PCBD1 | -0.35301 | 4.237125 |

| | | |
|---|---|---|
| NUCKS1 | 0.153234 | 4.229872 |
| NDE1 | 0.171735 | 4.220909 |
| PES1 | 0.120082 | 4.217017 |
| Isoform 2 of UPF0687 protein C20orf27 OS=Homo sapiens G | -0.18422 | 4.209545 |
| DYNC1LI1 | -0.1264 | 4.208691 |
| DYNC1LI2 | -0.06296 | 4.207203 |
| Isoform 4 of Acyl-coenzyme A thioesterase 9, mitochondrial OS=Homo sapier | 0.107095 | 4.206209 |
| NCL | 0.113963 | 4.198271 |
| SMAD4 | -0.31293 | 4.197665 |
| Isoform 3 of Protein YIPF5 OS=Homo sapien | 0.120082 | 4.196795 |
| HMGN2 | 0.630979 | 4.19376 |
| MAPRE2 | -0.1269 | 4.184568 |
| CTHRC1 | 0.68724 | 4.179382 |
| Isoform 3 of Nuclear pore complex protein Nup153 OS=Homo sapiens | 0.172194 | 4.178341 |
| NCBP1 | 0.151541 | 4.172884 |
| C14orf142 | 0.465969 | 4.171443 |
| ACO1 | -0.23118 | 4.165203 |
| cDNA FLJ36606 fis, clone TRACH2015654, highly similar to HEAT SHOCK 70 kD | -0.14482 | 4.164164 |
| RRAGD | 0.191536 | 4.160496 |
| LIMCH1 | -0.27498 | 4.156859 |
| DYNC1H1 | -0.09795 | 4.156415 |
| LOC51064 | 0.128659 | 4.155509 |
| BLVRB | 0.133566 | 4.142173 |
| PGCP | -0.25524 | 4.134253 |
| NDUFS8 | -0.26394 | 4.131174 |
| Isoform 4 of LIM domain and actin-binding protein 1 OS=Homo sapien | 0.813587 | 4.130206 |
| NAPG | -0.15948 | 4.128334 |
| Reticulon OS=Homo sapiens | 0.182084 | 4.127677 |
| MRC2 | -0.12522 | 4.127366 |
| OXR1 | 0.314468 | 4.122952 |
| SUCLG2 | 0.121306 | 4.120483 |
| RRBP1 | 0.151234 | 4.114477 |
| RBM8 | 0.273186 | 4.113895 |
| Isoform 2 of Reticulocalbin-2 OS=Homo sapie | -0.09132 | 4.11045 |
| FAM175B | -0.21477 | 4.109091 |
| SRP54 | 0.131576 | 4.10836 |
| SH3KBP1 | 0.103431 | 4.10818 |
| DMXL1 | 0.419801 | 4.105922 |
| Isoform 3 of Calmodulin-regulated spectrin-associated protein 1 OS=Homo sa | -0.14818 | 4.105691 |
| OGT | 0.220126 | 4.09062 |
| cDNA FLJ55296, highly similar to Homo sapiens WD repeat domain 42A (WDI | -0.22584 | 4.089159 |
| Isoform 2 of Elongation factor Ts, mitochondrial OS=Homo sapie | 0.512217 | 4.086219 |
| HMGCS1 | 0.118857 | 4.086075 |
| UBE2Z | -0.06226 | 4.080793 |
| AGPS | 0.126207 | 4.079674 |
| PTGR2 | -0.22415 | 4.077659 |
| Isoform 2 of ATPase family AAA domain-containing protein 3A OS=Homo sap | 0.129421 | 4.07057 |

| | | |
|---|---|---|
| DIS3 OS=Homo sapiens | 0.340773 | 4.069911 |
| SCOC | -0.11269 | 4.067249 |
| cDNA FLJ76732, highly similar to Homo sapiens TAO kinase 3 (TAOK3), mRNA | -0.03957 | 4.06017 |
| Isoform 3 of Echinoderm microtubule-associated protein-like 2 OS=Homo sa | -0.17127 | 4.045396 |
| Isoform 3 of Peptidyl-prolyl cis-trans isomerase E OS=Homo sapie | -0.18187 | 4.045192 |
| DPY30 | -0.17363 | 4.04075 |
| APBA1 | -0.3531 | 4.039108 |
| Isoform 3 of Unconventional myosin-Ic OS=Homo sapien | -0.11337 | 4.035678 |
| TMEM184C | 0.34205 | 4.033283 |
| COX6B1 | 0.126207 | 4.032349 |
| Isoform 2 of Creatine kinase U-type, mitochondrial OS=Homo sapiens | 0.216388 | 4.029572 |
| TMEM222 | -0.134 | 4.017681 |
| ADAM9 | -0.21543 | 4.015774 |
| SIN3A | 0.163094 | 4.004745 |
| PSPC1 | 0.06644 | 3.996284 |
| TUBG1 | 0.10099 | 3.995961 |
| Isoform 4 of YLP motif-containing protein 1 OS=Homo sapien | 0.112739 | 3.992391 |
| Epididymis tissue sperm binding protein Li 14m OS=Homo sapiens | 0.10616 | 3.988913 |
| COL5A1 | 0.220919 | 3.987584 |
| AHCY | -0.15527 | 3.985246 |
| CTTN | -0.05868 | 3.977646 |
| Uncharacterized protein OS=Homo sapiens | 0.124982 | 3.964017 |
| Tissue factor OS=Homo sapiens | -0.69175 | 3.959142 |
| Isoform 2 of Fasciculation and elongation protein zeta-2 OS=Homo sapie | -0.35766 | 3.95894 |
| UQCRH | 0.804915 | 3.958937 |
| PODXL | 0.619612 | 3.958606 |
| PTTG1IP | 0.321265 | 3.956004 |
| PSMD12 | -0.09845 | 3.951651 |
| TIPRL | -0.12522 | 3.951414 |
| Isoform 2 of Minor histocompatibility antigen H13 OS=Homo sapie | 0.203161 | 3.950469 |
| MRPL24 | 0.362539 | 3.949866 |
| Isoform 2 of Tetratricopeptide repeat protein 17 OS=Homo sapien | 0.203948 | 3.947843 |
| Putative uncharacterized protein (Fragment) OS=Homo sapiens | 1.657731 | 3.945116 |
| NUP88 | 0.153234 | 3.942529 |
| ILKAP | 0.647061 | 3.939843 |
| MYH9 | -0.08439 | 3.933265 |
| MRPL4 | 0.183644 | 3.930928 |
| AP complex subunit beta OS=Homo sapiens | -0.02089 | 3.929637 |
| NUDT1 | 0.262675 | 3.917566 |
| RPN2 | -0.06226 | 3.916042 |
| Isoform Long of Proteasome subunit alpha type-1 OS=Homo sapien | 0.117633 | 3.910319 |
| cDNA FLJ77762, highly similar to Homo sapiens cullin-associated and neddyla | -0.04435 | 3.910061 |
| cDNA FLJ76924, highly similar to Homo sapiens brix domain containing 1 (BX | -0.35364 | 3.896535 |
| TMEM205 | 0.140471 | 3.888784 |
| CWF19L1 | 1.220538 | 3.887716 |
| CPPED1 | -0.17531 | 3.885081 |
| CCAR1 | 0.494552 | 3.883499 |

| | | |
|---|---|---|
| Type II 3a-hydroxysteroid dehydrogenase variant OS=Homo sapiens | 0.681262 | 3.877737 |
| Isoform 7 of Tumor protein D54 OS=Homo sapiens | -0.12404 | 3.876746 |
| AP2M1 | -0.06058 | 3.867961 |
| cDNA FLJ46506 fis, clone THYMU3030752, highly similar to BTB/POZ domain- | 1.146506 | 3.8677 |
| cDNA, FLJ94314, highly similar to Homo sapiens FOS-like antigen 1 (FOSL1), n | 0.296721 | 3.866645 |
| MROH1 | -0.22767 | 3.863471 |
| PBXIP1 | -0.18305 | 3.855939 |
| ZPR1 | -0.0825 | 3.854852 |
| CHP | -0.09558 | 3.853924 |
| DDX21 | 0.250575 | 3.852791 |
| PSMB5 | -0.03239 | 3.852635 |
| PUM3 | -0.14178 | 3.849332 |
| Isoform 2 of Ankycorbin OS=Homo sapien | -0.17009 | 3.848824 |
| UGGT1 | 0.040262 | 3.846834 |
| VPS25 | -0.13064 | 3.83403 |
| DNAJA1 | 0.053099 | 3.829973 |
| CENPC | 0.150772 | 3.82663 |
| MGAT5 | -0.22818 | 3.819737 |
| YARS | 0.232152 | 3.817049 |
| VAPB | -0.02401 | 3.815821 |
| FARP1 | -0.39309 | 3.812356 |
| SOAT1 | 0.604603 | 3.812002 |
| NUDT21 | 0.091984 | 3.808072 |
| RIPK3 | -0.22415 | 3.799505 |
| TIGAR | -0.45677 | 3.799464 |
| MSI2 | -0.17548 | 3.793834 |
| ANKRD13A | -0.24458 | 3.79358 |
| cDNA FLJ76254, highly similar to Homo sapiens gamma-glutamyl hydrolase (( | -0.13114 | 3.78978 |
| GTPBP4 | -0.12048 | 3.781949 |
| H3F3A | -0.30827 | 3.779893 |
| ATG5 | 0.112739 | 3.775915 |
| TNFAIP2 | -0.17951 | 3.775377 |
| AKR1A1 | -0.03359 | 3.77404 |
| SSFA2 | 0.057947 | 3.773351 |
| Beta-hexosaminidase OS=Homo sapiens | -0.29248 | 3.770034 |
| Isoform 2 of Intraflagellar transport protein 20 homolog OS=Homo sapien | -0.1812 | 3.764436 |
| TEP1 | 0.362132 | 3.763073 |
| Isoform 2 of Cohesin subunit SA-2 OS=Homo sapien | -0.06583 | 3.760848 |
| IHPK1 | -0.22129 | 3.758626 |
| HIBADH | -0.17245 | 3.75527 |
| C10orf76 | -0.15999 | 3.747411 |
| HNRNPA3 | 0.162633 | 3.747203 |
| Isoform 2 of Protein SMG8 OS=Homo sapie | 0.255141 | 3.742192 |
| ARHGAP10 | 0.230559 | 3.734788 |
| MDH1 | -0.28146 | 3.733325 |
| SGPP1 | 0.253886 | 3.724621 |
| C4orf32 | 1.295389 | 3.722577 |

| | | |
|---|---|---|
| PLCH1 | 0.198979 | 3.722371 |
| DPP4 | -0.53879 | 3.720485 |
| Glutathione S-transferase OS=Homo sapiens | -0.14178 | 3.716131 |
| Isoform 2 of 28S ribosomal protein S27, mitochondrial OS=Homo sapiens | -0.111 | 3.715428 |
| NAPA | 0.106628 | 3.714737 |
| PLA2G12A | -0.2399 | 3.71027 |
| EIF2AK2 | -0.06871 | 3.701574 |
| BCAT2 | 0.108605 | 3.70103 |
| TBL3 | 0.177137 | 3.699343 |
| SFXN1 | 0.179152 | 3.699171 |
| PTPN2 | 0.319551 | 3.698537 |
| HSPA14 | 0.137249 | 3.693578 |
| SPATA18 | -0.15948 | 3.683257 |
| PLOD3 | 0.121306 | 3.676036 |
| PQBP1 | 0.294191 | 3.66779 |
| EPHA6 | -0.25459 | 3.667581 |
| Isoform 7 of Protein PRRC2C OS=Homo sapiens | 0.052362 | 3.663927 |
| PAIP1 | -0.12572 | 3.660678 |
| cDNA, FLJ94517, highly similar to Homo sapiens baculoviral IAP repeat-conta | -0.09795 | 3.65813 |
| ARL15 | -0.15308 | 3.656029 |
| IDH3A | 0.078591 | 3.650997 |
| PDLIM7 | -0.26161 | 3.646941 |
| CSTF3 | -0.147 | 3.645851 |
| cDNA FLJ61387, highly similar to Homo sapiens conserved nuclear protein NH | 0.273186 | 3.63651 |
| NES | -0.90364 | 3.632371 |
| AKR1C2 | 0.495876 | 3.6296 |
| LARP1 | -0.06106 | 3.629381 |
| cDNA FLJ61383, highly similar to Serine/threonine-protein kinase 24 (EC 2.7.1 | -0.08439 | 3.628473 |
| BASP1 | 0.60052 | 3.619987 |
| THRAP3 | 0.204279 | 3.618423 |
| EXOC8 | -0.12048 | 3.618071 |
| CLUH | 0.521108 | 3.610222 |
| ANKS1A | -0.19195 | 3.610038 |
| CHMP1A | 0.623716 | 3.609495 |
| Mprip-Ntrk1 fusion gene | -0.06128 | 3.608655 |
| GARS | 0.252632 | 3.606791 |
| Ribosomal protein S19 (Fragment) OS=Homo sapiens | 0.068396 | 3.604113 |
| Isoform 6 of ADP-ribosylation factor-binding protein GGA1 OS=Homo sapie | -0.07011 | 3.602823 |
| SAMM50 | -0.05559 | 3.596113 |
| FAM134A | 0.22466 | 3.592411 |
| MACF1 | -0.09677 | 3.592388 |
| NIPSNAP1 | -0.11643 | 3.588411 |
| Isoform 2 of ATP-dependent RNA helicase DDX54 OS=Homo sapien | 0.582879 | 3.586516 |
| ARF1 | -0.06226 | 3.5852 |
| Isoform 3 of Pyrroline-5-carboxylate reductase 1, mitochondrial OS=Homo sa | 0.422834 | 3.584001 |
| COMMD8 | -0.12572 | 3.569924 |
| FAM3C | 0.103717 | 3.564493 |

| | | |
|---|---|---|
| MRI1 | -0.14296 | 3.560698 |
| Isoform 5 of Microtubule-associated protein 4 OS=Homo sapie | 0.257651 | 3.556863 |
| LTB4DH | -0.18893 | 3.549456 |
| GALNT2 | -0.1748 | 3.544924 |
| WDR59 | 0.201463 | 3.544824 |
| GOLIM4 | 0.104185 | 3.544785 |
| cDNA, FLJ95064, highly similar to Homo sapiens nin one binding protein (NOI | 0.09611 | 3.54217 |
| STX4 | 0.081494 | 3.535851 |
| Isoform 2 of Protein disulfide-isomerase A6 OS=Homo sapien | -0.0832 | 3.530736 |
| THBS1 | -0.15527 | 3.527401 |
| EIF2B5 | -0.09607 | 3.527049 |
| Isoform 2 of 40S ribosomal protein S20 OS=Homo sapien | 0.068126 | 3.51666 |
| SF3A2 | 0.224319 | 3.511531 |
| ATG16L1 | 0.257651 | 3.510449 |
| DNAJC10 | 0.061113 | 3.509583 |
| HNRNPUL2 | 0.093954 | 3.505925 |
| H.sapiens ras-related Hrab3B protein OS=Homo sapiens | -0.42668 | 3.500629 |
| PON2 | -0.14346 | 3.499651 |
| SET | -0.18069 | 3.498321 |
| CREBBP | 1.174605 | 3.498259 |
| PMM1 | -0.2188 | 3.497992 |
| UQCRB | 0.240557 | 3.497693 |
| Isoform 5 of Rap1 GTPase-GDP dissociation stimulator 1 OS=Homo sapiens G | -0.10863 | 3.497437 |
| LSG1 | -0.09251 | 3.496528 |
| Isoform 2 of P2X purinoceptor 4 OS=Homo sapien | 0.336107 | 3.490967 |
| Importin subunit alpha OS=Homo sapiens | -0.0706 | 3.489921 |
| OCIAD1 | -0.14954 | 3.48807 |
| SAP18 | 0.16186 | 3.486787 |
| PDAP1 | 0.112739 | 3.486188 |
| PTMA | 1.197282 | 3.479725 |
| AKAP12 | 0.415903 | 3.478729 |
| RANBP3 | 0.423702 | 3.478093 |
| EXOC5 | -0.07943 | 3.476418 |
| METAP2 | 0.124982 | 3.472951 |
| cDNA, FLJ96114, highly similar to Homo sapiens bromodomain and WD repea | -0.07298 | 3.470423 |
| COL6A3 | 0.369835 | 3.464043 |
| TMED5 | -0.17127 | 3.46052 |
| HGS | -0.04603 | 3.457922 |
| cDNA FLJ31479 fis, clone NT2NE2001634, moderately similar to NADH-UBIQU | 0.177915 | 3.447488 |
| MRPS36 | 0.090765 | 3.444757 |
| GAPDH | -0.17363 | 3.439008 |
| cDNA, FLJ94988, highly similar to Homo sapiens protein phosphatase 2 (form | 0.494552 | 3.436258 |
| ASCC1 | -0.21829 | 3.433068 |
| PACSIN2 | 0.101743 | 3.432764 |
| TUBGCP6 | 0.196497 | 3.432586 |
| Isoform 4 of Protein-methionine sulfoxide oxidase MICAL1 OS=Homo sapiens | 0.158934 | 3.431998 |
| HAT1 | 0.131112 | 3.426689 |

| | | |
|---|---|---|
| SEC23A | 0.077375 | 3.426482 |
| SMARCA4 | 0.1417 | 3.42339 |
| SLC12A9 | -0.10507 | 3.422455 |
| cDNA FLJ76716, highly similar to Homo sapiens WD repeat domain 70 (WDR7 | 0.318279 | 3.407124 |
| STRN | 0.110294 | 3.40589 |
| HECTD3 | -0.21946 | 3.401872 |
| NOSIP | 0.110294 | 3.399726 |
| SLC16A2 | 0.235555 | 3.395965 |
| IFIT3 | -0.50646 | 3.394451 |
| HTRA1 | 0.559935 | 3.392957 |
| Structural maintenance of chromosomes protein OS=Homo sapiens | 0.261419 | 3.392624 |
| USMG5 | -0.18136 | 3.390735 |
| Isoform 2 of Cerebral cavernous malformations 2 protein OS=Homo sapie | 0.268963 | 3.389376 |
| IFT140 | 1.071682 | 3.38718 |
| Isoform B of Membrane cofactor protein OS=Homo sapie | -0.12048 | 3.386354 |
| ACTL6A | 0.188275 | 3.385161 |
| Isoform 2 of Adenylyl cyclase-associated protein 1 OS=Homo sapie | -0.18018 | 3.384457 |
| Isoform Non-brain of Clathrin light chain A OS=Homo sapie | 0.03931 | 3.383049 |
| RMDN1 | -0.09033 | 3.381847 |
| Isoform 2 of Gelsolin OS=Homo sapi | -0.1642 | 3.375877 |
| DBNL | 0.071297 | 3.37417 |
| cDNA FLJ45033 fis, clone BRAWH3019026, highly similar to Homo sapiens ec | -0.26096 | 3.373788 |
| OR1M1 | 0.326743 | 3.370939 |
| CRIP2 | 0.822679 | 3.367783 |
| C12orf57 | -0.13705 | 3.367306 |
| CBX1 | 0.251378 | 3.365996 |
| SRGAP1 | -0.08012 | 3.363039 |
| POTEF | 0.113496 | 3.340258 |
| RNF113A | -0.04793 | 3.338192 |
| CD248 | -0.3362 | 3.333178 |
| PSMD10 | -0.15898 | 3.332367 |
| EIF4E | 0.231807 | 3.332212 |
| STOML2 | 0.088329 | 3.329861 |
| PRPF4B | -0.18254 | 3.326548 |
| PDCL3 | 0.264741 | 3.325159 |
| FLAD1 | -0.17951 | 3.32211 |
| KIAA1143 | 0.927052 | 3.319133 |
| Isoform 2 of 26S proteasome non-ATPase regulatory subunit 11 OS=Homo sa | 0.032535 | 3.315256 |
| CRNKL1 | 0.267256 | 3.3141 |
| RBMX2 | 1.114628 | 3.307072 |
| RLTPR | 0.13035 | 3.300896 |
| XPO7 | -0.06583 | 3.299926 |
| cDNA FLJ77796, highly similar to Homo sapiens evolutionarily conserved G-pa | 1.005778 | 3.296843 |
| Isoform 4 of Pyruvate dehydrogenase E1 component subunit alpha, somatic f | 0.089078 | 3.293628 |
| APMAP | -0.07893 | 3.286643 |
| cDNA FLJ10153 fis, clone HEMBA1003417, highly similar to BAG family molec | -0.30764 | 3.285862 |
| EXOSC8 | -0.22466 | 3.282951 |

| | | |
|---|---|---|
| RBM45 | 1.161523 | 3.281608 |
| XAB2 | 0.887879 | 3.269667 |
| SERPINH1 | 0.200221 | 3.266404 |
| cDNA FLJ90740 fis, clone PLACE1011045, highly similar to Autophagy-related | -0.08558 | 3.265653 |
| HTATSF1 | 0.323366 | 3.265461 |
| SSR4 | -0.08181 | 3.263481 |
| DKFZp667H197 | 0.105406 | 3.26132 |
| Isoform 4 of Treacle protein OS=Homo sapien | 0.374541 | 3.256063 |
| cDNA FLJ10398 fis, clone NT2RM4000349, highly similar to Homo sapiens bas | -0.07893 | 3.255104 |
| Tetraspanin OS=Homo sapiens | 0.388716 | 3.253474 |
| ERMP1 | 0.257202 | 3.252815 |
| SHMT2 | 0.258907 | 3.251916 |
| cDNA, FLJ93580, highly similar to Homo sapiens TRAF family member-associa | 0.20892 | 3.251754 |
| RAD21 | 0.648439 | 3.251222 |
| PRPS1 | 0.228858 | 3.250671 |
| HDGFRP3 | -0.22884 | 3.242602 |
| WDR77 | -0.1269 | 3.24193 |
| LTBP3 | 0.22466 | 3.241289 |
| PDE1C | 0.27481 | 3.236307 |
| SUPT6H | 0.113206 | 3.225964 |
| cDNA FLJ12172 fis, clone MAMMA1000684, highly similar to Opioid growth f | 0.261419 | 3.224374 |
| IGF2BP1 | -0.24289 | 3.222749 |
| DNAJC25 | 0.217634 | 3.22169 |
| S100A6 | 0.182864 | 3.219975 |
| Phosphodiesterase 5A OS=Homo sapiens | -0.26277 | 3.218282 |
| Epsilon isoform of regulatory subunit B56, protein phosphatase 2A variant (F | -0.16066 | 3.217309 |
| SERPINB8 | -0.17598 | 3.217076 |
| Condensin complex subunit 2 OS=Homo sapiens | 0.588715 | 3.21366 |
| PARVA | -0.10389 | 3.209312 |
| cDNA, FLJ93454, highly similar to Homo sapiens BM88 antigen (BM88), mRNA | 0.839348 | 3.208815 |
| NDUFA4 | 0.120841 | 3.205296 |
| ANKLE2 | 0.148311 | 3.201179 |
| WIPI2 | -0.14464 | 3.199535 |
| cDNA FLJ56034, highly similar to 4-aminobutyrate aminotransferase, mitocho | -0.24523 | 3.198771 |
| PSPH | 0.404231 | 3.198427 |
| VDAC2 | -0.17178 | 3.197014 |
| COG1 | -0.10745 | 3.195044 |
| CAMKK2 | 0.181168 | 3.188204 |
| PDIA3 | 0.10099 | 3.187139 |
| CYB5A | -0.14886 | 3.184768 |
| LTBP2 | 0.404231 | 3.180962 |
| ZNF598 | 0.200221 | 3.174125 |
| Isoform 2 of Tether containing UBX domain for GLUT4 OS=Homo sapiens | 0.158162 | 3.172882 |
| KIF7 | 0.614147 | 3.172653 |
| CASC3 | 0.198523 | 3.170789 |
| cDNA FLJ59211, highly similar to Glucosidase 2 subunit beta OS=Homo sapier | 0.142625 | 3.169261 |
| PGPEP1 | -0.21074 | 3.168969 |

| | | |
|---|---|---|
| DHX37 | 0.295456 | 3.168894 |
| FOSL2 | 0.451541 | 3.159651 |
| UBE2E2 | 0.709607 | 3.15713 |
| IGBP1 | 0.226361 | 3.156441 |
| Isoform 2 of Presequence protease, mitochondrial OS=Homo sapiens | 0.144621 | 3.156359 |
| ZNF512 | 1.074841 | 3.156277 |
| TIMM10 | 0.302607 | 3.156177 |
| RIOK3 | 0.401207 | 3.155044 |
| cDNA FLJ76427, highly similar to Homo sapiens SH2 domain binding protein 1 | 0.190753 | 3.154404 |
| PPP6R1 | 0.110761 | 3.152997 |
| TLN1 | 0.081024 | 3.1529 |
| UGGT2 | 0.074943 | 3.14908 |
| hCG_17415 | -0.0937 | 3.147573 |
| ASCC3 | -0.0825 | 3.145415 |
| cDNA, FLJ96903 OS=Homo sapiens | -0.2494 | 3.14531 |
| HEBP2 | -0.43016 | 3.141966 |
| MTCH1 | -0.17884 | 3.141386 |
| TOMM34 | -0.14364 | 3.139275 |
| Isoform 6 of Calpastatin OS=Homo sapie | 0.354442 | 3.135407 |
| ACTN1 | -0.17834 | 3.135355 |
| CWC15 | 0.76435 | 3.13521 |
| Phosphoinositide phospholipase C OS=Homo sapiens | 0.172194 | 3.126541 |
| cDNA FLJ78655, highly similar to Homo sapiens exportin 5 (XPO5), mRNA OS= | -0.1901 | 3.124942 |
| HEL-S-34 | -0.13114 | 3.123689 |
| Isoform 3 of Rap guanine nucleotide exchange factor 1 OS=Homo sapiens | 0.876437 | 3.123446 |
| PGM2L1 | -0.10557 | 3.121001 |
| ARID1A | 0.557244 | 3.120305 |
| SFRS2 | 0.09855 | 3.11785 |
| ATP6V1C1 | -0.08131 | 3.115511 |
| MYL12A | -0.11574 | 3.115351 |
| cDNA FLJ30801 fis, clone FEBRA2001217, highly similar to Homo sapiens LIM | -0.26628 | 3.109256 |
| GSTP1 | 1.981903 | 3.108708 |
| HIST2H2AB | 0.374978 | 3.107773 |
| VPS26A | -0.21242 | 3.106808 |
| TMED4 | -0.07655 | 3.104921 |
| SLC25A1 | -0.12048 | 3.102468 |
| DIAPH2 | 0.126207 | 3.102349 |
| FAM20B | 0.126207 | 3.100779 |
| TSR3 | 0.405092 | 3.100223 |
| TNRC6B | 0.40337 | 3.100158 |
| PABPC1 | 0.104939 | 3.100123 |
| TOR1A | -0.09677 | 3.096655 |
| Uncharacterized protein (Fragment) OS=Homo sapiens | 0.116409 | 3.096636 |
| Isoform 2 of ATP synthase-coupling factor 6, mitochondrial OS=Homo sapien | -0.08181 | 3.09542 |
| ARHGAP1 | -0.1335 | 3.091593 |
| PLEC | 0.251378 | 3.081956 |
| NFYB | -0.24472 | 3.079391 |

| | | |
|---|---|---|
| Stearoyl-CoA desaturase variant (Fragment) OS=Homo sapiens | 0.233056 | 3.079255 |
| SFRS5 | 0.16803 | 3.077378 |
| MRPS35 | 0.131875 | 3.074063 |
| RPL38 | -0.13823 | 3.071671 |
| Isoform 2 of Insulin-like growth factor-binding protein 3 OS=Homo sapiens | 0.495876 | 3.0706 |
| cDNA, FLJ96764, highly similar to Homo sapiens sorting nexin 8 (SNX8), mRN/ | 0.082241 | 3.059892 |
| DKFZp686E1899 | -0.09558 | 3.057473 |
| Isoform 2 of Ubiquitin carboxyl-terminal hydrolase 36 OS=Homo sapien | 0.115652 | 3.057139 |
| VARS | -0.07228 | 3.056637 |
| QRICH1 | -0.14532 | 3.056072 |
| PDCD6 | -0.07774 | 3.051594 |
| MAPK12 | 0.499849 | 3.051183 |
| ZNF326 | 0.073728 | 3.049879 |
| DHRS7B | -0.24991 | 3.049755 |
| RHOC | -0.1115 | 3.044476 |
| ATP5H | -0.07417 | 3.043599 |
| EIF2S2 | 0.116409 | 3.042311 |
| HOMER3 | 0.110294 | 3.039831 |
| 9-Sep | 0.104185 | 3.037799 |
| RUVBL2 | -0.09508 | 3.03773 |
| CASP7 | 0.39776 | 3.037292 |
| PIK3CA variant protein | -0.19312 | 3.035008 |
| PROSC | -0.18422 | 3.025944 |
| HEL-S-102 | -0.21242 | 3.01842 |
| HEL-S-8a | -0.04435 | 3.017758 |
| cDNA, FLJ93703, highly similar to Homo sapiens putatative 28 kDa protein (L( | 0.291662 | 3.013251 |
| cDNA FLJ60091, highly similar to Hypoxia-inducible factor 1 alpha inhibitor(E( | -0.15477 | 3.012545 |
| MVP | 0.093203 | 3.004563 |
| CMPK1 | -0.30128 | 3.004485 |
| CYP4F12 | -0.28379 | 3.001161 |
| TOMM7 | 0.123756 | 2.997798 |
| RPN1 | 0.074943 | 2.99702 |
| cDNA FLJ78260, highly similar to Homo sapiens RNA binding motif protein 4, | 0.112739 | 2.996424 |
| SYBL1 | -0.16538 | 2.996064 |
| cDNA FLJ56047, highly similar to A kinase anchor protein 1, mitochondrial OS | -0.73977 | 2.995582 |
| HIST2H3A | -0.32521 | 2.994236 |
| HSD17B11 | -0.14582 | 2.99418 |
| COX17 | 0.088329 | 2.988659 |
| CDH2 | -0.07844 | 2.987309 |
| ICT1 | 0.210164 | 2.987245 |
| Isoform 2 of Dual specificity mitogen-activated protein kinase kinase 4 OS=H( | -0.18657 | 2.984133 |
| ACAT1 | 0.084675 | 2.976409 |
| Isoform 2 of Actin filament-associated protein 1 OS=Homo sapien | 0.671946 | 2.975014 |
| THRAP3 | 0.652576 | 2.973455 |
| HSPA9 | 0.112739 | 2.969344 |
| UNC93B1 | -0.21125 | 2.965218 |
| SPCS1 | 0.070554 | 2.960449 |

| | | |
|---|---|---|
| cDNA, FLJ93335, highly similar to Homo sapiens PRP3 pre-mRNA processing f | -0.09201 | 2.959936 |
| Glucan , branching enzyme 1 variant (Fragment) OS=Homo sapiens | 0.366828 | 2.958103 |
| OK/KNS-cl.6 | 0.121306 | 2.955971 |
| Acyl-coenzyme A oxidase OS=Homo sapiens | 0.559935 | 2.955095 |
| Isoform 4 of DNA-directed RNA polymerases I, II, and III subunit RPABC3 OS= | 0.838285 | 2.94819 |
| DNM1L | -0.05819 | 2.936149 |
| SLC16A3 | -0.39309 | 2.934077 |
| TNPO2 variant protein | 0.135257 | 2.930388 |
| Isoform 4 of Cyclin-K OS=Homo sapie | 0.201463 | 2.926942 |
| YWHAG | -0.04793 | 2.926803 |
| B4GALT7 | -0.13232 | 2.925356 |
| EPHX1 | 0.18736 | 2.917432 |
| VAC14 | 0.175442 | 2.916237 |
| LEPREL2 protein variant (Fragment) OS=Homo sapiens | 0.977008 | 2.913456 |
| TMEM49 | 0.24386 | 2.911547 |
| DYNC1I1 | 0.844124 | 2.908974 |
| LITAF | -0.49029 | 2.906359 |
| RNF181 | 0.383556 | 2.904634 |
| PI4K2A | -0.15359 | 2.903203 |
| RBM15 | 0.281561 | 2.901346 |
| FKBP14 | -0.18069 | 2.901298 |
| SMS | 0.177456 | 2.901245 |
| Isoform USP25m of Ubiquitin carboxyl-terminal hydrolase 25 OS=Homo sapie | 0.296721 | 2.901218 |
| cDNA, FLJ92106, highly similar to Homo sapiens adaptor-related protein com | 0.068396 | 2.898685 |
| PPL | 1.256186 | 2.893734 |
| FUNDC1 | -0.16235 | 2.89362 |
| Isoform 2 of Tropomyosin alpha-4 chain OS=Homo sapie | 0.093203 | 2.892283 |
| EFR3A | -0.1583 | 2.89085 |
| HEATR6 | -0.17716 | 2.889853 |
| NOMO1 | -0.06583 | 2.883753 |
| DPP7 | -0.27446 | 2.883586 |
| cDNA FLJ60317, highly similar to Aminoacylase-1 (EC 3.5.1.14) OS=Homo sap | 0.119323 | 2.881473 |
| LGALS1 | -0.08537 | 2.879038 |
| EIF3G | 0.168489 | 2.876069 |
| MICU1 | -0.11337 | 2.875537 |
| RBM10 | 0.966341 | 2.872707 |
| DNAJC17 | 1.037173 | 2.871925 |
| Isoform Beta-2 of Protein phosphatase 1B OS=Homo sapien | 0.136021 | 2.869339 |
| GNA11 | -0.08439 | 2.866172 |
| ATP5D | -0.10438 | 2.491168 |
| ARL1 | -0.34255 | 2.426705 |
| TWF1 | -0.1032 | 2.416538 |
| NPEPPS | -0.11811 | 2.394328 |
| NRBP2 | -0.12167 | 2.322768 |
| EPB41L2 | 0.220919 | 2.249316 |
| XPO6 | 0.238855 | 2.20829 |
| cDNA FLJ55767, highly similar to Lysophosphatidic acid receptor Edg-2 OS=H | 1.178259 | 2.178163 |

| | | |
|---|---|---|
| Isoform Non-brain of Clathrin light chain B OS=Homo sapie | -0.46024 | 2.154769 |
| DNAJC13 | -0.16302 | 2.136634 |
| HEL-S-45 | -0.87616 | 2.114761 |
| Calponin (Fragment) OS=Homo sapiens | -0.84303 | 2.095562 |
| COX2 | 0.177456 | 2.0631 |
| SMARCC2 | 0.454161 | 2.001783 |
| SDHB | 0.296721 | 1.955791 |
| GOLGA1 | 0.076904 | 1.95243 |
| hCG_2025883 | 0.537127 | 1.812555 |
| C8orf33 | -0.56996 | 1.782298 |
| CDK2AP1 | 0.520198 | 1.780276 |
| AGO2 | -0.16706 | 1.769668 |
| TRAPPC3 | 0.405092 | 1.754181 |
| PGAM5 | 0.744449 | 1.724969 |
| UBQLN2 | -0.15999 | 1.724372 |
| COLGALT1 | 0.100522 | 1.710442 |
| ATG3 | -0.10389 | 1.68942 |
| RPP38 | 0.611417 | 1.682987 |
| cDNA FLJ75881, highly similar to Homo sapiens transferrin receptor (p90, CD | -0.48391 | 1.672112 |
| SPAG9 | 0.181626 | 1.669201 |
| CCDC9 | 0.368113 | 1.656758 |
| ATRX | 0.669172 | 1.650209 |
| RAVER1 | 0.592368 | 1.625251 |
| ARFGAP2 | 0.36426 | 1.623663 |
| cDNA FLJ51221, highly similar to Ubiquitin-protein ligase RMA1 (EC 6.3.2.-) O | 0.955324 | 1.608254 |
| ECI1 | 0.104652 | 1.586563 |
| ABCF3 | -0.08656 | 1.585287 |
| BMS1 | 0.698406 | 1.581738 |
| RBMS2 | -0.36355 | 1.581132 |
| cDNA FLJ76871, highly similar to Homo sapiens DEAH (Asp-Glu-Ala-His) box p | 0.405092 | 1.565667 |
| TUBB2A | -0.35069 | 1.522494 |
| HEXB | -0.52725 | 1.507519 |
| USP19 | 0.479594 | 1.494354 |
| Isoform 2 of Secernin-1 OS=Homo sapien | -0.21477 | 1.493058 |
| cDNA FLJ51932 OS=Homo sapiens | 0.576116 | 1.490848 |
| HPS6 | -0.16824 | 1.482452 |
| UACA | -0.35248 | 1.473401 |
| MTX1 | 0.080554 | 1.466397 |
| LMOD1 | -0.56765 | 1.42799 |

| pvalue |
| --- |
| #VALUE! |
| #VALUE! |
| #VALUE! |
| #VALUE! |
| #VALUE! |
| #VALUE! |
| #VALUE! |
| 1.20699E-14 |
| 1.75061E-13 |
| 4.67843E-13 |
| 4.74067E-13 |
| 5.54849E-13 |
| 1.81255E-12 |
| 3.74896E-12 |
| 3.67489E-11 |
| 1.19062E-10 |
| 2.16704E-10 |
| 3.87615E-10 |
| 5.0965E-10 |
| 6.31023E-10 |
| 6.54919E-10 |
| 1.00892E-09 |
| 1.05191E-09 |
| 1.13269E-09 |
| 1.20251E-09 |
| 2.01258E-09 |
| 2.71619E-09 |
| 3.30238E-09 |
| 4.24927E-09 |
| 4.25659E-09 |
| 4.30844E-09 |
| 6.21357E-09 |
| 6.36372E-09 |
| 8.03212E-09 |
| 9.70492E-09 |
| 1.27753E-08 |
| 1.29715E-08 |
| 1.5035E-08 |
| 1.53018E-08 |
| 1.63263E-08 |
| 2.12605E-08 |
| 2.33243E-08 |
| 2.60887E-08 |
| 2.65062E-08 |
| 2.69535E-08 |
| 2.72196E-08 |

| |
|---|
| 2.75148E-08 |
| 2.96845E-08 |
| 3.00942E-08 |
| 3.10568E-08 |
| 3.12741E-08 |
| 3.1289E-08 |
| 3.40427E-08 |
| 3.5069E-08 |
| 3.51577E-08 |
| 3.79822E-08 |
| 4.38442E-08 |
| 4.45154E-08 |
| 4.72224E-08 |
| 4.93118E-08 |
| 4.94718E-08 |
| 5.49288E-08 |
| 5.50378E-08 |
| 5.53194E-08 |
| 5.59437E-08 |
| 6.35841E-08 |
| 6.59103E-08 |
| 7.02858E-08 |
| 7.4247E-08 |
| 7.49847E-08 |
| 7.79382E-08 |
| 8.28023E-08 |
| 8.60643E-08 |
| 8.66364E-08 |
| 9.10985E-08 |
| 9.52834E-08 |
| 9.76333E-08 |
| 1.09288E-07 |
| 1.11574E-07 |
| 1.14419E-07 |
| 1.20863E-07 |
| 1.28788E-07 |
| 1.40007E-07 |
| 1.76701E-07 |
| 1.93449E-07 |
| 2.07273E-07 |
| 2.09779E-07 |
| 2.27471E-07 |
| 2.46422E-07 |
| 2.49771E-07 |
| 2.52488E-07 |
| 2.54422E-07 |
| 2.59826E-07 |

| |
|---|
| 2.63268E-07 |
| 2.65083E-07 |
| 2.65501E-07 |
| 2.75763E-07 |
| 2.80193E-07 |
| 3.45943E-07 |
| 3.47173E-07 |
| 3.48309E-07 |
| 3.50207E-07 |
| 3.68763E-07 |
| 4.10965E-07 |
| 4.39515E-07 |
| 4.40948E-07 |
| 4.41201E-07 |
| 4.63409E-07 |
| 4.74638E-07 |
| 4.83118E-07 |
| 5.20693E-07 |
| 5.23789E-07 |
| 5.35564E-07 |
| 5.3925E-07 |
| 5.42446E-07 |
| 5.66233E-07 |
| 5.77738E-07 |
| 5.83889E-07 |
| 6.12853E-07 |
| 6.40644E-07 |
| 6.55946E-07 |
| 6.56209E-07 |
| 6.62439E-07 |
| 7.20308E-07 |
| 7.99017E-07 |
| 8.35023E-07 |
| 8.56926E-07 |
| 8.71836E-07 |
| 8.7456E-07 |
| 1.01488E-06 |
| 1.0525E-06 |
| 1.08021E-06 |
| 1.13883E-06 |
| 1.16347E-06 |
| 1.21723E-06 |
| 1.27112E-06 |
| 1.30115E-06 |
| 1.30451E-06 |
| 1.30958E-06 |
| 1.31862E-06 |

| |
|---|
| 1.35905E-06 |
| 1.43E-06 |
| 1.44293E-06 |
| 1.47522E-06 |
| 1.6112E-06 |
| 1.70689E-06 |
| 1.72996E-06 |
| 1.73308E-06 |
| 1.7789E-06 |
| 1.78572E-06 |
| 1.79959E-06 |
| 1.88333E-06 |
| 1.98861E-06 |
| 2.07118E-06 |
| 2.15576E-06 |
| 2.32646E-06 |
| 2.35442E-06 |
| 2.42212E-06 |
| 2.4231E-06 |
| 2.42538E-06 |
| 2.55575E-06 |
| 2.62739E-06 |
| 2.64666E-06 |
| 2.659E-06 |
| 2.67393E-06 |
| 2.72956E-06 |
| 2.7434E-06 |
| 2.78792E-06 |
| 2.80067E-06 |
| 2.81083E-06 |
| 2.89355E-06 |
| 2.90166E-06 |
| 2.92038E-06 |
| 2.97648E-06 |
| 2.98707E-06 |
| 3.09027E-06 |
| 3.36463E-06 |
| 3.38321E-06 |
| 3.38819E-06 |
| 3.45206E-06 |
| 3.46489E-06 |
| 3.46707E-06 |
| 3.50116E-06 |
| 3.62331E-06 |
| 3.71209E-06 |
| 3.71812E-06 |
| 3.86315E-06 |

| |
|---|
| 3.96108E-06 |
| 4.00217E-06 |
| 4.22769E-06 |
| 4.23136E-06 |
| 4.29206E-06 |
| 4.31841E-06 |
| 4.37391E-06 |
| 4.37585E-06 |
| 4.51411E-06 |
| 4.53983E-06 |
| 4.62042E-06 |
| 4.75625E-06 |
| 5.05591E-06 |
| 5.16186E-06 |
| 5.2164E-06 |
| 5.23764E-06 |
| 5.32921E-06 |
| 5.33013E-06 |
| 5.37504E-06 |
| 5.71625E-06 |
| 5.76722E-06 |
| 6.0251E-06 |
| 6.26211E-06 |
| 6.32022E-06 |
| 6.43514E-06 |
| 6.5494E-06 |
| 6.58468E-06 |
| 6.65094E-06 |
| 6.83116E-06 |
| 6.84109E-06 |
| 6.88178E-06 |
| 7.17235E-06 |
| 7.28283E-06 |
| 7.30427E-06 |
| 7.39871E-06 |
| 7.45116E-06 |
| 7.64725E-06 |
| 7.72531E-06 |
| 7.74221E-06 |
| 7.75335E-06 |
| 7.78727E-06 |
| 7.87045E-06 |
| 8.03127E-06 |
| 8.05596E-06 |
| 8.4092E-06 |
| 8.44448E-06 |
| 8.63321E-06 |

| |
|---|
| 8.72559E-06 |
| 8.79982E-06 |
| 8.98691E-06 |
| 9.3438E-06 |
| 9.36282E-06 |
| 9.41973E-06 |
| 9.48504E-06 |
| 9.53089E-06 |
| 9.53673E-06 |
| 9.60061E-06 |
| 9.84249E-06 |
| 9.88398E-06 |
| 1.01877E-05 |
| 1.03408E-05 |
| 1.06732E-05 |
| 1.19519E-05 |
| 1.21203E-05 |
| 1.22755E-05 |
| 1.22911E-05 |
| 1.23517E-05 |
| 1.23732E-05 |
| 1.31054E-05 |
| 1.32295E-05 |
| 1.32918E-05 |
| 1.3394E-05 |
| 1.4075E-05 |
| 1.45144E-05 |
| 1.46179E-05 |
| 1.49808E-05 |
| 1.50783E-05 |
| 1.51708E-05 |
| 1.52892E-05 |
| 1.54332E-05 |
| 1.58222E-05 |
| 1.58256E-05 |
| 1.60555E-05 |
| 1.66496E-05 |
| 1.68731E-05 |
| 1.69927E-05 |
| 1.71422E-05 |
| 1.76063E-05 |
| 1.76277E-05 |
| 1.80053E-05 |
| 1.81435E-05 |
| 1.82921E-05 |
| 1.83314E-05 |
| 1.85648E-05 |

| |
|---|
| 1.86807E-05 |
| 1.89511E-05 |
| 1.99559E-05 |
| 1.99694E-05 |
| 2.0093E-05 |
| 2.01247E-05 |
| 2.10487E-05 |
| 2.129E-05 |
| 2.15547E-05 |
| 2.216E-05 |
| 2.22754E-05 |
| 2.32938E-05 |
| 2.35449E-05 |
| 2.37021E-05 |
| 2.39011E-05 |
| 2.39762E-05 |
| 2.40356E-05 |
| 2.40609E-05 |
| 2.48982E-05 |
| 2.52109E-05 |
| 2.53353E-05 |
| 2.54661E-05 |
| 2.59529E-05 |
| 2.63937E-05 |
| 2.65451E-05 |
| 2.66543E-05 |
| 2.71004E-05 |
| 2.78971E-05 |
| 2.81125E-05 |
| 2.83307E-05 |
| 2.86101E-05 |
| 2.8647E-05 |
| 2.92292E-05 |
| 2.93148E-05 |
| 2.97392E-05 |
| 2.9776E-05 |
| 3.00408E-05 |
| 3.0585E-05 |
| 3.05936E-05 |
| 3.07268E-05 |
| 3.09874E-05 |
| 3.14002E-05 |
| 3.14226E-05 |
| 3.16084E-05 |
| 3.1623E-05 |
| 3.19027E-05 |
| 3.27214E-05 |

| |
|---|
| 3.29365E-05 |
| 3.31125E-05 |
| 3.47116E-05 |
| 3.47178E-05 |
| 3.50758E-05 |
| 3.56659E-05 |
| 3.71732E-05 |
| 3.75503E-05 |
| 3.75672E-05 |
| 3.80519E-05 |
| 3.83877E-05 |
| 3.87967E-05 |
| 3.96844E-05 |
| 4.07777E-05 |
| 4.11488E-05 |
| 4.26461E-05 |
| 4.36925E-05 |
| 4.38441E-05 |
| 4.47124E-05 |
| 4.47709E-05 |
| 4.48038E-05 |
| 4.48392E-05 |
| 4.56901E-05 |
| 4.57196E-05 |
| 4.57228E-05 |
| 4.58209E-05 |
| 4.59734E-05 |
| 4.72549E-05 |
| 4.74771E-05 |
| 4.78646E-05 |
| 4.87891E-05 |
| 4.92465E-05 |
| 5.02747E-05 |
| 5.02756E-05 |
| 5.02797E-05 |
| 5.098E-05 |
| 5.15388E-05 |
| 5.24086E-05 |
| 5.38456E-05 |
| 5.39041E-05 |
| 5.47546E-05 |
| 5.5906E-05 |
| 5.59698E-05 |
| 5.69602E-05 |
| 5.69648E-05 |
| 5.78207E-05 |
| 5.79262E-05 |

| |
|---|
| 5.89017E-05 |
| 6.013E-05 |
| 6.06713E-05 |
| 6.17241E-05 |
| 6.18457E-05 |
| 6.20579E-05 |
| 6.22001E-05 |
| 6.33475E-05 |
| 6.34359E-05 |
| 6.35631E-05 |
| 6.40089E-05 |
| 6.53781E-05 |
| 6.61635E-05 |
| 6.63222E-05 |
| 6.71608E-05 |
| 6.7384E-05 |
| 6.83592E-05 |
| 6.85229E-05 |
| 6.91041E-05 |
| 6.96853E-05 |
| 6.97566E-05 |
| 6.99022E-05 |
| 7.20821E-05 |
| 7.34086E-05 |
| 7.39308E-05 |
| 7.40958E-05 |
| 7.4416E-05 |
| 7.45285E-05 |
| 7.45819E-05 |
| 7.53439E-05 |
| 7.57734E-05 |
| 7.68285E-05 |
| 7.69316E-05 |
| 7.75443E-05 |
| 7.77873E-05 |
| 7.79184E-05 |
| 7.79508E-05 |
| 7.83571E-05 |
| 7.83988E-05 |
| 8.11672E-05 |
| 8.14407E-05 |
| 8.19938E-05 |
| 8.20209E-05 |
| 8.30246E-05 |
| 8.32389E-05 |
| 8.36259E-05 |
| 8.50021E-05 |

| |
|---|
| 8.51313E-05 |
| 8.56546E-05 |
| 8.70623E-05 |
| 9.0075E-05 |
| 9.01172E-05 |
| 9.10436E-05 |
| 9.13887E-05 |
| 9.21133E-05 |
| 9.26225E-05 |
| 9.2822E-05 |
| 9.34173E-05 |
| 9.60106E-05 |
| 9.6433E-05 |
| 9.89135E-05 |
| 0.000100859 |
| 0.000100934 |
| 0.000101767 |
| 0.000102586 |
| 0.0001029 |
| 0.000103456 |
| 0.000105282 |
| 0.000108638 |
| 0.000109865 |
| 0.000109916 |
| 0.000109916 |
| 0.00011 |
| 0.000110661 |
| 0.000111776 |
| 0.000111837 |
| 0.000112081 |
| 0.000112236 |
| 0.000112761 |
| 0.000113471 |
| 0.000114149 |
| 0.000114857 |
| 0.00011661 |
| 0.000117239 |
| 0.000117588 |
| 0.000120902 |
| 0.000121327 |
| 0.000122937 |
| 0.00012301 |
| 0.000126901 |
| 0.000129186 |
| 0.000129504 |
| 0.000130292 |
| 0.000130768 |

| |
|---|
| 0.000132514 |
| 0.000132817 |
| 0.000135531 |
| 0.000135613 |
| 0.000135942 |
| 0.00013694 |
| 0.000139335 |
| 0.000139684 |
| 0.000139983 |
| 0.000140349 |
| 0.000140399 |
| 0.000141471 |
| 0.000141637 |
| 0.000142287 |
| 0.000146545 |
| 0.00014792 |
| 0.000149063 |
| 0.000151448 |
| 0.000152388 |
| 0.000152819 |
| 0.000154044 |
| 0.000154169 |
| 0.000155571 |
| 0.00015867 |
| 0.000158685 |
| 0.000160756 |
| 0.00016085 |
| 0.000162263 |
| 0.000165215 |
| 0.000165999 |
| 0.000167527 |
| 0.000167735 |
| 0.000168252 |
| 0.000168519 |
| 0.000169811 |
| 0.000172014 |
| 0.000172555 |
| 0.000173441 |
| 0.000174331 |
| 0.000175683 |
| 0.000178891 |
| 0.000178977 |
| 0.000181054 |
| 0.000184167 |
| 0.000184788 |
| 0.000188529 |
| 0.000189419 |

| |
|---|
| 0.000189509 |
| 0.000190334 |
| 0.000192251 |
| 0.000192563 |
| 0.000192869 |
| 0.000194863 |
| 0.000198805 |
| 0.000199054 |
| 0.000199828 |
| 0.000199907 |
| 0.0002002 |
| 0.000202499 |
| 0.000207368 |
| 0.000210845 |
| 0.000214887 |
| 0.00021499 |
| 0.000216807 |
| 0.000218435 |
| 0.00021972 |
| 0.000220786 |
| 0.000223359 |
| 0.000225454 |
| 0.000226021 |
| 0.000230935 |
| 0.000233146 |
| 0.000234639 |
| 0.000234757 |
| 0.000235248 |
| 0.00023989 |
| 0.000240756 |
| 0.000240951 |
| 0.000245345 |
| 0.000245449 |
| 0.000245757 |
| 0.000246232 |
| 0.000247291 |
| 0.000248821 |
| 0.000249561 |
| 0.000253447 |
| 0.000255616 |
| 0.00025563 |
| 0.000257982 |
| 0.00025911 |
| 0.000259896 |
| 0.000260615 |
| 0.0002692 |
| 0.000272588 |

| |
|---|
| 0.000274981 |
| 0.00027742 |
| 0.000282192 |
| 0.000285152 |
| 0.000285217 |
| 0.000285243 |
| 0.000286966 |
| 0.000291171 |
| 0.000294621 |
| 0.000296892 |
| 0.000297133 |
| 0.000304327 |
| 0.000307942 |
| 0.00030871 |
| 0.000309326 |
| 0.000311943 |
| 0.00031577 |
| 0.000316482 |
| 0.000317453 |
| 0.000317498 |
| 0.000317693 |
| 0.000317912 |
| 0.000318099 |
| 0.000318766 |
| 0.000322874 |
| 0.000323653 |
| 0.000325035 |
| 0.000325997 |
| 0.000326446 |
| 0.000331341 |
| 0.000332102 |
| 0.000332588 |
| 0.000333874 |
| 0.00033655 |
| 0.000338514 |
| 0.000343524 |
| 0.000346322 |
| 0.0003484 |
| 0.000356871 |
| 0.000359122 |
| 0.000363909 |
| 0.00036622 |
| 0.00036892 |
| 0.000369178 |
| 0.000369329 |
| 0.00036983 |
| 0.000374379 |

| |
|---|
| 0.000374557 |
| 0.000377233 |
| 0.000378046 |
| 0.00039163 |
| 0.000392744 |
| 0.000396395 |
| 0.000398358 |
| 0.000401823 |
| 0.000403227 |
| 0.000404616 |
| 0.000404926 |
| 0.000406692 |
| 0.000407966 |
| 0.000410034 |
| 0.000410815 |
| 0.000411945 |
| 0.000412614 |
| 0.000413953 |
| 0.000415101 |
| 0.000420846 |
| 0.000422503 |
| 0.000422875 |
| 0.000425658 |
| 0.000428762 |
| 0.000429234 |
| 0.00043053 |
| 0.000433472 |
| 0.000456816 |
| 0.000458995 |
| 0.000464325 |
| 0.000465192 |
| 0.000465359 |
| 0.000467885 |
| 0.000471468 |
| 0.000472978 |
| 0.00047631 |
| 0.000479586 |
| 0.000483887 |
| 0.000485177 |
| 0.000493092 |
| 0.000500154 |
| 0.000501273 |
| 0.000504844 |
| 0.000508595 |
| 0.000516841 |
| 0.000517771 |
| 0.000521254 |

| |
|---|
| 0.000522868 |
| 0.000537443 |
| 0.000541497 |
| 0.000542434 |
| 0.000542674 |
| 0.000545154 |
| 0.000547873 |
| 0.000554546 |
| 0.000555771 |
| 0.000557861 |
| 0.000558709 |
| 0.000559866 |
| 0.000560075 |
| 0.000560762 |
| 0.000561474 |
| 0.000572003 |
| 0.000572888 |
| 0.000573735 |
| 0.000580354 |
| 0.000594341 |
| 0.000596521 |
| 0.000598757 |
| 0.00060022 |
| 0.000602595 |
| 0.000604948 |
| 0.000606305 |
| 0.00060663 |
| 0.000611421 |
| 0.000617573 |
| 0.00061828 |
| 0.000623309 |
| 0.000629247 |
| 0.000631633 |
| 0.000632745 |
| 0.000633247 |
| 0.000635311 |
| 0.000638198 |
| 0.00064833 |
| 0.000649922 |
| 0.000653479 |
| 0.000659232 |
| 0.000669692 |
| 0.000671612 |
| 0.000671966 |
| 0.000674856 |
| 0.000677235 |
| 0.00067769 |

0.000677807
0.000692387
0.000696418
0.000697523
0.000697656
0.000697787
0.000697948
0.000699771
0.000700803
0.000703078
0.000703235
0.000709447
0.000711913
0.00071546
0.000715633
0.000721165
0.000722128
0.000725647
0.000732139
0.000732226
0.000732471
0.000747239
0.000749995
0.000752161
0.000752582
0.000756832
0.000758046
0.000762343
0.00076646
0.000766742
0.000777578
0.000778561
0.000780238
0.000781973
0.000785378
0.000789827
0.000790044
0.000792905
0.00079392
0.00079404
0.000794104
0.00080047
0.000800505
0.00080275
0.000809855
0.000828026
0.000832931

| |
|---|
| 0.000833192 |
| 0.000836801 |
| 0.000843212 |
| 0.000847869 |
| 0.000849963 |
| 0.00087118 |
| 0.000876046 |
| 0.00087672 |
| 0.000877734 |
| 0.000878876 |
| 0.000887986 |
| 0.000888826 |
| 0.0008915 |
| 0.000891754 |
| 0.000902659 |
| 0.000904485 |
| 0.000907171 |
| 0.000912365 |
| 0.000916644 |
| 0.00091679 |
| 0.000917715 |
| 0.000922555 |
| 0.000942011 |
| 0.000958473 |
| 0.000959936 |
| 0.000969949 |
| 0.000971528 |
| 0.000989549 |
| 0.000989726 |
| 0.000997329 |
| 0.001005083 |
| 0.001006885 |
| 0.001008268 |
| 0.001009105 |
| 0.001010224 |
| 0.00101336 |
| 0.001013491 |
| 0.001026458 |
| 0.001029653 |
| 0.001029804 |
| 0.001037211 |
| 0.001055824 |
| 0.00105922 |
| 0.001063028 |
| 0.001073138 |
| 0.001083383 |
| 0.001095345 |

| |
|---|
| 0.00109664 |
| 0.001101279 |
| 0.001106699 |
| 0.001108931 |
| 0.001126703 |
| 0.001158381 |
| 0.001163919 |
| 0.001173847 |
| 0.0011832 |
| 0.001183578 |
| 0.001187529 |
| 0.001209394 |
| 0.001212727 |
| 0.001220518 |
| 0.001225893 |
| 0.001233178 |
| 0.001240628 |
| 0.001245563 |
| 0.001249675 |
| 0.001255028 |
| 0.001255169 |
| 0.001255323 |
| 0.001255399 |
| 0.001262744 |
| 0.001277221 |
| 0.001277557 |
| 0.001281495 |
| 0.001285729 |
| 0.001288686 |
| 0.001306915 |
| 0.001307415 |
| 0.001313794 |
| 0.00132118 |
| 0.001330242 |
| 0.001331872 |
| 0.001340582 |
| 0.001342996 |
| 0.001351019 |
| 0.001360905 |
| 0.003227247 |
| 0.003743645 |
| 0.003832318 |
| 0.004033411 |
| 0.004755897 |
| 0.005632282 |
| 0.00619028 |
| 0.006634937 |

| |
|---|
| 0.007002143 |
| 0.00730072 |
| 0.007677845 |
| 0.008024871 |
| 0.008647697 |
| 0.009959019 |
| 0.011071554 |
| 0.01115759 |
| 0.015397302 |
| 0.016508277 |
| 0.016585319 |
| 0.016995426 |
| 0.017612436 |
| 0.018837844 |
| 0.018863757 |
| 0.019478615 |
| 0.020444666 |
| 0.020749762 |
| 0.021275924 |
| 0.021419013 |
| 0.022041557 |
| 0.022376456 |
| 0.023700017 |
| 0.023786871 |
| 0.024645952 |
| 0.025908181 |
| 0.025984429 |
| 0.026197612 |
| 0.026234209 |
| 0.027185204 |
| 0.030026621 |
| 0.031079999 |
| 0.032036537 |
| 0.032132277 |
| 0.032296222 |
| 0.032926714 |
| 0.033620076 |
| 0.034166736 |
| 0.037325858 |
